# 338 coleopteran genomes reveal exceptional rearrangement variation compared to other insect orders

**DOI:** 10.64898/2026.07.17.739156

**Authors:** Arif Maulana, Aleksandra Bliznina, Sam Ebdon, Jessica T. Thorpe, Julia Gries, Karin Näsvall, Duane D. McKenna, Ksenia Krasheninnikova, Camilla A. Santos, Karen Brooks, Dominic E. Absolon, Danil Zilov, Darwin Tree of Life Consortium, Kamil S. Jaron, Joana I Meier

## Abstract

Chromosomal rearrangements reshape genome architecture, but rate variation across the tree of life remains poorly understood. Analysing 338 beetle (Coleoptera) genomes, we inferred eight ancestral linkage groups. No species retained these linkage groups, whereas the most frequent karyotypes of 10 or 11 chromosomes evolved at least 65 times through independent rearrangements. Rearrangement rates vary widely across lineages, with exceptionally high rates associated with major species radiations. Comparisons to 340 fly and 210 butterfly and moth genomes reveal the highest rates of long-term persistent rearrangements in beetles and that the beetle X chromosome traces back to the ancestor of Holometabola. We identified frequent X chromosome changes, mostly representing recent autosomal-X fusions, but no fissions. For autosomes, key rearrangement predictors include chromosome size, repeats and centromere position.

## Introduction

In eukaryotes, chromosomes are the physical units storing, organising, and transmitting genetic information. Changes in chromosome structure, collectively termed chromosomal rearrangements, can have profound consequences for genome function, adaptation and speciation (*1*, *2*). The rates of large-scale chromosomal rearrangements, such as fissions, fusions, and translocations, vary widely among lineages (*3*, *4*). Chromosome-scale comparative studies have revealed that ancestral linkage groups can remain conserved over hundreds of millions of years in some animal lineages, whereas others show extensive reorganisation of ancestral chromosomal structure (*5–7*). The processes determining how these contrasting patterns arise and accumulate over macroevolutionary time, why particular chromosomes are more prone to rearrangement, and the macroevolutionary significance of this variation remain unclear.

Several mechanisms and genomic features have been proposed to influence chromosomal rearrangement dynamics, including chromosome size, sex linkage, recombination landscape, repeat content, gene density, segmental duplications, centromere architecture, and nuclear organisation (*8–19*). However, empirical evidence remains limited. Recent large-scale work on 210 Lepidoptera genomes has shown that chromosome size, repeat content, and sex linkage are correlated with the probability of particular rearrangement types, but did not reveal any correlates of lineage-specific variation in rearrangement rates (*20*). Comparable large-scale analyses remain rare, leaving the generality of these predictors and their relevance to long-term macroevolutionary patterns poorly resolved. Broad comparative studies based on cytogenetic data have revealed large variation in the speed of chromosome number evolution (e.g., *9*, *21*), but they cannot determine homology of rearranged chromosomes, addressing whether similar karyotypes arose through the same or different rearrangement events, or identifying genomic features that predict rearrangements. The increasing availability of high-quality chromosome-scale reference genomes, together with advances in comparative genomic tools, now makes it possible to reconstruct ancestral linkage groups and investigate the genomic determinants of rearrangement across much deeper evolutionary timescales (*22–26*).

Coleoptera (beetles) comprise an extremely species-rich animal order, with over 400,000 described species, approximately 20% of all known species, and spanning more than 300 million years of evolutionary history (*27–31*). Cytogenetic studies across thousands of species have revealed substantial karyotypic variation, including wide variation in chromosome number (haploid number, *n* = 3–39), suggesting numerous chromosomal fissions and fusions (*32*). This long history encompasses survival through major biotic crises, repeated colonization of terrestrial and aquatic habitats, and major diet transitions between plants, fungi, wood, litter, carrion or prey. Beetles therefore provide an unusually rich system for asking whether chromosomal evolution is a largely neutral background process or whether its tempo and constraints vary with the ecological and genomic transitions that shaped a major animal radiation. However, chromosome evolution in beetles remains poorly understood beyond chromosome counts (*33*, *34*). Recent comparative genomic studies have begun to reconstruct chromosomal synteny across species, revealing generally conserved linkage patterns alongside extensive lineage-specific structural reorganisation (*35*, *36*), but these analyses have been taxonomically restricted with a relatively small number of species, limiting their power to assess broad-scale rearrangement patterns and underlying causes.

Here, we analyse 338 chromosome-scale beetle genomes, including 231 assemblies generated by the Darwin Tree of Life Project (*37*) and 107 from other initiatives (e.g., (*38, 39*) (Table S1)). We reconstructed the ancestral linkage groups (ALGs) of Coleoptera, i.e., sets of genes that were linked on ancestral chromosomes, to characterise the patterns, constraints, and temporal dynamics of chromosomal rearrangements at a macroevolutionary scale. We then asked whether rearrangement dynamics are predictable from intrinsic genomic features, whether similar karyotypes reflect shared ancestry or repeated convergence, and whether bursts of chromosomal change coincide with major ecological radiations. We further assess how these processes may have contributed to beetle diversification and compare our findings with chromosomal rearrangement patterns reported in other major insect orders, including flies (Diptera) (*40*) and butterflies and moths (Lepidoptera) (*20*).

### Eight ancestral linkage groups reveal deep macrosyntenic conservation in beetles

We analysed 338 high-quality beetle genomes representing 18 of 25 (72%) superfamilies and 52 of 178 (29%) families, together with 16 outgroup species from Strepsiptera, Neuroptera, Megaloptera and Raphidioptera, to investigate chromosome evolution across Coleoptera (Fig. 1A). The dataset captures substantial variation in karyotype and genome size, largely consistent with previous cytological studies (*32*), ranging from *n* = 5 in *Endomychus coccineus* to *n* = 39 in *Euwallacea fornicatus* (*41*), and from 100.37 Mb (*Metoecus paradoxus*) to 2.59 Gb (*Agriotes lineatus*) representing an approximately 26-fold variation in genome size (Fig. S1; Table S1). We found no association between genome size and chromosome number across sampled species (PGLS: *n =* 355; t = -0.238, *p =* 0.812, adjusted R^2^ = -0.0027, λ = 0.971; Fig. S2).

**Fig. 1.**
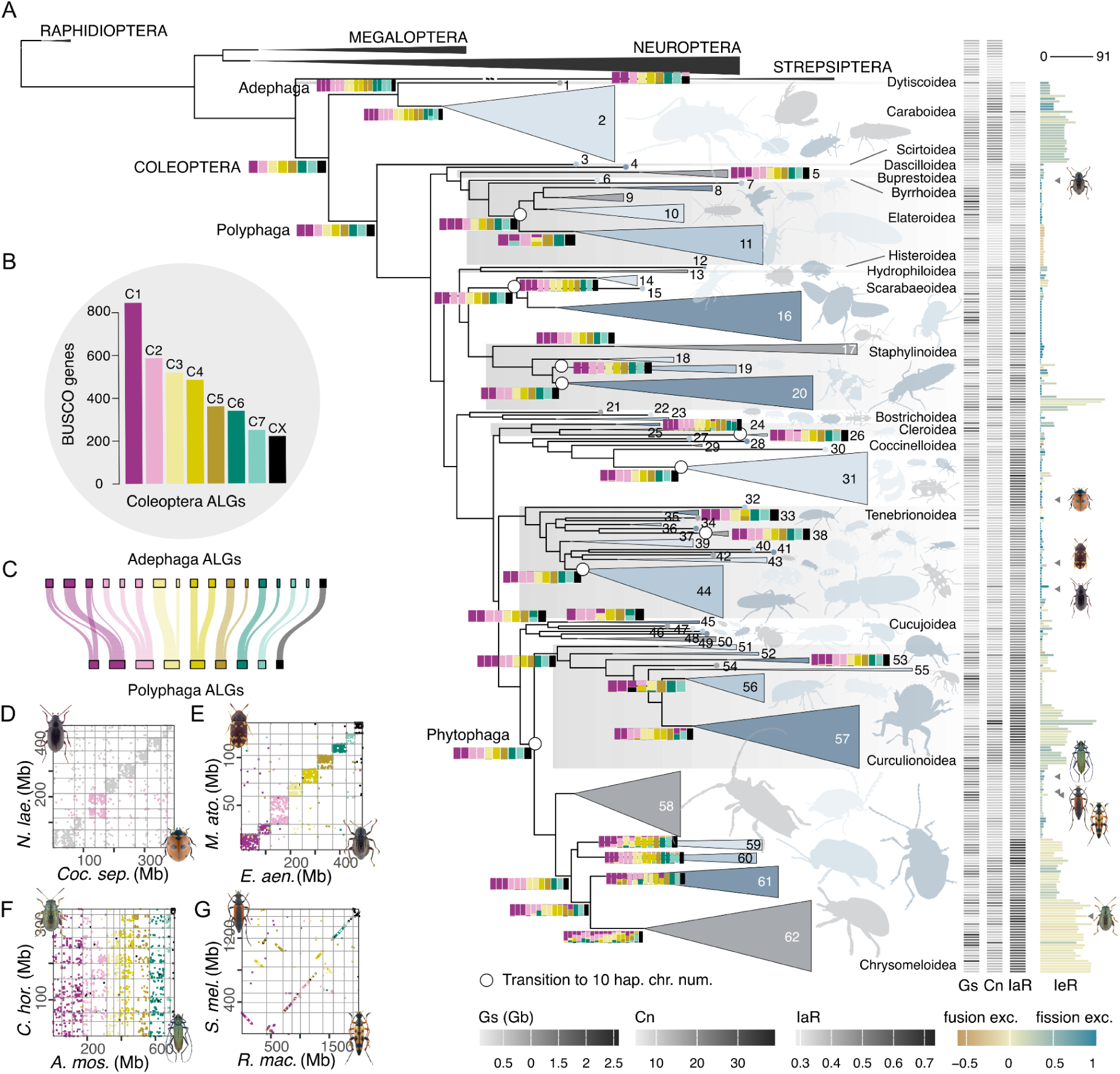
Ancestral linkage groups (ALGs) and chromosomal evolution in beetles. (A) Family-level tree showing reconstructed chromosomes and rearrangement events above the family level. Alternating grey shading indicates different superfamilies, and members of the same families or clades were collapsed and coloured in alternating shades of blue. Linkage groups at internal nodes are simplified and visualised as coloured rectangles, with the colours showing the proportion of genes assigned to each ALG. Chromosomes are separated by white gaps. Changes in ALG configuration are shown only for families or clades with more than one representative species. Open circles indicate internal nodes at which a karyotype of 10 haploid chromosomes evolved independently from the nine polyphagan ALGs. These recurrent transitions are described in more detail in Fig. S14. On the right side of the phylogeny, a heatmap shows genome sizes (Gs), haploid chromosome numbers (Cn), and gene order fragmentation index (*FI*) as a measure of intrachromosomal rearrangements (IaR) for each species. The barplot on the far right shows the total number of interchromosomal rearrangements (IeR) since the origin of beetles and is coloured according to the ratio of the numbers of fissions and fusions. **(B) Ancestral karyotype of Coleoptera**. Size distribution of the ancestral linkage group (ALG) by number of BUSCO genes. The sex-linked CX is the smallest ALG. **(C) Sankey plot of Adephaga and Polyphaga ALGs**. Each ribbon is proportional to the number of genes shared among the chromosomes in the two ALG being compared, providing visual support for the inferred eight beetle ALGs (Fig. S12). **(D) Repeated evolution of a haploid chromosome number of ten involving the same chromosome in different families**. Oxford dot plot (ODP) comparing two genomes from different superfamilies with 10 haploid chromosome count, *Coccinella septempunctata* (Coccinellidae) and *Nalassus laevioctostriatus* (Tenebrionidae), indicating independent fission involving C2 with different chromosomal breakpoints. Only the relevant chromosomes are highlighted on the ODP plot. **(E) Conservation of beetle ALGs in extant genomes**. ODP between two distantly related beetle genomes from different infraorders, *Elmis aenea* (Elateriformia) and *Mycetophagus atomarius* (Cucujiformia), showing that beetle ALGs have remained remarkably conserved across more than 200 million years of divergence. Each dot represents a BUSCO gene, coloured by its assigned ALGs. **(F) Genome comparisons of lineages with high rearrangement rates**. ODP comparing two genomes from the same superfamily, Chrysomeloidea, showing extensive genome reorganisation between the cerambycid *Aromia moschata* and the chrysomelid *Chaetocnema hortensis*. **(G) Fusion and fission signatures in recent rearrangements**. ODP showcasing the genomes of two closely related cerambycid beetles, *Rutpela maculata* and *Stenurella melanura,* with chromosomal breakpoints from fission and subsequent fusions resembling breakage and joining that is consistent with breakpoints near the centromere. 1, Dytiscidae; 2, Carabidae; 3, Scirtidae; 4, Dascillidae; 5, Buprestidae; 6, Elmidae; 7, Eucnemidae; 8, Lampyridae; 9–10 Elateridae; 11, Cantharidae; 12, Histeridae; 13, Hydrophilidae; 14, Lucanidae; 15, Geotrupidae; 16, Scarabaeidae; 17, Leiodidae; 18–20, Staphylinidae; 21, Dermestidae; 22, Bostrichidae; 23, Ptinidae; 24, Biphyllidae; 25, Cleridae; 26, Melyridae; 27, Bothrideridae; 28, Latridiidae; 29, Endomychidae; 30–31, Coccinellidae; 32, Ripiphoridae; 33, Scraptiidae; 34, Salpingidae; 35, Oedemeridae; 36, Pyrochroidae; 37, Anthicidae; 38, Meloidae; 39, Zopheridae; 40, Melandyridae; 41, Ciidae; 42, Tetratomidae; 43, Mycetophagidae; 44, Tenebrionidae; 45, Silvanidae; 46, Cucujidae; 47, Cryptophagidae; 48, Monotomidae; 49, Kateretidae; 50, Nitidulidae; 51, Anthribidae; 52, Attelabidae; 53, Apionidae; 54, Erirhinidae; 55–57, Curculionidae; 58, Cerambycidae; 59–62, Chrysomelidae: sagrine, eumolpine, Chrysomelinae, Galerucinae, respectively. Photo credits: Udo Schmidt and John Hallmén.

We reconstructed a species phylogeny from conserved single-copy orthologues (i.e., BUSCO genes) and used it as the backbone for ancestral linkage group and rearrangement inference. The topology is broadly consistent with prevailing Coleoptera phylogenies (*29*), with minor differences in the placement of selected superfamilies and families (Fig. S3). Using a parsimony-based method which progressively reconstructs ancestral linkage groups on a tree implemented in Syngraph (*23*), we inferred eight ancestral linkage groups in the last common ancestor of Coleoptera, hereafter termed coleopteran ALGs (C1–C7 and CX) (Figs. 1B; S4–5; Table S2–4). As CX is part of the sex chromosome in all but one species (*Aleochara curtula*), we infer that it represents the ancestral sex-linked chromosome. Approximately 97% of BUSCO genes (3,603 of 3,729) were assigned to these eight ALGs, which are ordered by size, with CX being the smallest. We found that the previously proposed nine ancestral elements (Stevens elements: SA–SH and SX) (*36*), correspond closely to the ALGs of the suborder Polyphaga (Fig. S6; Table S5). However, in the suborder Adephaga and in the strepsipteran outgroup, the two Stevens elements SA and SF are linked, and we thus infer that they correspond to a single ALG (C1) in the ancestor of Coleoptera (Fig. S7). The robustness of these assignments was supported by 10,000 Syngraph bootstrap replicates, which recovered clear BUSCO co-assignment blocks corresponding to the inferred eight ALGs (Fig. S8).

This ALG reconstruction framework allowed us to study how the ALGs changed across beetle lineages and over time via chromosomal rearrangements (Fig. S9). We found that beetle genomes fall into six broad categories in regard to their genome evolution (Fig. S10): (I) largely syntenic karyotypes differing from the coleopteran ALGs only by fissions ancestral to the entire polyphagan suborder; (II) category I with additional fissions and occasional fusions; (III) fusion-dominated rearrangements; (IV) fission-dominated rearrangements; (V) extensive fissions and fusions with limited gene order reshuffling; and (VI) extensive inter- and intrachromosomal reorganisation involving both changes in karyotype and gene order. Phylogenetic ordination analyses indicate significant phylogenetic structure in chromosomal-rearrangement profiles, with the dominant axis of variation corresponding to a broad continuum of karyotype conservation and reorganisation (Fig. S11).

### Ancient fissions and divergent rearrangement dynamics in the early evolution of beetles

Beetle chromosomes underwent several fissions early in the divergence of Adephaga and Polyphaga, the two major beetle suborders, such that no extant genome in our dataset fully retains the inferred coleopteran ALG configuration (Figs. 1C; S12). In Polyphaga, we inferred an ancient fission of C1 (forming C1a and C1b) on the branch leading to the last common ancestor of the suborder. The resulting derived configuration of nine polyphagan ALGs is still retained in 15 extant species across deeply divergent polyphagan lineages, indicating long-term conservation following an early structural change (Fig. S13). We also identified 16 additional species with nine chromosomes that were inferred to have arisen through different rearrangement histories. In total, a nine-chromosome karyotype evolved independently 16 times.

A chromosome count of *n* = 10 is the most common karyotype in our dataset, and has been reported in over 20% of described beetle karyotypes (*32*), all in Polyphaga. We find that this chromosome count evolved independently at least 36 times (101 species). Of these, 17 transitions (79 species) resulted from a single fission of the polyphagan ALG configuration. This chromosomal configuration most commonly evolved through fission of C2 (forming C2a and C2b) in 15 transitions. In two additional transitions, the rearranged chromosome involved either C1a (forming C1c and C1d) or C4 (forming C4a and C4b), with different breakpoints producing chromosomes with distinct gene compositions (Figs. 1D; S14). In 42 species, this was followed by further rearrangement to form an 11-chromosome karyotype, the second most common configuration in beetles, mostly within Curculionoidea. This configuration evolved 29 times and, again, most commonly involved fission at C1a, C2, or C4 (Fig. S15). Ornstein-Uhlenbeck (OU) (*42*) and bayou (*43*) analyses supported broad constraints around low chromosome numbers, with inferred optima near *n* = 10–11. However, we found no evidence for a universal pull toward *n* = 10, as fewer transitions moved toward *n* = 10 than away from it (91 vs. 134; *p =* 0.998). Instead, *n* = 10 was produced more often than expected in calibrated simulations (31 observed origins; simulated median = 19 [11–29], *p =* 0.0136) and was relatively stable once reached (exit rate: 19.0% vs. 43.0%; Fisher’s exact *p =* 3.0 × 10^-10^; branch-length GLM OR = 0.31, *p =* 2.03 × 10^-8^; permutation *p =* 0.0001) (Fig. S16). We found that chromosome number reduction due to fusion-biased rearrangement was rare. Rare examples include ancient fusions shared by all Cantharidae species (Fig. S17). Thus, the most common chromosome count in Polyphaga is not a single conserved ancestral state, but has been repeatedly formed via fissions of a limited set of large ancestral linkage groups. Chromosome number alone therefore masks distinct evolutionary histories that produced similar karyotypic endpoints.

In contrast, Adephaga exhibits substantially elevated chromosomal dynamism early in its evolution compared to Polyphaga (Figs. S11; S18). We inferred nine ancestral fission events on the branch leading to the last common ancestor of Adephaga, involving all coleopteran ALGs except CX and resulting in 17 linkage groups. The two largest coleopteran ALGs (C1 and C2) were each fragmented into three chromosomes. This early burst of rearrangement was followed by numerous successive fissions and fusions, likely involving chromosomal arm exchange (Fig. S18), resulting in no extant genome retaining the inferred adephagan ALG configuration.

### Highly variable tempo of chromosome evolution across beetle lineages

The ancient fissions inferred on the branches leading to the last common ancestors of Adephaga and Polyphaga have shaped chromosome evolution across beetles and 33% of species show limited additional interchromosomal change (at most three rearrangements) and thus have retained remarkably conserved chromosome-scale synteny over 200 Mya (Figs. 1A, E; S9). A further 26% of species showed moderate rearrangement, with up to nine inferred events. Against this background of high karyotype conservation, we identified at least 21 lineages (41% of species) that experienced more than ten rearrangements. Of these highly rearranged species, 80% (113 species), showed exceptionally extensive chromosomal reorganisation. These include the suborder Adephaga, Staphylininae, one subclade within Curculionidae, and Chrysomelidae (Figs. 1A, F; S18–21). The extent and nature of reorganisation differ among these lineages. In Adephaga and Curculionidae, extensive fissions and fusions have occurred while ancestral linkage group identity has remained largely preserved, suggesting rearrangements involving whole chromosomes or chromosome arms. In other lineages, ancestral domains are extensively intermingled across chromosomes, indicating more complex histories involving nested fusions, fissions and intrachromosomal reshuffling (i.e., fusion-with-mixing (*5*)). The occurrence of high rearrangement rates in both Adephaga and species-rich polyphagan lineages suggests that chromosomal lability is not tied to a single ecology or life-history syndrome, but has been recruited or tolerated in different evolutionary contexts.

### Extensive intrachromosomal rearrangement across most beetle species

To quantify intrachromosomal rearrangements, we reconstructed the ancestral gene order using AGORA (*22*) (Fig. S22) and calculated a gene-order fragmentation index (*FI*; see Methods) to assess how strongly the ancestral gene order is broken up in extant genomes. *FI* showed a strong phylogenetic signal and was positively associated with log-transformed interchromosomal rearrangement counts (PGLS: β = 0.0189, *p =* 1.29 × 10^-7^, λ = 1), indicating that inter- and intrachromosomal rearrangement rates are correlated. Nevertheless, many beetle species with few interchromosomal changes still showed high *FI*, indicating that substantial gene-order reshuffling can accumulate even in lineages with relatively stable chromosome-scale organisation (Figs. 1A; S23; Table S6).

The gene order conservation is highest (low *FI*) in Adephaga and lowest (high *FI*) in Chrysomelidae, Coccinellidae, and Curculionidae (Fig. 1C). To assess whether gene order reshuffling is mostly localised or involves shuffling across ALGs, we additionally assessed intrachromosomal rearrangement with the domain fragmentation index (*DFI*; see Methods). Several pairs of families showed comparable *FI* but substantially different domain fragmentation (Fig. S24; Table S7). For example, Chrysomelidae and Curculionidae had nearly identical *FI* (median difference = 0.001 [-0.071–0.074]), but Curculionidae showed lower domain fragmentation, meaning that they show less shuffling across ALGs (median difference = 0.059 [0.005–0.244]; posterior directional probability = 0.988). In many curculionid species with chromosomal fusions, intrachromosomal rearrangements are largely confined within ancestral chromosomal domains. In contrast, in lineages such as Chrysomelidae and Staphylinidae, ancestral chromosomal domains are extensively shuffled, consistent with repeated cycles of nested fusions, fissions, and gene order reshuffling. Chrysomelidae (leaf beetles) shows both some of the most extensive intra- and interchromosomal rearrangement and some of the greatest variability in rearrangement levels among beetles (Fig. S21). For example, the degree of interchromosomal rearrangement between two species within Chrysomelidae can exceed that observed between species from different beetle suborders (Fig. 1F).

### Chromosomal rearrangement bursts coincide with major beetle radiations

Chromosomal rearrangements are thought to promote speciation, e.g., by decreasing hybrid fitness or reducing recombination between co-adapted genes (*2*, *3*). Chromosomal rearrangement rates varied substantially across beetle lineages with clade-specific rate ratios ranging from 0.08–18.17-fold relative to their background rates. We identified strong increases in rearrangement rate in seven clades, including Lucanidae, Staphylinoidea, Staphylinidae, Cucujiformia, Phytophaga, Curculionidae, and Chrysomelidae. These shifts include the Phytophaga, the clade comprising Chrysomeloidea and Curculionoidea, which represents the largest radiation of plant-feeding beetles and one of the largest radiations of phytophagous animals. We then tested if beetle clades with increased chromosomal rearrangement rates exhibit increased diversification rate using estimates obtained from McKenna et al. 2019 (*29*). We found a positive association between log rearrangement-rate ratio and log diversification-rate ratio (PGLS: β = 0.0373, t = 2.56, *p =* 0.0145; Fig. S25). Thus, clades with larger increases in chromosomal rearrangement rate tended to show larger increases in diversification rate after accounting for phylogenetic non-independence among clades. Several clades showed high diversification rates without strong evidence of accelerated chromosomal rearrangement, consistent with the expectation that diversification can be promoted by multiple processes.

Low extinction rates combined with high speciation rates have been proposed to underlie the diversification of beetles (*27*, *28*, *31*, *44*). Across the phylogeny, shifts in rearrangement dynamics partially coincide with major ecological and phenotypic transitions. In our reconstruction, lineage such as Phytophaga, underwent additional stepwise chromosomal rearrangements during the radiation of angiosperms (Fig. S26). This period also coincides with evidence of horizontal gene transfer associated with the evolution of plant feeding (*29*, *45*, *46*), suggesting a potential link between genomic innovation, ecological transition, and chromosomal reorganisation.

This correspondence does not imply that chromosomal rearrangements alone drive phytophagan diversification. Rather, it suggests that major beetle radiations may have involved coupled changes in ecological opportunity, gene content and chromosomal architecture. In this view, chromosomal rearrangements may have acted alongside plant-associated innovations, including horizontally-acquired plant cell wall-degrading enzymes (*29*), by altering linkage relationships, recombination landscapes or reproductive compatibility during the expansion of specialised herbivorous lineages.

### Chromosome size and centromere architecture shape chromosomal rearrangements in beetles

In lineages with rearrangements, we tested whether chromosomal features predict which chromosomes are most likely to undergo fission or fusion. We first tested whether chromosome size influences rearrangement frequency, as previously observed in butterflies and moths (*20*). We used the number of BUSCO genes as a size proxy (Fig. S27) so that we could also include rearrangements in ancestral karyotypes. By identifying chromosomes involved in fusions and fissions along each branch in the phylogeny, we found that the chromosomes that fused were on average 2.26× smaller prior to the fusion than non-rearranged chromosomes, indicating that small chromosomes are most likely to fuse. In contrast, chromosomes about to undergo fissions were 1.65× larger than non-rearranged chromosomes and 3.73× larger than fusion contributors (LMM: F = 2947, *p* < 2.2 × 10^-16^; Fig. 2B), indicating that large chromosomes are most likely to undergo fission. Fission products that subsequently fused were smaller than fission-only products, with minor and major products being 0.39× and 0.68× the size of their fission-only counterparts, respectively (both *p* < 0.0001; Fig. S28). Similarly, the number of fusions a chromosome was involved in decreased with chromosome size (Poisson GLMM: β = -0.090, χ^2^ = 20.27, *p =* 6.71 × 10^-6^; rate ratio = 0.91), whereas the number of fissions a chromosome was involved in increased with chromosome size (Poisson GLMM: β = 0.396, χ^2^ = 42.96, *p =* 5.59 × 10^-11^; rate ratio = 1.49, Figs. 2A; S29). In addition to fusions involving fission products, one-to-one fusions between non-rearranged chromosomes were also common and predominantly involved smaller non-rearranged chromosomes (Figs. 2D; S27–28). All pairwise combinations of coleopteran ALGs were observed to fuse, suggesting that no ALG pair is strictly constrained from fusing (Fig. S32). These results suggest that either large chromosomes are more likely to undergo fission and small ones to fuse, or that these rearrangements are most likely to become fixed because they maintain roughly even chromosome sizes.

**Fig. 2.**
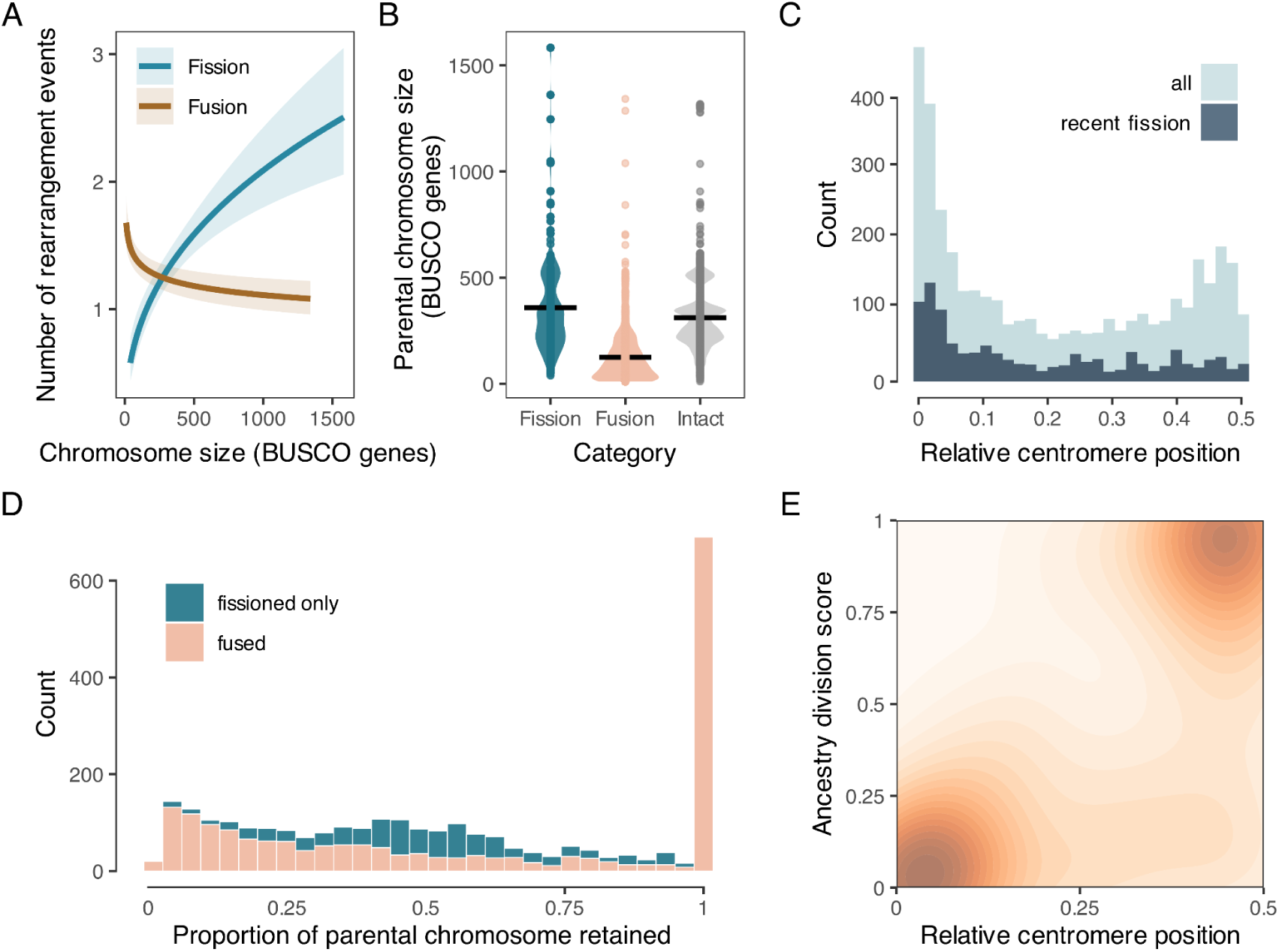
Pattern of chromosomal rearrangements in beetles. (**A**) **Predicted numbers of fission and fusion events per chromosome as a function of chromosome size.** The fitted line shows the predicted relationship between chromosome size (log-transformed) and the number of fission events, estimated using a Poisson generalised linear mixed-effects model (GLMM) with shaded area indicates the 95% confidence interval. (**B**) **Sizes of chromosomes involved in rearrangements compared with those of non-rearranged chromosomes**. The number of BUSCO genes was used as a proxy to chromosome size. For each rearrangement event recorded at an internal node of the phylogeny, chromosome size refers to the inferred size of the parental chromosome before rearrangement, based on ancestral chromosome reconstructions generated using Syngraph. (**C**) **Distribution of centromere positions among chromosomes that have recently undergone fission (dark blue) and all chromosomes (light blue)**. The x-axis indicates the relative position of the centromere along a chromosome, with 0.5 indicating a metacentric position. Recent fissions tend to produce acrocentric chromosomes. (**D**) **Distribution of rearranged chromosome fragments according to the proportion of the parental chromosome retained in each fragment, decomposed by rearrangement history.** The proportion of parental chromosomes retained was calculated as the number of BUSCO genes in each fragment divided by the number of BUSCO genes in its parental chromosome before rearrangement. Values close to 1 indicate that nearly the entire parental chromosome was retained, whereas lower values indicate that only a fraction of the parental chromosome was retained. Blue represents fragments generated by fission that were not subsequently involved in fusion, whereas salmon represents parental chromosome fragments involved in fusion. The high frequency of fusion observations with BUSCO proportions close to 1 indicates that, although most fusion events involved products of previous fissions, direct fusions involving previously non-rearranged chromosomes were also common. (**E**) **Relationship between centromere positions among recently fused chromosomes and the ancestry division score (ADS).** ADS is a measure of ALG separation by centromeres. A score of 0 indicates that distinct ALGs are completely separated by the centromere. This measure is plotted against the relative centromere position, showing that chromosomes in which different ancestries are almost perfectly separated following fusion are nearly always metacentric. These chromosomes may have formed through fusions between acrocentric chromosomes produced by fissions.

Some of the size contrasts among chromosomes involved in fusions and fissions may also be due to compositional differences associated with chromosome length. Smaller chromosomes tended to have higher GC content and coding density than larger ones, with negative size correlations observed in 80.2% and 82.9% of species, respectively (median ρ = -0.600 and -0.643; Fig. S33). Repeat density (including satellite repeats) showed a weaker and more heterogeneous autosomal pattern across superfamilies (median ρ = -0.143; 57.8% negative; Fig. S34). Given its relative length, the sex-linked ALG CX had slightly lower GC (estimate = -0.00648 [-0.0111 – -0.00186], *p* = 0.00608) and coding density (estimate = -0.113 [-0.152 – -0.0747], *p* = 1.56 × 10^-8^) than expected, but significantly lower overall repeat density (estimate = -0.205 [-0.249 – -0.160], *p* = 2.57 × 10^-18^; Fig. S35–36), suggesting that additional evolutionary forces may shape sex-linked chromosome composition.

The repeat density is elevated at both terminal and specific internal regions of chromosomes, likely corresponding to pericentromeric regions (Figs. S37–38). This architecture is important for interpreting rearranged chromosomes, because fusion of chromosomes that have undergone fission can bring repeat-rich terminal or centromere-associated regions into internal positions near the fusion point. In Lepidoptera, repeat landscapes of fused chromosomes can shift over time toward expectations based on their chromosome length. Fissions can instead generate smaller, repeat-rich chromosomes (*20*). However, such changes in repeat content after rearrangement appear less clear in beetles. Although fused chromosomes tended to show a weak initial increase in repeat content followed by reduction over time (β = -0.033, *p =* 0.046), fissions showed no association (β = 0.006, *p =* 0.590) (Figs. S39–40). This could partly be explained by the three times lower chromosome number of beetles compared with Lepidoptera, resulting in weaker chromosome-size contrasts after rearrangement. In agreement with cytological observations (*47*), fusions in beetles often involve chromosomes that have already experienced fission, so recently fused chromosomes may inherit repeat-rich regions near the fusion point (Figs. S41–42). These patterns may also reflect differences in centromere organisation between holocentric Lepidoptera and monocentric beetles, which could affect how repeat-rich regions are retained, repositioned, or remodelled after rearrangement. This makes it difficult to distinguish ancestral repeat landscapes from post-rearrangement remodelling. Therefore, while chromosome length likely contributes to compositional differences after rearrangement, repeat-density evolution in beetles may also depend on rearrangement history, lineage-specific repeat dynamics, and the age of the fusion or fission events.

Repeats may also be directly relevant to the origin of rearrangements. Genome size was strongly correlated with total transposable element (TE) content (PGLS: t = 24.64, *p =* 2 × 10^-16^, adjusted R^2^ = 0.6421, λ = 1; Figs. S43–44). In permutation tests, satellite repeats were consistently enriched at both fusion and fission breakpoints, with the strongest signal in fixed 100-kb windows around breakpoints (2.30-fold enrichment for fusions and 2.69-fold enrichment for fissions; *p* < 0.01; Fig. S45). Because satellite repeats are often associated with centromeric and pericentromeric regions, their enrichment at breakpoints suggests that centromeres or their associated repeat landscapes may contribute to chromosomal breakage and fusion in beetles, such as Robertsonian fissions, fusions or translocations seen in several other monocentric lineages (*48–52*). The frequent occurrence of fusions without extensive mixing of chromosomal segments from different ancestral origins suggests that similar centromere-associated processes may operate in beetles. If satellite-enriched breakpoints reflect centromere-associated regions, chromosomes that have recently undergone fission or fusion are expected to show biases in centromere position.

To test this hypothesis, we inferred putative centromere locations in 157 genomes. We found that chromosomes that have recently undergone fission were strongly shifted toward more terminal centromere positions, consistent with an enrichment of acrocentric or telocentric configurations (median relative centromere position: recent fission = 0.126 vs. 0.326–0.420 for other categories; permutation *p =* 2.5 × 10^-4^ for all recent-fission comparisons; Figs. 2C; S46), consistent with Robertsonian fission (e.g., Fig. S47). In contrast, fused chromosomes were mostly metacentric, comparable to background chromosomes, consistent with predominantly Robertsonian fusions (Figs. 2E; S48). Supporting this interpretation, fused chromosomes with putative centromeres located closer to the chromosome midpoint showed stronger separation of distinct ancestral segments on either side of the centromere (Spearman’s ρ = 0.726, *p* < 2.2 × 10^-16^), indicating that the centromere in fused metacentric chromosomes is typically located near the boundary between ancestral chromosome segments. This model is further supported by the enrichment of satellite repeats at the ends of fission-derived chromosomes and at fusion points, suggesting that fusions often involve centromeric or repeat-rich chromosome ends (e.g., Figs. 2E; S45).

### Sex chromosome changes in Coleoptera

Sex chromosomes were identified in 147 of 338 beetle species where a male individual had been sequenced (Table S8). Regardless of whether a species exhibits a highly reorganised genome or a more conserved chromosomal structure, the ancestrally sex-linked ALG CX has remained unchanged and retained the role of the X chromosome in 116 species, while the remaining 31 species represented 25 independent cases of neo-X chromosomes (Fig. 3; Table S9). In 21 cases, CX included additional ancestrally autosomal chromosomal fragments (8), or entire chromosomes (13). We identified three species with two X chromosomes: *Quedius lateralis*, *Cryptophagus acutangulus* and *Philonthus cognatus* (Fig. 3). In all three cases, one of the two X chromosomes is predominantly of CX origin, while the other X chromosome is one of the typically autosomal linkage groups. Multiple X chromosomes can result from Y-autosome fusions (*47*). However, in *C. acutangulus* the situation is more complex: both X chromosomes contain small fractions of ancestry from the other linkage group, consistent with fusion followed by intrachromosomal rearrangement and subsequent fission. Interestingly, in all three cases, the Y shows little homology with either of the X chromosomes, except for two small regions in *P. cognatus* (Fig. S49). Finally, we detected only a single instance of true sex chromosome turnover in *Aleochara curtula*, where the neo-X chromosome consists of C1 segments, while CX is a standalone autosome (Fig. S50).

**Fig. 3.**
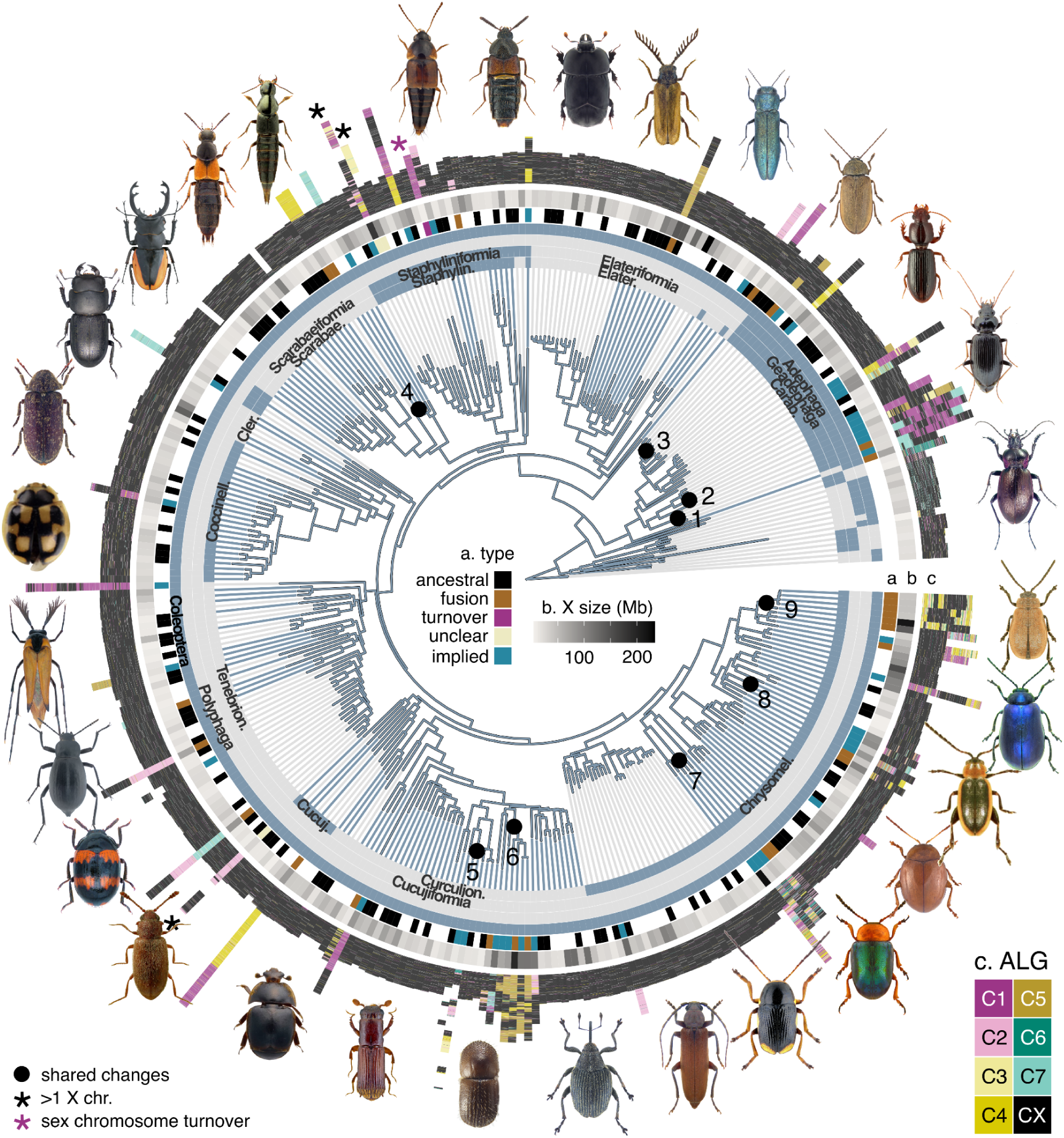
Sex chromosome evolution in beetles. Phylogeny of beetles showing the occurrence of sex chromosome changes. The inner layers, including the tip extension lines, indicate taxonomic assignments across taxonomic ranks, from family, as shown by alternating shades of blue of the phylogeny tip extension lines, to superfamily, infraorder, and order in the outermost layer. The circular layers above it show (a) the type of sex chromosome changes if sex chromosomes are characterised, or “implied” if sex chromosomes are unknown, but the canonical beetle sex chromosome CX rearranged (b) X chromosome size, and (c) painting of the X (if known) or CX-associated (if implied) chromosomes using ALGs. The painting is at single-gene resolution, with each gene represented by an individual stacked line. For species with more than one X or CX-assicuated chromosome, the X chromosomes are separated by white space. Numbered black circles on the tree indicate sex chromosome transitions shared among multiple species of the same subgenus (3. *Pterostichus*), genus (1. *Carabus*; 2. *Nebria*; 4. *Dorcus*; 5. *Euwallacea*; 7. *Cryptocephalus*; 8. *Longitarsus*), two genera (9. *Diorhabda* + *Galeruca*), or a subfamily (6. Ceutorhynchinae). The beetles shown in the plot are representative examples of species with sex chromosome changes. Photo credits: Udo Schmidt, Lech Borowiec, Yves Bousquet, David Ignace, Guillaume Jacquemin, Ilya Zabaluev, Sang Woo Jung, David Crossley (UK Crown), and Damian F. Gomez.

In addition to the 25 transitions with genomically confirmed X chromosomes, we identified 30 additional transitions in which the canonical beetle X chromosome (CX) rearranged, of which 27 represent fusions. In three transitions, the CX underwent fission. *Blaps rhynchoptera* features a simple fission of CX (Fig. 3) (*53*). As previously reported (*41*), two members of the genus *Euwallacea* show multiple fissions of CX into three and five chromosomes, respectively, which do not have clear 1:1 correspondence with each other indicating independent fission events. *Sphaeroderma testaceum* shows a combination of fission of CX and fusion of both halves to an autosome. In all these cases, it is unclear if the CX-derived chromosomes are involved in sex determination as the genomes were produced from female individuals.

We found no clear preferential fusion of CX with specific autosomal ALGs across all 55 transitions (Fig. S32). The unfused CX chromosomes are often acrocentric, in strong contrast with the typically metacentric neo-X chromosomes. The conserved acrocentric nature of CX chromosomes may promote frequent formations of neo-X chromosomes. However, despite relatively frequent CX-autosome fusions, CX fuses less often than autosomes after accounting for size (odds ratio = 0.23 [0.15–0.36], χ^2^ = 52.66, *p =* 3.97 × 10^-13^; Fig. S51A).

Most of the sex chromosome changes were observed in a single species, with only eight transitions shared by at least two species. The oldest change is a fusion of CX to C5 in the common ancestor of the subfamily Ceutorhynchinae. The remaining seven shared changes occurred at the genus or sub-genus level. We tested whether X-linked rearrangements were disproportionately restricted to recent branches of the phylogeny. In branch-length-weighted simulations, X-linked fissions and fusions were both enriched on terminal branches relative to a null model in which events occur uniformly along the tree. In contrast, autosomal rearrangements were not enriched on terminal branches under the same null model (Fig. S51B). Lack of persistence over deep evolutionary time suggests that sex chromosome rearrangements may be selected against in the long term, for example through reduced lineage persistence or increased extinction risk. A similar pattern has been observed in Lepidoptera, where sex chromosomes show no fissions but frequent, recent fusions between the Z chromosome and autosomes (*20*). This suggests that limited long-term persistence of sex chromosome rearrangements may be widespread, although it contrasts with Diptera, where the ancestral X chromosome is a dynamic chromosome (*40*).

Fusion of largely intact X chromosomes with autosomal segments results in disproportionately large X chromosomes relative to autosomes, even when compared with autosomes derived from fusion events (Fig. S52), thereby exacerbating the size contrast with the typically strongly reduced Y chromosome (Fig. S53). Across species with X-autosome fusion, we found only eleven putative neo-Y chromosomes, suggesting that the Y chromosome might degenerate extremely fast. The Y chromosomes in beetles overall show very few homologous genes (Fig. S54), which is in agreement with previously reported absence of sex-determining genes on the Y (*54*) and highlights its dispensability (*55*). This dispensability of the Y chromosome suggests an X:A ratio mechanism of sex determination (*54*).

The X chromosome evolution in Coleoptera shows both similarities and differences from other insect orders. In Diptera (*40*), Lepidoptera (*20*) and Coleoptera, sex chromosomes change most frequently through fusion, but only in Coleoptera do these fusions frequently involve small fragments of various ancestral origins. There is only one known case of sex chromosome turnover in Coleoptera and none in Lepidoptera, while in Diptera, multiple sex chromosome turnovers, sometimes shared by entire families, have been observed (*40*, *56*). Finally, acquisition of multiple X/Z chromosomes through Y/W fusion seems to be present in all three orders, but more frequently in Lepidoptera and Coleoptera than in Diptera, consistent with the cytological literature (*32*). Whereas all coleopteran CX fusions are very recent at most at subfamily level, in both Lepidoptera (*20*) and Diptera (*40*), some sex chromosome changes are old, occurring in the common ancestors of families, or even infraorders.

### Deep conservation of the ancestral holometabolan X chromosome

The origin of the insect X chromosome is thought to predate the origin of insects and to have persisted for more than 450 million years (*57*). In line with this hypothesis, we find that the conservation of the X chromosome extends beyond Coleoptera. The beetle ancestral X (CX) is exclusively and consistently found within X-linked chromosomes of outgroup taxa, with lineage-specific fusions observed in twisted-wing parasites (Strepsiptera) and lacewings (Neuroptera). This pattern suggests that CX might represent the closest approximation of the ancestral X chromosome of Neuropteroidea, which also includes alderflies (Megaloptera) and snakeflies (Raphidioptera) (Fig. S55). Using recently reconstructed ancestral linkage groups in Lepidoptera (*20*) and Diptera (*40*), we tested whether CX represents the ancestral X chromosome of Holometabola (clade of all insects that undergo metamorphosis). We find that a substantial proportion of the dipteran sex-linked ALG (D6) is homologous to C7 and CX (Fig. 4A), consistent with previous suggestions that the reduced X chromosome in Diptera represents a derived state associated with increased sex chromosome turnover (*57*). Previous work proposed that X chromosome reduction occurred after the divergence of flies (Diptera) and the common ancestor of scorpionflies (Mecoptera) and fleas (Siphonaptera) (*58*). Our results instead suggest that this reduction occurred even earlier, likely before the origin of Antliophora (flies, scorpionflies and fleas). Although the X chromosomes of the scorpionfly *Panorpa germanica* and the flea *Ctenocephalides felis* retain homology to CX, this homology comprises only a small fraction of the chromosome (Fig. S56). These findings support the hypothesis that CX represents the ancestral X chromosome of Holometabola, and that the reduction of X chromosome happened in the common ancestor of flies, scorpionflies, and fleas, with lineage-specific fusions occurring in scorpionflies and fleas.

**Fig. 4.**
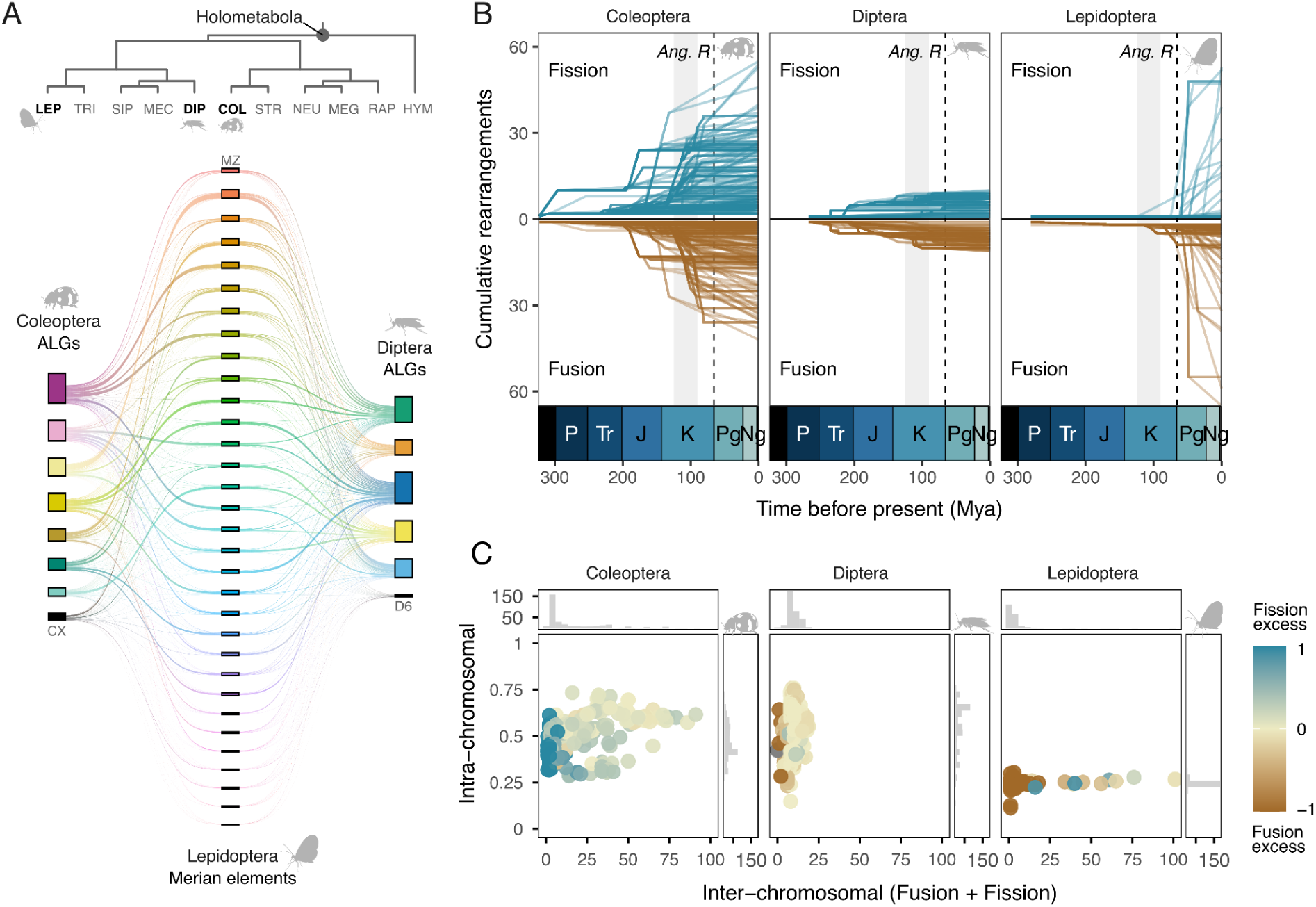
Distinct patterns of chromosomal evolution across three major insect orders. (**A**) **Cross-painting of ancestral elements among coleopteran ALGs, dipteran ALGs, and lepidopteran ALGs (Merian elements).** Phylogeny of Holometabola rooted with Hymenoptera (HYM) showing the relationships among the compared orders. RAP, Raphidioptera; MEG, Megaloptera; NEU, Neuroptera; STR, Strepsiptera; COL, Coleoptera; DIP, Diptera; MEC, Mecoptera; SIP, Siphonaptera; TRI, Trichoptera; LEP, Lepidoptera. Each coloured box represents an ancestral element, and each line represents an orthologous gene linking its assignments across orders. The thickness of the ribbons connecting elements is therefore proportional to the number of orthologous genes shared between them. Elements are coloured according to the coleopteran ALGs and the original descriptions of the lepidopteran Merian elements and dipteran ALGs. Sex-linked ancestral elements are indicated as CX, D6, and MZ. Elements are ordered from top to bottom as C1–C7 and CX for Coleoptera, D1–D6 for Diptera, and MZ followed by M1–M31 for Lepidoptera. (**B**) **Temporal dynamics of chromosomal evolution (fusion and fission) in beetles, flies, and butterflies.** For each species, cumulative trajectories are shown separately for fissions above zero and fusions below zero, from the origin of each order to the present. Rearrangement events occurring along branches between parent and descendant nodes were accumulated at the descendant nodes, and node ages were calibrated using published divergence-time estimates for major taxonomic groups. Ang. R, Angiosperm radiation; dotted lines, K-Pg boundary. (**C**) **Characteristics of chromosomal rearrangements across the three insect orders.** For each species, the total number of interchromosomal rearrangements, defined as the sum of fusion and fission events since the origin of each order to the present, is plotted against the intrachromosomal rearrangement index. Points are coloured according to the difference between the numbers of fissions and fusions, with positive values indicating a fission excess and negative values indicating a fusion excess. Marginal histograms show the distributions of interchromosomal and intrachromosomal rearrangements within each order.

### Contrasting modes of genome structure evolution across major insect orders

As detailed above, our study confirms earlier findings (*57*) that the ancestral X chromosome of insects is remarkably conserved. Next, we tested whether autosomes show similarly high levels of conservation across insects by comparing chromosomal rearrangement dynamics among beetles, flies and butterflies and moths. These insect orders represent three of the four most species-rich lineages, each characterised by major shifts in diversification rates (*59*). A first comparative insight from 12 insect genomes found extensive chromosomal rearrangements and suggested only minimal constraint on gene order except for specific functional clusters (*60*). This could imply that macro-syntenic relationships are limited but can still be detectable across deeply diverged insect lineages.

By identifying homologous BUSCO genes assigned to ALGs across beetles, 340 flies (*40*), and 210 Lepidoptera (*20*), we find that despite large differences in ALG number, genes belonging to the same linkage group in one order tend to remain associated in others (Figs. 4A; S57). This pattern is particularly apparent when viewed from the perspective of Lepidoptera ALGs (31 Merian elements), where most elements correspond to a single ALG in Diptera (6 ALGs) and Coleoptera (8 ALGs) (Fig. S58). In contrast, there is no clear one-to-one correspondence between coleopteran and dipteran ALGs, suggesting extensive large-scale rearrangements along the branches separating these three orders. As additional high-quality genomes and reconstructed ancestral linkage groups become available across other insect orders, resolving the ALGs of Insecta should become increasingly feasible.

We found strong differences in the temporal dynamics of chromosomal rearrangement across insect orders (Fig. 4B). In beetles, chromosomal evolution reflects a combination of the retention of older stepwise rearrangements and the accumulation of more recent events, resulting in cycles of relative stasis punctuated by bursts of change (Fig. S59). In contrast, most rearrangements in flies reflect the fixation of older, stepwise events, whereas butterfly and moth genomes are predominantly conserved and show recent rearrangements in only a few lineages. We also observe systematic differences in the types of rearrangements involved (Fig. 4C). Beetles show a bias toward fission, butterflies and moths toward fusion, and flies exhibit an intermediate pattern. Gene order rearrangement is much more common in both beetles and flies (particularly inversions, in line with, e.g., (*61*, *62*)), than in butterflies and moths whose genomes show remarkable conservation of gene order, deviating from patterns observed in most other insect taxa (*35*).

### Conclusions

Analysing 338 beetle (Coleoptera) genomes and comparing them to 340 fly and 210 butterfly and moth genomes, revealed that the exceptionally species-rich order of beetles shows both high rates of persistent interchromosomal rearrangements and intrachromosomal rearrangements. Yet, we find high variation among beetle lineages with some showing no rearrangements since the ancestral rearrangements in the Adephaga and Polyphaga ancestry, and others showing exceptionally high rates of rearrangements. The association with high rearrangement rates and speciation rates suggests that chromosomal fissions and fusions might contribute to speciation by reducing hybrid fitness. The link of plant-eating beetles showing particularly high rates of rearrangements could also indicate that they take up genotoxic plant compounds accelerating the rearrangement rates. Inferring the causality between these associations will be promising follow-up studies.

Thanks to openly-accessible genomes by the Darwin Tree of Life Project (*37*) and other projects, we could compare three major insect orders, revealing which coleopteran patterns are likely general across insects or order-specific. For instance, the association of small chromosomes with fusions and large chromosomes with fissions seems to be a general pattern across Coleoptera and Lepidoptera. We find that the X chromosome of beetles and the Z chromosome almost never undergo fission, suggesting a very strong constraint, perhaps due to dosage compensation, whereas in Diptera, the ancestral X chromosome is less constrained. On the other hand, fusions of CX are all very recent in Coleoptera, whereas they can be older in Lepidoptera and Diptera, showing different constraints on fusions and fissions. The rates and particularly the temporal patterns of chromosomal rearrangements vary greatly across orders. Coleopteran chromosomal fissions and fusions seem to persist longer, leading to a steady increase in rearrangements over time in many lineages, whereas in Lepidoptera, rearrangements are restricted to recent clades and in Diptera, fusions and particularly fissions less common, although some of them are shared by very large numbers of species. In terms of intrachromosomal gene order reshuffling, likely due to inversions or translocations, beetles show a similarly wide variation across lineages as Diptera, whereas Lepidoptera show very conserved gene order. With the upcoming large-scale comparative publications of vertebrates by VGP (*63*) and much more up-scaled analyses of lepidopteran genomes by Project Psyche (*24*), we will be able expand these comparisons across animals and with additional upcoming initiatives on other phyla across the tree of life. These comparisons will also allow us to identify drivers of the differences among orders, such as differences in meiosis, chromatin structure or reproduction.

## Materials and Methods

### Chromosomal-level genome assemblies

We used publicly available chromosomal-level genome assemblies for Coleoptera and selected outgroups available in NCBI up to 7 February 2026. Assemblies were downloaded using NCBI Datasets (v.16.13.0) (*64*). The final dataset comprised 354 assemblies, representing 338 coleopteran species and 16 outgroup species. General assembly information and associated references are provided in Table S1. From an initial set of 340 coleopteran genomes, two assemblies were excluded after quality assessment: *Laparocerus anagae* and *Pogonus chalceus* (*65*) because the X chromosome was largely absent from both assemblies. A large proportion of the retained assemblies (68%) were generated by the Darwin Tree of Life Project (DToL) (*37*). For all DToL genomes, the primary assembly was used for downstream analyses.

The dataset included representatives of 18 of the 25 recognised superfamilies. No genomes were available for the early-diverging suborders Archostemata and Myxophaga. Sixteen species were included as outgroups for ancestral reconstruction. These included the strepsipteran genomes *Stylops aterrimus* (*66*) and *Xenos peckii* (*67*), representing the closest outgroups to Coleoptera, together with representatives of Megaloptera, Neuroptera, and Raphidioptera.

We tested the association between genome size and chromosome number using phylogenetic generalised least squares (PGLS). We fitted PGLS models using ‘pgls’ in the R package ‘caper’, with chromosome number modelled as the response variable and log10-transformed genome size as the predictor. Pagel’s λ was estimated by maximum likelihood to model phylogenetic covariance.

To assess the genome quality, we ran BUSCO (v.6.0.0) analyses using metaeuk mode and the coleoptera_odb12 dataset (*68*, *69*). The coleopteran genomes had an average BUSCO completeness score of 96.22 ± 3.69%, whereas the outgroup genomes 79.05 ± 13.84%. Most BUSCO genes were located on chromosomes, with an average of 99.28 ± 0.06%. The summary statistics of these BUSCO runs are included in Table S1. We combined BUSCO results and genome coverage information to manually assess whether chromosome-level sequences were labelled as unlocalised scaffolds, as indicated by unusually high numbers of complete single-copy BUSCO genes, and whether scaffolds likely belonging to the same chromosome had been treated as separate chromosomes. These manual adjustments of the genomes are explained in Supplementary Text 1. In addition, we also used this information to identify sex-linked chromosomes and to correct the assignment of the marked X/Y chromosomes in the genome publication page when necessary (see below).

### Identifying sex-linked chromosomes

To investigate sex chromosome evolution, X and Y chromosomes were identified using a combination of sex annotations from genome metadata, karyotype information from the Coleoptera Karyotype Database (*32*), and read coverage analyses based on raw reads data. Reads were mapped to the genome using pbmm2 (v.1.13.1) (*70*) for PacBio HiFi reads, or HISAT2 (v.2.2.1) for RNA-seq data (*71*). Coverage depth was calculated using mosdepth (v.0.3.3) (*72*), and sex chromosomes were inferred based on reduced coverage in males (i.e., approximately half the depth observed for autosomes). In cases where genomes were assembled from female specimens, the X chromosome could not be confidently identified and was annotated accordingly throughout our analyses. We also identified several instances where the sex indicated in genome metadata was inconsistent with the sex inferred from coverage patterns. In such cases, we relied on the latter (coverage-based inference) as the more accurate source of information. The list of sex-linked chromosomes identified and applicable corrections on the published information are available in Table S8. The table also includes the known sex-determination system for each species, where available, alongside the relevant cytological literature (*32*).

### Reconstructing backbone phylogenetic tree

Single-copy orthologues were identified within each genome using BUSCO (v.6.0.0), in ‘metaeuk’ mode and employing the coleoptera_odb12 database (*68*). We used the custom script busco2fasta.py (https://github.com/lstevens17/busco2fasta) to find 3,729 BUSCO genes present in at least 90% of the genomes. These protein sequences were aligned using MAFFT (v.7.525) (*73*) and trimmed with trimAl (v.1.4.1) (*74*) with parameters -gt 0.8 -st 0.001 -resoverlap 0.75 -seqoverlap 80. A total of 3,153 alignments passed the thresholds. The filtered alignments were concatenated into a supermatrix using catfasta2phyml.py (https://github.com/nylander/catfasta2phyml) and used to reconstruct a phylogenetic tree using IQ-TREE (v.3.0.1) (*75*) under the Q.insect+I+G4 substitution model, selected as the best-fitting model using ModelFinder (*76*, *77*), incorporating 1,000 ultrafast bootstrap replicates. The resulting tree was rooted using *Xanthostigma xanthostigma* (Raphidioptera) as the outgroup.

We compared our reconstructed tree with a widely accepted Coleoptera phylogeny (*44*). The topology of our tree was largely consistent with this published tree for the families included in our dataset, except for differences in the placement of Coccinelloidea and Tenebrionoidea. Additional differences were observed in the placement of some families within Tenebrionoidea and in the position of Cryptophagidae within Cucujoidea.

### Inferring ancestral linkage groups

Ancestral linkage groups (ALGs) in Coleoptera were inferred using Syngraph (*23*), based on single-copy orthologues identified by BUSCO and a reconstructed phylogeny comprising 338 coleopteran and 16 outgroup genomes. BUSCO genes located on unlocalised scaffolds and the Y chromosome were excluded from this and all subsequent analyses. Some unlocalised scaffolds from a few species were included when there was evidence that they represented unannotated chromosomes (Supplementary Text 1). As a consequence, manual corrections to the inferred rearrangement events were required for these species, as described in Supplementary Text 2. Syngraph uses a parsimony-based, syntenic edit-distance framework to reconstruct ancestral linkage groups across a phylogeny and infer inter-chromosomal rearrangements along the tree. It represents genomes using the co-occurrence of orthologous markers on the same chromosome or linkage group, without using their order, and infers events such as fissions, fusions, and optionally reciprocal translocations.

We used a threshold of ten orthologues, -m 10, representing the minimum number of markers required to support a rearrangement, together with an algorithm restricted to detecting fusion and fission events, -r 2. The -m parameter was selected following an empirical evaluation of ALG inference across major internal phylogenetic nodes using values ranging from 1 to 100 (Figs. S4–S5). In general, Syngraph produced meaningful results when -m was neither too low nor too high. At values below -m 5, the number of inferred ALGs decreased substantially, often producing uninformative solutions in which only a single ALG was recovered (Fig. S4). Between -m 5 and -m 25, the number of inferred ALGs was relatively stable, with seven ALGs recovered at -m 5–6 and -m 12–23, and eight ALGs recovered at -m 7–11 and -m 24–25. Beyond -m 25, the number of inferred ALGs fluctuated more strongly, coinciding with a decrease in the number of markers assigned to each group.

We constrained the likely number of Coleoptera ALGs to eight, corresponding to the stable range recovered at -m 7–11, for two reasons. First, when mapping ALG definitions onto extant genomes, markers located on sex chromosomes were often underrepresented or entirely absent at -m values above 11 (Fig. S4A), despite previous studies showing high synteny conservation across Coleoptera X chromosomes. Second, manual inspection showed that two ALGs that were inferred as distinct at -m 7–11, but merged at -m 4–6, were rarely located on the same chromosome across extant genomes. In the few exceptions, such as Cantharidae, the markers did not intermix and likely reflected chromosomal arm fusion. This observation ruled out -m 4–6 as the optimal range and reinforced our choice of the -m 7–11 range, supporting a model with eight Coleoptera ALGs. We selected -m 10 within this range for downstream Coleoptera ALG inference. The defined Coleoptera ALGs from this inference are available in Table S2.

Using the selected parameters, we also performed manual checks for internal nodes that were variable across the -m 7–11 range. Some nodes produced uninformative rearrangement inferences, erroneously reducing chromosome number by merging multiple chromosomes into one or two inferred linkage groups, which we considered unlikely (Fig. S5). These problematic nodes occurred within Chrysomeloidea, a lineage known to be highly rearranged. We therefore performed a separate Syngraph analysis for Chrysomeloidea using higher -m values to reduce noise from small, likely erroneous rearrangements inferred under lower thresholds. We selected -m 24 as the optimal threshold because it minimised erroneous exclusion of Coleoptera ALG markers while improving the inferred ALG structure based on manual inspection. Rearrangement events inferred from this separate Chrysomeloidea analysis were used to replace the corresponding events from the initial Coleoptera-wide run. The final table listing all ALGs definitions for all internal phylogenetic nodes are available in Table S3, and the inferred rearrangement events in Table S4.

We assessed the robustness of BUSCO assignment to ALGs using 10,000 bootstrap resampling of BUSCO markers. In each replicate, the full BUSCO marker set was resampled with replacement to generate a marker list of the same size as the original set. Syngraph was then rerun using the same parameters as the final analysis (-m 10 -r 2 -a quick). We counted, for each BUSCO pair, the number of replicates in which both markers were assigned to the same ALG, generating a pairwise bootstrap co-assignment matrix. Under bootstrap sampling with replacement, a marker is expected to be sampled in approximately 63.21% of replicates, and a pair of markers is expected to co-occur in approximately 39.96% of replicates. We therefore used the lower-tail 1×10^−6^ quantile of *B*(*n*, 0.3996), where n is the number of bootstrap replicates, as the threshold for high-confidence pairwise co-assignment.

### Comparing inferred ALGs with Stevens elements

We also compared our inferred Coleoptera ALGs with the previously proposed beetle ancestral linkage groups, termed Stevens elements (*36*). Because no publicly available gene or orthologue membership table exists for each Stevens element, and because the methods used to define Stevens elements differ from those used in this study, a direct one-to-one comparison was not possible. However, the Stevens elements were defined using the *Tribolium castaneum* genome assembly Tcas5.2 (*78*), in which 10 of the 11 chromosomes were each assigned to a Stevens element. We therefore assigned all BUSCO genes located on each of these chromosomes to the corresponding Stevens element and used this chromosome-based assignment as an approximate basis for comparison (Table S5). Using this approach, we compared the correspondence between Stevens elements and the Coleoptera ALGs inferred in this study, and evaluated the differences between them to assess whether Stevens elements represent coleopteran ALGs.

### Phylogenetic tests of chromosome-number recurrence and stability

Because *n* = 10 is the most common chromosome number in beetles and was inferred to have evolved independently multiple times, we tested whether this recurrence reflects broader evolutionary constraint around *n* = 10, repeated convergence towards this state, or increased stability once *n* = 10 is reached. To evaluate whether chromosome number showed evidence of constraint around low values, we fitted continuous-trait models of chromosome-number evolution to the species tree (*42*). Tip chromosome numbers were treated as continuous values for this analysis. We compared Brownian motion (BM), Ornstein–Uhlenbeck (OU), and early-burst (EB) models using AICc in R package ‘geiger’ (*42*). We additionally used bayou to fit reversible-jump Bayesian OU models and to evaluate whether chromosome number was consistent with one or more inferred optima (*43*). Analyses were run using the reparameterised OU model. Priors were placed on OU half-life, stationary variance, number of shifts, shift locations, and θ values. Four independent MCMC chains were run for 1,000,000 generations each, with 30% discarded as burn-in. We summarised the posterior distribution of the root/background optimum and calculated the posterior probability that at least one inferred optimum fell within *n* = 10 ± 1.

We next asked whether chromosome-number transitions were biased towards *n* = 10. We used Syngraph-inferred chromosome-number histories to construct a branch-level transition table. For each branch, we recorded the parent chromosome number, child chromosome number, and whether the branch changed chromosome number. We classified directional changes according to whether they moved closer to or farther from *n* = 10. Branches with no change in distance from *n* = 10 were excluded from the toward/away comparison. A one-sided exact binomial test was used to test whether transitions toward *n* = 10 were more frequent than transitions away from *n* = 10.

To test whether *n* = 10 was stable once reached, we compared the frequency of chromosome-number change on branches starting at *n* = 10 with branches starting from all other chromosome numbers. We used Fisher’s exact test to compare exit rates between *n* = 10 and non-10 states. We also fitted a logistic regression with chromosome-number change as the response and starting state as the predictor. To account for branch length, we fitted an additional logistic regression including branch length from the time-calibrated tree: changed_n ∼ start_n10 + branch_length. Where changed_n indicates whether parent and child chromosome numbers differed, and start_n10 indicates whether the branch started at *n* = 10. We compared exit rates among nearby chromosome numbers *n* = 9, 10, 11, and 12. Pairwise Fisher’s exact tests were used to compare the exit rate of *n* = 10 with each nearby state. *P*-values were adjusted for multiple testing using the Benjamini–Hochberg procedure. To evaluate whether the reduced exit rate from *n* = 10 could arise from random placement of *n* = 10 states across branches, we performed a permutation test. The start_n10 labels were randomly permuted across branches while keeping branch lengths and observed chromosome-number changes fixed. For each permutation, we refitted the branch-length logistic model and recorded the odds ratio associated with the permuted *n* = 10 label. The observed odds ratio was then compared with the permutation null distribution. The adjusted permutation *p*-value was calculated as: (sum(perm_or ≤ observed_or) + 1) / (nperm + 1).

Finally, we tested whether *n* = 10 originated more often than expected, we simulated chromosome-number histories using an empirical multi-state transition matrix. The matrix was constructed from Syngraph-inferred parent-to-child chromosome-number transitions, using all observed chromosome-number states rather than a binary *n* = 10 / non-10 coding. No-change transitions were removed from the off-diagonal matrix, and a small pseudocount was added to off-diagonal entries to avoid zero-probability transitions. The transition matrix was scaled so that simulated histories produced a number of chromosome-number changes comparable to the observed branch history. We then simulated chromosome-number histories on the time-calibrated beetle tree using phytools::sim.history (*79*). For each simulated history, we counted the number of branches in which *n* = 10 originated, defined as a transition from a non-10 parent state to an *n* = 10 child state. The observed number of *n* = 10 origins was compared with the simulated null distribution. The adjusted simulation permutation *p-*value was calculated as: (sum(perm_or ≤ observed_or) + 1) / (nperm + 1).

### Reconstructing ancestral gene order

Ancestral gene order in the last common ancestor of beetles was inferred using AGORA (*22*). AGORA input files were prepared with a custom script that converted complete BUSCO gene information from 354 BUSCO tables into AGORA-compatible gene lists, orthology groups, and the required species tree file. The resulting orthology groups and species tree were then used as input for ‘agora-generic.py’. All 3,722 BUSCO genes were included in the reconstruction of the last common ancestor of Coleoptera. Of these, AGORA placed 3,504 genes into 156 contiguous ancestral regions (CARs) (Table S6). Almost all CARs (except 9 CARs) contained orthologues mapping to a single Coleoptera ALG, indicating that the AGORA reconstruction was broadly consistent with the Coleoptera ALGs inferred using Syngraph. However, because the CARs were highly fragmented, we did not use the AGORA output to define the exact gene order within Coleoptera ALGs. Instead, we used the AGORA reconstruction as an approximate ancestral reference for estimating the degree of intrachromosomal rearrangement relative to the most recent inferable ancestral gene-order configuration.

To quantify the degree of intrachromosomal rearrangement relative to the inferred ancestral gene-order reconstruction, we calculated a gene-order fragmentation index (*FI*) for each species (Table S7). FI was defined as:

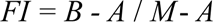

where *B* is the number of observed gene-order blocks on the extant chromosome, *A* is the number of AGORA ancestral units (CARs and singletons) represented on that chromosome, and *M* is the number of informative BUSCO markers. This index rescales fragmentation between 0 to 1, where *FI* = 0 indicates that all informative ancestral adjacencies are conserved, whereas *FI* = 1 indicates complete fragmentation of the ancestral gene order.

To distinguish local gene-order reshuffling from broader mixing of ancestral chromosomal domains, we additionally quantified domain fragmentation index (*DFI*) on fusion chromosomes. Domain fragmentation was calculated from the number of uninterrupted runs of ancestral domains along each extant chromosome. For a chromosome with *D* ancestral domains, *R* observed domain runs and *M* informative markers, domain fragmentation was calculated as:

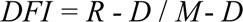

where *DFI* = 0 indicates that each ancestral domain occurs as a single contiguous block, and higher values indicate increasing intermixing of ancestral domains. To reduce sensitivity to assembly noise, ancestral domains represented by fewer than 10 markers or less than 2% of markers on a chromosome were excluded, and isolated short runs were smoothed when flanked by longer runs of the same domain.

We fitted order-specific PGLS models to test the association between intrachromosomal rearrangement (*FI*) and interchromosomal rearrangement, using *FI* as the response and log1p-transformed interchromosomal rearrangement as the predictor. Phylogenetic covariance was modelled using Pagel’s λ, estimated by maximum likelihood. PGLS models were fitted using ‘gls’ in the R package ‘nlme’ with a Pagel correlation structure implemented in ‘ape’ (*80*). Phylogenetic signal in *FI* and interchromosomal rearrangement was assessed using Pagel’s λ and Blomberg’s K with phylosig in ‘phytools’ (*79*).

To test whether lineages with similar *FI* differed in ancestral-domain mixing, we fitted Bayesian phylogenetic beta-regression models using brms (*81*). Domain fragmentation was modelled as the response, with family and scaled FI as predictors, and a phylogenetic random effect based on the species covariance matrix. Pairwise posterior contrasts in predicted *DFI* were then calculated between families. Family pairs were considered to have comparable *FI* when the absolute posterior median *FI* difference was ≤0.05 and then 95% credible interval overlapped zero.

### Ordination of chromosomal-evolution profiles

To summarise broad patterns of chromosomal evolution across species, we performed ordination analyses using summary variables derived from the ALG reconstruction. These variables included ALG integrity scores (AIS), ALG composition scores (ACS), inferred fission and fusion counts, and the proportion of intrachromosomal reorganisation. Variables were centred and scaled prior to ordination to account for differences in measurement scale.

ALG integrity scores (AIS) were calculated as the mean fraction of markers from each Coleoptera ALG retained on individual extant chromosomes. ALG composition scores (ACS) were calculated as the mean fraction of each chromosome composed of markers from a given ALG. AIS captures the degree to which each ancestral linkage group remains intact, whereas ACS captures the degree to which extant chromosomes are dominated by a single ancestral linkage group. To reduce noise from small translocations involving only a few genes, chromosome–ALG combinations were excluded when either the fraction of the ALG represented on that chromosome or the fraction of the chromosome composed of that ALG was below 0.2.

We first used t-distributed stochastic neighbour embedding (t-SNE) to visualise broad similarities among species in the ordination profiles. The t-SNE analysis was performed using the ‘Rtsne’ package in R with two dimensions, a perplexity of 20, 10,000 iterations. To assess whether chromosomal-evolution profiles were structured by shared ancestry, we performed complementary phylogenetic ordination analyses using ‘gm.prcomp’ in R (*82*). These included phylogenetic PCA and phylogenetically aligned component analysis (PACA). Phylogenetic PCA was used to visualise major axes after accounting for phylogenetic non-independence, and PACA was used to identify axes most strongly aligned with phylogenetic signals. We further evaluated the distribution of phylogenetic signals across components using Blomberg’s K and assessed the association between phylogenetic covariance and PACA scores using phylogenetic partial least squares. To test whether chromosomal-evolution profiles differed among families, we used residual randomisation in a phylogenetic RRPP framework with family as the grouping factor.

### Transposable elements, segmental duplications, and gene annotation

Transposable elements (TEs) were annotated for each genome using the EDTA pipeline (v.2.2.2) (*83*) with parameters –anno 1 and –sensitive 1. Segmental duplications were detected using BISER (v.1.4), utilising the repeat-masked genome assemblies. The softmasking of the genomes was done using ‘make_masked.pl’ script from EDTA. Segmental duplications were defined as regions larger than 1 kb, exhibiting at least 90% sequence identity, after excluding bases attributed to known high-copy-number repeats. The genes in each genome were annotated using Helixer (v.0.3.5) (*84*).

### Detecting potential centromeric regions

Candidate centromere regions were annotated using the Nextflow version of the Centromere Analysis Pipeline (CAP; v.1.0) (*85*), that integrates context-tree weighting algorithm (CTW) and distribution of tandem repeats annotated with TRASH2 (https://github.com/vlothec/TRASH_2) along chromosomes to calculate centromeric scores. For each genome, optional inputs of transposable element annotations generated with EDTA (*83*) and gene annotations predicted by Helixer (*84*) were provided for additional genomic context. CAP outputs of centromere prediction scores, repeat-family summaries, and distribution plots of chromosome-wide genomic features were manually inspected to identify candidate centromere-associated repeat regions. Chromosome regions were prioritised as candidate centromeric regions if they had a combination of some or all of the factors: 1. decreased CTW, 2. change in GC-content (drop or increased), 3. clustering of specific, preferably shared across all or most chromosomes, satellite repeats, 4. peak in density of transposable elements, and 5. drop in gene content. In cases where no clustering of satellite repeats and/or changes in GC-content were observed, centromeres were assumed to be TE-based and the approximate location was assumed to be around a peak of TEs and a drop in gene density.

### Analysing potential constraints in chromosomal rearrangement: chromosome size

To compare chromosome size across rearrangement outcomes, chromosomes were classified as fissioned, fused, or non-rearranged. Fissioned chromosomes represent parental chromosome size before splitting, fused chromosomes represent parental chromosome size participating in fusion events, and non-rearranged represent chromosomes with no fission or fusion on the corresponding branch. Differences in chromosome size among the three classes were tested using a linear mixed model on log-transformed BUSCO counts, with rearrangement class as a fixed effect and parent and child nodes as random intercepts. Pairwise contrasts among classes were estimated using Tukey-adjusted post hoc comparisons with emmeans, and effect sizes were reported as back-transformed size ratios.

Fusion can involve previously non-rearranged chromosomes and also following fission events. To compare the difference between chromosomes that followed subsequent fusion after fission with only fission, we tested using linear mixed models on log-transformed BUSCO counts. The model included fission-product rank, fate, and their interaction as fixed effects, with parent and child nodes as random intercepts. To compare the difference in chromosome size between fusing non-rearranged chromosomes with non-rearranged chromosomes, we compared between non-fissioned fusion contributors and non-rearranged chromosomes, with the parent node included as a random intercept. Post hoc contrasts were estimated using emmeans, and effect sizes were reported as back-transformed size ratios.

We tested whether chromosome size predicted rearrangement intensity using Poisson generalised linear mixed models implemented in ‘glmmTMB’. Chromosome size was measured as BUSCO count and log-transformed before analysis. Fission events were defined as parental chromosomes contributing to more than one descendant chromosome, and fusion events as descendant chromosomes receiving contributions from more than one parental chromosome. For fission, we modelled fission intensity as a function of log-transformed chromosome size, with parent and child nodes included as random intercepts. For fusion, we modelled fusion intensity as a function of log-transformed contributing chromosome size, with parent node, child lineage, and fusion event identity included as random intercepts. Model significance was assessed using likelihood-ratio tests comparing full models with null models lacking the chromosome-size term but retaining the same random-effect structure. Effect sizes are reported as model coefficients and rate ratios. Models were checked for convergence and overdispersion.

To test whether fusion partners were non-random with respect to ancestral linkage group identity, we recorded all unordered pairs of ALG categories contributing to each fusion event. ALG categories were defined as C1–C7 and CX; contributors classified as mixed-origin were excluded from the primary test because they do not represent a single ancestral linkage group. We generated a null distribution by permuting ALG labels among fusion contributors while preserving the number of contributors per fusion event and the overall abundance of each ALG category. For each permuted dataset, unordered co-fusion pair counts were recalculated. Observed pair counts were compared with 10,000 permuted datasets to estimate expected counts, enrichment values, and empirical *p*-values. *P*-values were adjusted across pairwise comparisons using the Benjamini–Hochberg procedure. Enrichment was reported as log2(observed/expected), with values below zero indicating depleted pairings and values above zero indicating enriched pairings.

### Analysing potential constraints in chromosomal rearrangement: genomic features

Chromosome size is often correlated with genomic feature density. To test whether genomic features were associated with chromosome size, we calculated feature densities in 100 kb windows and summarised them at the chromosome level. Windows were generated using ‘bedtools makewindows’, and GC content was calculated for each window using ‘bedtools nuc’ in BEDTools v.2.31.1 (*86*). Coding density was calculated as the proportion of each window covered by CDS annotations. For each genome, the longest isoform per gene was retained using AGAT (https://github.com/oushujun/EDTA). CDS features were extracted from the filtered GFF3 file, overlapping CDS intervals were merged using bedtools merge, and coverage across 100 kb windows was calculated using ‘bedtools coverage’.

Repeat density was calculated from EDTA repeat annotations using ‘bedtools coverage’ after merging overlapping repeat intervals. Repeat classes were grouped into major subclasses, including Gypsy, Copia, Other LTR, LINE, SINE, Penelope/DIRS, TIR, Helitron, Polinton, Other DNA, YR/Crypton, and unclassified or other repeats. Satellite repeats were identified from TRASH (*87*) annotations and analysed separately. Total repeat plus satellite density was calculated by merging EDTA (*83*) repeat intervals with satellite-repeat intervals before estimating coverage in each window. For each species and chromosome, we calculated mean GC content, coding density, repeat plus satellite density, individual repeat-class density, and satellite repeat density across 100 kb windows. Chromosome length was expressed as the proportion of total assembled chromosome length within each species. In addition, we calculated feature densities in 100 proportional windows per chromosome to compare chromosome-position landscapes across chromosomes. All associated summary statistics tables resulting from these analyses are available in Table S10–13).

We included only chromosomes with clear ancestral linkage group assignments. Chromosomes were assigned to an ALG when at least 80% of mapped BUSCO markers belonged to a single ALG and the second-largest ALG contribution was ≤5%. Chromosomes were retained only when the corresponding ALG was represented once per species, except for C1, which was allowed to occur as two chromosomes because of ancient fissions and was split into large and small C1 classes based on marker-content similarity. CX chromosomes were identified from ALG assignment and excluded from autosomal trend tests, but plotted separately for comparison.

For each species, we tested whether proportional chromosome length was associated with chromosome-level feature density using Spearman rank correlations across retained autosomal chromosomes. Tests were performed separately for GC content, coding density, total repeat plus satellite density, each repeat class, and satellite repeats. Species with fewer than four retained autosomal chromosomes were excluded from these tests. *P*-values were adjusted across species using the Benjamini–Hochberg procedure. For repeat-class analyses, correction was applied separately within each repeat class.

To assess whether repeat-density deviation in rearranged chromosomes varied with rearrangement age, we used the age of the ancestral reference node associated with each retained rearrangement. Rearrangements were restricted to cases in which the closest comparator species belonged to the same family as the focal species. For fusion events, repeat-density deviation was calculated as the repeat density of the whole fused chromosome minus the repeat density of the closest matched chromosome corresponding to each ancestral component. For fission events, deviation was calculated as the length-weighted repeat density of the fissioned chromosome unit minus the repeat density of the closest matched chromosome. Temporal trends were tested using linear mixed models with species as a random effect. The fusion model included ancestral reference-node age and fusion component size class as fixed effects. The fission model included ancestral reference-node age and the number of fissioned chromosome pieces as fixed effects.

### Analysing potential constraints in chromosomal rearrangement: repeat enrichment analyses

We tested whether repeat content was enriched near inferred chromosome rearrangement regions using same-chromosome permutation tests. Repeat density was measured from 100 kb genomic windows. For total repeat enrichment, we used the combined repeat + satellite density track. For repeat-class enrichment, we repeated the analysis separately for satellite repeats and each transposable-element class, including Gypsy, Copia, Other_LTR, LINE, SINE, Penelope_DIRS, TIR, Helitron, Polinton, Other_DNA, YR_Crypton, and Unclassified_or_other.

For each observed rearrangement-associated interval, we calculated repeat density as the length-weighted mean of all overlapping 100 kb windows. To generate a null distribution, each observed interval was randomly relocated within the same species and chromosome while preserving its interval length. For each permutation, one matched random interval was sampled for every observed interval, and the length-weighted mean repeat density across all intervals was calculated. This was repeated 1,000 times, with p-values calculated as: *p* = (1 + number of null values ≥ observed value) / (N + 1), where N is the number of permutations. Repeat-class Repeat-class p-values were corrected using the Benjamini-Hochberg procedure within each interval definition.

For fusion events, we tested repeat enrichment using two classes of breakpoint intervals. First, we defined a BUSCO-marker interval as the region between the nearest syntenic BUSCO markers assigned to the two fused ancestral components. Only simple fusion events with a reliable fusion component boundary were included. We defined fixed windows around the inferred fusion boundary at ±100 kb, ±500 kb, and ±1 Mb. For fission events, we used the closest non-rearranged comparison chromosome to infer the likely breakpoint-bearing end of each fission product. Shared BUSCO markers were mapped between each focal fission product and the closest non-rearranged chromosome. The order of fission products along the closest chromosome was used to identify terminal pieces, and marker-order correlation was used to account for possible orientation reversals. Predicted breakpoint-bearing ends were then tested using a BUSCO-marker terminal interval, defined from the terminal BUSCO marker to the chromosome end, and fixed terminal windows of ±100 kb, ±500 kb, and ±1 Mb. As a control for general chromosome-end repeat enrichment, we also tested both ends of all fission products using the same fixed window sizes.

Enrichment was calculated as the difference between the observed mean repeat density in rearrangement-associated intervals and the mean repeat density expected from same-size random intervals sampled from the same species and chromosome. For fission events, we additionally calculated breakpoint-specific enrichment by subtracting the enrichment observed at both fission-product ends from the enrichment observed at the predicted breakpoint-bearing ends.

### Analysing potential constraints in chromosomal rearrangement: centromere

To identify rearrangements consistent with centric fusion, we quantified how strongly the centromere separated ancestral chromosome identities within each extant chromosome. For chromosomes composed primarily of two ancestral linkage groups, A and B, we calculated an ancestry division score, ADS, using the proportional representation of each ancestry on either side of the centromere:

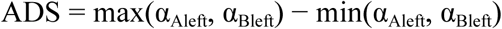

or equivalently,

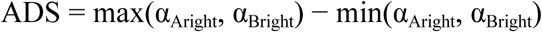

where α_Aleft_ and α_Bleft_ are the proportions of markers assigned to ancestral linkage groups A and B located to the left of the centromere, and α_Aright_ and α_Bright_ are the corresponding proportions to the right of the centromere. Higher ADS values indicate stronger separation of ancestral identities on opposite chromosome arms, as expected under centric fusion, whereas lower values indicate greater intermixing of ancestral identities across the centromere. Predicted centromere positions are available in Table S14.

### Identifying changes and constraints of sex chromosome rearrangement

We tested whether X-linked chromosomes differed from autosomes in their involvement in chromosome fusions using a binomial generalised linear mixed model implemented in ‘glmmTMB’. Fusion involvement was modelled as a binary response for each chromosome–branch comparison. The model included log-transformed chromosome size and X-linkage status as fixed effects, with parent and child nodes included as random intercepts. The contribution of X-linkage was assessed by comparing models with and without the X-linkage term using a likelihood-ratio test, and effect size was reported as an odds ratio. For visualization, average model-adjusted probabilities of fusion involvement were obtained by predicting each observed chromosome–branch record under autosomal and X-linked status while retaining chromosome size and fitted parent and child random effects, and then averaging predictions across records.

To test whether sex-chromosome changes were unusually concentrated on shallow branches of the phylogeny, we combined reconstructed sex-chromosome fissions and fusions into a single category and counted each affected branch once. Autosomal-only rearrangement branches were defined as branches containing reconstructed fissions or fusions but no sex-chromosome change. We then used a branch-length-weighted null model with 100,000 replicates, in which the observed number of affected branches in each category was randomly assigned to phylogenetic branches with probabilities proportional to branch length. For each replicate, we recorded the number of assigned changes occurring on terminal branches. One-sided *p*-values were calculated as the proportion of simulations with at least as many terminal-branch changes as observed.

To test whether CX chromosomes differed from autosomes in genomic composition after accounting for chromosome size, we fitted linear mixed models for each genomic feature. The response variable was the logit-transformed chromosome-level feature density. Proportional chromosome length and CX identity were included as fixed effects, with species included as a random intercept. Models with and without the CX term were compared using likelihood-ratio tests. This analysis was applied to GC content, coding density, total repeat plus satellite density, individual repeat classes, and filtered satellite repeat density. *P*-values for CX effects were adjusted using the Benjamini–Hochberg procedure within feature groups. Negative CX coefficients indicate lower feature density on CX chromosomes relative to autosomes of comparable proportional chromosome length, whereas positive coefficients indicate higher feature density on CX chromosomes.

### Testing association of rearrangement rates and diversification rates

To test whether increases in chromosomal rearrangement rates were associated with increases in diversification rate, we compared rate changes at the level of predefined beetle clades. For each branch in the time-calibrated phylogeny, the number of inferred chromosomal rearrangements was calculated as the sum of fission and fusion events. Branch-specific rearrangement rates were then calculated as the number of rearrangements divided by branch length. For each focal clade, rearrangement counts and branch length were summed across all descendant branches, and the focal rearrangement rate was calculated as the total number of rearrangements divided by the total descendant branch time. This focal rate was compared with the corresponding parent-background rate, defined as the rearrangement rate across the remaining branches descending from the nearest sampled parent clade after excluding the focal clade.

Rearrangement-rate shifts were tested using negative-binomial generalised linear models, with the number of rearrangements as the response and log branch length as an offset. Rearrangement-rate ratios were calculated as the focal clade rearrangement rate divided by the parent-background rearrangement rate. Clades were classified as having increased rearrangement rates when the negative-binomial test was significant after Benjamini–Hochberg correction and the estimated rate ratio was at least two-fold.

Diversification rates for beetle clades were obtained from McKenna et al. (2019) (*29*). To make the diversification-rate comparison equivalent to the rearrangement-rate comparison, we calculated a diversification-rate ratio for each focal clade as the diversification rate of the focal clade divided by the diversification rate of its nearest sampled parent clade.

We then tested whether increases in rearrangement rate predicted increases in diversification rate using node-level phylogenetic generalised least squares. Phylogenetic non-independence among clades was accounted for using a covariance matrix derived from shared branch length among the corresponding clade-defining nodes in the time-calibrated phylogeny. The final model used log diversification-rate ratio as the response variable and log rearrangement-rate ratio as the predictor.

### Comparing dynamics of chromosomal rearrangements across insect orders

To allow cross-order comparison, we used publicly available phylogenies and ALG reconstructions for Lepidoptera (*20*) and Diptera (*40*). For Diptera, additional reconstruction of the ancestral gene order was performed using AGORA (*22*), following the same procedure described in the previous section.

To test differences in the temporal dynamics of chromosomal rearrangement accumulation across the three orders, we calibrated the phylogenies by aligning the median divergence time of a shared node between the calibrated phylogenies from published studies (McKenna et al. 2019 (*29*) for Coleoptera, Bertone et al. 2008 (*88*) for Diptera, and Kawahara et al. 2029; 2023 (*89*, *90*) for Lepidoptera; Table S15) and the phylogenies used for ALG inference. To examine the association between interchromosomal rearrangements (cumulative fusions and fissions) and changes in gene order, we calculated fragmentation index (*FI*) as a proxy for the degree of intrachromosomal rearrangement relative to the inferred ancestral gene order of each order.

To assess the extent to which Coleoptera ALGs correspond to those inferred in other holometabolous insect orders, we compared the ALG assignments of orthologous genes across Coleoptera, Lepidoptera, and Diptera. Orthology between the three BUSCO gene sets (“coleoptera_odb12”, “diptera_odb12” and “lepidoptera_odb10”) was inferred using OrthoFinder (v.3.0.1.b1) (*91*). For each single-copy orthologue shared between either by all or by one of the pairs, we recorded its assignment to a Coleoptera ALG, a Lepidoptera Merian element/ALG assignment (M1–M31, MZ), and a Diptera ALG assignment (D1–D6), where available. The final correspondence table is available in Table S16.

To test whether ALG correspondence among orders exceeded random expectation, we compared ALG assignments of homologous BUSCO genes shared between each pair of orders. Correspondence was quantified using normalised mutual information (NMI), which treats ALG assignments as categorical partitions of the same set of genes. For each comparison, we generated a null distribution by randomly shuffling ALG labels in one order among BUSCO genes while preserving ALG size distributions. This procedure was repeated 9,999 times. Empirical *p*-values were calculated as the proportion of randomised datasets with NMI greater than or equal to the observed NMI, using a +1 correction.

As a complementary directional test, we asked whether genes assigned to each source ALG were more unevenly distributed across target ALGs than expected under random assignment. For each source ALG, we calculated the variance in the distribution of its homologous BUSCO genes across target ALGs. We then generated a null distribution by randomly assigning the same number of genes to target ALGs with probabilities proportional to target ALG sizes. This procedure was repeated 9,999 times for each source ALG and directional comparison. Empirical *p*-values were calculated as the proportion of simulations with variance greater than or equal to the observed variance, using a +1 correction, and were adjusted for multiple testing using the Benjamini–Hochberg procedure.

## Acknowledgments

We thank Charlotte Wright and Alexander Mackintosh for their valuable insights on reconstructing ancestral linkage groups and quantifying chromosomal rearrangements; Martin Wagah and Clothilde Chenal for their suggestions on the transposable elements annotation; and members of the Meier and Jaron Groups and the rest of the Tree of Life programme for helpful discussions and constructive criticism. This research was funded by the Wellcome Trust 220540/Z/20/A. For the purpose of Open Access, the author has applied a CC BY public copyright licence to any Author Accepted Manuscript version arising from this submission. The Darwin Tree of Life Project is funded by the Wellcome Trust through a Discretionary Award (218328/Z/19/Z) and a Bridge Award (226458/Z/22/Z) to the partnership and core funding to the Sanger Institute (206194), and by in-kind support from the partner institutions.

## Author contributions

Conceptualization: AM, JIM, KSJ

Methodology: AM, AB, SE, JG, JT, KN, JIM, KSJ

Investigation: AM, AB, SE, JT, KN, KSJ, KK, CS, KB, DEA, DZ

Visualization: AM, AB, SE, KN, KSJ

Funding acquisition: JIM, KSJ

Project administration: AM

Supervision: SE, JIM, KSJ

Writing – original draft: AM

Writing – review & editing: AM, AB, SE, DDM, JG, KN, JT, JIM, KSJ

## Data and materials availability

All data and code used in the analysis are publicly accessible at https://github.com/Obscuromics/coleoptera-ALGs.

## Supplementary Material

Materials and Methods

Supplementary Text

Figs. S1 to S59

Tables S1 to S16

References (*92–95*)

Data S1 to S2

