## Supplementary materials for "338 coleopteran genomes reveal exceptional rearrangement variation compared to other insect orders"

**Authors:** Arif Maulana[1, 2], Aleksandra Bliznina[1], Sam Ebdon[1], Jessica T. Thorpe[1], Julia Gries[1, 3], Karin Näsvall[1], Duane D. McKenna[4, 5], Ksenia Krasheninnikova[1], Camilla Santos[1], Karen Brooks[1], Dominic E. Absolon[1], Danil Zilov[1], Darwin Tree of Life Consortium[†], Kamil S. Jaron[1, ‡, *], Joana I. Meier[1, 6, ‡]

**Affiliations:**

[1]Tree of Life Programme, Wellcome Sanger Institute; Wellcome Genome Campus, Hinxton, Cambridge CB10 1SA, United Kingdom.

[2]Darwin College, University of Cambridge; Silver Street, Cambridge CB3 9EU, United Kingdom.

[3]Trinity College, University of Cambridge; Trinity Street, Cambridge CB2 1TQ, United Kingdom.

[4]Department of Biological Sciences, University of Memphis; Memphis TN 38152, United States of America.

[5]Center for Biodiversity Research, University of Memphis; Memphis TN 38152, United States of America.

[6]Department of Zoology, University of Cambridge; Downing Street, Cambridge CB2 3EJ, United Kingdom.

[†]Collective authorship: https://zenodo.org/records/7105029

[‡]these authors contributed equally

**The PDF file includes:**

Supplementary Text
Figs. S1 to S59
Tables S1 to S16

**Other Supplementary Materials for this manuscript include the following:**

Data S1 to S2

Supplementary Text

Manual adjustments of individual assemblies

*Philonthus cognatus* (GCA_932526585.2)
Both the X (OW052243.1) and Y (OW052250.1) chromosomes are, in fact, X chromosomes. They are labelled X1 and X2 in the updated assembly, which is currently being incorporated into the databases and will appear as icPhiCogn1.3. In addition, the Y chromosome was missing from the original assembly and has now been newly assembled and added to the chromosome list.

*Quedius lateralis* (GCA_964338995.1)
Based on BUSCO analysis, we propose that OZ202132.1 should be treated as X2 rather than Y, as currently indicated on the genome assembly page. This chromosome contains 221 genes, 200 of which are complete BUSCO genes, making it unlikely to be a Y chromosome.

*Tribolium confusum* (GCA_019155225.1)
LG7 and LG9 are missing from the assembly. Based on BUSCO analyses, four unlocalised scaffolds contain hundreds of BUSCO genes: JAGFVK010000006.1, JAGFVK010000007.1, JAGFVK010000009.1, and JAGFVK010000010.1. Based on coleopteran ALG painting and comparison with the close relative *Tribolium castaneum* (GCF_031307605.1), we propose that JAGFVK010000006.1 and JAGFVK010000007.1 are likely linked and together represent LG7, whereas JAGFVK010000009.1 and JAGFVK010000010.1 are likely linked and together represent LG9. We therefore consider this species to have nine chromosomes.

*Onthophagus taurus* (GCA_036711975.1)
Based on BUSCO analysis, two unlocalised scaffolds, NW_027248940.1 and NW_027248941.1, contain hundreds of BUSCO genes. Coleopteran ALG painting also indicates that the genes located on these scaffolds are assigned to C7, which is missing from all other chromosomes in the assembly. Based on comparison with the closest relative, *Onthophagus binodis* (GCA_054643895.1), these two scaffolds are probably linked and together form a single chromosome. In addition, two chromosomes in the assembly, CM071725.1 and CM071730.1, contain no BUSCO genes and have low coverage similar to that of the X chromosome, CM071729.1. This may indicate a fragmented assembly of the X or Y chromosome.

*Xenos peckii* (GCA_040167675.1)
This strepsipteran genome is a scaffold-level assembly. In the original publication, it was unclear which scaffolds corresponded to the contig names reported in the paper. We found that JAWUEG010000002.1–JAWUEG010000012.1 contained the highest proportions of BUSCO genes and considered most of these scaffolds to represent chromosomes.

The expected chromosome number is eight, suggesting that two chromosomes are fragmented into two and three scaffolds, respectively. The three fragmented scaffolds are treated as X-linked, as suggested in the original publication, and most likely correspond to JAWUEG010000009.1, JAWUEG010000010.1, and JAWUEG010000011.1. This X-linked assignment is supported by coleopteran ALG painting and comparison with the close relative *Stylops aterrimus* (GCA_965240395.1).

*Gnatocerus cornutus* (GCA_029298725.1)

Based on coverage data, we found that the unlocalised scaffold JARHXT010000016.1 has coverage equivalent to that of the X chromosome, CM055518.1. It contains 170 BUSCO genes, 168 of which are duplicated. None of these genes are present on the X chromosome. When analysed using coleopteran ALG painting, the genes on this scaffold were assigned to ALGs other than CX, predominantly C6. It remains unclear whether this scaffold represents the Y chromosome or an X2 chromosome.

*Dendarus foraminosus* (GCA_965152765.1)
Based on BUSCO analysis, we found that the chromosome assigned as Y, OZ224577.1, contains 35 complete single-copy BUSCO genes, most of them of CX origin. No coverage data are available for this assembly. This may indicate either misassembly of the X chromosome or a genuine chromosomal fission of the X chromosome.

*Lathrobium brunnipes* (GCA_965240315.1)
Based on BUSCO analysis, we found that the chromosome assigned as Y, OZ251362.1, contains 36 complete single-copy BUSCO genes, most of them of CX origin. Coverage data show that this chromosome has coverage similar to that of the X chromosome, OZ251361.1. This may indicate either misassembly of the X chromosome or a genuine chromosomal fission of the X chromosome.

*Phosphuga atrata* (GCA_944588485.1)
Based on BUSCO analysis, we found that the chromosome assigned as Y, OX155909.1, contains 29 complete single-copy BUSCO genes, all of them of CX origin. Coverage data show that this chromosome has coverage similar to that of the X chromosome, OX155907.1. This may indicate either misassembly of the X chromosome or a genuine chromosomal fission of the X chromosome.

*Diorhadba carinulata* (GCF_026250575.1)
Based on coverage data, we assigned NC_079460.1 as the X chromosome instead of NC_079472.1, as currently indicated on the genome assembly page.

*Platystomos albinus* (GCA_964106875.1)
On the genome assembly page, OZ066731.1 was assigned as the X chromosome based on homology with icEneSepi1.1 (GCA_963920635.1). However, this assignment does not appear to be consistent with the accompanying description. If homology was used as the basis for the assignment, OZ066730.1 is the more likely X chromosome. However, no coverage data are available to support this interpretation.

*Aethina tumida* (GCF_024364675.1)
Based on coverage data, we assigned NC_065442.1 as the Y chromosome and excluded it from the Syngraph analysis. We therefore consider this assembly to contain only seven chromosomes.

Manual adjustments of chromosomal rearrangement inference

As described above, three assemblies contain chromosomes that are fragmented across multiple scaffolds: *Tribolium confusum, Onthophagus taurus, and Xenos peckii*. Because these chromosomes are included in the reconstruction of chromosomal rearrangements along the phylogenetic tree, their pseudo-fragmentation would inflate the inferred number of chromosomal fissions. We therefore manually assessed and corrected the number of rearrangements on the branches descending from the internal nodes leading to the lineages in which these apparent fissions occurred. The relevant internal nodes are n535 for *Tribolium confusum*, n327 for *Onthophagus taurus,* and n7 for *Xenos peckii.*

We also made one modification to n125, which corresponds to the common ancestor of Phytophaga. Rather than retaining the nine chromosomes inferred to have been conserved since the emergence of Polyphaga, our manual assessment and comparative analyses suggest that n125 had 10 chromosomes. These chromosomes correspond to the ancestral linkage groups inferred for Chrysomeloidea (n158) and Curculionoidea (n159). The ancestral linkage groups reconstructed for these two nodes are largely identical. The reason n125 was not initially inferred to have 10 chromosomes is that the reconstruction of n158, representing Chrysomeloidea, was performed separately from the initial Syngraph analysis that included all species (see Materials and Methods).

#### Supplementary Figures

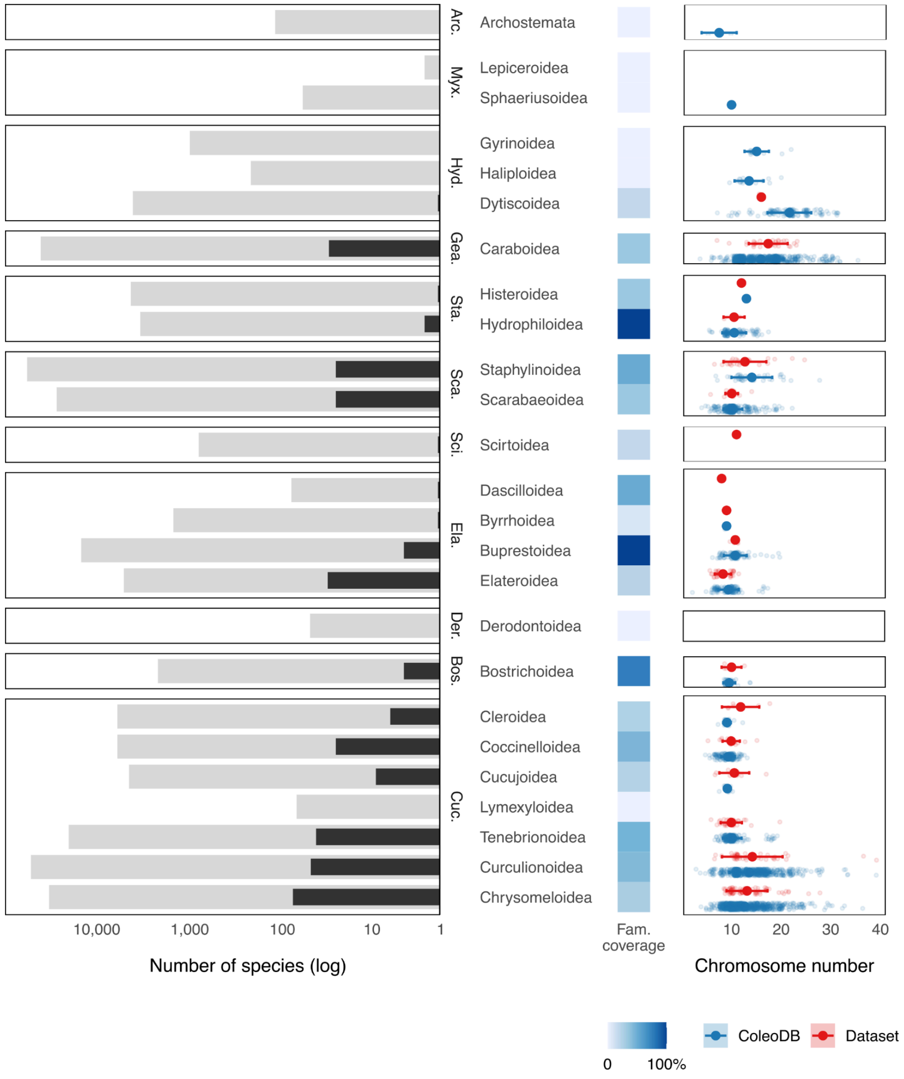


**Fig. S1.**

**Taxonomic sampling and karyotypic diversity represented in the dataset.** The left panel shows the number of species included in this study per superfamily, shown in dark grey, compared with estimates of the total number of described species per superfamily, shown in light grey (*92–94*). Boxes indicate different infraorders and the x-axis is shown on a log scale. The heatmap in the middle shows the relative proportion of families represented in the dataset for each superfamily. The heatmap in the middle showing the relative proportion of families represented in the dataset per superfamilies. The right panel shows the diversity of karyotype counts represented in the dataset, shown in red, compared with all known beetle karyotypes, shown in blue (*32*).

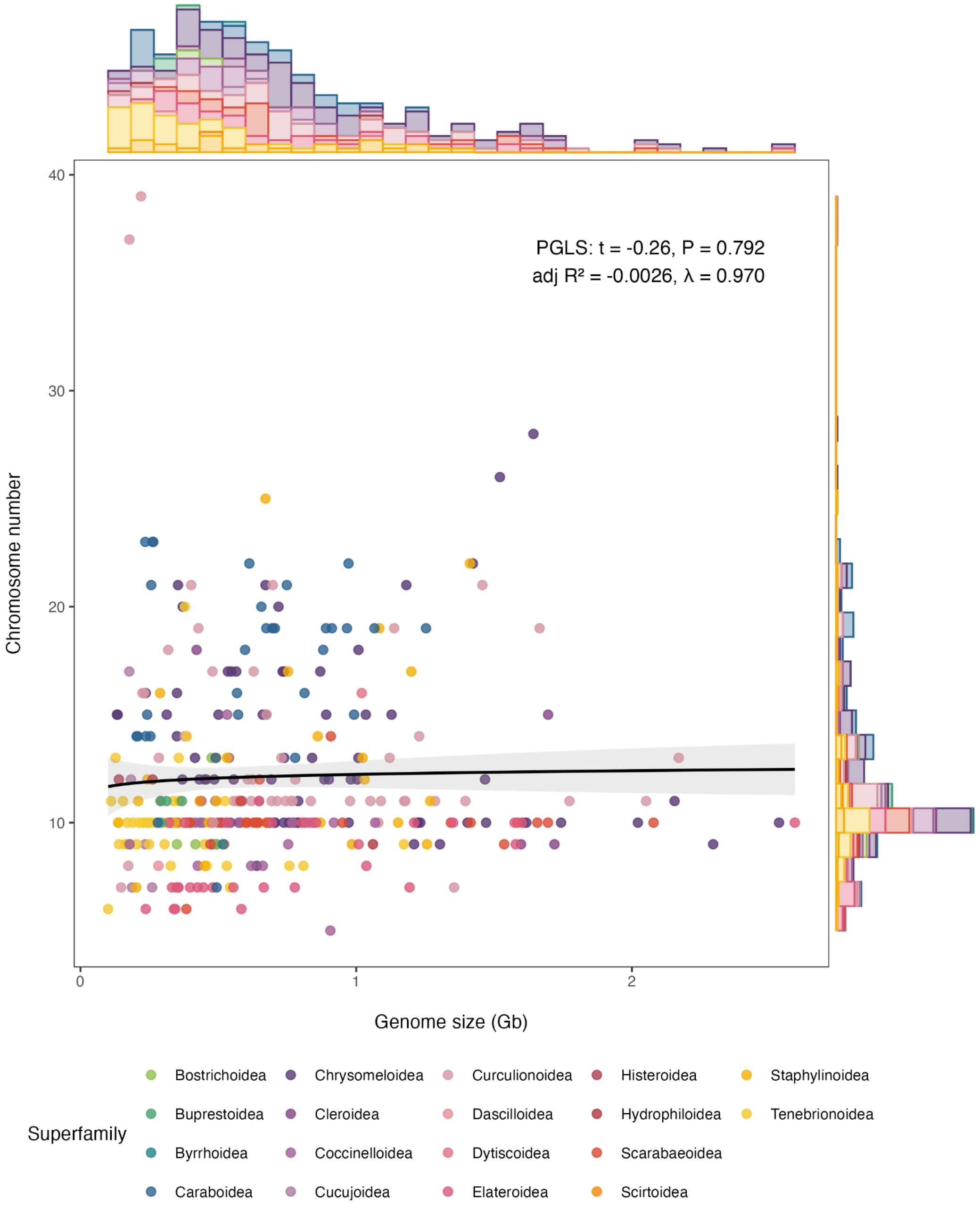


**Fig. S2.**

**Association between chromosome number and genome size in beetles.** Each point represents one species and is coloured according to the superfamily. Marginal histograms show the distributions of genome size and chromosome number across sampled species. The black line represents the phylogenetic generalised least squares regression (PGLS) fit, with the grey shaded area indicating the confidence interval. PGLS analysis showed no significant association between genome size and chromosome number across the sampled species (PGLS: $n = 355$; $t = -0.238$, $p = 0.812$, adjusted $R^2 = -0.0027$, $\lambda = 0.971$).

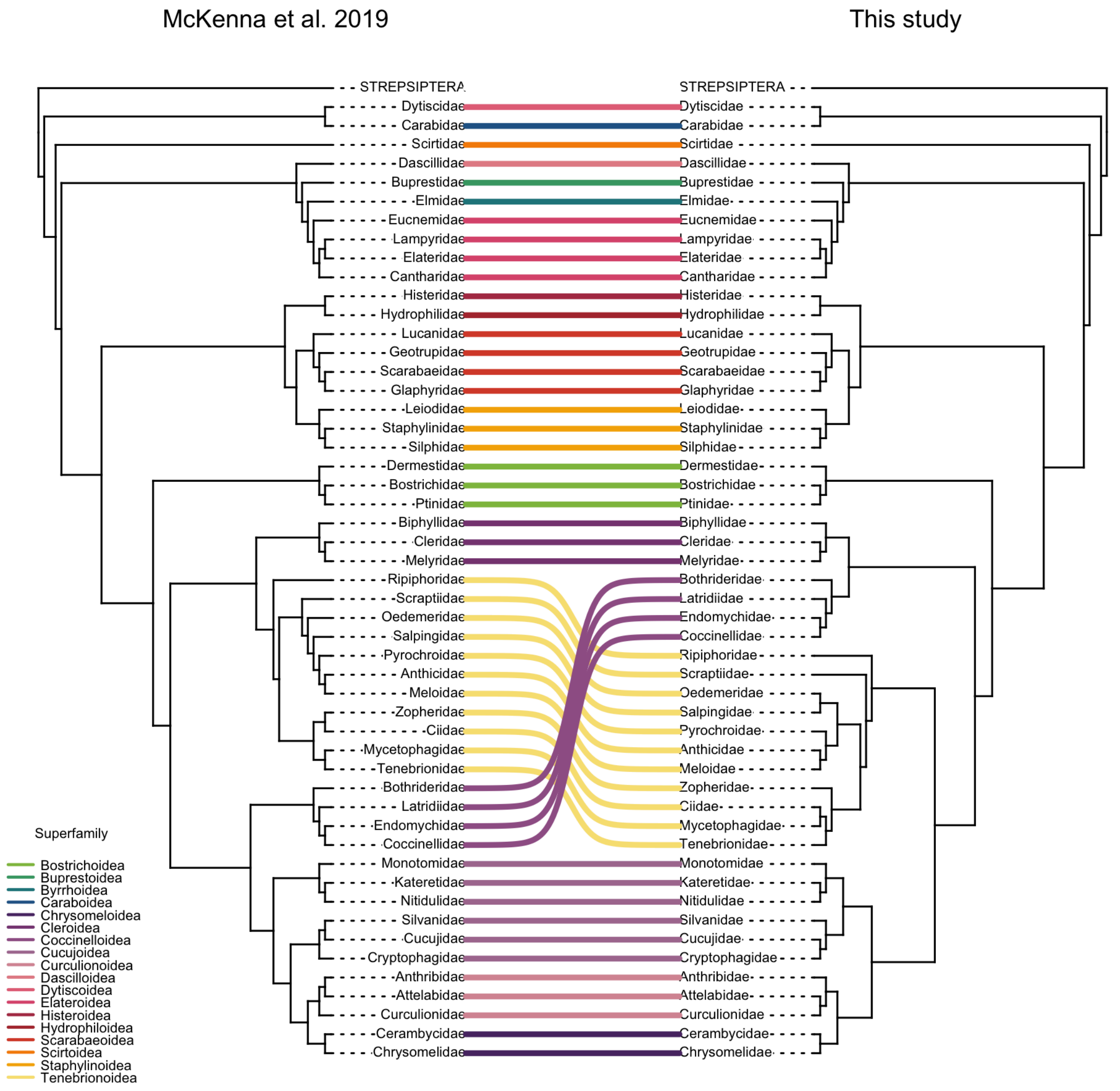


**Fig. S3.**

**Comparison of the topology of the phylogeny in this paper with widely used published beetle phylogeny.** The phylogeny reconstructed in this study was compared with the widely used phylogeny of (*29*), which includes broader family-level representation. Only families present in both phylogenies are shown. Although several deep relationships within Coleoptera remain debated (*30*, *95*), the topology of the current phylogeny is largely congruent with (*29*) at deeper superfamily- and family-level relationships, except for the positions of Coccinelloidea and Tenebrionoidea. Additional differences are observed in the placement of some families within Tenebrionoidea and in the position of Cryptophagidae within Cucujoidea.

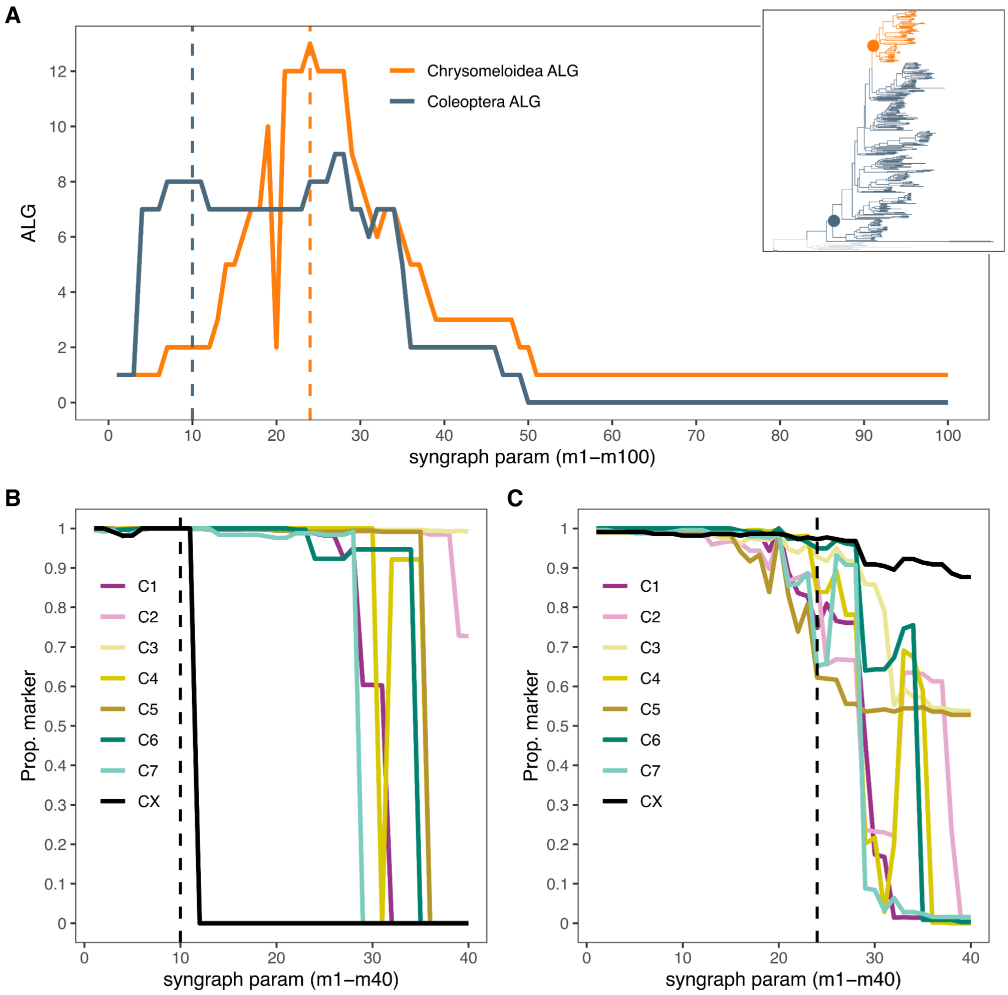


**Fig. S4.**

**Sensitivity of inferred ancestral linkage groups to Syngraph parameter choice. A.** Number of inferred ancestral linkage groups across Syngraph parameter value for Coleoptera and Chrysomeloidea ALG inference. A separate analysis was performed for Chrysomeloidea using higher parameter values to obtain more informative resolution. Dashed vertical lines indicate the parameter values selected for downstream analyses: m10 for Coleoptera ALGs and m24 for Chrysomeloidea ALGs. **B.** Proportion of markers retained across Syngraph parameters for Coleoptera ALGs. **C.** Proportion of markers retained across Syngraph parameters for Chrysomeloidea ALGs, coloured by Coleoptera ALG identity. The selected parameter values balance linkage group stability and marker retention.

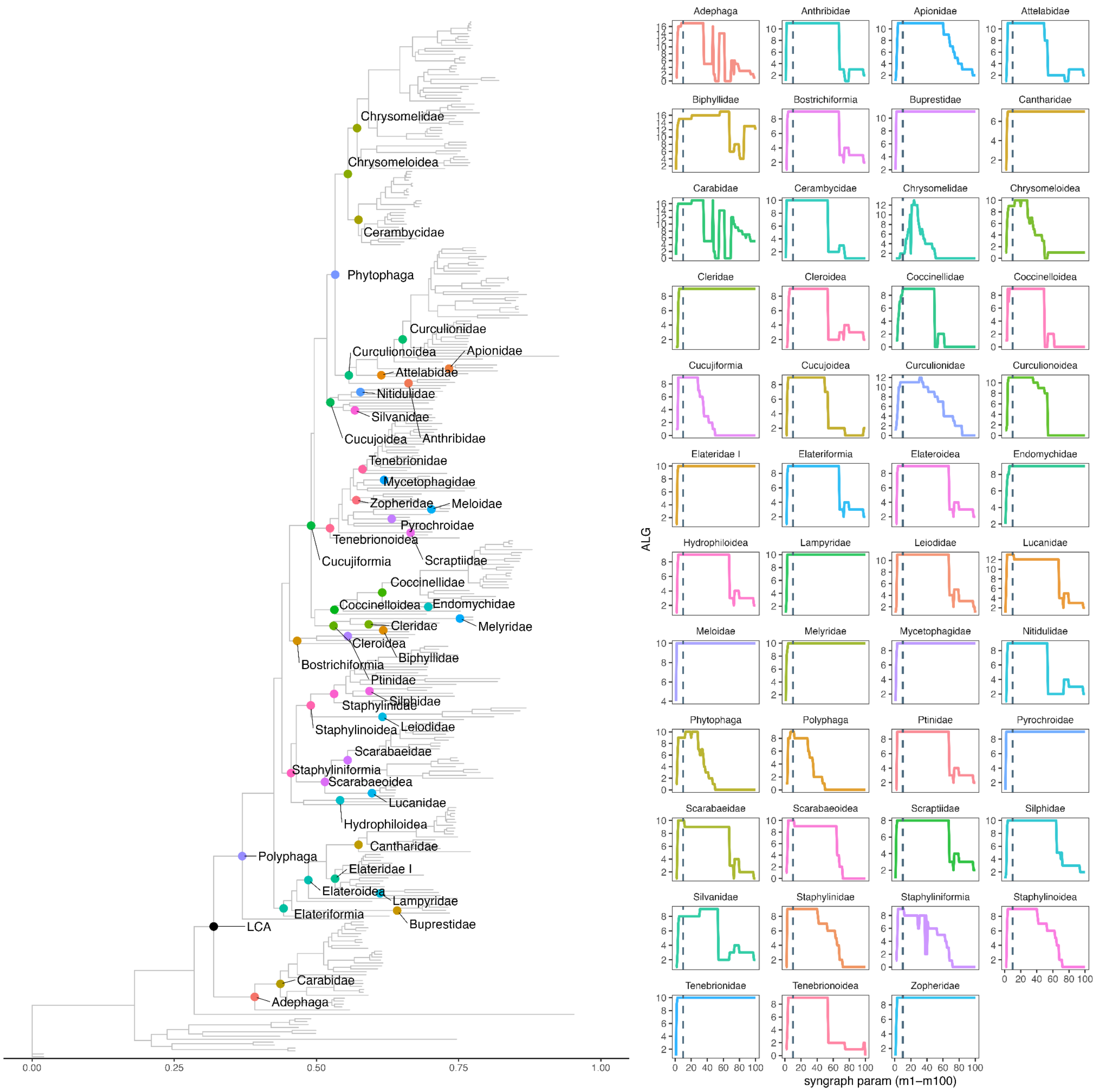


### Fig. S5.

**Sensitivity of inferred ancestral linkage groups to Syngraph parameter choice across major taxonomic lineages.** The number of inferred ALGs is shown across Syngraph parameter values for nodes representing the ancestors of infraorders, superfamilies, and families included in the dataset. Only lineages represented by at least two species are shown. Dashed vertical lines indicate the selected parameter value, m10, which was used for downstream inference of Coleoptera ALGs. A separate analysis was performed for Chrysomeloidea using higher parameter values, with m24 selected for downstream analyses, as shown in Fig. S4. The corresponding nodes are shown on the phylogeny to the left, which was inferred from a supermatrix of 3,153 BUSCO genes using IQ-TREE. Two Strepsiptera species with long branches were truncated from the tree to improve visibility.

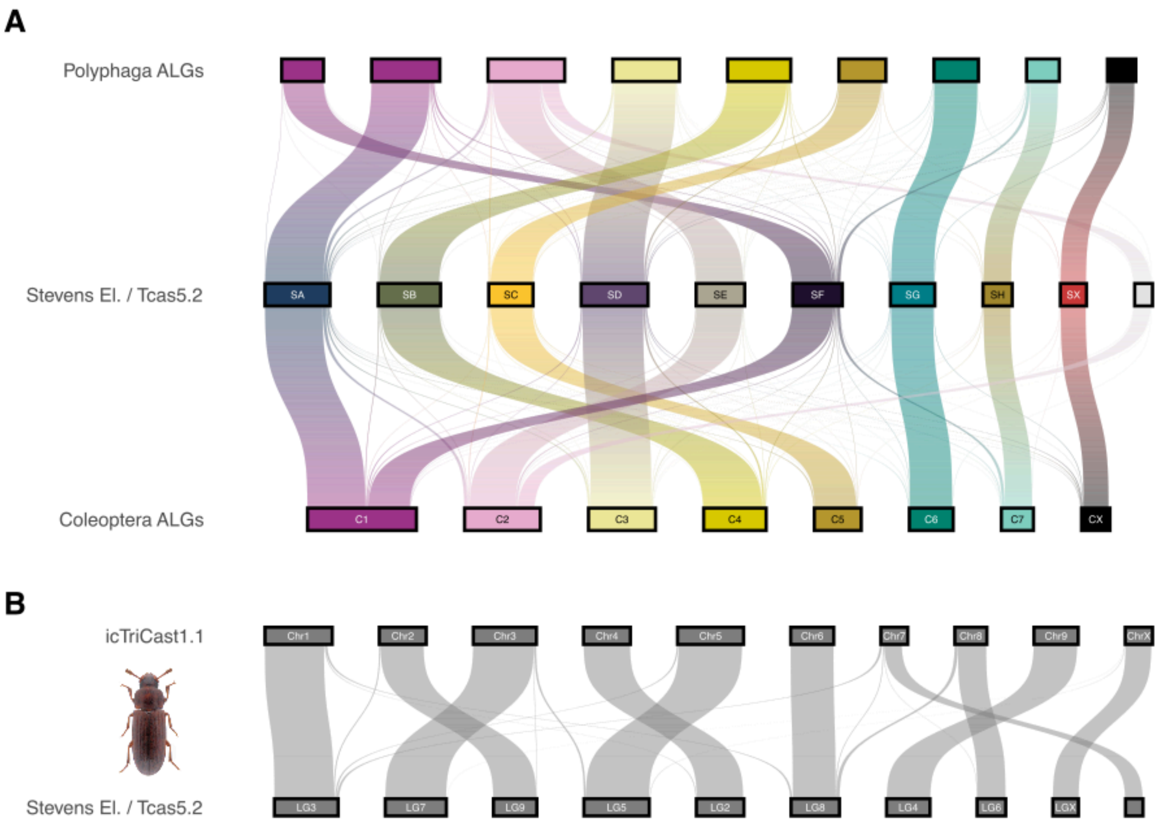


**Fig. S6.**

**Comparison of coleopteran ALGs with Stevens elements and polyphagan ALGS. A.** Sankey plot showing the correspondence between the coleopteran ancestral linkage groups (ALGs) reconstructed in this study, Stevens elements (*36*) and polyphagan ALGs. Stevens elements SA and SF were treated as a single element in our reconstruction. The Stevens elements broadly approximate the ancestral linkage groups of the last common ancestor of Polyphaga, although a large portion of C2 is absent. Stevens elements are coloured as in the original publication (*36*) **B.** Sankey plot comparing the *Tribolium castaneum* assembly used to infer Stevens elements (Tcas5.2; GCA_000002335.3), with the *T. castaneum* assembly included in this study (icTriCast1.1; GCF_031307605.1). The two assemblies are largely congruent, with minor differences in the positions of some BUSCO genes. Photo credits: *T. castaneum*, Udo Schmidt.

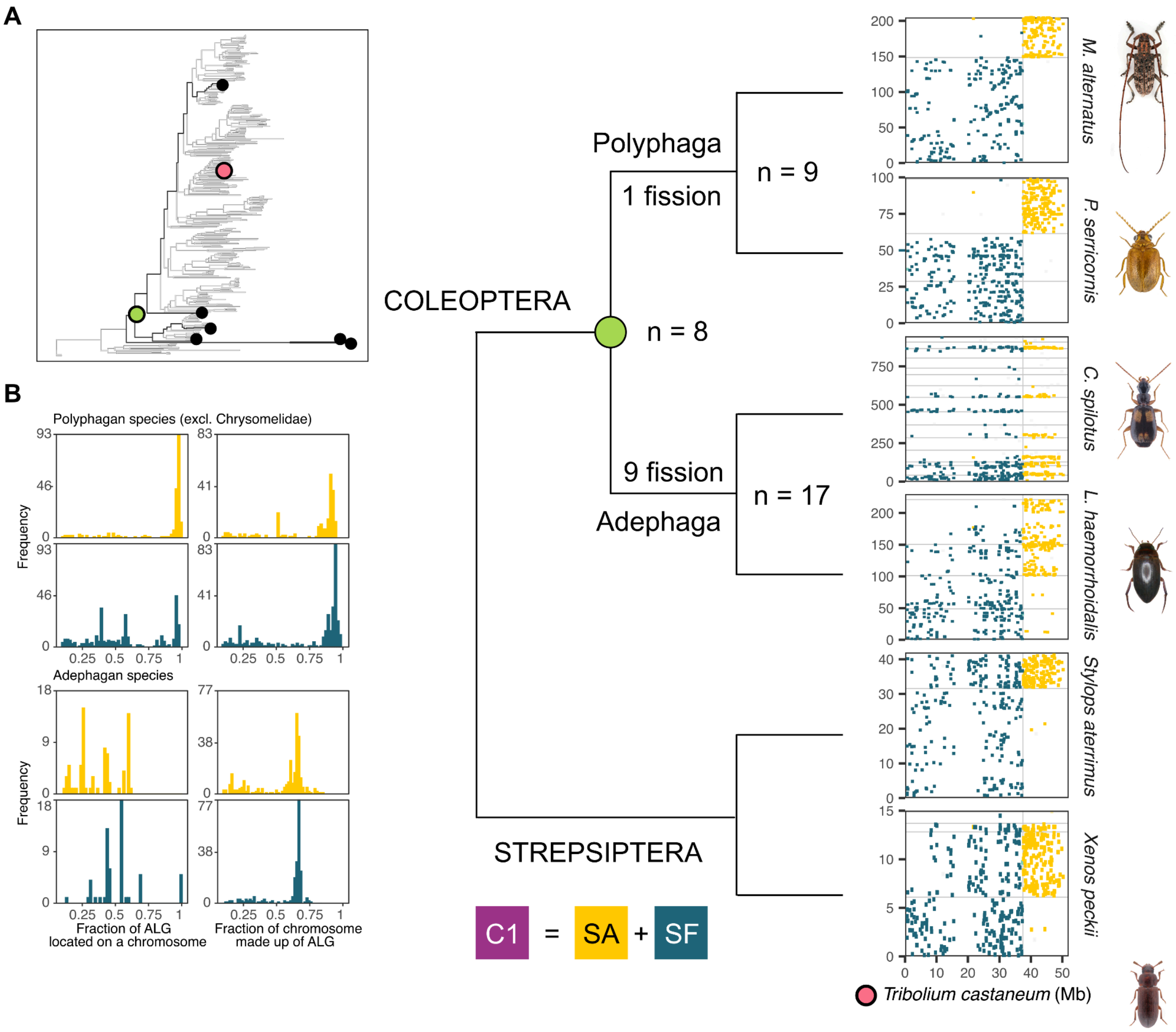


**Fig. S7.**

**Stevens elements do not represent Coleoptera ancestral linkage groups. A.** Oxford dot plots for four beetle species, including two Adephaga and two Polyphaga, and two strepsipteran outgroup species, mapped against *Tribolium castaneum* and coloured according to Stevens element identity (*36*). Only the two *T. castaneum* chromosomes corresponding to Stevens elements SA and SF are shown. For visibility, SA and SF are not coloured according to the original colour scheme used in Fig. S6. In our inference, these two Stevens elements are recovered as a single ancestral linkage group, C1. **B.** Histograms showing the fraction of each ALG located on a chromosome (left) and the fraction of each chromosome composed of a given ALG (right), for SA (yellow) and SF (blue) across Polyphaga (excluding Chrysomelidae) and Adephaga, but see Fig. S12. A clear signal separating SA and SF is observed only among polyphagan species, indicating that these elements likely reflect a polyphagan-specific subdivision rather than distinct Coleoptera ancestral linkage groups. Photo credits: Udo Schmidt and Hyunkyu Jang.

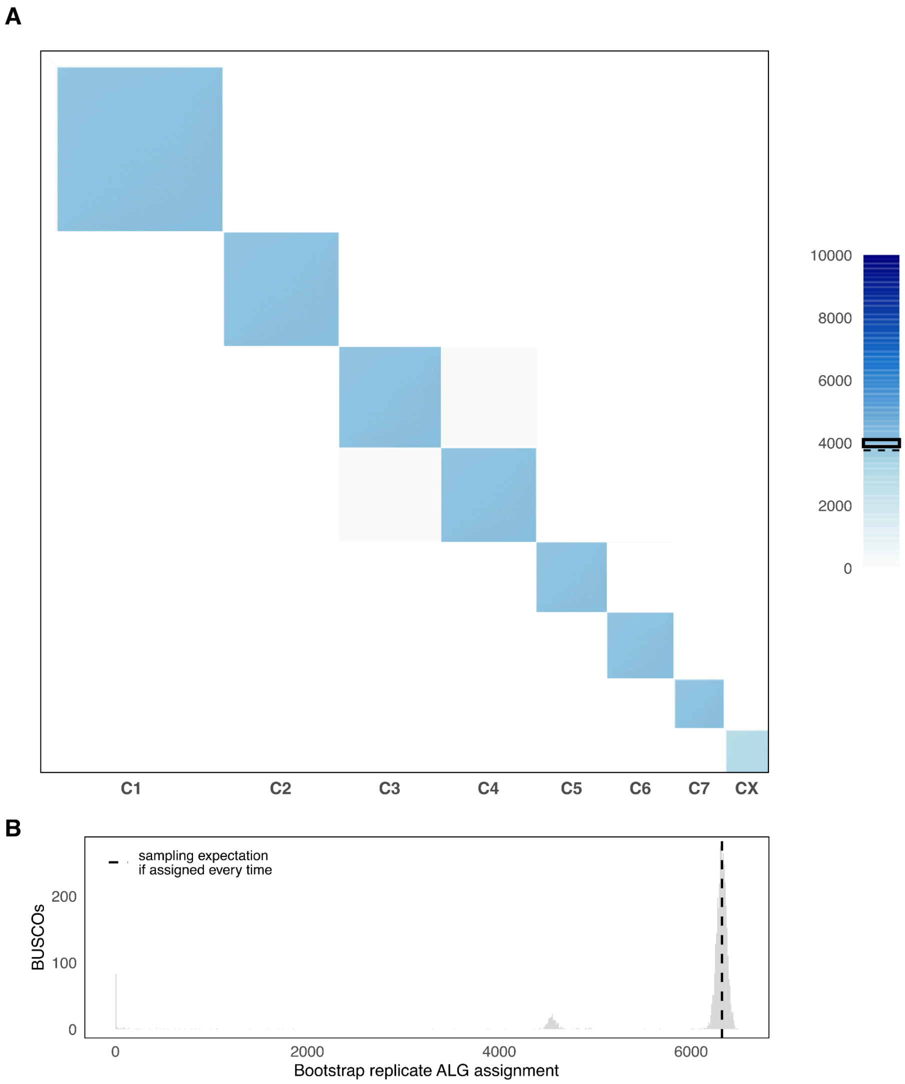


**Fig. S8.**

**Bootstrap support for reconstructed Coleoptera ALGs. A.** Pairwise co-assignment support of BUSCO genes into inferred ALGs over 10,000 bootstrap replicates of Syngraph. Each cell shows the number of bootstrap replicates in which two BUSCOs were inferred as members of the same linkage group. Diagonal blocks represent groups of BUSCOs that were consistently co-assigned across bootstrap replicates. The square on the legend corresponds to the 0.01–0.99 quantiles of the binomial distribution describing the expected sampling variation for perfectly consistent marker pairs under bootstrap resampling. **B.** Frequency of BUSCO assignment to any ALG across 10,000 bootstrap replicates. The dashed black line indicates the expected number of assignments for a BUSCO sampled and assigned whenever present under bootstrap resampling with replacement. Most assigned markers fall near this sampling expectation (except CX: ~4,500), indicating that they were assigned to a linkage group in all or nearly all bootstrap replicates in which they were sampled.

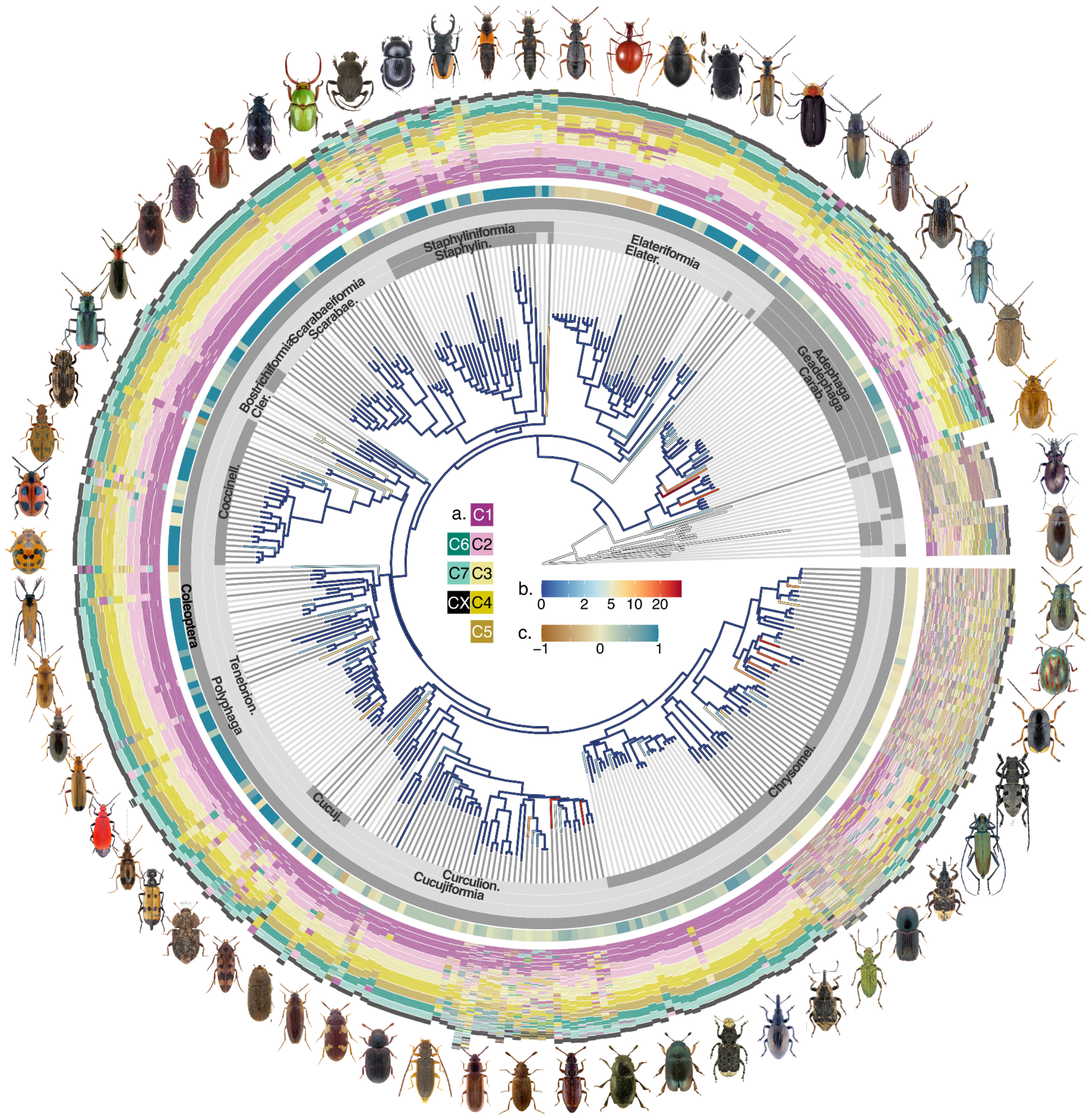


**Fig. S9.**

**Ancestral linkage groups painting across 338 species of Coleoptera.** Beetle phylogeny showing the number of chromosomal rearrangements and the painting of extant chromosomes according to coleopteran ancestral linkage groups (ALGs). The inner annotation layers, including the tip-extension lines, indicate taxonomic assignments at successive ranks. Families are distinguished by alternating blue shades along the tip-extension lines, followed outward by superfamily, infraorder, and order in the outermost taxonomic layer. The outermost layer beyond these annotations shows ALG painting, with chromosomes separated by thin white lines. Each chromosome is painted at single-gene resolution, with each gene represented by an individual stacked line. a, color legend for coleopteran ALG painting; b, number of rearrangement events; c, fusion excess (-1) or fission excess (1). Photo credits: Udo Schmidt, Lech Borowiec, JL Tachet, David Navrátil, Fred Chevaillot, Siga, Cosmin Manci, John Hallmén, Igor Souza-Gonçalves, Alex Hyde, H. Coiffait, David Ignace, Yves Bousquet.

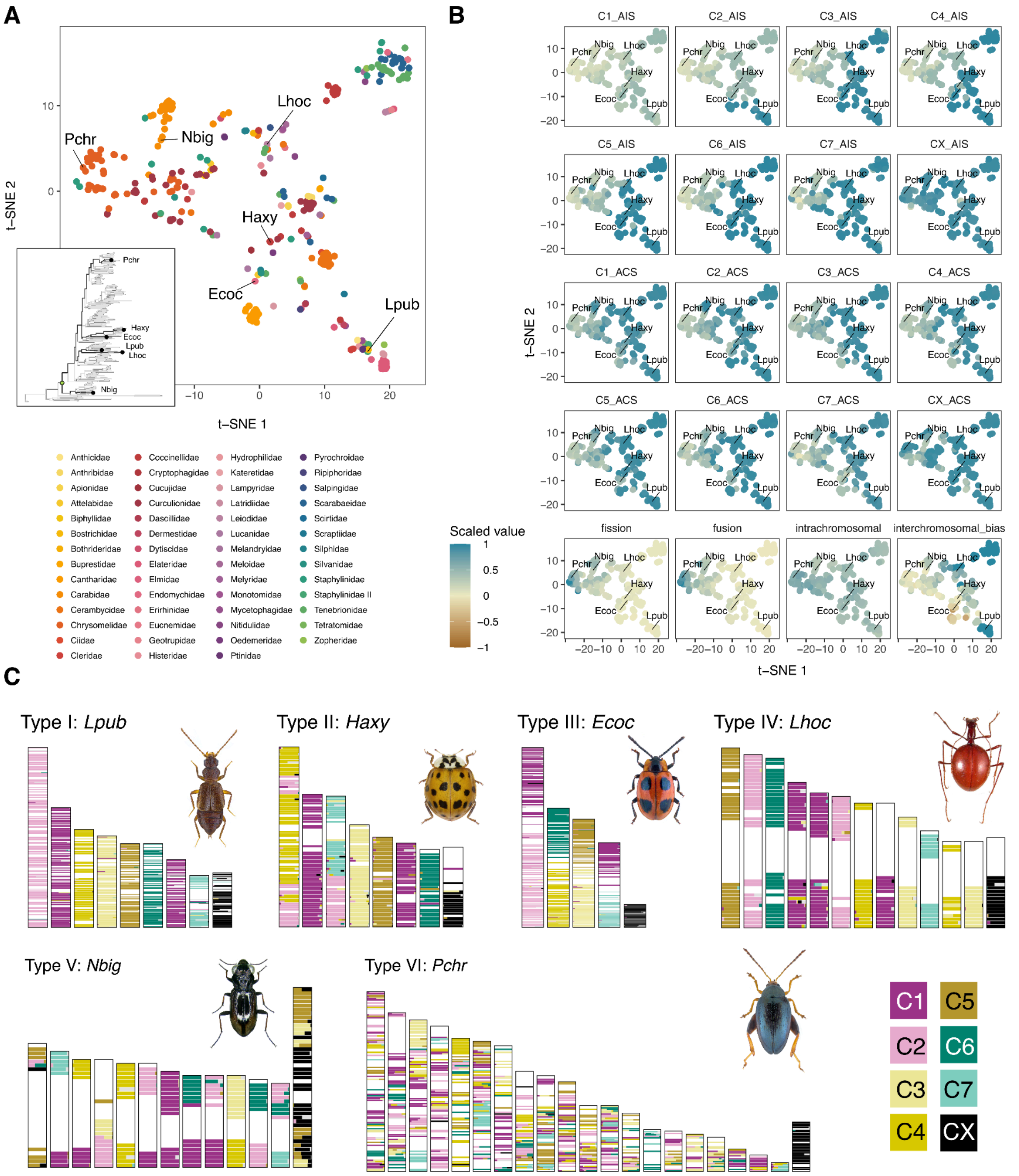


**Fig. S10.**

**Broad chromosomal evolution pattern in beetles. A.** t-SNE projection of beetle genomes based on ALG integrity scores (AIS), ALG composition scores (ACS), inferred total fission and fusion counts since the last common ancestor, interchromosomal rearrangement bias, and intrachromosomal rearrangement index. Each point represents one species and is coloured by family. Representative species corresponding to the six broad chromosomal evolution types are labelled, and their positions are shown in the inset phylogeny. **B.** Projection of scaled input variables onto the same t-SNE space. C1_AIS–C7_AIS and CX_AIS represent Coleoptera ALG integrity scores, with higher values indicating that an ALG remains highly intact and lower values indicating greater fragmentation across chromosomes. C1_ACS–C7_ACS and CX_ACS represent Coleoptera ALG contribution scores, with higher values indicating that chromosomes are dominated by markers from a single ALG and lower values indicating that chromosomes contain a more mixed composition of markers from multiple ALGs. Note that interchromosomal rearrangement bias ranges from -1 to 1, with negative values indicating more fusions than fissions meanwhile all other variables range from 0 to 1. **C.** ALG chromosomal painting for six representative species representing six broad chromosomal evolution types. Each vertical bar represents one chromosome, and each horizontal segment shows the proportional contribution of each ALG within a 1 Mb window. Photo credits: Udo Schmidt and H. Coiffait.

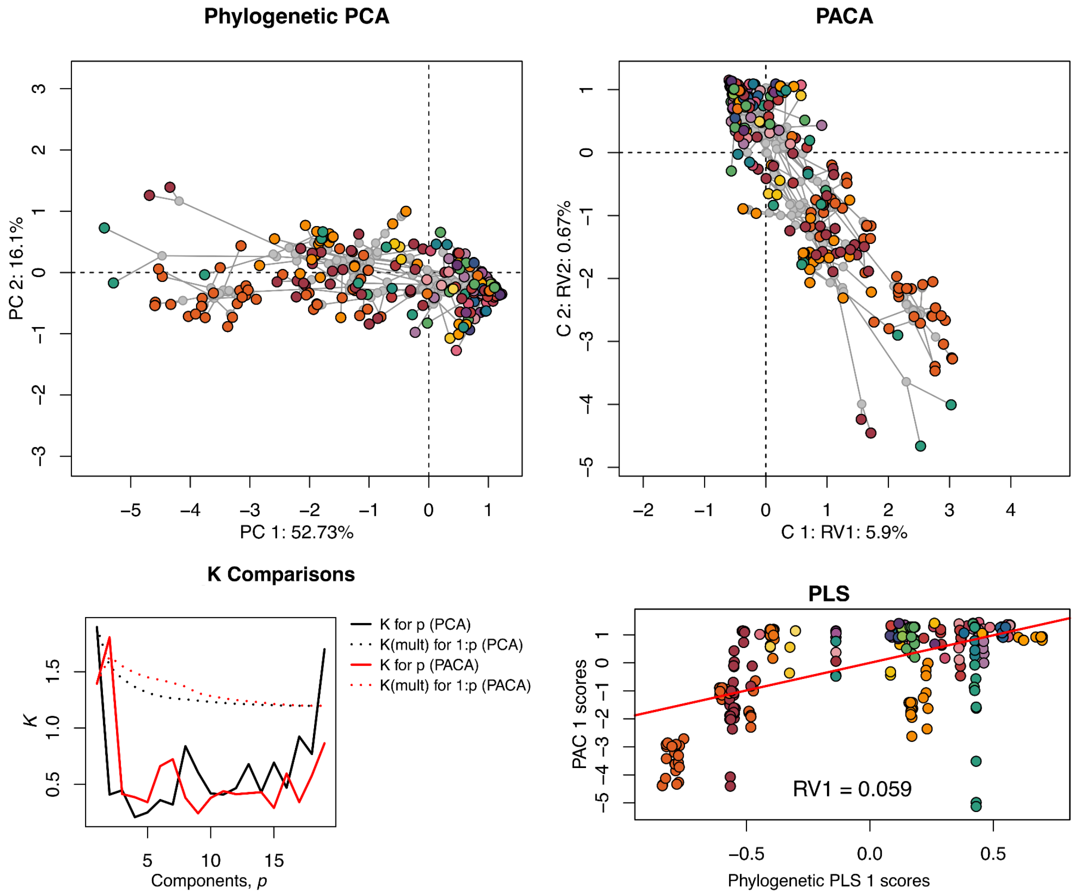


**Fig. S11.**

**Phylogenetic structure in multivariate chromosomal-evolution profiles.** Phylogenetic PCA and phylogenetically aligned component analysis (PACA) were performed using data used for t-SNE ordination plot in Fig. S10. Phylogenetic PCA summarises major axes of chromosomal variation after accounting for phylogenetic non-independence, whereas PACA identifies axes most strongly aligned with phylogenetic signals. The K comparison shows the distribution of phylogenetic signals across components for phylogenetic PCA and PACA, and the PLS panel shows the association between phylogenetic covariance and PACA scores. These analyses indicate that chromosomal-evolution profiles are phylogenetically structured but form a continuous landscape rather than discrete classes. Each point represents one species and is coloured by family with the same scheme as Fig. S10.

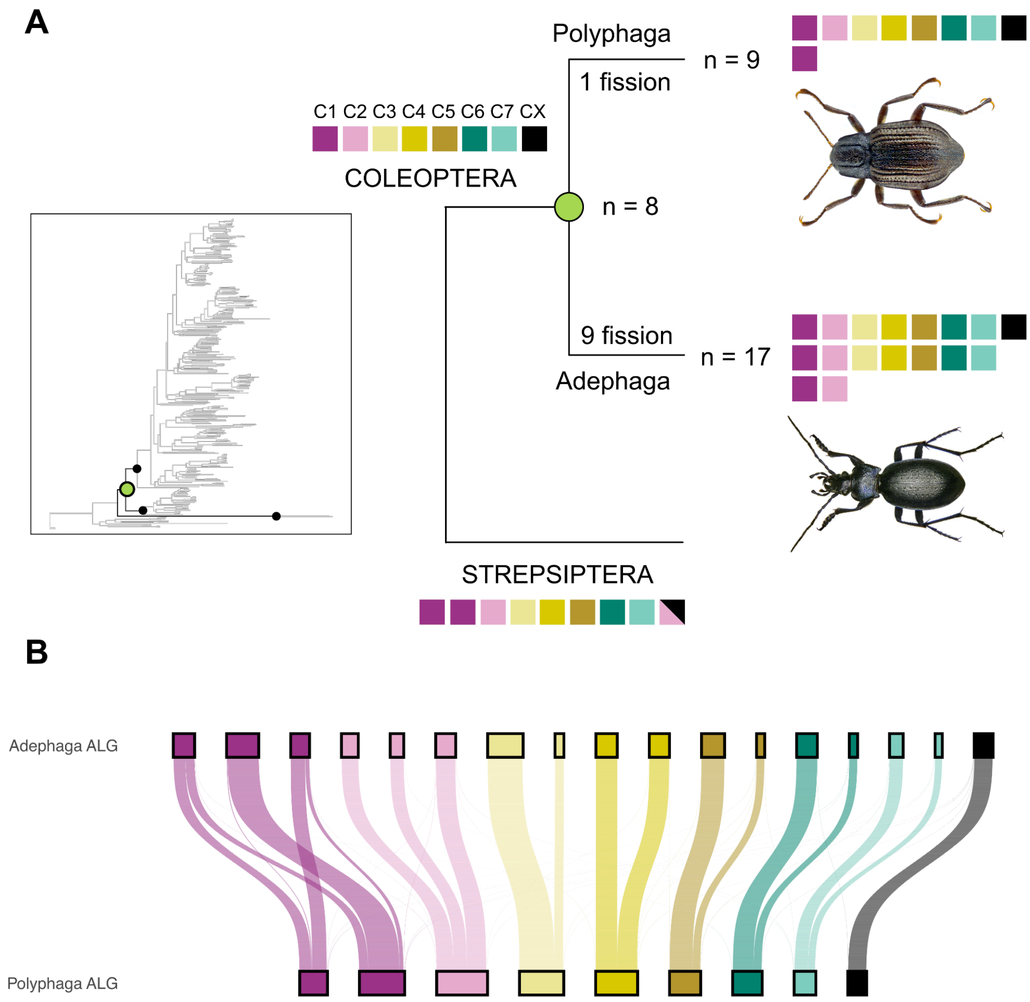


**Fig. S12.**

**Ancient fissions shaped the early evolution of beetle chromosomes. A.** Schematic representation of chromosomal rearrangement events during the divergence of the two largest beetle suborders, Adephaga and Polyphaga. The two suborders experienced lineage-specific fissions from the eight beetle ALGs (C1–C7 and CX), resulting in $n = 9$ in the polyphagan ancestor and $n = 17$ in the adephagan ancestor. Each coloured block represents one Coleoptera ALG. These contrasting early fission events produced distinct ancestral chromosomal configurations in Adephaga and Polyphaga, such that no extant species is expected to retain the complete ancestral coleopteran configuration. **B.** Sankey plot showing the correspondence between Adephaga ALGs and Polyphaga ALGs. Photo credits: Udo Schmidt.

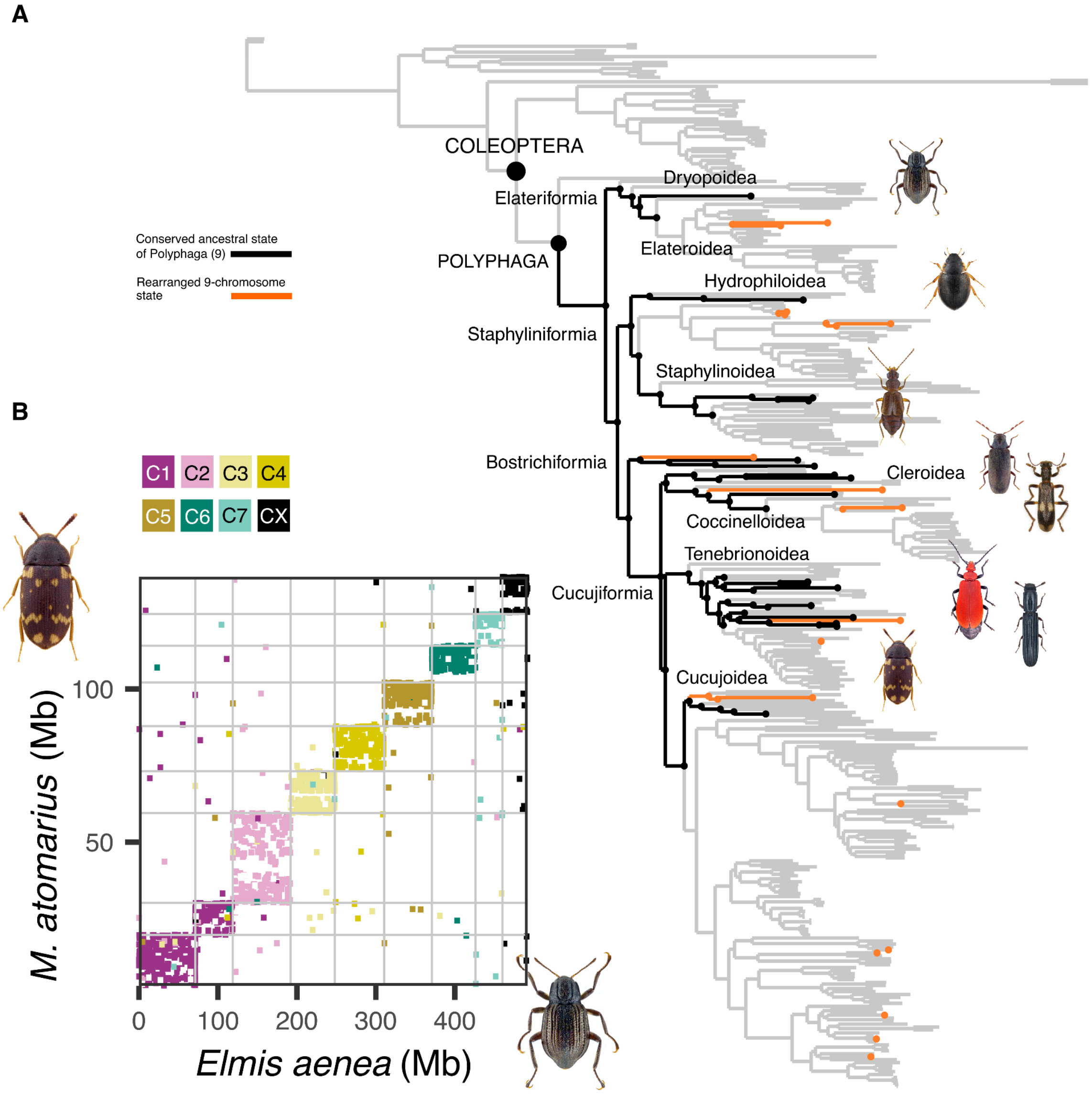


**Fig. S13.**

**Deeply conserved synteny of the ancestral polyphagan chromosome configuration. A.** Conservation of the nine inferred ancestral chromosomes of Polyphaga across extant beetle lineages. Black nodes and branches indicate internal branches and descendant lineages with no inferred interchromosomal rearrangements or chromosome-number changes relative to the polyphagan ancestral configuration. Orange branches indicate lineages with nine chromosomes whose current karyotypes were inferred to have arisen through subsequent rearrangements. **B.** Despite deep conservation of chromosome-scale synteny, extant species retaining this configuration show substantial gene-order reshuffling as shown in the example oxford dot plot of *Elmis aenea* and *Mycetophagus atomarius*. Photo credits: Udo Schmidt, John Hallmén, Fred Chevaillot, Alex Hyde.

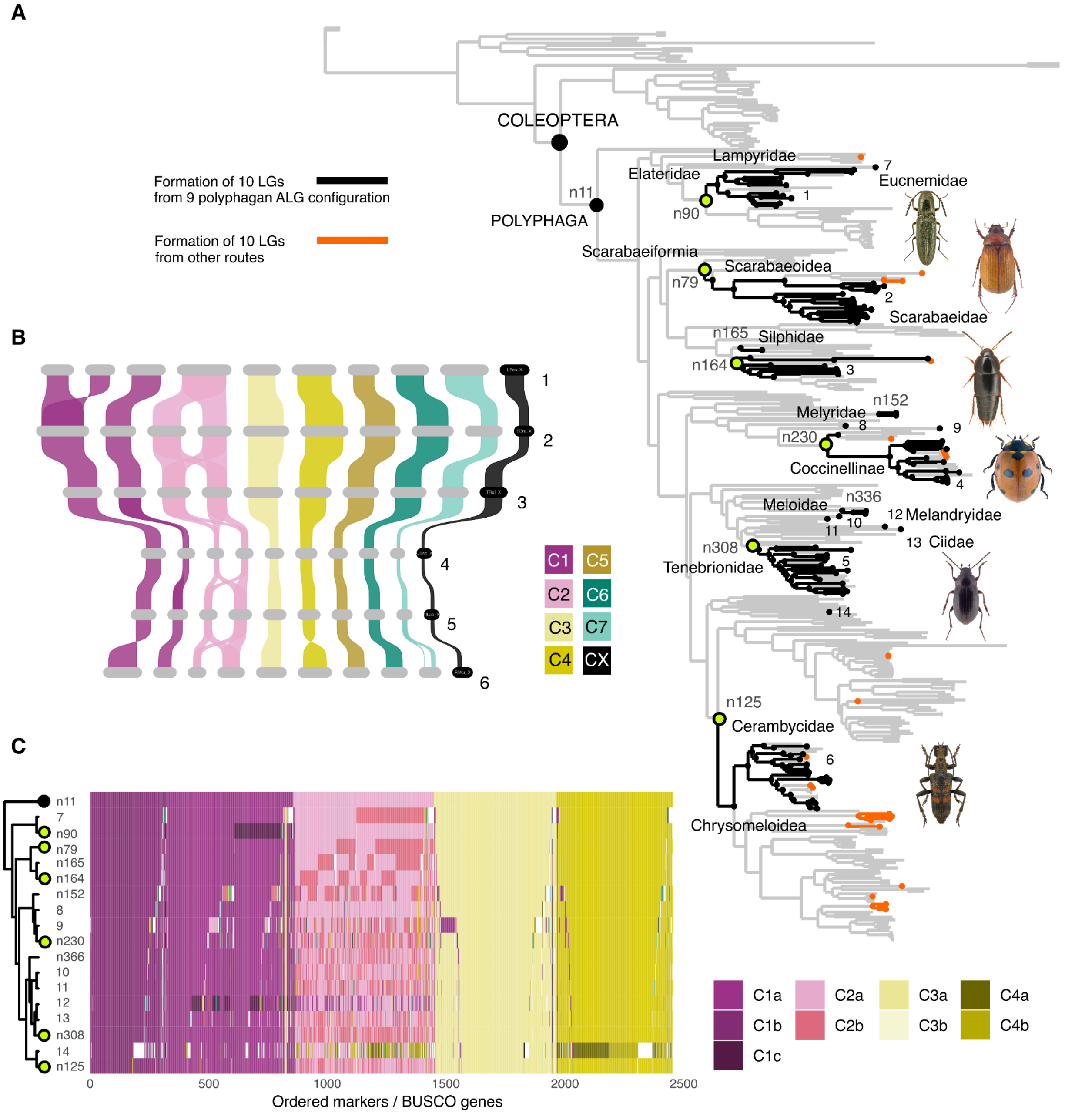


**Fig. S14.**

**Transition towards the most common 10-chromosome karyotype occurred multiple times in beetles. A**. Multiple origins of the 10 chromosome karyotype across beetle phylogeny. Independent transitions from 9 polyphagan ALGs to a 10-LGs configuration occurred 17 times, with an additional 20 transitions to 10 LGs inferred through other rearrangement routes. Black branches indicate lineages retaining the inferred 10-LGs configuration without subsequent interchromosomal rearrangements. Orange branches indicate 10-LGs lineages whose karyotypes were inferred to have undergone additional rearrangements after the initial transition. Numbers on the tree correspond to representative species shown in the riparian plot in panel B and the ALG tile plot in panel C. Green nodes mark six representative major ancestral nodes selected for detailed comparison in panel C. **B**. Riparian plot of six representative extant species that retain this 10-ALG karyotype without subsequent interchromosomal rearrangements. Horizontal bars indicate different chromosomes and the vertical ribbons connect the position of BUSCO genes between species. BUSCO links assigned to the same ALG were merged into broad syntenic ribbons when they formed contiguous or near-contiguous blocks, regardless of their relative gene order between species and were coloured according to Coleoptera ALG. Singleton and small blocks were removed to improve visibility. **C**. ALG tile plot for 19 representative ancestral nodes/species with independently evolved 10-LGs configurations. BUSCO markers are ordered along the y axis. Colours indicate the polyphagan ALG components involved in the transition, showing that superficially similar 10-LGs configurations can be produced by rearrangements involving different gene sets. 1, *Limonius poneli*; 2, *Serica brunnea*; 3, *Tachinus rufipes*; 4, *Coccinella septempunctata*; 5, *Nalassus laevioctostriatus*; 6, *Rhagium mordax*; 7, *Microrhagus pygmaeus*; 8, *Endomychus armeniacus*; 9, *Serangium japonicum*; 10, *Pycnomerus fuliginosus*; 11, *Tarphius canariensis*; 12, *Orchesia undulata*; 13, *Cis bilamellatus*; 14, *Librodor japonicus*. Photo credits: Lech Borowiec.

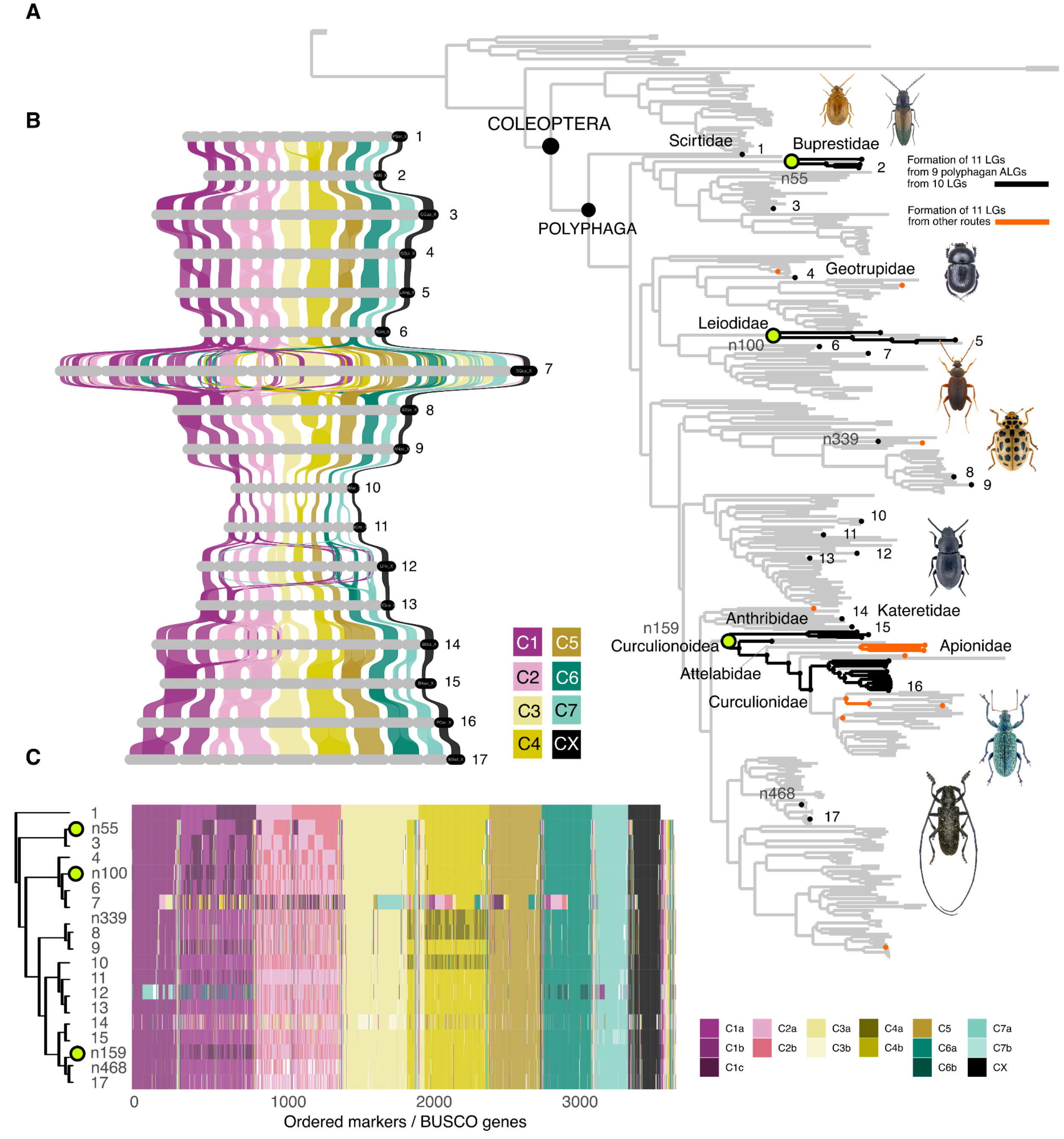


**Fig. S15.**

**The evolution of the 11-chromosome karyotype in beetles. A.** Multiple origins of the 11-chromosome karyotype across the beetle phylogeny. Black branches indicate lineages inferred to have formed 11 LGs either directly from the 9-ALG polyphagan ancestral configuration, or 10-LGs configuration. Orange branches indicate formation of 11 LGs through other rearrangement routes. Labels on the tree correspond to representative species shown in the riparian plot in panel B and the ALG tile plot in panel C. **B.** Riparian plot of 17 representative extant species or lineages with 11-ALG configurations. Horizontal bars indicate different chromosomes, and the vertical ribbons connect the positions of BUSCO genes between species. BUSCO links assigned to the same ALG were merged into broad syntenic ribbons when they formed contiguous or near-contiguous blocks, regardless of their relative gene order between species, and were coloured according to Coleoptera ALG. Singleton and small blocks were removed to improve visibility. **C.** ALG tile plot comparing marker composition across the representative 11-ALG node/species shown in panel B and highlighted in panel A. BUSCO markers are ordered along the y-axis. Colours indicate ALG assignments, showing that superficially similar 11-ALG configurations differ in the underlying gene sets involved. 1, *Prionocyphon serricornis*; 2, *Agrilus biguttatus*; 3, *Ctenicera cuprea*; 4, *Geotrupes spiniger*; 5, *Leonhardella angulicollis*; 6, *Anthobium unicolor*; 7, *Scaphidium quadrimaculatum*; 8, *Adalia decempunctata*; 9, *Anisosticta novemdecimpunctata*; 10, *Hycleus marcipoli*; 11, *Bitoma crenata*; 12, *Lagria hirta*; 13, *Dailognatha quadricollis*; 14, *Brachypterus glaber*; 15, *Brassicogethes aeneus*; 16, P*olydrusus cervinus*; 17, *Monochamus sutor*. Photo credits: Udo Schmidt.

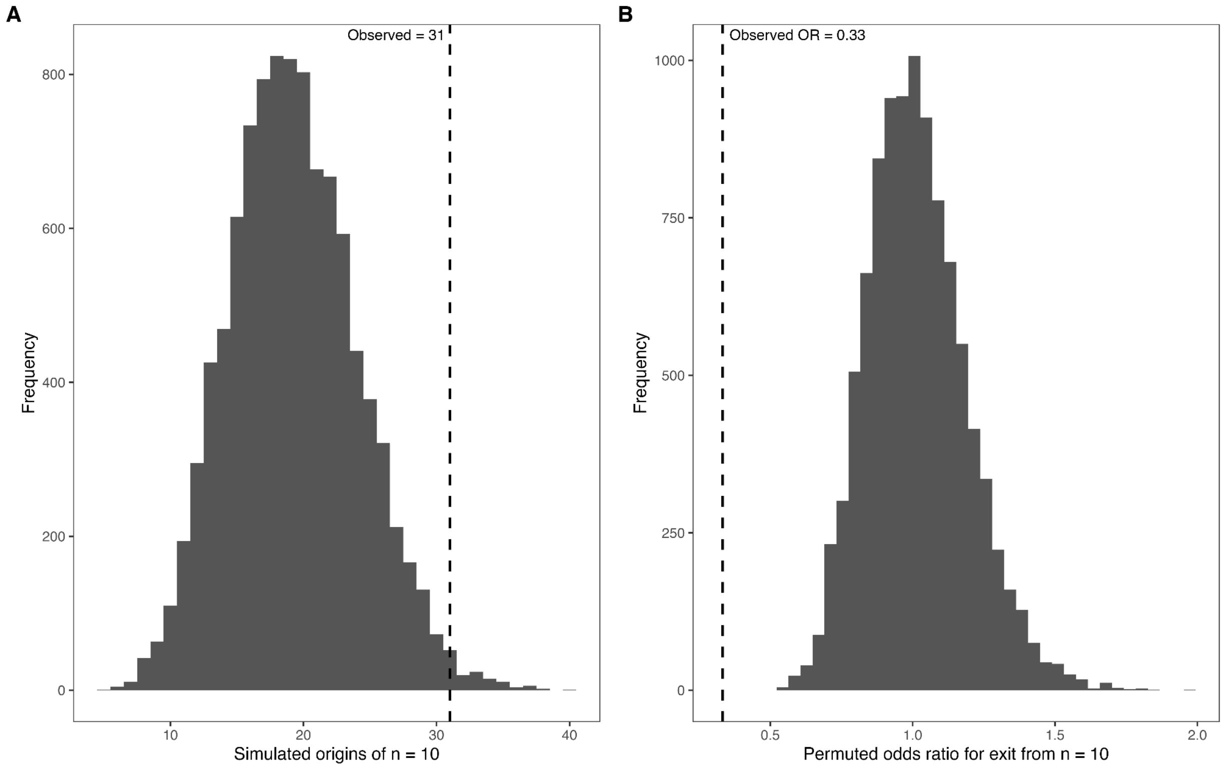


**Fig. S16.**

**Recurrence and stability of the 10-chromosome karyotype in beetles.**

**A.** Null distribution of the number of independent origins of $n = 10$ from calibrated empirical transition simulations. Simulations were scaled to match the observed number of chromosome-number changes. The dashed line shows the observed number of origins, 31, compared with a simulated median of 19 and 95% interval of 11–29. **B.** Permutation null distribution of the odds ratio for chromosome-number change on branches starting at $n = 10$ after accounting for branch length. The dashed line shows the observed odds ratio, 0.33, compared with a permutation 95% interval of 0.71–1.40.

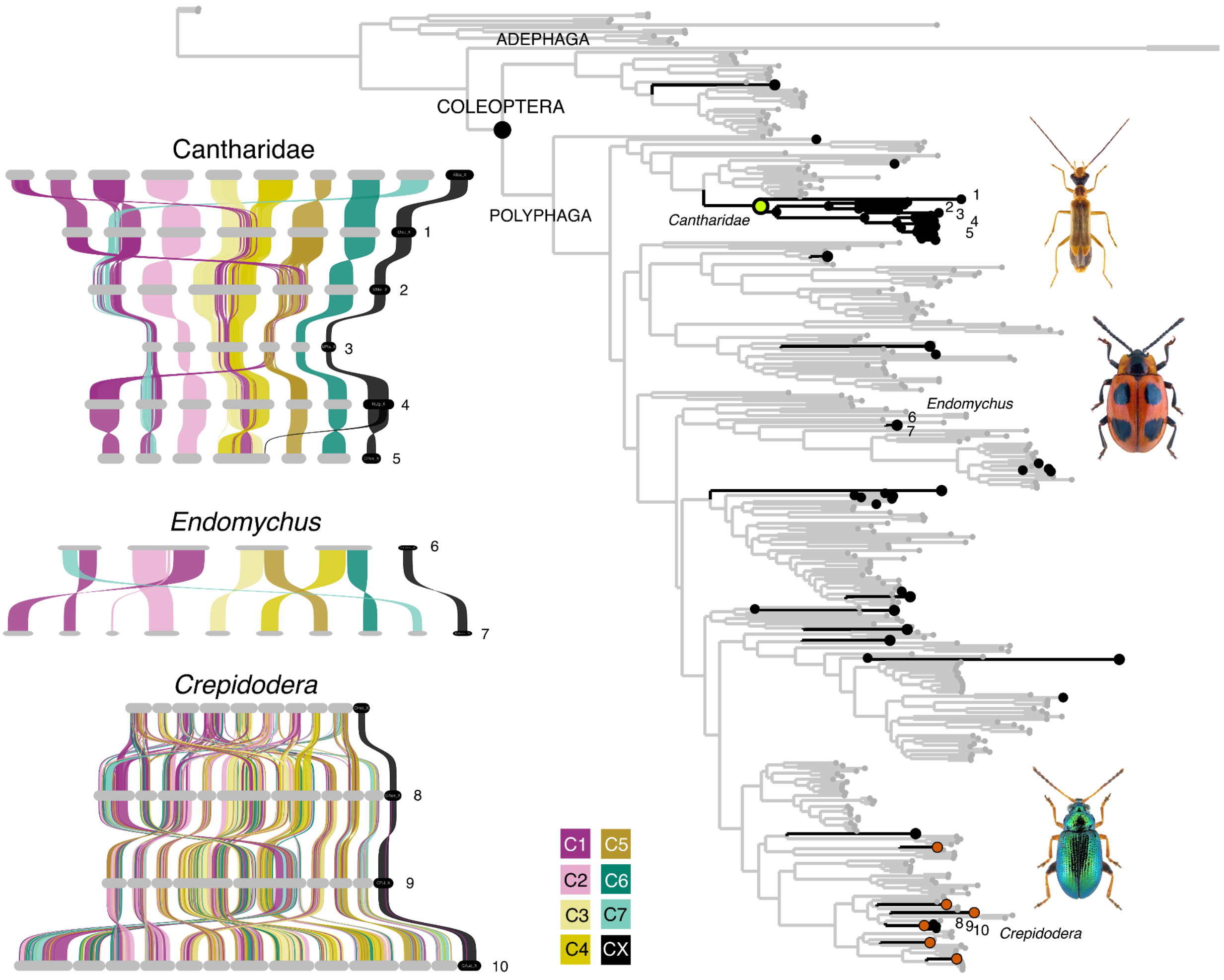


**Fig. S17.**

**Fusion-biased rearrangements are rare across beetle lineages. A.** Phylogenetic distribution of fusion-biased chromosome rearrangement lineages in Coleoptera. Highlighted branches mark inferred origins where the cumulative state first becomes fusion-biased and where cumulative rearrangement history is fusion-biased, defined as cumulative fusion events exceeding cumulative fission events from the last common ancestor of beetles. Nodes highlighted in black indicate lineages or species with chromosome number less than nine (less than Polyphaga ALGs) meanwhile orange points indicate nodes or tips that are fusion biased but have chromosomes more than nine. Numbered tips correspond to representative taxa shown in the adjacent synteny plots. Green node marks Cantharidae, the only family of beetles in our dataset that shares a fusion-biased rearrangement pattern. Left panels show representative chromosome-scale synteny comparisons for selected taxa, with ribbons coloured by ancestral linkage group. 1, *Ichthyurus bourgeoisi*; 2, *Malthodes minimus*; 3, *Malthinus flaveolus*; 4, *Rhagonycha lignosa*; 5, *Cantharis rustica*; 6, *Endomychus coccineus*; 7, *Endomychus armeniacus*; 8, *Crepidodera aurea*; 9, *Crepidodera fulvicornis*; 10, *Crepidodera aurata*. Photo credits: Udo Schmidt.

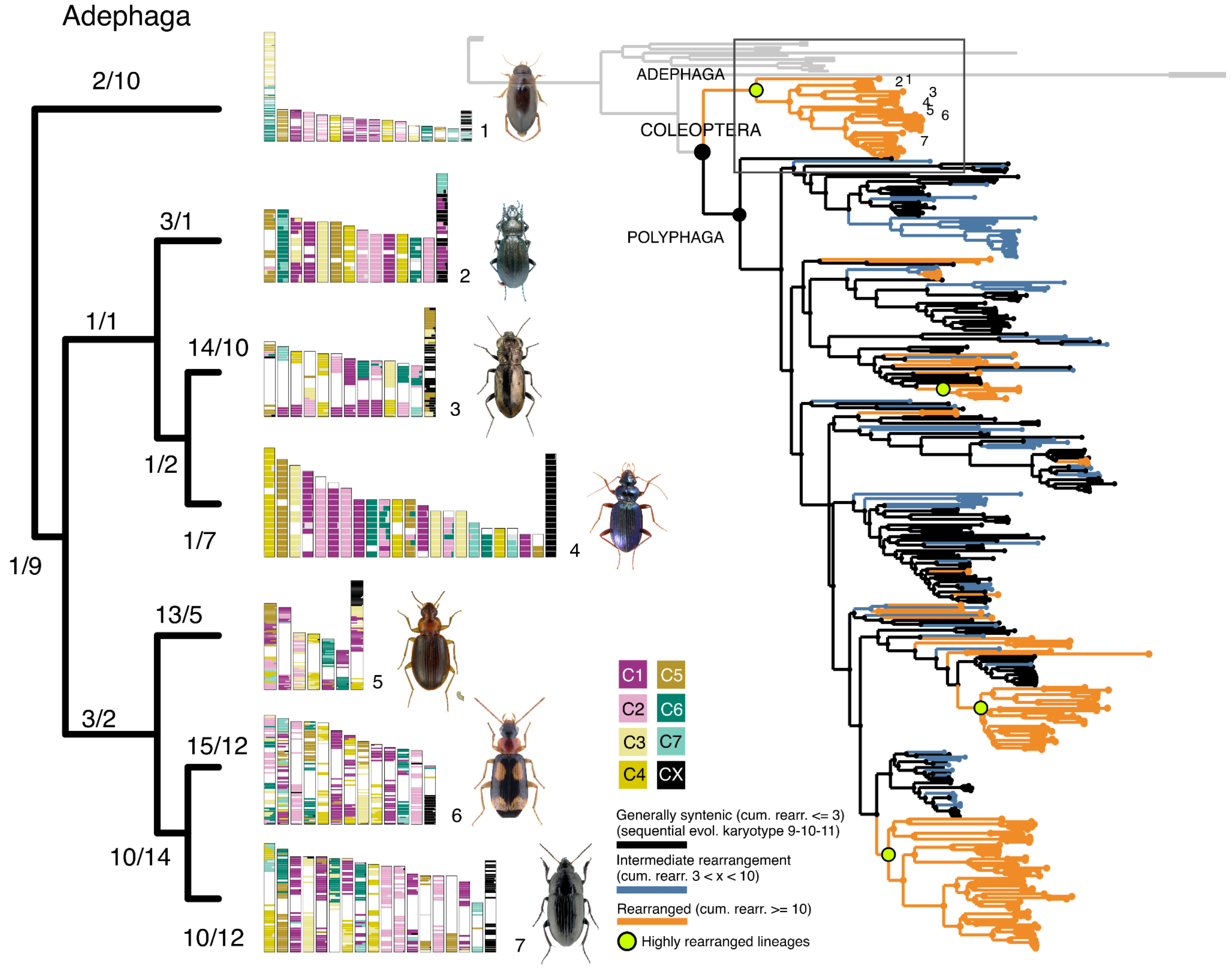


**Fig. S18.**

**Highly rearranged beetle karyotypes evolved repeatedly across major beetle lineages: Adephaga.** Phylogenetic distribution of rearrangement categories across Coleoptera, with Adephaga highlighted. Generally syntenic lineages (black) were defined as those with fewer than three cumulative rearrangements and a sequential karyotype transition involving 9, 10, or 11 chromosomes. Intermediate lineages (blue) were defined as those with three to fewer than ten cumulative rearrangements. Highly rearranged lineages (orange) were defined as those with at least ten cumulative rearrangements from the beetle ancestor node n6 to the focal node or tip. Highlighted green nodes mark the most highly rearranged clades. These highly rearranged lineages show distinct patterns of karyotype evolution, ranging from extensive simple fissions to putative chromosome-arm exchanges and combinations of repeated fusions and fissions with highly reshuffled gene order. Labels on the tree correspond to representative species shown in the subset phylogeny and ALG painting plot. ALG painting of representative adephagan species. Each vertical bar represents one chromosome, and each horizontal segment shows the proportional contribution of each ALG within a 1 Mb window. Left-side branch labels indicate inferred fusion/fission counts along collapsed branches connecting the selected taxa. Horizontal bars indicate different chromosomes, and the vertical ribbons connect the positions of BUSCO genes between species. BUSCO links assigned to the same ALG were merged into broad syntenic ribbons when they formed contiguous or near-contiguous blocks, regardless of their relative gene order between species, and were coloured according to Coleoptera ALG. Singleton and small blocks were removed to improve visibility. 1, *Liopterus haemorrhoidalis*; 2, *Carabus staehlini*; 3, *Notiophilus biguttatus*; 4, *Leistus spinibarbis*; 5, *Ocys tachysoides*; 6, *Dromius quadrimaculatus*; 7, *Amara aenea*. Photo credits: Udo Schmidt, David Ignace, Dmitriy V Obydov, Göran Liljeberg, Patrick Deyroze, Guillaume Jacquemin.

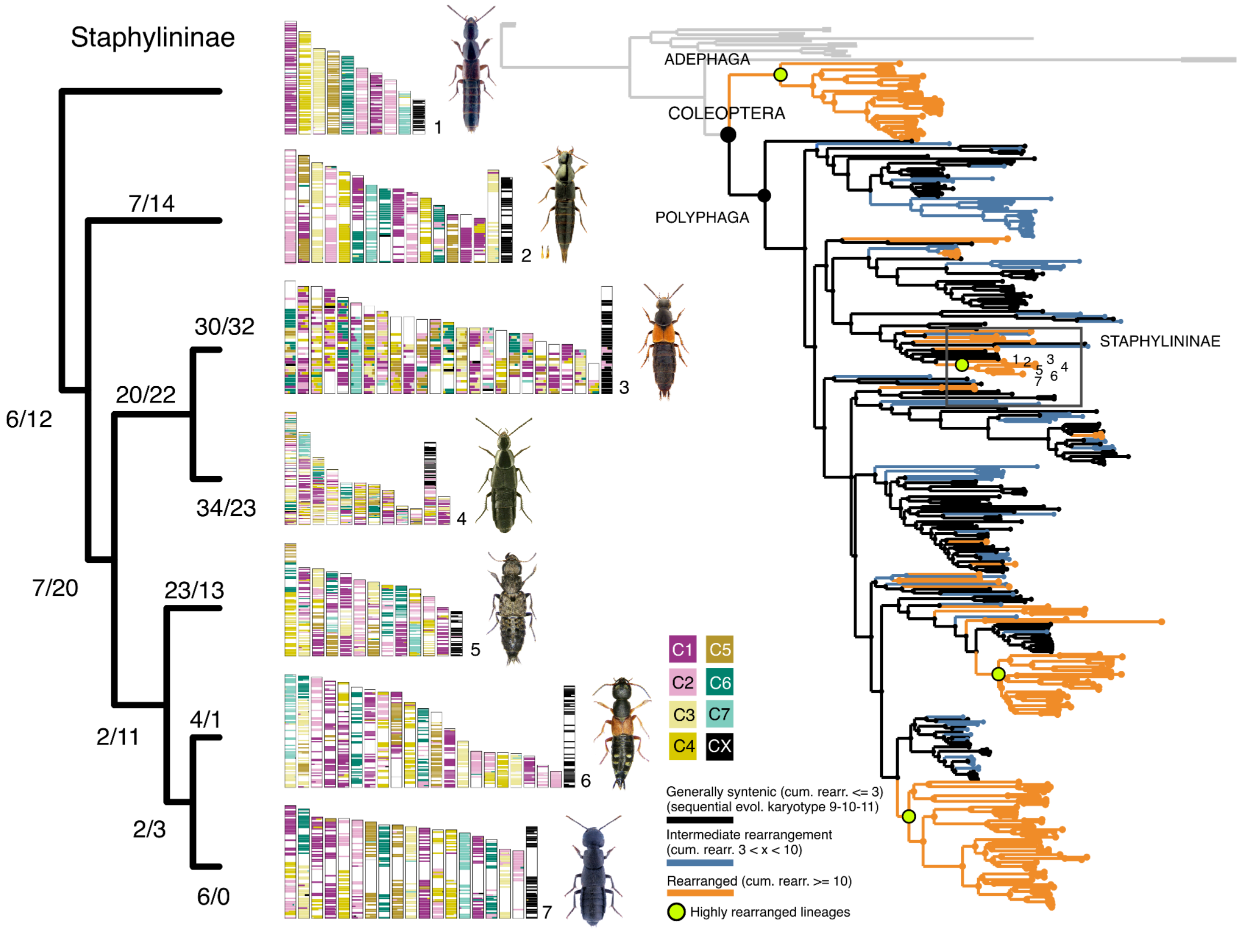


**Fig. S19.**

**Highly rearranged beetle karyotypes evolved repeatedly across major beetle lineages: Staphylininae.** Phylogenetic distribution of rearrangement categories across Coleoptera, with Staphylininae highlighted. Generally syntenic lineages (black) were defined as those with fewer than three cumulative rearrangements and a sequential karyotype transition involving 9, 10, or 11 chromosomes. Intermediate lineages (blue) were defined as those with three to fewer than ten cumulative rearrangements. Highly rearranged lineages (orange) were defined as those with at least ten cumulative rearrangements from the beetle ancestor node n6 to the focal node or tip. Highlighted green nodes mark the most highly rearranged clades. These highly rearranged lineages show distinct patterns of karyotype evolution, ranging from extensive simple fissions to putative chromosome-arm exchanges and combinations of repeated fusions and fissions with highly reshuffled gene order. Labels on the tree correspond to representative species shown in the subset phylogeny and ALG painting plot. ALG painting of representative staphylinine species. Each vertical bar represents one chromosome, and each horizontal segment shows the proportional contribution of each ALG within a 1 Mb window. Left-side branch labels indicate inferred fusion/fission counts along collapsed branches connecting the selected taxa. Horizontal bars indicate different chromosomes, and the vertical ribbons connect the positions of BUSCO genes between species. BUSCO links assigned to the same ALG were merged into broad syntenic ribbons when they formed contiguous or near-contiguous blocks, regardless of their relative gene order between species, and were coloured according to Coleoptera ALG. Singleton and small blocks were removed to improve visibility. 1, *Othius punctulatus*; 2, *Quedius lateralis*; 3, *Philonthus spinipes*; 4, *Philonthus cognatus*; 5, *Ontholestes murinus*; 6, *Staphylinus erythropterus*; 7, *Ocypus olens*. Photo credits: Udo Schmidt, Lech Borowiec, Ilya Zabaluev.

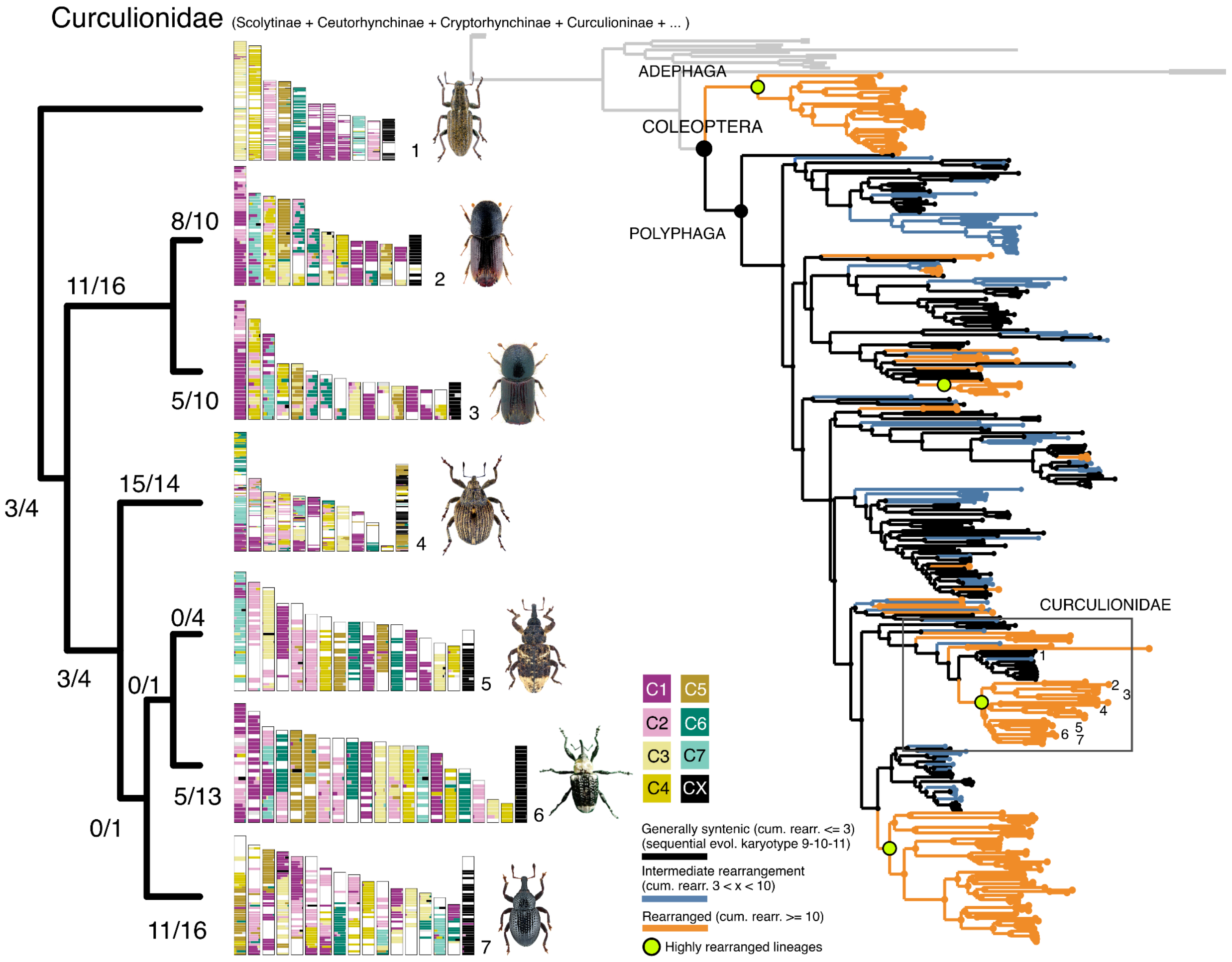


**Fig. S20.**

**Highly rearranged beetle karyotypes evolved repeatedly across major beetle lineages: Curculionidae.** Generally syntenic lineages (black) were defined as those with fewer than three cumulative rearrangements and a sequential karyotype transition involving 9, 10, or 11 chromosomes. Intermediate lineages (blue) were defined as those with three to fewer than ten cumulative rearrangements. Highly rearranged lineages (orange) were defined as those with at least ten cumulative rearrangements from the beetle ancestor node n6 to the focal node or tip. Highlighted green nodes mark the most highly rearranged clades. These highly rearranged lineages show distinct patterns of karyotype evolution, ranging from extensive simple fissions to putative chromosome-arm exchanges and combinations of repeated fusions and fissions with highly reshuffled gene order. Labels on the tree correspond to representative species shown in the subset phylogeny and ALG painting plot. ALG painting of representative curculionine species. Each vertical bar represents one chromosome, and each horizontal segment shows the proportional contribution of each ALG within a 1 Mb window. Left-side branch labels indicate inferred fusion/fission counts along collapsed branches connecting the selected taxa. Horizontal bars indicate different chromosomes, and the vertical ribbons connect the positions of BUSCO genes between species. BUSCO links assigned to the same ALG were merged into broad syntenic ribbons when they formed contiguous or near-contiguous blocks, regardless of their relative gene order between species, and were coloured according to Coleoptera ALG. Singleton and small blocks were removed to improve visibility. 1, *Sitona lineatus*; 2, *Ips sexdentatus*; 3, *Ips typographus*; 4, *Stenocarus ruficornis*; 5, *Cryptorhynchus lapathi*; 6, *Eucryptorrhynchus brandti*; 7, *Leiosoma deflexum*. Photo credits: Udo Schmidt and Huijuan Li.

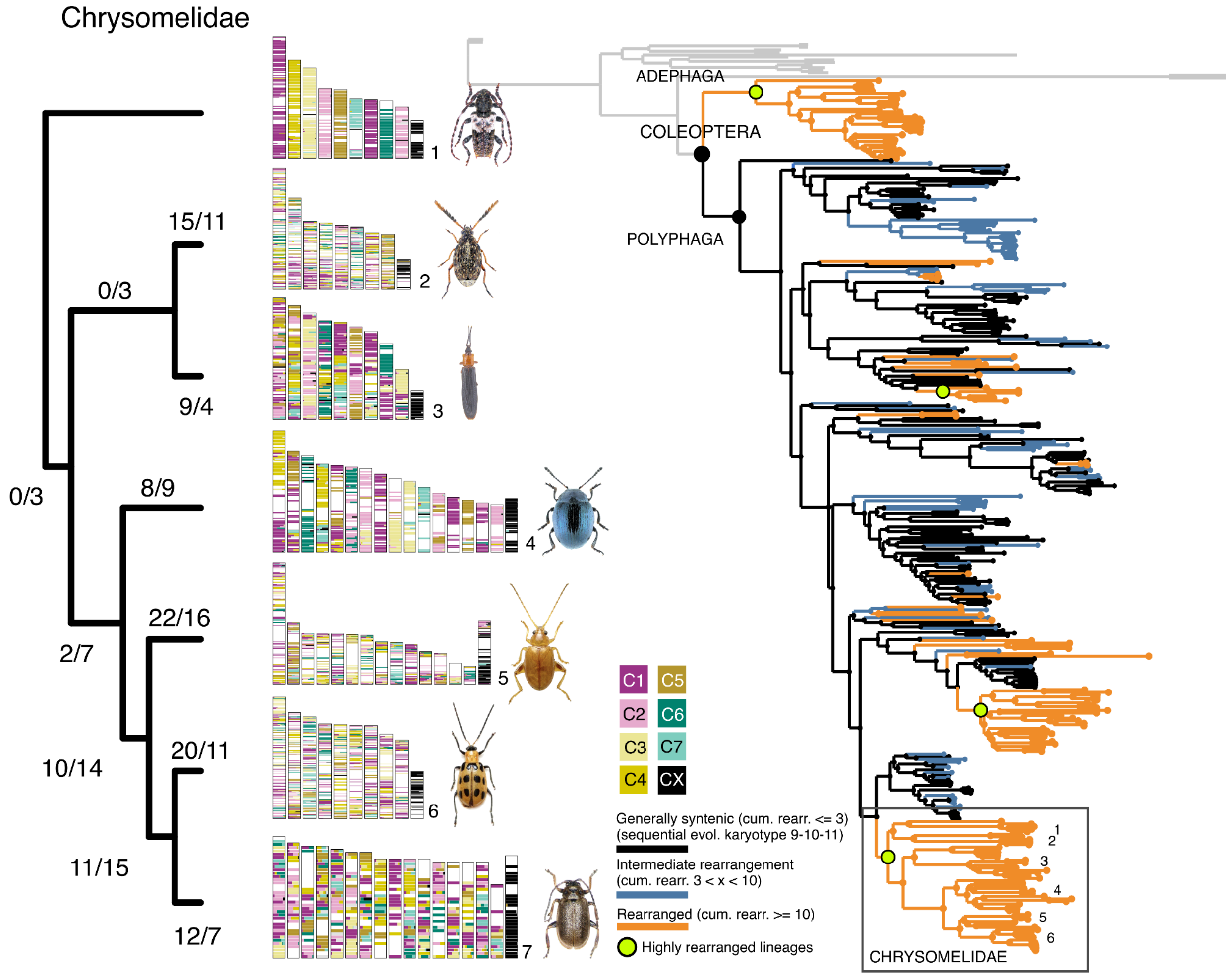


**Fig. S21.**

**Highly rearranged beetle karyotypes evolved repeatedly across major beetle lineages: Chrysomelidae.** Phylogenetic distribution of rearrangement categories across Coleoptera, with Chrysomelidae highlighted. Generally syntenic lineages (black) were defined as those with fewer than three cumulative rearrangements and a sequential karyotype transition involving 9, 10, or 11 chromosomes. Intermediate lineages (blue) were defined as those with three to fewer than ten cumulative rearrangements. Highly rearranged lineages (orange) were defined as those with at least ten cumulative rearrangements from the beetle ancestor node n6 to the focal node or tip. Highlighted green nodes mark the most highly rearranged clades. These highly rearranged lineages show distinct patterns of karyotype evolution, ranging from extensive simple fissions to putative chromosome-arm exchanges and combinations of repeated fusions and fissions with highly reshuffled gene order. Labels on the tree correspond to representative species shown in the subset phylogeny and ALG painting plot. ALG painting of representative chrysomeline species. Each vertical bar represents one chromosome, and each horizontal segment shows the proportional contribution of each ALG within a 1 Mb window. Left-side branch labels indicate inferred fusion/fission counts along collapsed branches connecting the selected taxa. Horizontal bars indicate different chromosomes, and the vertical ribbons connect the positions of BUSCO genes between species. BUSCO links assigned to the same ALG were merged into broad syntenic ribbons when they formed contiguous or near-contiguous blocks, regardless of their relative gene order between species, and were coloured according to Coleoptera ALG. Singleton and small blocks were removed to improve visibility. 1, *Pogonocherus hispidulus*; 2, *Bruchidius varius*; 3, *Brontispa longissima*; 4, *Phaedon cochleariae*; 5, *Longitarsus flavicornis*; 6, *Diabrotica undecimpunctata*; 7, *Lochmaea capreae*. Photo credits: Udo Schmidt and Alexis Vincent.

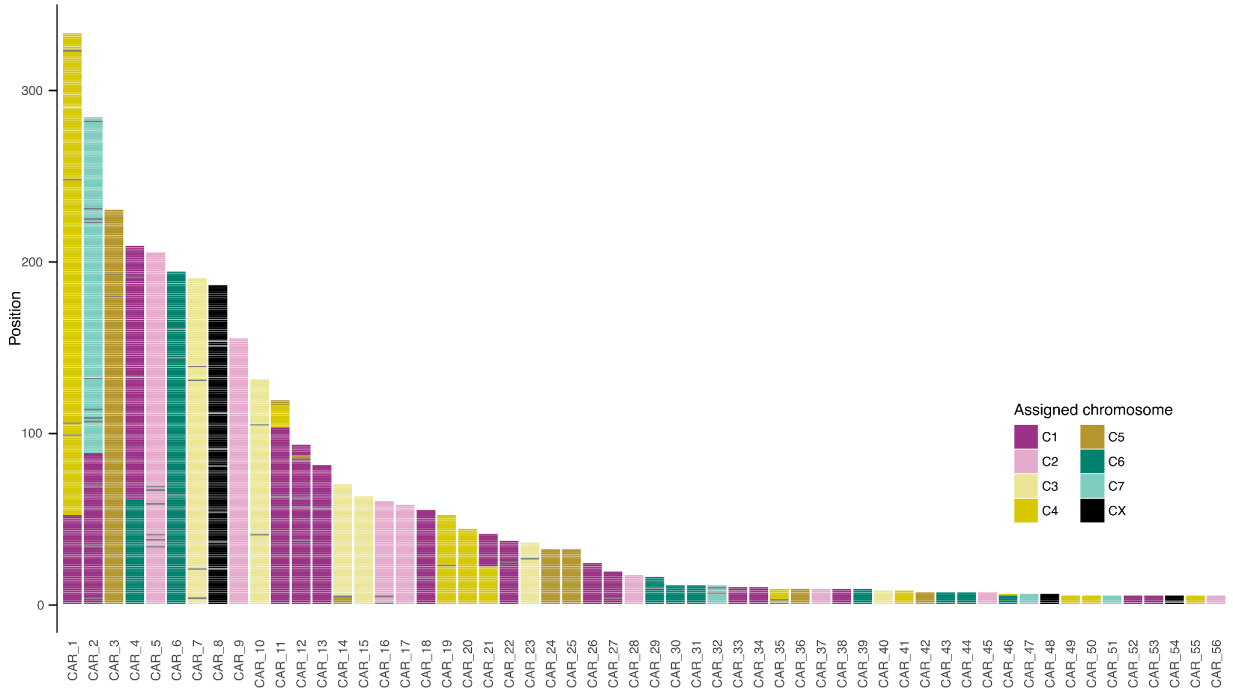


**Fig. S22.**

**Ancestral gene order reconstruction of coleopteran ALGs inferred using AGORA.** Each row represents a contiguous ancestral region (CAR) inferred by AGORA and containing at least five BUSCO markers. Marker positions are shown along the x-axis, and each tile represents one BUSCO gene, coloured according to its assigned Coleoptera ALG. Most CARs were assigned to a single ALG, although some contained more than one ALG. In these cases, the boundaries between ALGs were generally clear, suggesting that multi-ALG CARs may reflect artefacts of the AGORA (*22*) reconstruction.

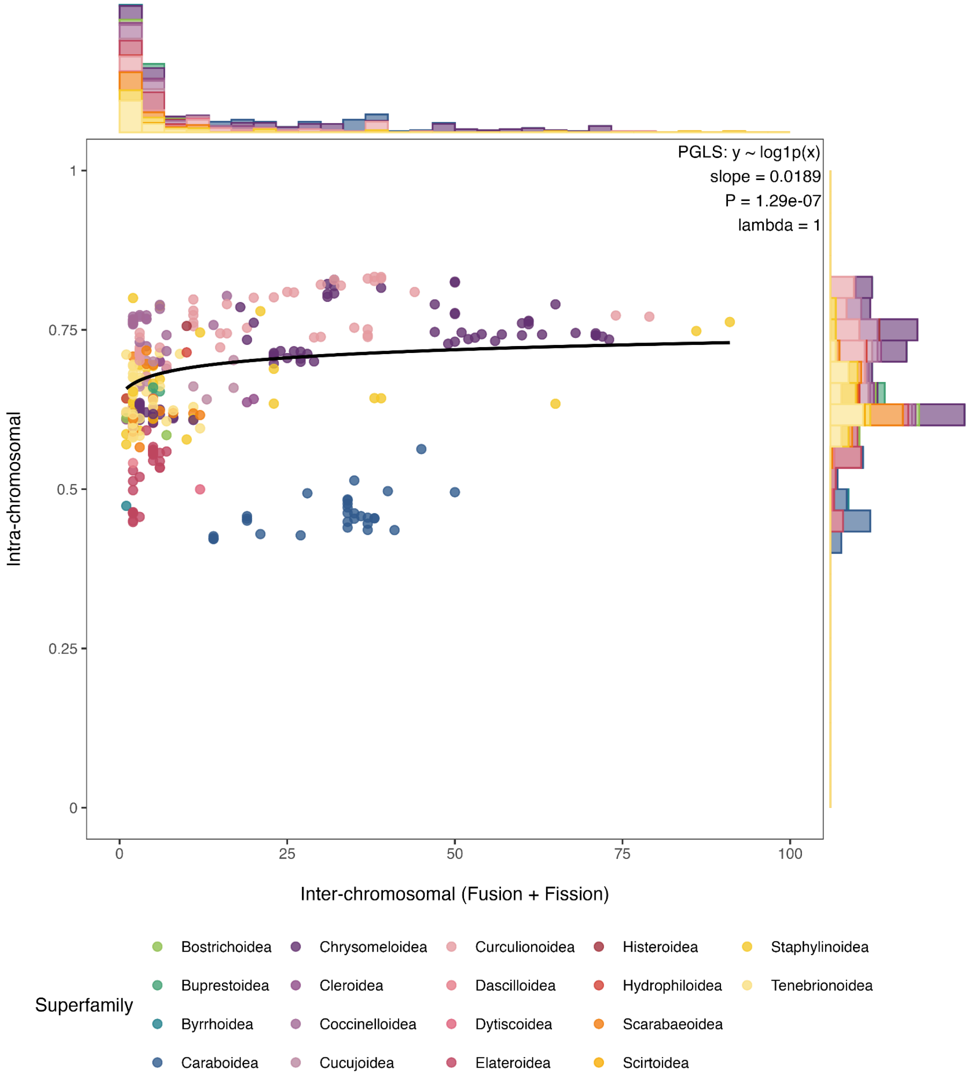


**Fig. S23.**

**Association between intra- and interchromosomal rearrangement in Coleoptera.** Each point represents one beetle species, with interchromosomal rearrangement count plotted on the x axis and intrachromosomal rearrangement index plotted on the y axis. Interchromosomal rearrangement was defined as the total number of rearrangements inferred from Syngraph reconstruction, whereas intrachromosomal rearrangement was estimated using gene-order fragmentation index (*FI*) based on AGORA reconstruction. Points are coloured by superfamily. The black line shows the fitted PGLS relationship from the model intrachromosomal ~ log1p(interchromosomal), accounting for shared ancestry among species using Pagel's λ. The positive fitted relationship indicates that beetle species with more interchromosomal rearrangements tend to show higher intrachromosomal rearrangement, although many species with low interchromosomal counts still show variable intrachromosomal rearrangement.

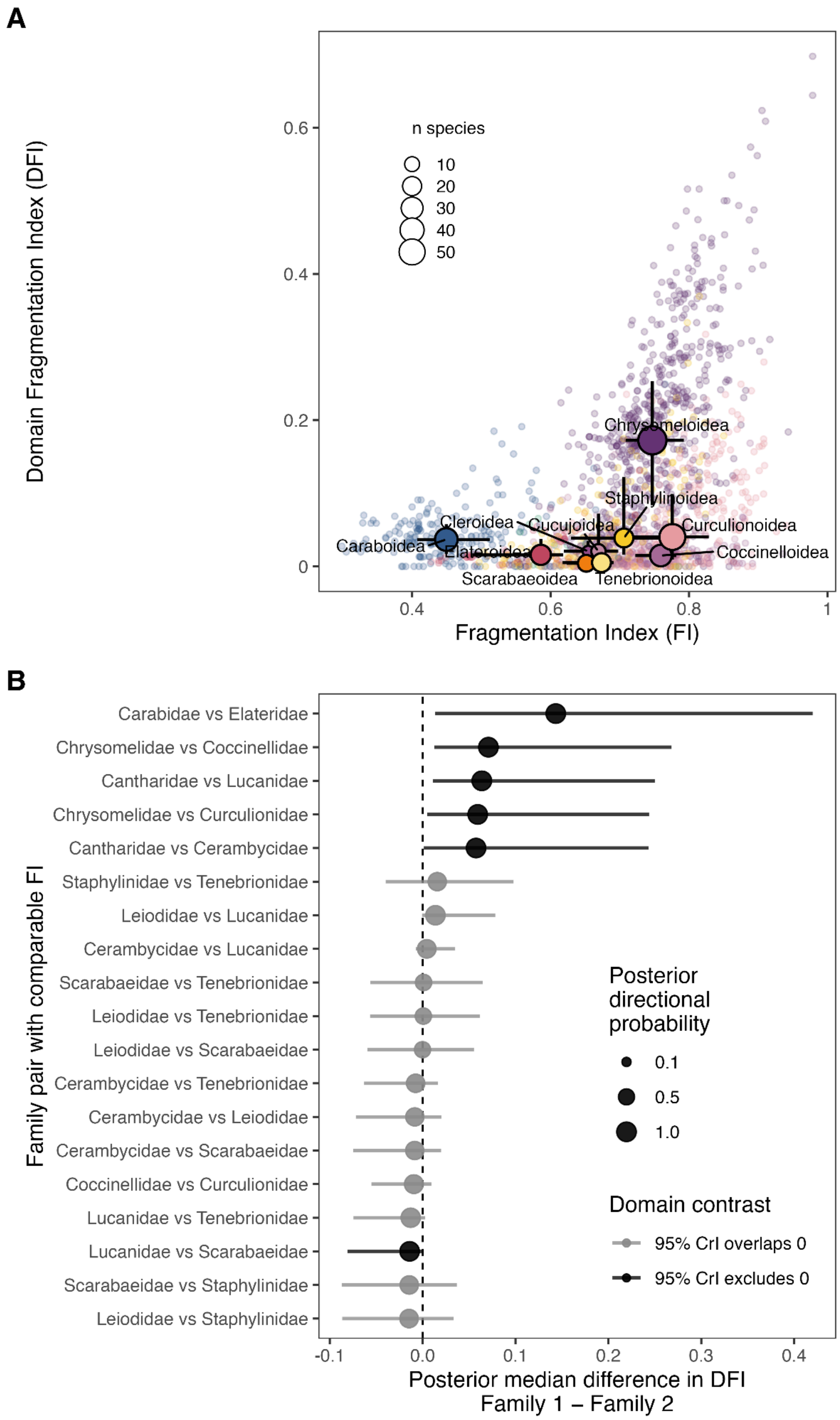


**Fig. S24.**

**Similar gene-order fragmentation can correspond to different ancestral-domain mixing patterns. A.** Relationship between chromosome-level gene-order fragmentation index (*FI*) based on AGORA and domain fragmentation index (*DFI*) for fused chromosomes. Faint points represent individual fusion chromosomes, coloured by superfamily. Large points show superfamily medians, with horizontal and vertical bars indicating interquartile ranges. Point size indicates the number of species represented in each superfamily. **B.** Pairwise posterior contrasts in predicted domain fragmentation among family pairs with comparable *FI*. Comparable FI was defined as an absolute posterior median FI difference ≤ 0.05 with a 95% credible interval overlapping zero. Points show posterior median domain-fragmentation differences, and horizontal bars show 95% credible intervals. Black points/bars indicate contrasts where the 95% credible interval excludes zero; grey indicates contrasts where it overlaps zero. Point size shows the posterior directional probability. Positive values indicate higher domain fragmentation in the first family of the pair, and negative values indicate higher domain fragmentation in the second family. Only superfamilies or families with at least five species were highlighted.

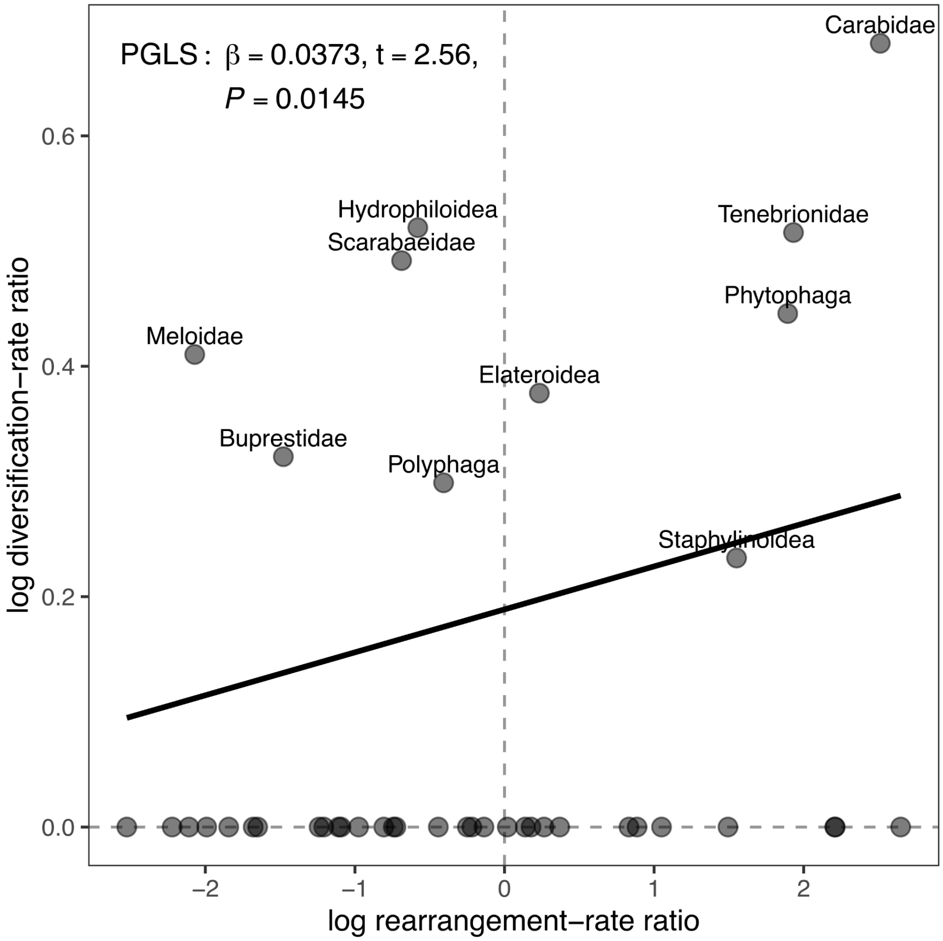


**Fig. S25.**

**Phylogenetic association between chromosomal rearrangement-rate increases and diversification-rate increases across beetle clades.** Each point represents a clade-level comparison. The x-axis shows the log rearrangement-rate ratio, calculated as the rearrangement rate of the focal clade relative to its phylogenetic background. The y-axis shows the log diversification-rate ratio, calculated as the diversification rate of the focal clade relative to its parent diversification regime. Dashed lines indicate no change relative to background or parent rate. The fitted line shows the node-level PGLS relationship, accounting for phylogenetic non-independence among clades. Increases in chromosomal rearrangement rate were significantly associated with increases in diversification rate ($\beta = 0.0373$, $t = 2.56$, $p = 0.0145$).

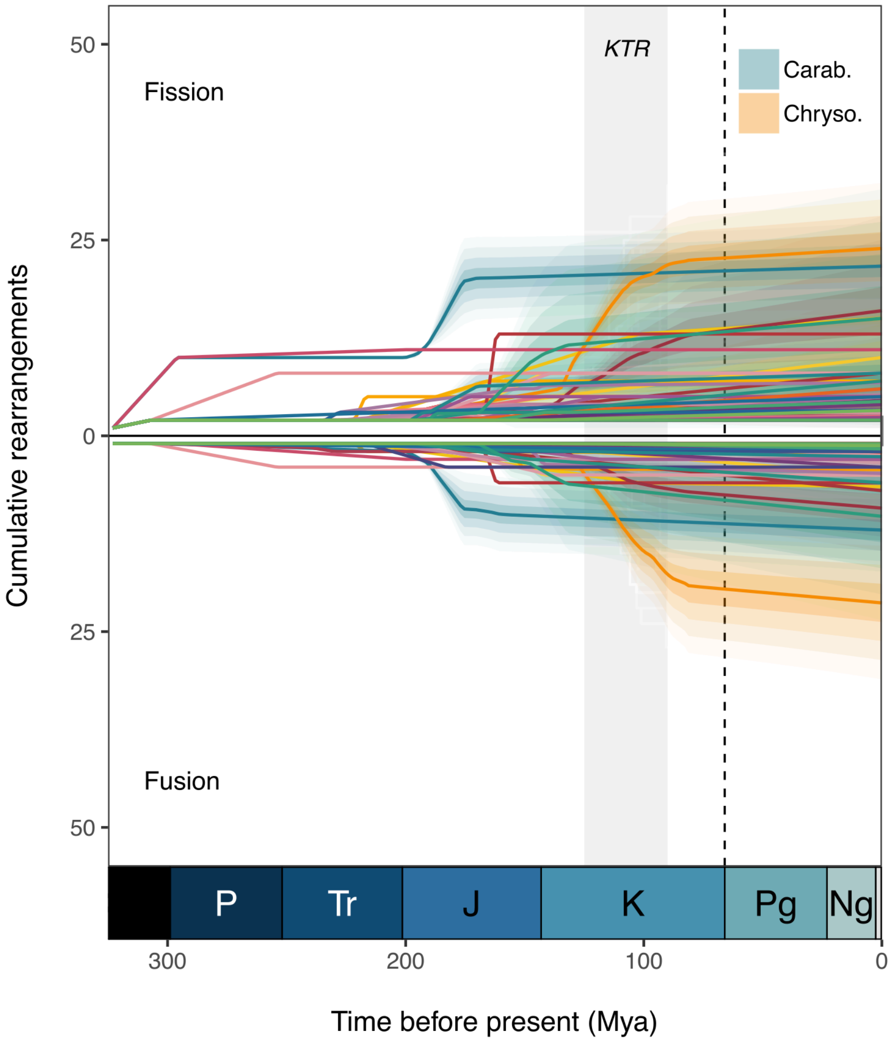


**Fig. S26.**

**Temporal dynamics of fission and fusion events across beetle superfamilies.**
Cumulative rearrangement trajectories through time for each beetle superfamily. Positive values indicate cumulative fission events, whereas negative values indicate cumulative fusion events. Each line represents the interpolated rearrangement trajectory for one superfamily estimated using functional data analysis. Shaded regions indicate uncertainty around selected trajectories, with Caraboidea and Chrysomeloidea highlighted. The geological timescale is shown below the plot, with KTR (Angiosperm radiation) shown, and the dashed vertical line marks the K–Pg boundary.

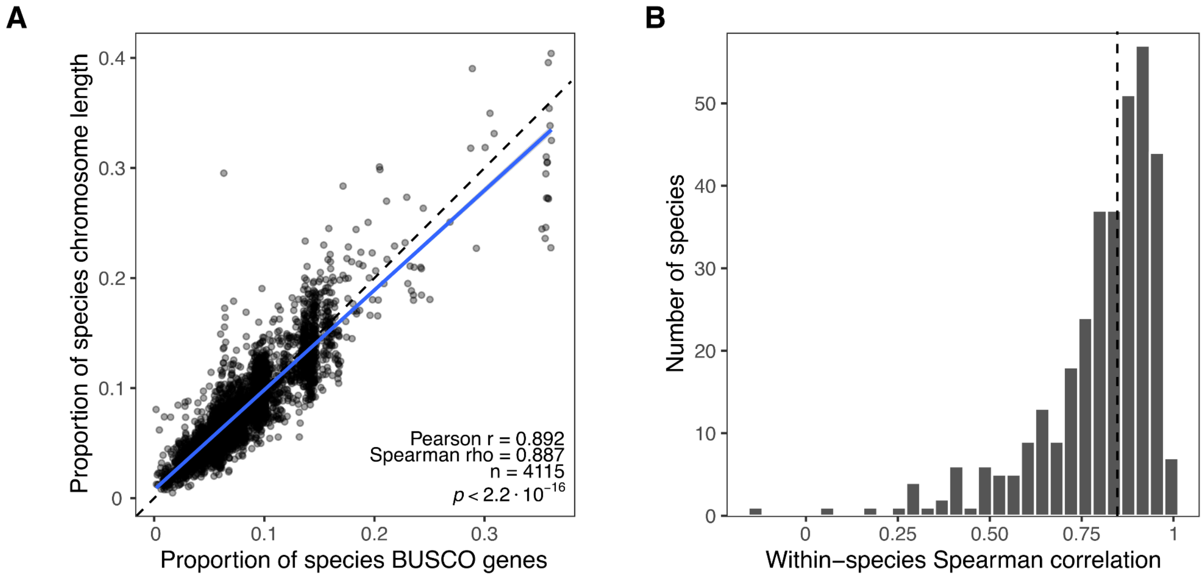


**Fig. S27.**

**Validation of chromosome BUSCO content as a proxy for relative chromosome size. A.** The proportion of each species' BUSCO genes assigned to a chromosome was strongly correlated with the chromosome's proportion of total assembled chromosome length (Pearson's $r = 0.892$, Spearman's $\rho = 0.887$, $n = 4{,}115$ chromosomes). The dashed line represents one-to-one correspondence. **B.** Distribution of Spearman correlations calculated separately within each species between chromosome BUSCO count and physical chromosome length. The dashed line indicates the median species-level correlation.

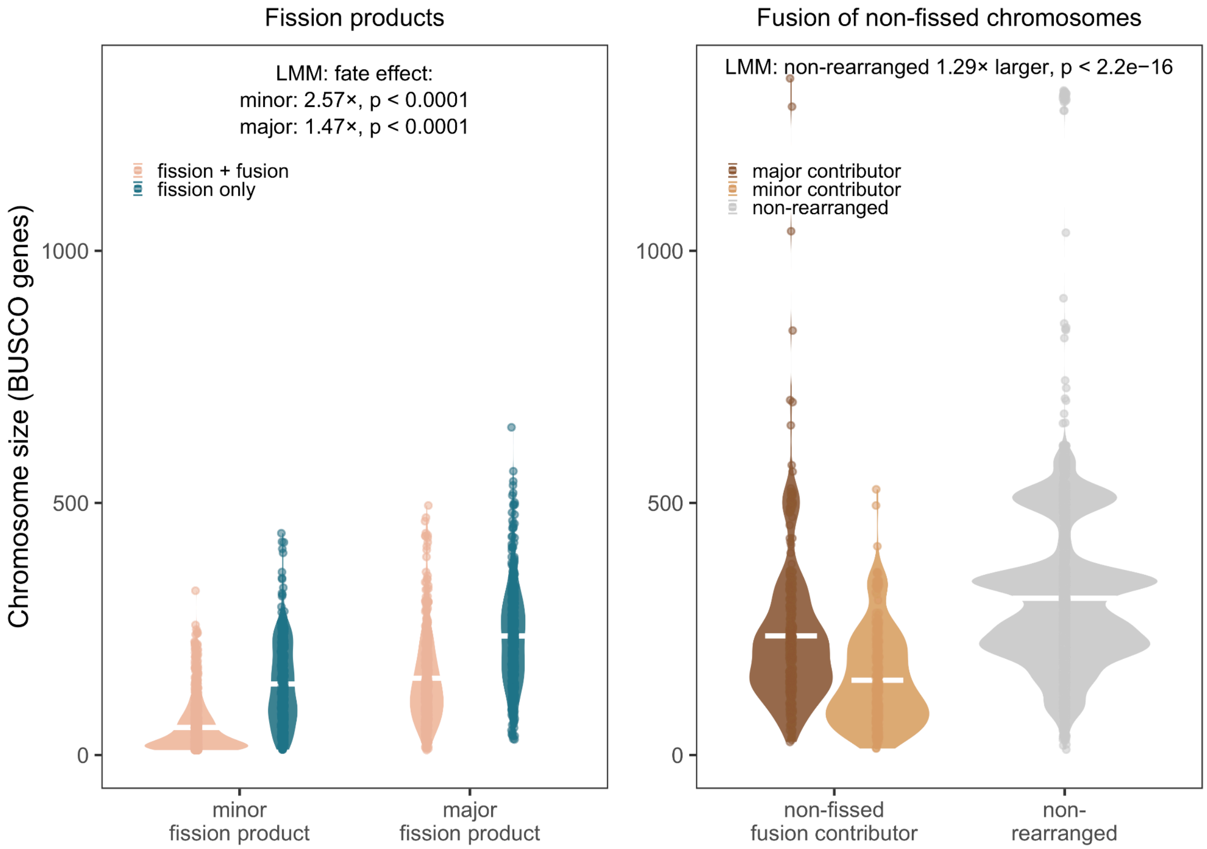


**Fig. S28.**

**Chromosome sizes across rearrangement outcomes**. Chromosome sizes, measured by BUSCO marker count, across different rearrangement outcomes. The left panel shows the size distribution of fission products. Within each fission event, products were classified as minor or major fragments according to their relative size. Fission products that subsequently fused were smaller than fission-only products in both minor and major classes. The right panel shows the size distributions of chromosomes contributing to fusion events without prior fission. Fusion contributors were classified as minor or major according to their relative contribution within each fusion event and were smaller than non-rearranged chromosomes. Linear mixed models were fitted to log-transformed BUSCO counts, with parent and/or child lineages included as random effects. White horizontal bars indicate group means.

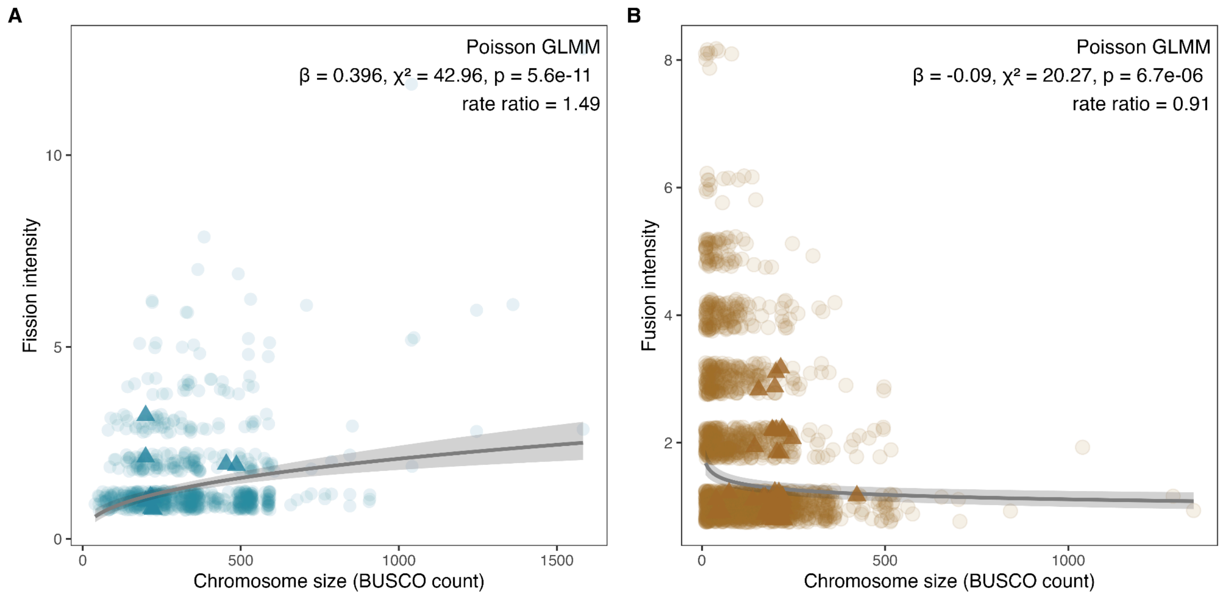


**Fig. S29.**

**Chromosome size predicts rearrangement intensity**. Relationship between chromosome size, measured as BUSCO marker count, and inferred rearrangement counts. **A.** Fission intensity increases with chromosome size, indicating that larger chromosomes tend to undergo more extensive fission. **B.** Fusion intensity decreases with chromosome size, indicating that smaller chromosomes tend to be involved in more fusion events. Each point represents an individual chromosome. Grey lines and shaded bands show fixed-effect predictions and 95% confidence intervals from Poisson GLMMs with parent and child lineages included as random intercepts. For fusion, fusion event identity was also included as a random intercept. circles = autosomes; triangles = X chromosomes.

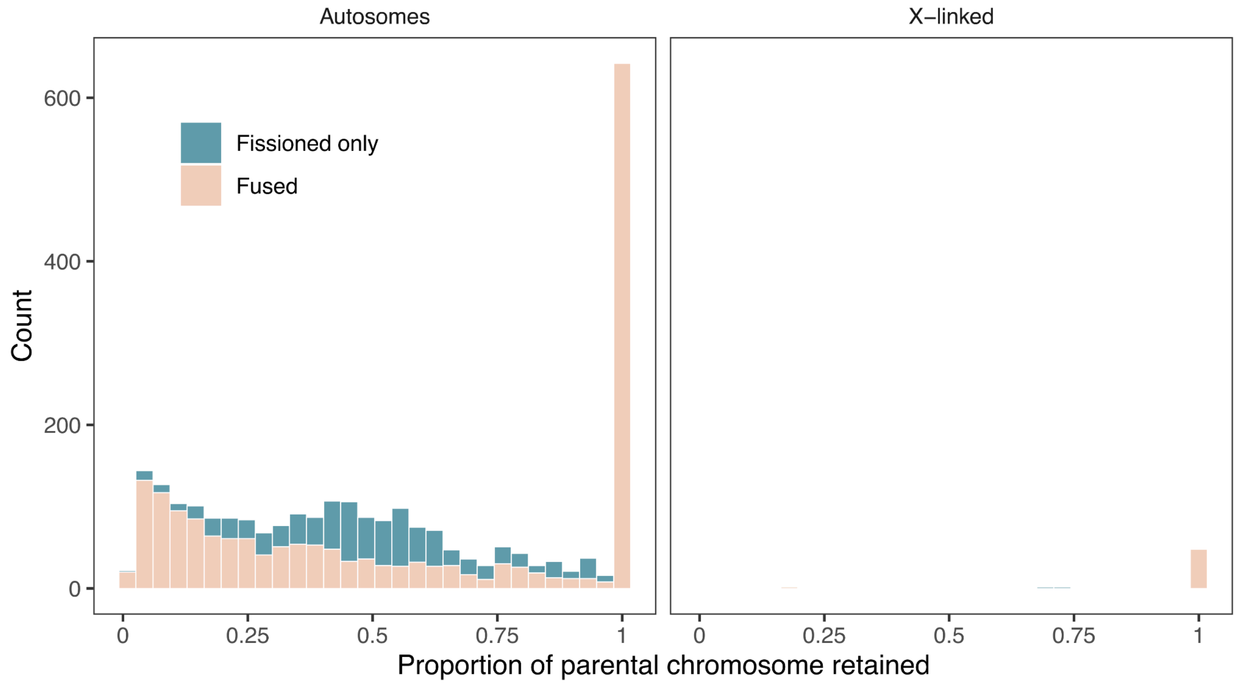


**Fig. S30.**

**Relative sizes of chromosomes involved in fission-only and fusion-associated rearrangements.** Histograms show the size of rearranged chromosomes relative to their inferred parental chromosomes, measured as the proportion of BUSCO markers retained from the parental chromosome. Fissioned-only chromosomes represent fragments produced by fission. Fusion-associated chromosomes include both previously fissioned chromosomes that later fused and fusion of non-rearranged parental chromosomes, the latter appearing at a relative size of 1. Histograms are shown separately for autosomes and X-linked chromosomes.

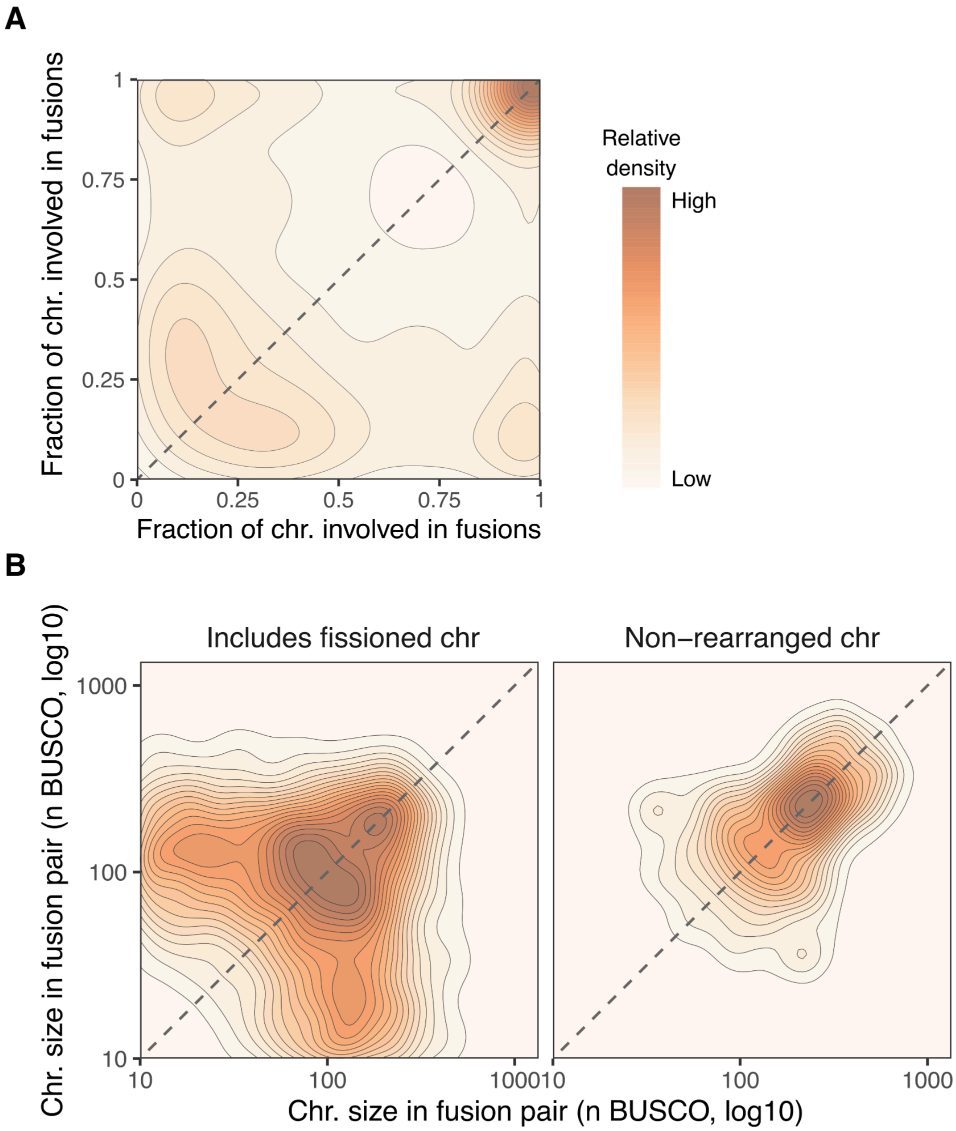


**Fig. S31.**

**Relative and absolute sizes of chromosomes involved in fusion events.** Density plots showing the size relationships between chromosomes involved in fusion events, after excluding weakly supported fusion pairs. **A.** Relative sizes of chromosome pairs involved in fusion events, measured as the fraction of the parental chromosome retained by each fusion partner. Density near the upper-right corner indicates fusion between two largely unrearranged chromosomes, whereas density near the upper-left and lower-right corners indicates fusion between a large, largely unrearranged chromosome and a smaller fission-derived fragment. This shows that many fusion events involve either two non-rearranged chromosomes or one non-rearranged chromosome and a smaller fragment generated by prior fission. **B.** Absolute sizes of chromosomes in fusion pairs, measured as BUSCO marker counts on a log10 scale. Fusion pairs are separated according to whether they include a fission-derived chromosome or consist only of non-rearranged chromosomes. Dashed lines indicate equal size between fusion partners.

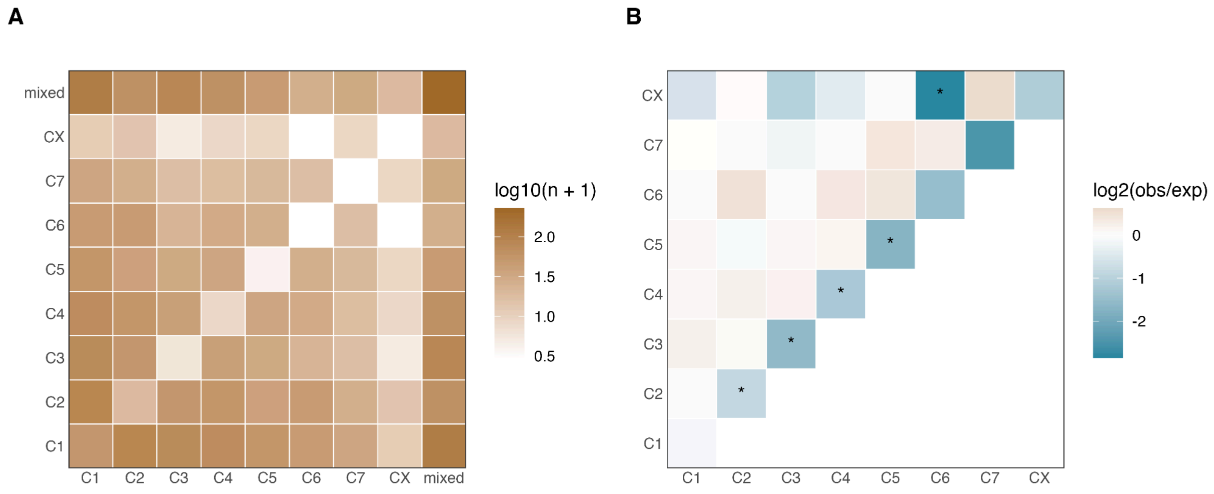


**Fig. S32.**

**Co-fusion frequencies among Coleoptera ancestral linkage groups. A.** Observed counts of fusion-partner pairs among chromosomes classified by their dominant Coleoptera ALG, including mixed-origin chromosomes. Mixed-origin chromosomes represent fusion contributors without a single dominant ALG assignment. **B.** Enrichment and depletion of co-fusion pairs after excluding mixed-origin contributors. Tile colour shows log2 ratio of observed to expected pair frequency from permutation tests in which ALG labels were shuffled among fusion contributors while preserving fusion-event structure and overall ALG abundance. Blue indicates pairs occurring less often than expected, and white indicates values close to expectation. Asterisks mark pairs significant after Benjamini–Hochberg correction. Same-ALG pairs were generally depleted, suggesting that fragments derived from the same ancestral linkage group rarely re-fuse.

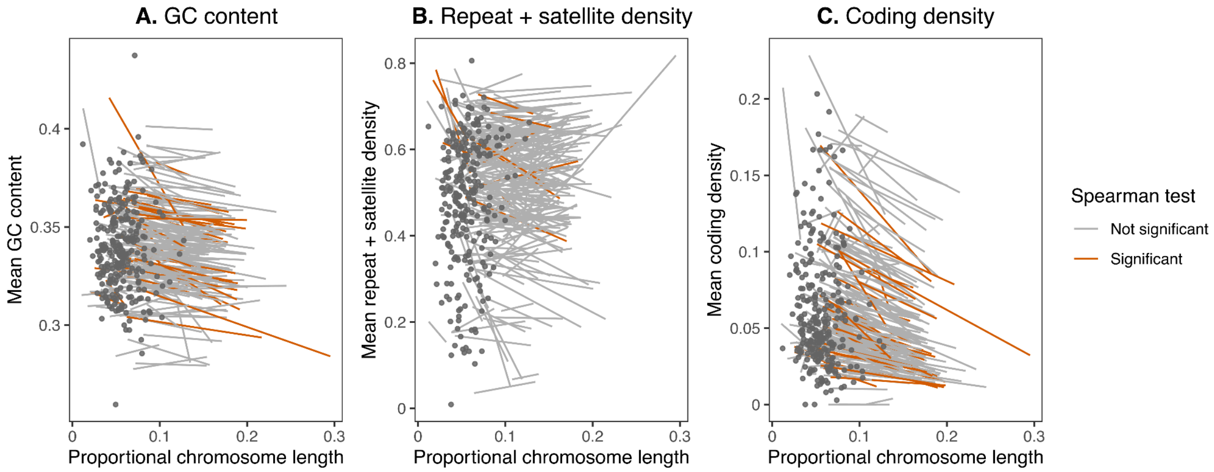


**Fig. S33.**

**Associations between relative chromosome size and genomic features.** For each species, Spearman correlations were calculated across non-rearranged autosomes only between proportional chromosome length (length divided by genome size) and mean GC content, repeat plus satellite density, or coding density. Each line represents the fitted visual trend for a single species. Orange lines indicate significant Spearman correlations after Benjamini–Hochberg correction, whereas grey lines indicate non-significant correlations. CX chromosomes (shown as dark grey points) and rearranged autosomes were excluded from the correlation analyses and fitted trends. Not significantly different correlation might be due to the small number of chromosomes available per species. Most species showed negative Spearman correlations (GC: 80.2%, median $\rho = -0.600$; coding density: 82.9%, median $\rho = -0.643$), although only 34/217 and 32/217 species, respectively, remained significant after BH correction. Repeat plus satellite density showed a weaker and more heterogeneous autosomal pattern, with 11/211 species significant after correction and a modest negative median correlation (median $\rho = -0.143$; 57.8% negative).

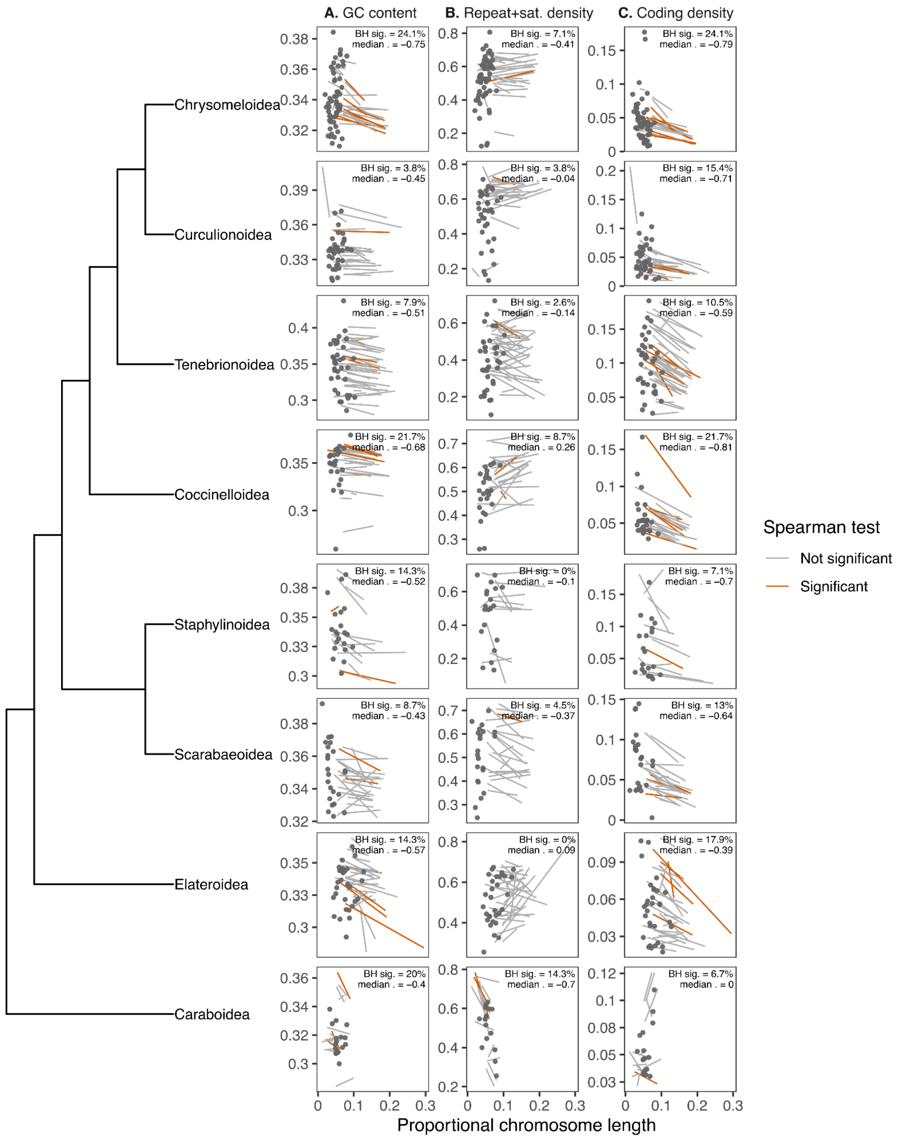


**Fig. S34.**

**Correlation of genomic features with chromosome size in beetle superfamilies.** For each species, Spearman correlations were calculated across non-rearranged autosomes only between proportional chromosome length (length divided by genome size) and mean GC content, repeat plus satellite density, or coding density. Each line represents the fitted visual trend for a single species. Line colour indicates whether the within-species Spearman correlation was significant after Benjamini–Hochberg correction. CX chromosomes (shown as dark grey points) and rearranged autosomes were excluded from the correlation analyses and fitted trends. Text within each panel reports the percentage of significant autosomal species-level correlations and the median Spearman correlation coefficient for that superfamily. Only superfamilies represented by at least eight species after excluding CX chromosomes and rearranged autosomes are shown.

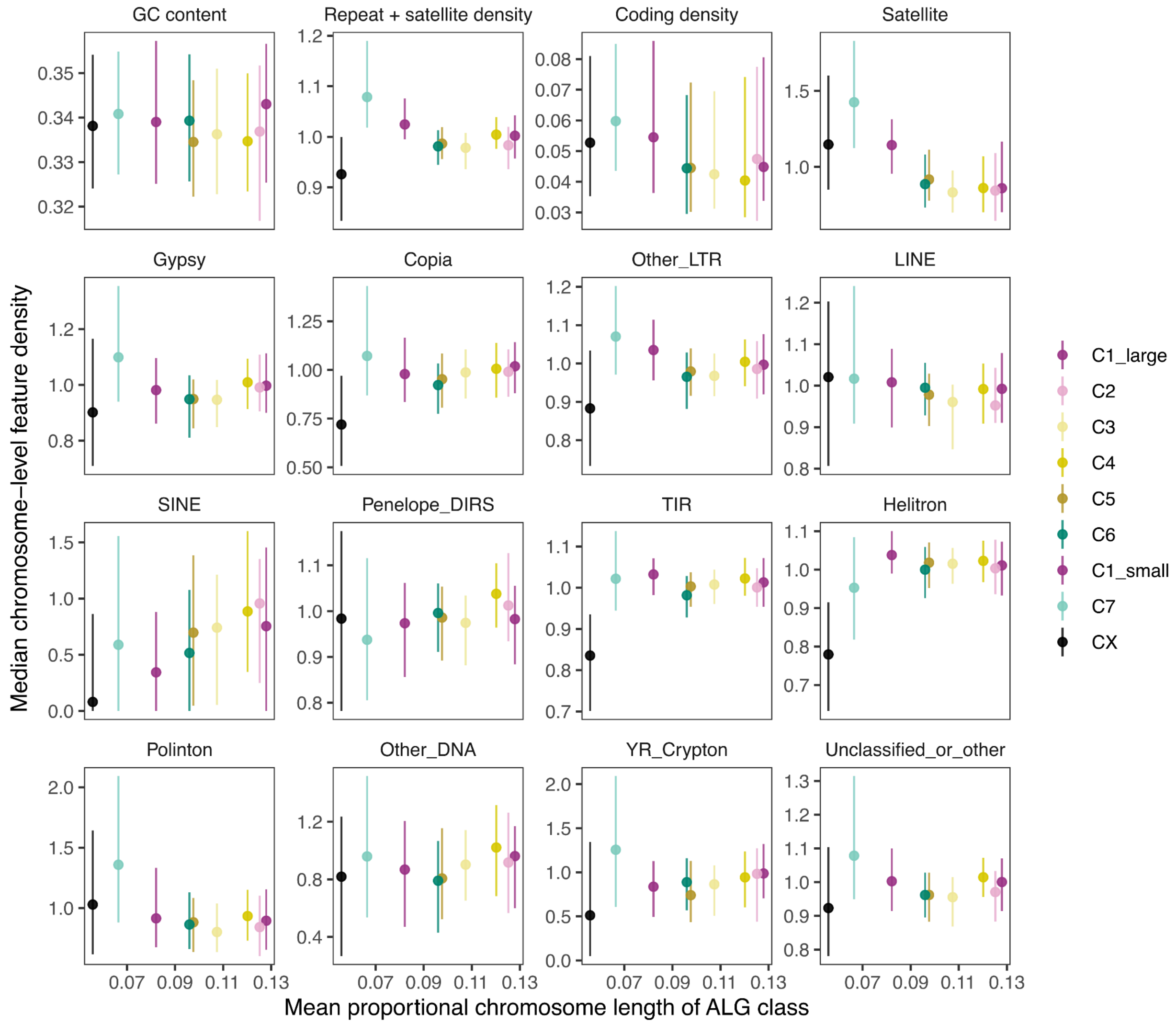


**Fig. S35.**

**Relationship between genomic features and repetitive element density and proportional chromosome length for each Coleoptera ALG.** Mean proportional chromosome length, calculated as chromosome length divided by genome size, was compared with the mean density of genomic features and repeat classes for each Coleoptera ALG. To enable comparison between species with different average repeat densities, the density of each repetitive element class was scaled by the mean repeat density of the genome. Only chromosomes that had not undergone fusion or fission were included. Points and error bars show the mean and confidence interval for each ALG and are coloured according to Coleoptera ALG.

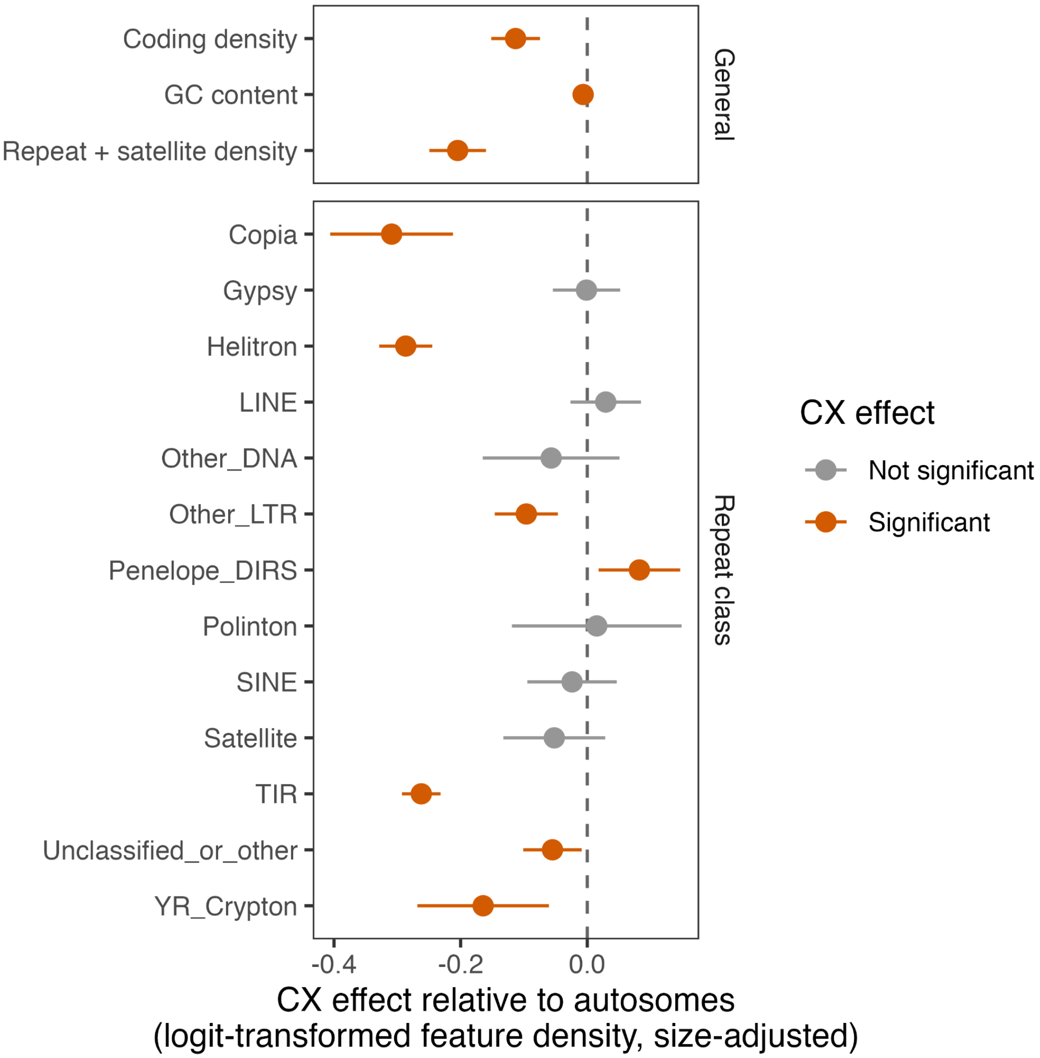


**Fig. S36.**

**Size-adjusted CX effects on genomic features.** Size-adjusted CX effects on genomic feature density. For each feature, linear mixed models were fitted with logit-transformed chromosome-level feature density as the response, proportional chromosome length and CX identity as fixed effects, and species as a random intercept. Points show the estimated CX effect relative to autosomes after accounting for chromosome size. Horizontal bars show approximate 95% confidence intervals. Negative values indicate lower feature density on CX, whereas positive values indicate higher feature density on CX. Orange points indicate significant CX effects after Benjamini–Hochberg correction; grey points indicate non-significant effects.

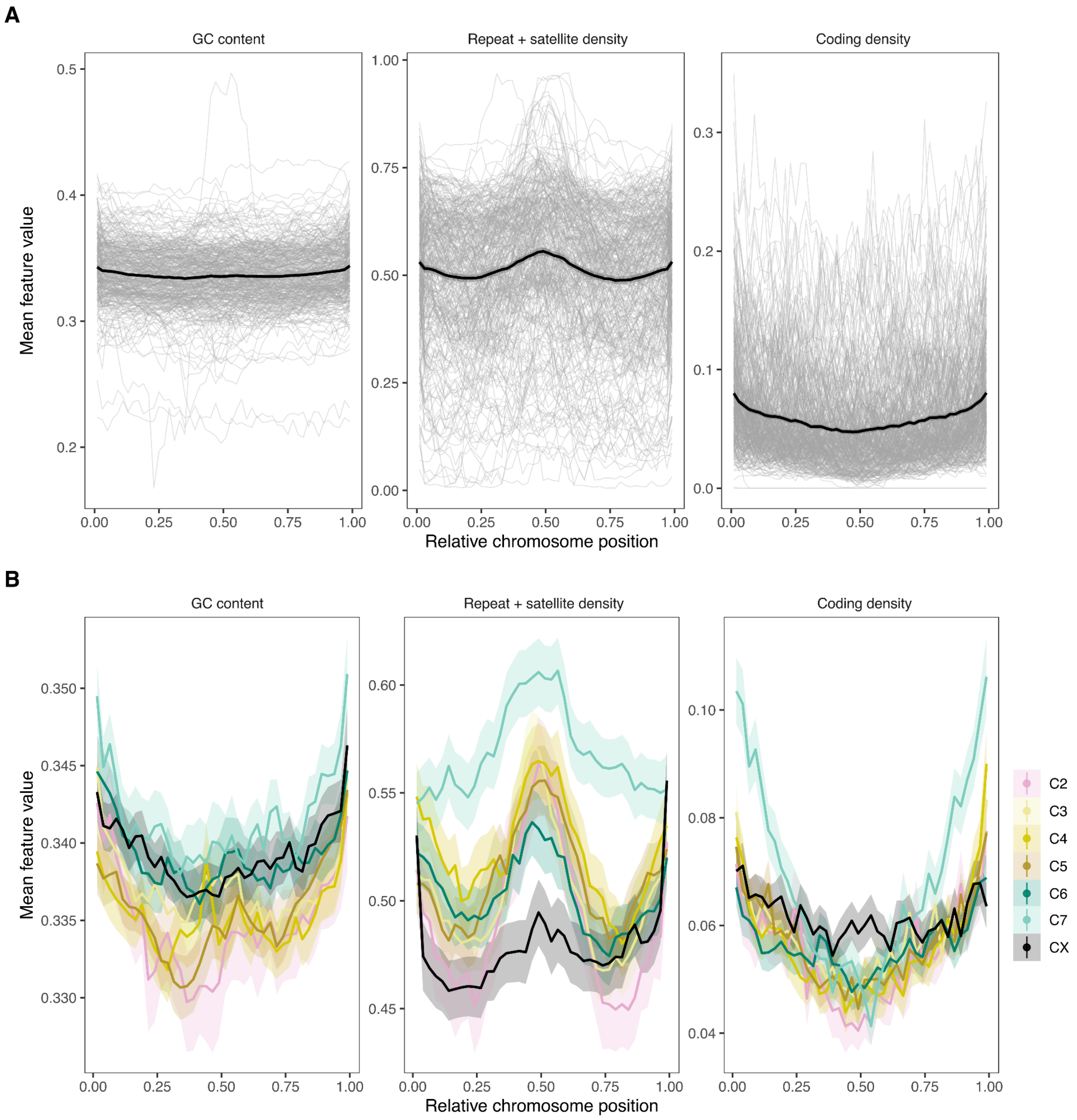


**Fig. S37.**

**Sequence patterns in coleopteran chromosomes. A.** Chromosome-position landscapes for GC content, repeat plus satellite density, and coding density across beetles. Each chromosome was divided into 100 windows, with relative chromosome position scaled from 0 to 1. Grey lines represent individual chromosome-level binned landscapes, black lines show the mean across chromosomes, and shaded bands indicate ±1 SE across chromosome-level means. **B.** ALG-specific chromosome-position landscapes for the same features, using intact chromosomes assigned to a single Coleoptera ALG. Lines show mean binned feature values for each Coleoptera ALG across relative chromosome position, and shaded bands indicate ±1 SE across chromosome-level means. C1 was omitted from panel B as it was ancestrally rearranged to all extant species.

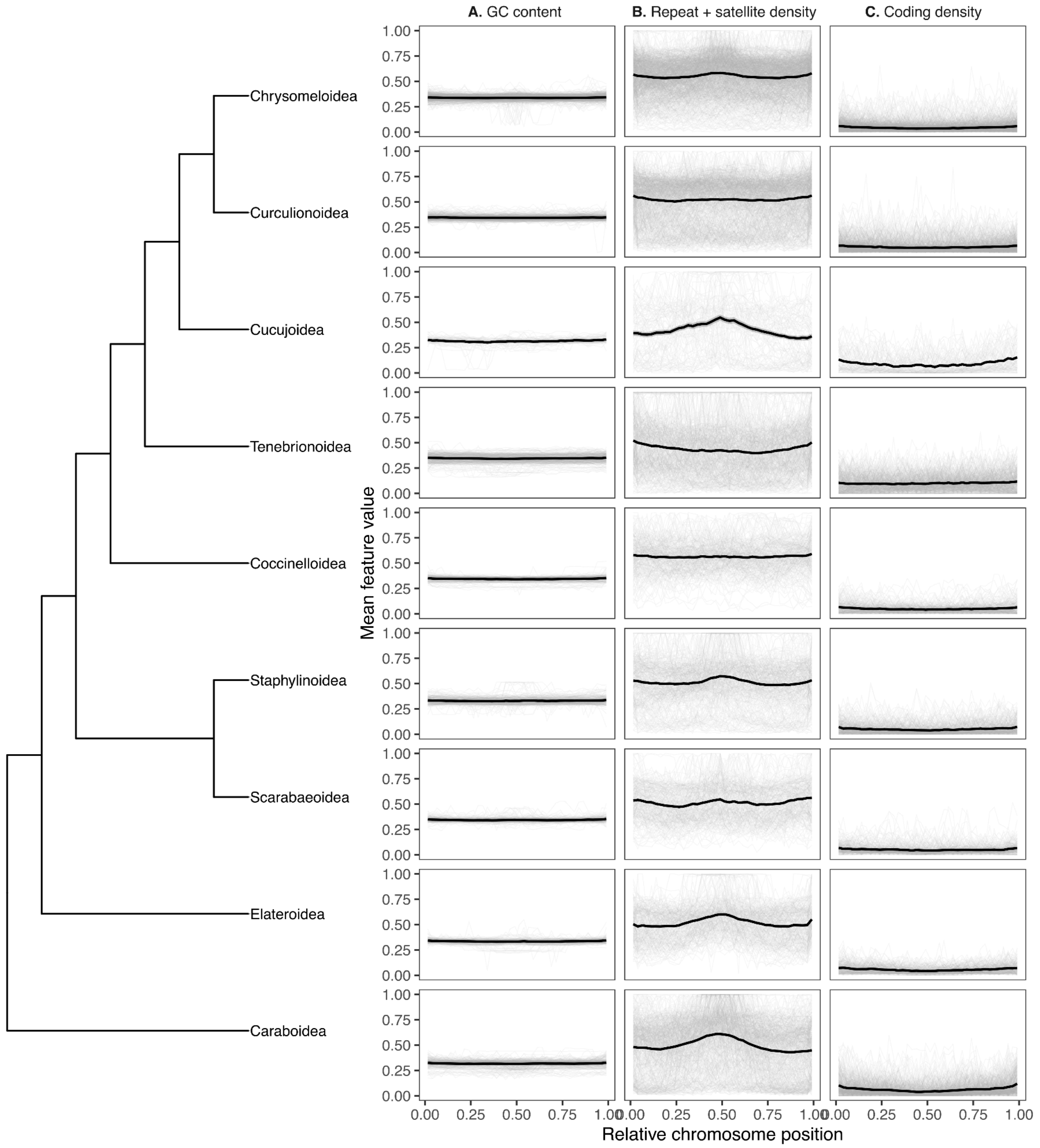


**Fig. S38.**

**Sequence patterns in coleopteran chromosomes across beetle superfamilies.**
Chromosome-position landscapes for GC content, repeat plus satellite density, and coding density across beetle superfamilies. Each chromosome was divided into 100 windows, with relative chromosome position scaled from 0 to 1. Grey lines represent individual chromosome-level binned landscapes, black lines show the mean across chromosomes, and shaded bands indicate ±1 SE across chromosome-level means. Only superfamilies represented by at least eight species are shown.

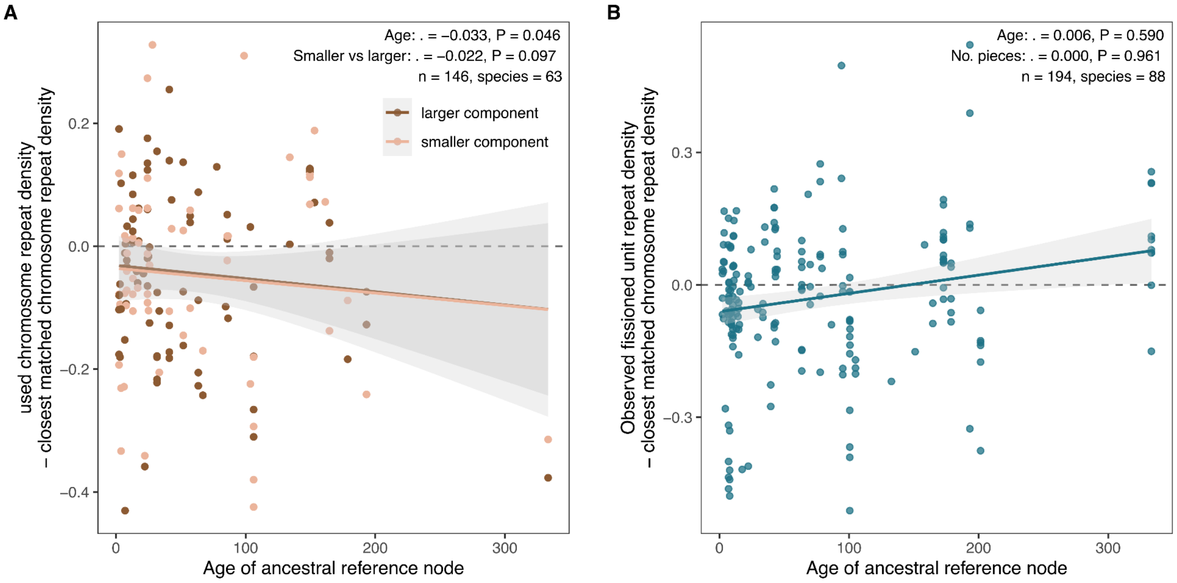


**Fig. S39.**

**Temporal association between rearrangement age and repeat-density deviation**. Repeat-density deviation was calculated relative to the closest matched chromosome from a comparator species within the same genus or subfamily. Points show individual events, with dashed horizontal lines indicating no deviation from the closest matched chromosome. **A.** Fusion comparisons. The y-axis shows the fused chromosome repeat density minus the closest matched chromosome repeat density. Points are coloured by fusion component size class, and fitted lines show linear trends for larger and smaller components separately. Fusion showed a weak negative association with ancestral reference-node age after family filtering ($\beta$ = -0.033, $p$ = 0.046), while smaller components tended to have lower deltas than larger components but not significantly ($\beta$ = -0.022, $p$ = 0.097; $n$ = 146, 63 species). **B.** Fission comparisons. The y-axis shows repeat density of the observed fissioned unit minus the closest matched chromosome repeat density. Fission showed no significant association with ancestral reference-node age after family filtering ($\beta$ = 0.006, $p$ = 0.590), and the number of fission pieces was not associated with repeat-density deviation ($\beta$ = 0.000, $p$ = 0.961; $n$ = 194, 88 species). Shaded ribbons indicate 95% confidence intervals around linear fits.

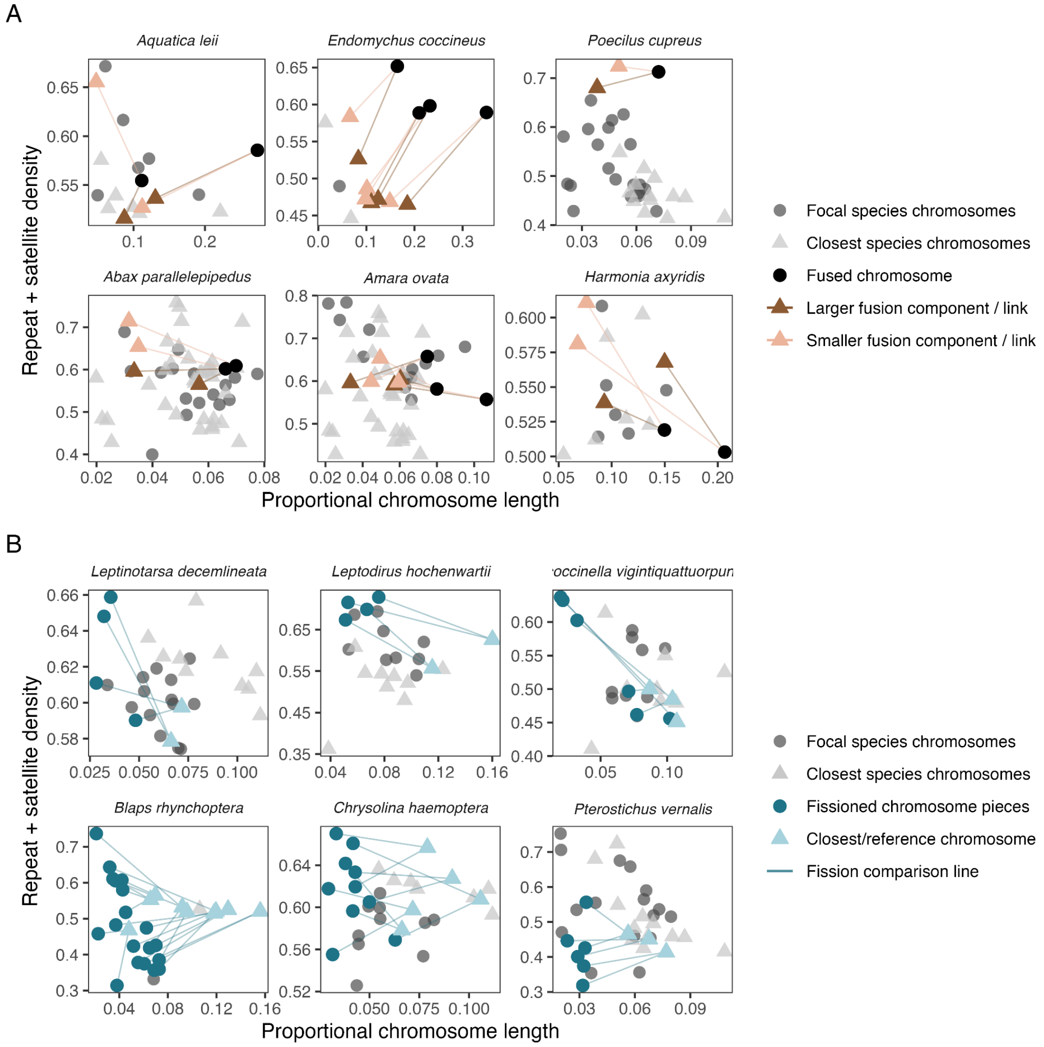


**Fig. S40.**

**Relationships between chromosome size and repeat density in representative rearrangement events.** Examples of selected fusion and fission events representing both recent and ancient rearrangements. Each facet shows proportional chromosome length against repeat plus satellite density for one focal rearranged species and its closest same-genus or same-subfamily comparator. Grey circles indicate chromosomes from the focal rearranged species, and grey triangles indicate chromosomes from the comparator species. **A.** Fusion examples. Black points mark the observed fused chromosomes. Coloured triangles mark the closest matched comparator chromosomes corresponding to the larger and smaller inferred fusion components, with lines connecting each matched component to the observed fused chromosome. Ancient fusions are shown in the top row and recent fusions in the bottom row. **B.** Fission examples. Dark blue points mark individual fission-derived chromosome fragments in the focal species. Light blue triangles mark the closest matched comparator chromosome, and lines connect the fission-derived fragments to that reference chromosome. Recent fissions are shown in the top row and ancient fissions in the bottom row.

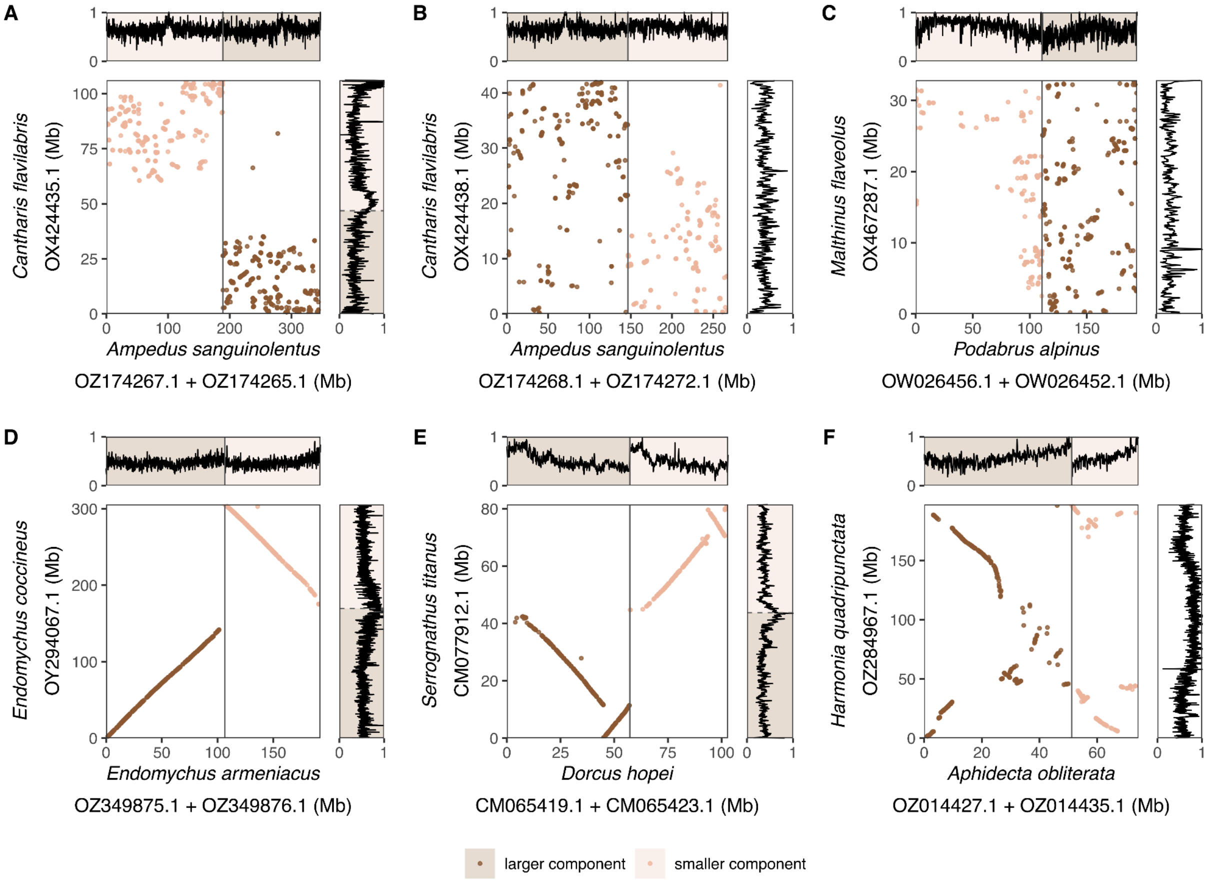


**Fig. S41.**

**Consequences of chromosome fusion in beetles.** Oxford dot plots and coverage tracks showing selected chromosome fusion examples across beetle lineages. In each panel, the focal fused chromosome is plotted on the y-axis, and the corresponding unfused chromosomes from the closest matched comparison species are concatenated on the x-axis. Points represent shared syntenic BUSCO markers, coloured by fusion components: larger ancestral component in brown and smaller ancestral component in pale orange. Solid vertical lines mark boundaries between concatenated comparison chromosomes. The top and right marginal tracks show repeat + satellite density along the comparison and focal chromosomes, respectively. Panels A–C show stepwise sequential fusions involving 6 chromosomes across three different timeframes in Cantharidae. From the oldest in A to the youngest in C. Panels D–F show three additional cases of young (D), moderately old (E) and older fusion event (F).

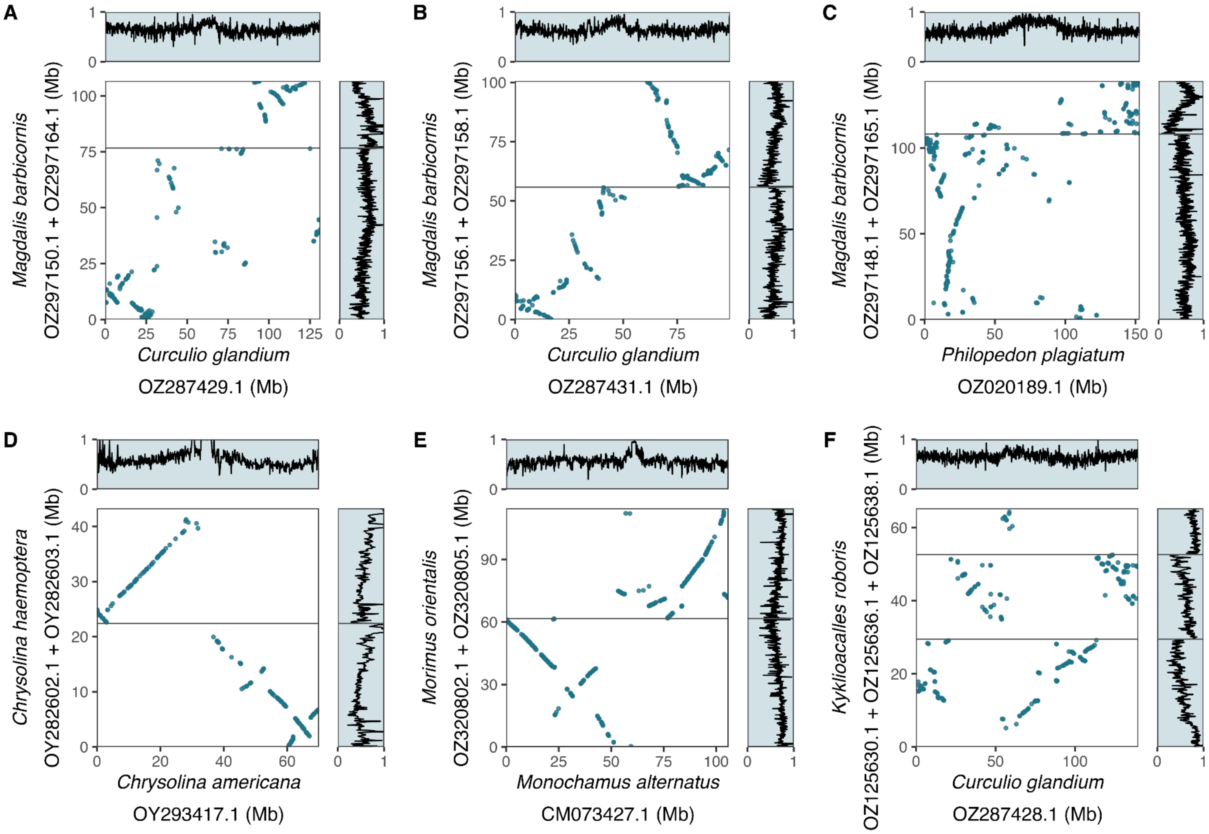


**Fig. S42.**

**Consequences of chromosome fission in beetles.** Oxford dot plots and coverage tracks showing selected chromosome fission examples across beetle lineages**.** In each panel, the closest matched unfissioned chromosome is plotted on the x-axis, and the focal fission products are concatenated on the y-axis. Points represent shared syntenic BUSCO markers between the focal fission products and the matched reference chromosome. Solid horizontal lines mark boundaries between concatenated fission products. The top and right marginal tracks show repeat + satellite density along the reference chromosome and focal fission products, respectively. Panels A–C show stepwise sequential fissions involving 6 chromosomes across three different timeframes in Curculionidae. From the youngest in A to the oldest in C. Panels D–F show three additional cases of young (D), moderately old (E) and older fusion event (F).

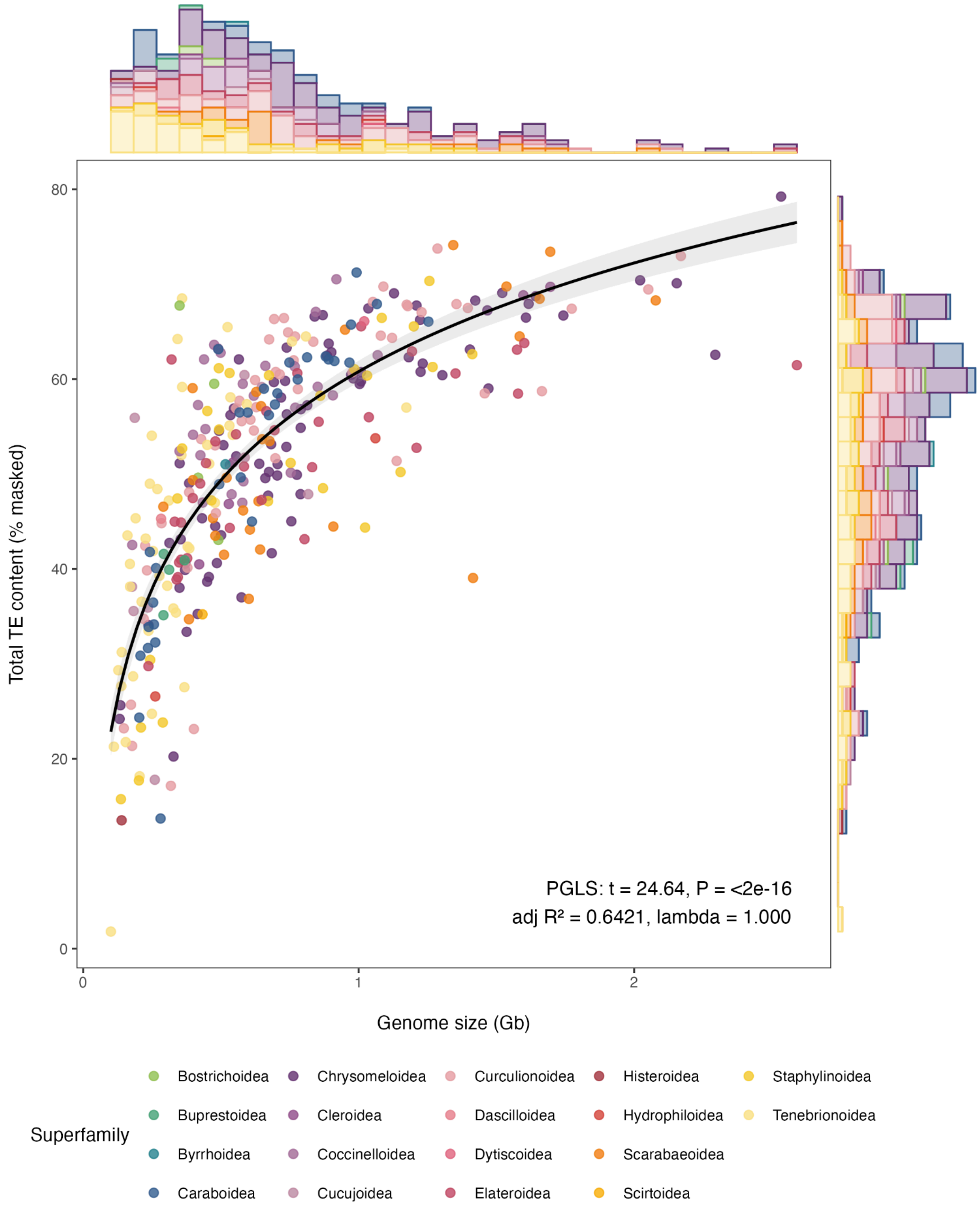


**Fig. S43.**

**Relationship between genome size and transposable element content across beetle genomes.** Scatter plot showing the relationship between genome size and total transposable element (TE) content across sampled beetle genomes. Each point represents one species and is coloured by superfamily. Marginal histograms show the distributions of genome size and TE content. The black line shows the fitted phylogenetic generalised least squares (PGLS) regression, with the grey shaded area indicating the confidence interval. PGLS analysis indicates a strong positive association between genome size and TE content across beetles.

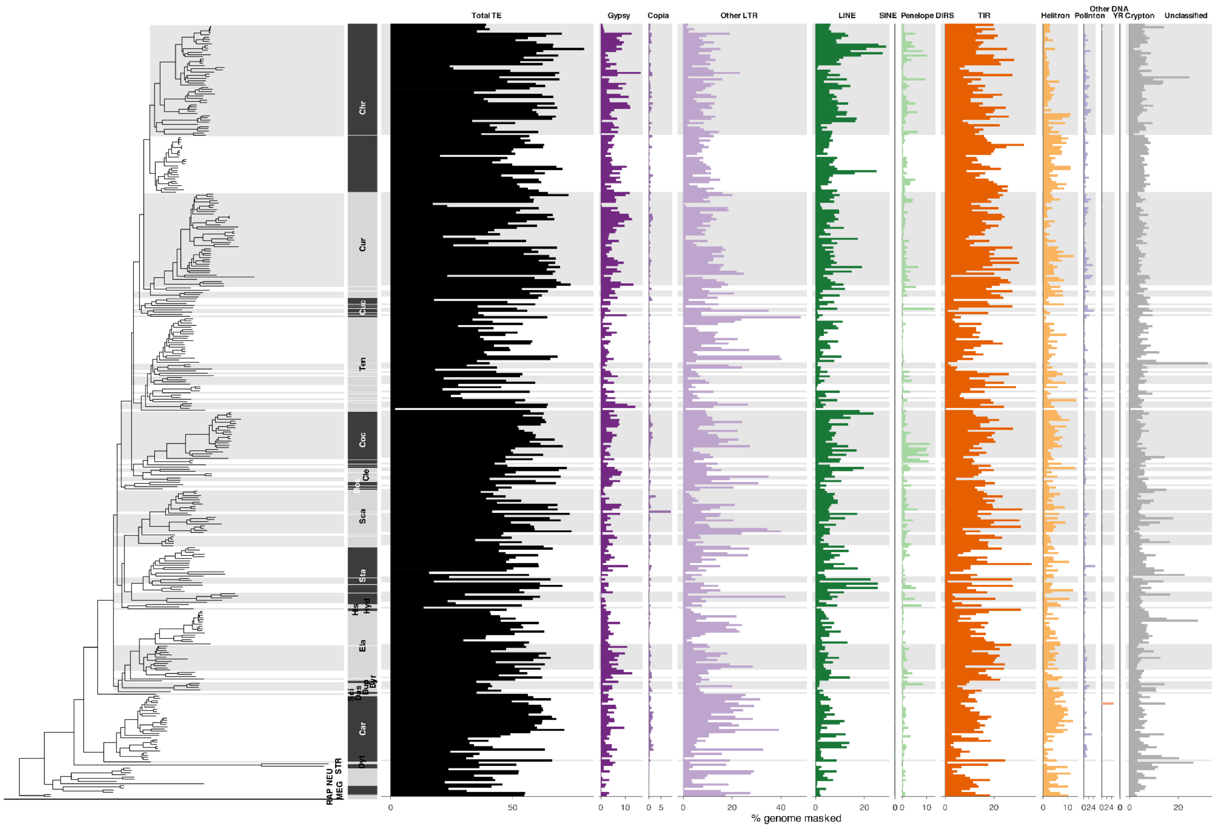


**Fig. S44.**

**Transposable element content across beetle genomes.** The left panel shows the beetle phylogeny, with alternating black and grey blocks denoting superfamilies. Horizontal grey shading across the phylogeny and TE panels indicates family-level groupings. The right panels show the percentage of each genome masked by all TEs combined (Total TE) and by major TE classes, including LTR retrotransposons, non-LTR retrotransposons, DNA transposons, and unclassified repeats. Each horizontal bar represents one species and bar length indicates the percentage of the genome masked by each TE class.

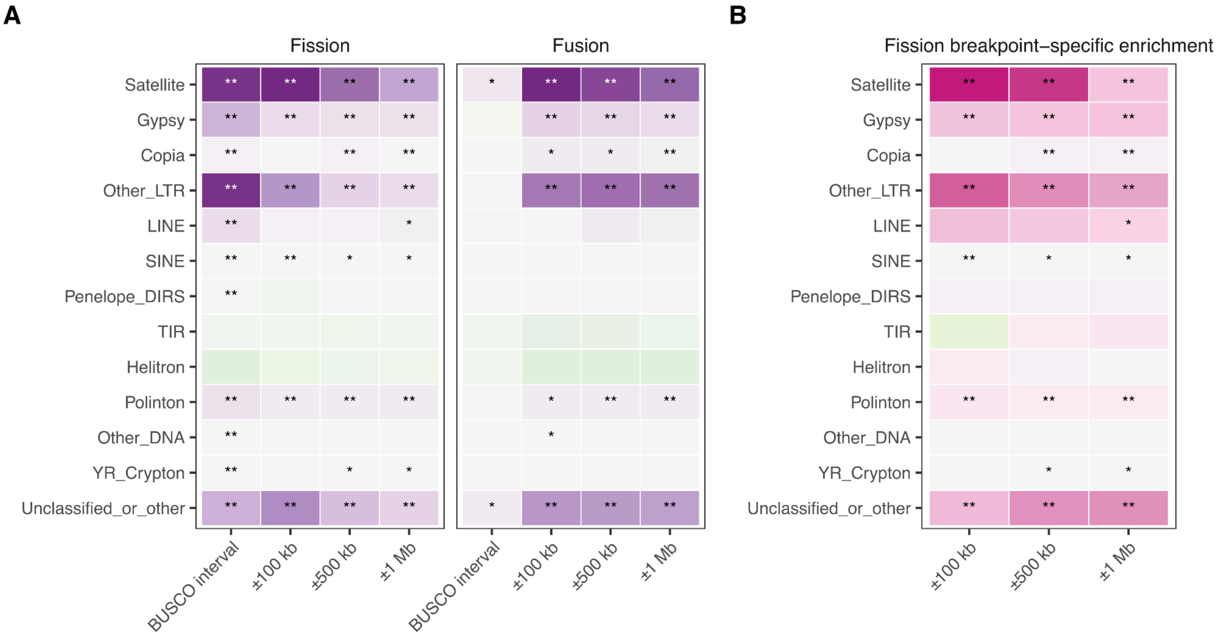


**Fig. S45.**

**Repeat-class enrichment around inferred rearrangement regions.** Permutation tests comparing observed repeat density in rearrangement-associated intervals to same-size random intervals sampled from the same species and chromosome. **A.** Repeat-class enrichment for fission and fusion regions. For fission, intervals include the BUSCO-marker-defined terminal interval and fixed windows around predicted breakpoint-bearing fission-product ends. For fusion, intervals include the BUSCO-marker-defined interval and fixed windows around inferred fusion boundaries. Colour shows enrichment calculated as observed minus null repeat density. **B.** Fission breakpoint-specific enrichment, calculated as enrichment at predicted fission breakpoint ends minus enrichment at both fission-product ends. Positive values indicate repeat classes more enriched at predicted breakpoint ends than expected from general chromosome-end enrichment. Asterisks indicate Benjamini-Hochberg-corrected permutation significance within each interval definition: * adjusted $p < 0.05$, ** adjusted $p < 0.01$.

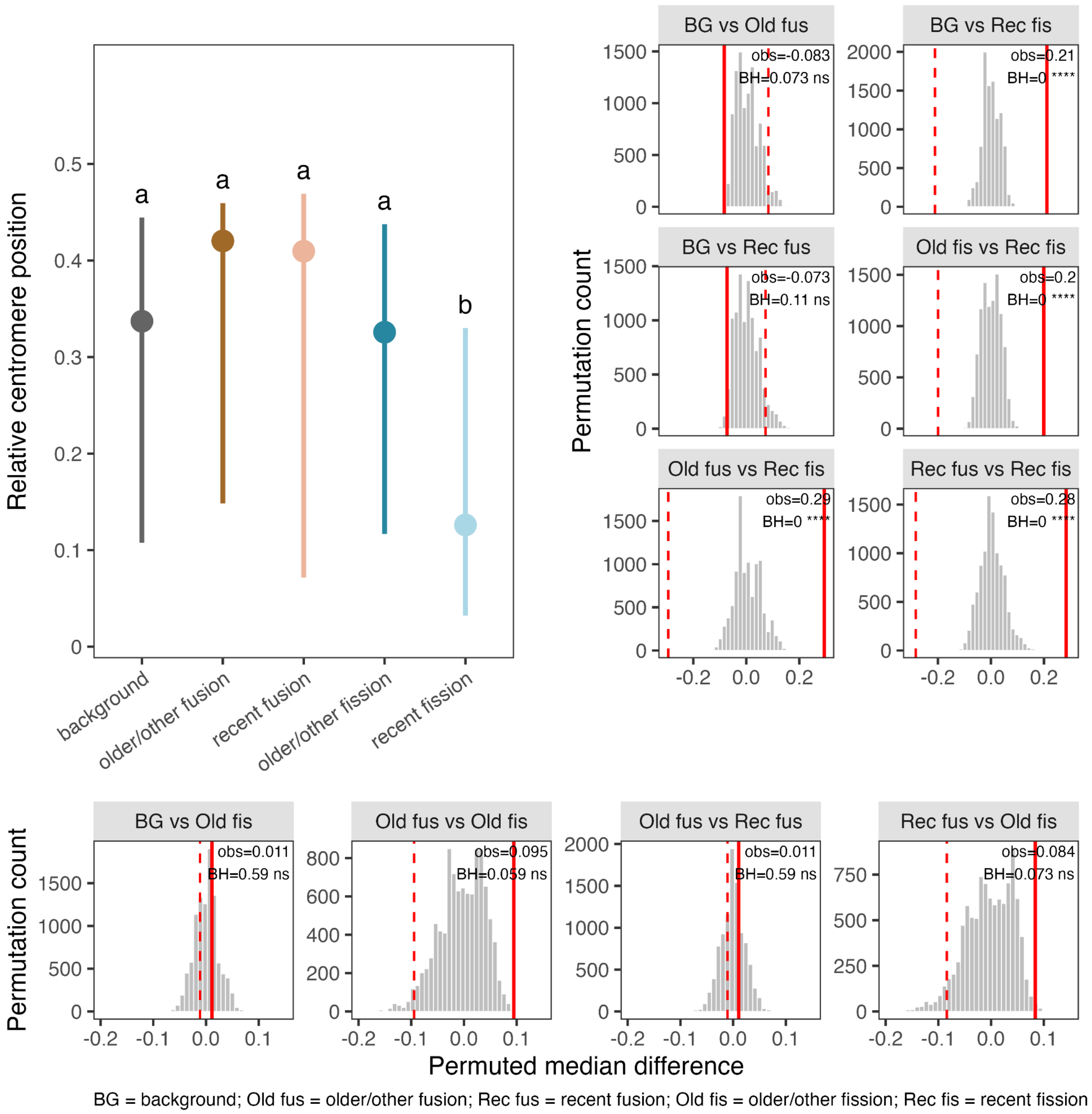


**Fig. S46.**

**Relative centromere position across rearrangement categories.** Top left, median relative centromere position for each rearrangement category. Vertical lines show the interquartile range. Different letters indicate significant differences among groups from permutation tests of median differences after Benjamini-Hochberg correction. Top right and bottom panels show permutation null distributions for pairwise median differences. Red solid lines indicate the observed median difference, and red dashed lines indicate the opposite tail for two-sided tests. Recent fission chromosomes differed significantly from all other categories, whereas the remaining categories did not differ significantly from one another. BG, background; Old fus, older/other fusion; Rec fus, recent fusion; Old fis, older/other fission; Rec fis, recent fission.

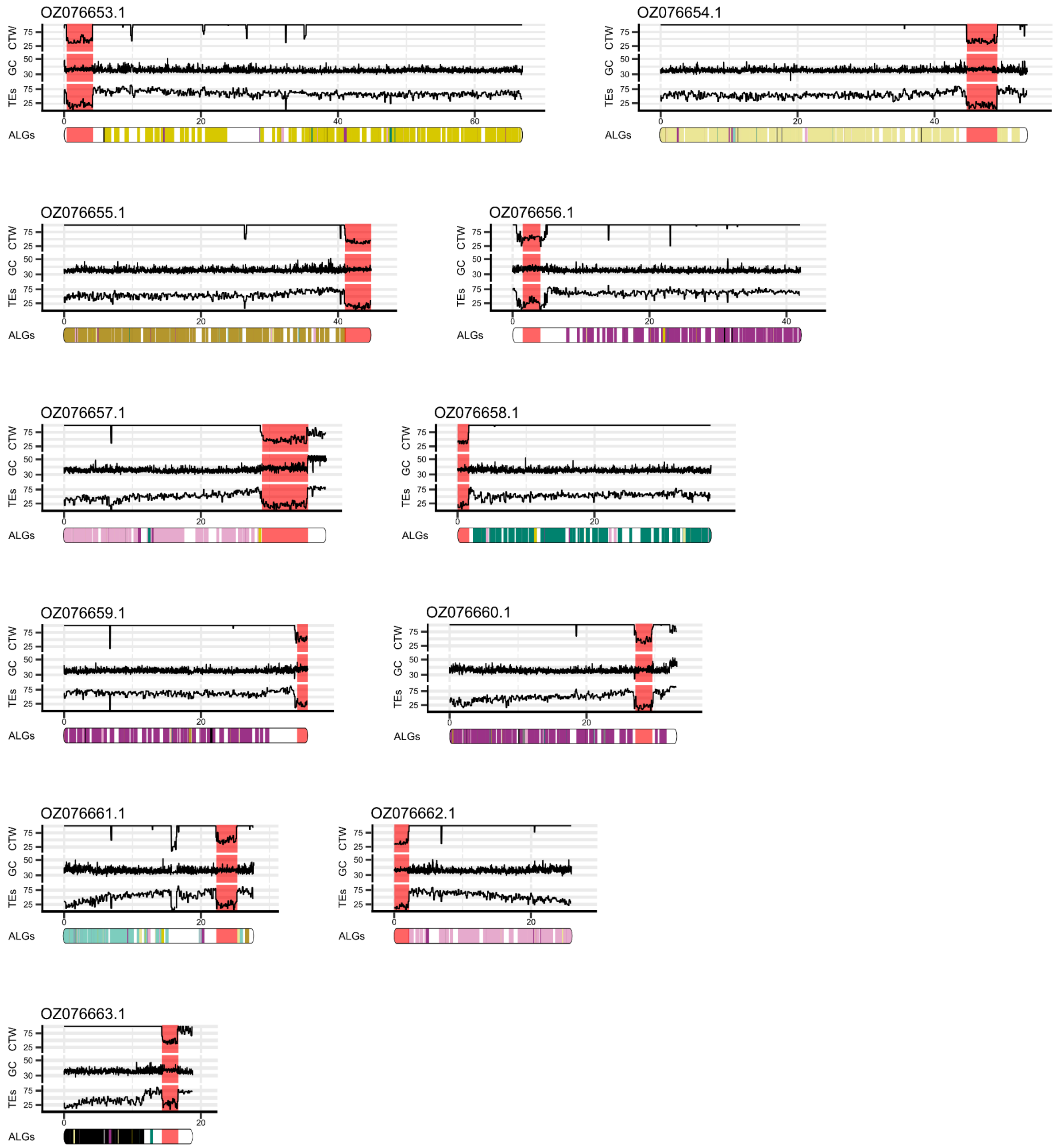


**Fig. S47.**

**Example of putative centric fission in *Anisosticta novemdecimpunctata*.** Chromosomes are painted according to Coleoptera ALG identity. Multiple chromosomes with the same predominant ALG identity and terminal or near-terminal centromeres are consistent with possible centric fission. Inferred centromere positions are highlighted in red, and vertical bars within each chromosome show chromosome position of BUSCO genes coloured according to their ALG assignment.

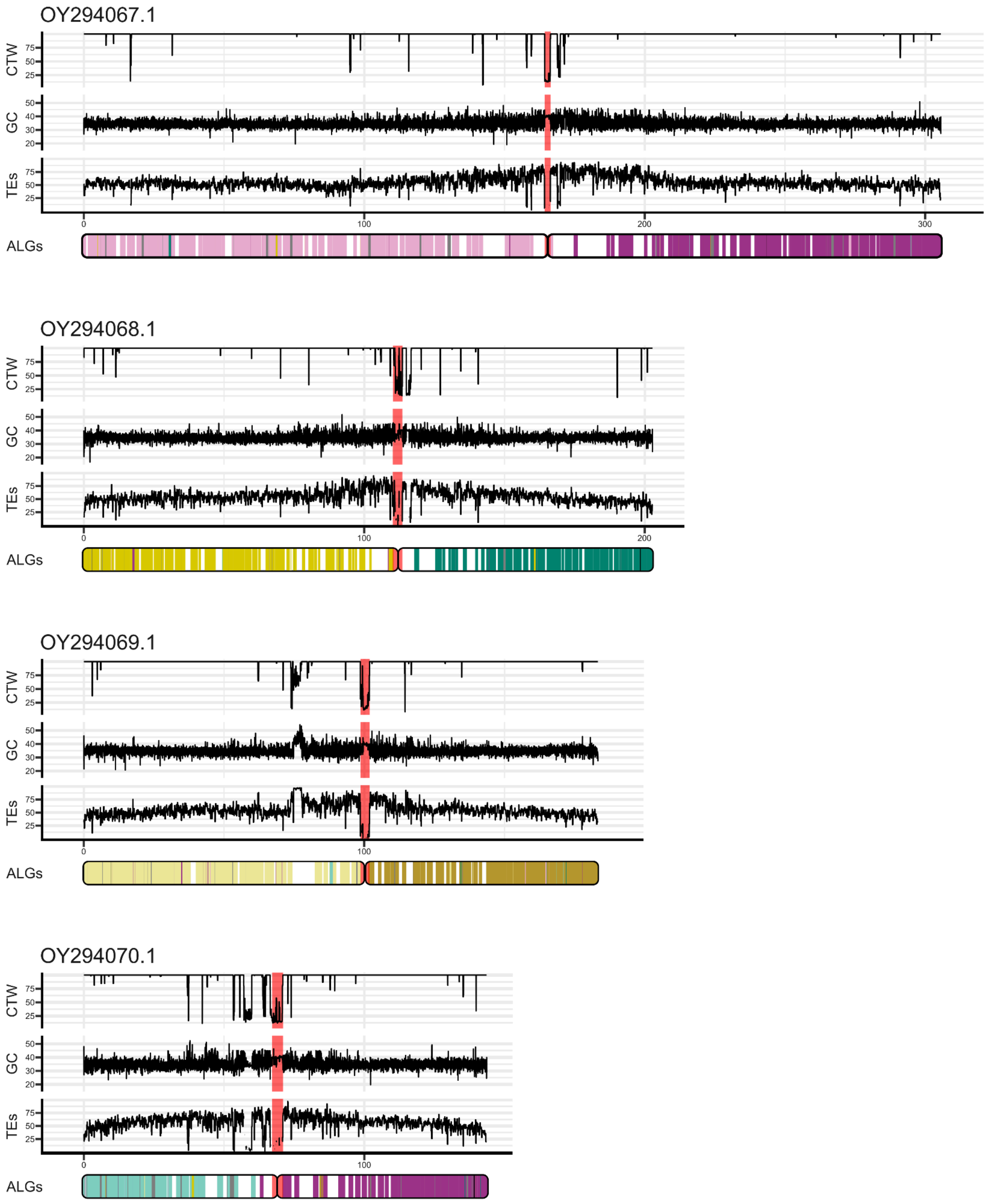


**Fig. S48.**

**Example of putative centric fusion in *Endomychus coccineus*.** Chromosomes are painted according to the inferred chromosomal composition of the most recent inferable ancestor of this species. Several chromosomes contain more than one predominant ancestral linkage identity, separated around the centromere, a pattern consistent with possible centric fusion. Inferred centromere positions are highlighted in red, and vertical bars within each chromosome show chromosome position of BUSCO genes coloured according to their ALG assignment.

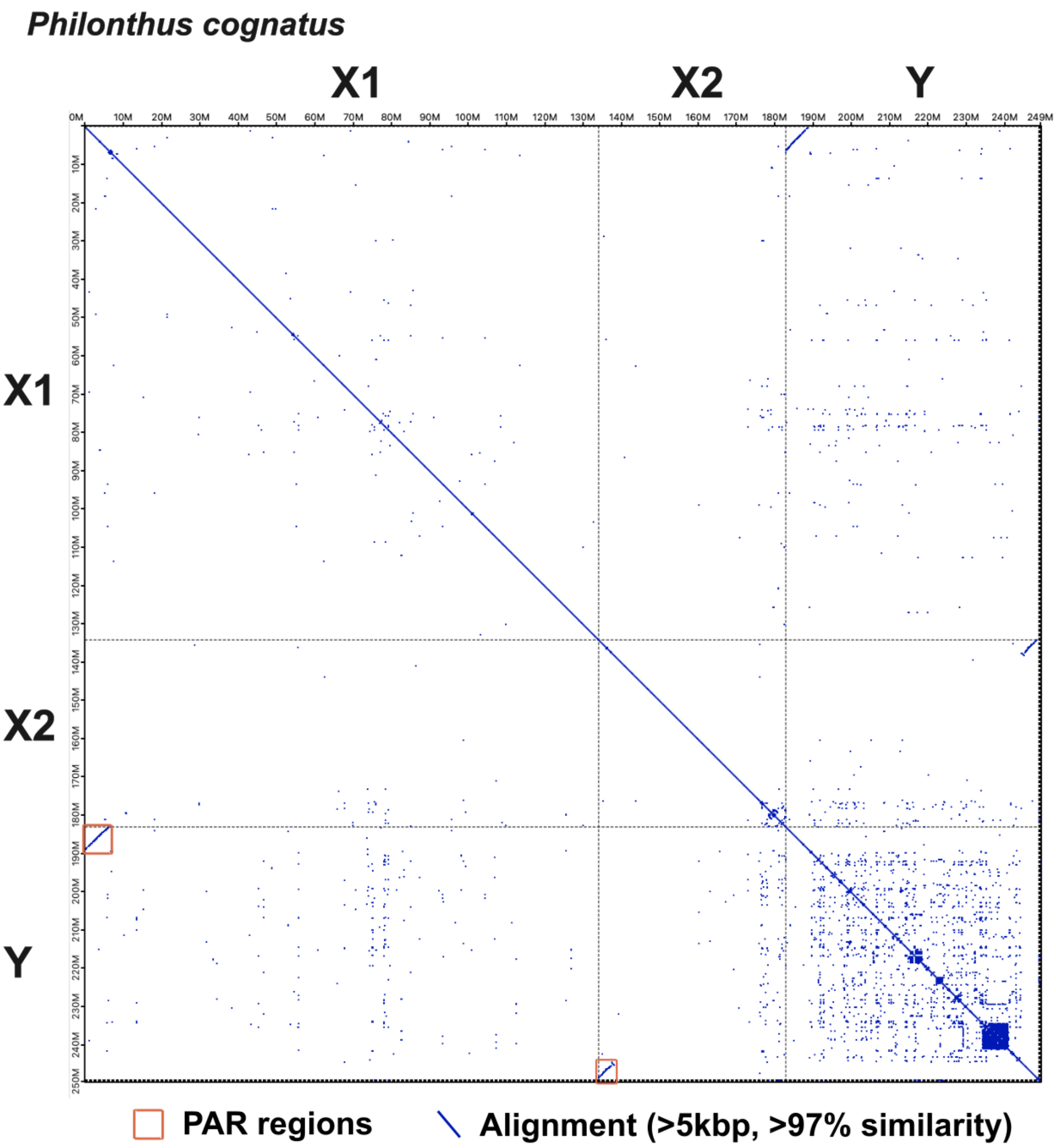


**Fig. S49**

**Homologous regions visualised of sex chromosomes shown via self-alignment of *Philonthus cognatus* using corrected assembly.** The whole genome alignment was performed using FastGA (https://github.com/thegenemyers/FASTGA) and visualised using ALNview (https://github.com/thegenemyers/ALNVIEW) showing only alignments >5kbp with >97% similarity. Only the three sex chromosomes are shown. For purposes of this supplementary materials, the colours of the ALNview were inverted (software uses dark mode). The alignments of the PAR regions are highlighted in orange squares and have approximately 6.11 Mbp (X1/Y) and 4.37 Mbp (X2/Y) in size respectively.

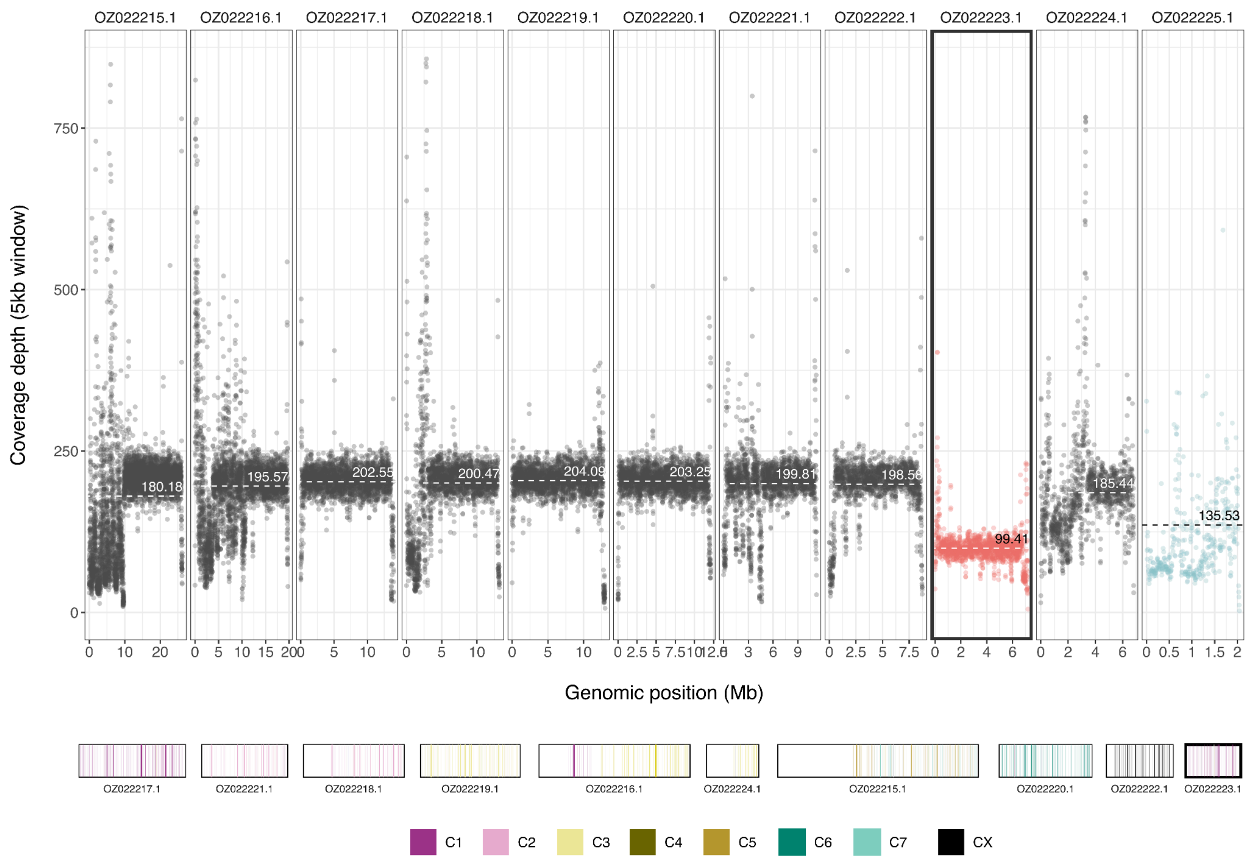


**Fig. S50**

**Putative rare example of sex chromosome turnover in beetles.** Coverage of *Aleochara curtula* male reads. Red points and the highlighted box indicate a rare, previously unreported case of sex chromosome turnover in beetles, while blue points indicate the Y chromosome. The bottom panel shows ALG painting for each chromosome. The canonical ancestral beetle X chromosome, OZ022222.1, no longer functions as the X chromosome and shows coverage comparable to that of the autosomes. In contrast, OZ022223.1, which is partially composed of C1, has become the new X chromosome and shows the expected reduced coverage in males.

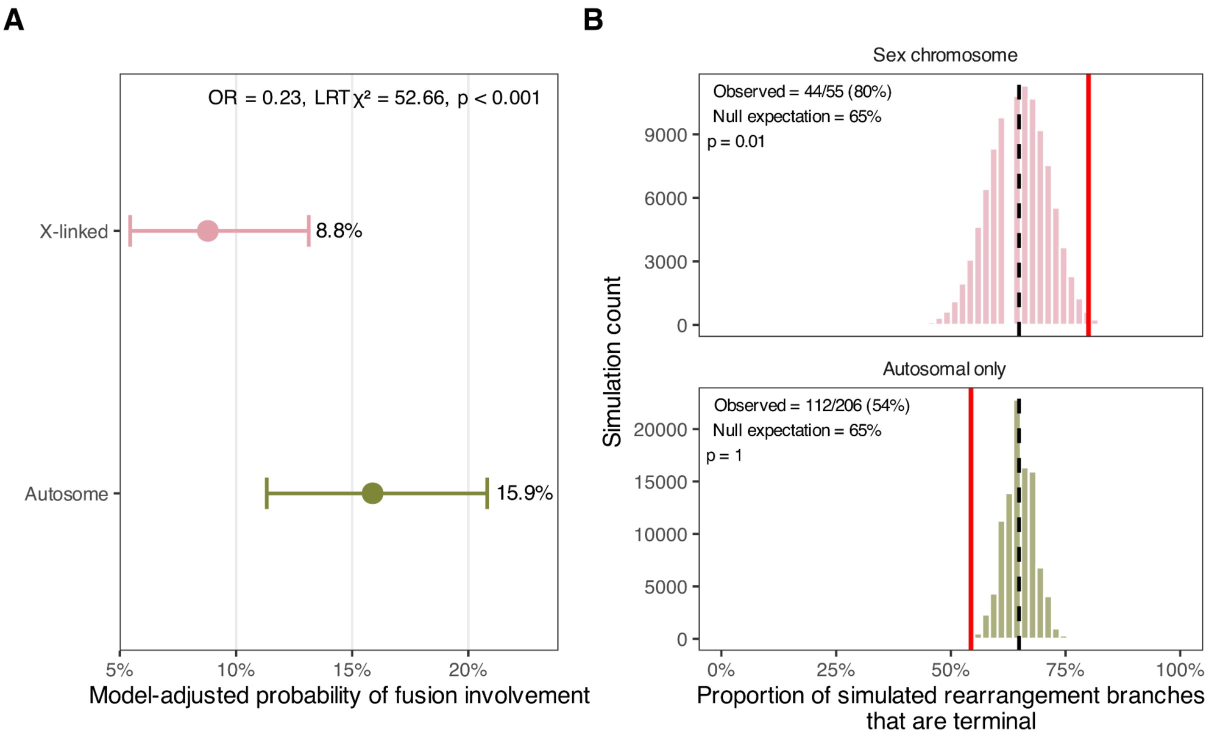


**Fig. S51.**

**X-linked chromosomes show distinct rearrangement involvement and persistence. A.** A. Average model-adjusted probability of fusion involvement for autosomal and X-linked chromosomes. Predictions were generated from a binomial GLMM including log-transformed chromosome size and X-linkage status as fixed effects, with parent and child nodes as random intercepts, and were averaged across the observed chromosome–branch records while retaining fitted random effects. Horizontal intervals show uncertainty from simulation of the fixed-effect coefficients. The annotation reports the conditional odds ratio for X-linkage. X-linked chromosomes had lower odds of fusion involvement than autosomes.

**B.** Branch-length-weighted simulations testing whether branches with sex-chromosome changes or autosomal-only rearrangements were enriched among terminal branches. For each category, the observed number of change branches was randomly assigned 100,000 times, with branch-selection probabilities proportional to branch length. Dashed lines show the null expectation and red lines show the observed proportion of terminal branches. *P*-values indicate the fraction of simulations with at least as many terminal-branch changes as observed. Sex-chromosome changes were enriched on terminal branches, whereas autosomal-only rearrangements were not.

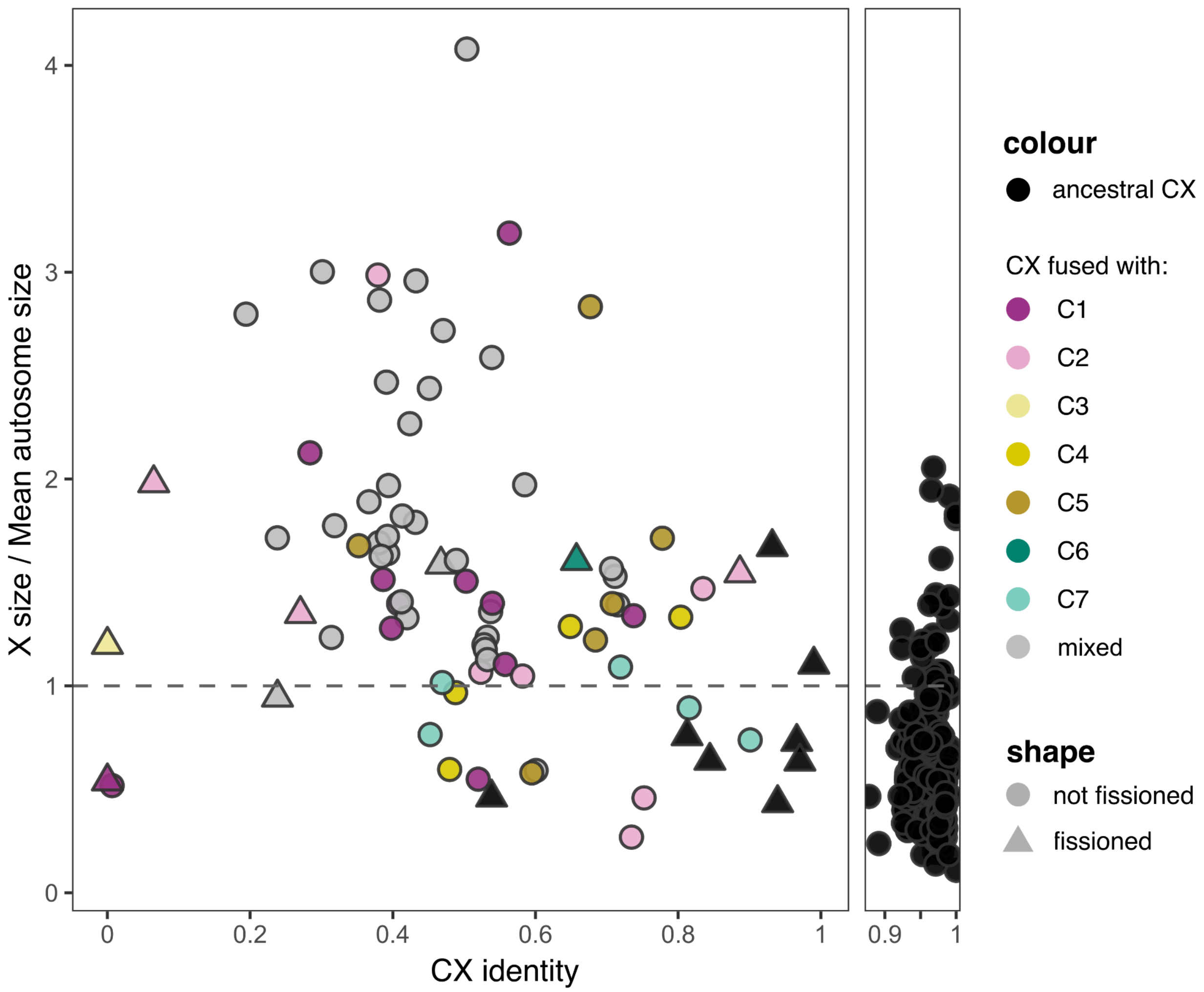


**Fig. S52.**

**Formation of disproportionately large X chromosomes relative to autosomes.** X chromosomes are coloured black when classified as ancestral, defined as ≥80% CX identity and with no more than 15 translocated genes from any other single ALG. X chromosomes are coloured according to the most dominant non-CX ALG when that ALG is the sole additional element and contributes more than 15 genes. X chromosomes are coloured grey ("mixed") when multiple non-CX ALGs are present, each contributing more than 15 genes. Cases in which CX is split into multiple X chromosomes due to fission(s) are indicated by triangles.

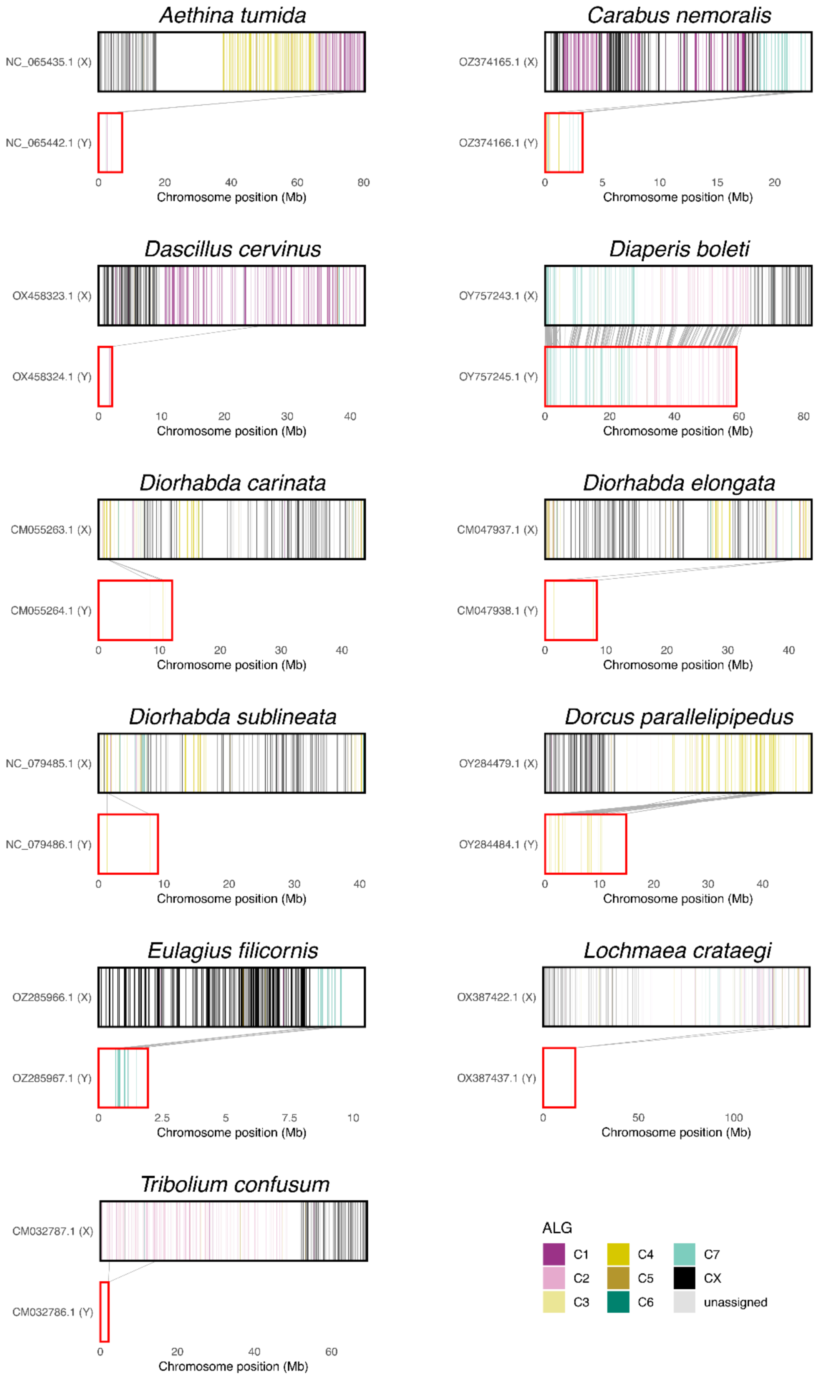


**Fig. S53.**

**Evolution of neo-Y chromosomes in beetles.** Among 76 species with Y chromosomes assembled and confirmed by read coverage in males, only 33 species have at least one gene on the Y. Among those 33, 11 species shown here are putative candidates of the neoY chromosome, which we defined as Y chromosomes that share genes with a neo-X. For each species, the X chromosome is shown above the corresponding Y chromosome. BUSCO genes are coloured according to their coleopteran ALG assignment. Grey lines connect BUSCO orthologues detected on both the X and Y chromosomes, illustrating retained gene correspondence between the X and Y. Chromosome lengths are plotted to scale within each species, with genomic position shown in megabases. X chromosomes are outlined in black and Y chromosomes in red.

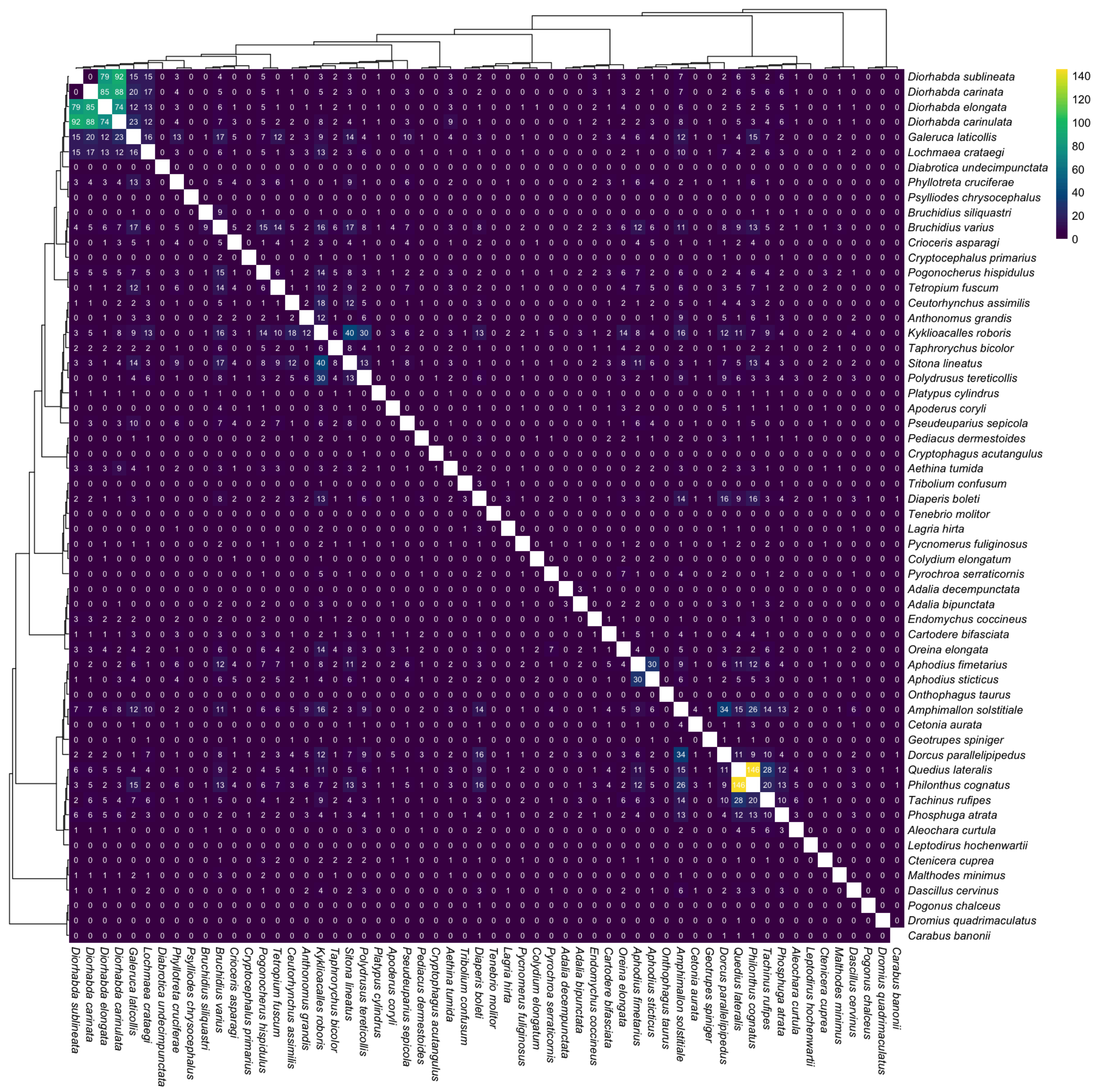


**Fig. S54.**

**Numbers of homologous genes shared between Y chromosomes.** Heatmap cells show the number of orthogroups shared between each pair of Y chromosomes. The number of shared genes between Y chromosomes shows non-negligible gene overlap only in cases of shared formation of neo-Y chromosomes.

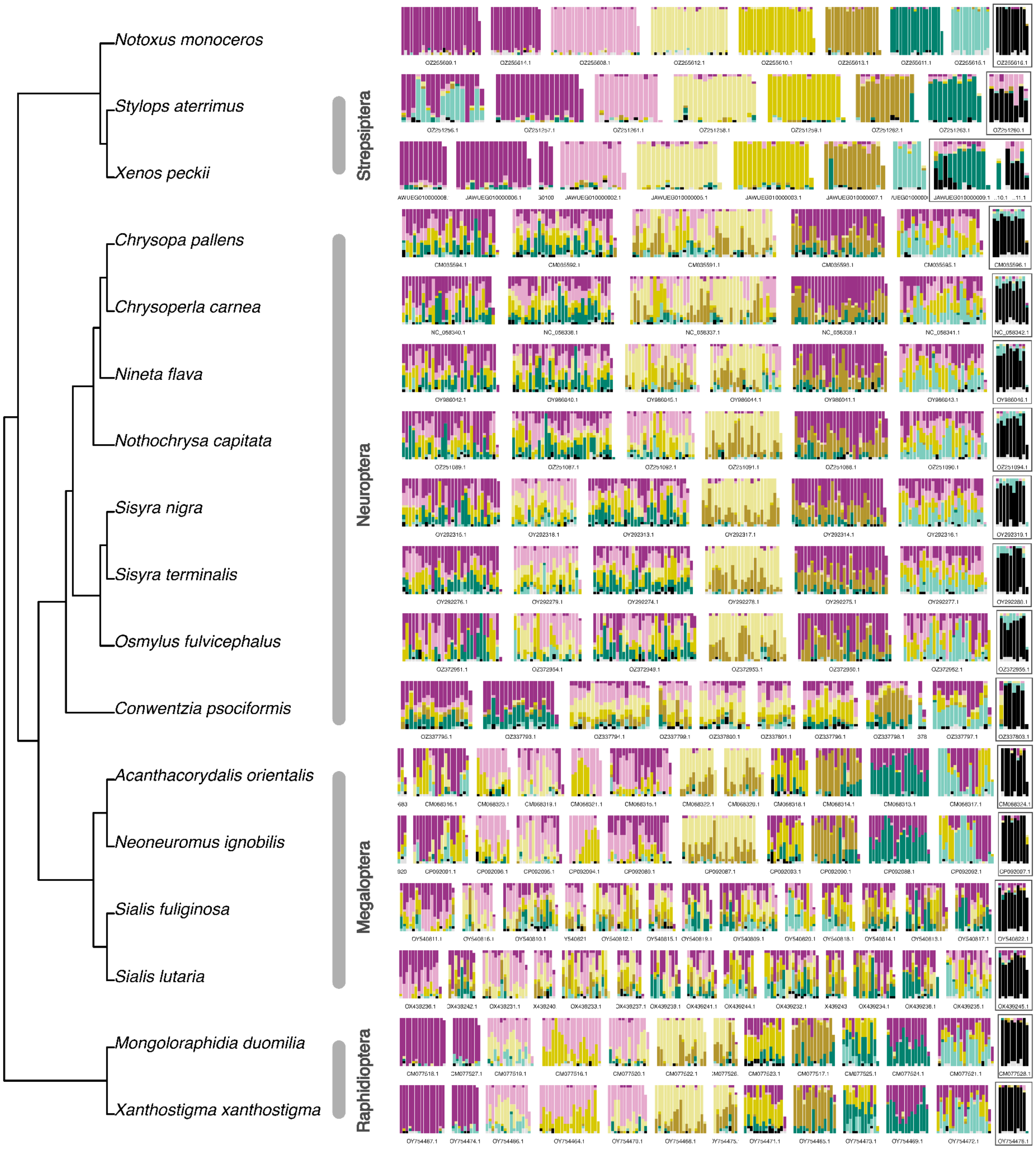


**Fig. S55.**

**Conservation of the X chromosome across Neuropteroidea.** Synteny plots across Neuropteroidea, including Coleoptera (represented by *Notoxus monoceros*), Strepsiptera, Neuroptera, Megaloptera and Raphidioptera. Chromosomes are arranged horizontally within each species and separated by white gaps. Each chromosome is divided into consecutive 20-gene windows, with colours indicating the proportional contribution of BUSCO genes assigned to each coleopteran ALG. The chromosome outlined in black denotes the inferred X chromosome. For *Xenos peckii*, *Chrysopa pallens*, *Nothochrysa capitata*, and *Xanthostigma xanthostigma*, the

X chromosome was identified from male assemblies. The X chromosome of *X. peckii* is represented by three scaffolds because a chromosome-level assembly is not available for this species. Black segments represent genes assigned to the ancestral coleopteran X chromosome, CX. Their strong enrichment on the inferred X chromosomes indicates that X-chromosome identity is broadly conserved across Neuropteroidea, beyond Coleoptera.

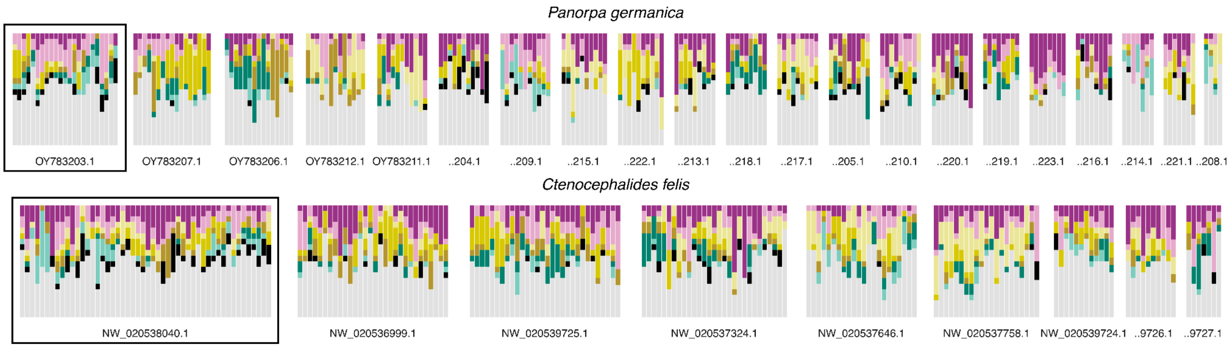


**Fig. S56.**

**Crosspainting beetle ALGs to Mecoptera and Siphonaptera.** Chromosomes of *Panorpa germanica* (Mecoptera) and *Ctenocephalides felis* (Siphonaptera) are arranged horizontally within each species and separated by white gaps. Each chromosome is divided into consecutive 20-gene windows, with colours indicating the proportional contribution of BUSCO genes assigned to each coleopteran ALG. Diptera odb12 BUSCO genes identified in the two assemblies were matched to orthologous Coleoptera odb12 BUSCO genes using correspondences inferred with OrthoFinder. Diptera BUSCOs with an identified Coleoptera orthologue were then coloured according to the coleopteran ALG assignment of that orthologue. Chromosomes outlined in black denote the inferred X chromosomes.

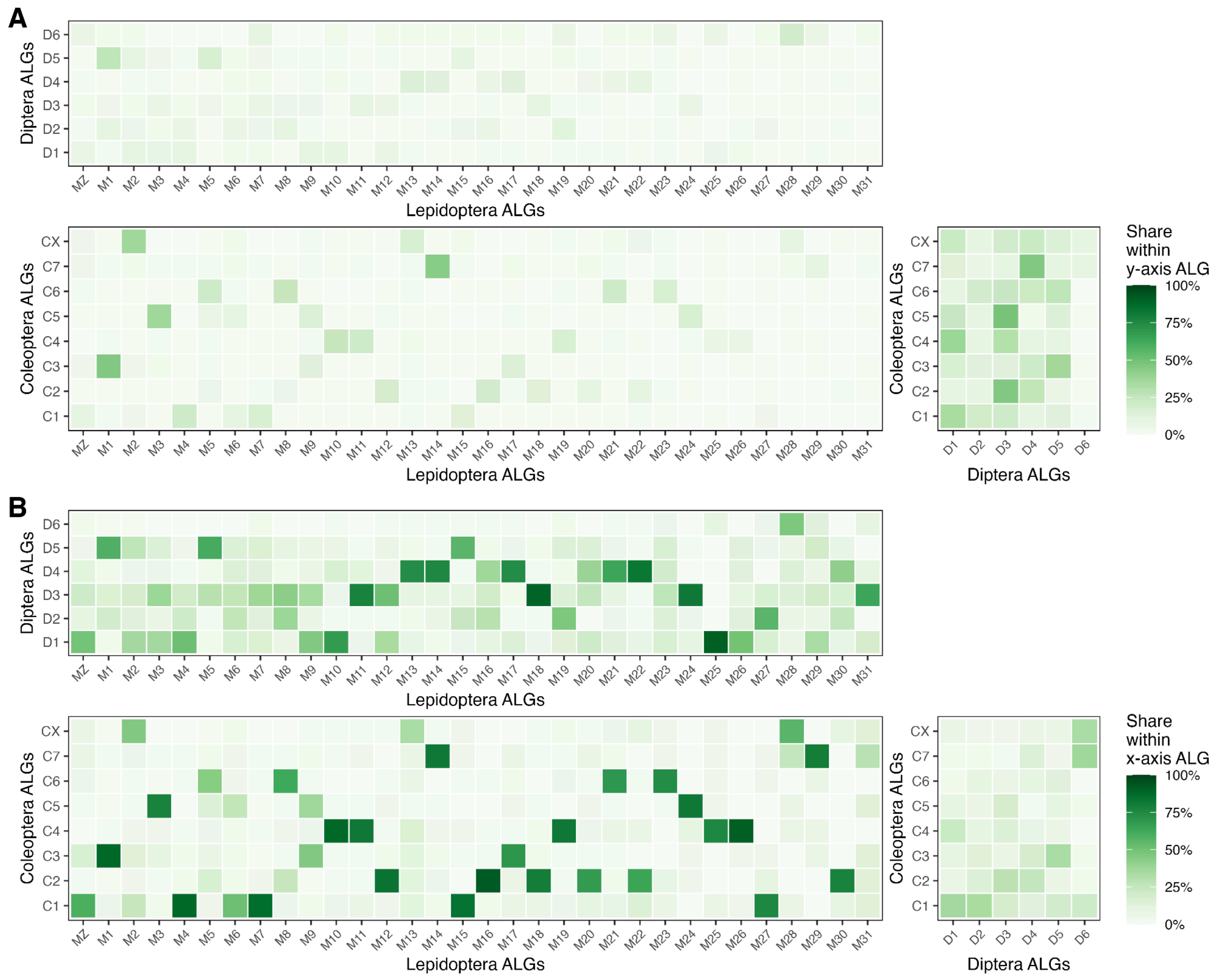


**Fig. S57.**

**Correspondence among ancestral linkage groups across holometabolan insects.**
Heatmaps show pairwise correspondence among inferred ancestral linkage groups (ALGs) of Lepidoptera, Diptera, and Coleoptera based on homologous BUSCO genes. **A**. For each ALG on the y-axis, colours indicate the proportion of its BUSCO genes assigned to each ALG on the x-axis. **B**. For each ALG on the x-axis, colours indicate the proportion of its BUSCO genes assigned to each ALG on the y-axis. Darker colours indicate higher proportions. Thus, panel A shows the composition of y-axis ALGs, whereas panel B shows the distribution of x-axis ALGs across y-axis ALGs.

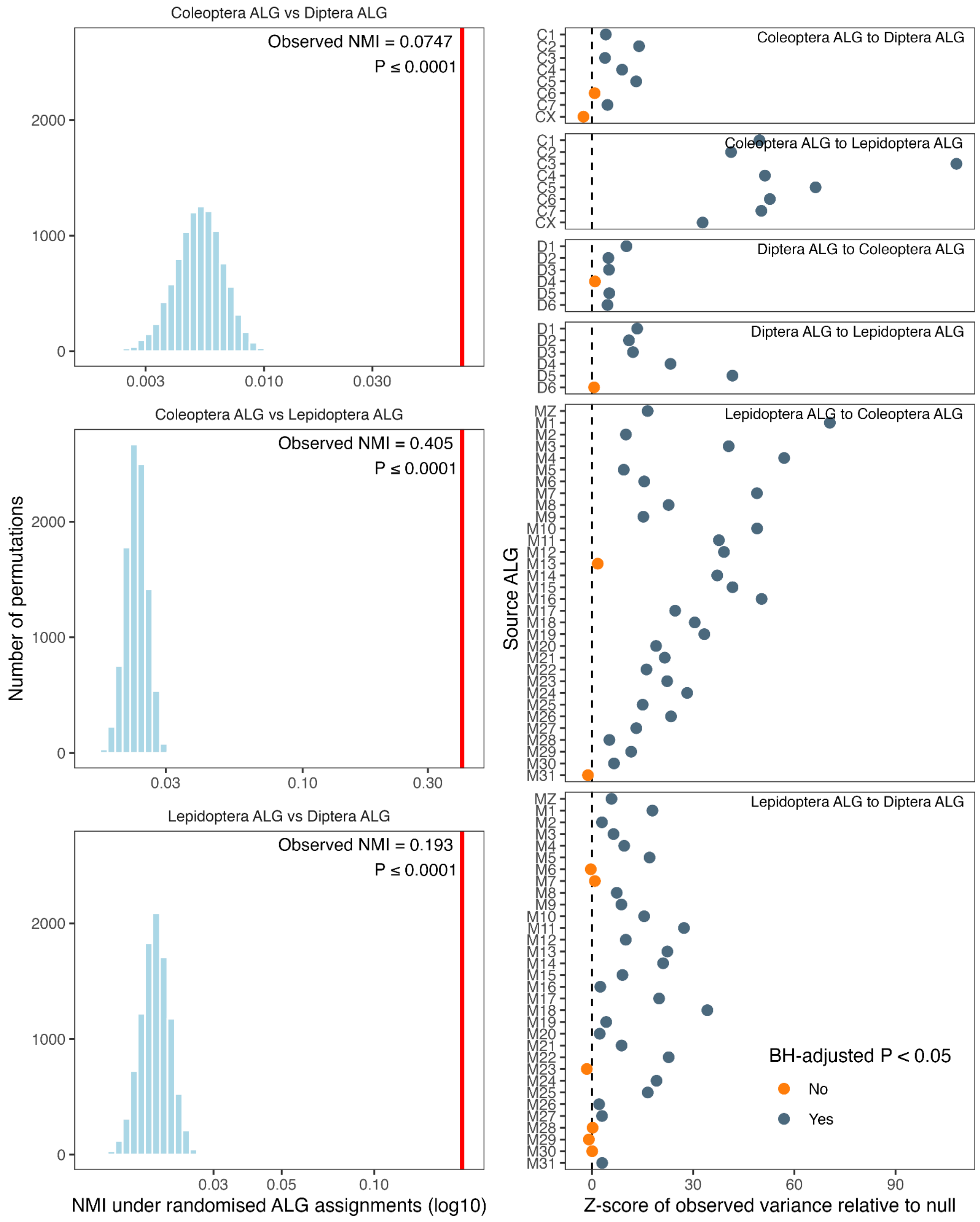


**Fig. S58.**

**Permutation tests of ALG correspondence among insect orders. A.** Null distributions show normalised mutual information (NMI) values obtained after randomly shuffling ALG labels among homologous BUSCO genes while preserving ALG size distributions. Vertical lines indicate the observed NMI for each pairwise comparison. In all comparisons, the observed ALG correspondence was far greater than expected under random shuffling, with no randomised replicate reaching the observed value across 9,999 permutations. **B.** Per-ALG clustering tests showing Z-scores of the observed variance in the distribution of each source ALG's BUSCO genes across target ALGs relative to null expectations. Black points indicate source ALGs significant after Benjamini–Hochberg correction; grey points indicate non-significant ALGs.

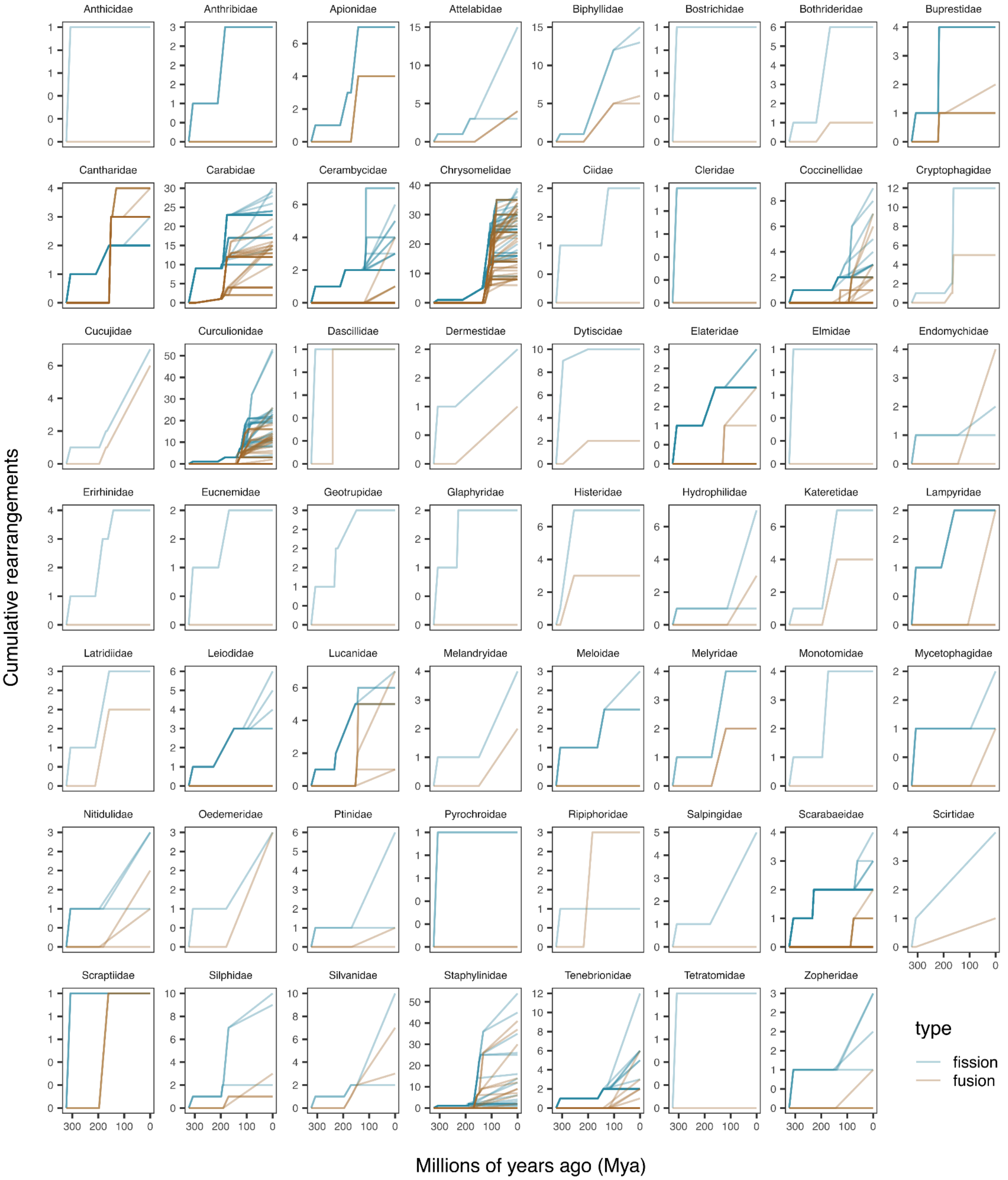


**Fig. S59.**

**Family-level temporal dynamics of chromosomal rearrangement accumulation.**
Facet plots showing cumulative inferred fission and fusion events through time for each beetle family. Each line represents a lineage-specific rearrangement trajectory within a family, with blue lines indicating fissions and orange lines indicating fusions. The x-axis shows time before present in millions of years, and the y-axis shows the cumulative number of rearrangements.

#### Supplementary Tables

**Table S1. Genome assembly metadata, assembly quality metrics, and BUSCO completeness statistics for the 354 genomes included in this study.**

For each genome analysed, the table provides the assembly identifiers and sources, taxonomic classification, genome size and assembly metrics, BUSCO summary statistics, and citations for previously published genomes.

**Table S2. Coleoptera ancestral linkage group definitions.**

This table lists BUSCO gene assignments to the ancestral linkage groups of beetles, here termed Coleoptera ALGs, C1–C7 and CX, based on Syngraph reconstruction (-m10). Approximately 97% of BUSCO genes, 3,603 of 3,729, were assigned to these eight ALGs. The ALGs are ordered by size, with CX being the smallest. CX corresponds to the inferred ancestral sex chromosome of beetles.

**Table S3. Ancestral linkage group inference for all phylogenetic nodes.**

For each internal node in the phylogeny, the table provides the inferred ALGs and the BUSCO genes belong to it based on the output of the Syngraph tabulate function (-m10). For all descendant nodes of Chrysomeloidea, the inference was from a separate run using higher parameters (-m24). The results of the two tables were merged.

**Table S4. Chromosomal rearrangement events inference along the phylogenetic tree.**

For each branch along the phylogeny, the table reports the number of rearrangement events (fusion and fission) and the resulting change in chromosome number between an ancestor and its descendant(s), inferred from Syngraph.

**Table S5. Correspondence between Coleoptera ALGs and Stevens elements.**

This table compares the Coleoptera ALGs inferred in this study with previously proposed Stevens elements. BUSCO genes located on *Tribolium castaneum* Tcas5.2 chromosomes were assigned to the corresponding Stevens elements and compared with their assignments to Coleoptera ALGs.

**Table S6. Ancestral gene order reconstructions for all phylogenetic nodes.**

For each internal node in the phylogeny, the table provides the inferred contiguous ancestral regions (CARs) and the gene order reconstructions of BUSCO genes belong to it based on the output of the AGORA.

**Table S7. Intrachromosomal rearrangement estimates of extant genomes relative to the ancestral gene order reconstruction of beetle ALGs.**

For each extant genome and chromosome, the fragmentation index (FI) and domain fragmentation index (DFI) are provided as estimates of intrachromosomal (gene-order) rearrangements and for local gene-order reshuffling within fused domains in fused chromosomes, respectively.

**Table S8. Sex chromosome assignments for species included in this study.**

For each genome, the table lists the sex chromosome assignments identified in this study based on male sequencing-read coverage, the number and size of the X chromosome(s), the number and proportion of CX genes relative to genes of other ancestries, and the type of rearrangement, if any, that occurred on these chromosomes.

**Table S9. Sex chromosome changes identified in this study.**

For each genome, the table lists the sex chromosome changes identified in this study and briefly describes the evidence used for their assignment. "Genomes" refers to cases in which known sex chromosomes were identified based on male sequencing coverage mapped to the genome assemblies. "Implied" refers to genomes without known sex chromosomes that nevertheless contain the ancestral sex chromosome composition (CX). "Species-specific" refers to cases in which a sex chromosome transition occurs in only one species, while genomes from other members of the same genus are available. "Singleton" refers to cases in which a transition occurs in only one species and no close relatives are available, making it impossible to assess the age of the transition.

**Table S10. Genomic correlates of chromosome length across non-rearranged beetle chromosomes.**

For each species, the table lists the strength of correlation (Spearman's rank) between GC content/repeat density/coding density and proportional chromosome length. Only non-rearranged autosomes (i.e., have not undergone fusion or fission) in the 338 beetle genomes were included in these statistics. Spearman's Rank correlation coefficients (ρ) and *p*-values obtained by two-sided Spearman's correlation test are indicated.

**Table S11. Summary of feature density per beetle ALGs across the dataset.**

The table specifies the average feature density for a set of features (GC content, repeat density, coding density, and the density of each major repeat class) per beetle ALG. Only chromosomes corresponding to intact beetle ALGs (i.e., have not undergone fusion or fission) in the 338 beetle genomes were included in these statistics.

**Table S12. Feature statistics for all 4,091 chromosomes in the dataset.**

For each chromosome, the table includes the corresponding species name, assigned dominant beetle ALGs for chromosomes that have not undergone fusion and fission, GC content, repeat density, coding density, and the density of each major repeat class.

**Table S13. Proportion of each of the 354 genomes annotated by major TE subclasses.**

The table details the proportion of each genome, comprising 338 beetle genomes and 16 outgroup genomes, annotated by major TE subclasses using EDTA. These comprise LTR retrotransposons (Gypsy, Copia, and Other LTR), non-LTR retrotransposons (LINE, SINE, and Penelope DIRS), DNA transposons (TIR, Polinton, and Other DNA), rolling-circle transposons (Helitron), tyrosine recombinase elements (YR Crypton), and unclassified or other repeat categories.

**Table S14. Predicted centromere positions of the 338 beetle genomes in the dataset.**

Predicted centromere coordinates and relative centromere positions are shown for each chromosome. Rearrangement categories were assigned to either recent fission, recent fusion, older/other fission, older/other fusion, and background chromosomes. The ancestry division score (ADS) is reported where applicable for fused chromosomes with two dominant ancestral linkage identities; higher values indicate stronger separation of ancestral identities on opposite sides of the predicted centromere.

**Table S15. Age estimates for major phylogenetic nodes used in temporal analyses of rearrangement dynamics in three largest insect orders: beetles, flies, and butterflies.**

For each insect order included in the comparative temporal analyses, the table lists the estimated age of each major phylogenetic node in million years ago (Mya), the corresponding taxonomic level, the clade represented by that node, and the source used for the age estimate.

**Table S16. Correspondence of coleopteran ALGs with dipteran ALGs and lepidopteran ALGs (Merian elements).**

The table provides the correspondence between BUSCO genes from the odb12 Coleoptera dataset, odb12 Diptera dataset, and odb10 Lepidoptera dataset. For each orthologue, the table lists the corresponding Coleoptera ALG, Diptera ALG, and Merian element assignments, where available.

**Supplementary Data**

Data S1.
Schematic representation of all reconstructed ancestral chromosome configurations across internal nodes and extant genomes since the last common ancestor of beetles, ‘painted’ with Coleoptera ALGs.

Data S2.
Coleoptera ALG paintings for each chromosome of each species in the dataset.
