## Supplementary figures and images for "338 coleopteran genomes reveal exceptional rearrangement variation compared to other insect orders"

### plot.syngraph_windowbp_1Mb_rel_vertical_chrsize_Adalia_bipunctata.png

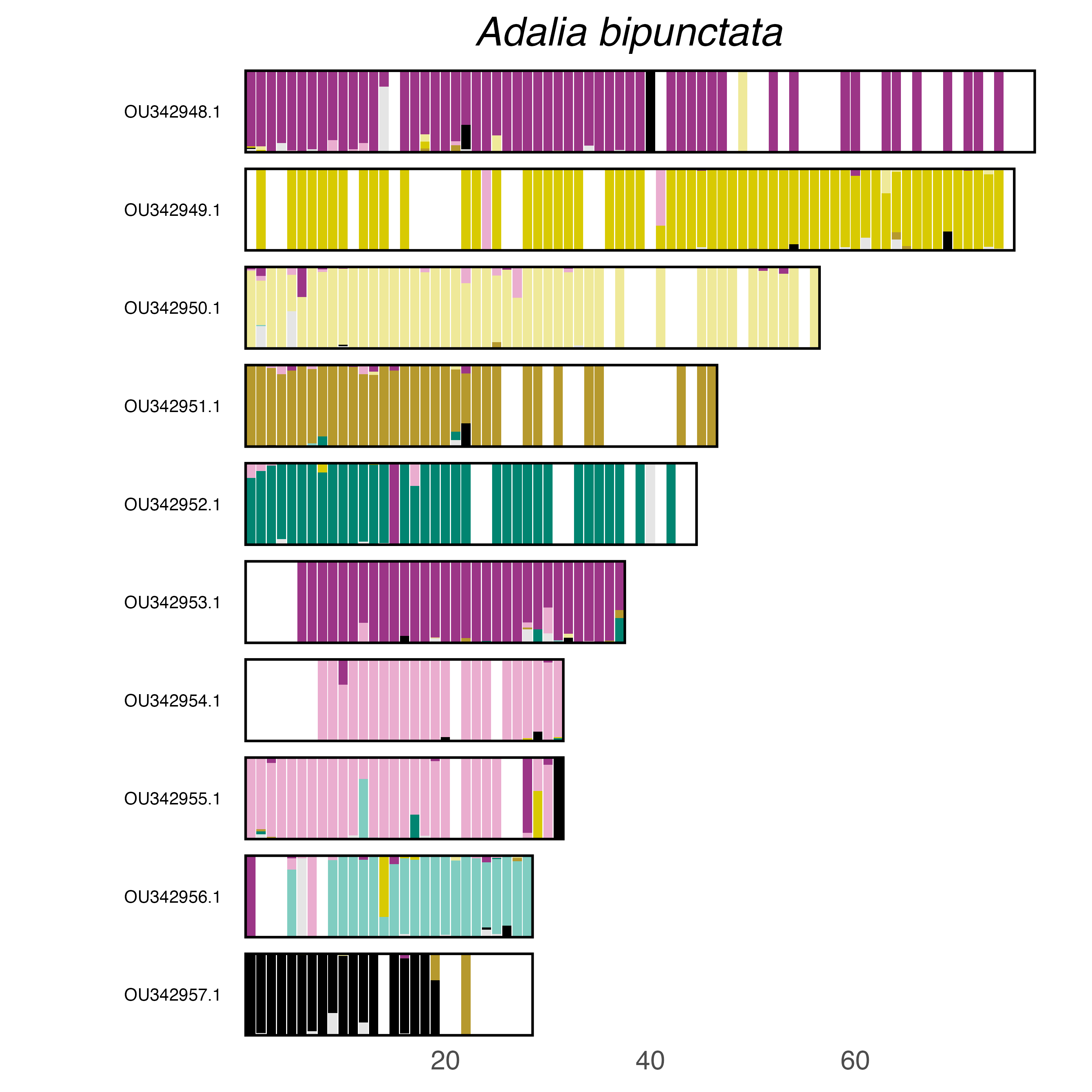

### plot.syngraph_windowbp_1Mb_rel_vertical_chrsize_Adalia_decempunctata.png

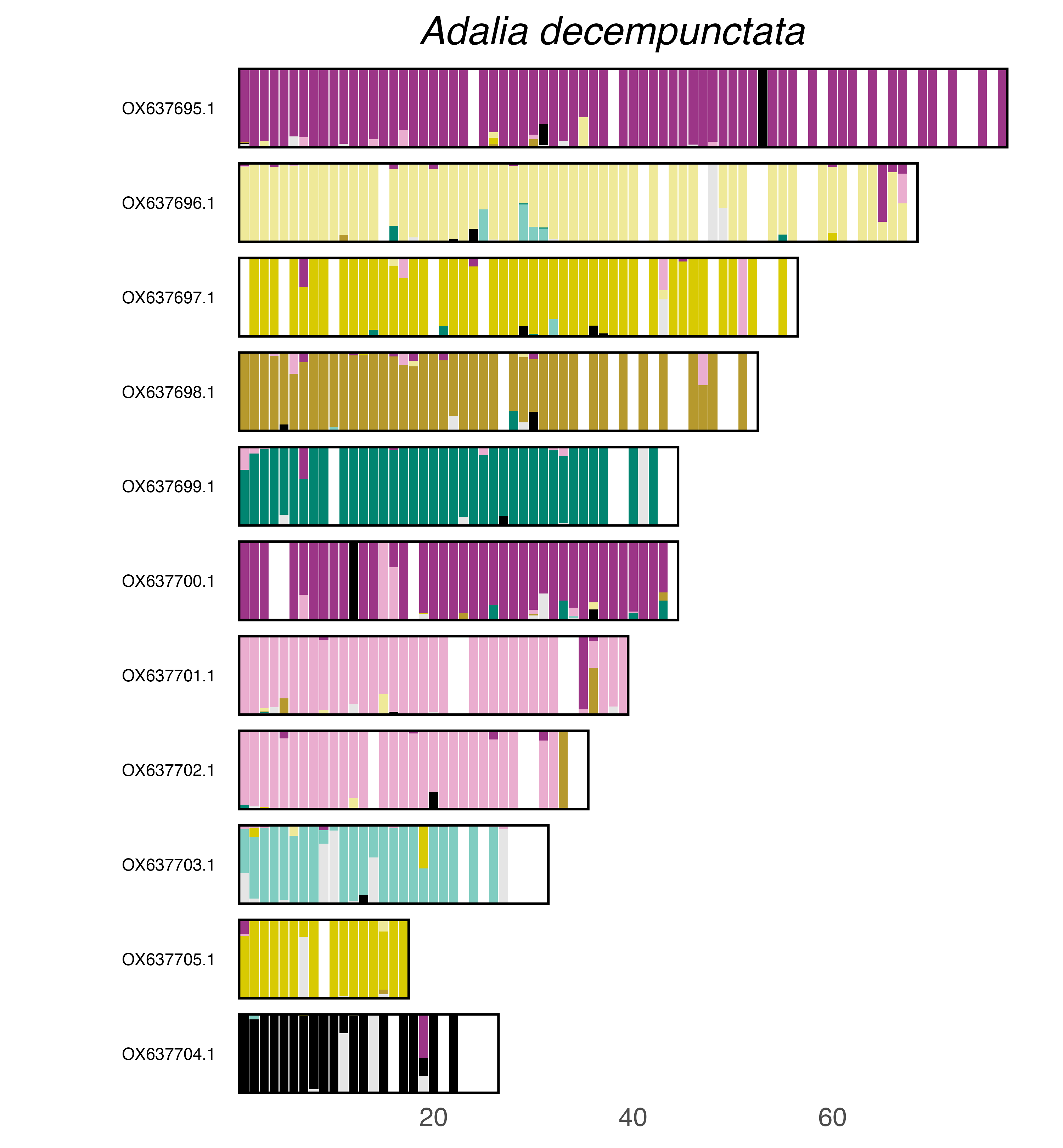

### plot.syngraph_windowbp_1Mb_rel_vertical_chrsize_Amara_aenea.png

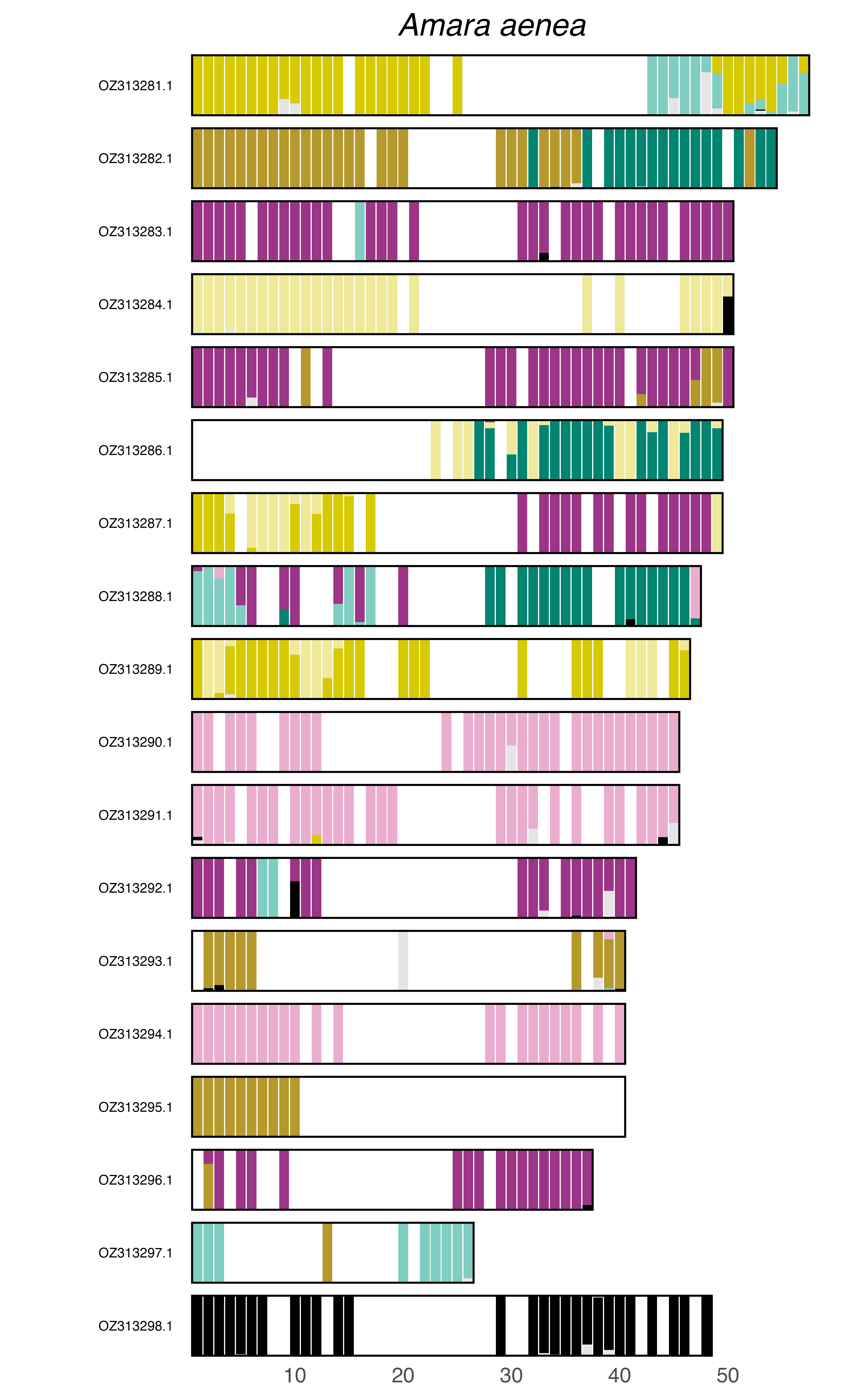

### plot.syngraph_windowbp_1Mb_rel_vertical_chrsize_Anobium_punctatum.png

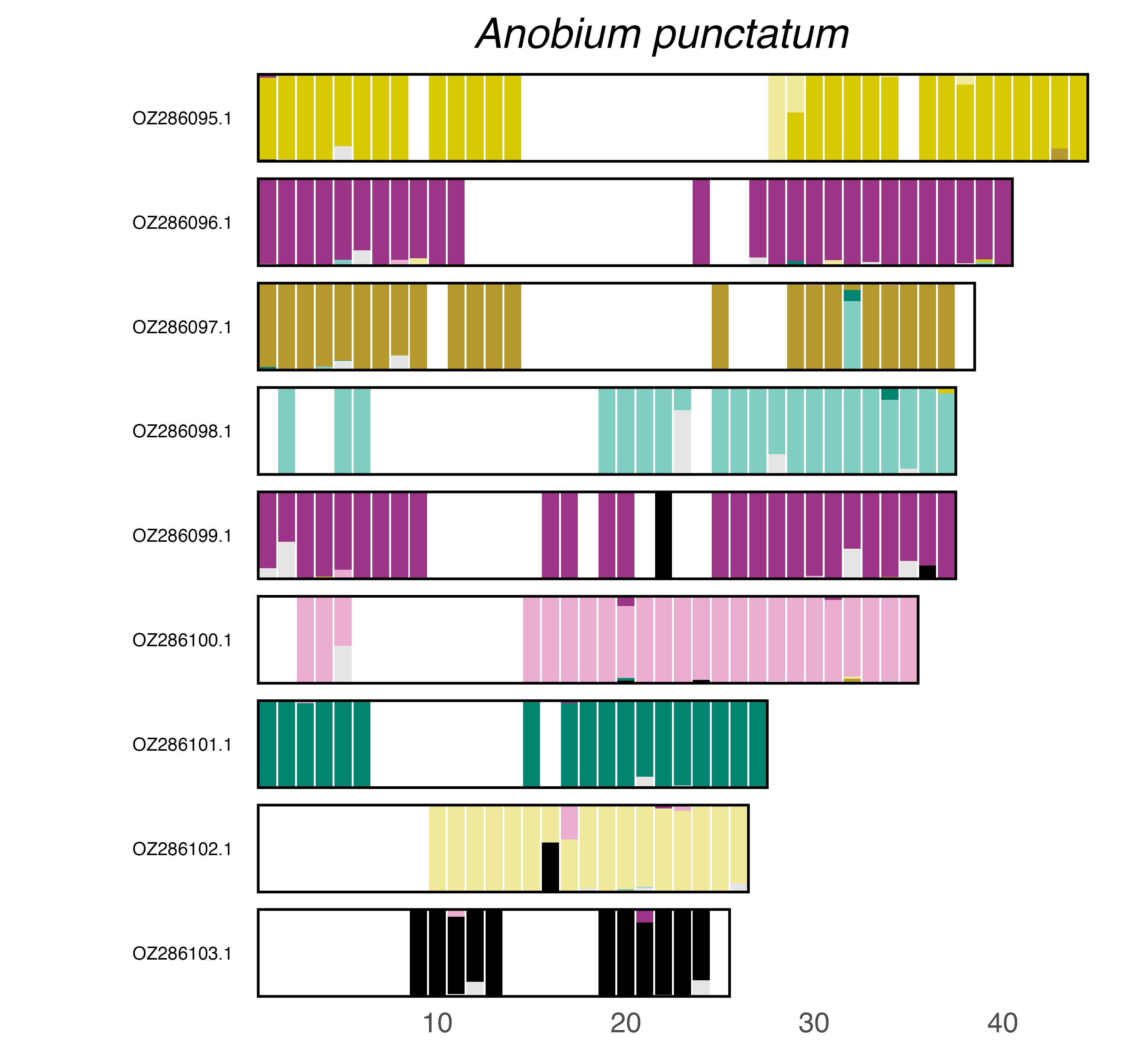

### plot.syngraph_windowbp_1Mb_rel_vertical_chrsize_Anoplophora_chinensis.png

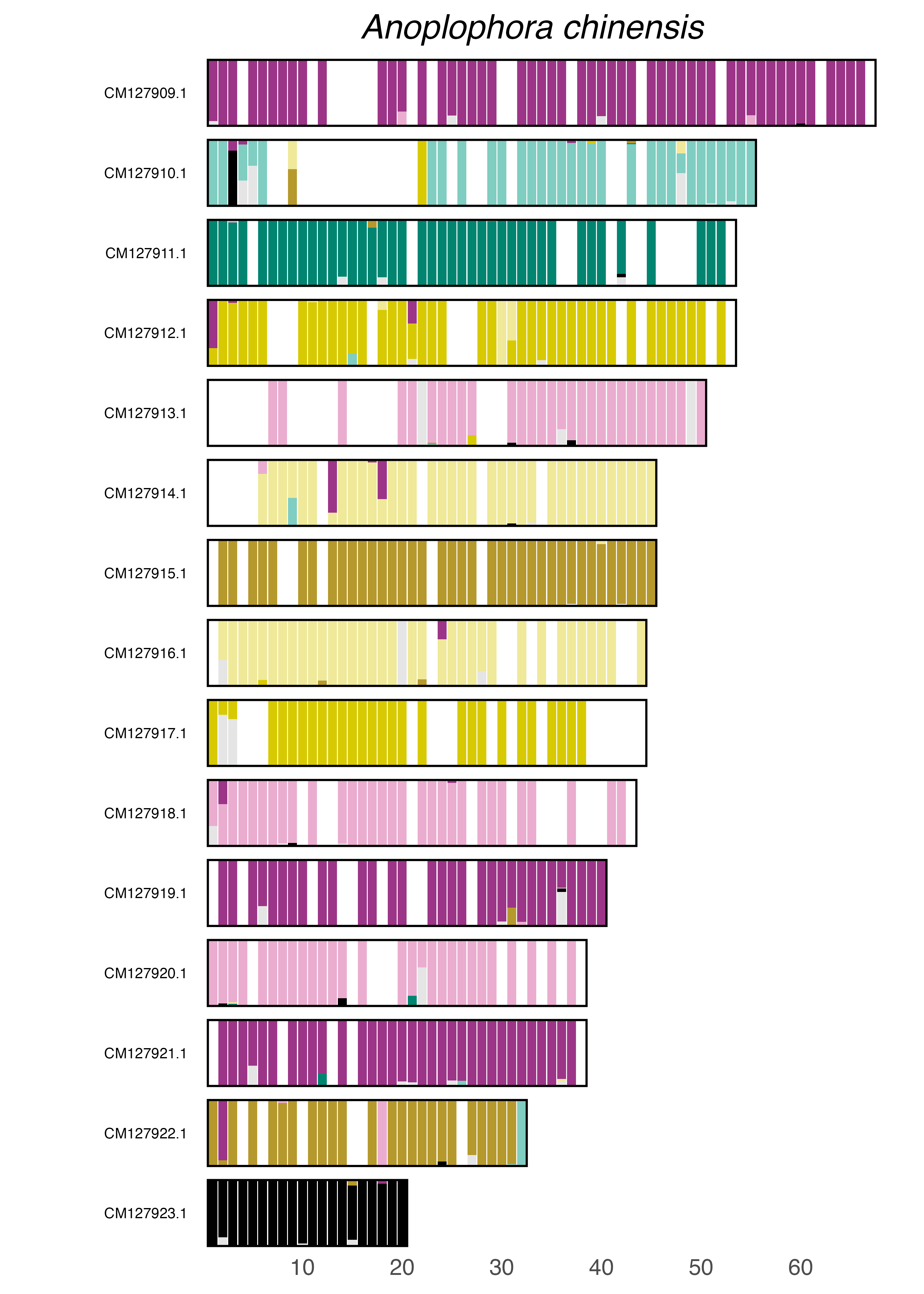

### plot.syngraph_windowbp_1Mb_rel_vertical_chrsize_Anoplophora_malasiaca.png

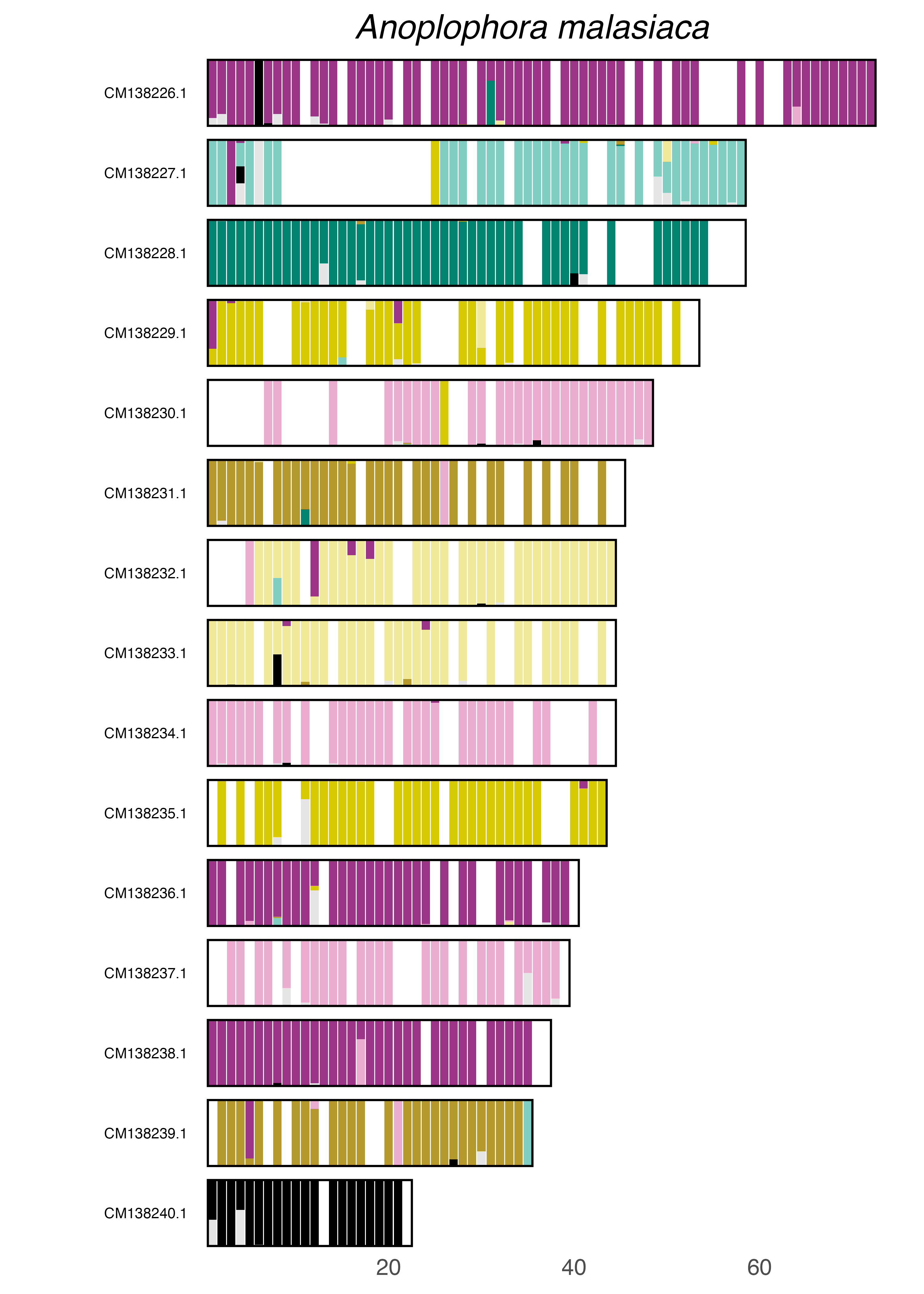

### plot.syngraph_windowbp_1Mb_rel_vertical_chrsize_Anthocomus_fasciatus.png

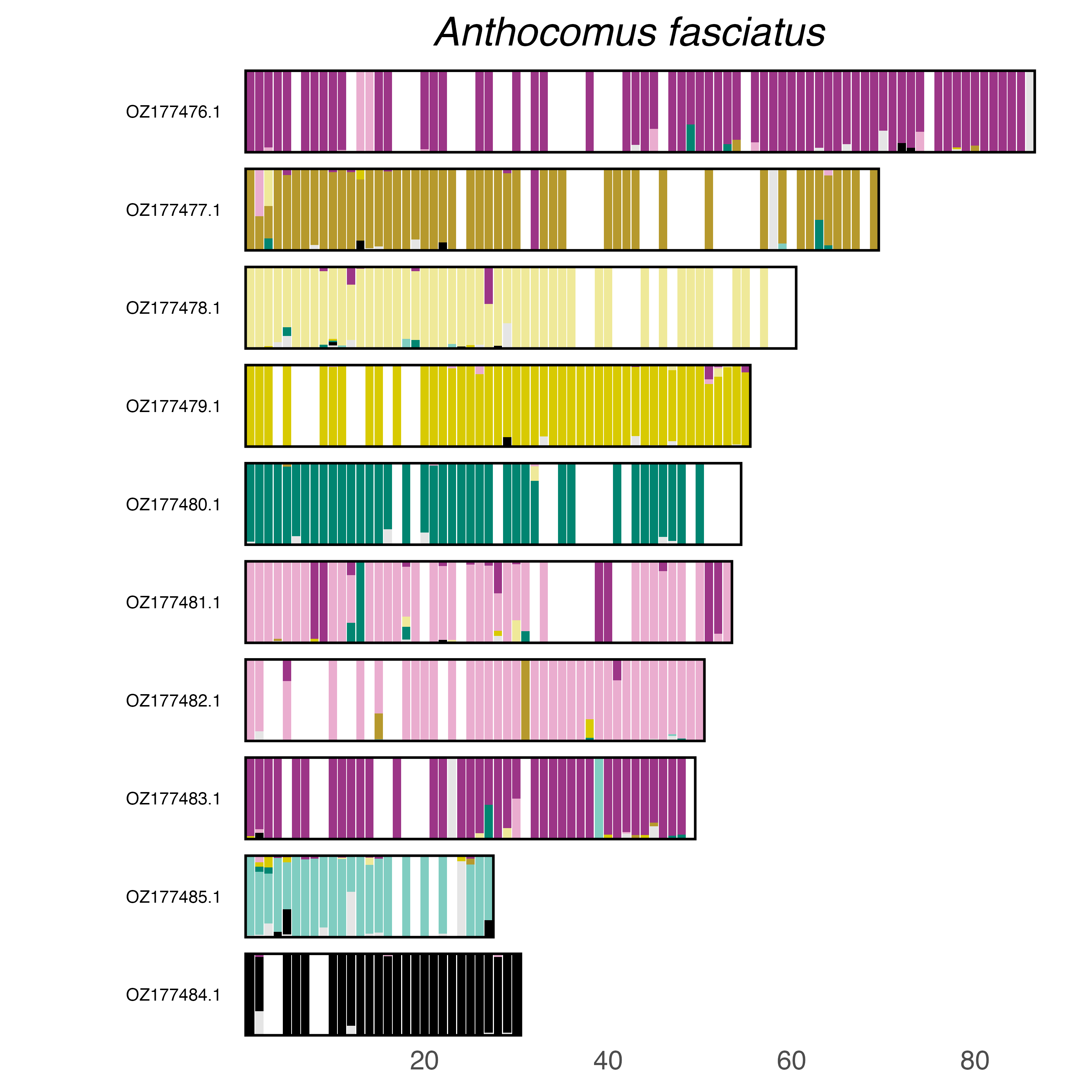

### plot.syngraph_windowbp_1Mb_rel_vertical_chrsize_Arhopalus_rusticus.png

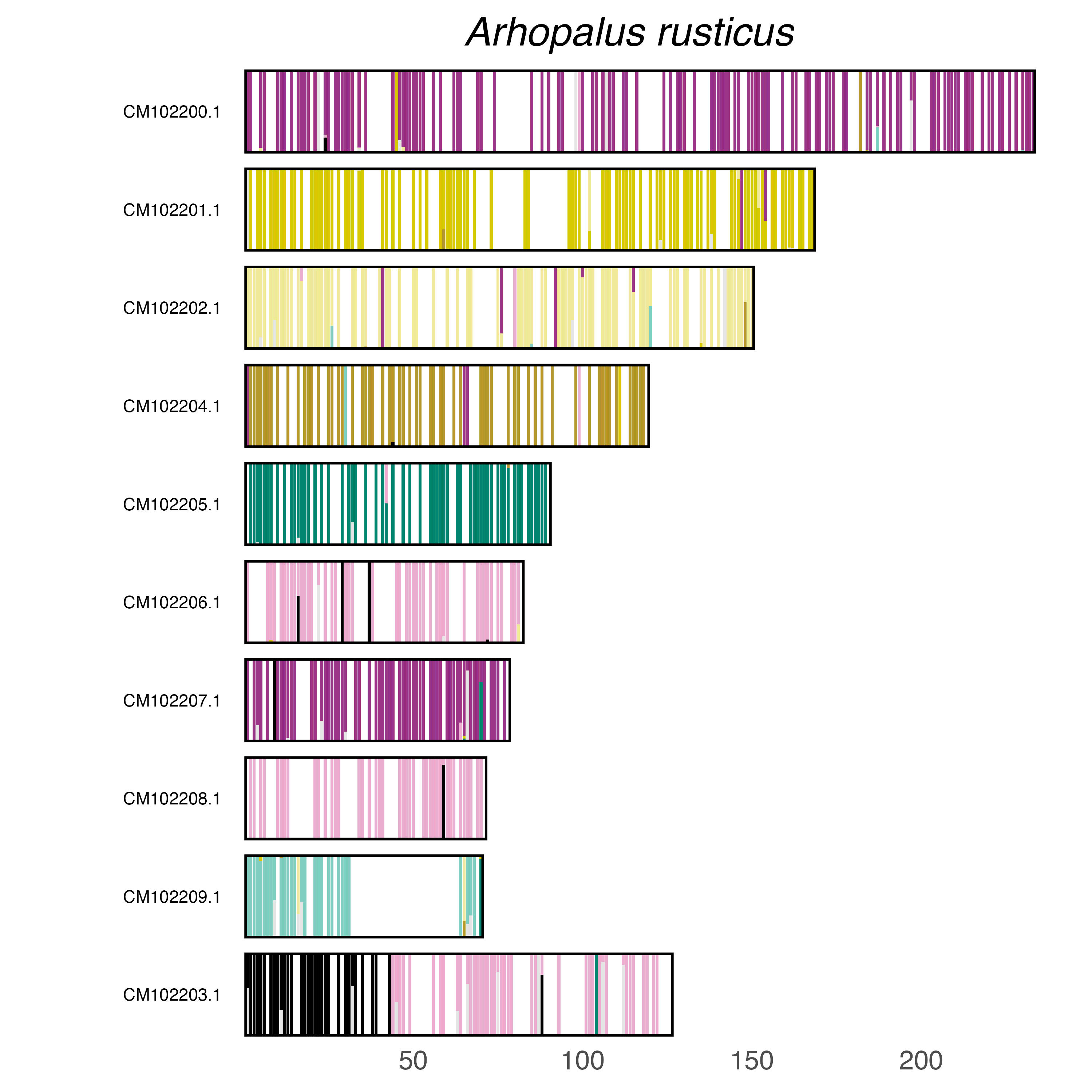

### plot.syngraph_windowbp_1Mb_rel_vertical_chrsize_Arhopalus_unicolor.png

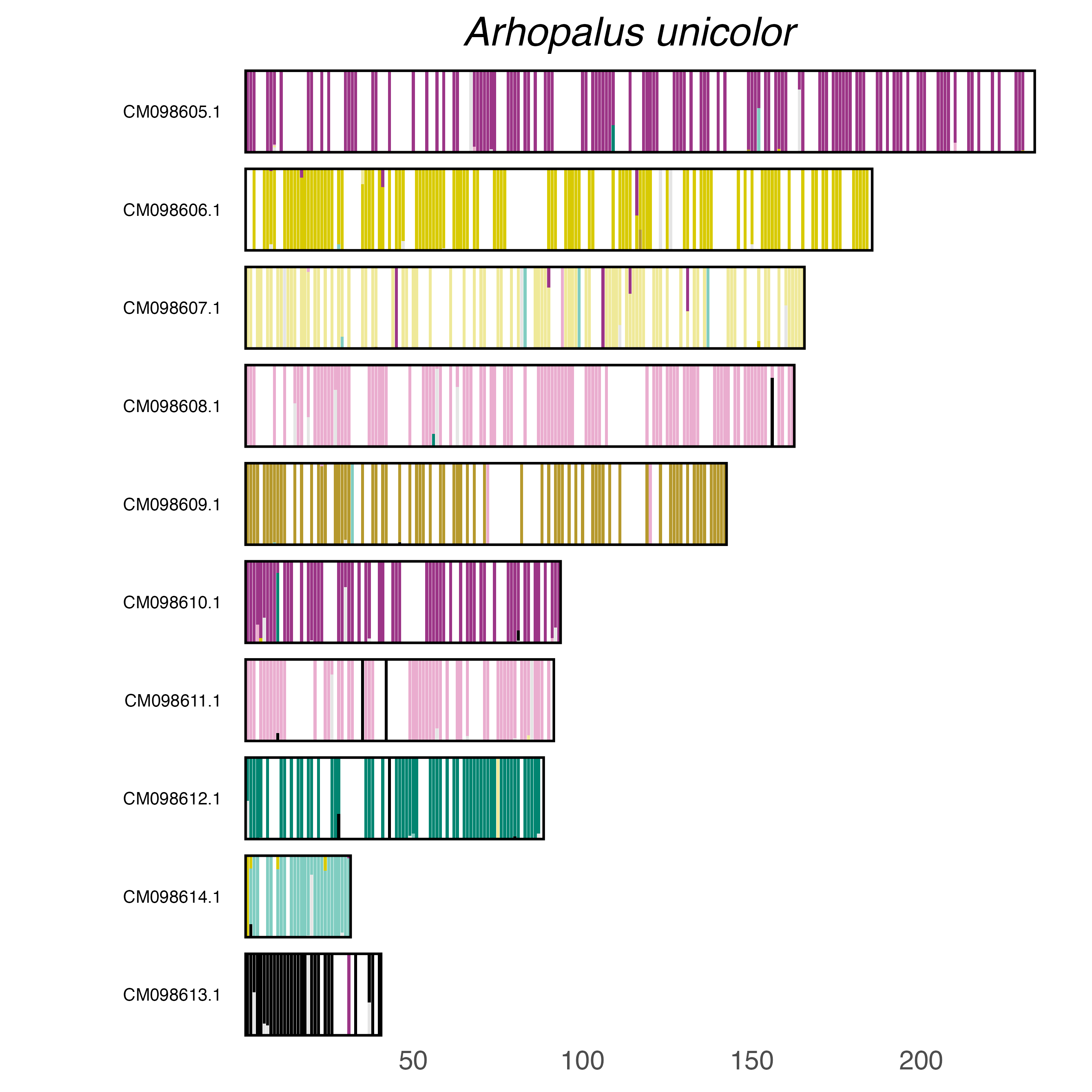

### plot.syngraph_windowbp_1Mb_rel_vertical_chrsize_Bitoma_crenata.png

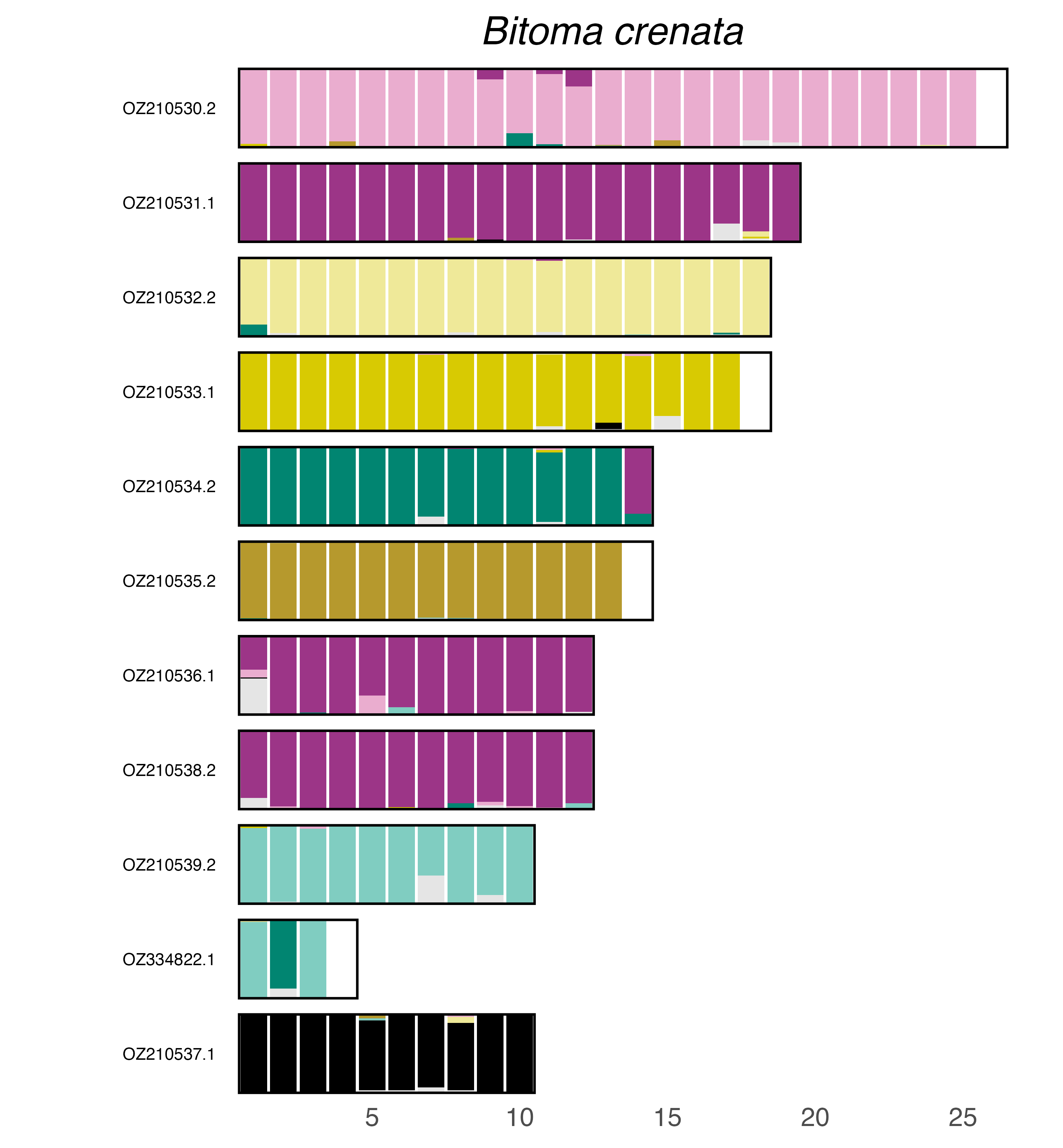

### plot.syngraph_windowbp_1Mb_rel_vertical_chrsize_Brachypterus_glaber.png

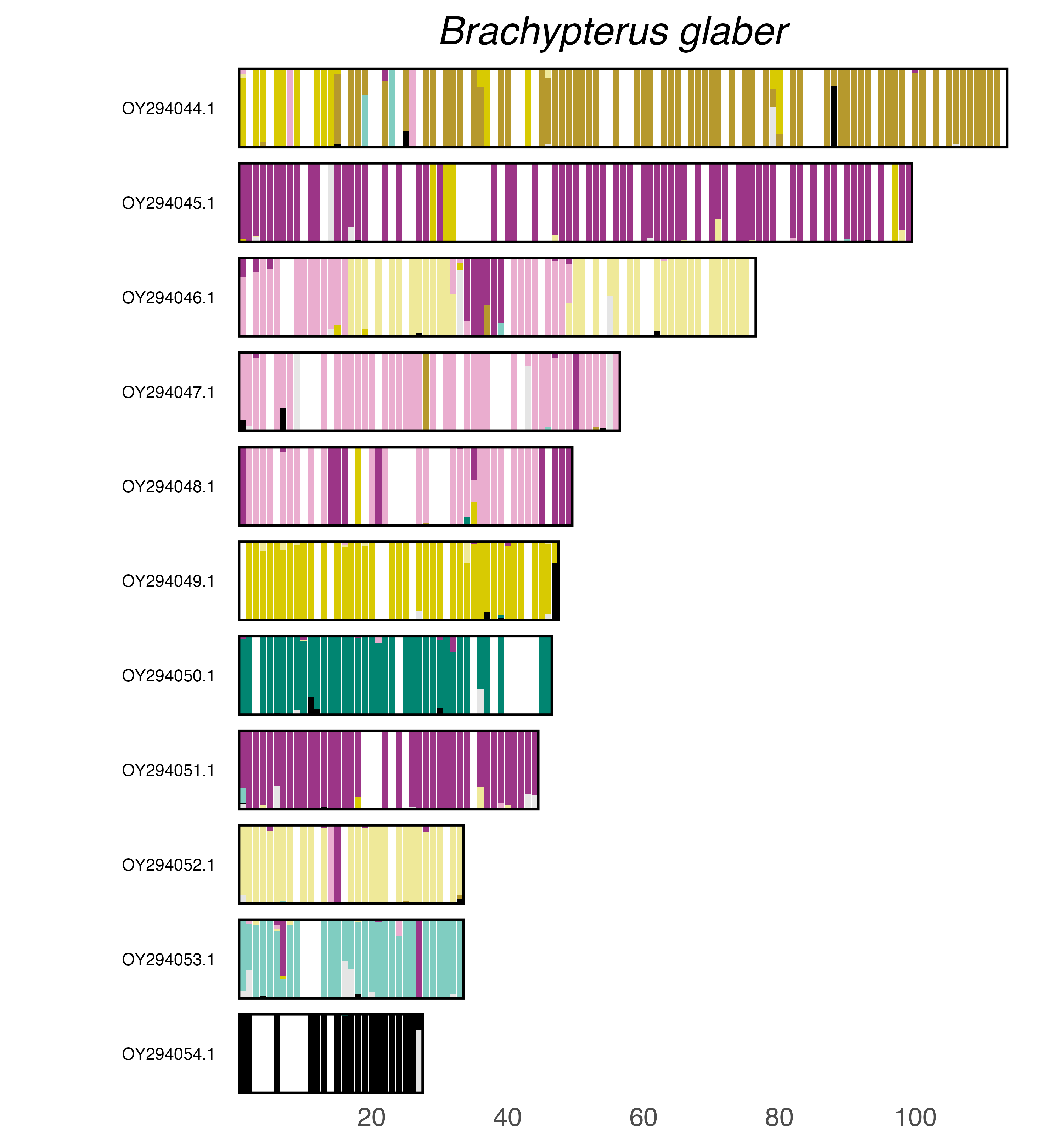

### plot.syngraph_windowbp_1Mb_rel_vertical_chrsize_Brassicogethes_aeneus.png

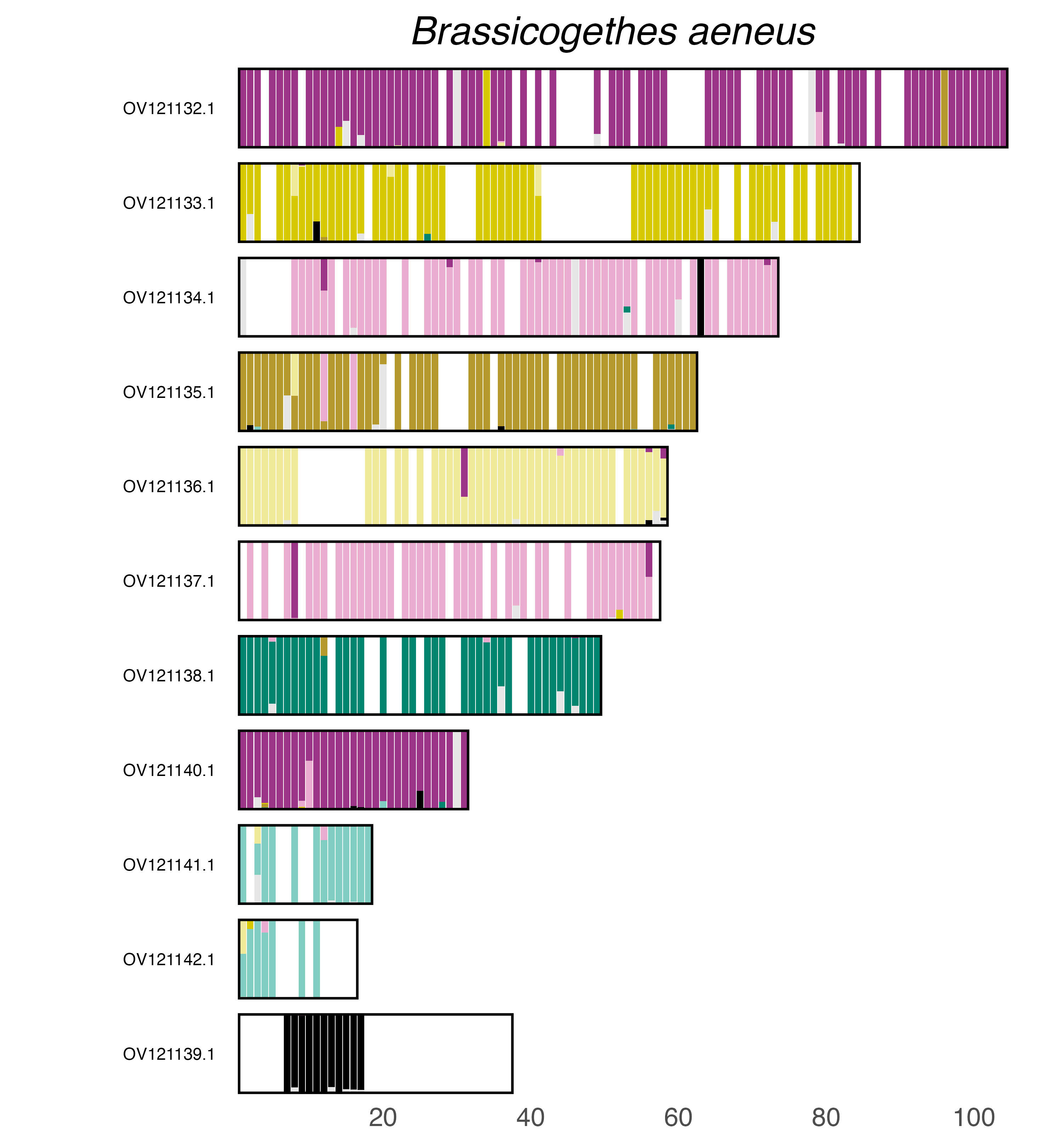

### plot.syngraph_windowbp_1Mb_rel_vertical_chrsize_Calathus_melanocephalus.png

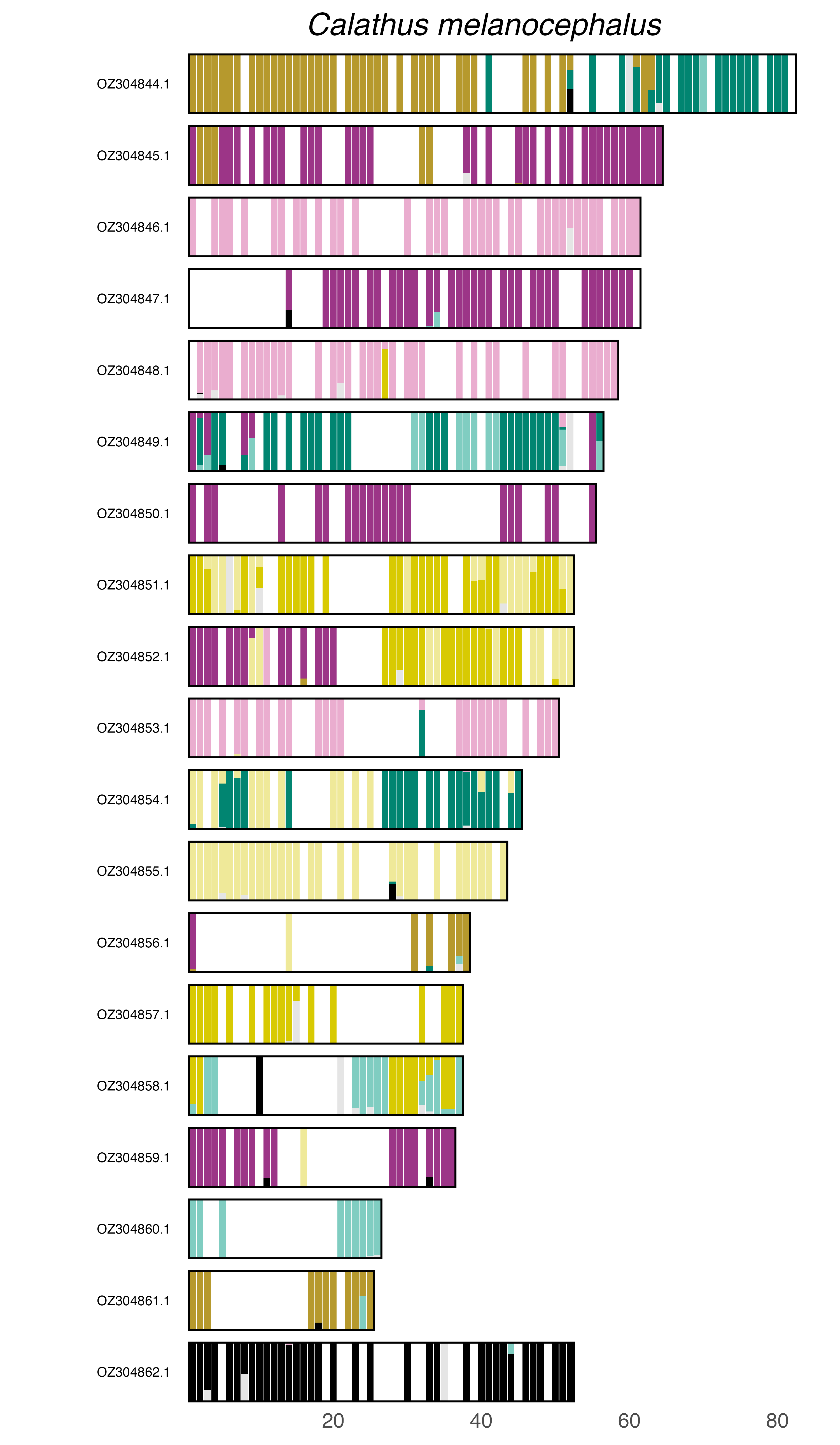

### plot.syngraph_windowbp_1Mb_rel_vertical_chrsize_Calodromius_spilotus.png

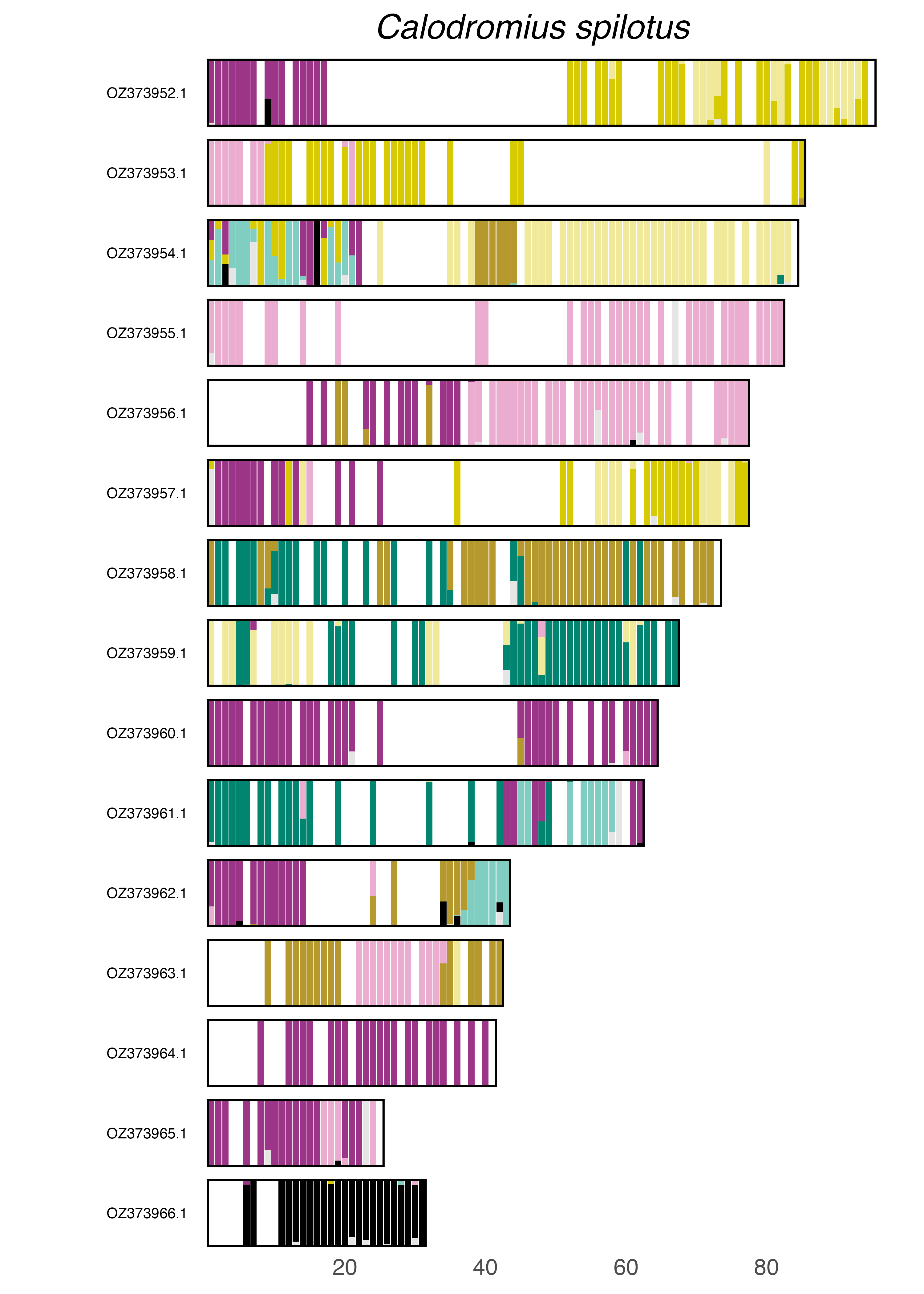

### plot.syngraph_windowbp_1Mb_rel_vertical_chrsize_Cantharis_flavilabris.png

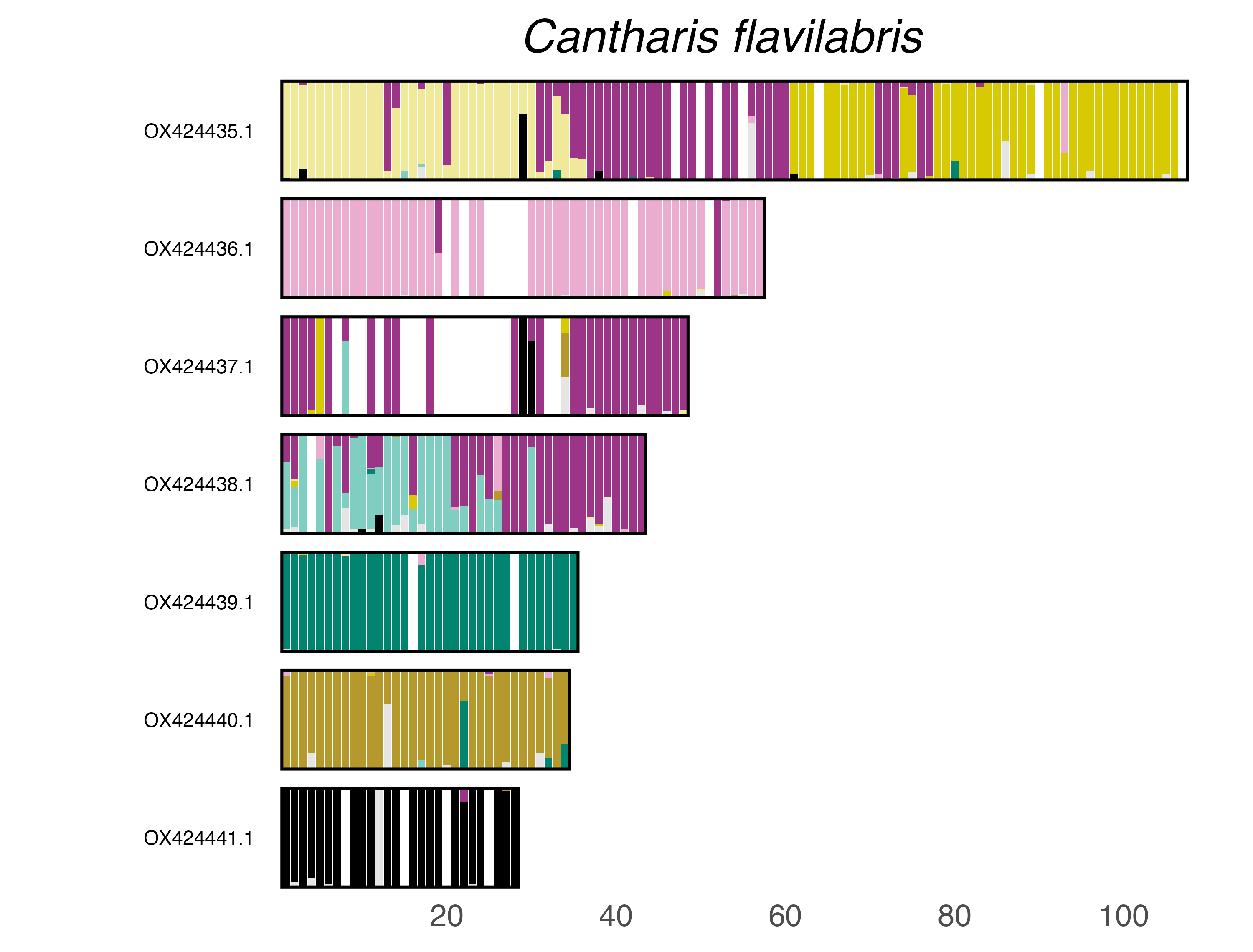

### plot.syngraph_windowbp_1Mb_rel_vertical_chrsize_Carabus_banonii.png

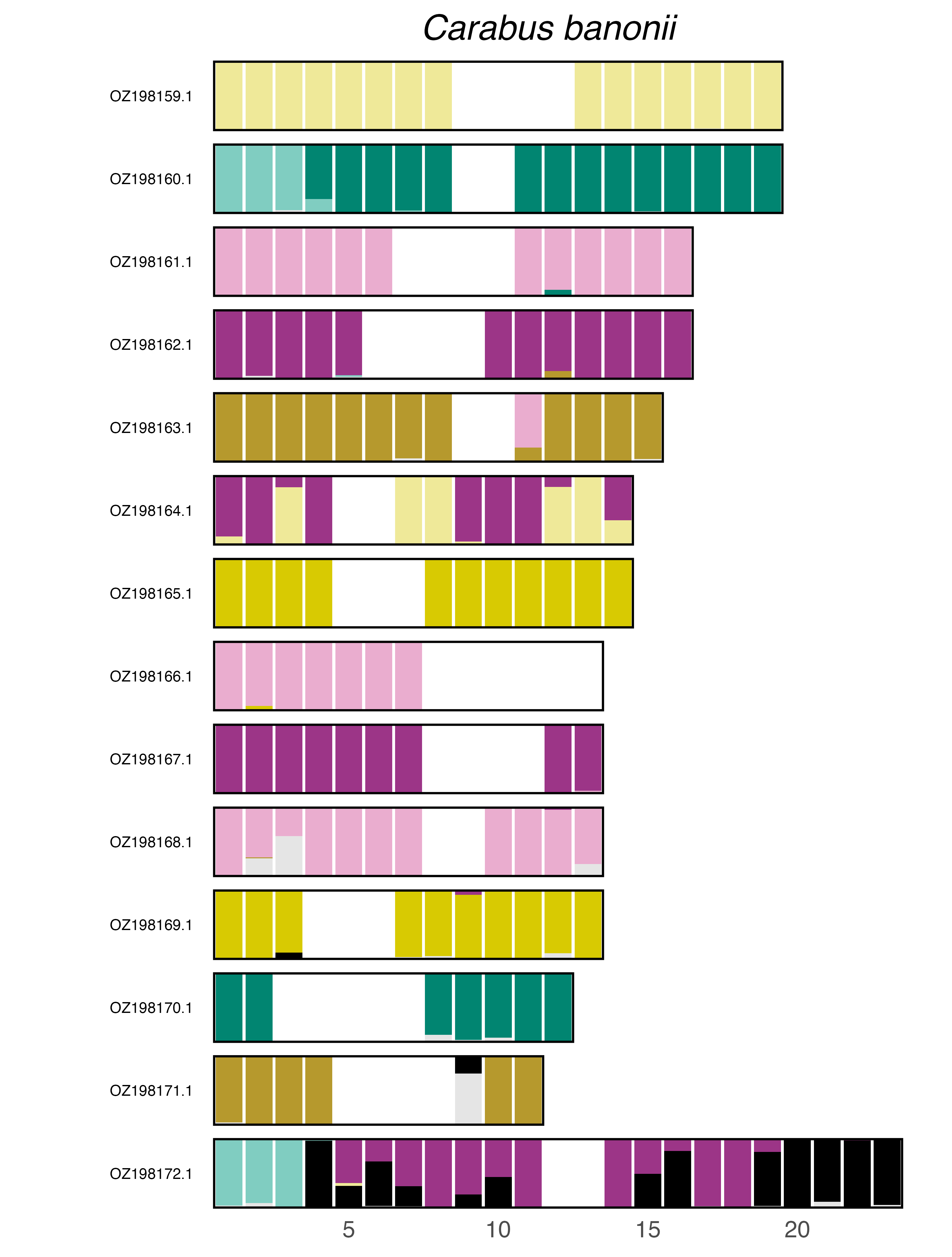

### plot.syngraph_windowbp_1Mb_rel_vertical_chrsize_Carabus_nemoralis.png

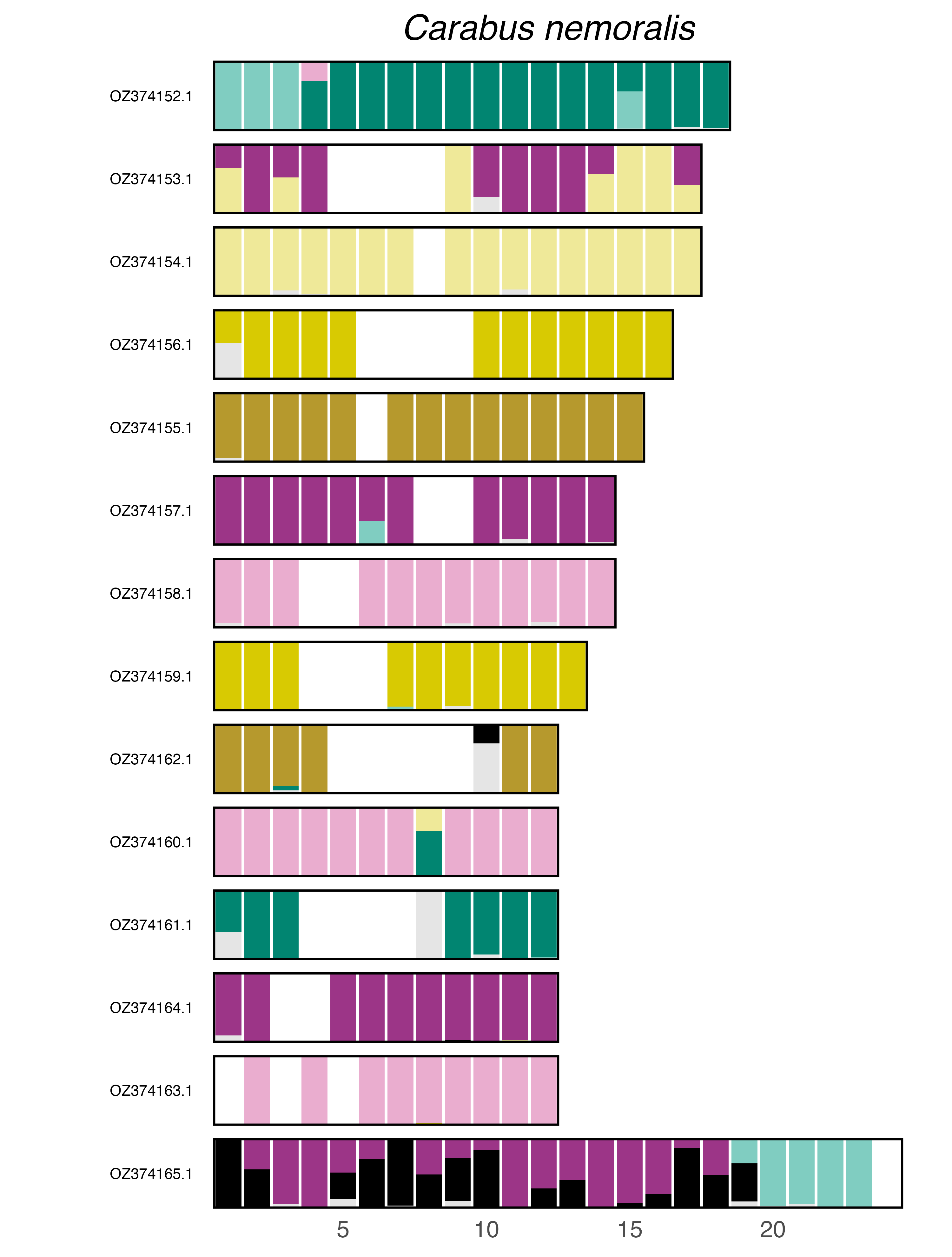

### plot.syngraph_windowbp_1Mb_rel_vertical_chrsize_Carabus_problematicus.png

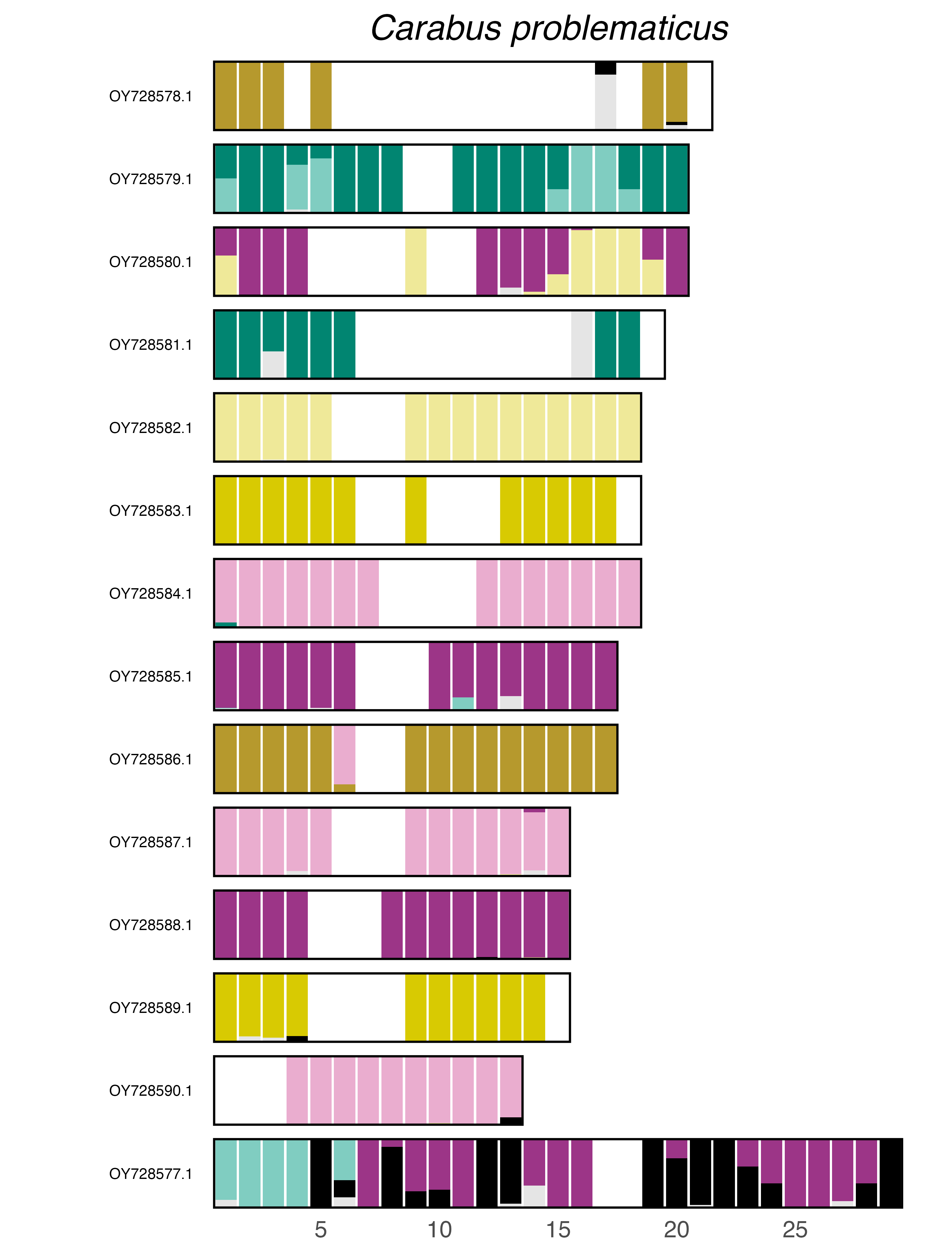

### plot.syngraph_windowbp_1Mb_rel_vertical_chrsize_Carabus_staehlini.png

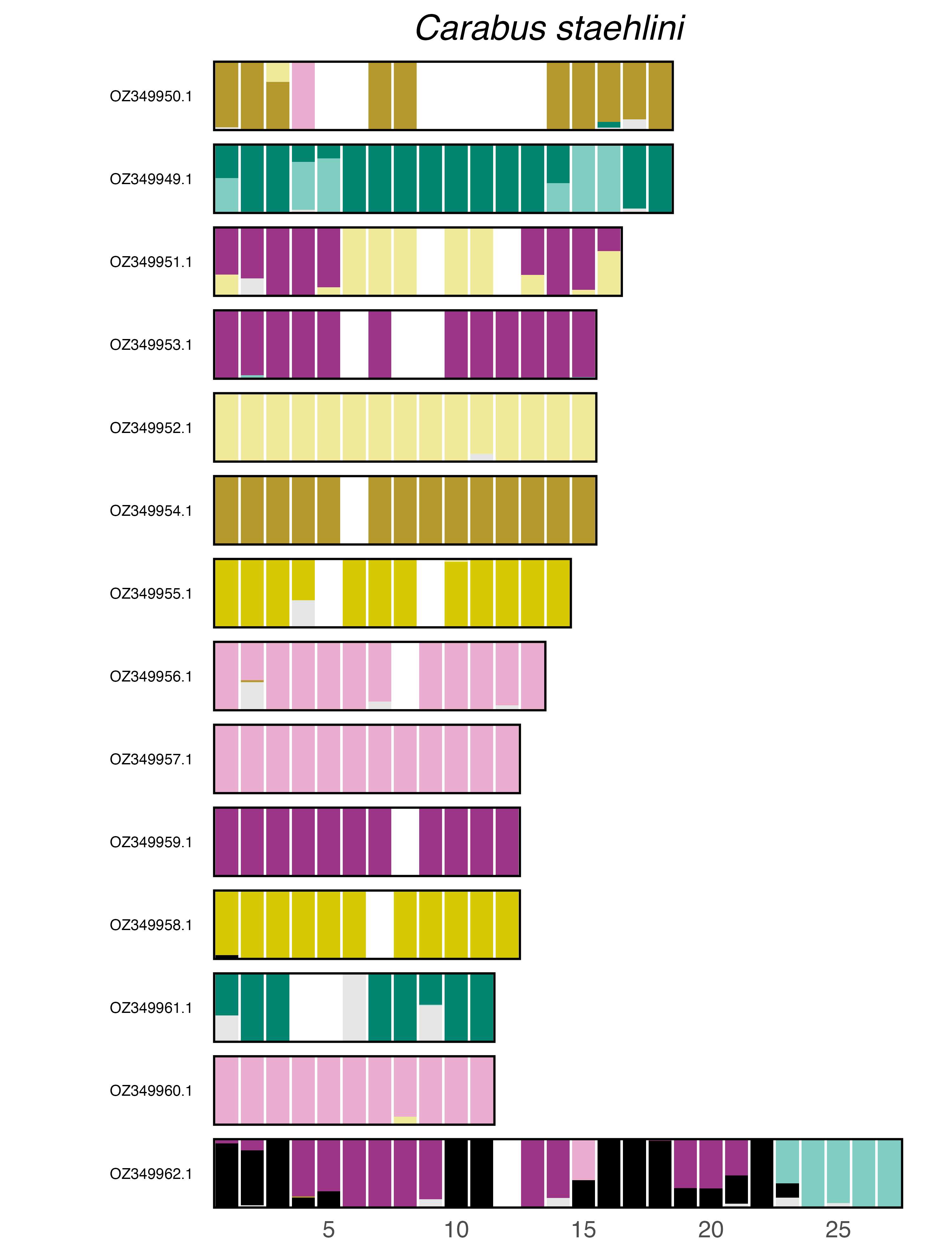

### plot.syngraph_windowbp_1Mb_rel_vertical_chrsize_Cartodere_bifasciata.png

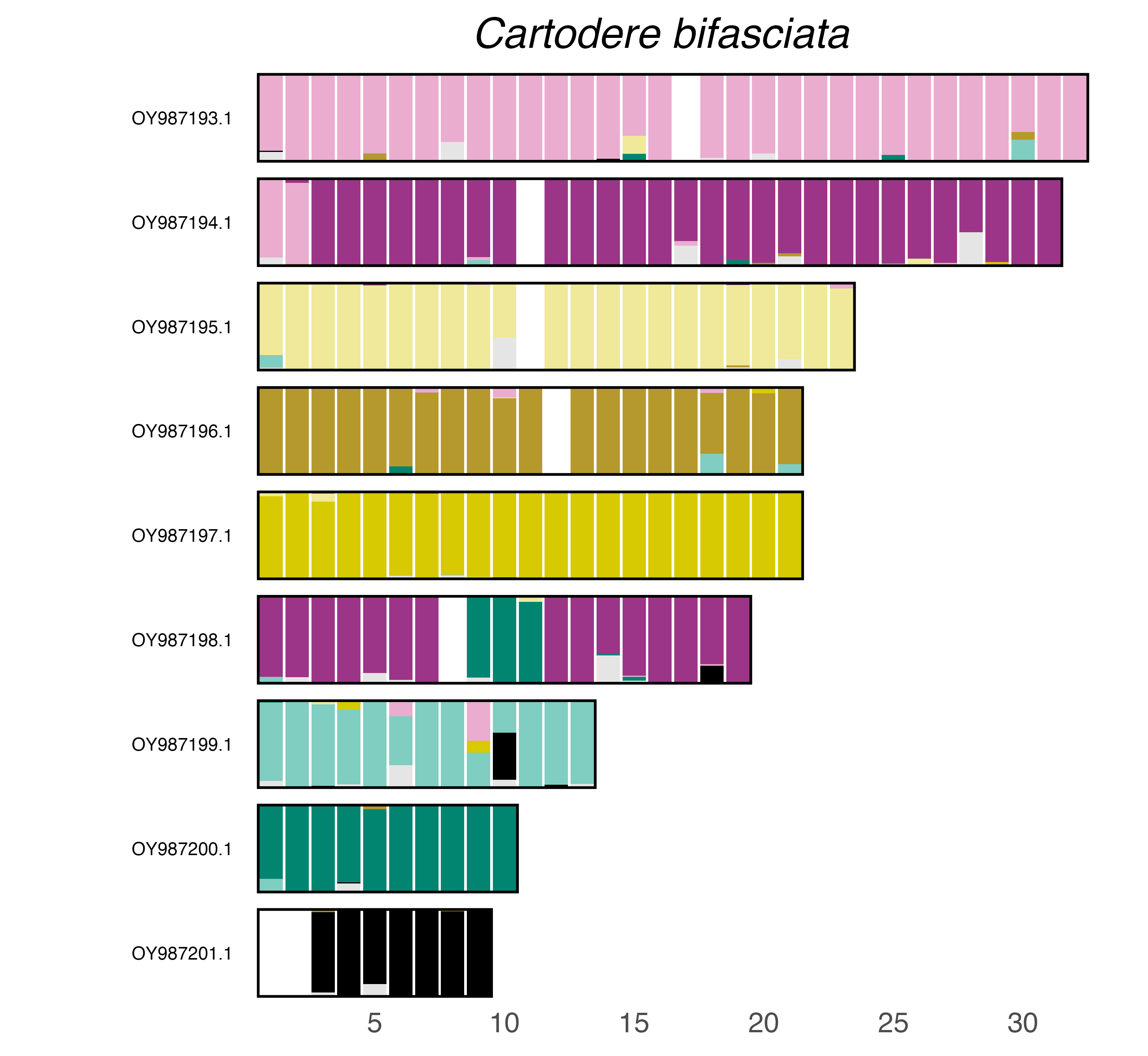

### plot.syngraph_windowbp_1Mb_rel_vertical_chrsize_Cetonia_aurata.png

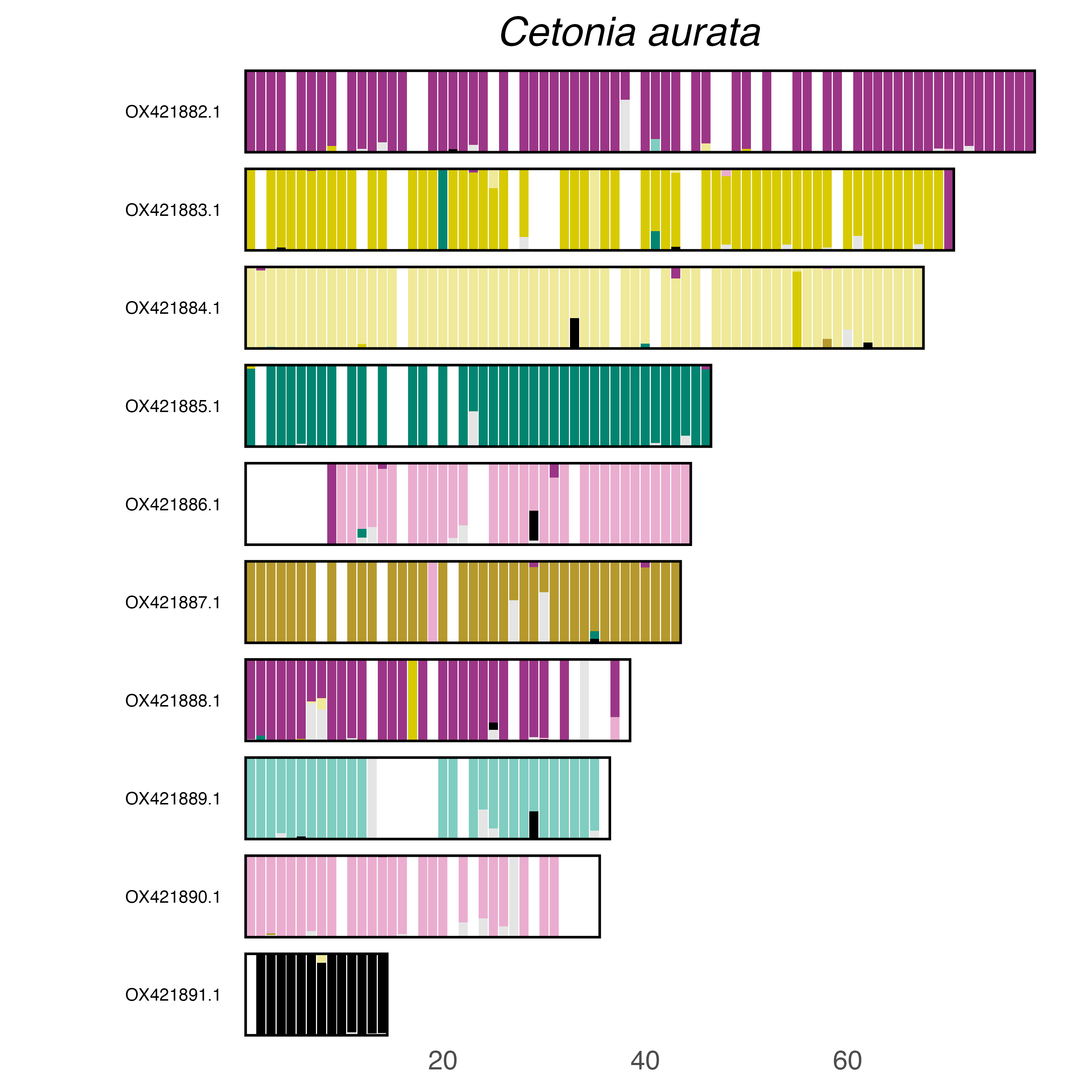

### plot.syngraph_windowbp_1Mb_rel_vertical_chrsize_Ceutorhynchus_obstrictus.png

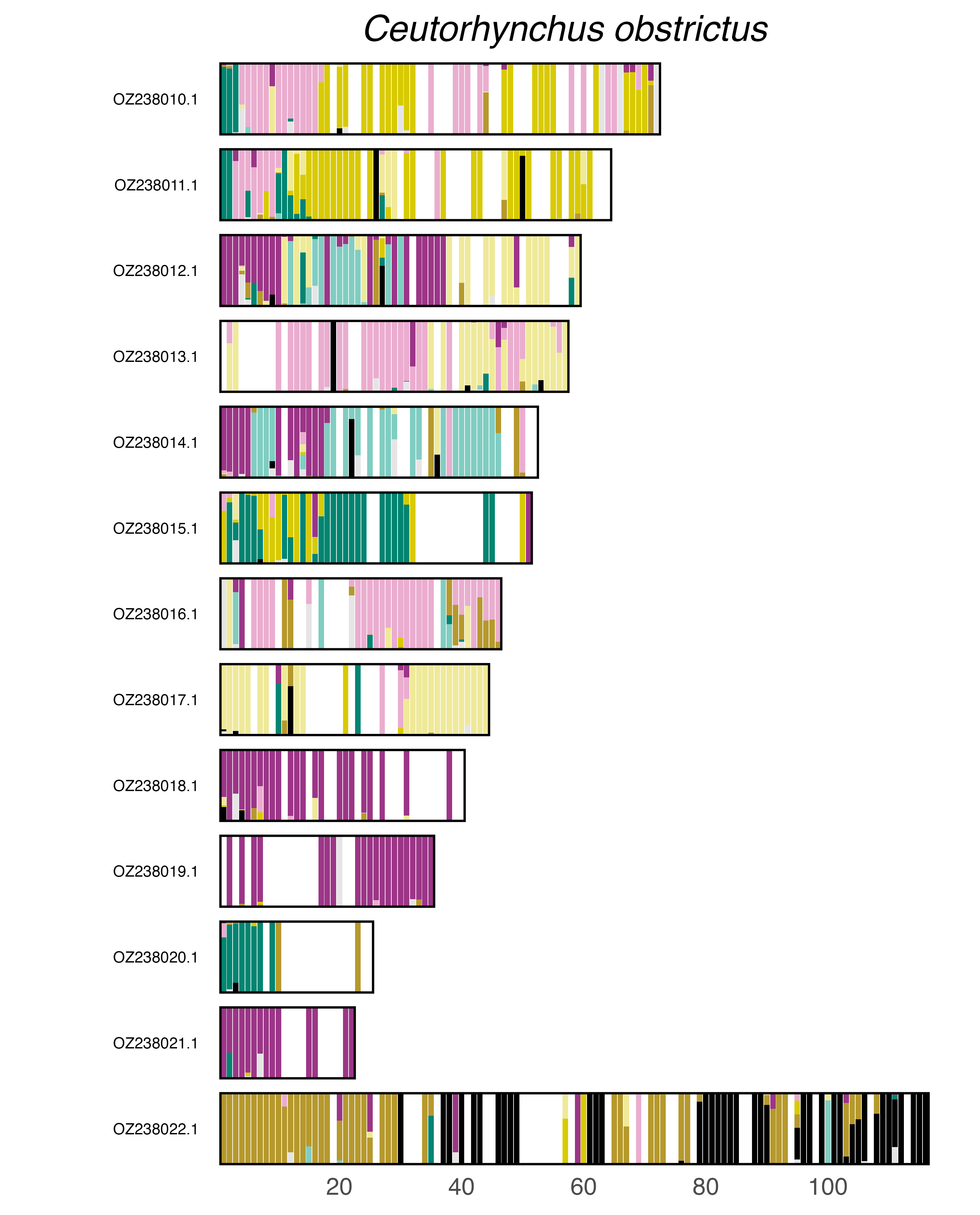

### plot.syngraph_windowbp_1Mb_rel_vertical_chrsize_Chrysomela_saliceti.png

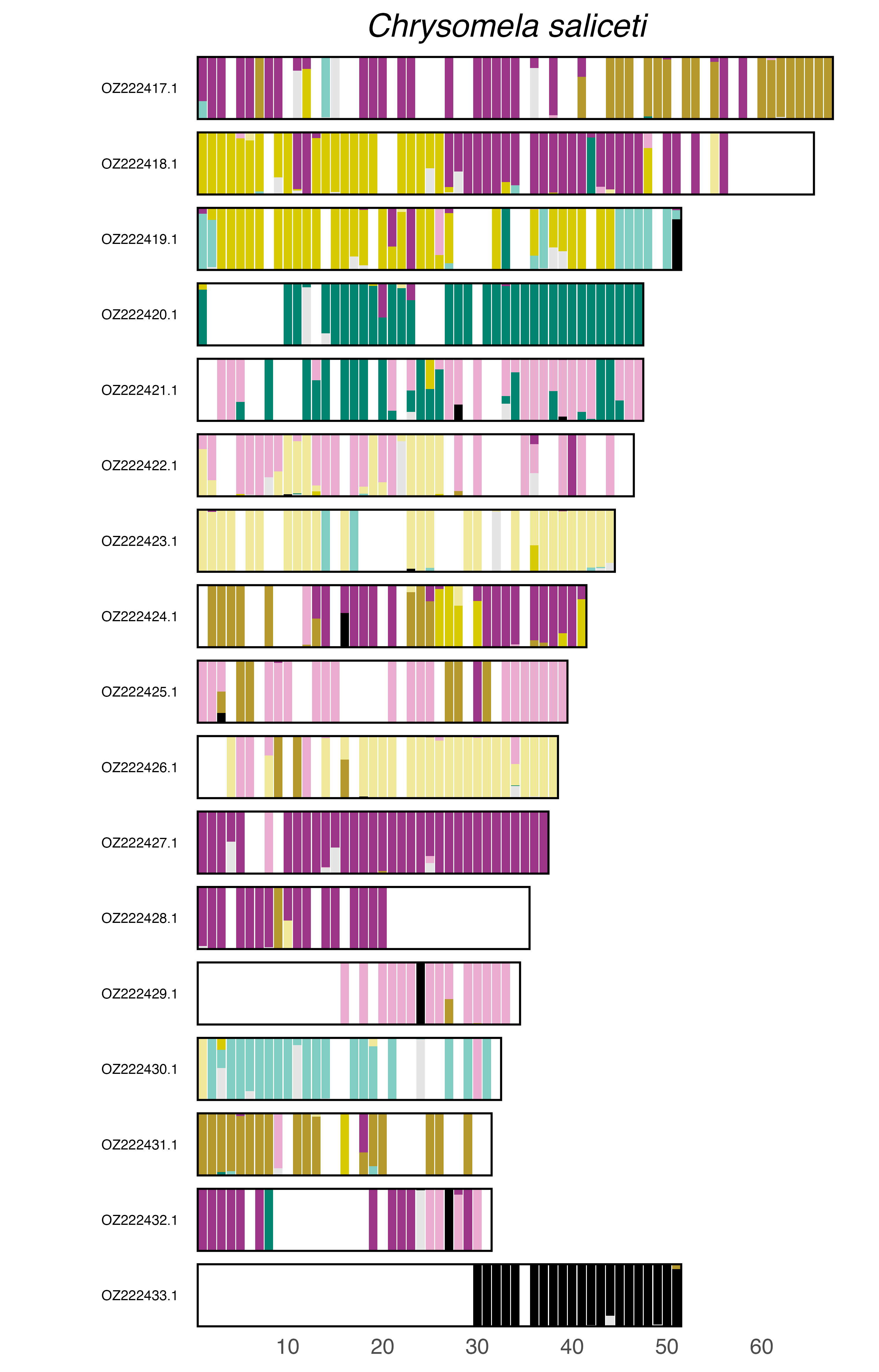

### plot.syngraph_windowbp_1Mb_rel_vertical_chrsize_Cis_bilamellatus.png

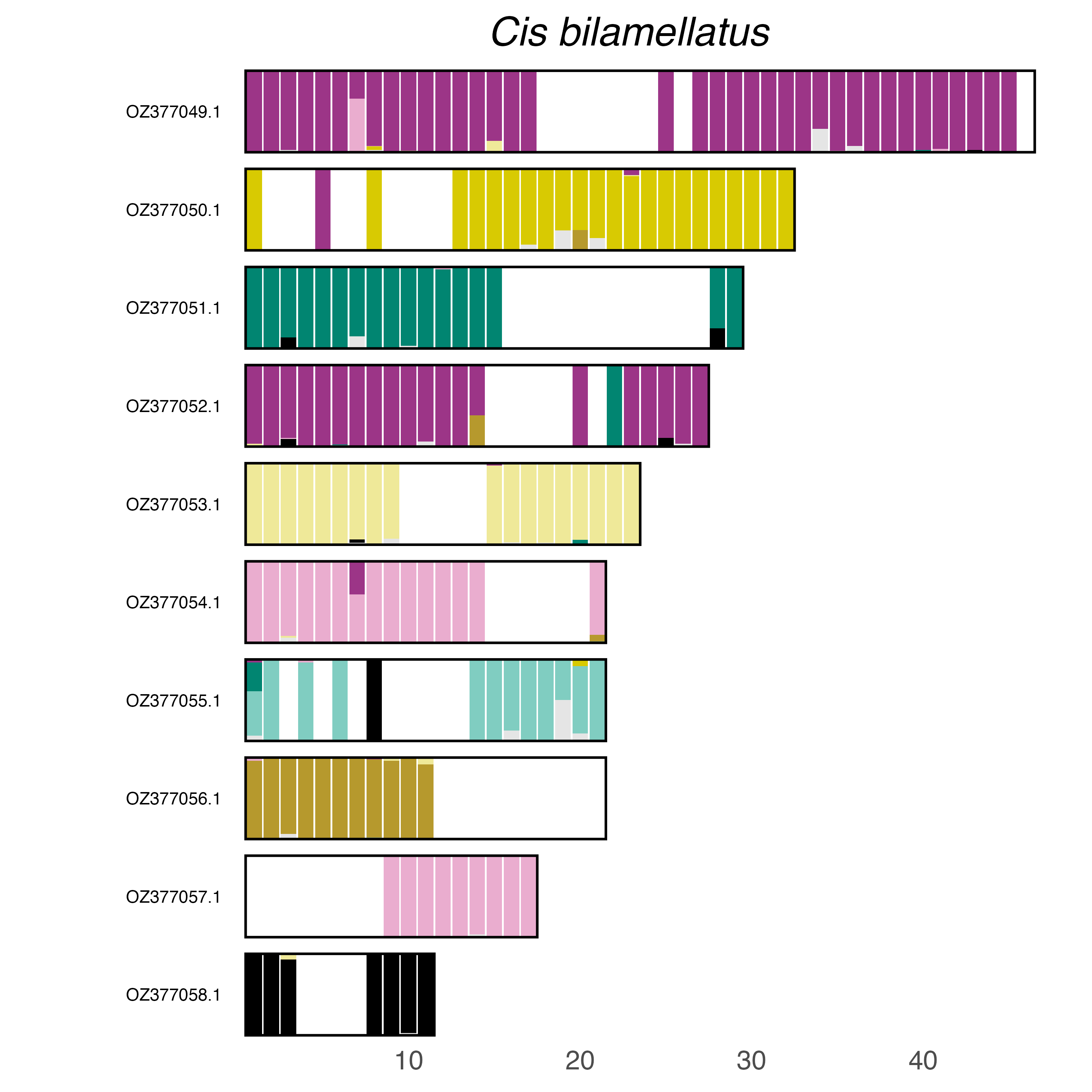

### plot.syngraph_windowbp_1Mb_rel_vertical_chrsize_Crepidodera_fulvicornis.png

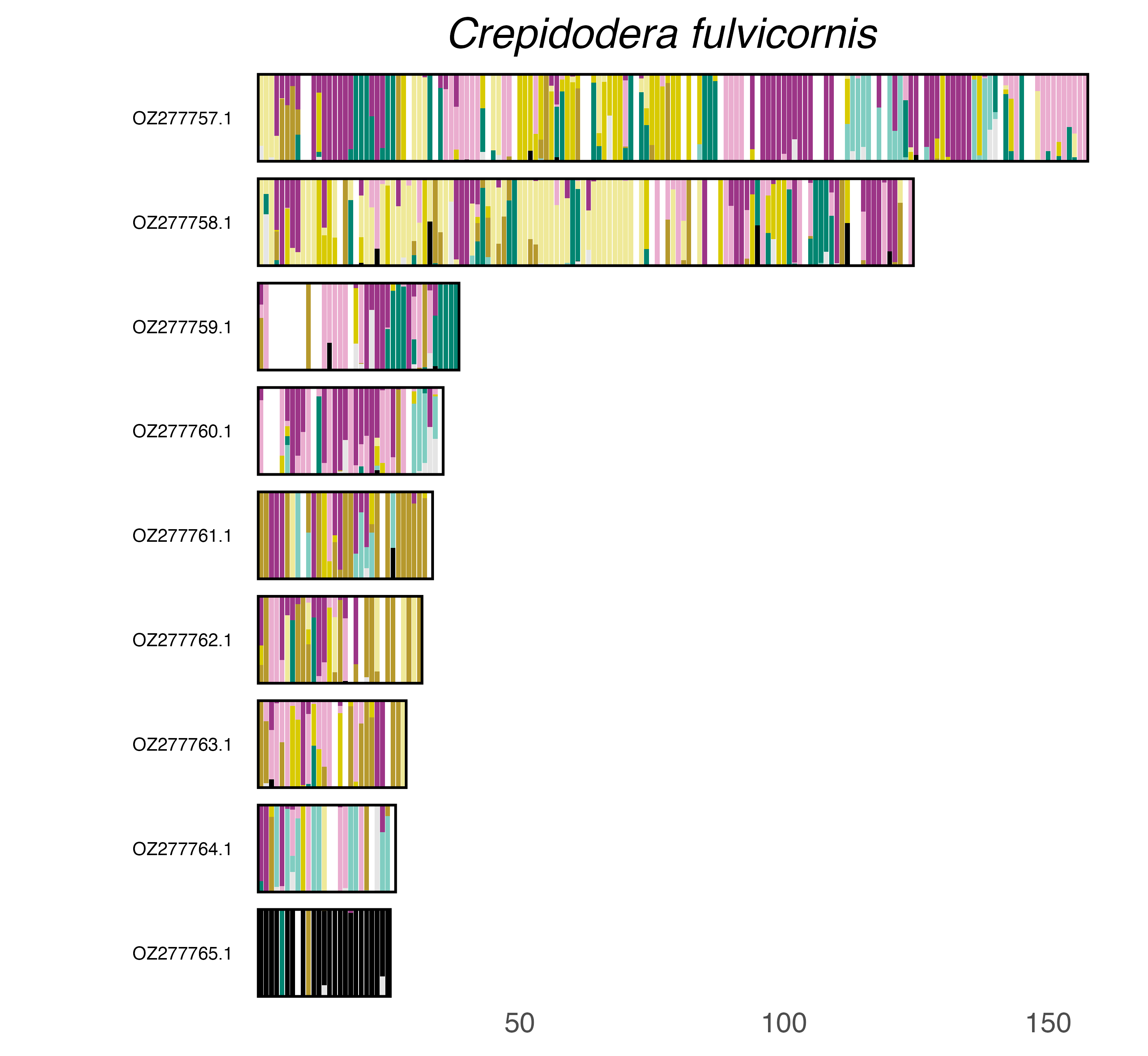

### plot.syngraph_windowbp_1Mb_rel_vertical_chrsize_Ctenicera_cuprea.png

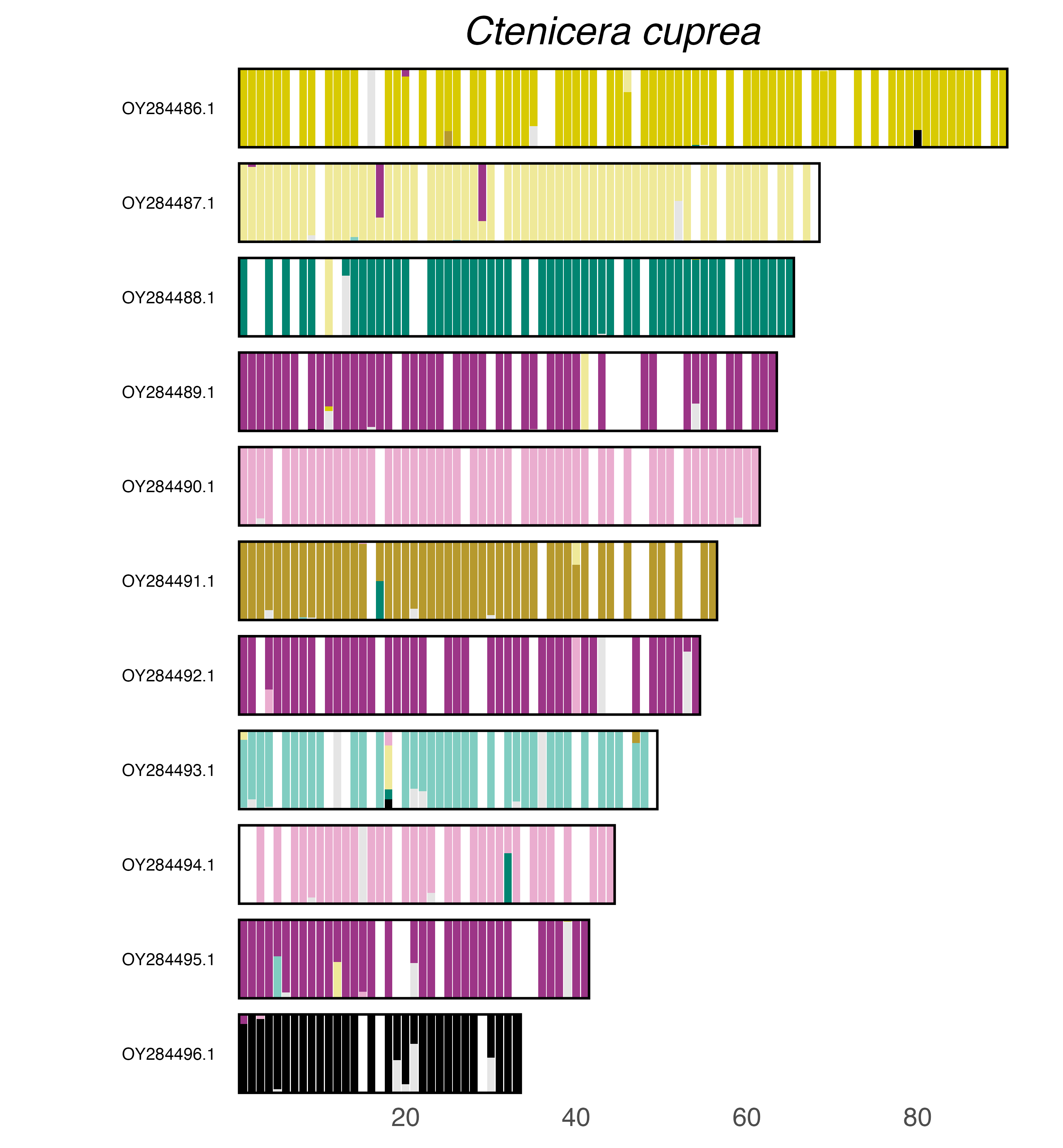

### plot.syngraph_windowbp_1Mb_rel_vertical_chrsize_Curculio_glandium.png

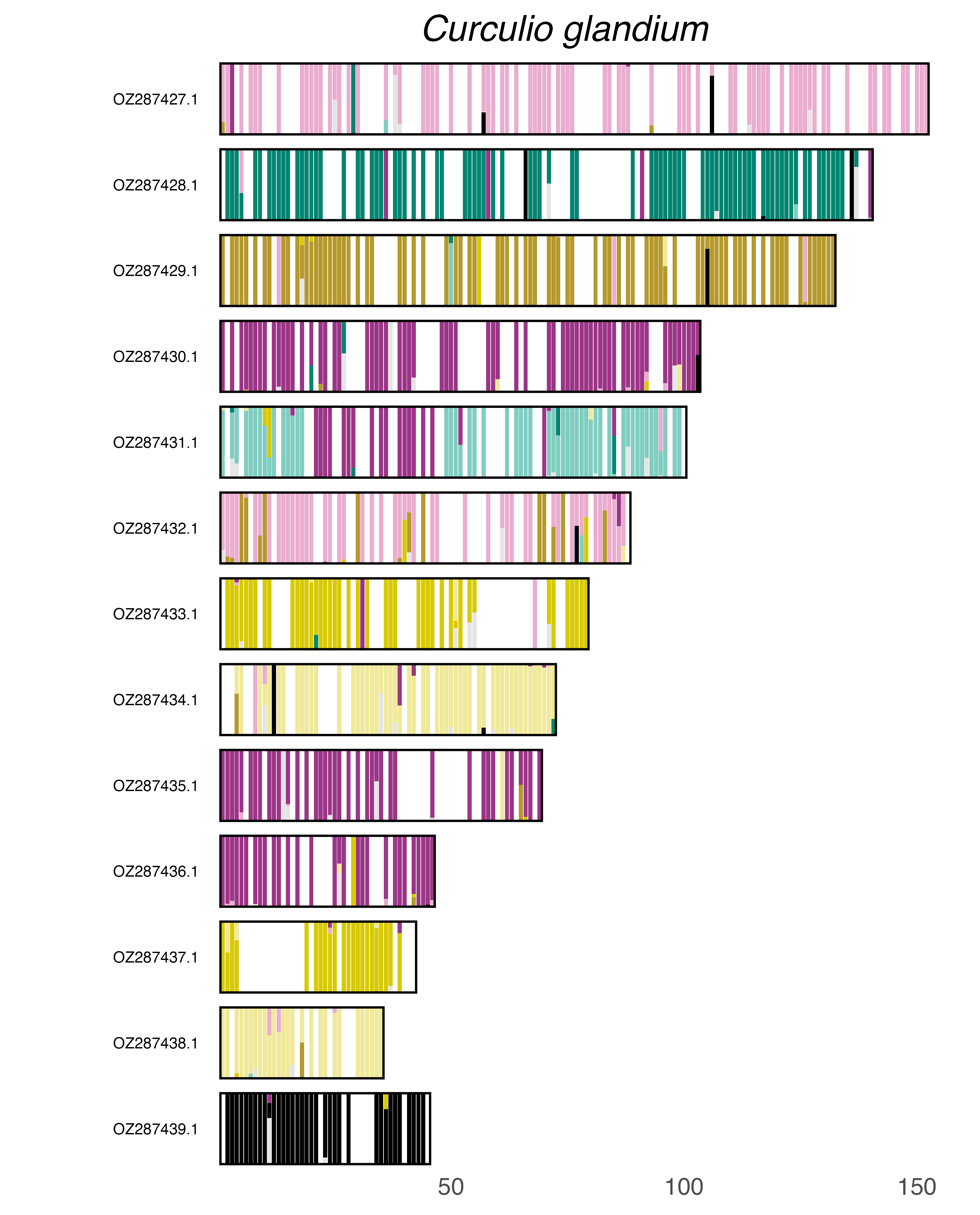

### plot.syngraph_windowbp_1Mb_rel_vertical_chrsize_Dascillus_cervinus.png

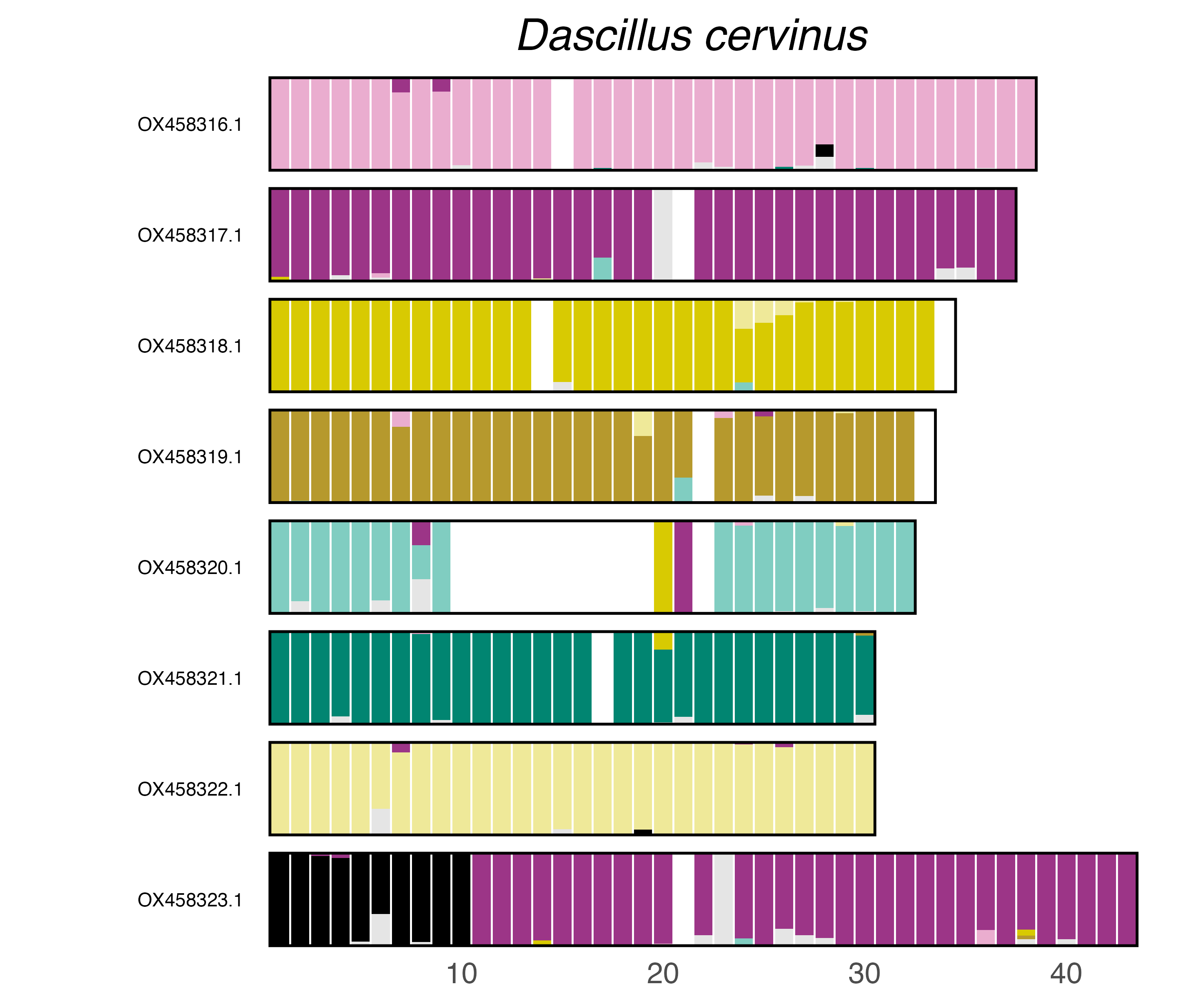

### plot.syngraph_windowbp_1Mb_rel_vertical_chrsize_Diabrotica_undecimpunctata.png

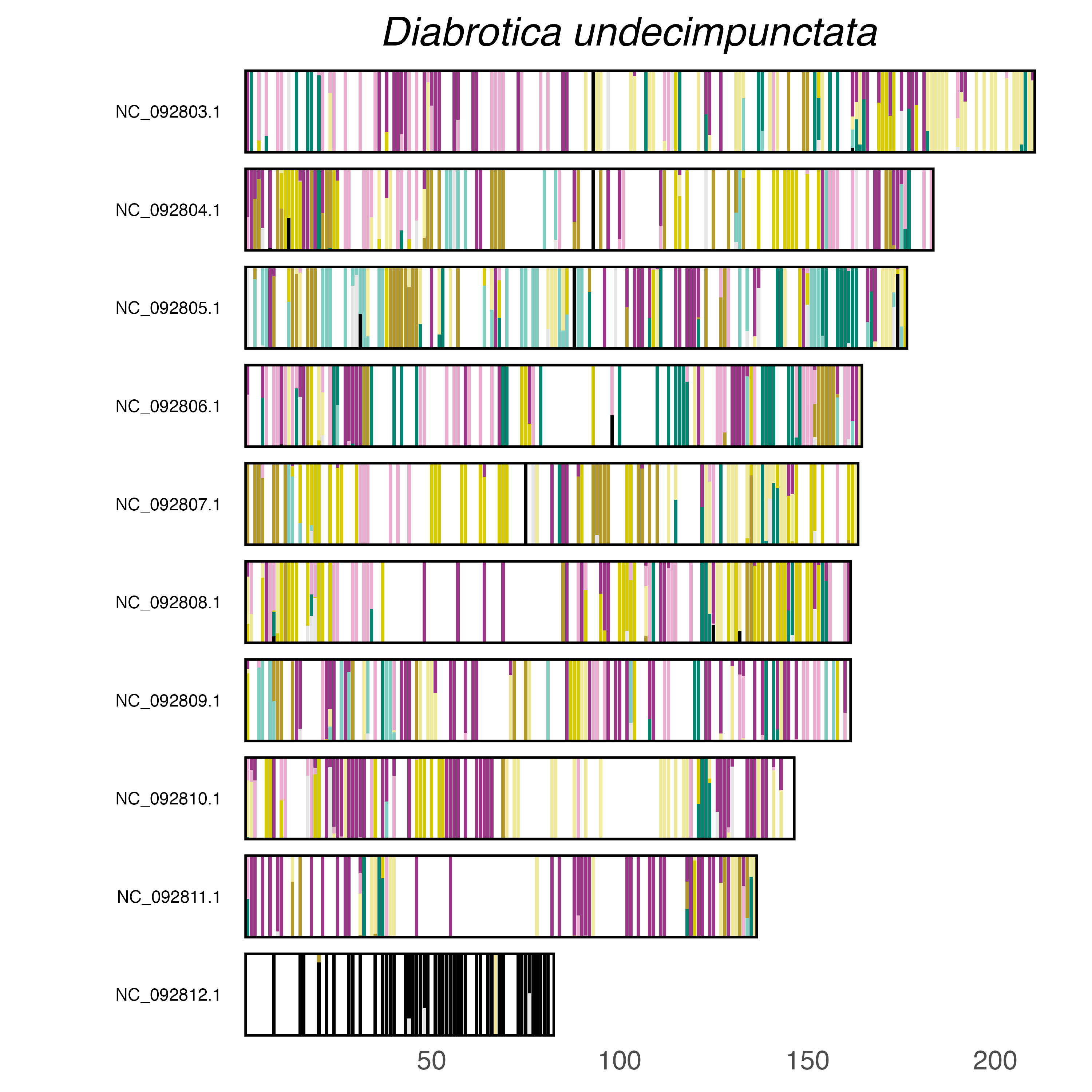

### plot.syngraph_windowbp_1Mb_rel_vertical_chrsize_Diorhabda_sublineata.png

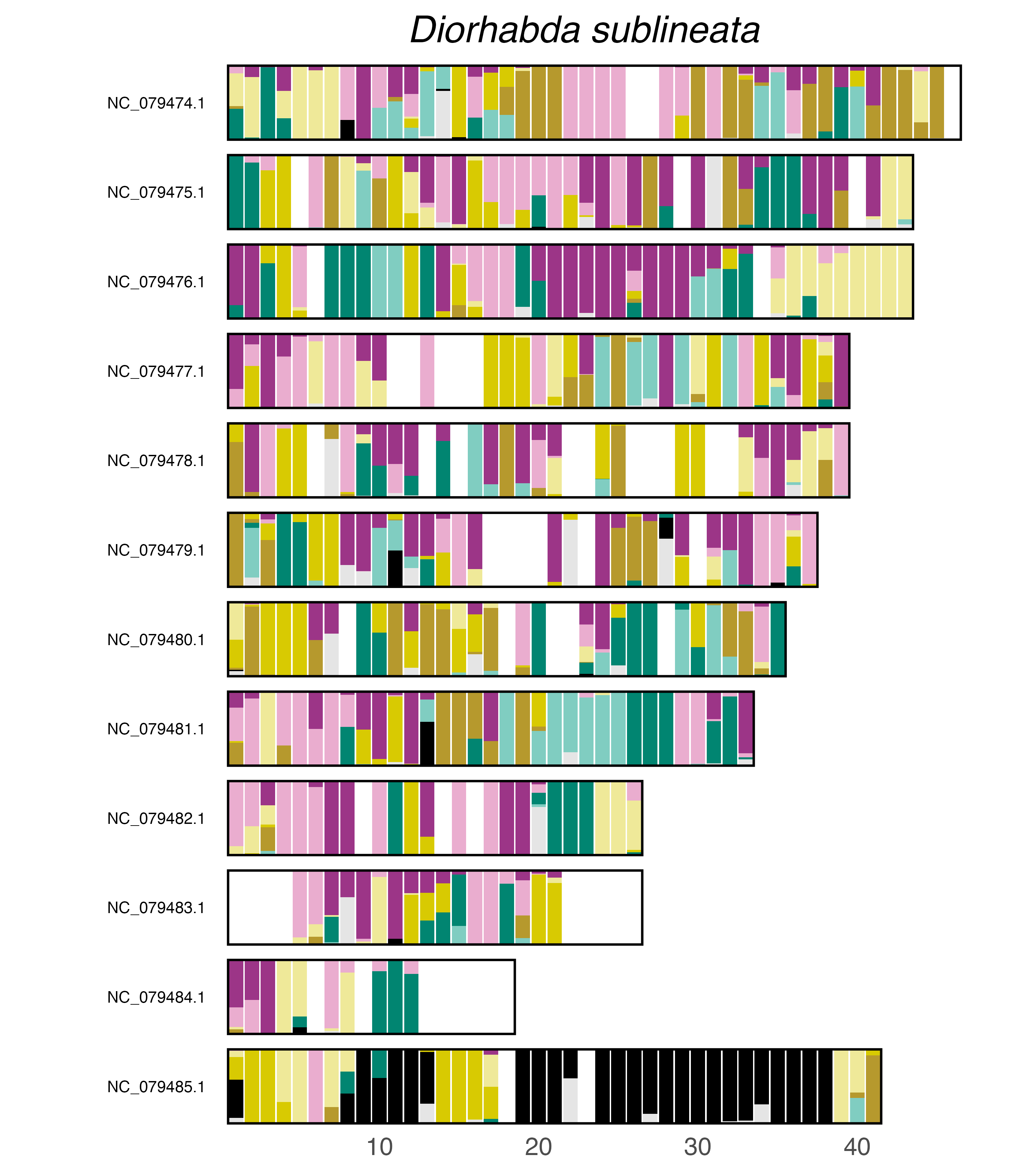

### plot.syngraph_windowbp_1Mb_rel_vertical_chrsize_Dorysthenes_granulosus.png

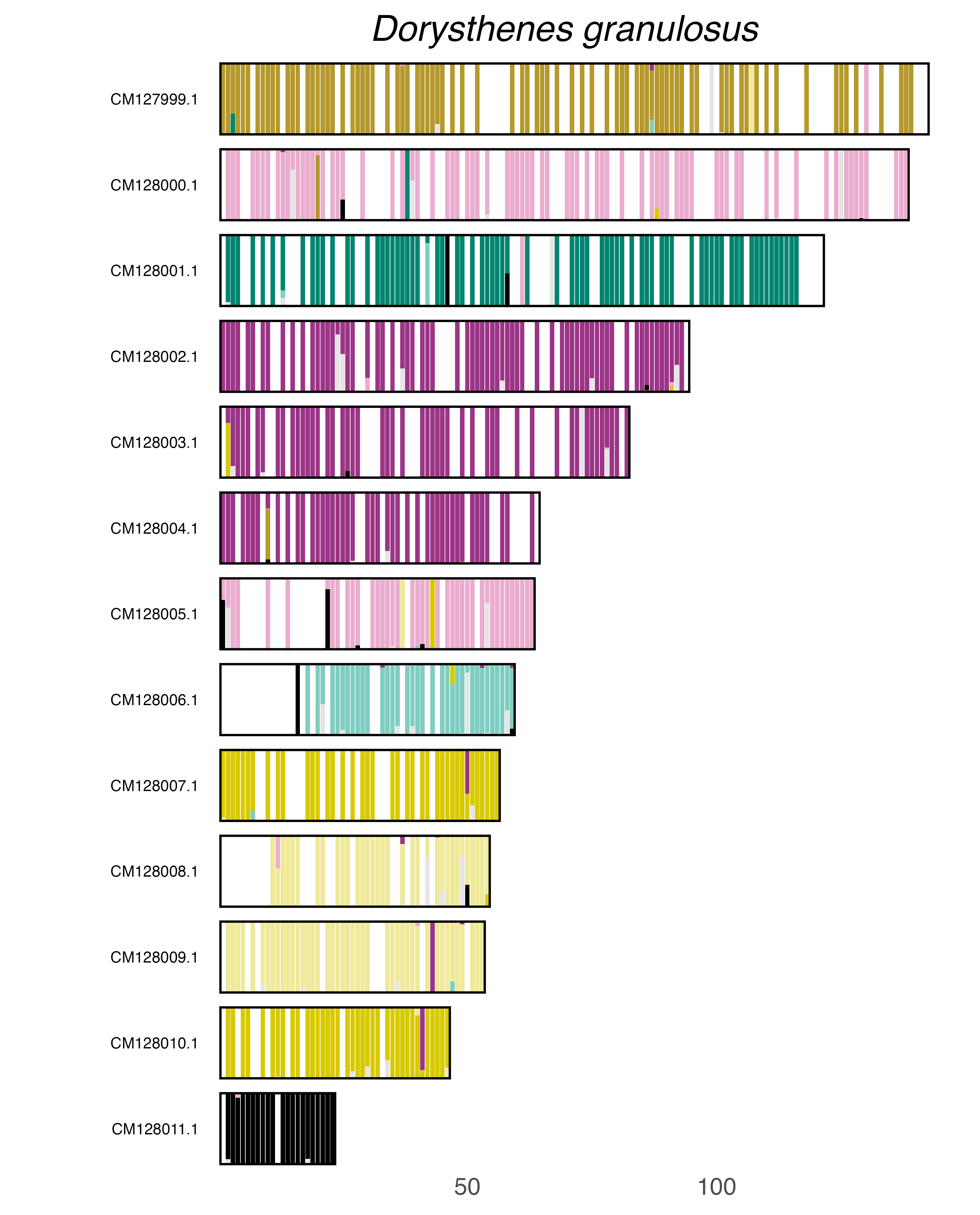

### plot.syngraph_windowbp_1Mb_rel_vertical_chrsize_Drusilla_canaliculata.png

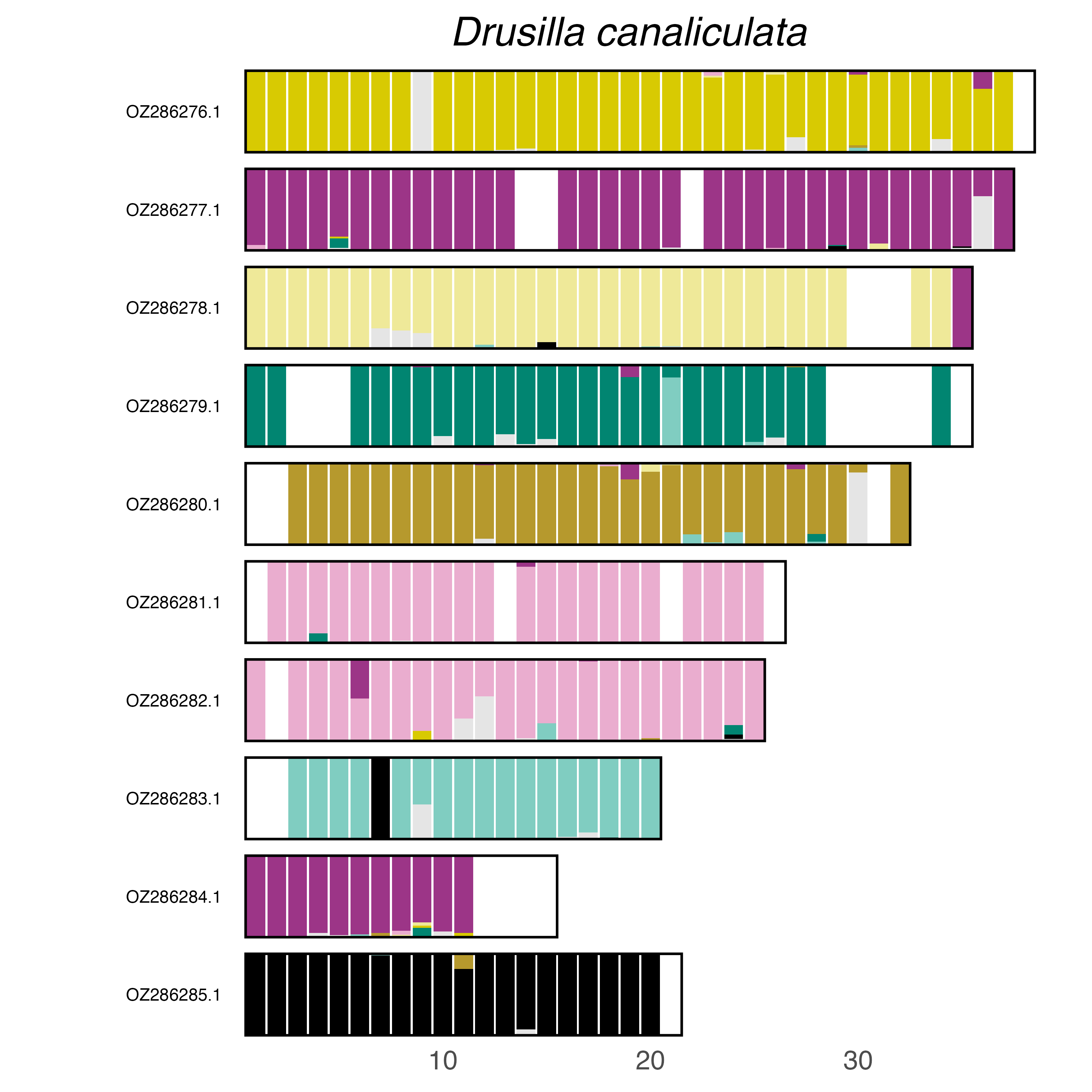

### plot.syngraph_windowbp_1Mb_rel_vertical_chrsize_Eledona_agricola.png

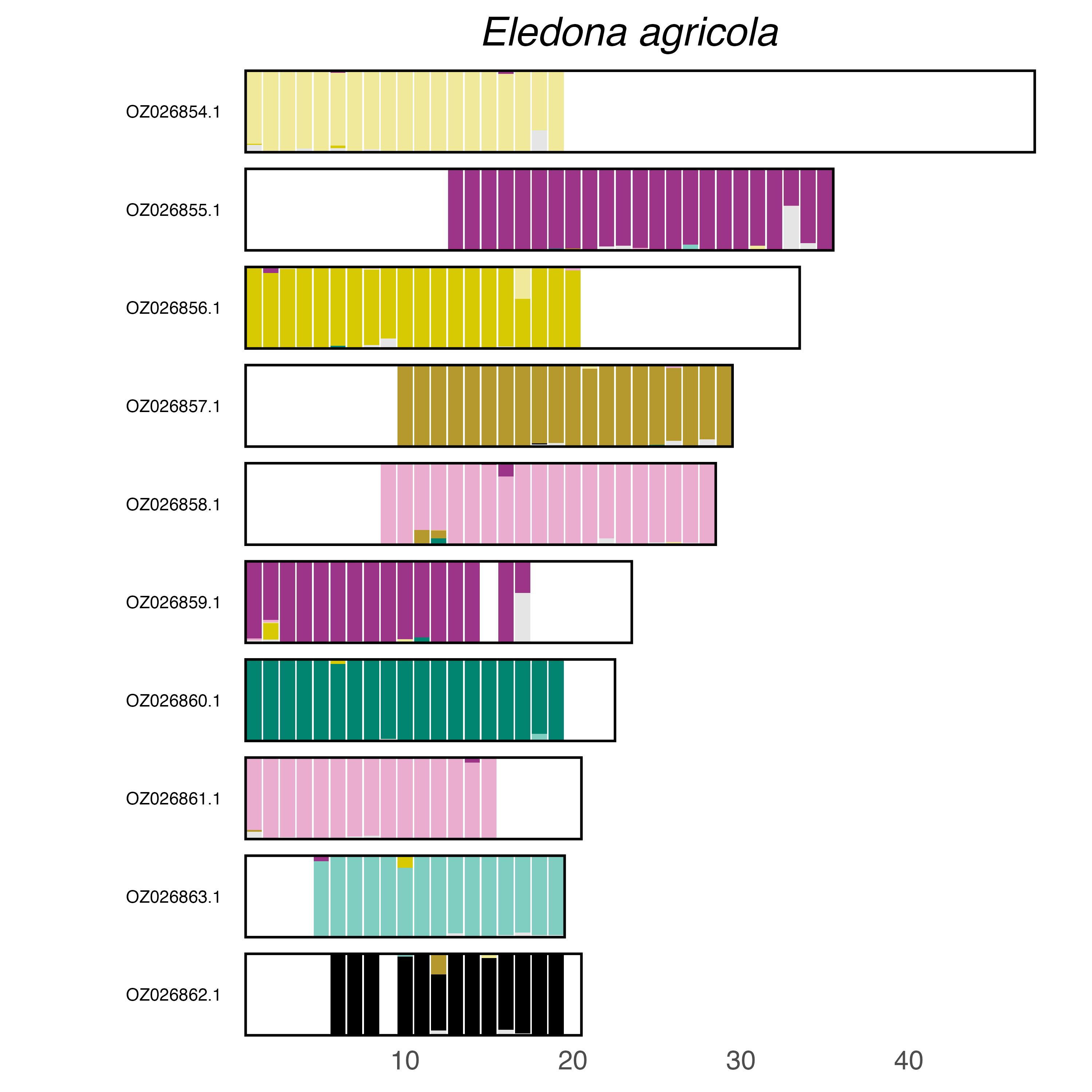

### plot.syngraph_windowbp_1Mb_rel_vertical_chrsize_Eucryptorrhynchus_brandti.png

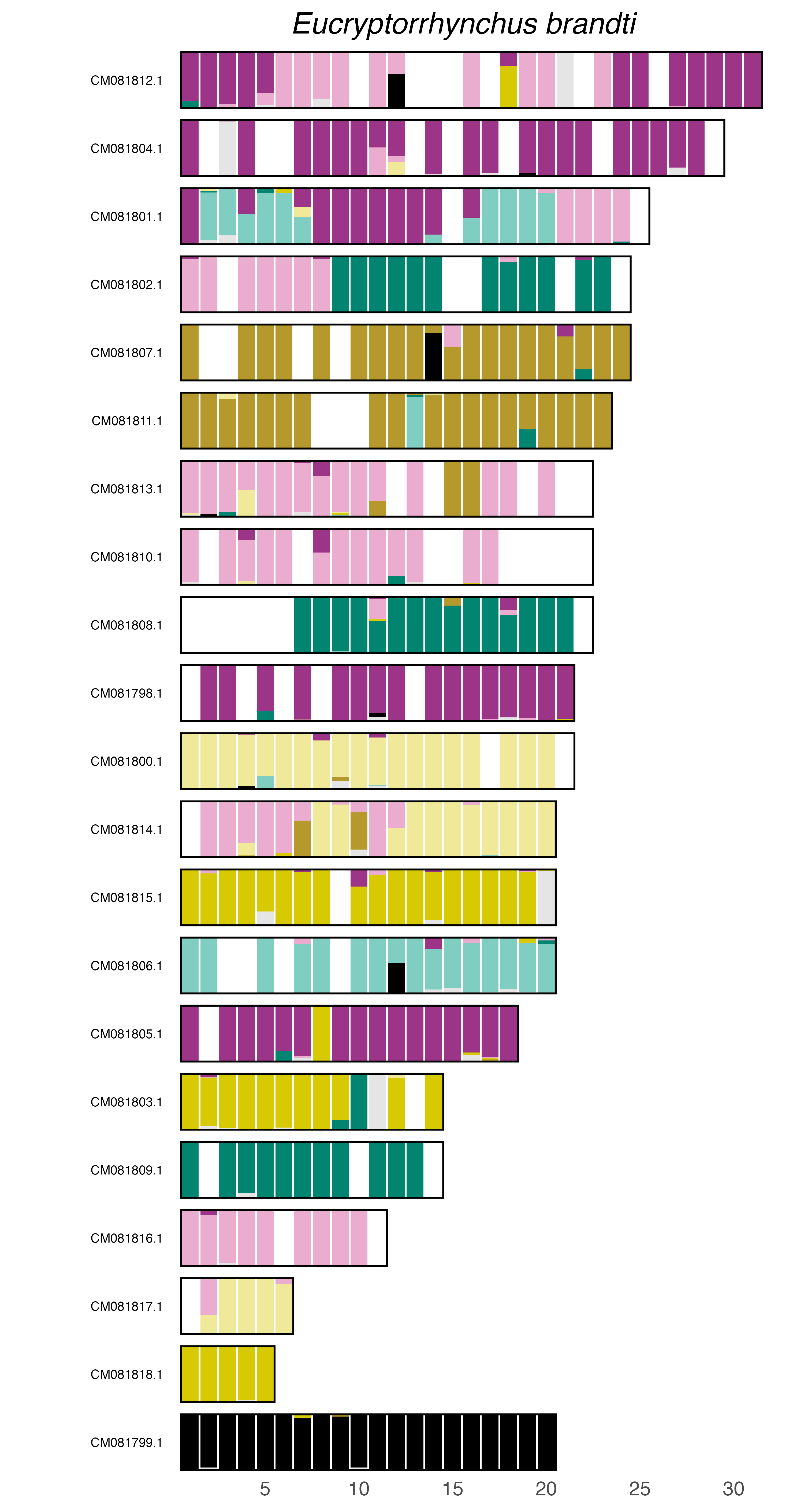

### plot.syngraph_windowbp_1Mb_rel_vertical_chrsize_Euwallacea_similis.png

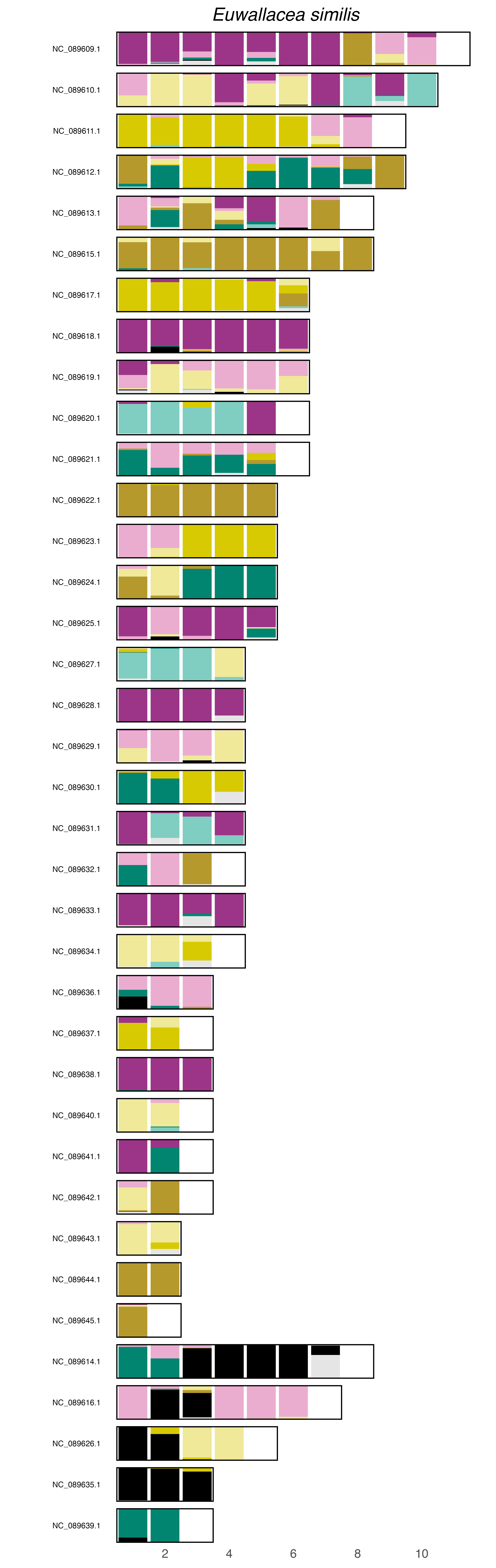

### plot.syngraph_windowbp_1Mb_rel_vertical_chrsize_Galeruca_laticollis.png

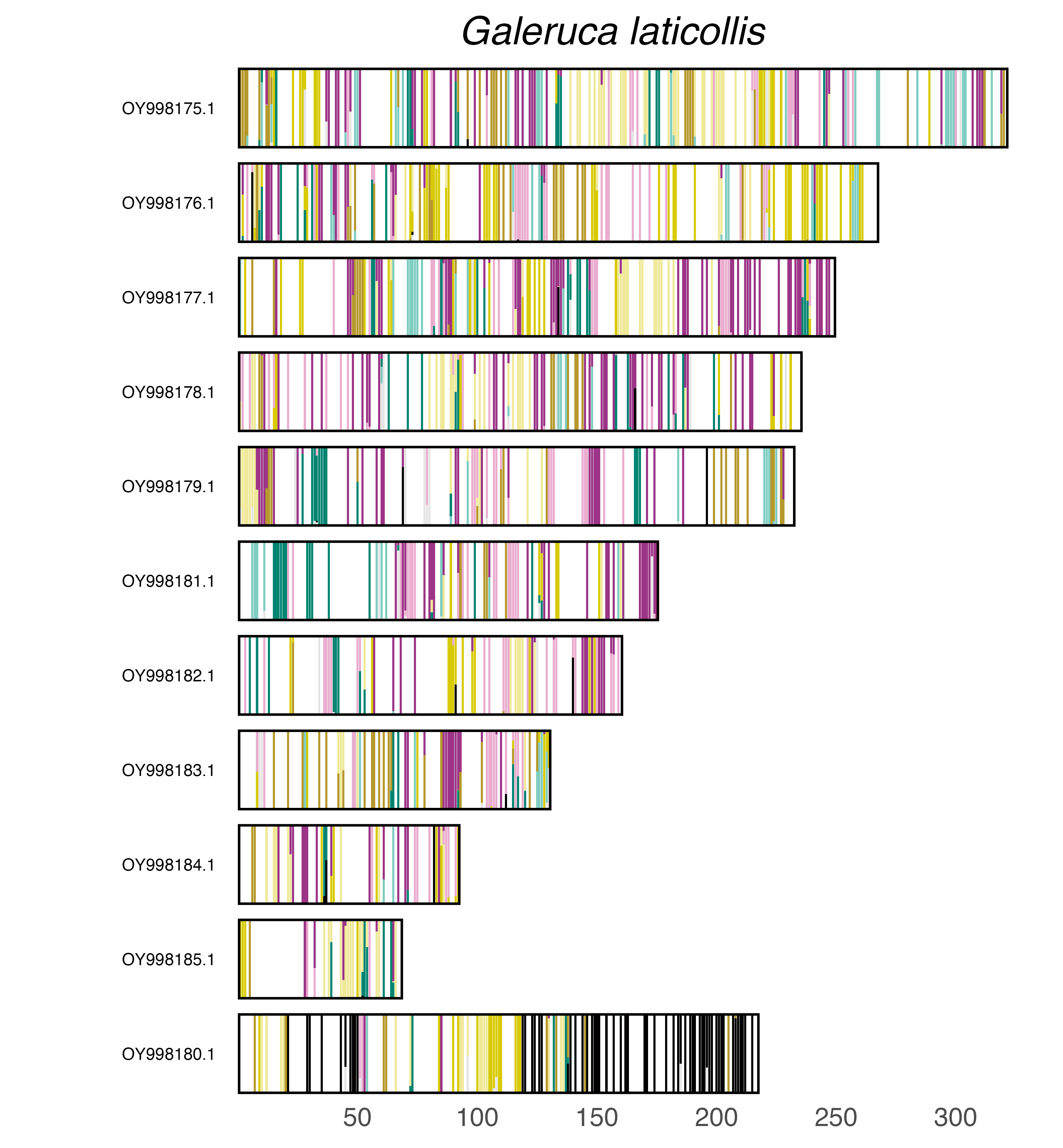

### plot.syngraph_windowbp_1Mb_rel_vertical_chrsize_Geotrupes_spiniger.png

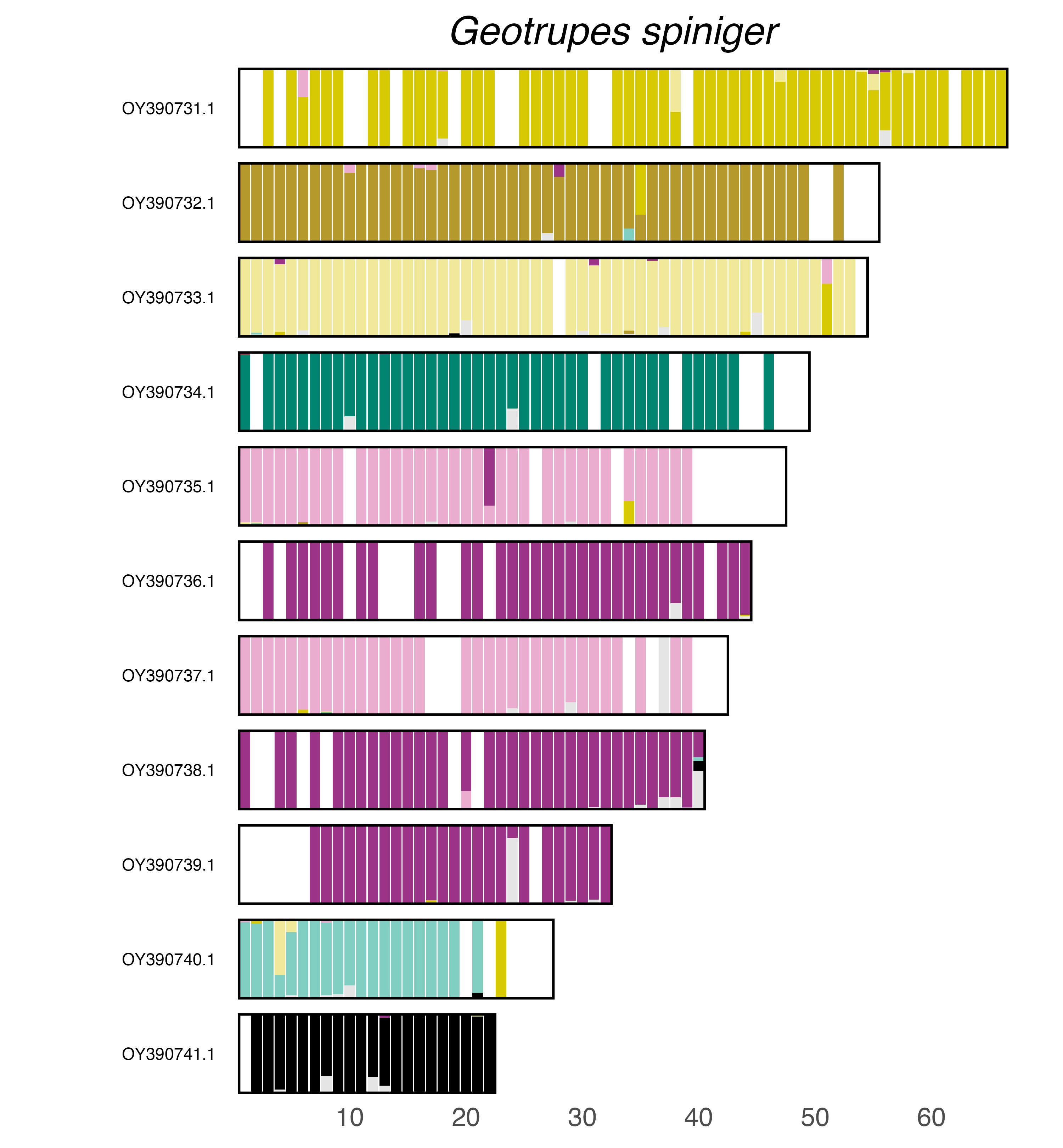

### plot.syngraph_windowbp_1Mb_rel_vertical_chrsize_Gonodera_luperus.png

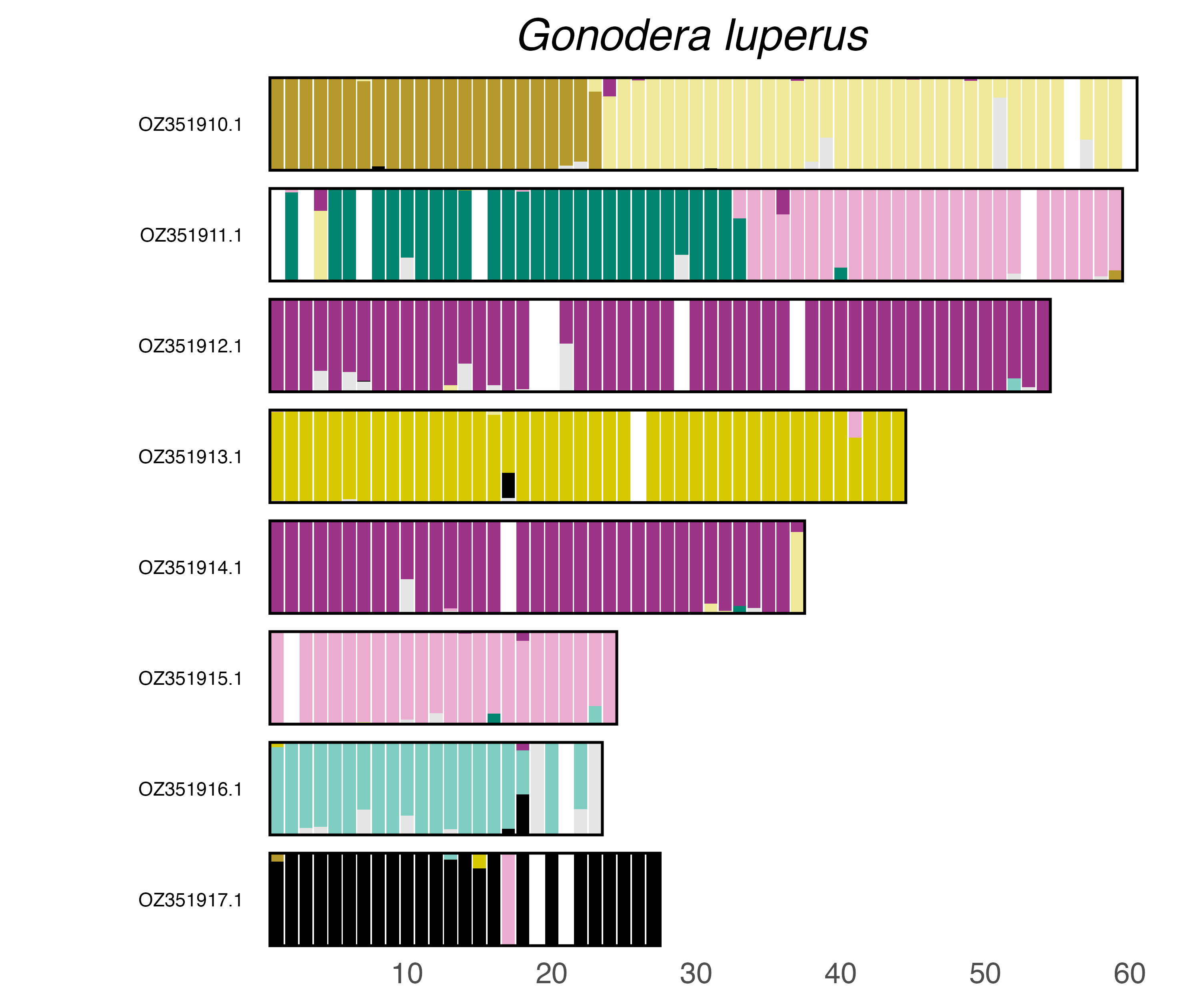

### plot.syngraph_windowbp_1Mb_rel_vertical_chrsize_Harpalus_anxius.png

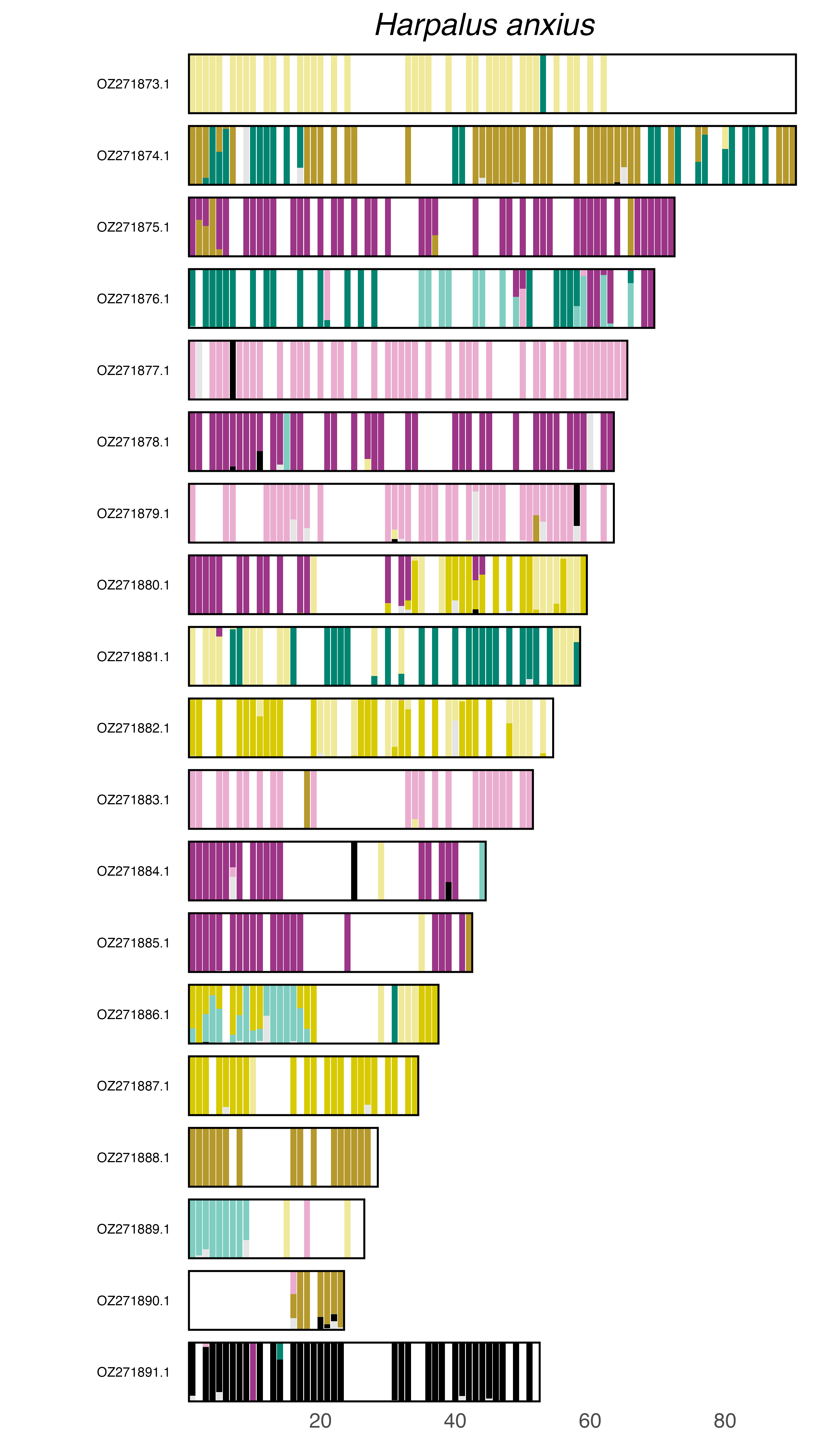

### plot.syngraph_windowbp_1Mb_rel_vertical_chrsize_Harpalus_rufipes.png

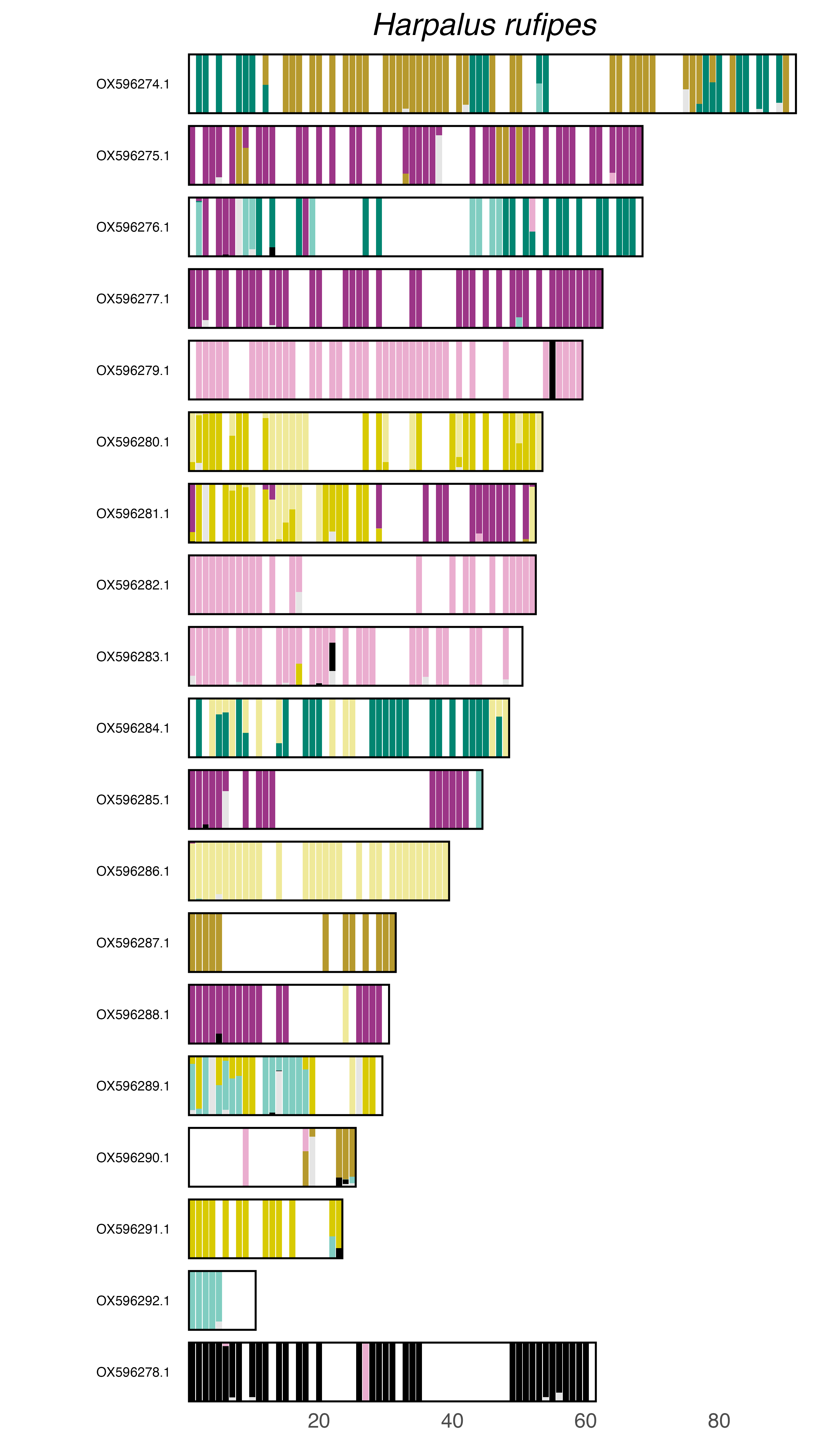

### plot.syngraph_windowbp_1Mb_rel_vertical_chrsize_Hippodamia_variegata.png

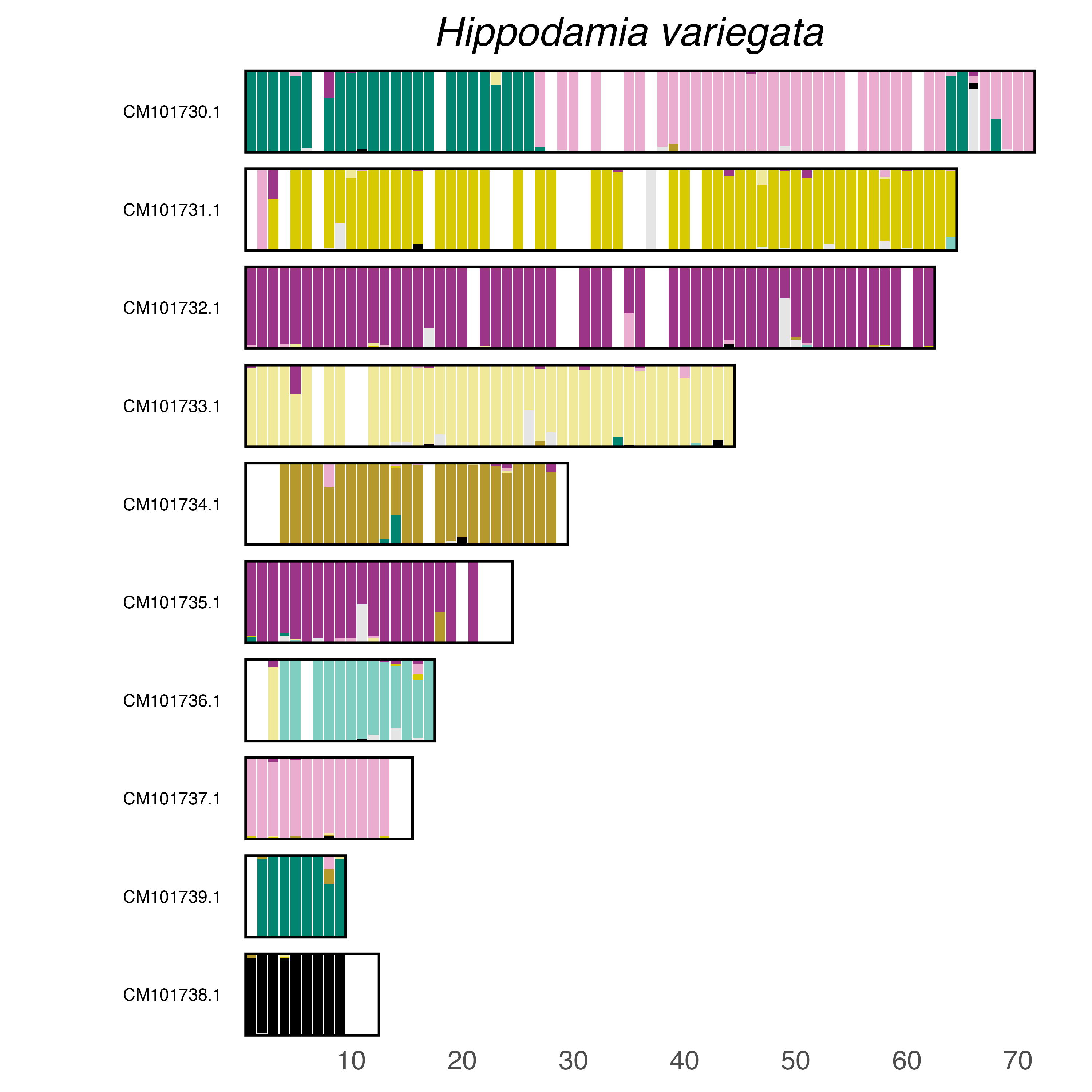

### plot.syngraph_windowbp_1Mb_rel_vertical_chrsize_Hololepta_plana.png

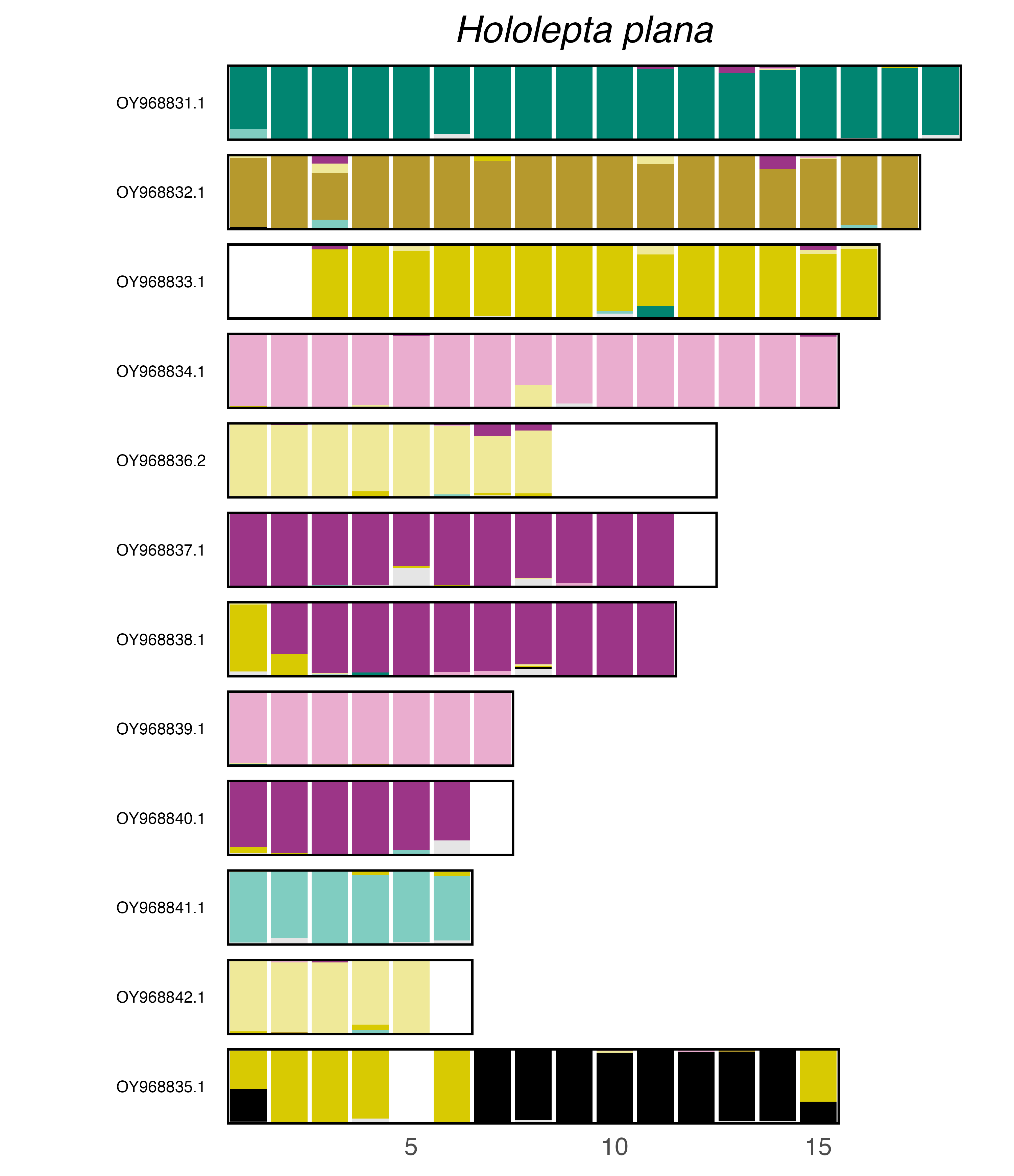

### plot.syngraph_windowbp_1Mb_rel_vertical_chrsize_Holotrichia_oblita.png

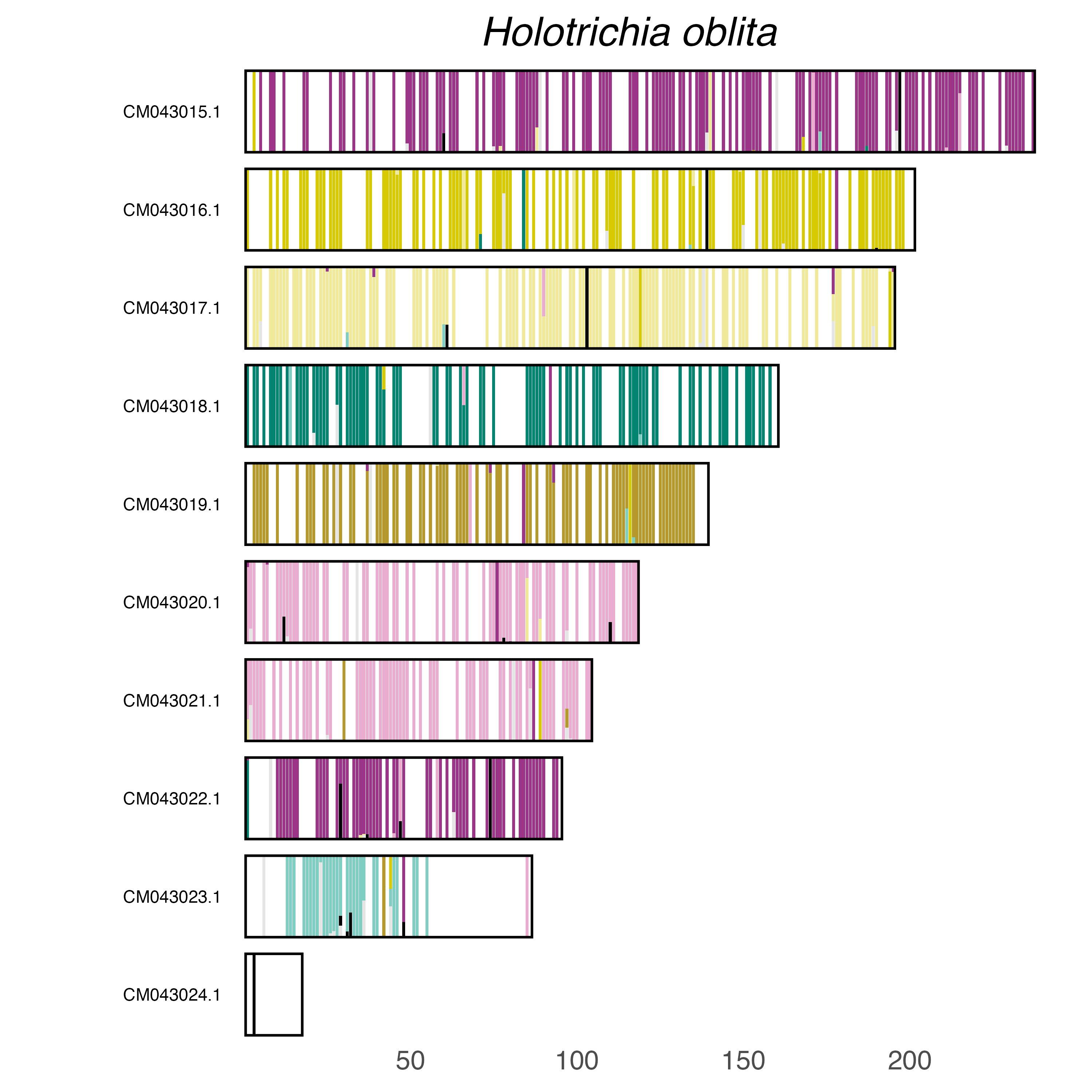

### plot.syngraph_windowbp_1Mb_rel_vertical_chrsize_Hycleus_marcipoli.png

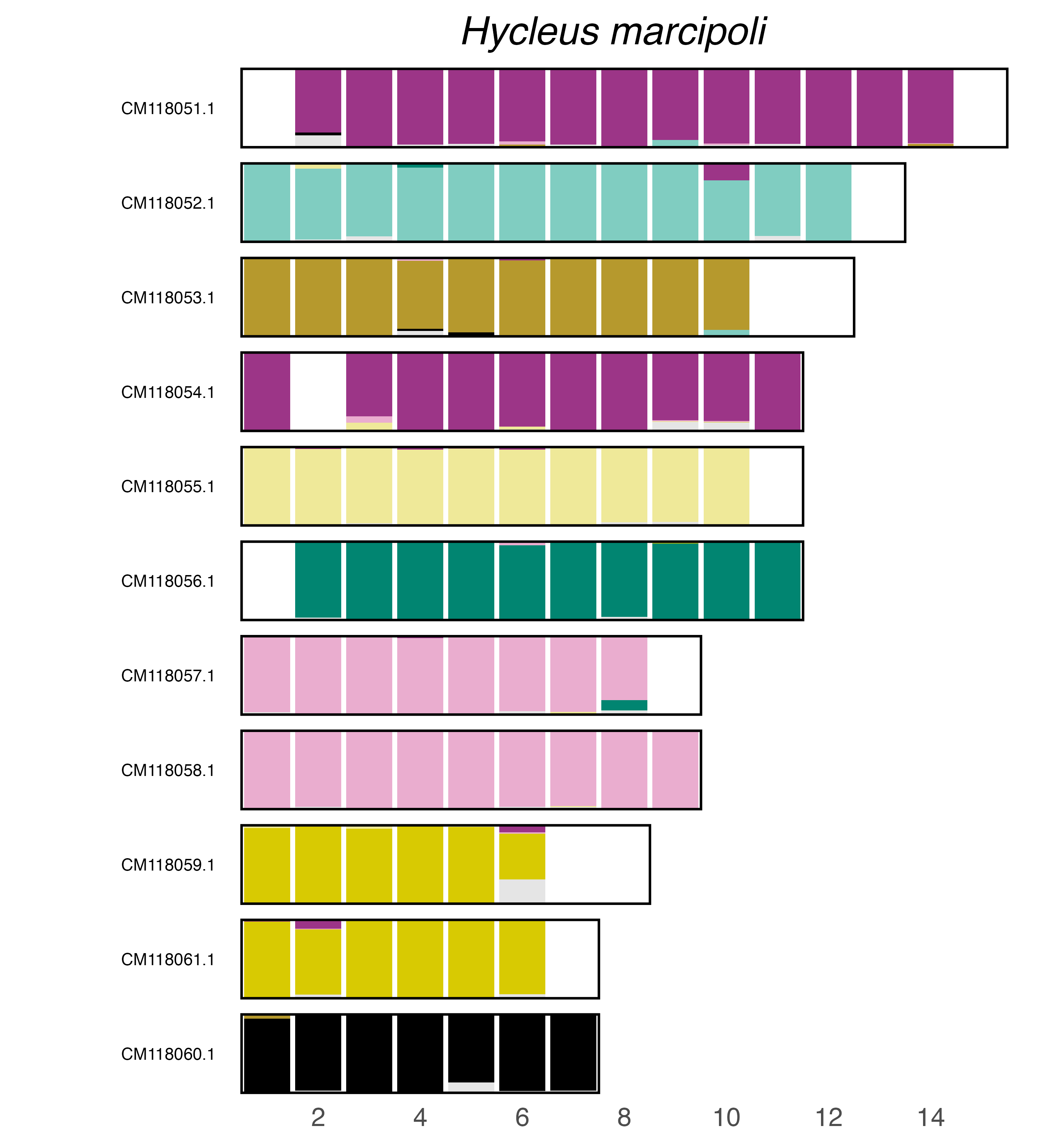

### plot.syngraph_windowbp_1Mb_rel_vertical_chrsize_Ichthyurus_bourgeoisi.png

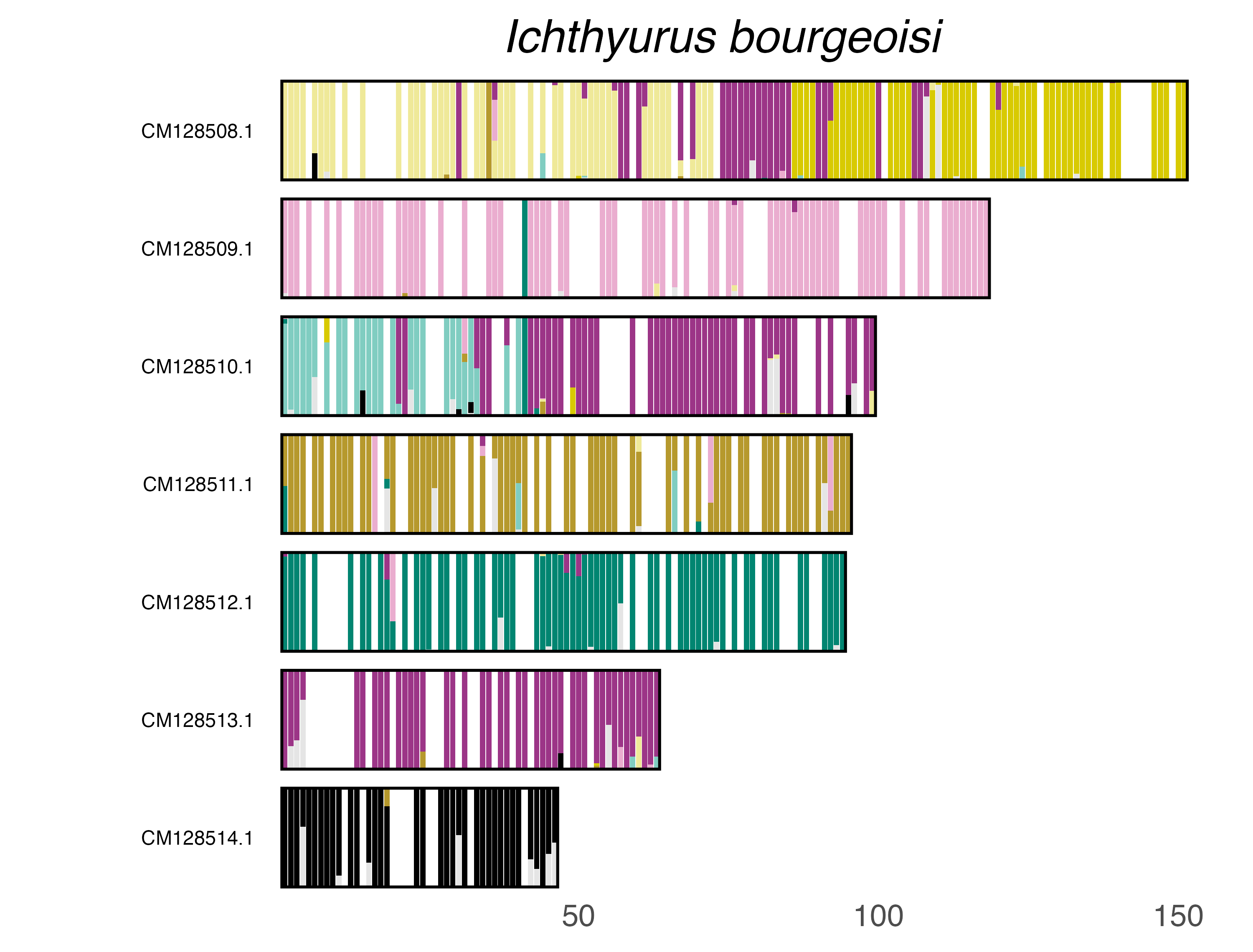

### plot.syngraph_windowbp_1Mb_rel_vertical_chrsize_Lagria_hirta.png

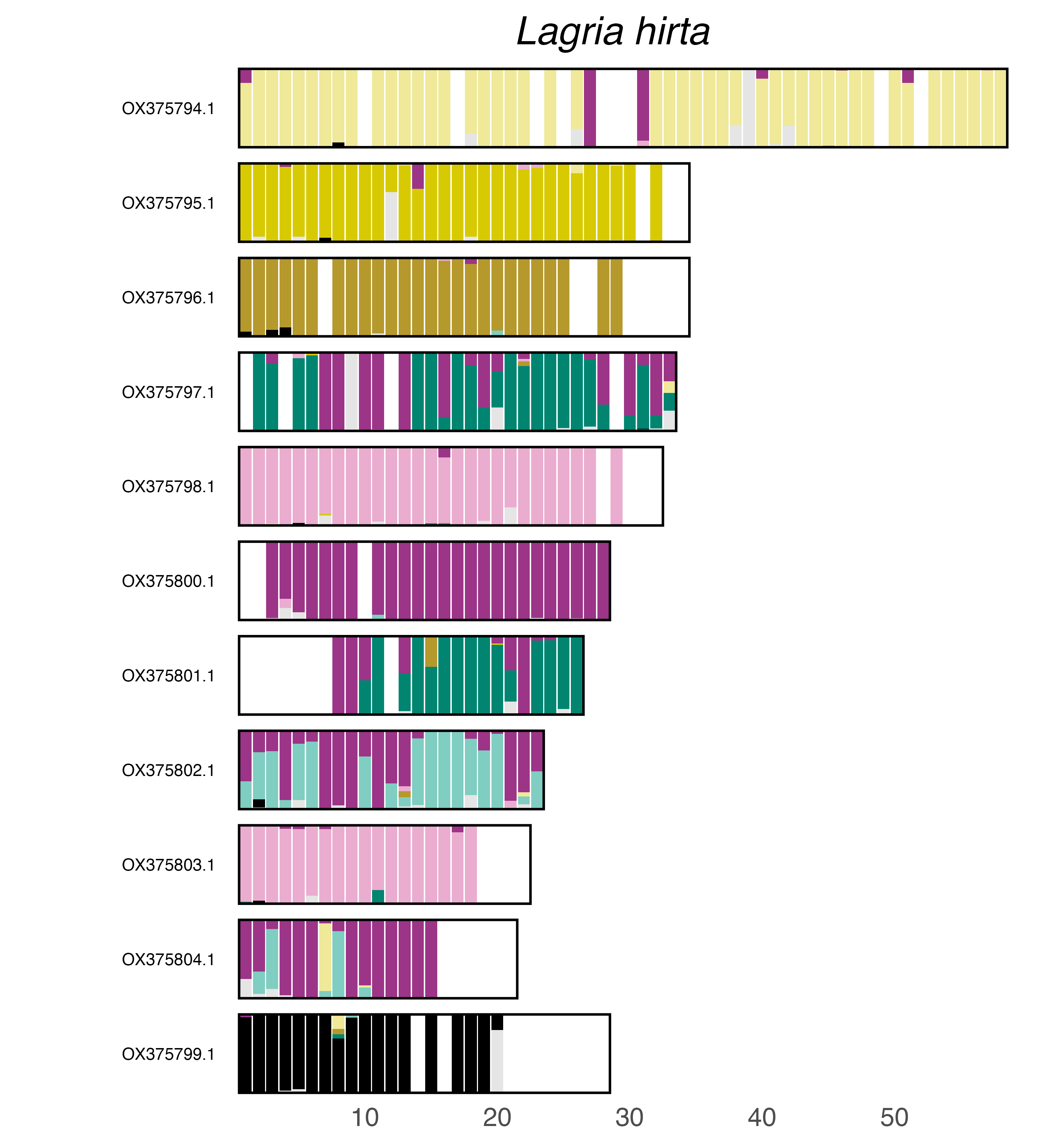

### plot.syngraph_windowbp_1Mb_rel_vertical_chrsize_Latheticus_oryzae.png

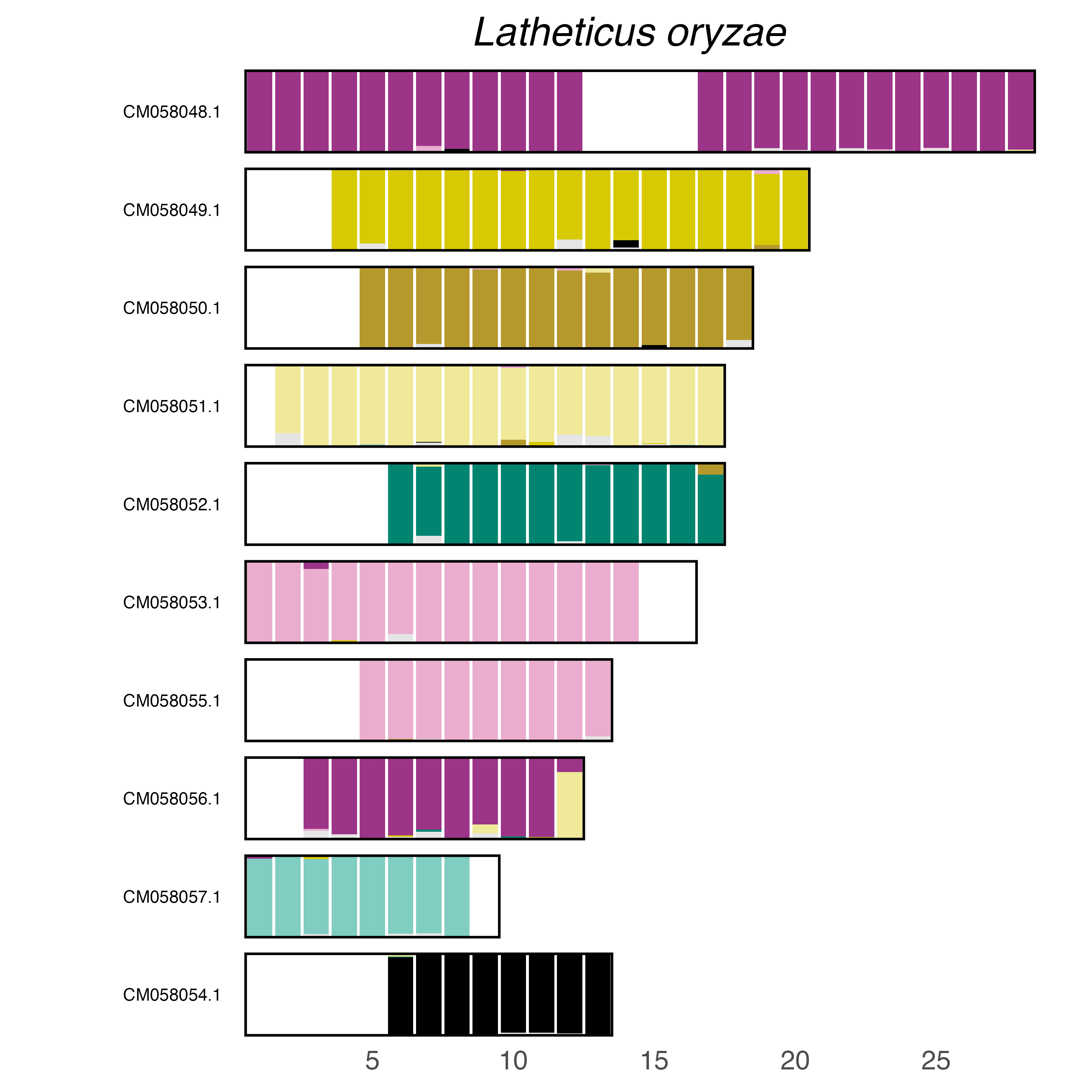

### plot.syngraph_windowbp_1Mb_rel_vertical_chrsize_Leistus_rufomarginatus.png

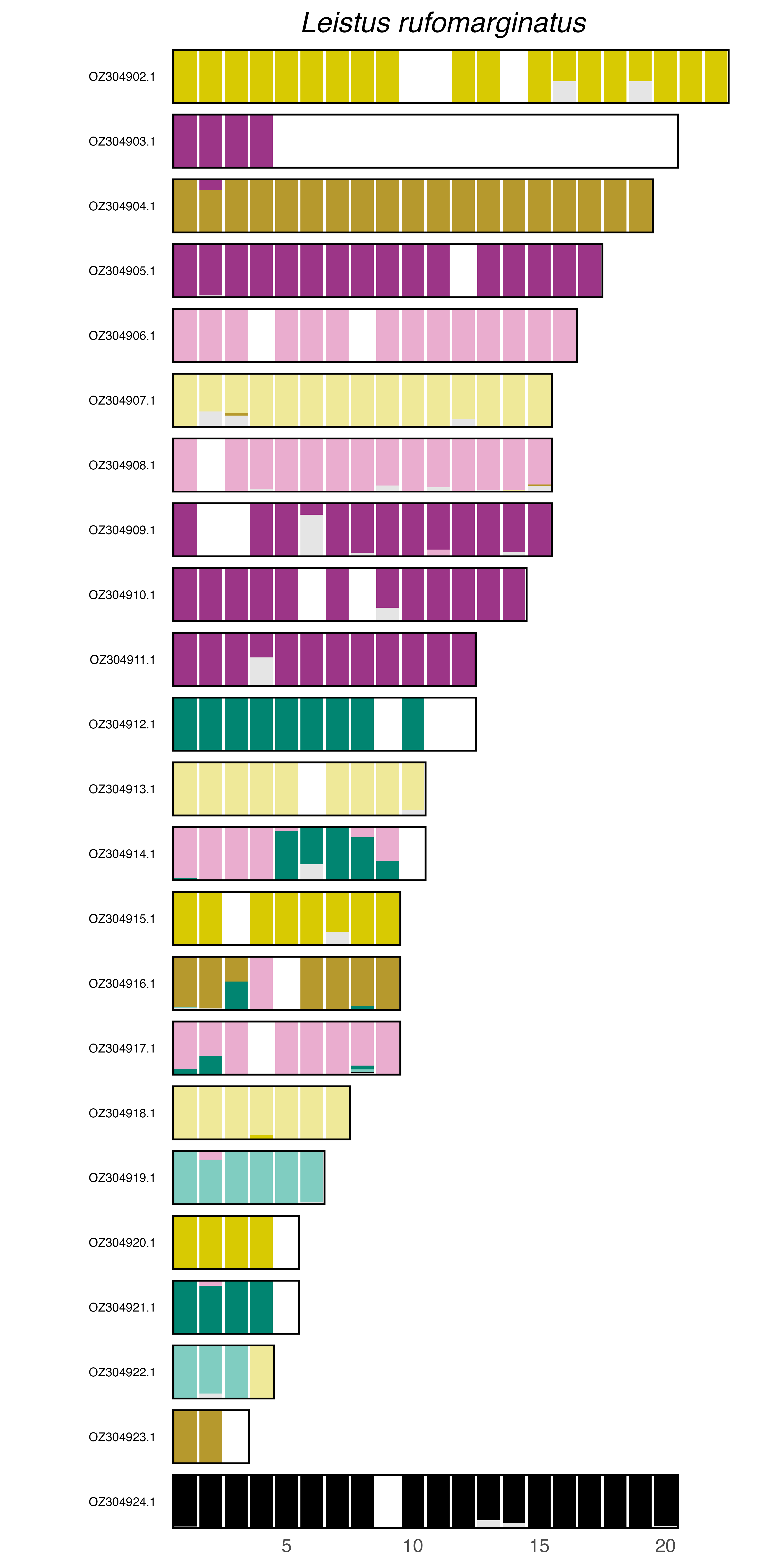

### plot.syngraph_windowbp_1Mb_rel_vertical_chrsize_Leonhardella_angulicollis.png

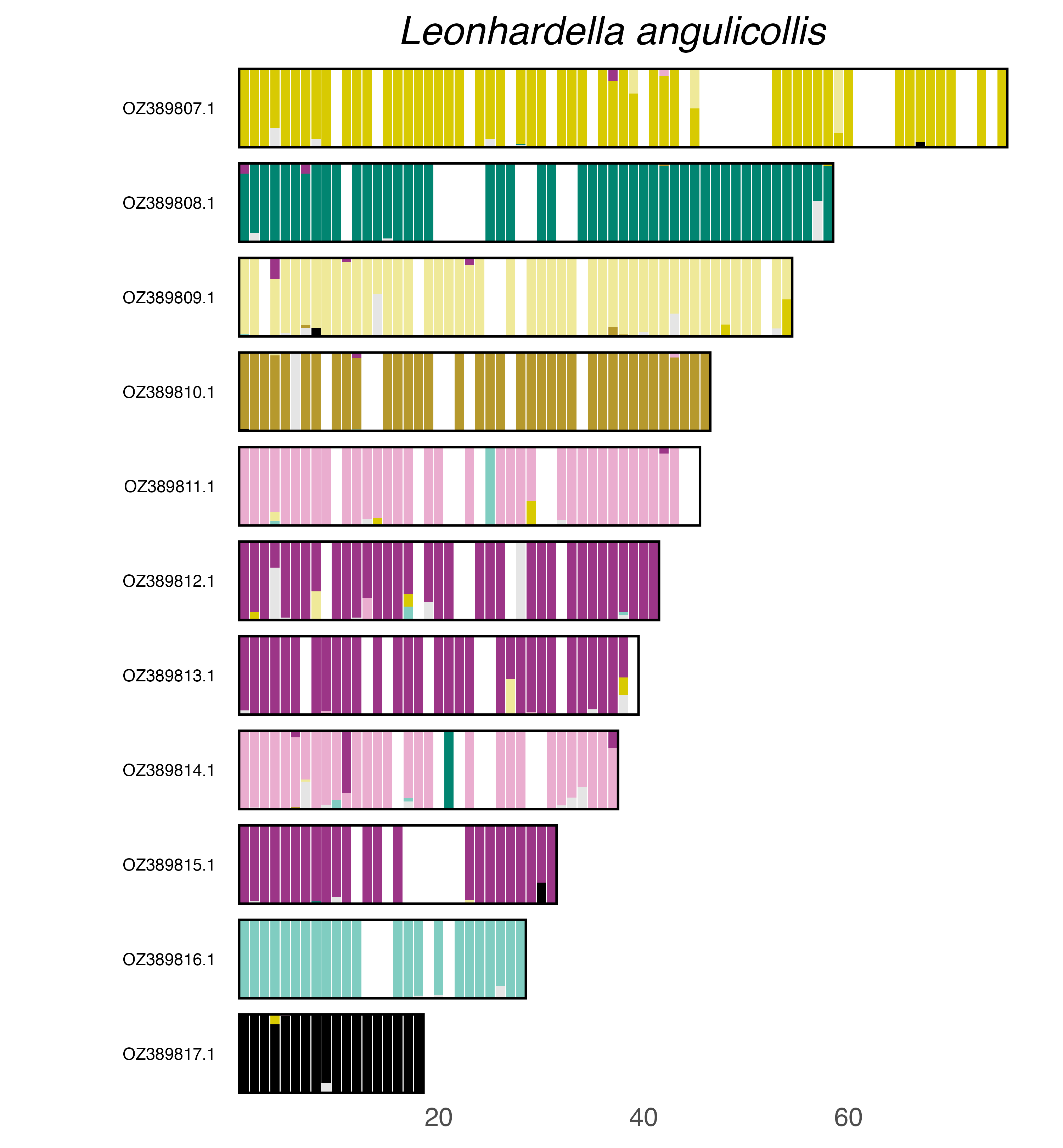

### plot.syngraph_windowbp_1Mb_rel_vertical_chrsize_Limonius_poneli.png

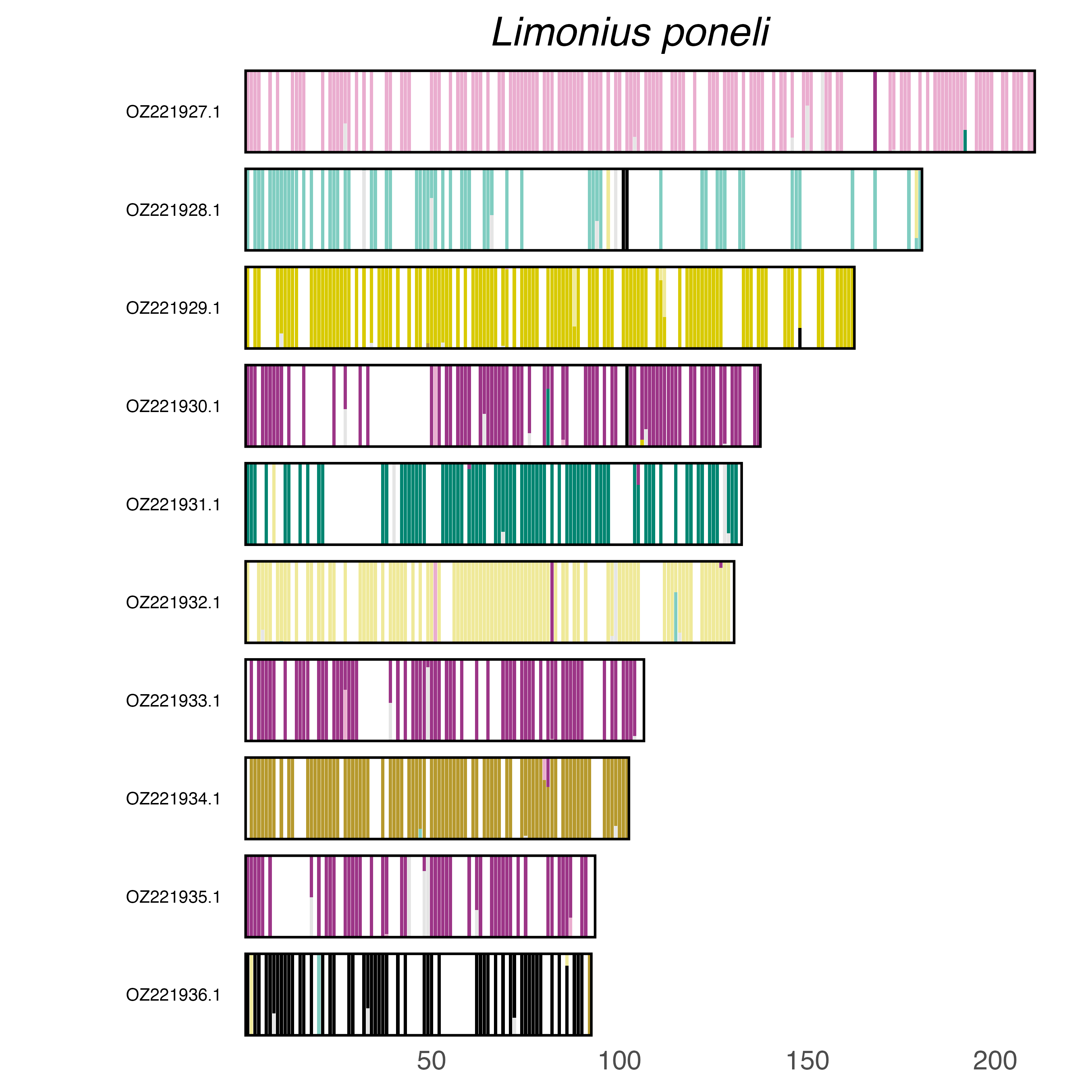

### plot.syngraph_windowbp_1Mb_rel_vertical_chrsize_Liopterus_haemorrhoidalis.png

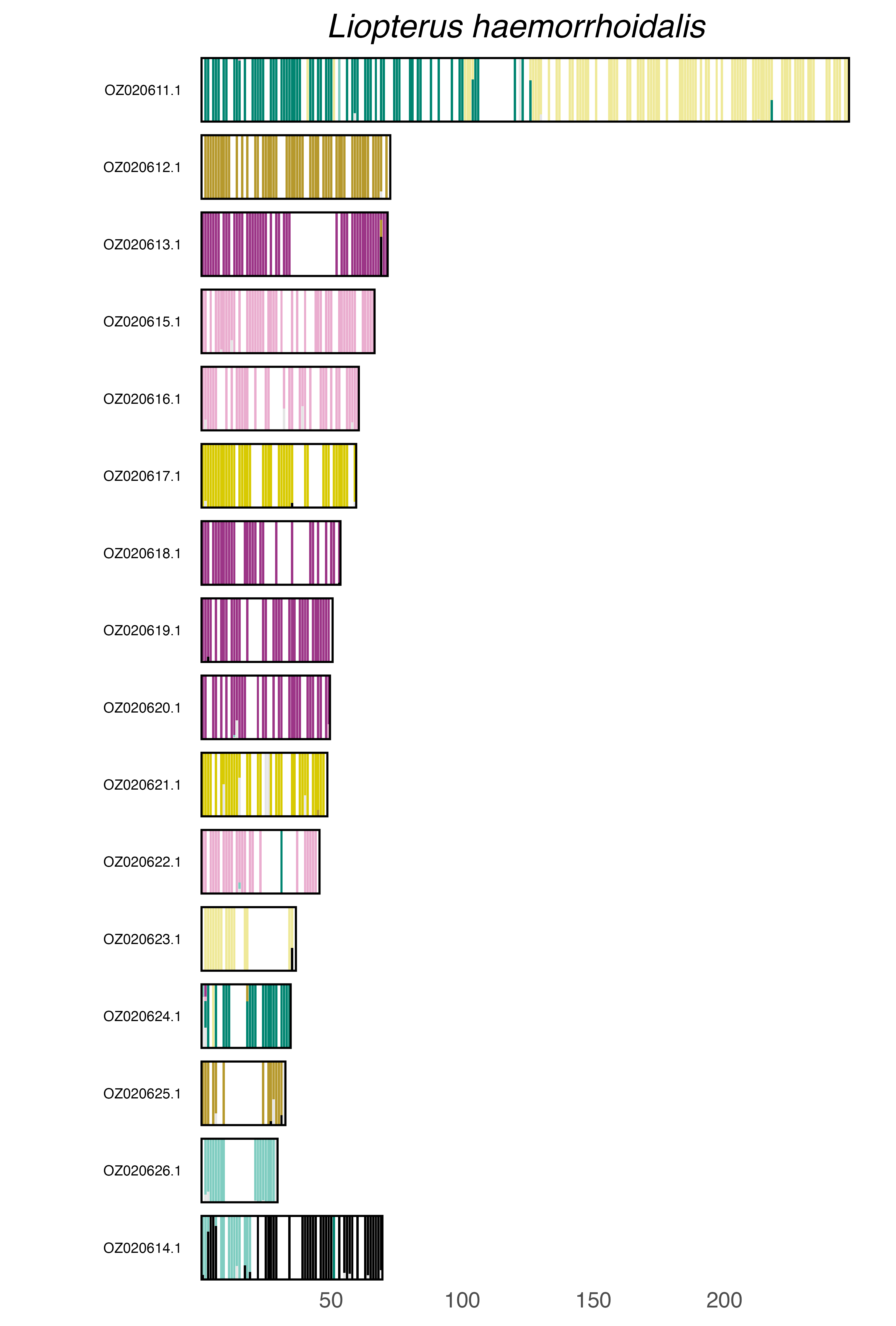

### plot.syngraph_windowbp_1Mb_rel_vertical_chrsize_Lochmaea_crataegi.png

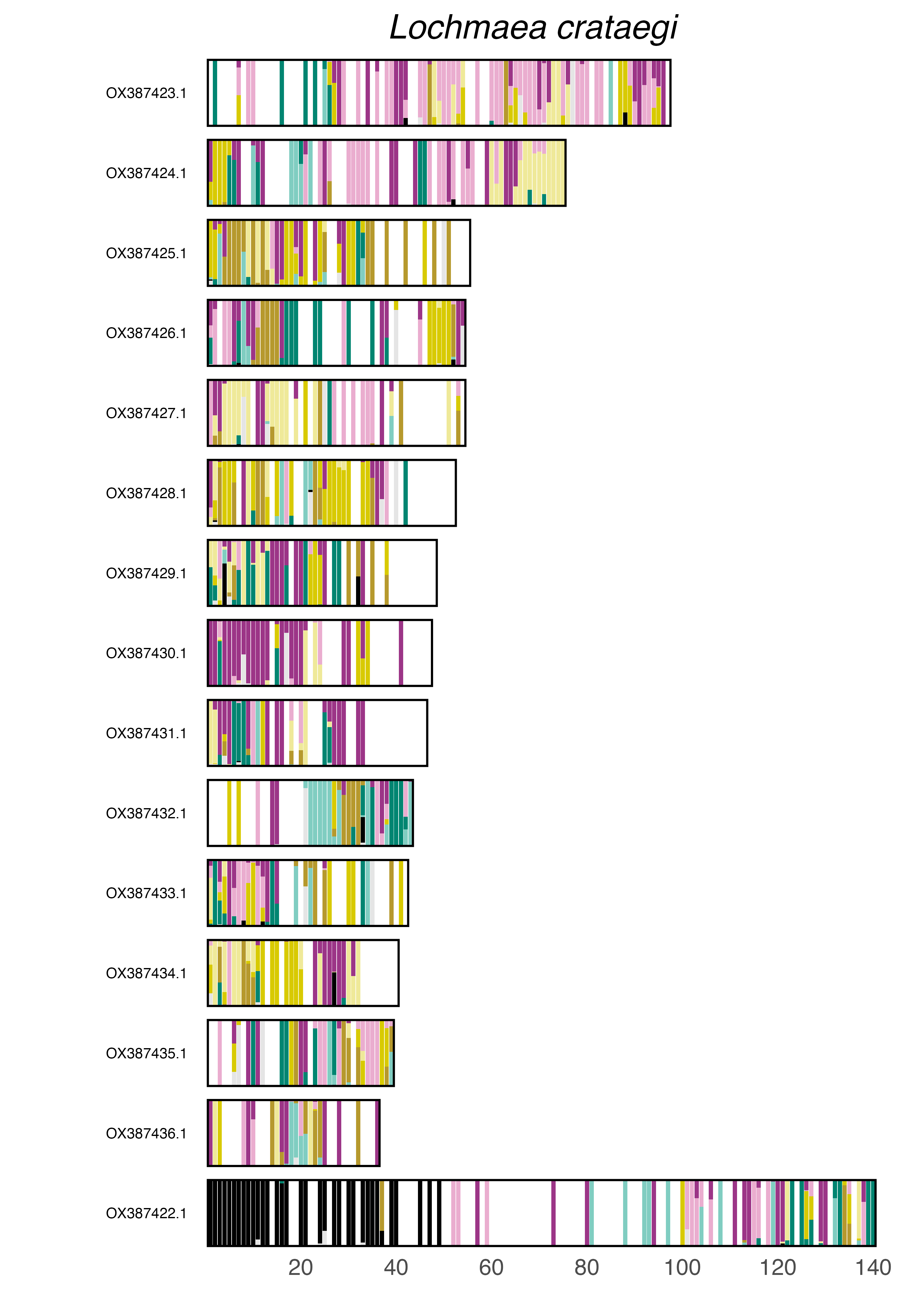

### plot.syngraph_windowbp_1Mb_rel_vertical_chrsize_Luciola_italica.png

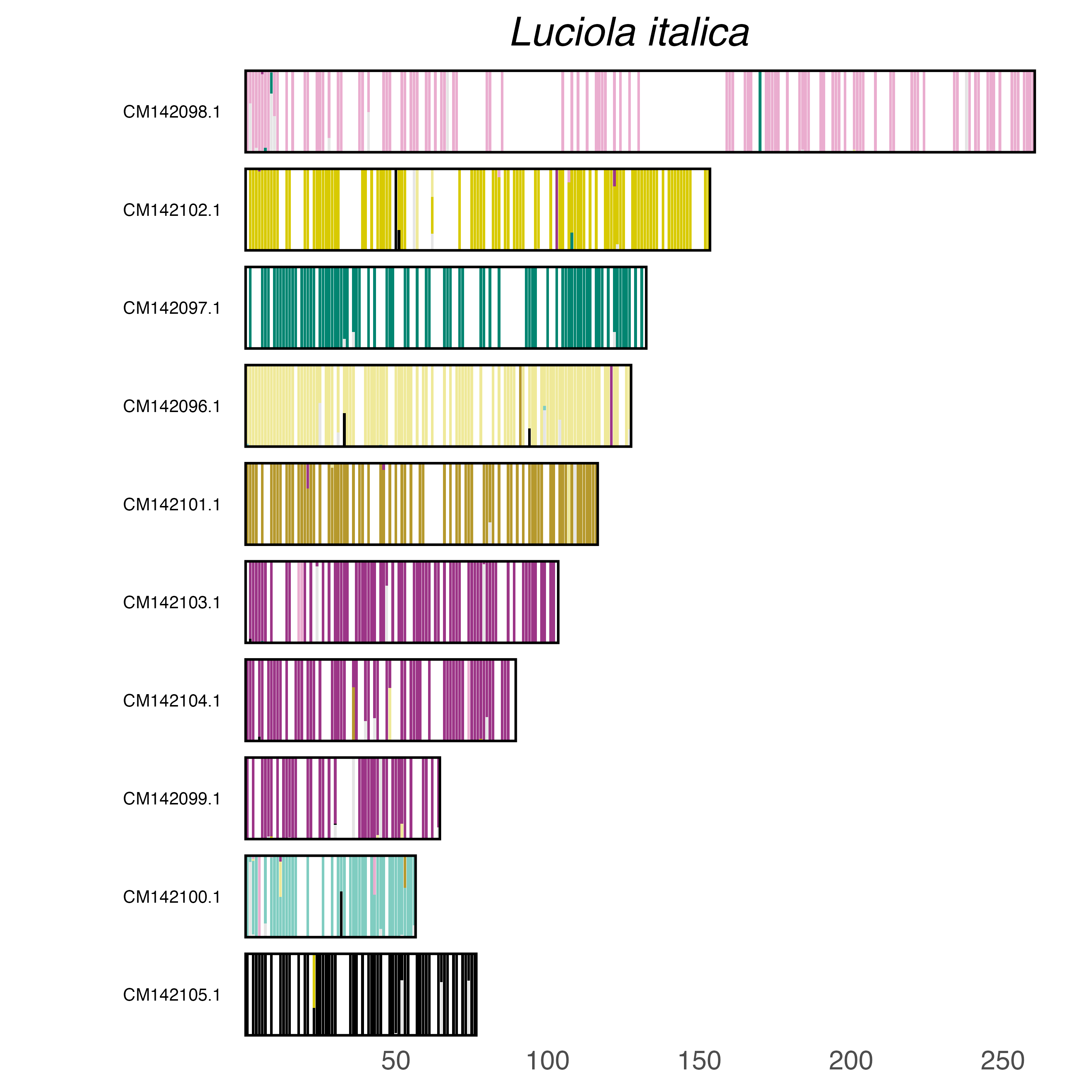

### plot.syngraph_windowbp_1Mb_rel_vertical_chrsize_Melanotus_villosus.png

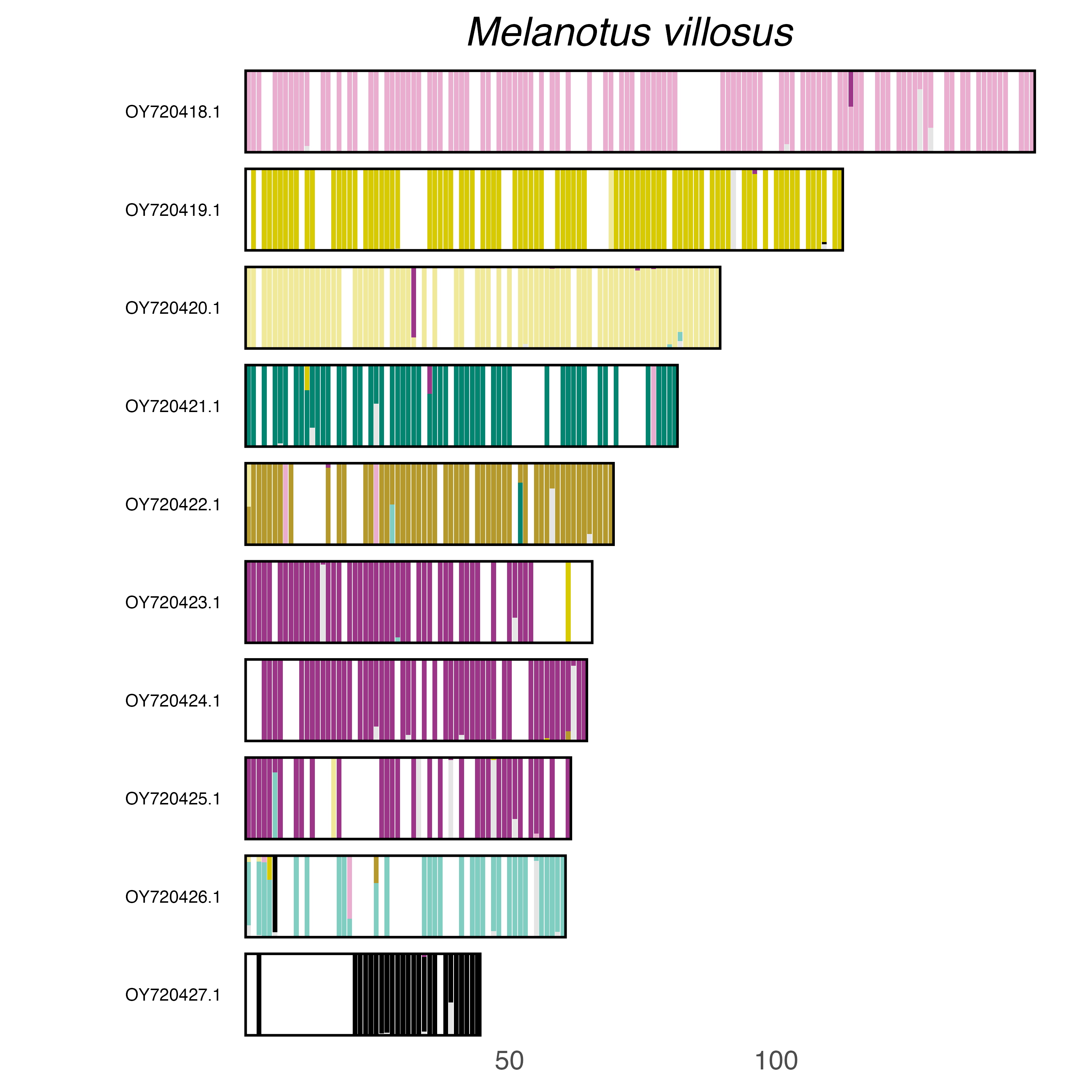

### plot.syngraph_windowbp_1Mb_rel_vertical_chrsize_Melinopterus_prodromus.png

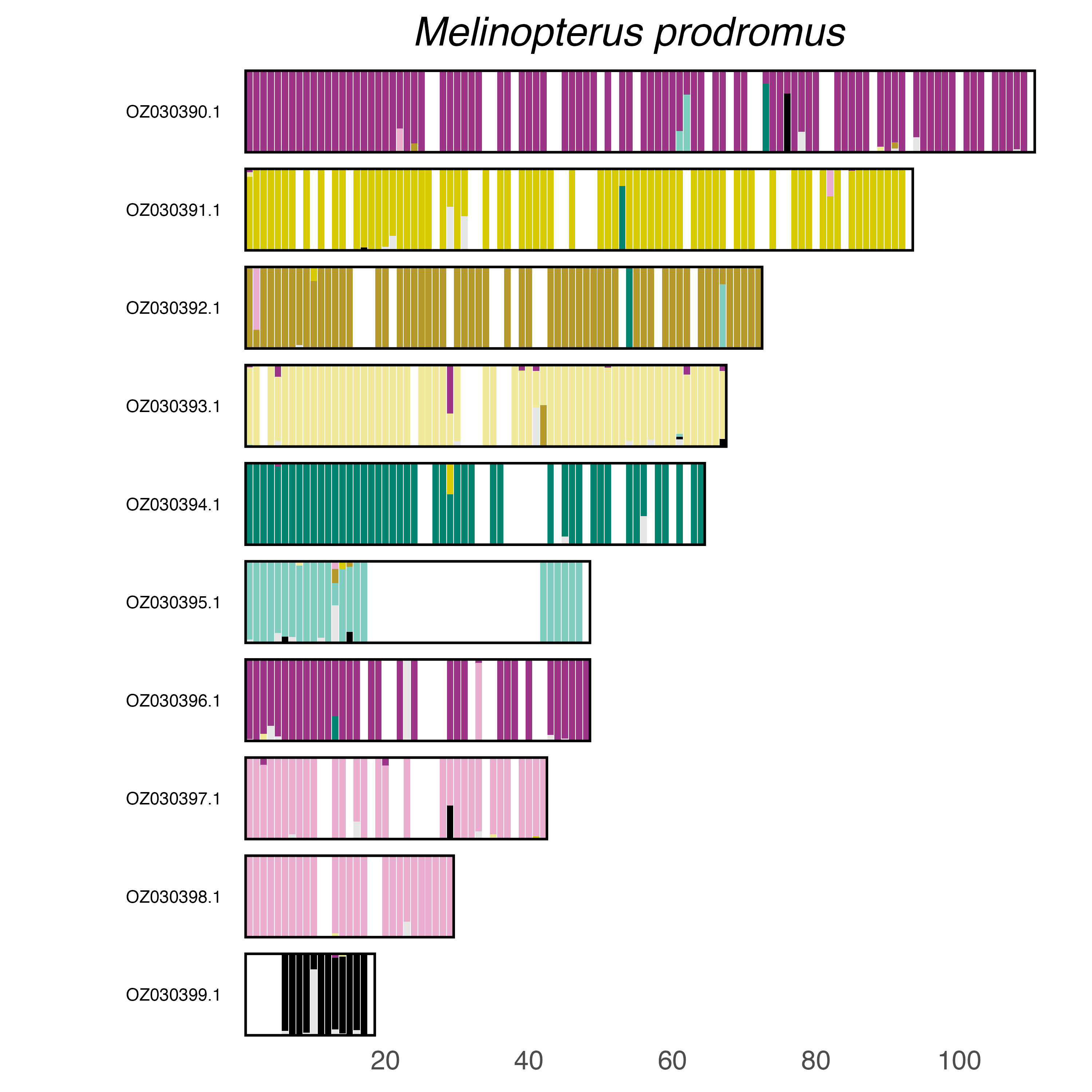

### plot.syngraph_windowbp_1Mb_rel_vertical_chrsize_Melolontha_melolontha.png

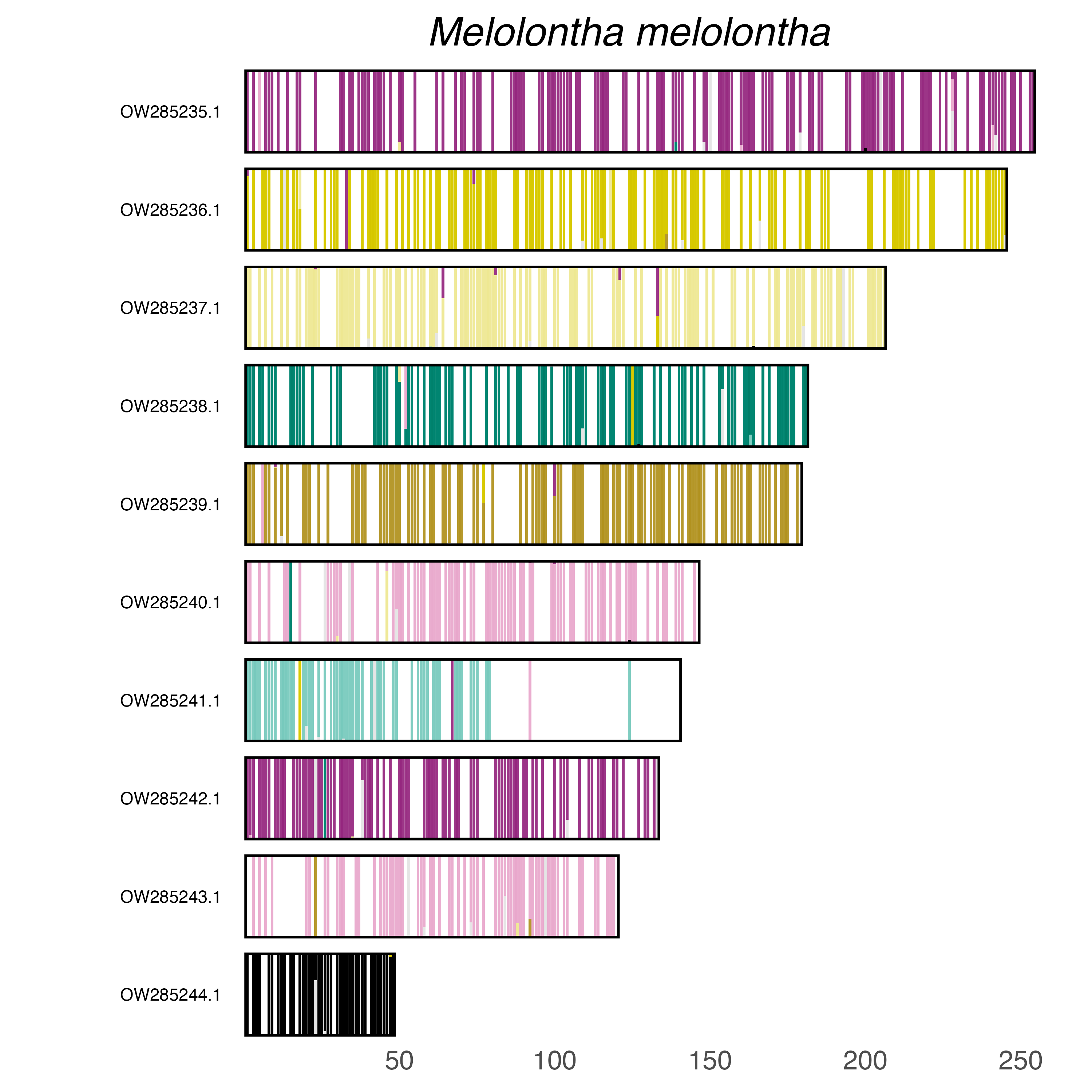

### plot.syngraph_windowbp_1Mb_rel_vertical_chrsize_Microrhagus_pygmaeus.png

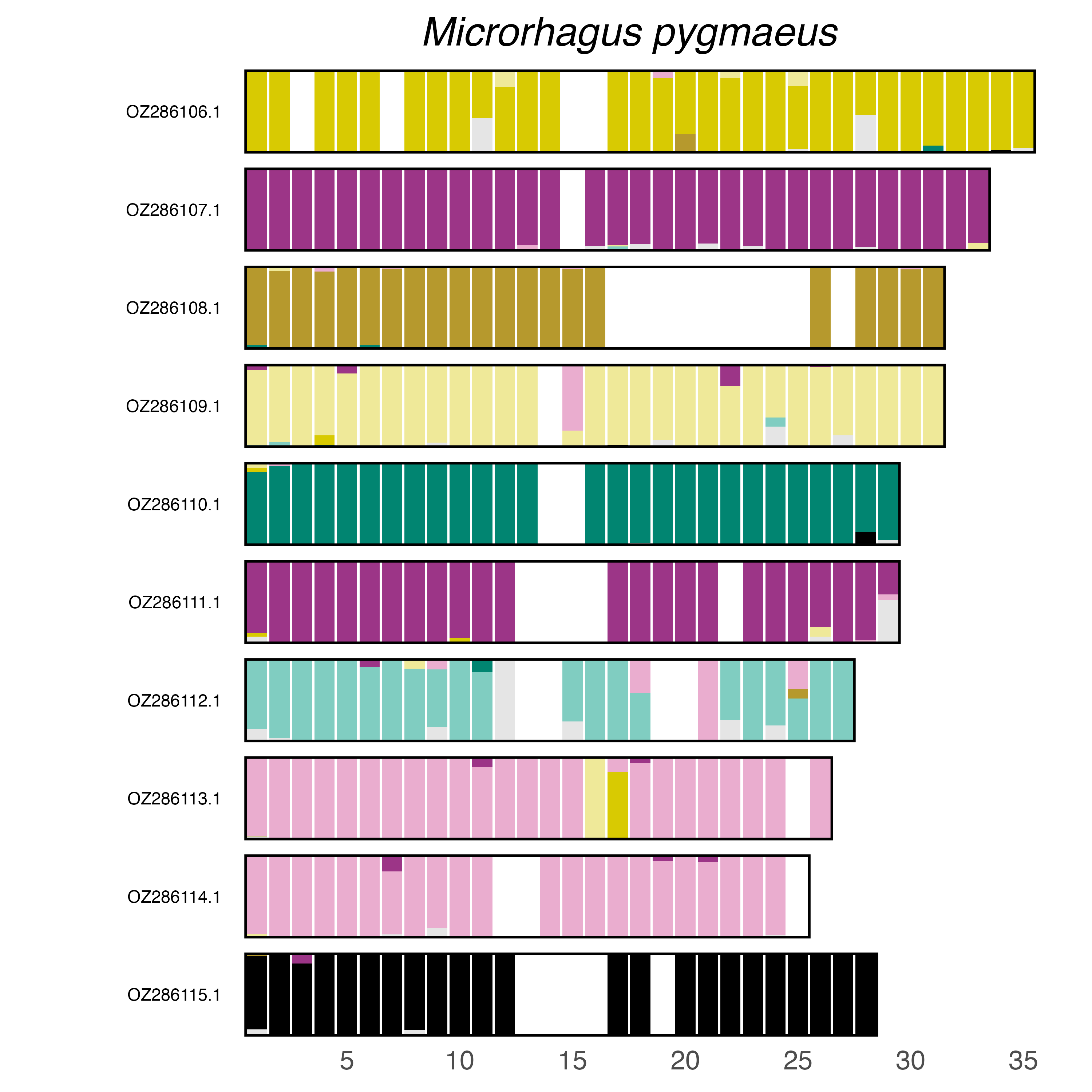

### plot.syngraph_windowbp_1Mb_rel_vertical_chrsize_Mongoloraphidia_duomilia.png

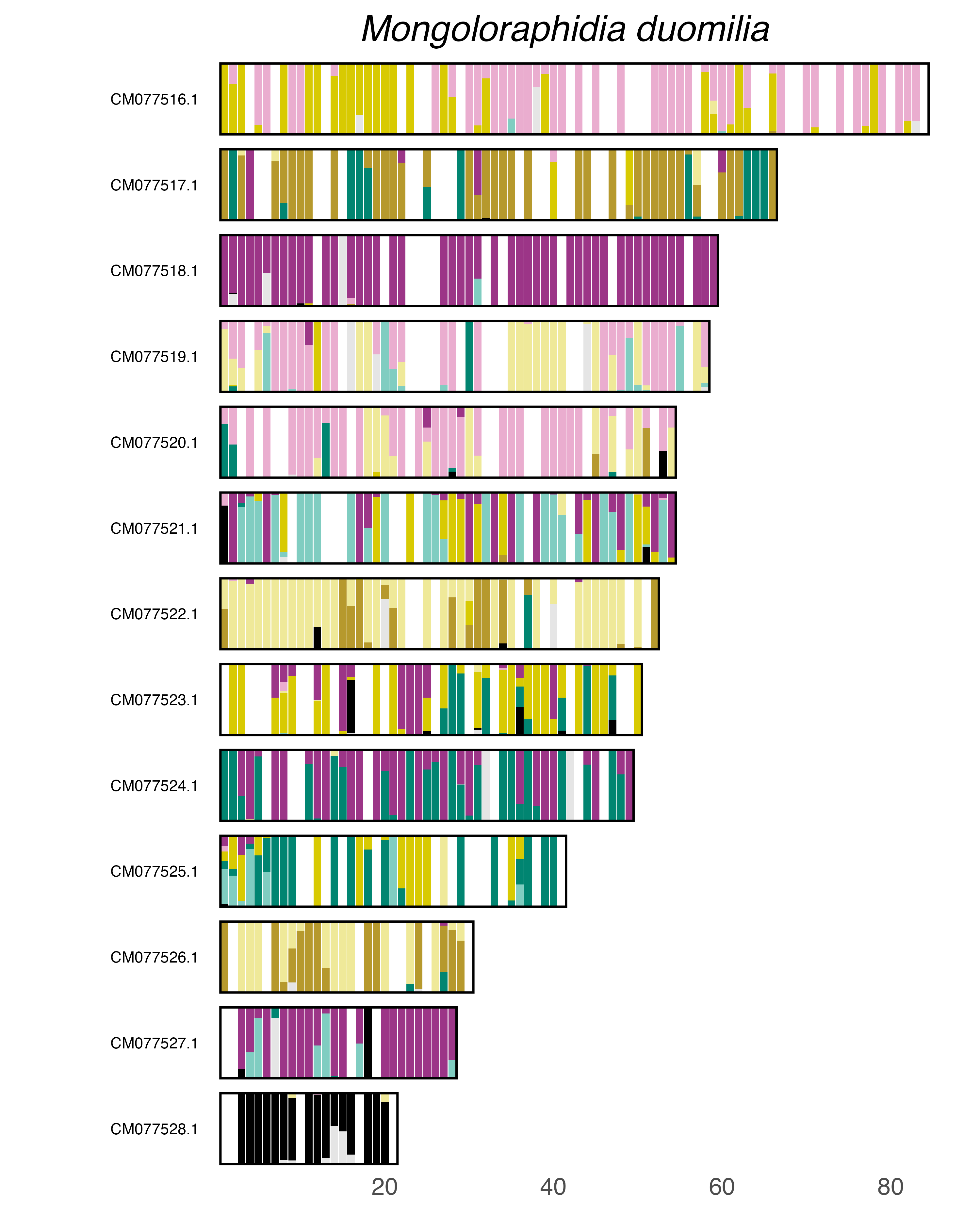

### plot.syngraph_windowbp_1Mb_rel_vertical_chrsize_Monotoma_picipes.png

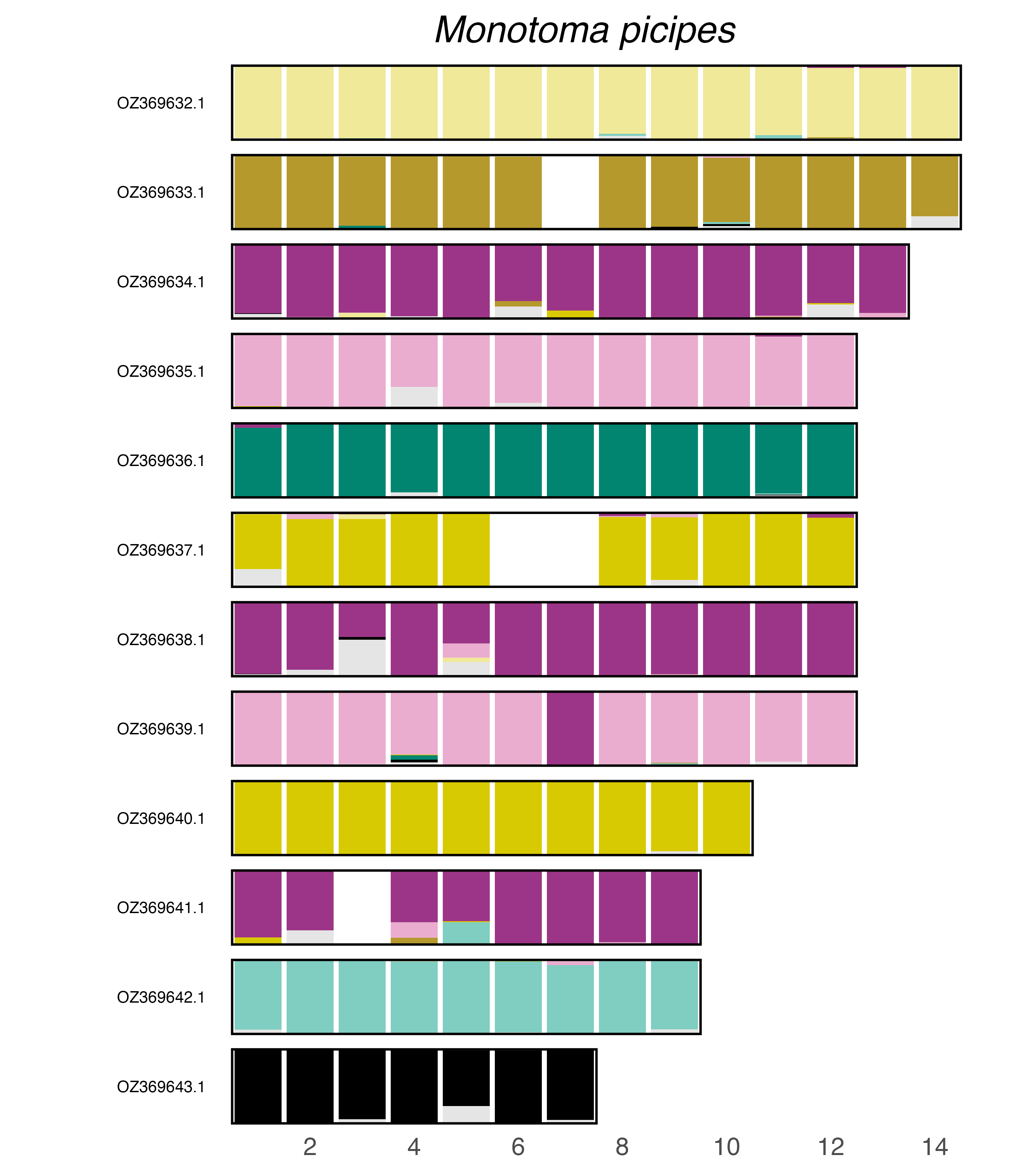

### plot.syngraph_windowbp_1Mb_rel_vertical_chrsize_Mycetophagus_atomarius.png

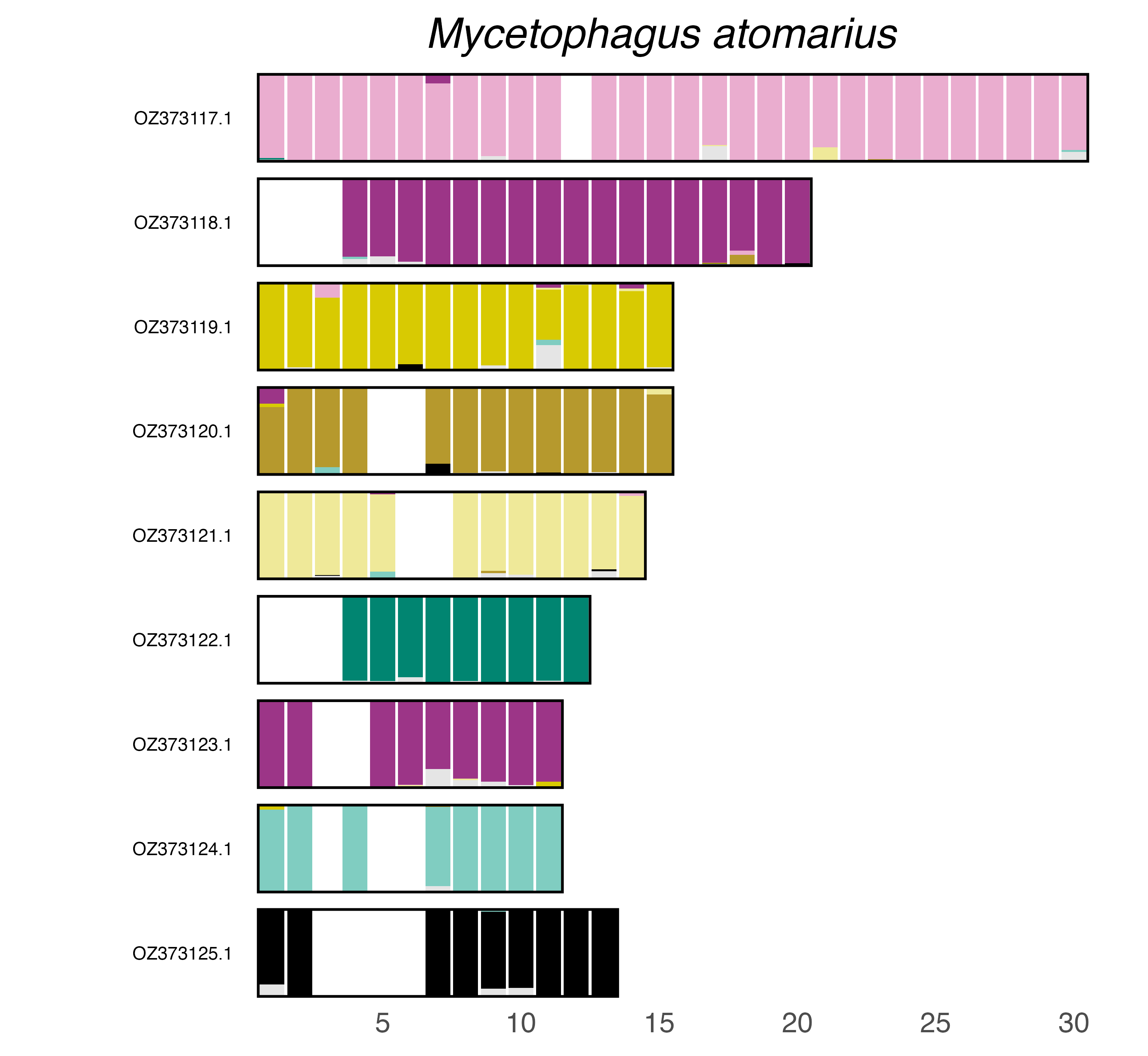

### plot.syngraph_windowbp_1Mb_rel_vertical_chrsize_Mycetophagus_piceus.png

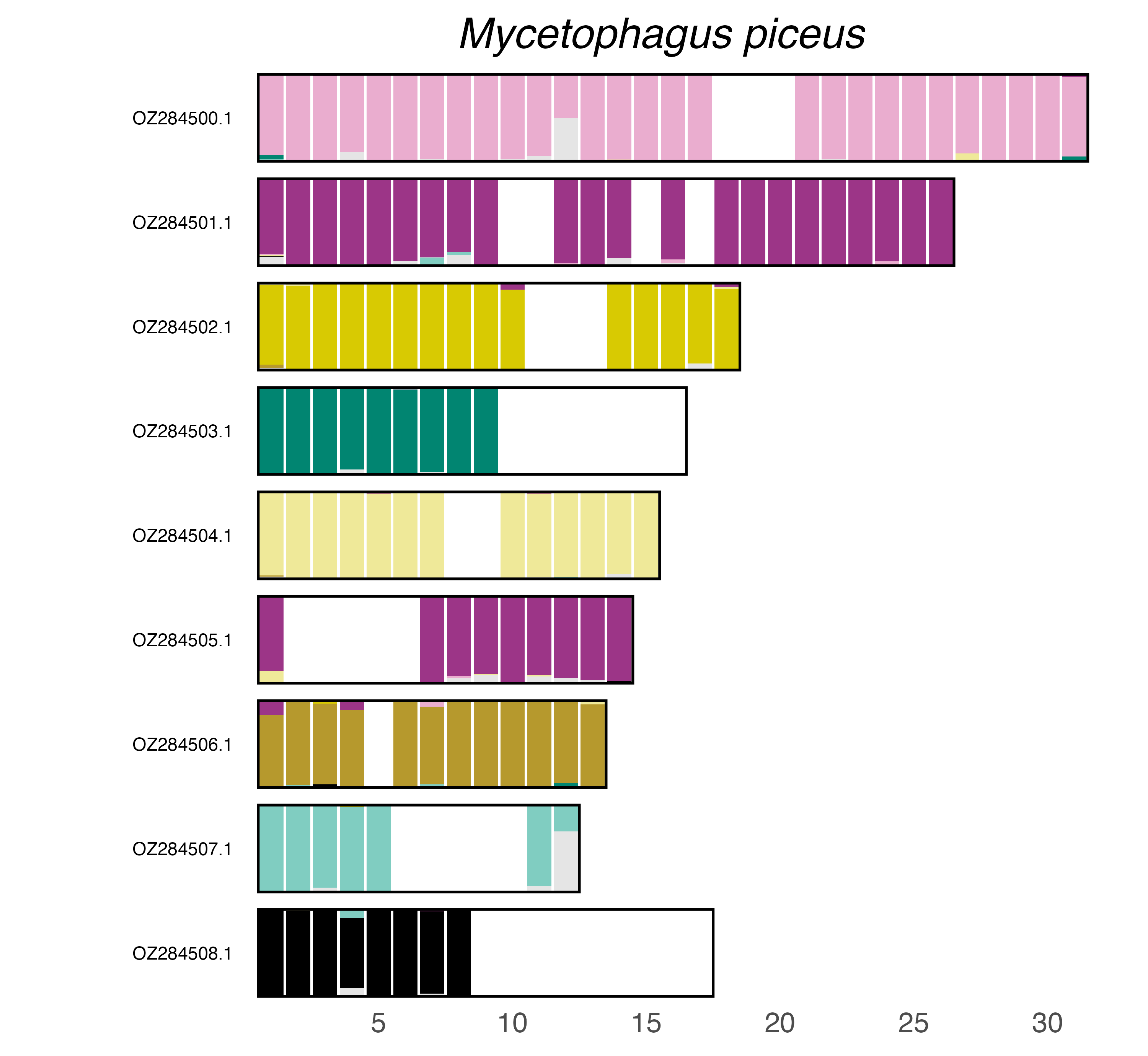

### plot.syngraph_windowbp_1Mb_rel_vertical_chrsize_Mylabris_sibirica.png

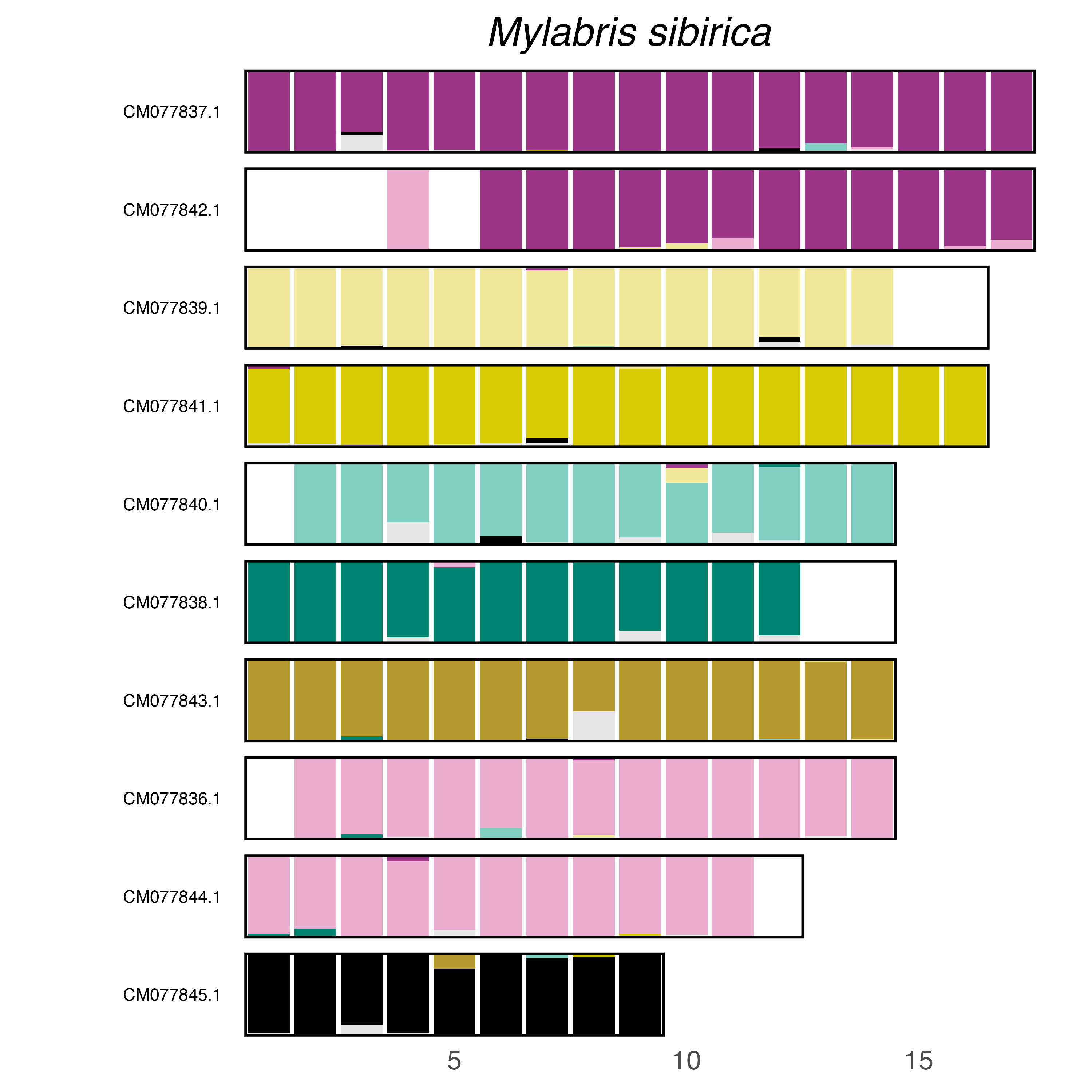

### plot.syngraph_windowbp_1Mb_rel_vertical_chrsize_Nadezhdiella_cantori.png

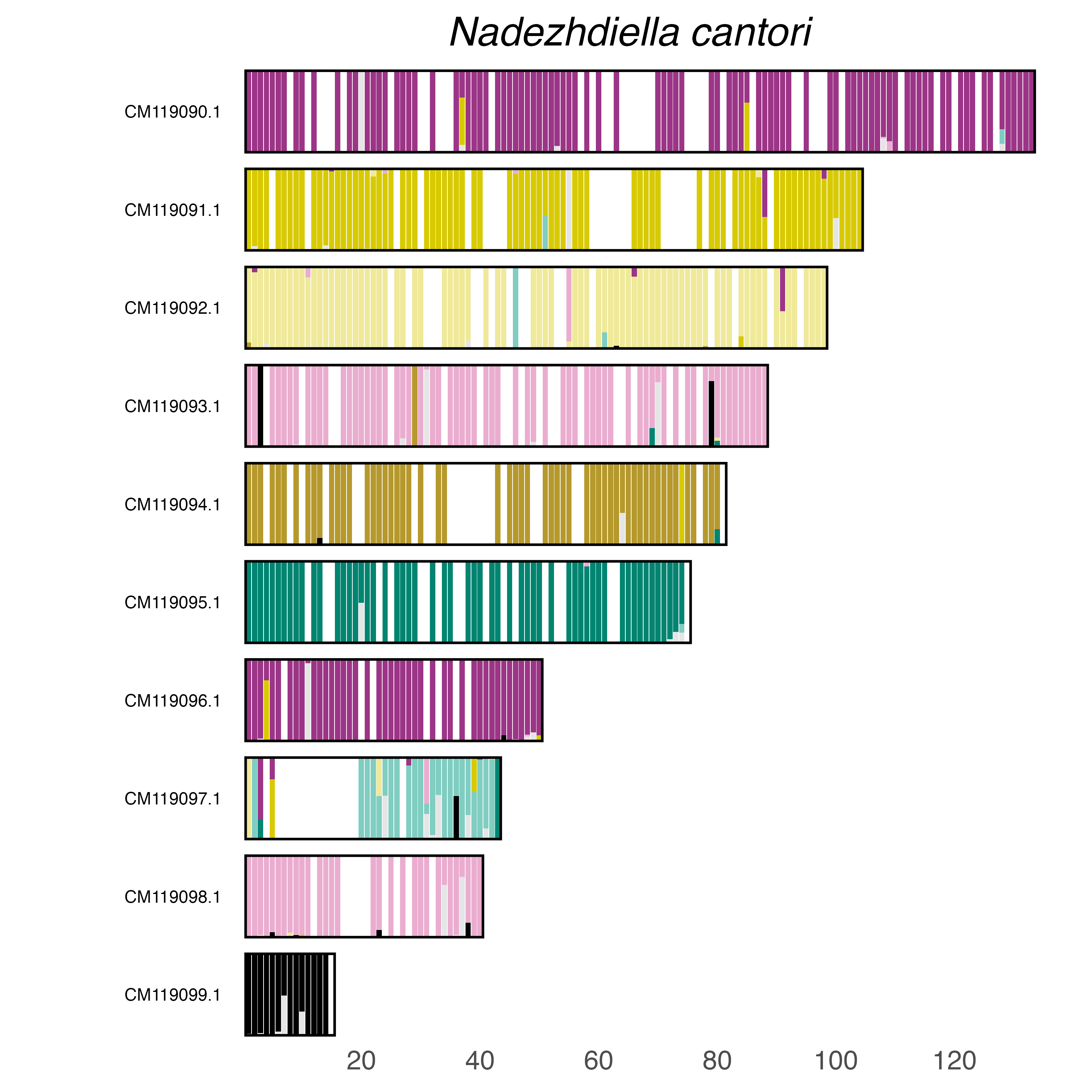

### plot.syngraph_windowbp_1Mb_rel_vertical_chrsize_Nebria_brevicollis.png

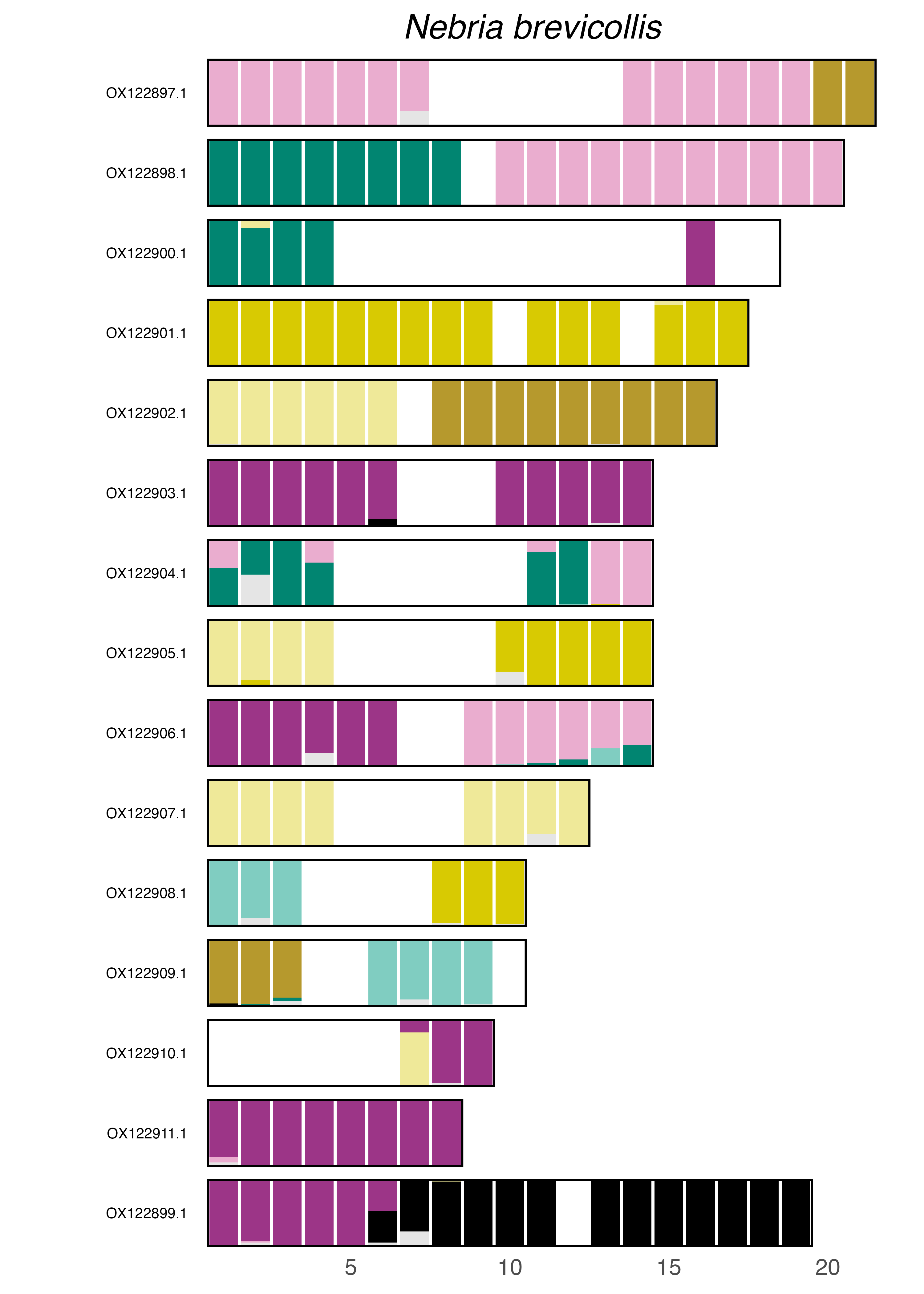

### plot.syngraph_windowbp_1Mb_rel_vertical_chrsize_Nineta_flava.png

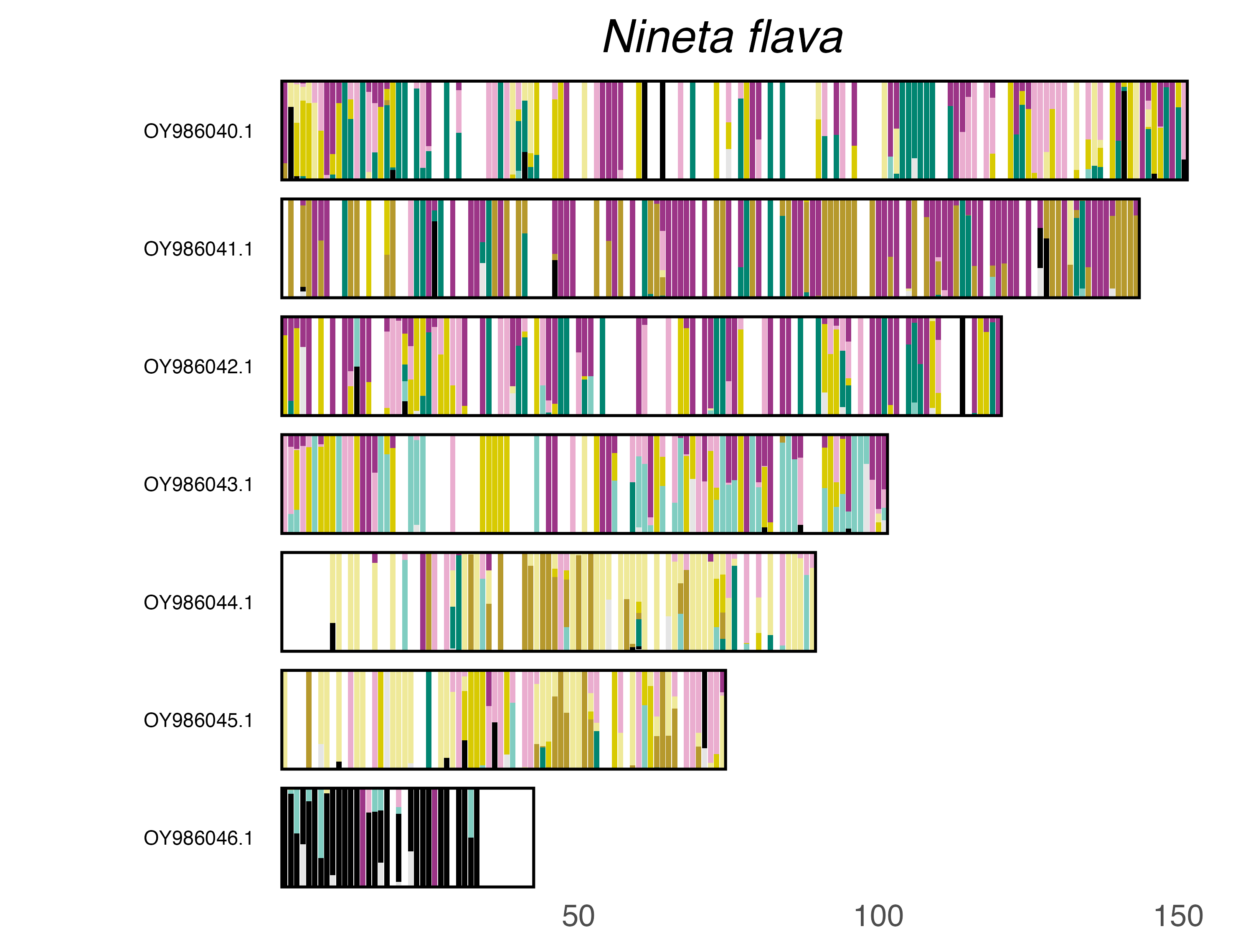

### plot.syngraph_windowbp_1Mb_rel_vertical_chrsize_Notoxus_monoceros.png

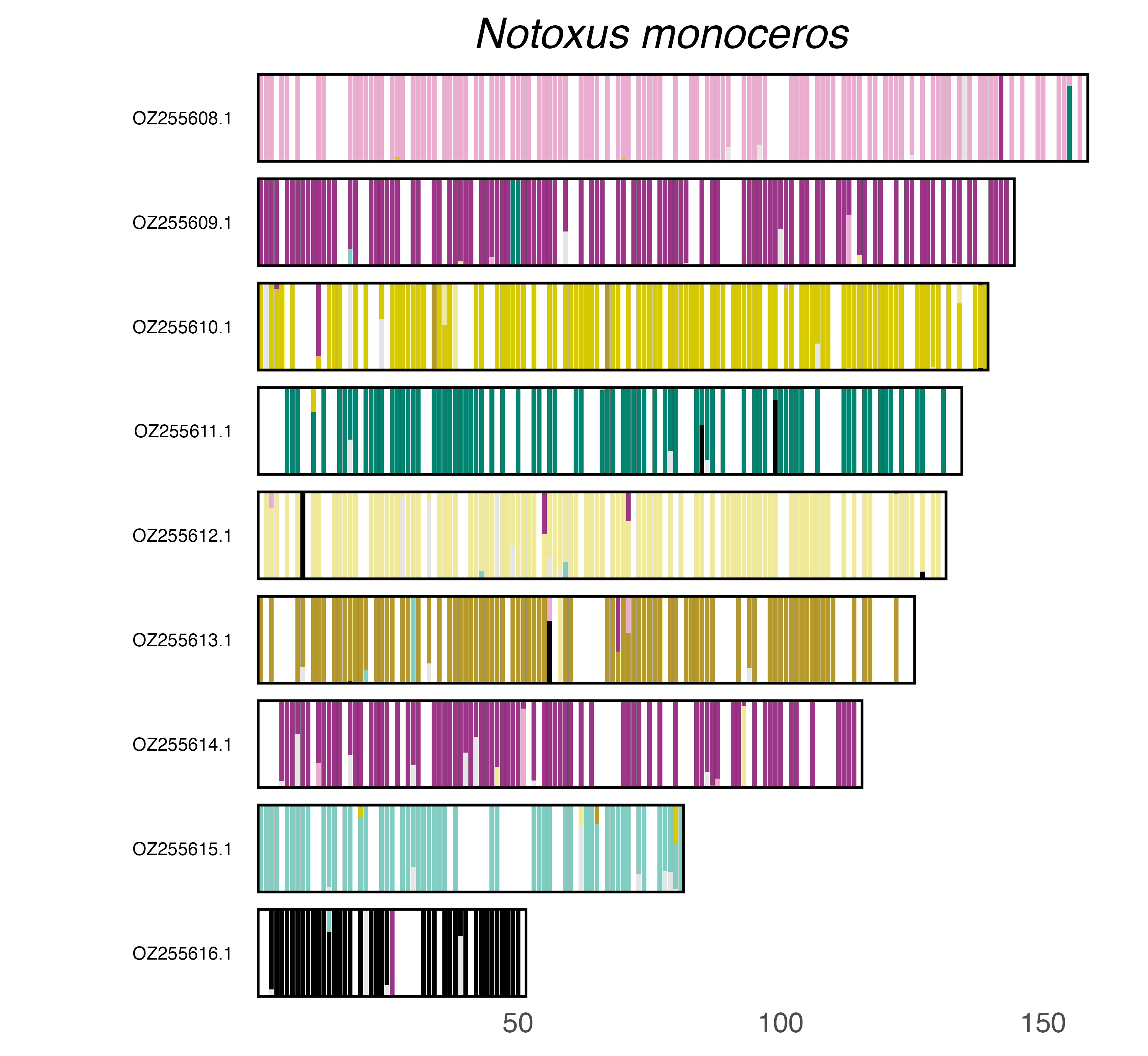

### plot.syngraph_windowbp_1Mb_rel_vertical_chrsize_Ocypus_olens.png

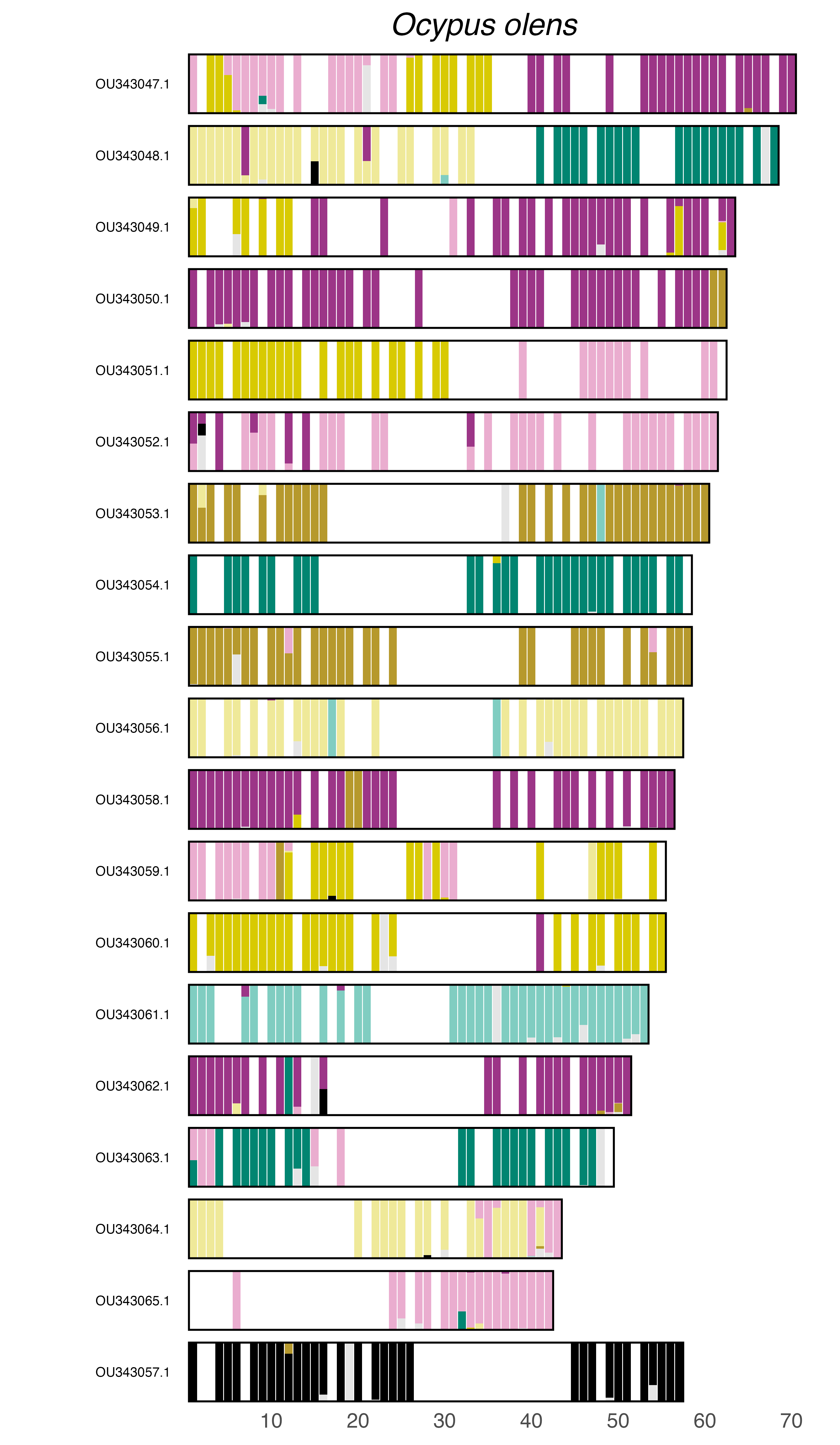

### plot.syngraph_windowbp_1Mb_rel_vertical_chrsize_Odontolabis_cuvera.png

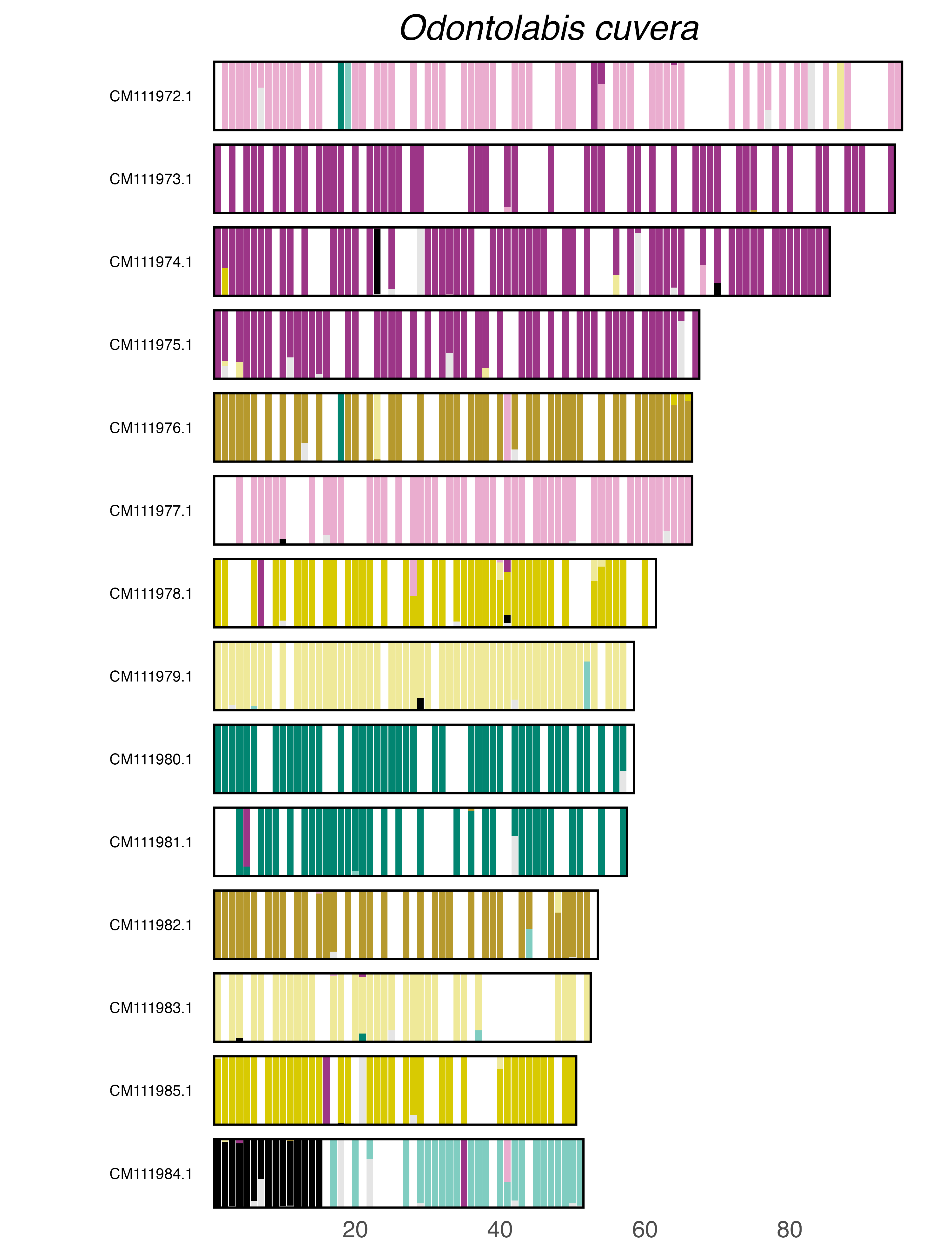

### plot.syngraph_windowbp_1Mb_rel_vertical_chrsize_Onthophagus_binodis.png

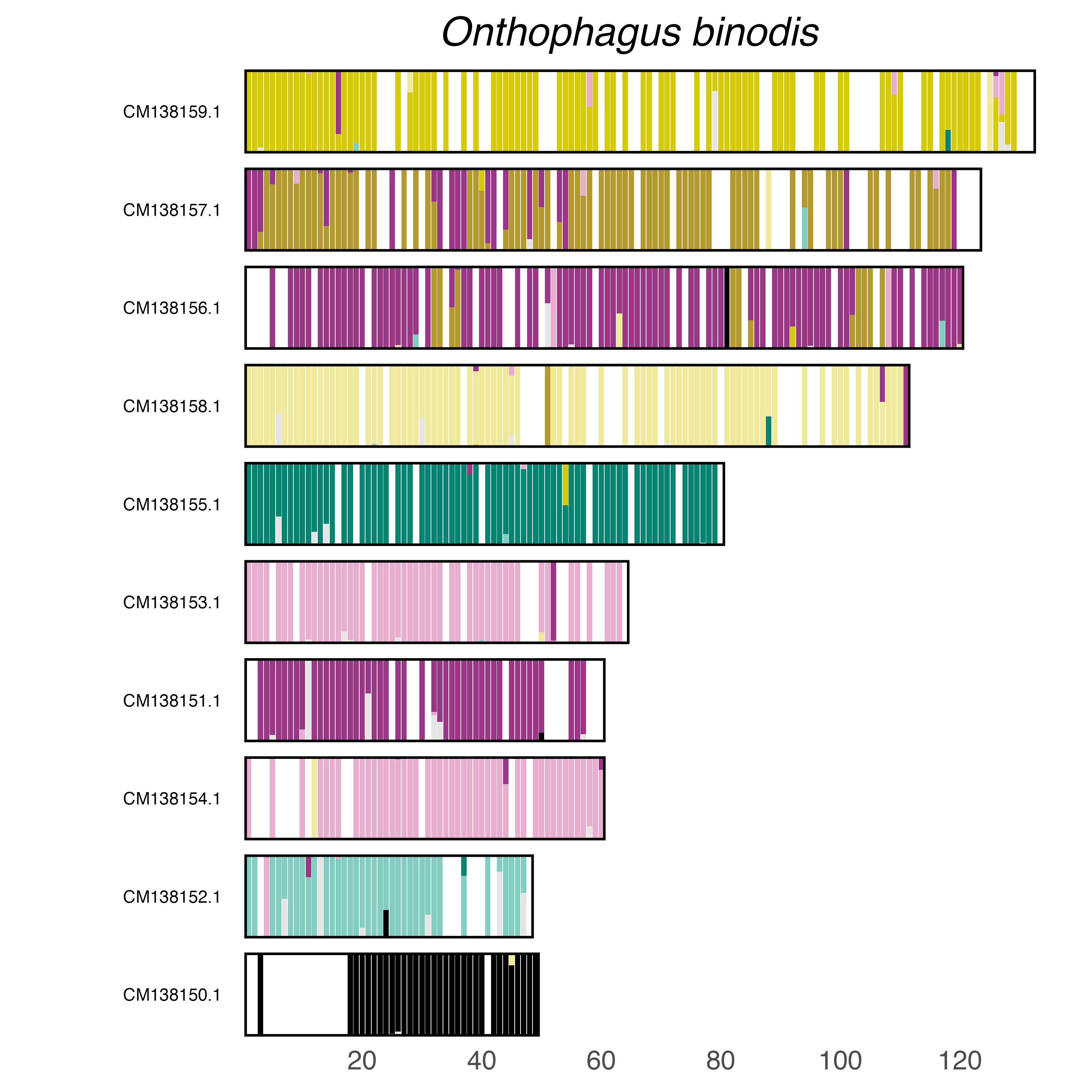

### plot.syngraph_windowbp_1Mb_rel_vertical_chrsize_Ophraella_communa.png

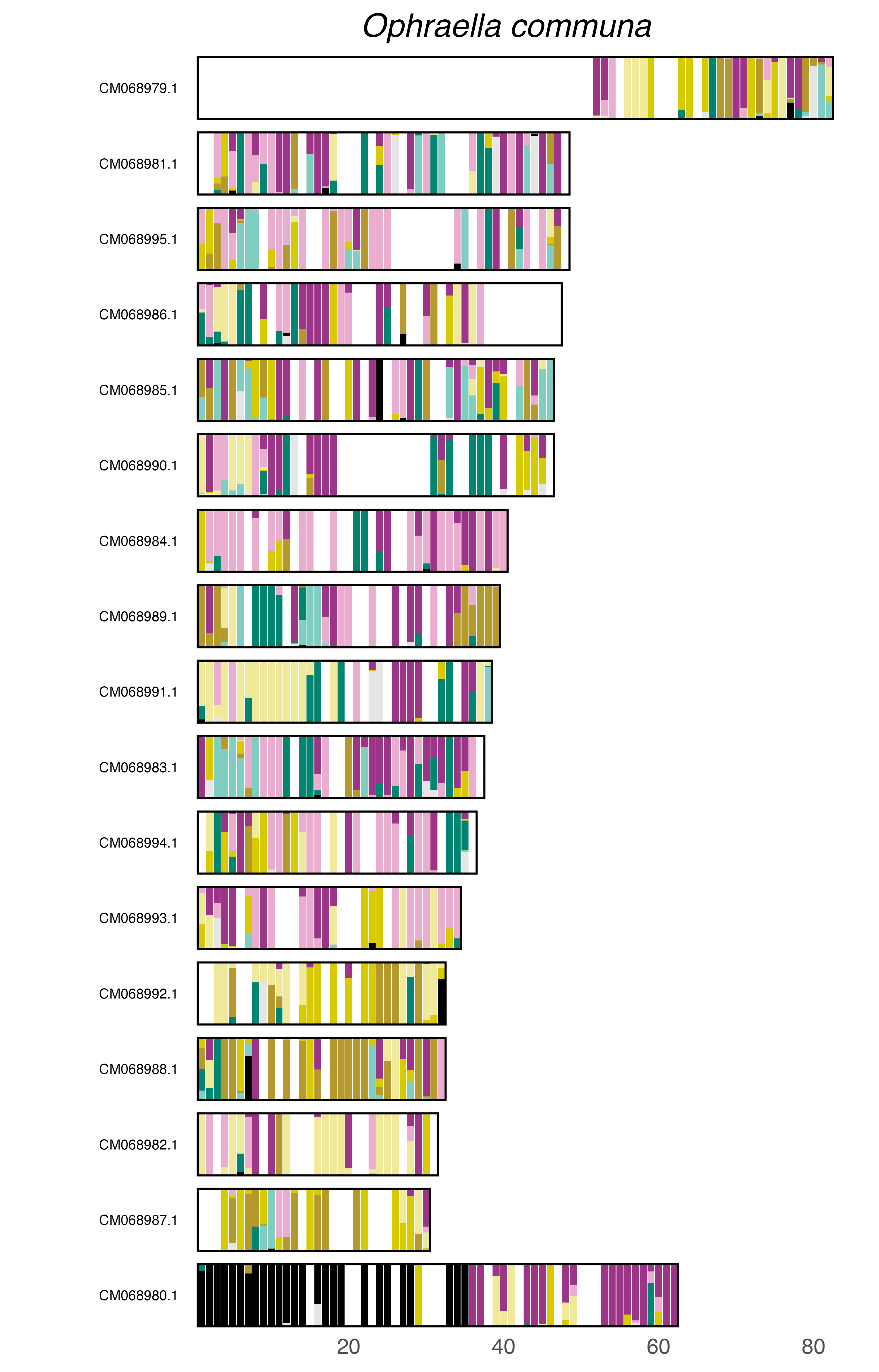

### plot.syngraph_windowbp_1Mb_rel_vertical_chrsize_Orchestes_rusci.png

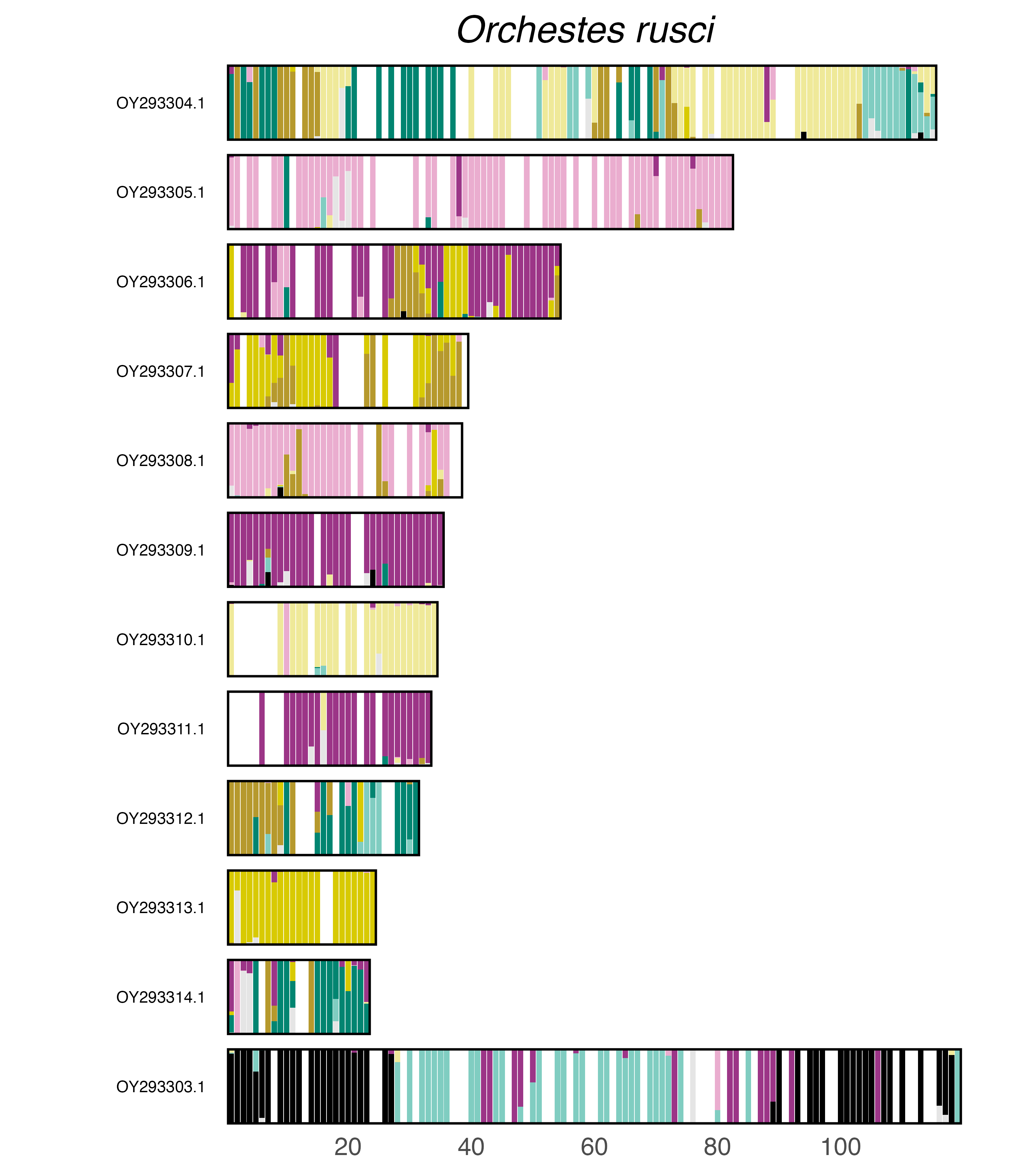

### plot.syngraph_windowbp_1Mb_rel_vertical_chrsize_Phaedon_cochleariae.png

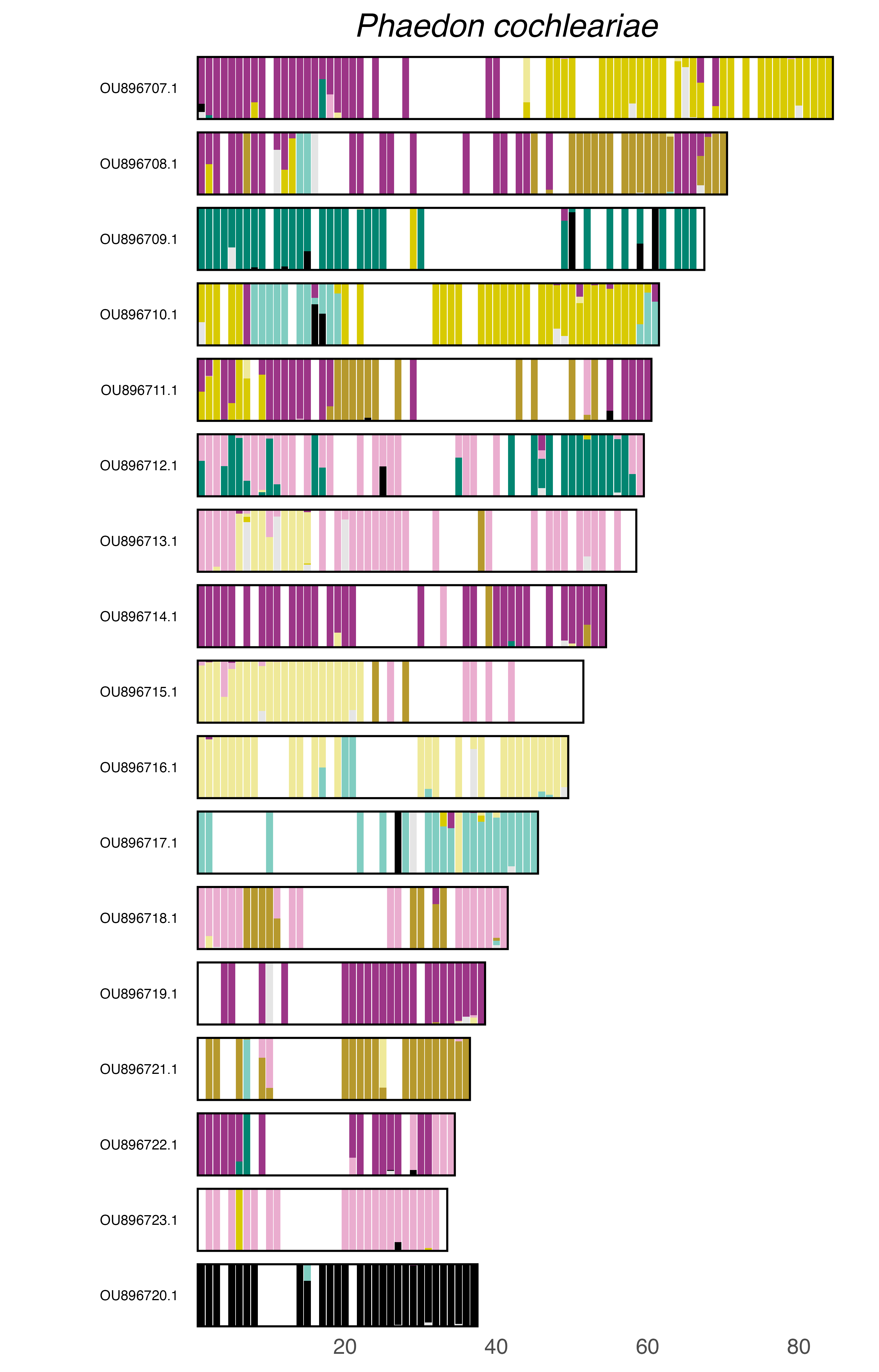

### plot.syngraph_windowbp_1Mb_rel_vertical_chrsize_Phaleria_cadaverina.png

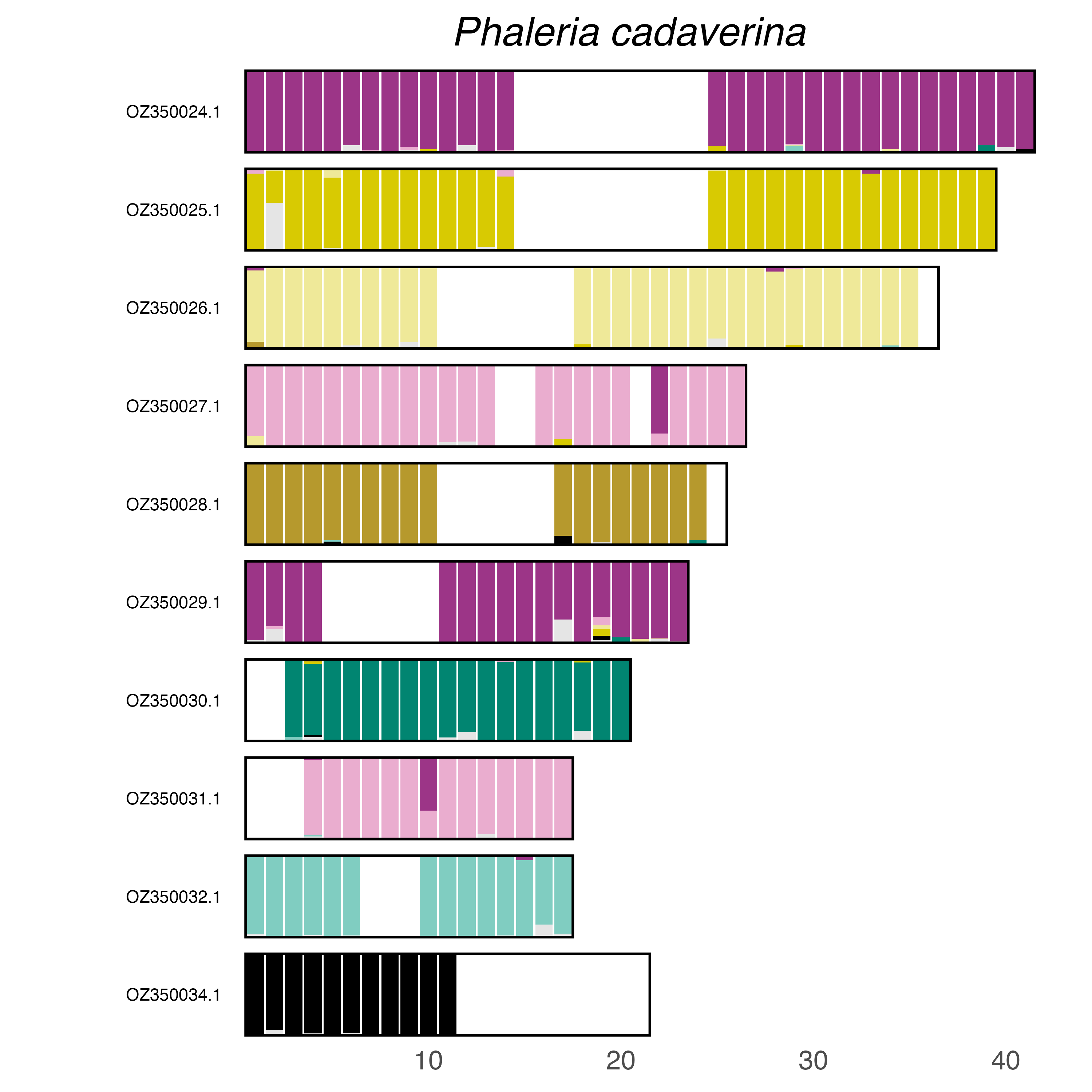

### plot.syngraph_windowbp_1Mb_rel_vertical_chrsize_Philonthus_cognatus.png

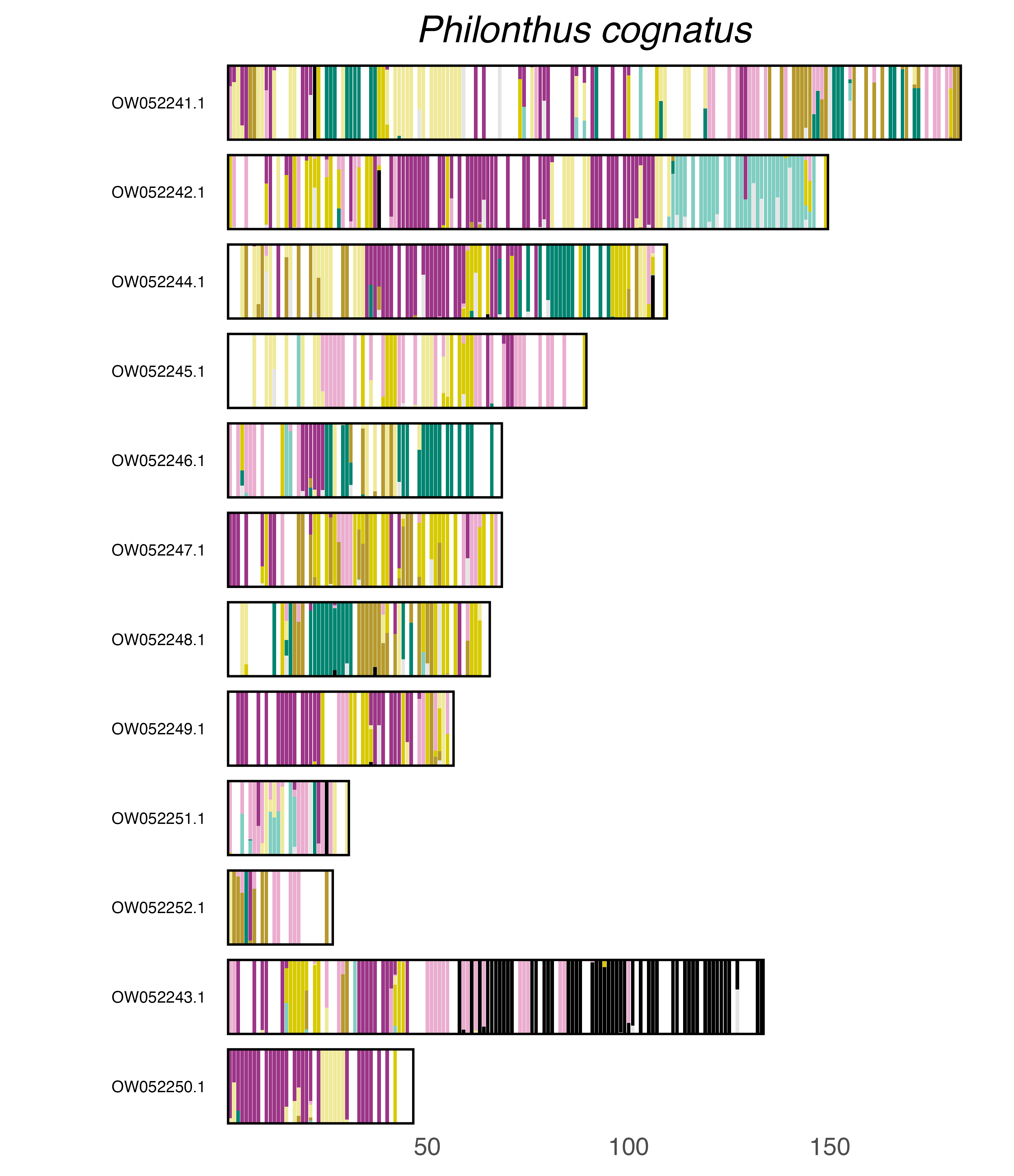

### plot.syngraph_windowbp_1Mb_rel_vertical_chrsize_Phratora_vitellinae.png

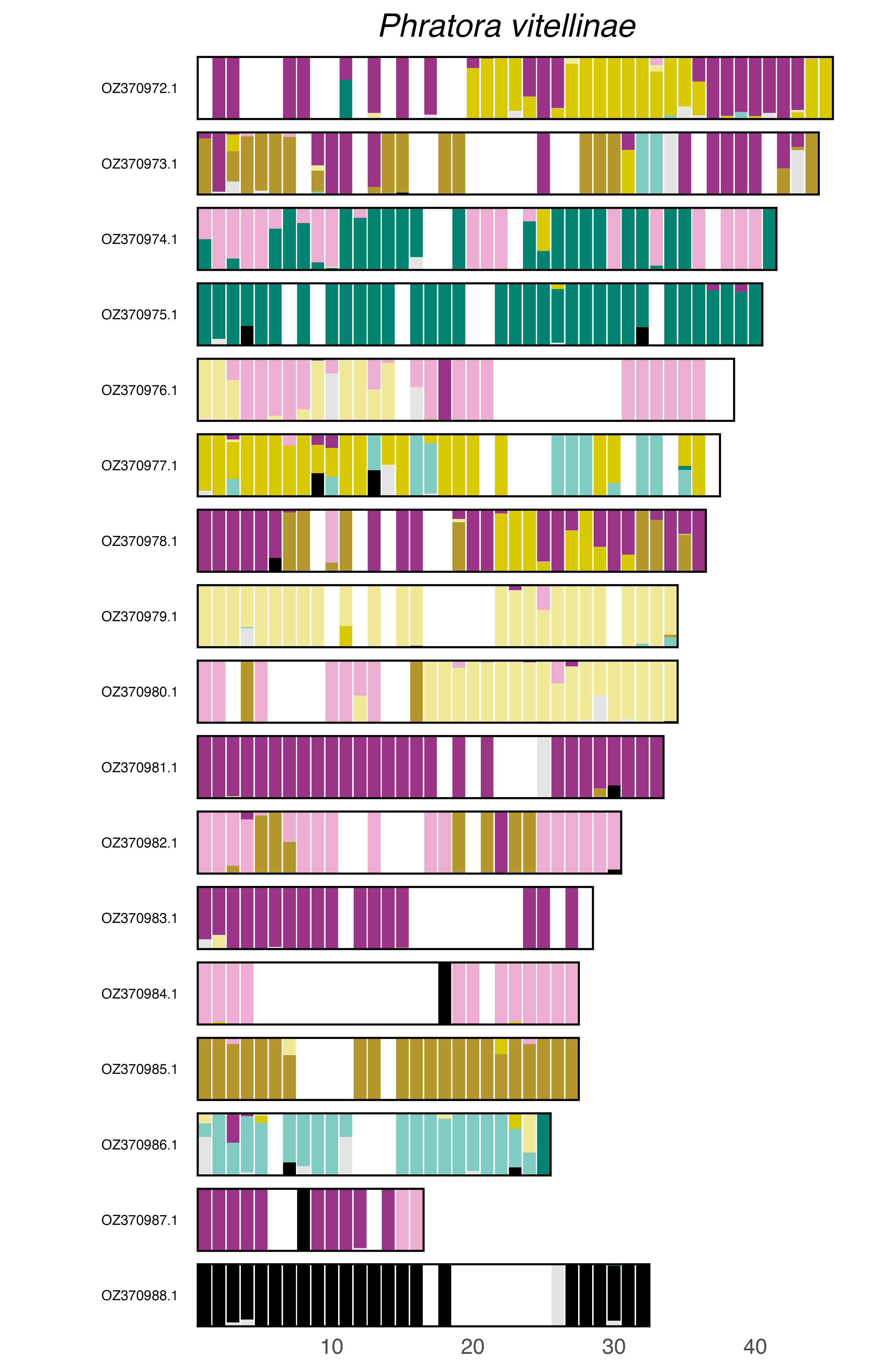

### plot.syngraph_windowbp_1Mb_rel_vertical_chrsize_Phyllobius_argentatus.png

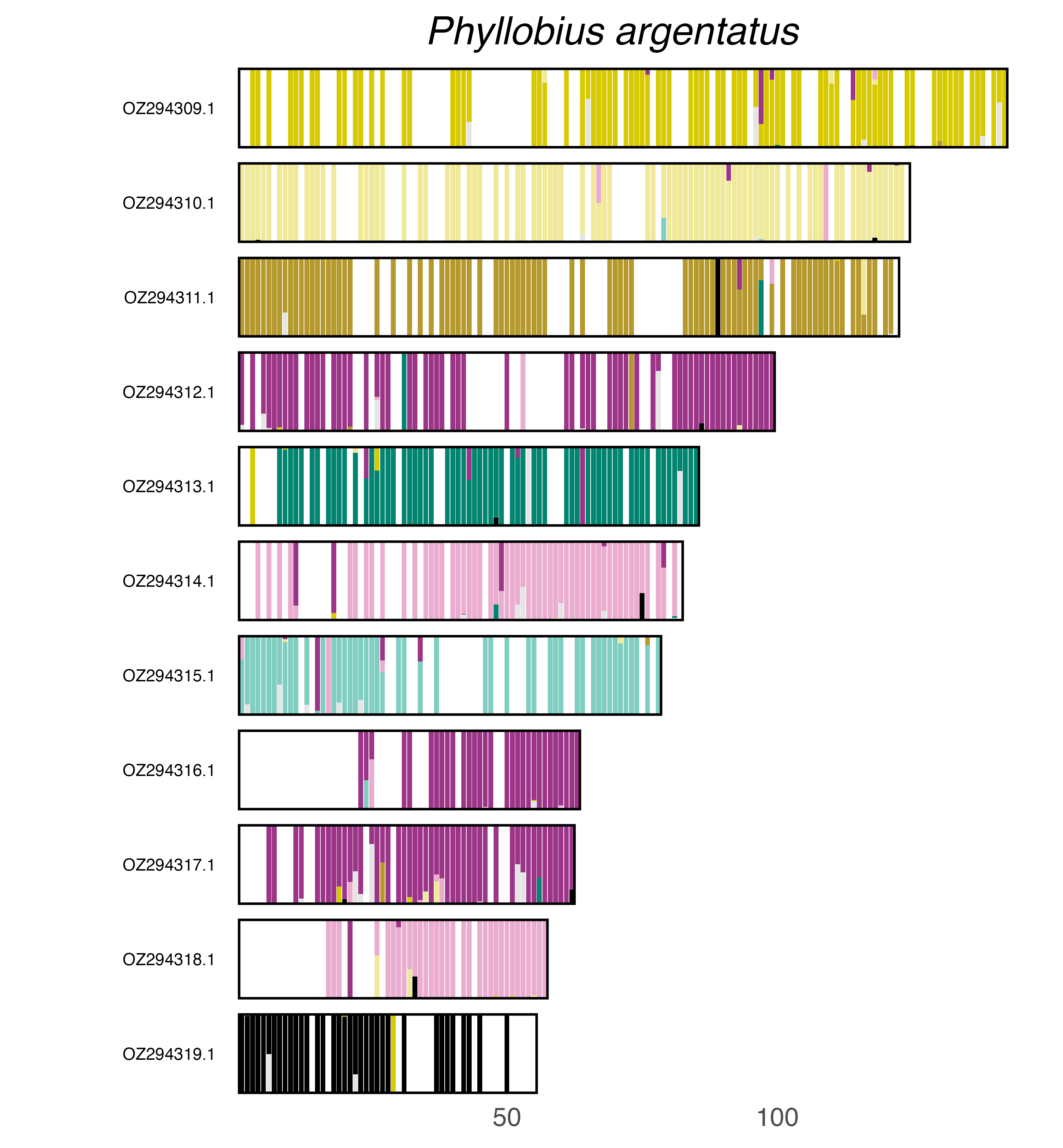

### plot.syngraph_windowbp_1Mb_rel_vertical_chrsize_Phyllobius_virideaeris.png

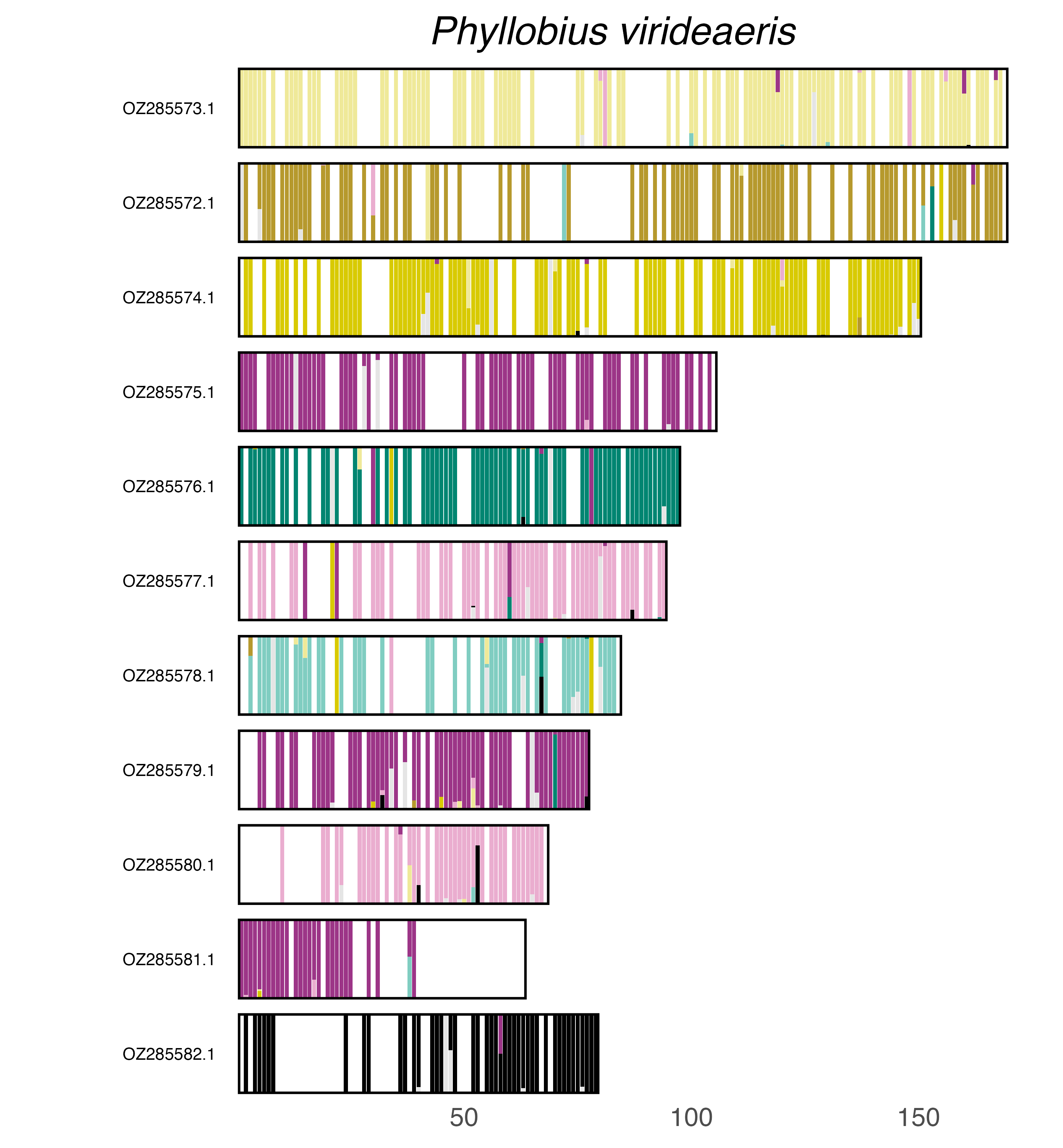

### plot.syngraph_windowbp_1Mb_rel_vertical_chrsize_Plagiodera_versicolora.png

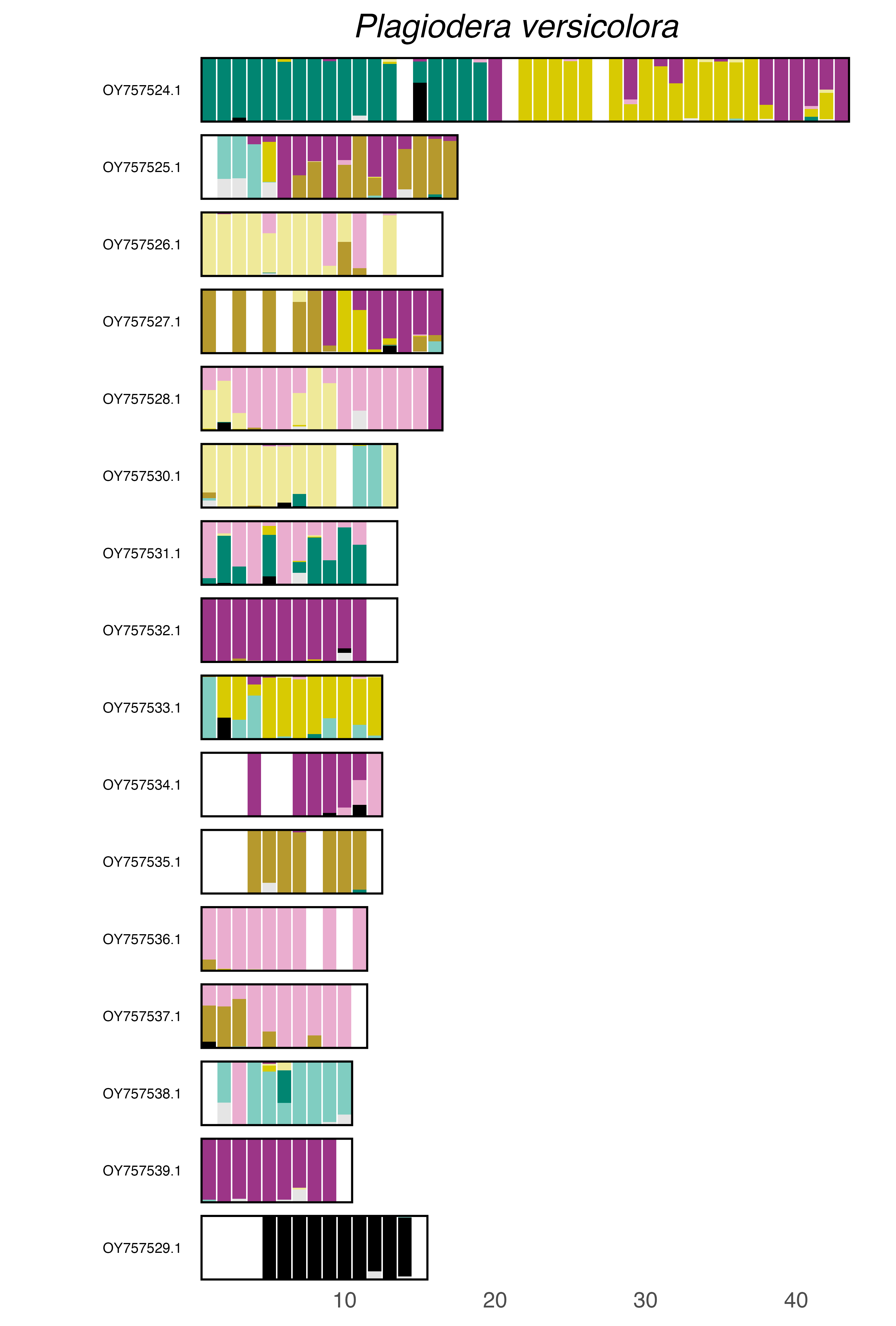

### plot.syngraph_windowbp_1Mb_rel_vertical_chrsize_Platypus_cylindrus.png

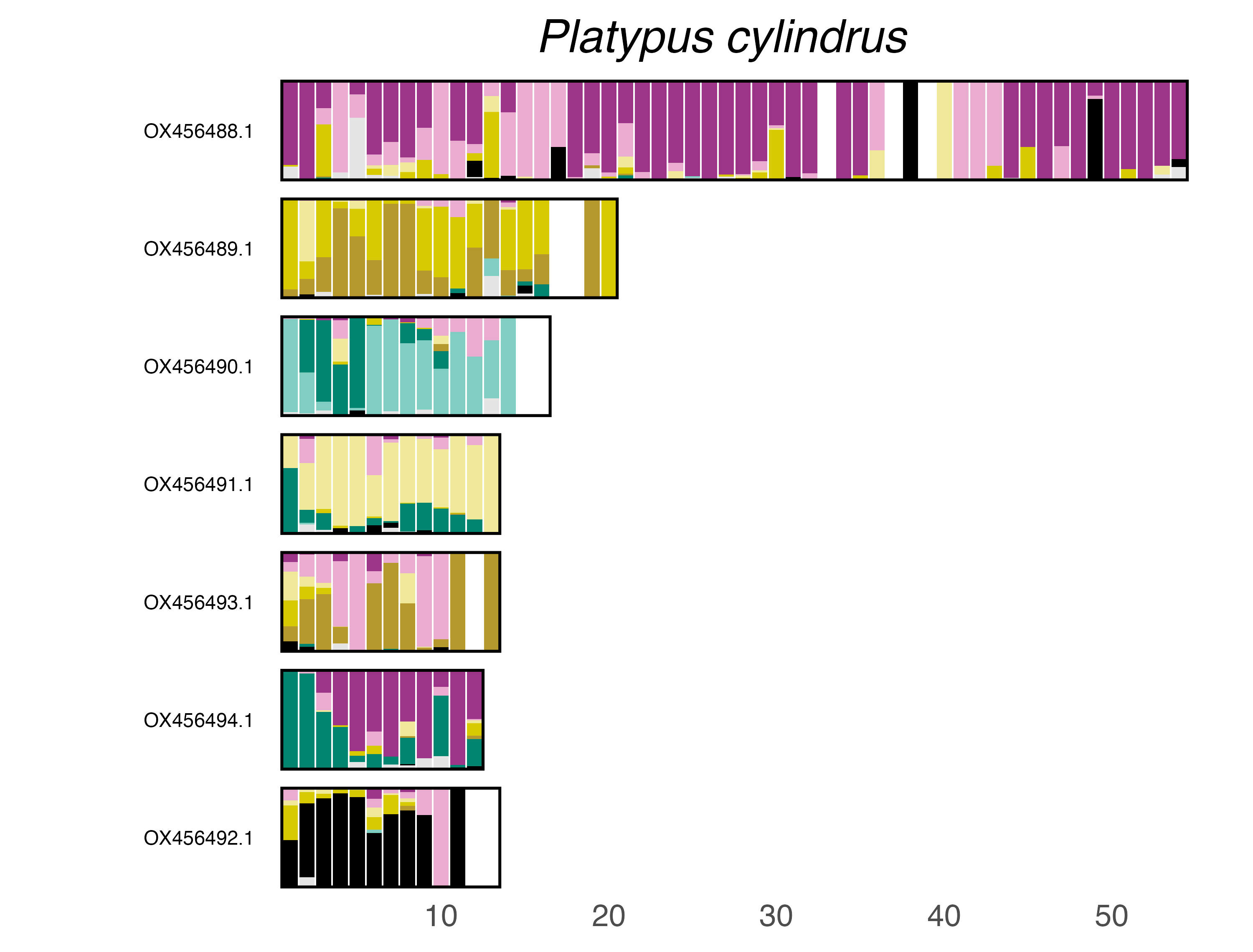

### plot.syngraph_windowbp_1Mb_rel_vertical_chrsize_Pogonus_chalceus.png

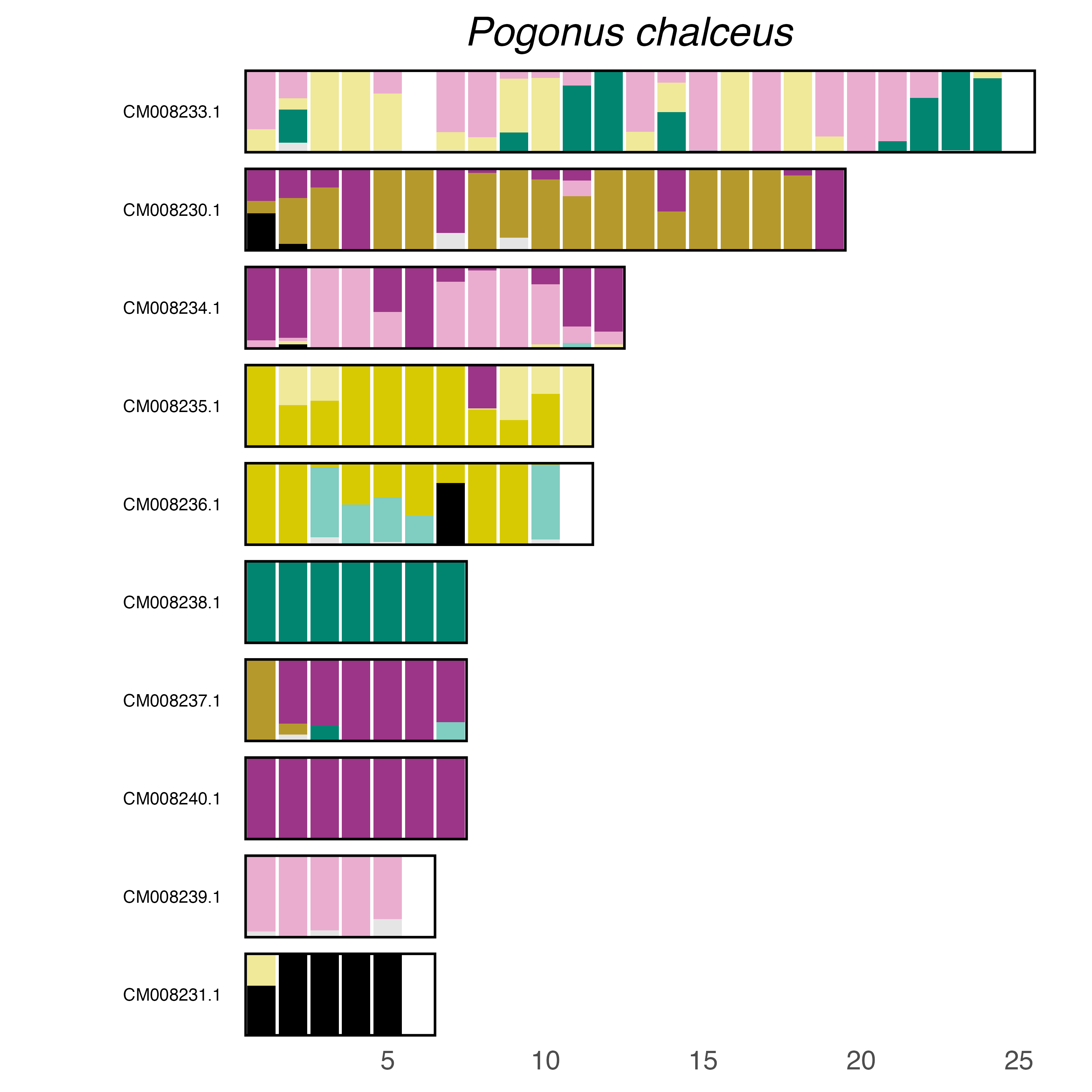

### plot.syngraph_windowbp_1Mb_rel_vertical_chrsize_Polydrusus_cervinus.png

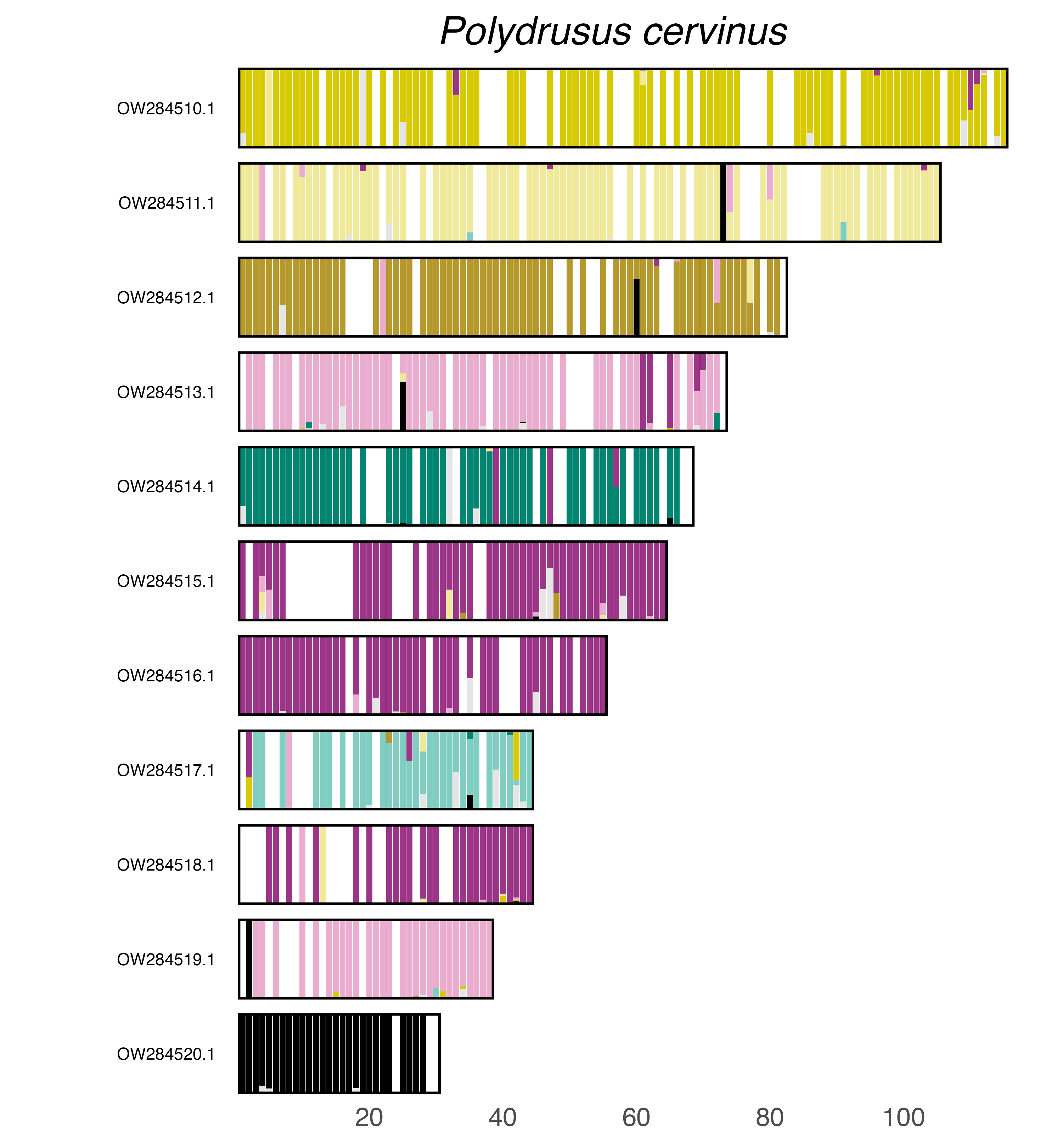

### plot.syngraph_windowbp_1Mb_rel_vertical_chrsize_Prionocyphon_serricornis.png

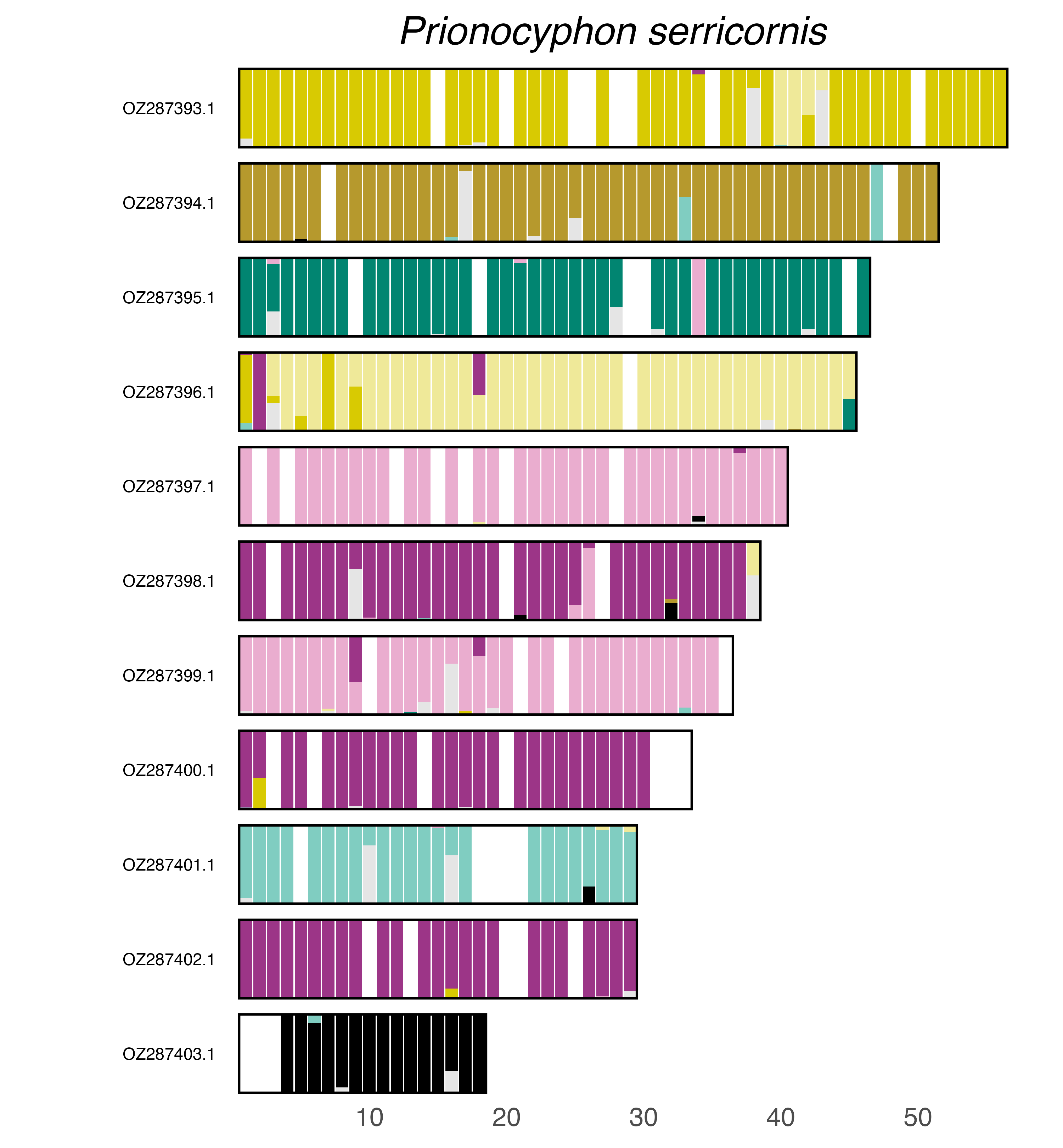

### plot.syngraph_windowbp_1Mb_rel_vertical_chrsize_Propylea_japonica.png

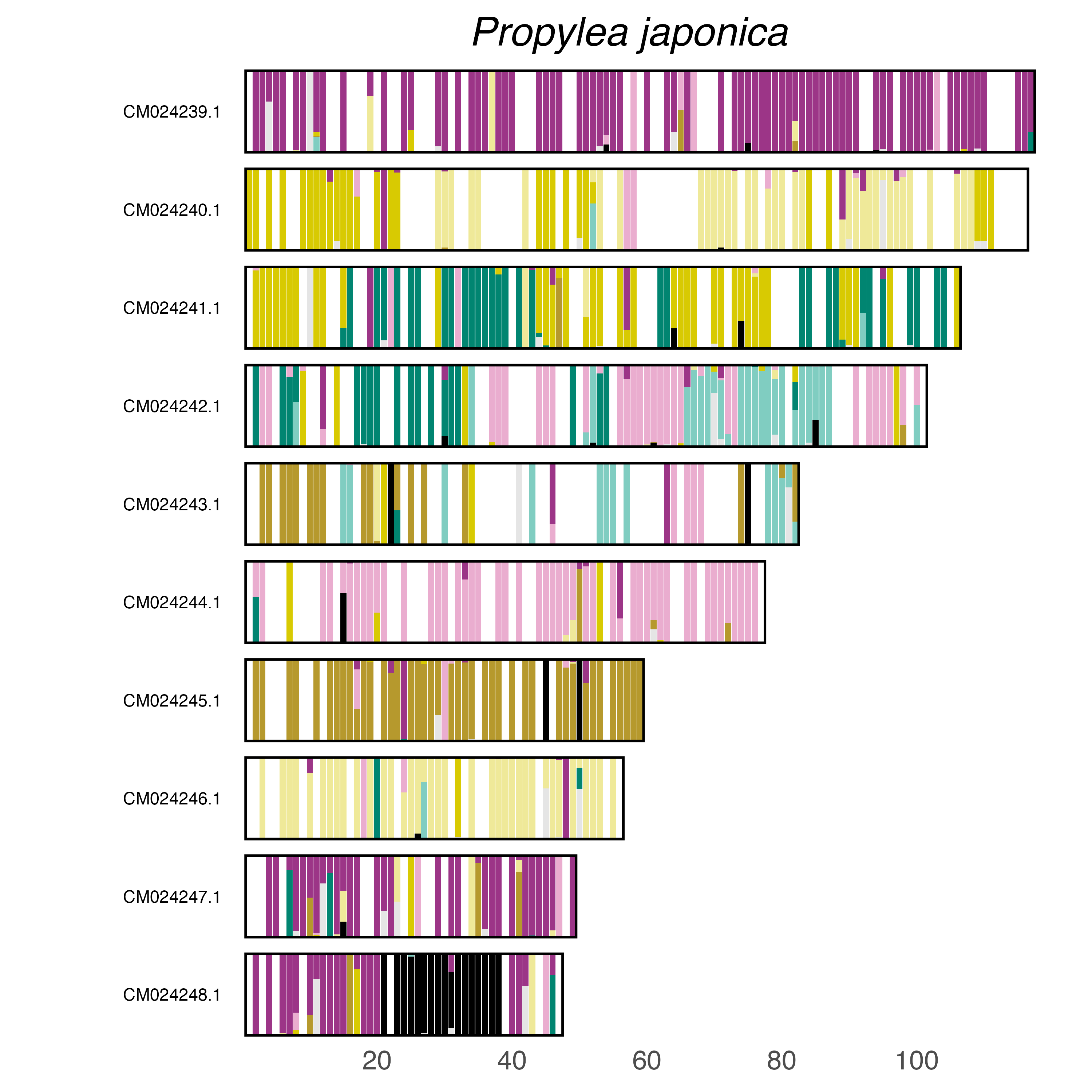

### plot.syngraph_windowbp_1Mb_rel_vertical_chrsize_Propylea_quattuordecimpunctata.png

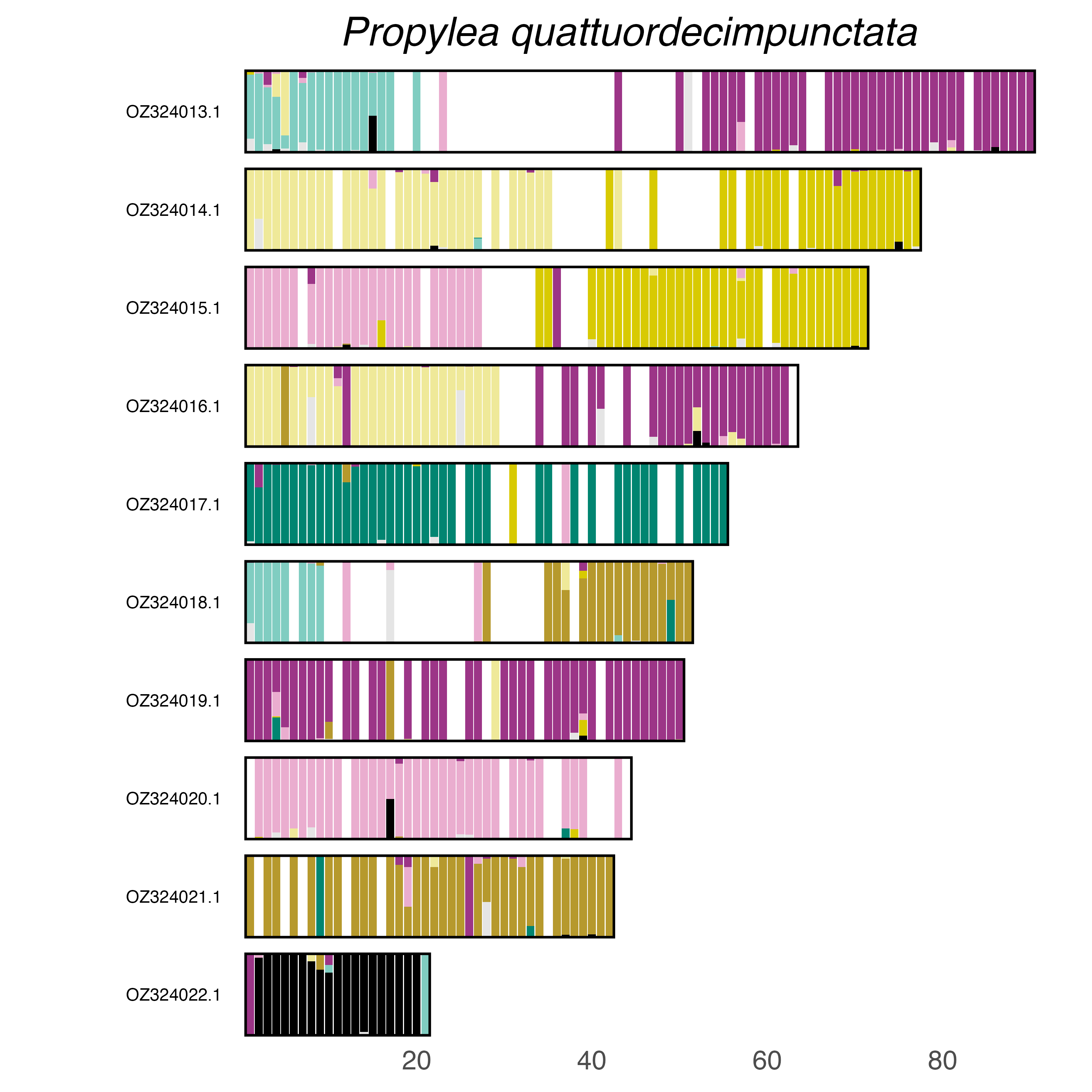

### plot.syngraph_windowbp_1Mb_rel_vertical_chrsize_Prosopocoilus_inquinatus.png

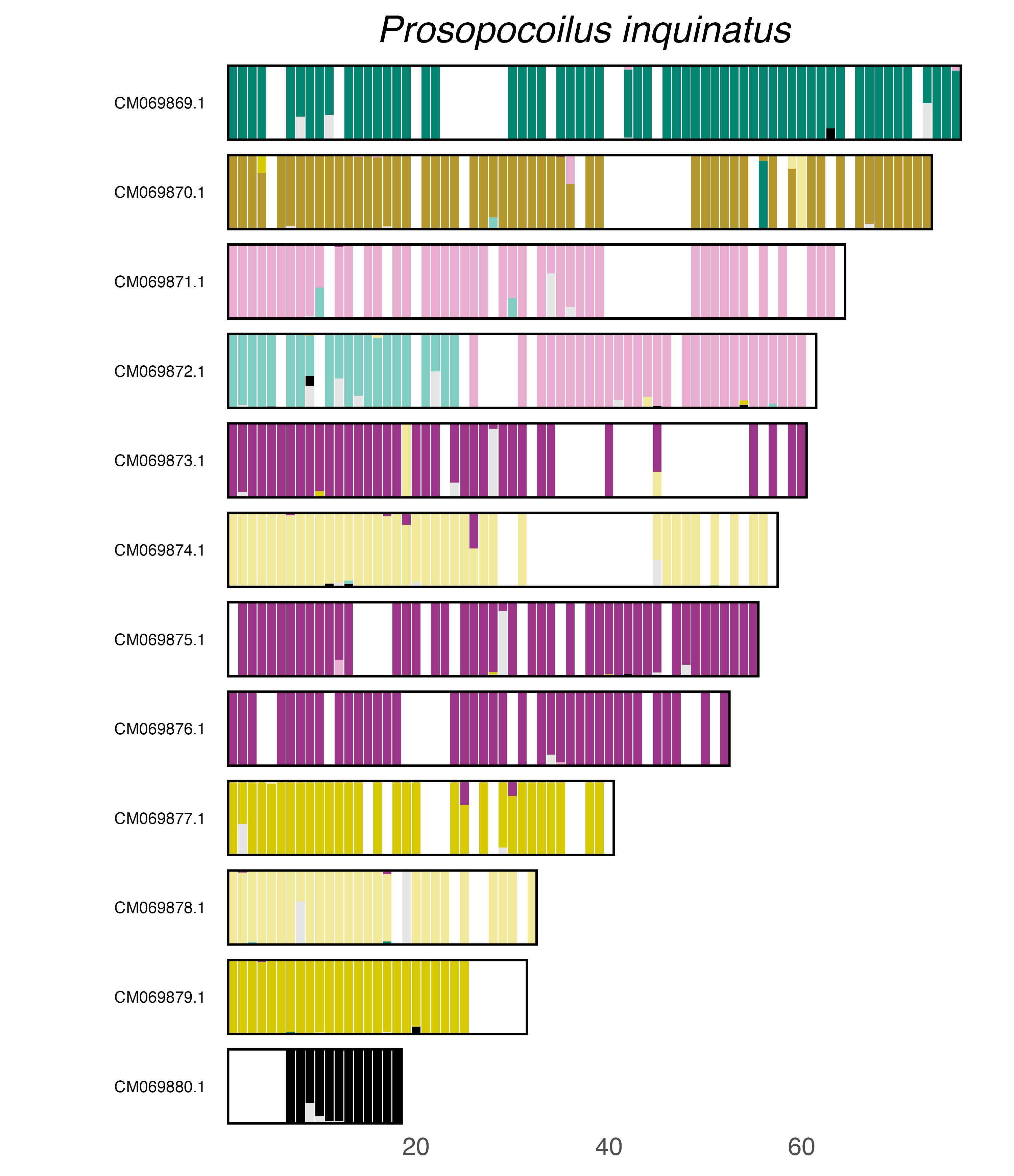

### plot.syngraph_windowbp_1Mb_rel_vertical_chrsize_Psylliodes_chrysocephalus.png

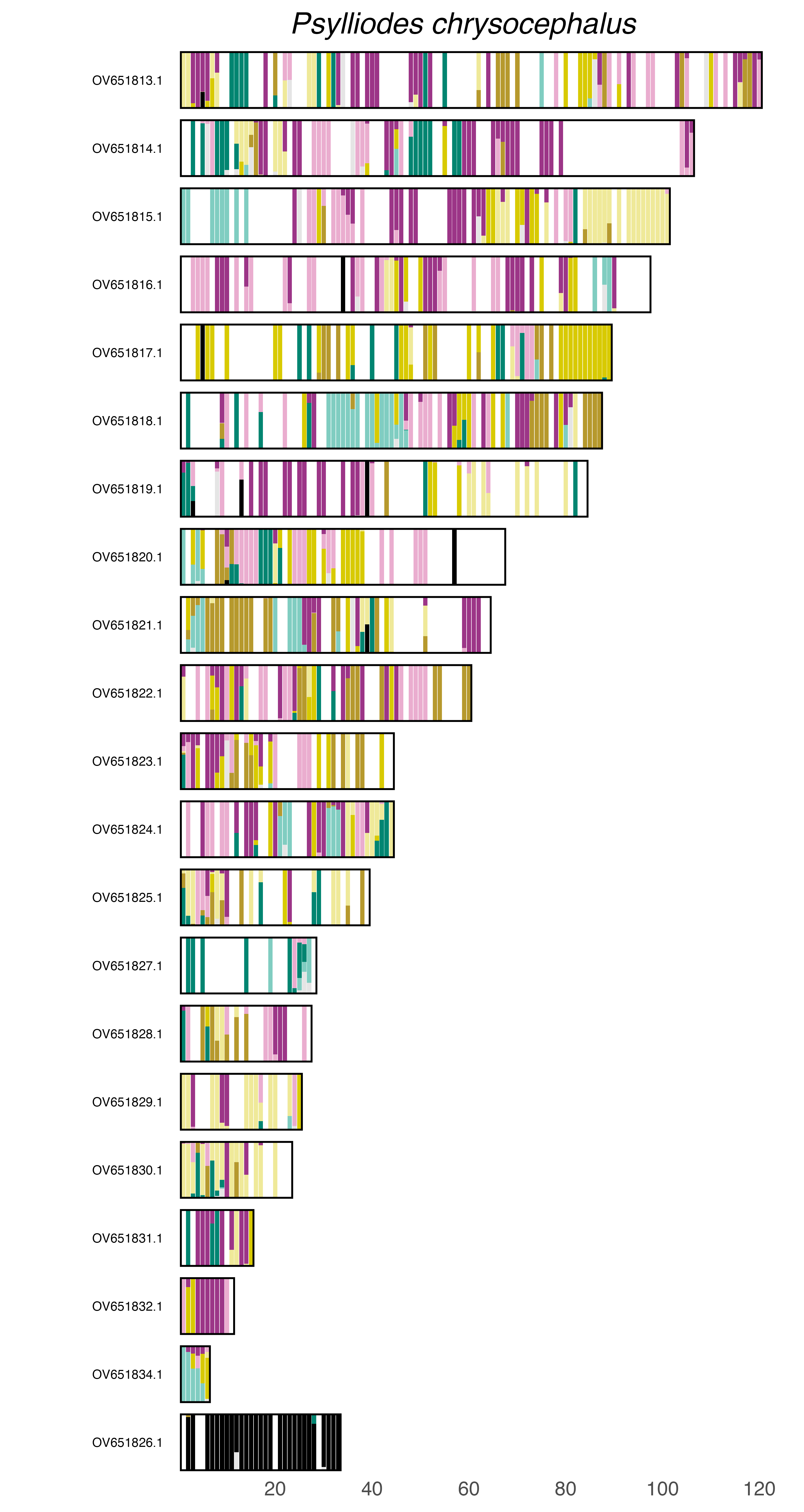

### plot.syngraph_windowbp_1Mb_rel_vertical_chrsize_Pterostichus_diligens.png

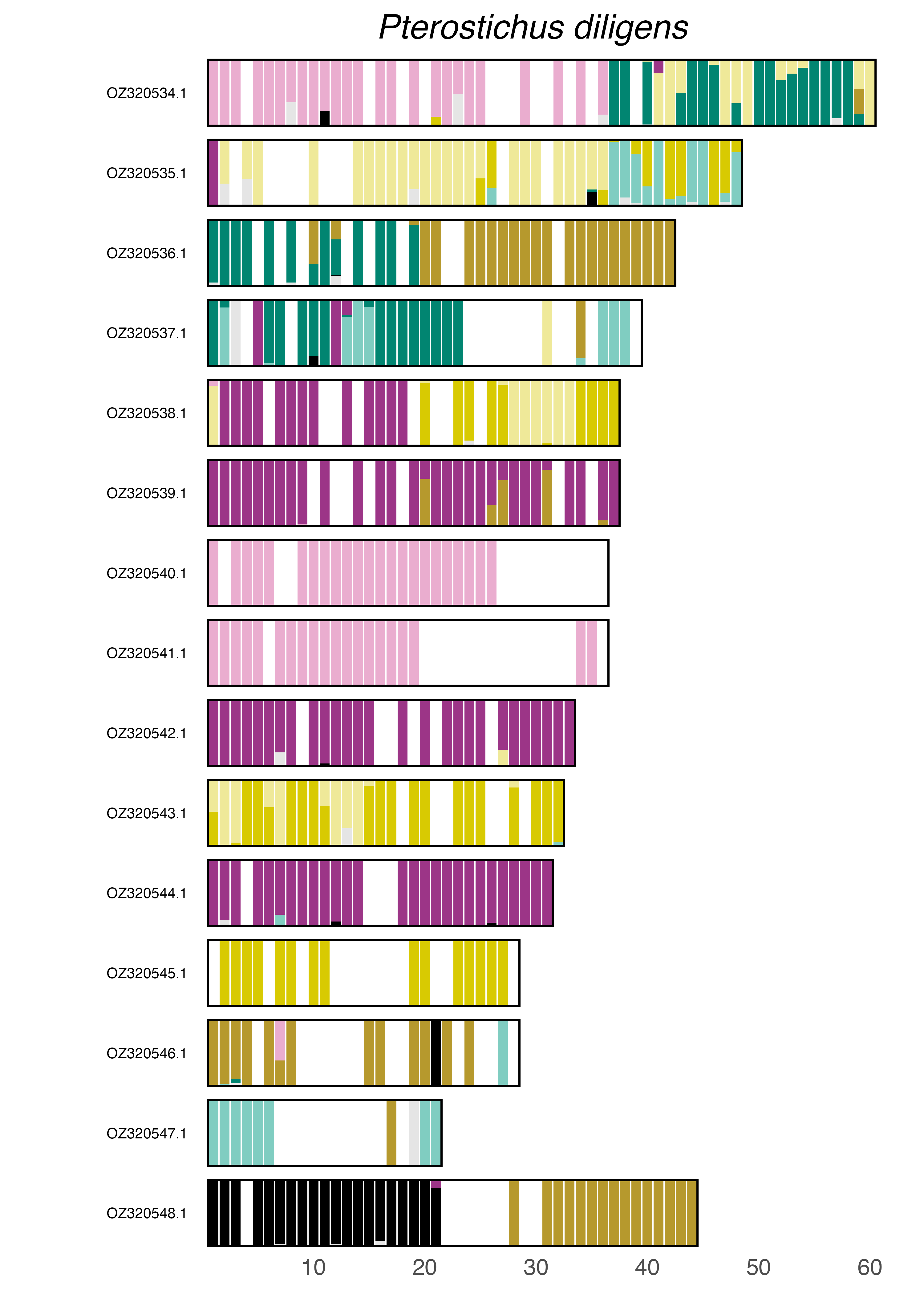

### plot.syngraph_windowbp_1Mb_rel_vertical_chrsize_Pyrocoelia_pectoralis.png

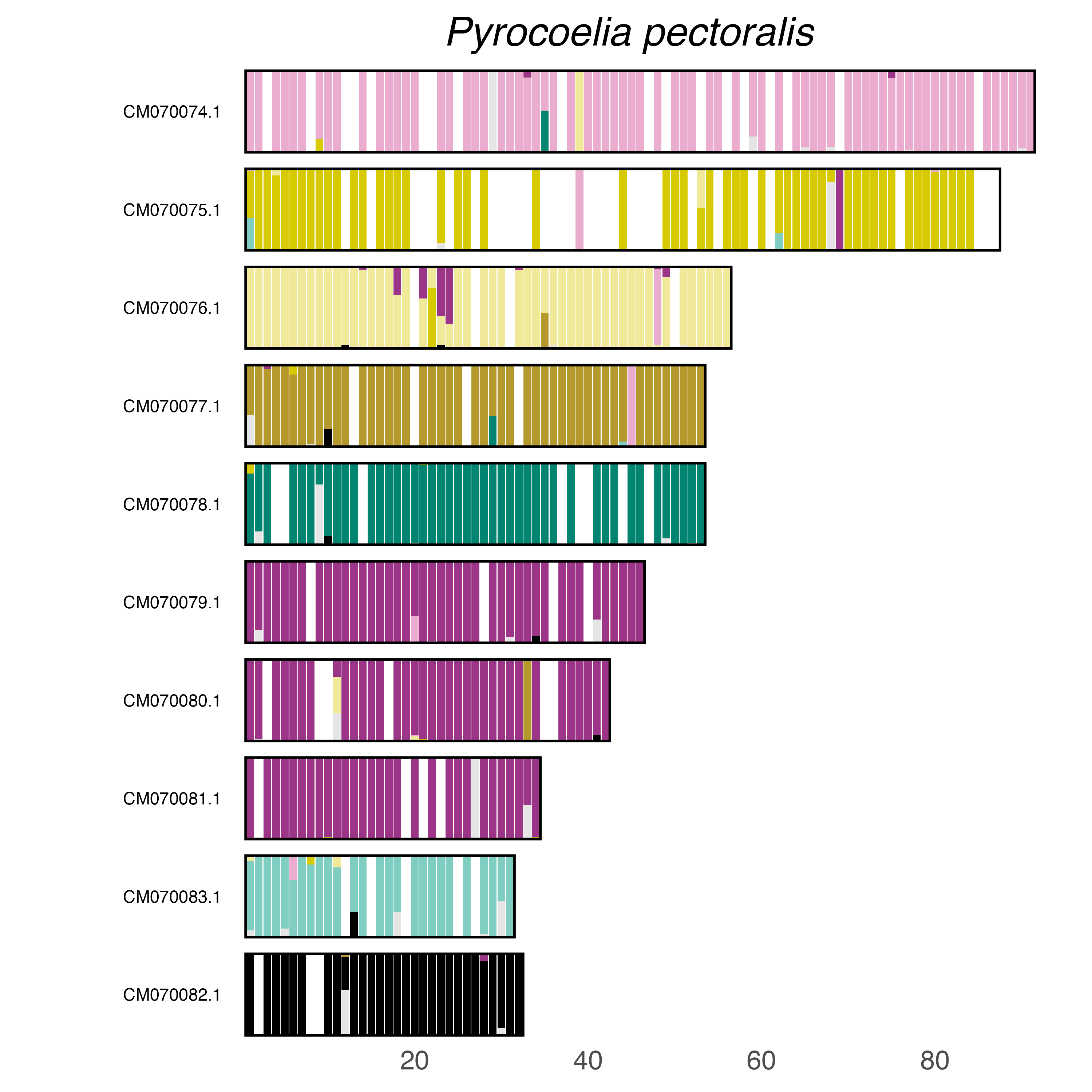

### plot.syngraph_windowbp_1Mb_rel_vertical_chrsize_Quedius_lateralis.png

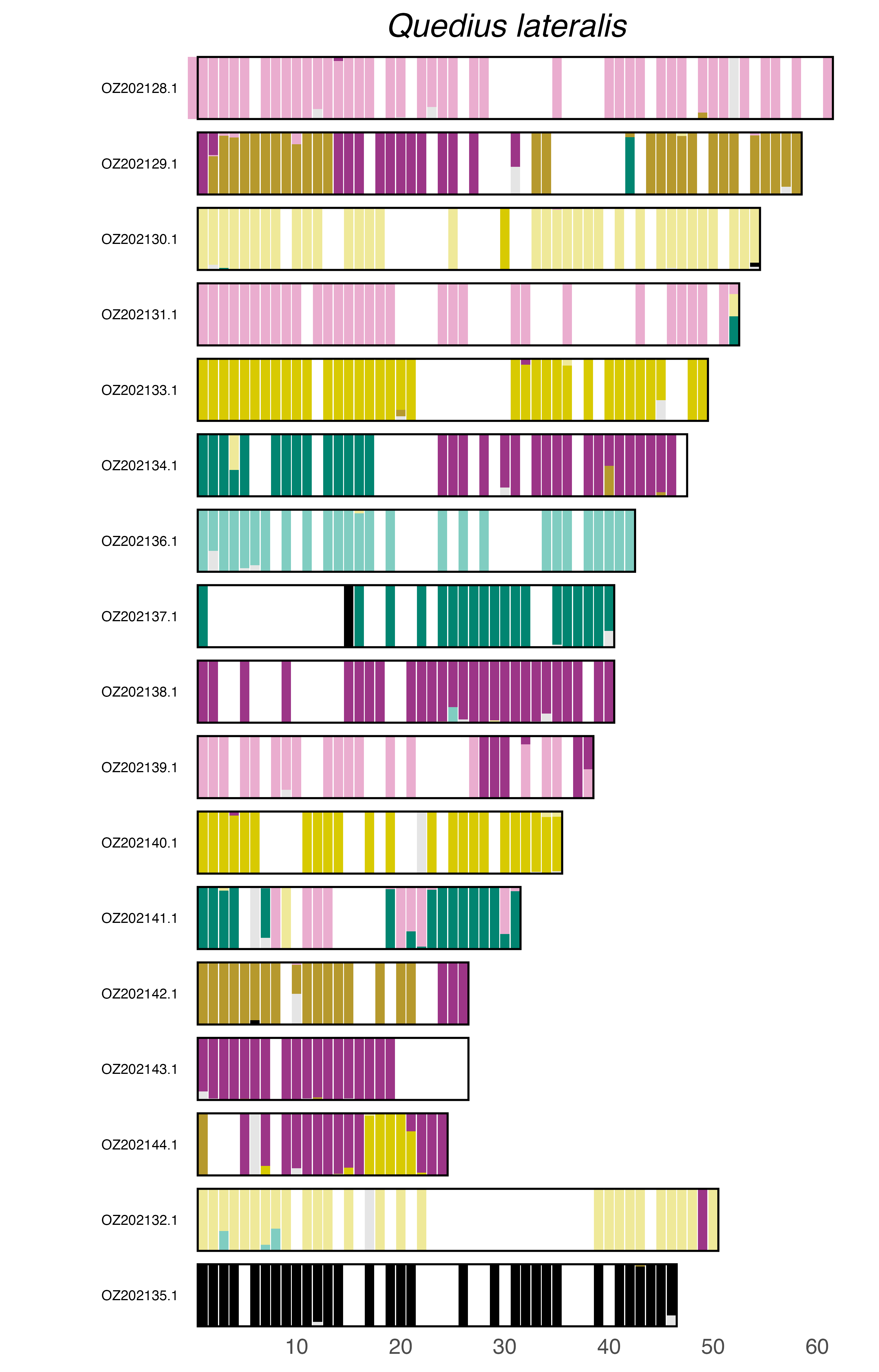

### plot.syngraph_windowbp_1Mb_rel_vertical_chrsize_Rhyzopertha_dominica.png

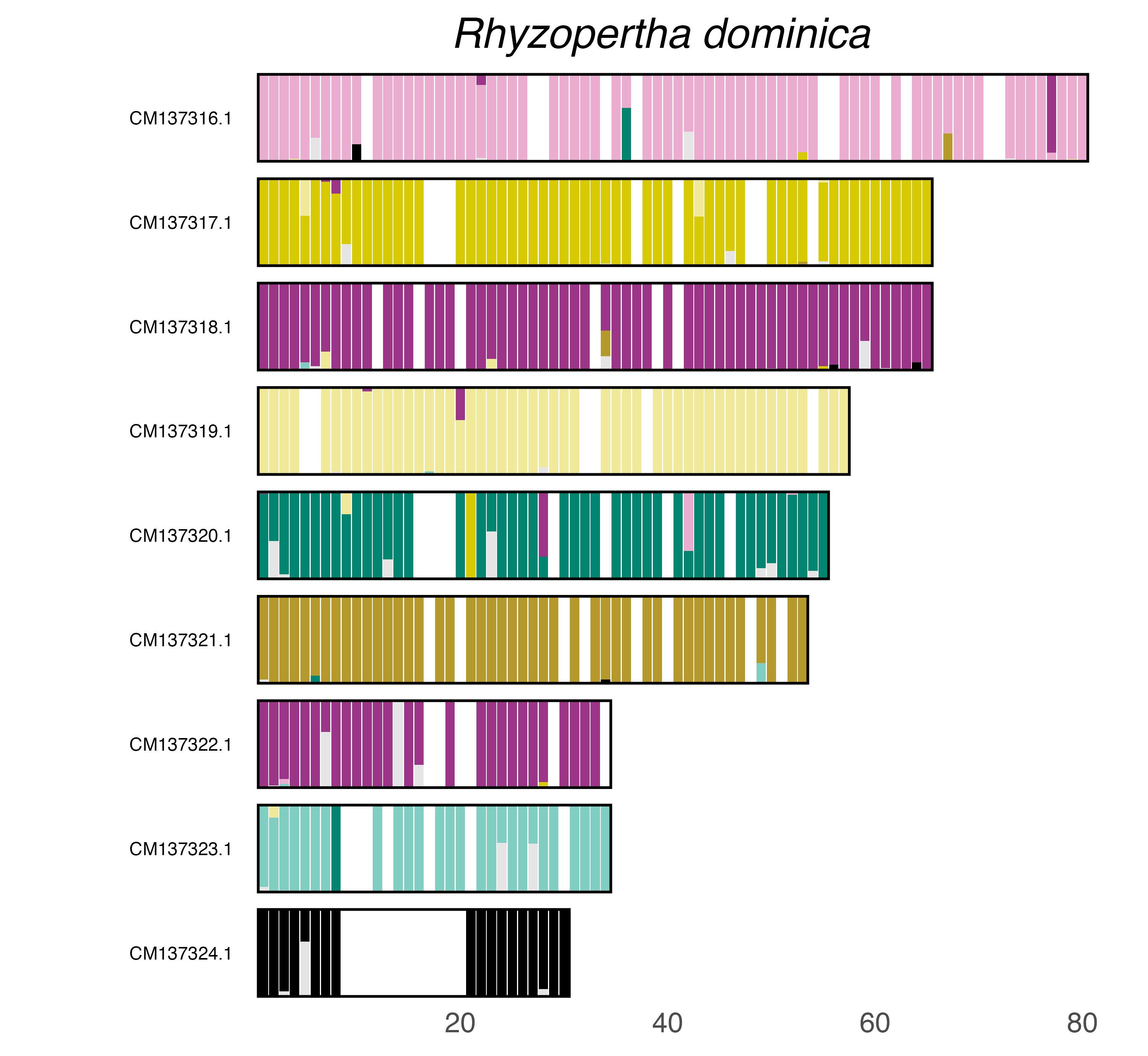

### plot.syngraph_windowbp_1Mb_rel_vertical_chrsize_Sisyphus_schaefferi.png

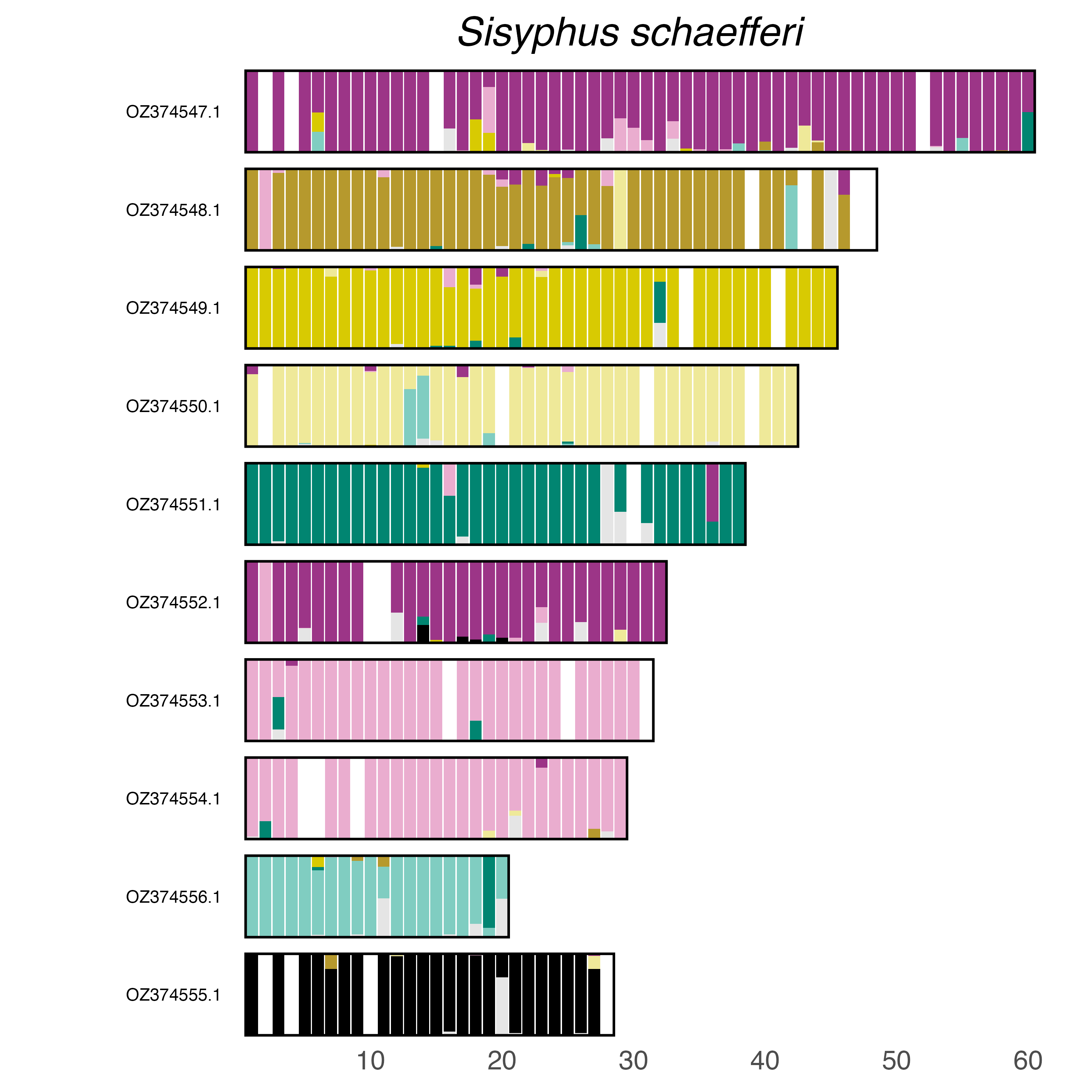

### plot.syngraph_windowbp_1Mb_rel_vertical_chrsize_Sisyra_terminalis.png

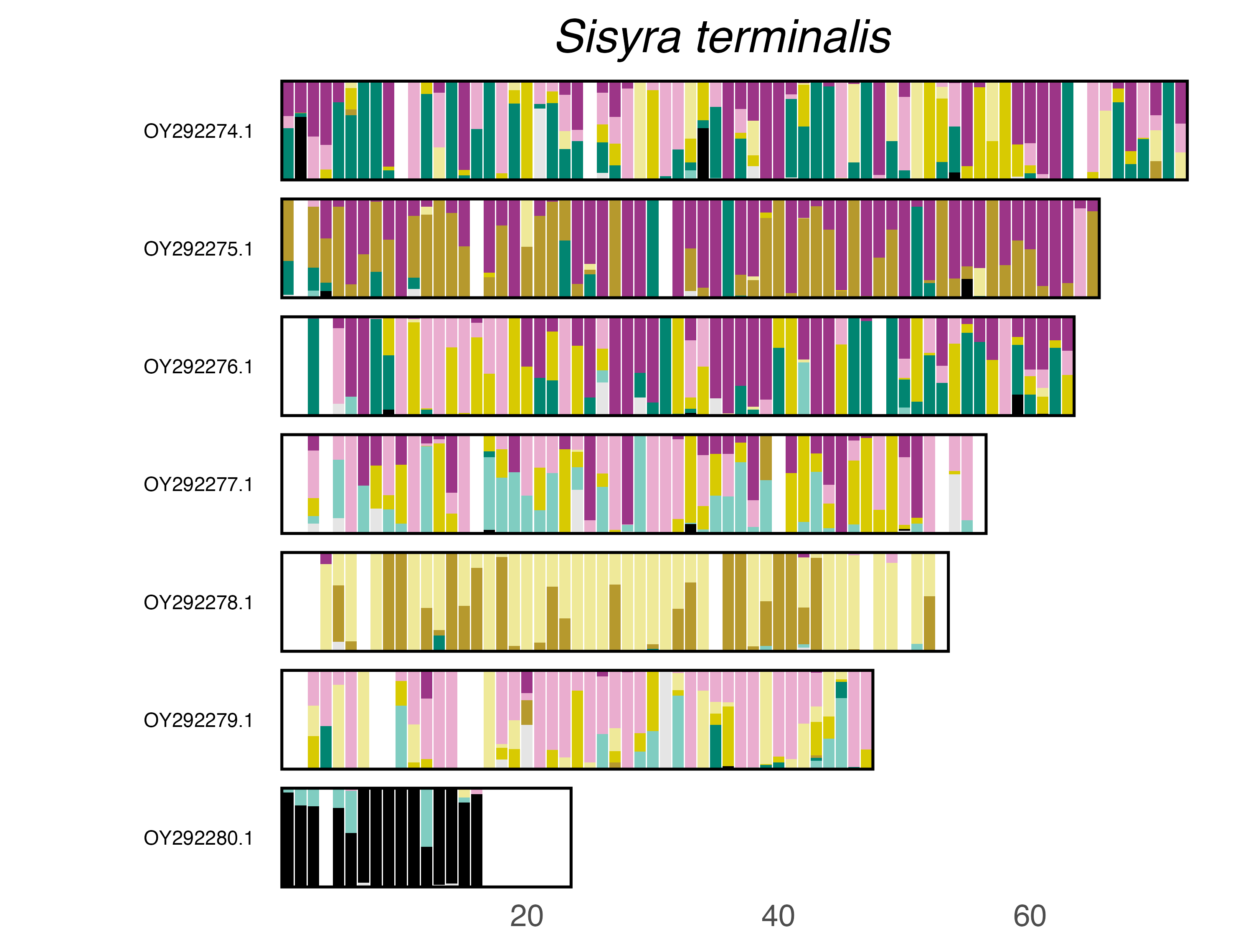

### plot.syngraph_windowbp_1Mb_rel_vertical_chrsize_Tachinus_rufipes.png

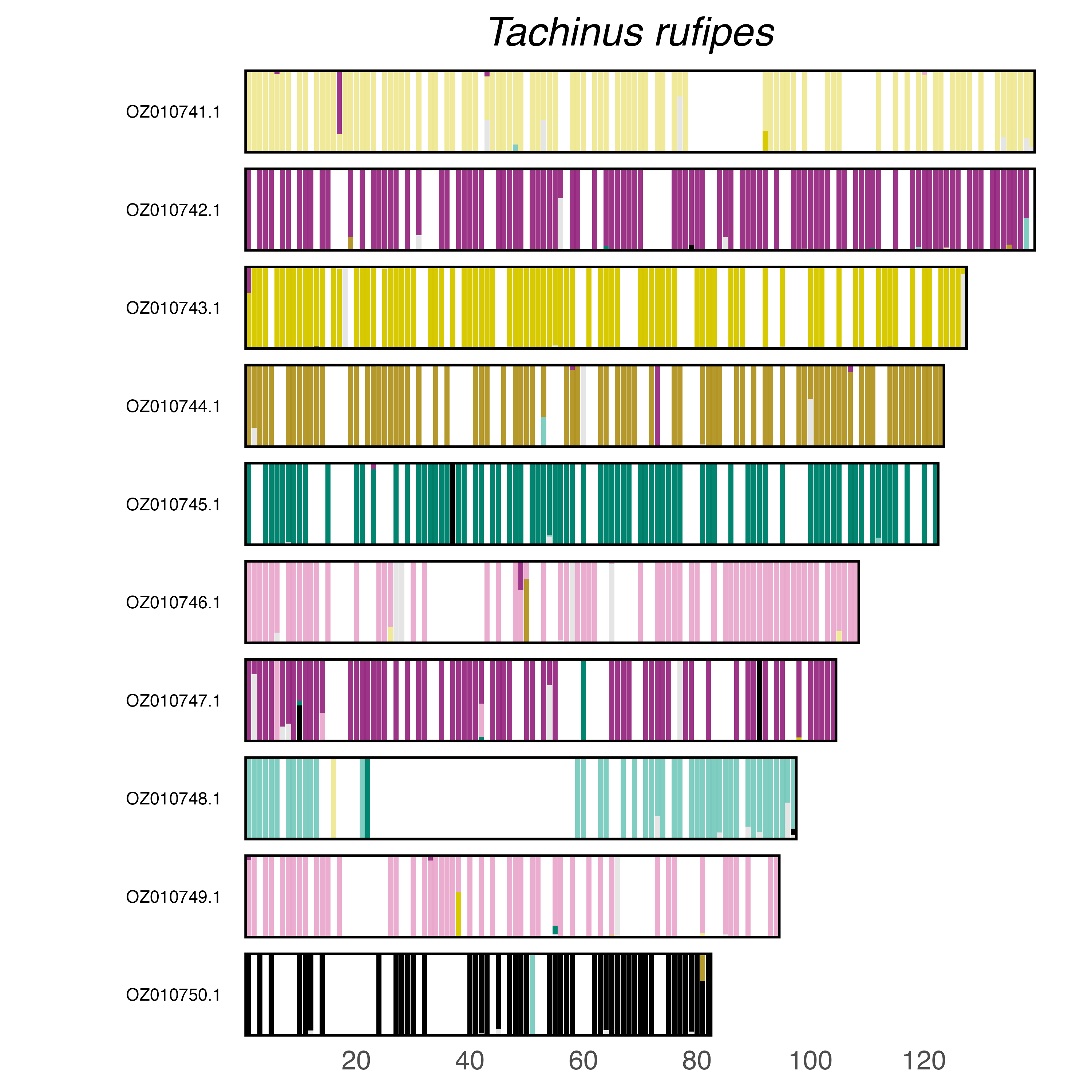

### plot.syngraph_windowbp_1Mb_rel_vertical_chrsize_Tachyporus_hypnorum.png

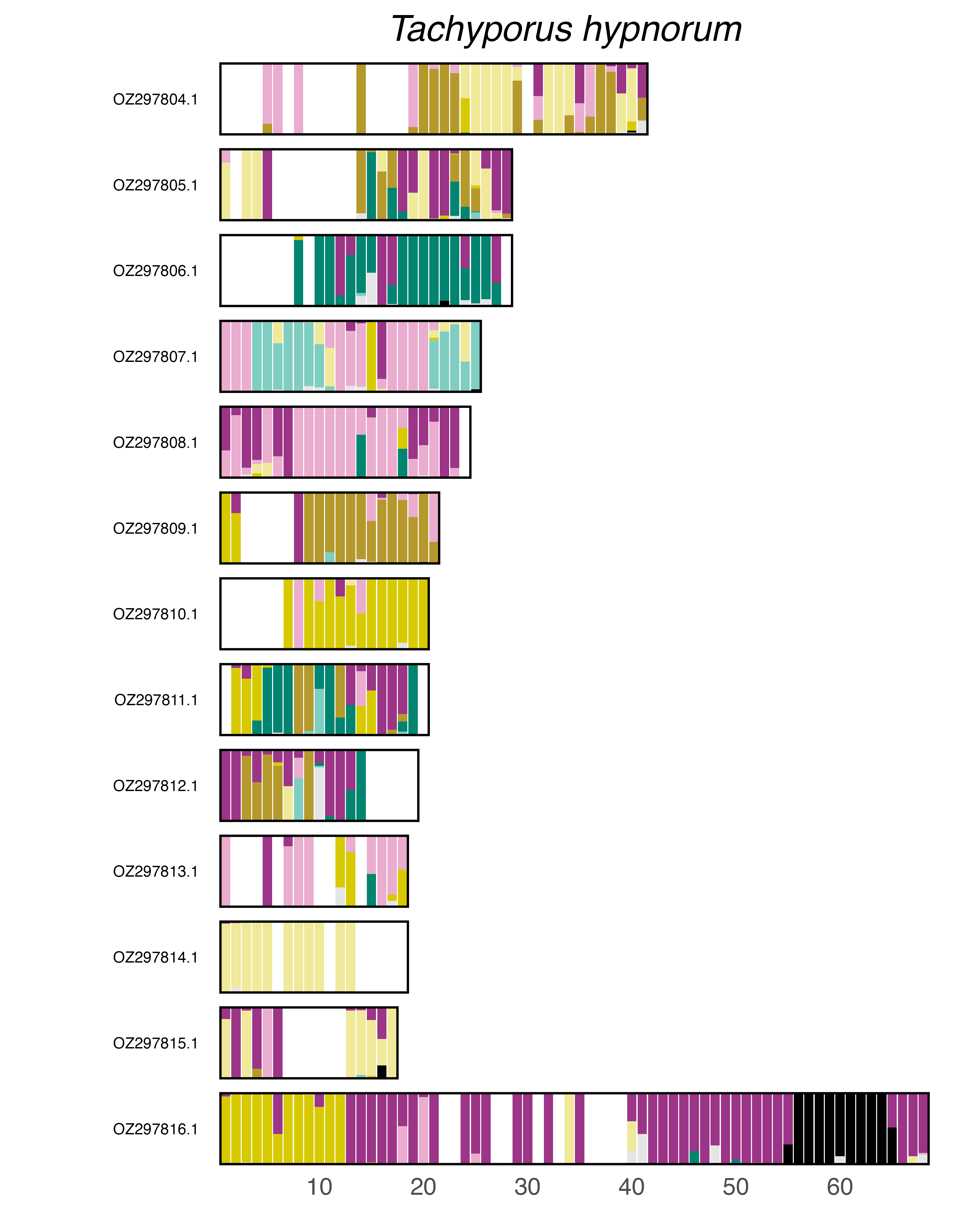

### plot.syngraph_windowbp_1Mb_rel_vertical_chrsize_Tenebrio_molitor.png

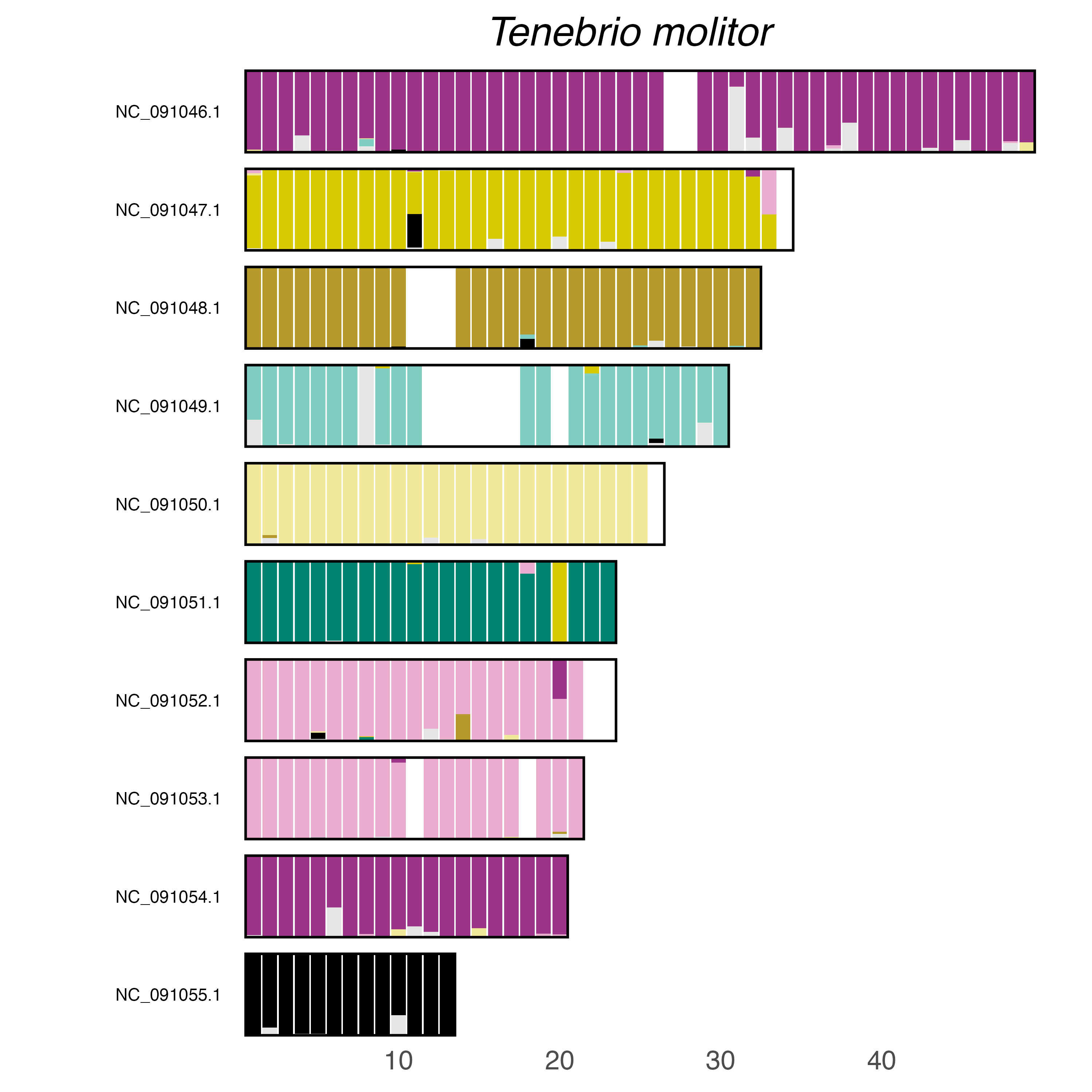

### plot.syngraph_windowbp_1Mb_rel_vertical_chrsize_Tetraopes_tetrophthalmus.png

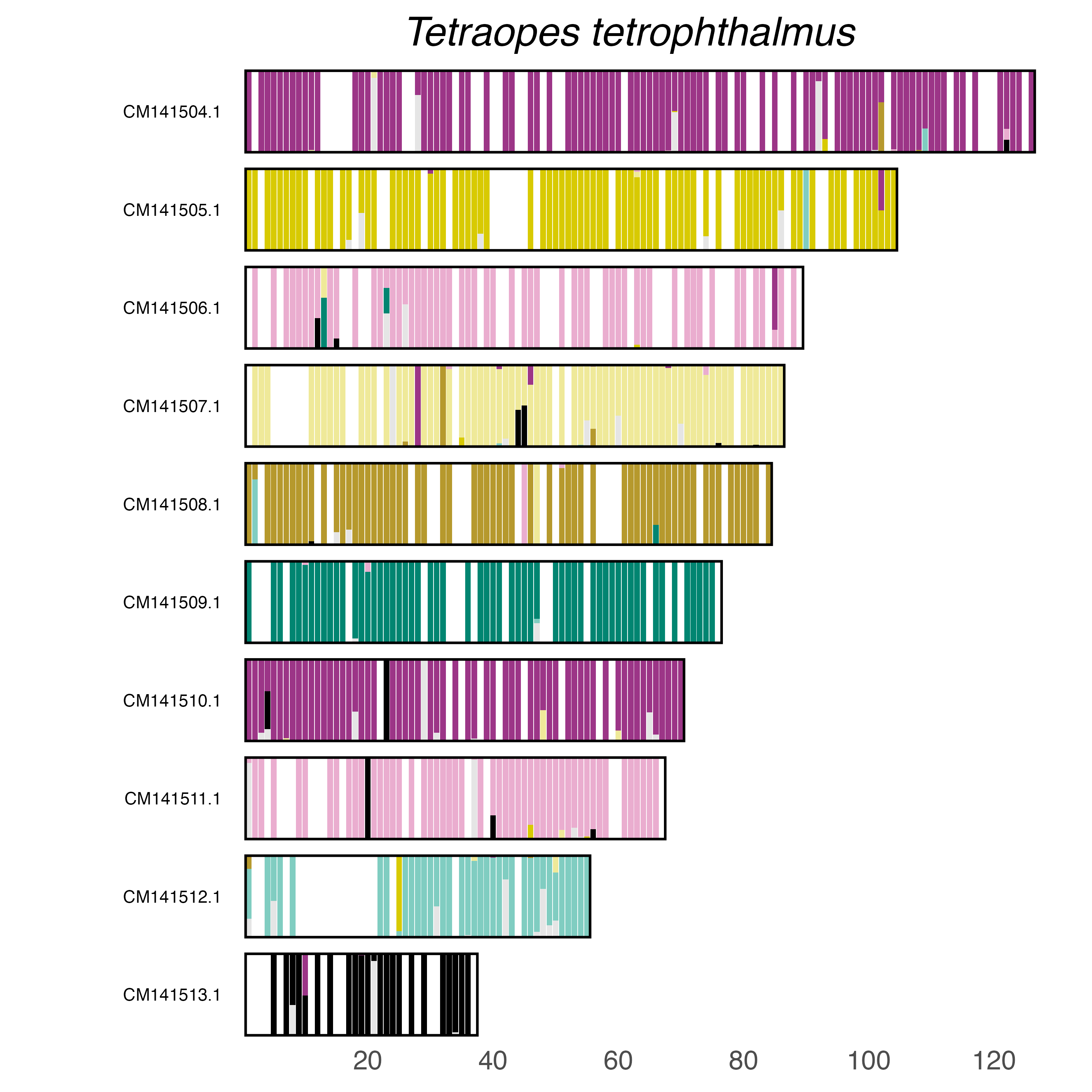

### plot.syngraph_windowbp_1Mb_rel_vertical_chrsize_Tomicus_yunnanensis.png

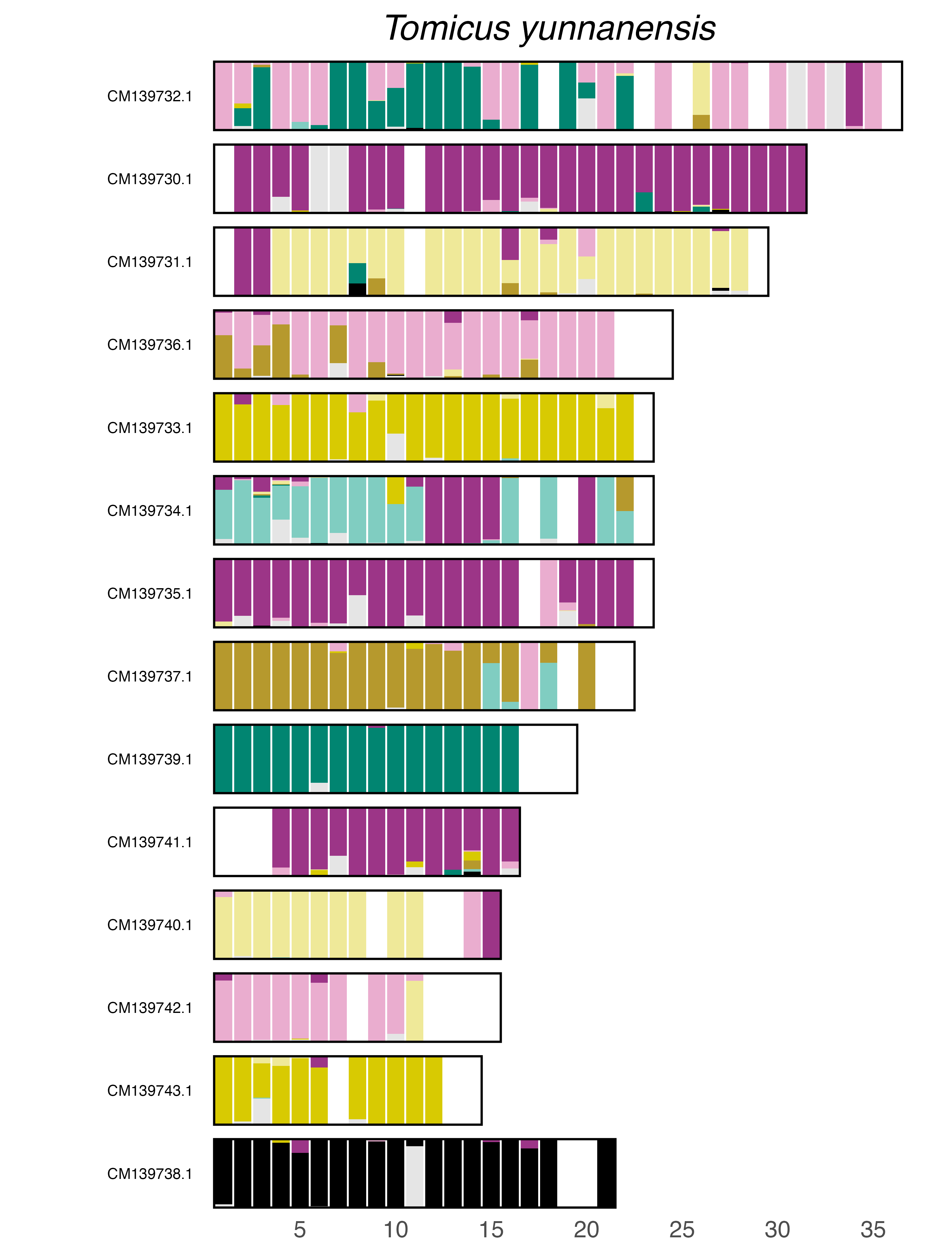

### plot.syngraph_windowbp_1Mb_rel_vertical_chrsize_Tribolium_confusum.png

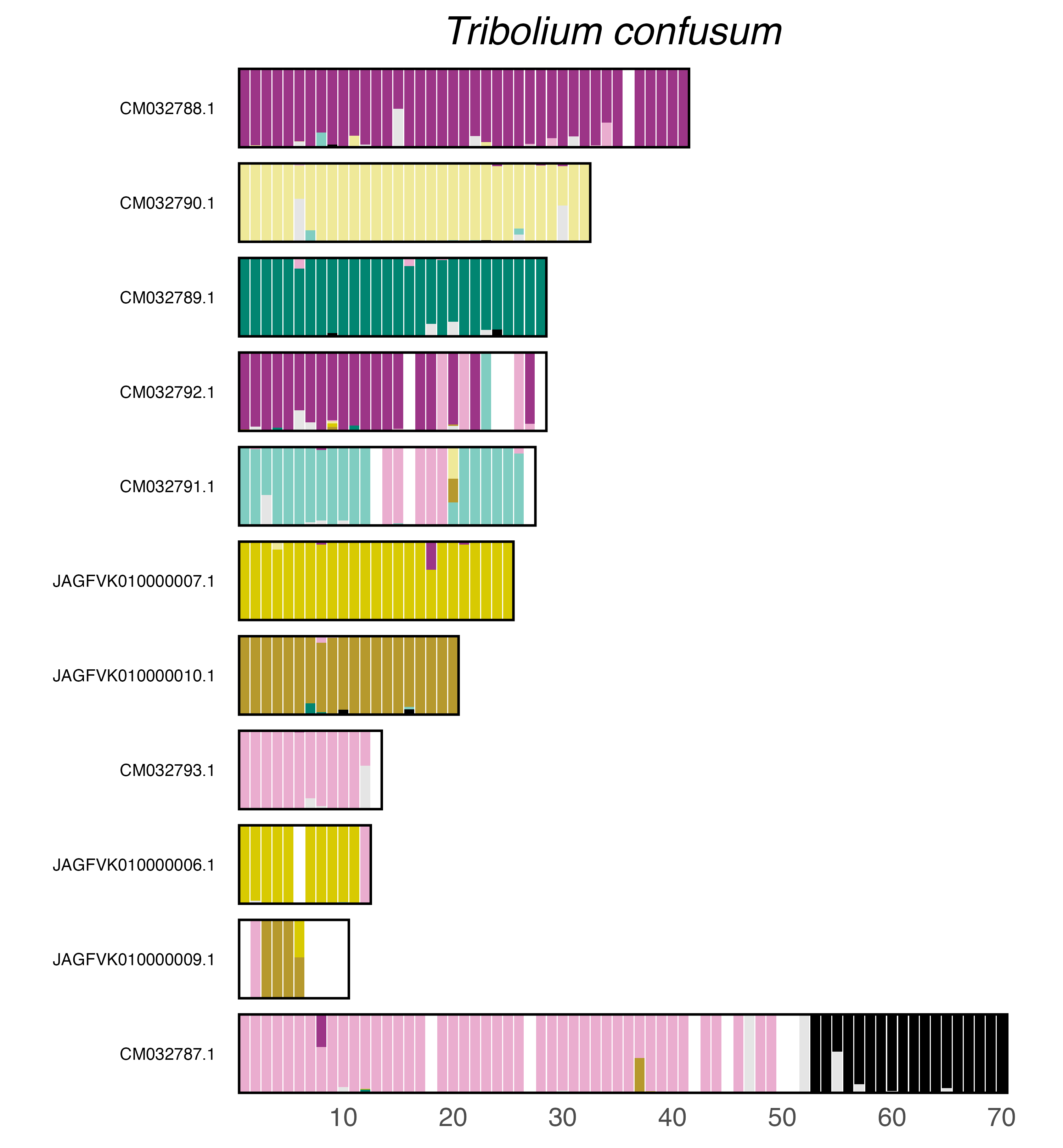

### plot.syngraph_windowbp_1Mb_rel_vertical_chrsize_Xestobium_rufovillosum.png

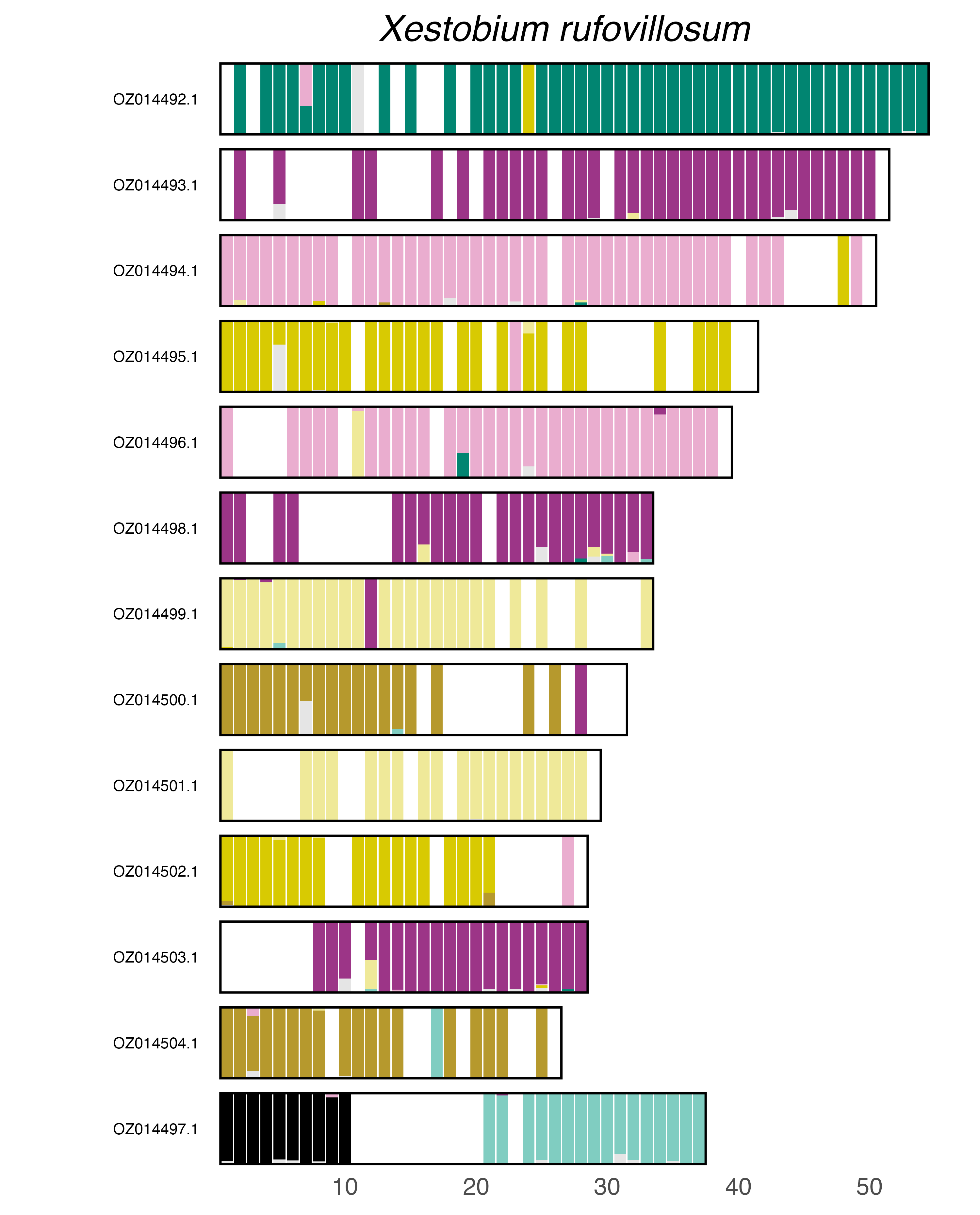

### Supplementary data S1

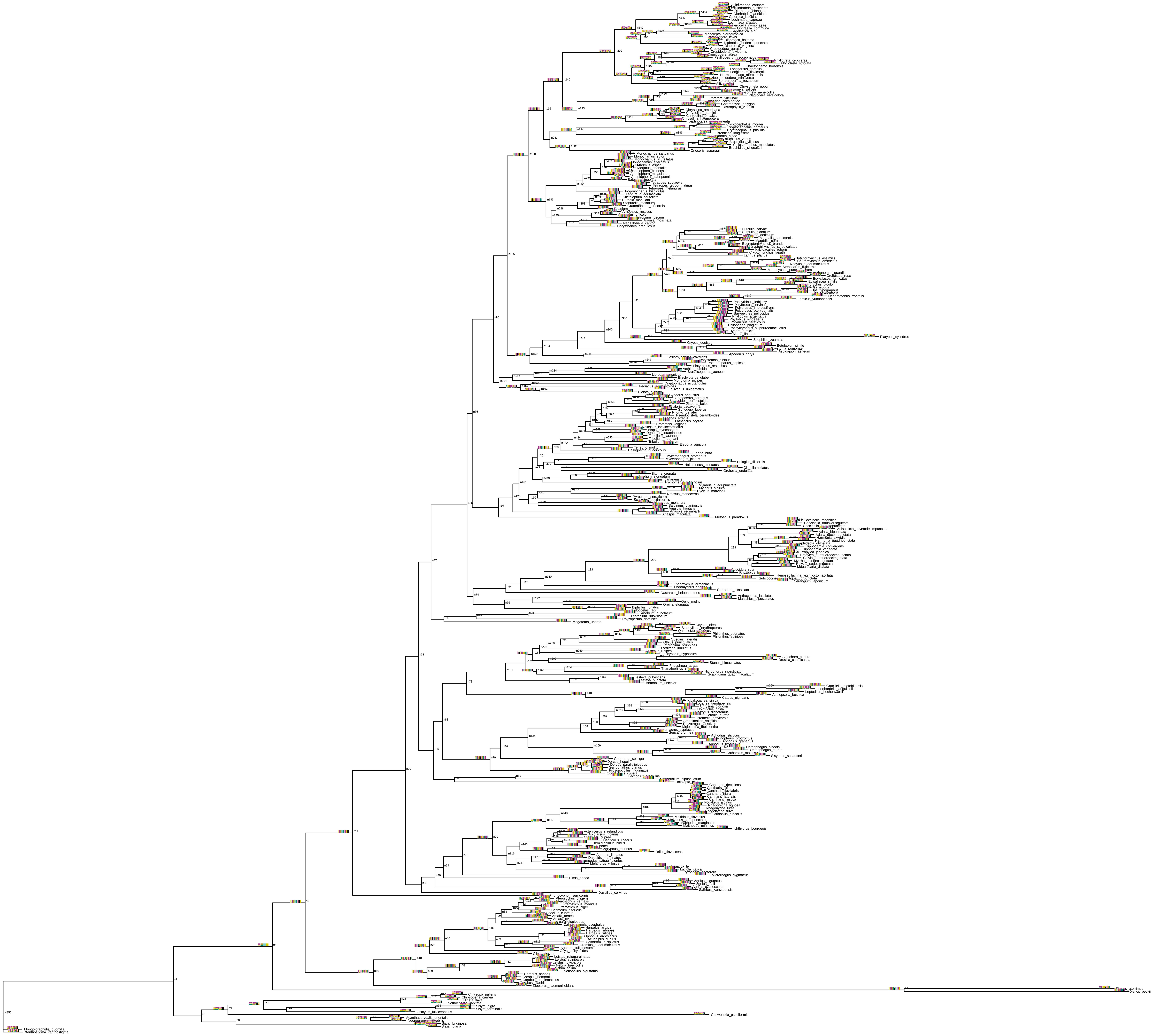
