## Supplementary figures and images for "338 coleopteran genomes reveal exceptional rearrangement variation compared to other insect orders"

### plot.syngraph_windowbp_1Mb_rel_vertical_chrsize_Philonthus_spinipes.png

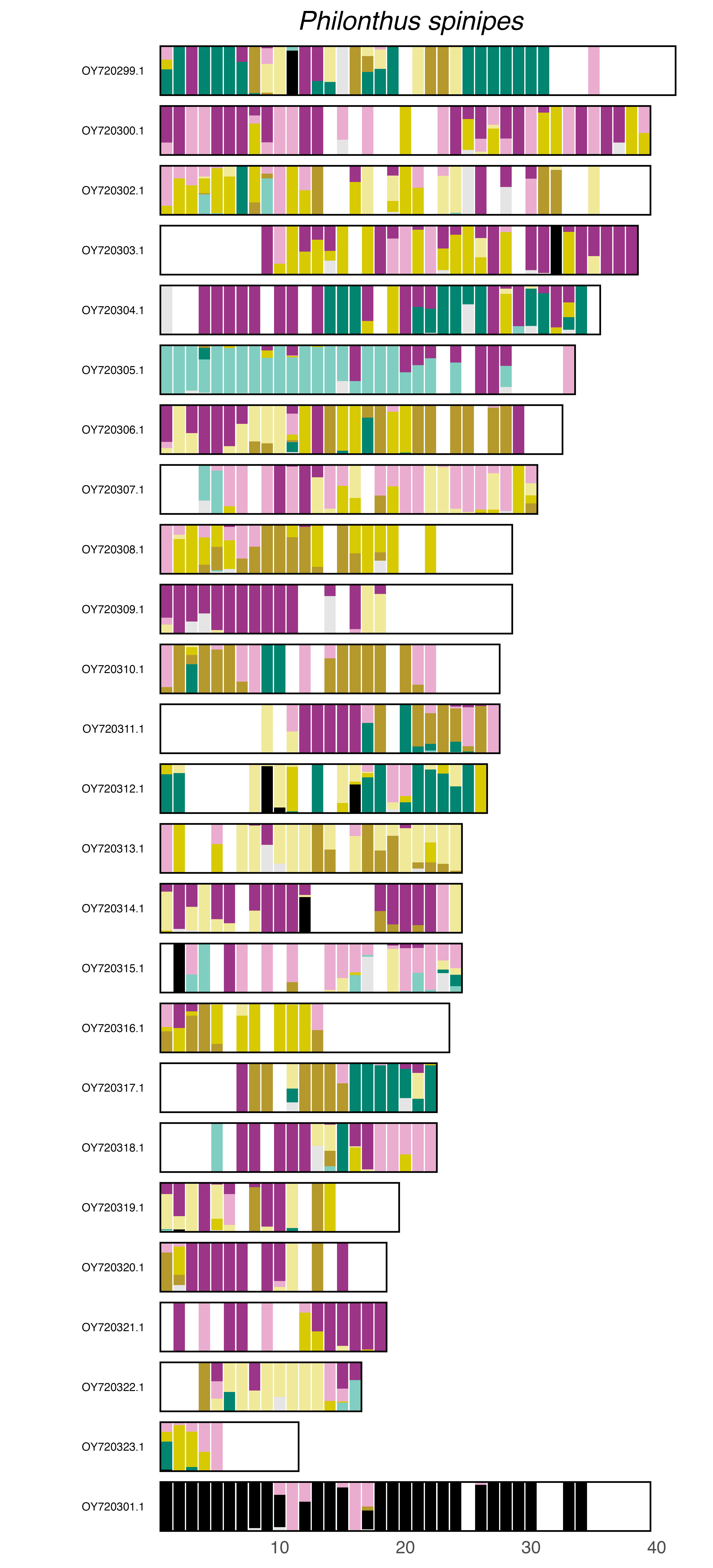
