## Supplementary figures and images for "338 coleopteran genomes reveal exceptional rearrangement variation compared to other insect orders"

### plot.syngraph_windowbp_1Mb_rel_vertical_chrsize_Abax_parallelepipedus.png

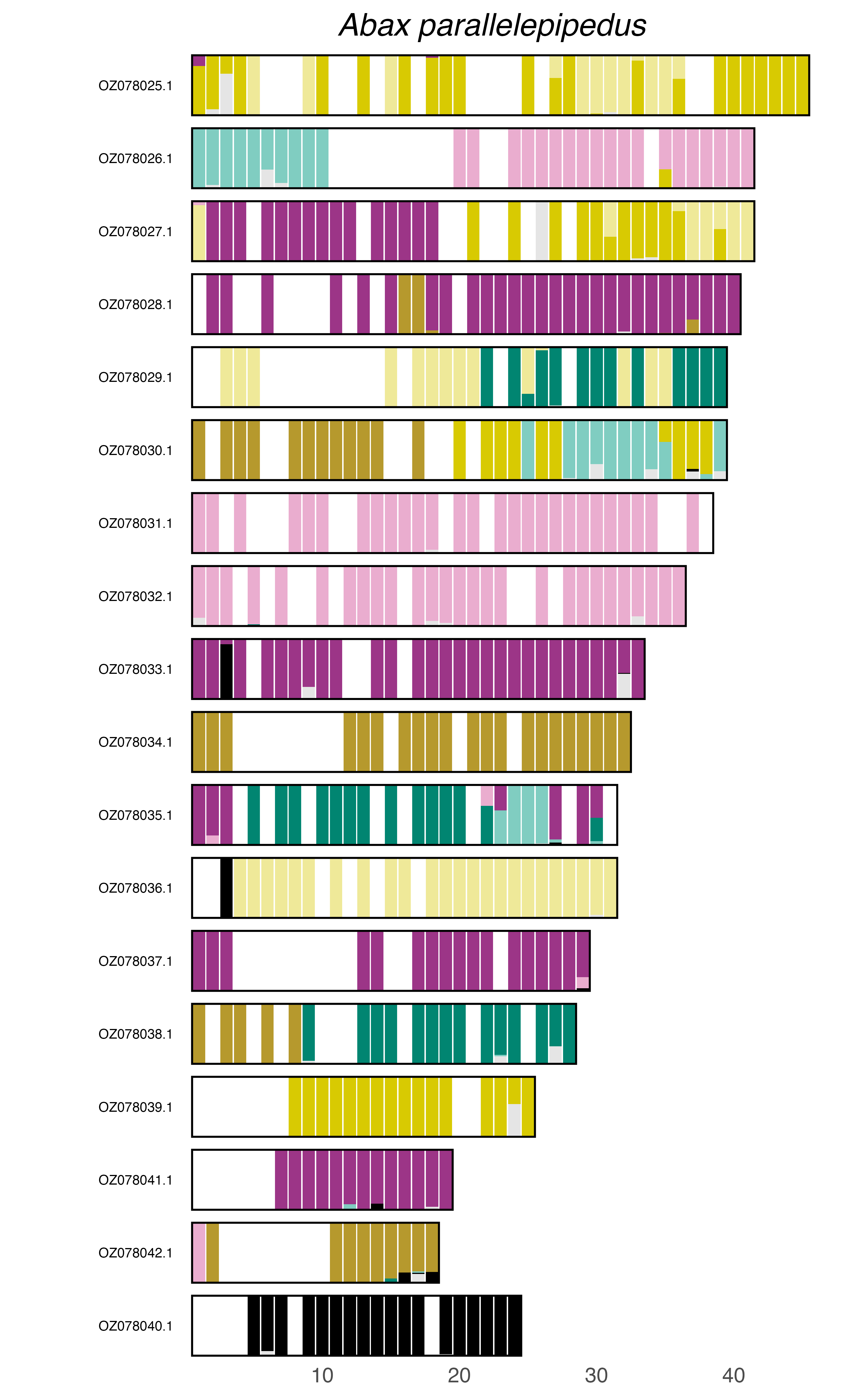

### plot.syngraph_windowbp_1Mb_rel_vertical_chrsize_Agriotes_lineatus.png

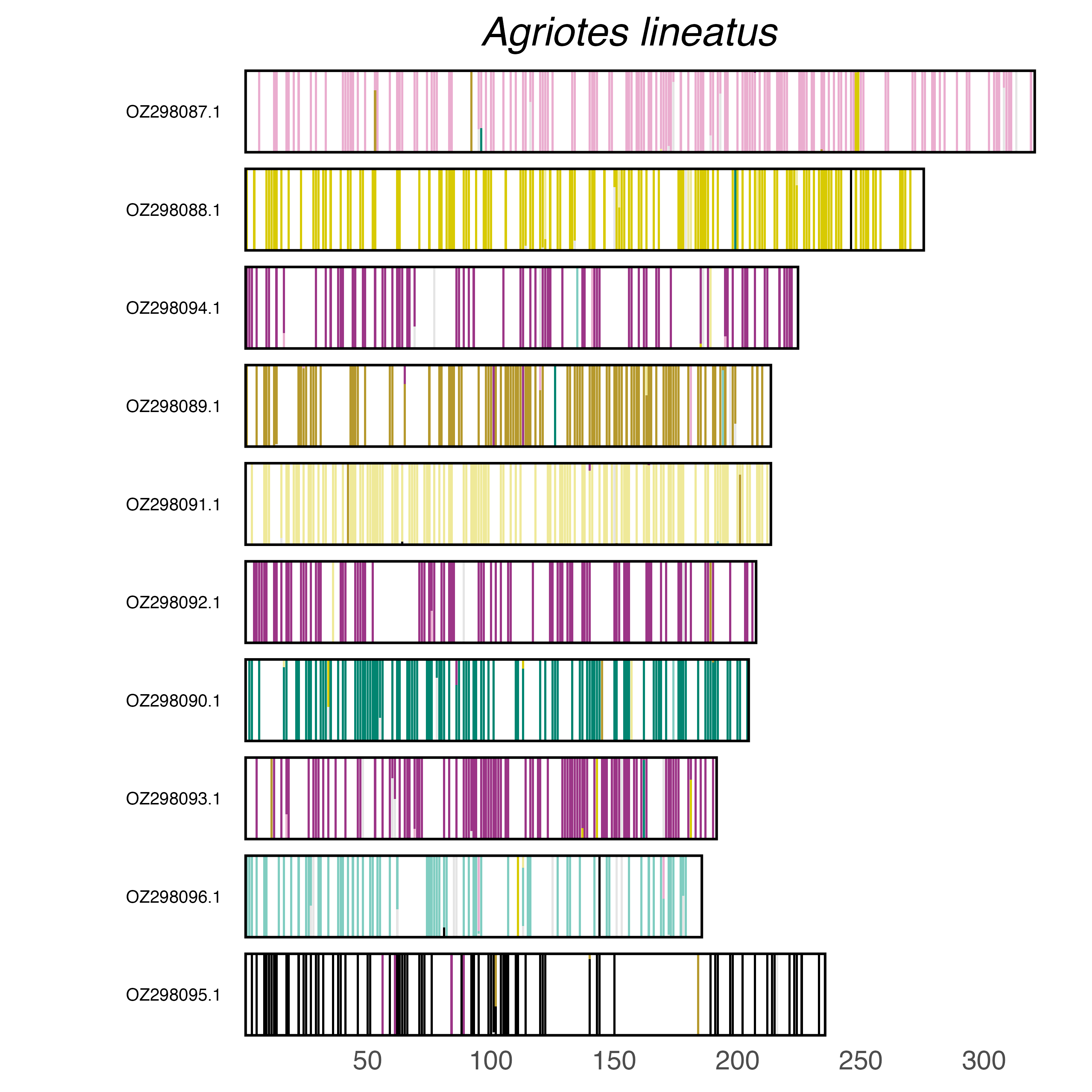

### plot.syngraph_windowbp_1Mb_rel_vertical_chrsize_Agrypnus_murinus.png

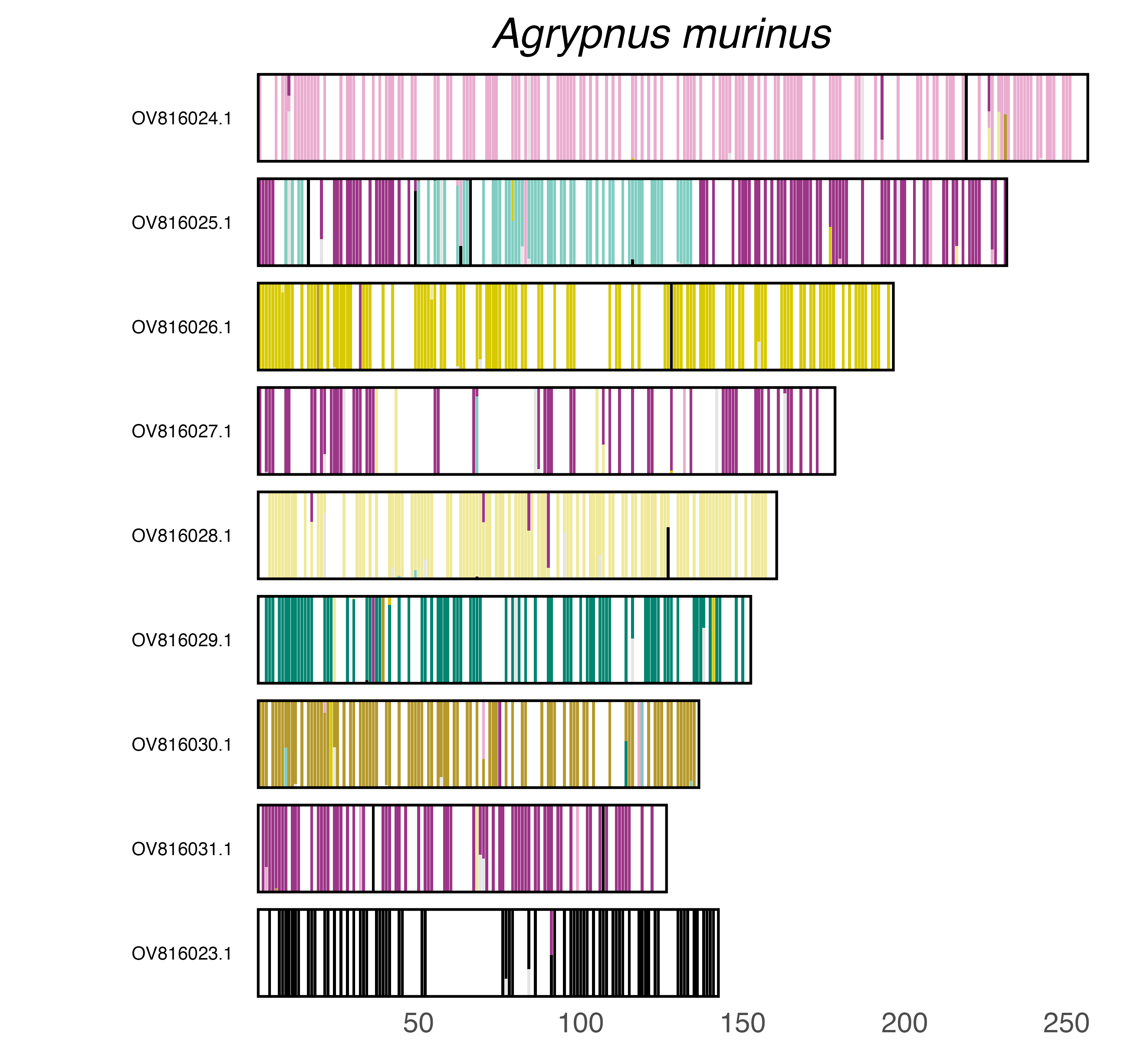

### plot.syngraph_windowbp_1Mb_rel_vertical_chrsize_Aleochara_curtula.png

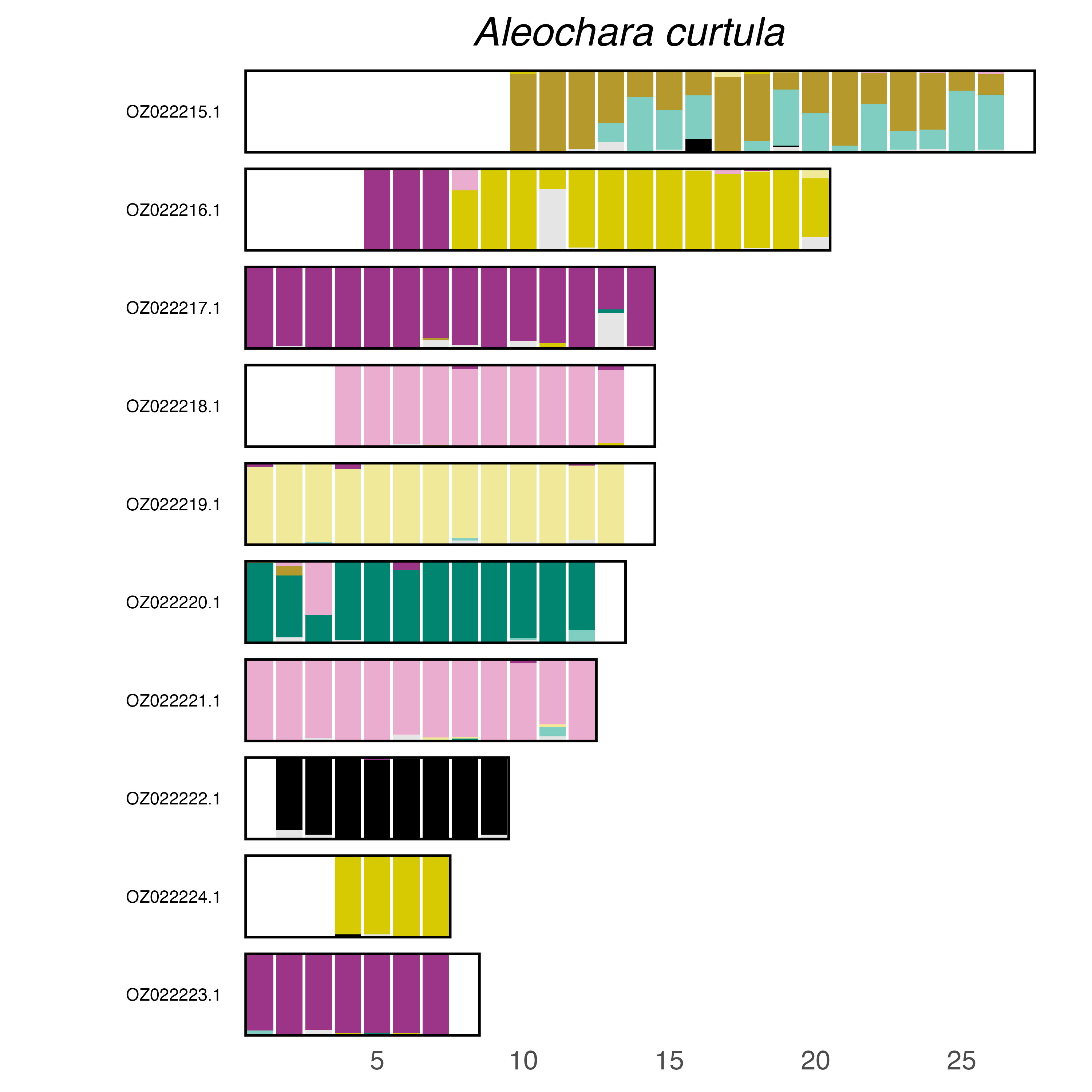

### plot.syngraph_windowbp_1Mb_rel_vertical_chrsize_Amara_ovata.png

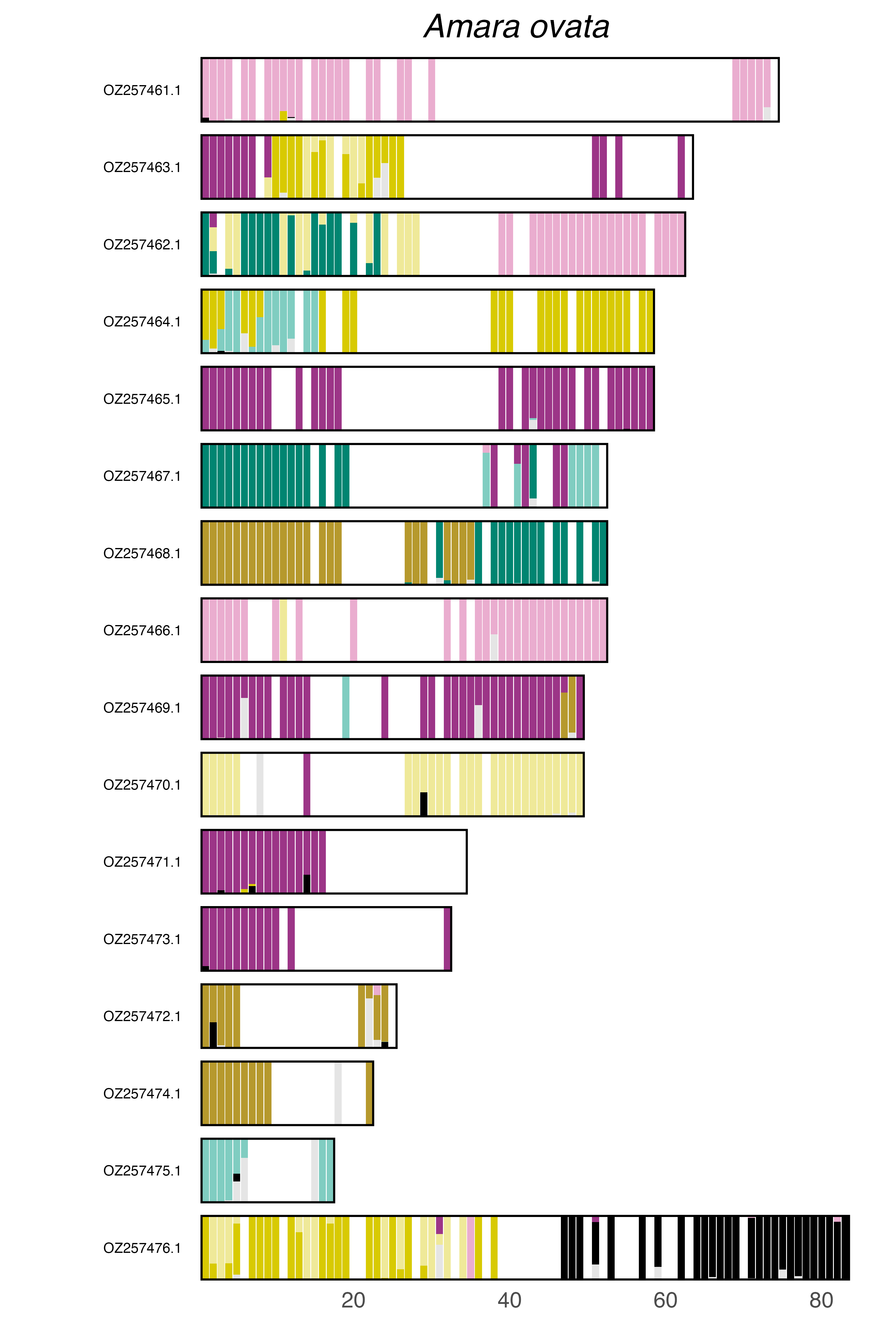

### plot.syngraph_windowbp_1Mb_rel_vertical_chrsize_Anaspis_frontalis.png

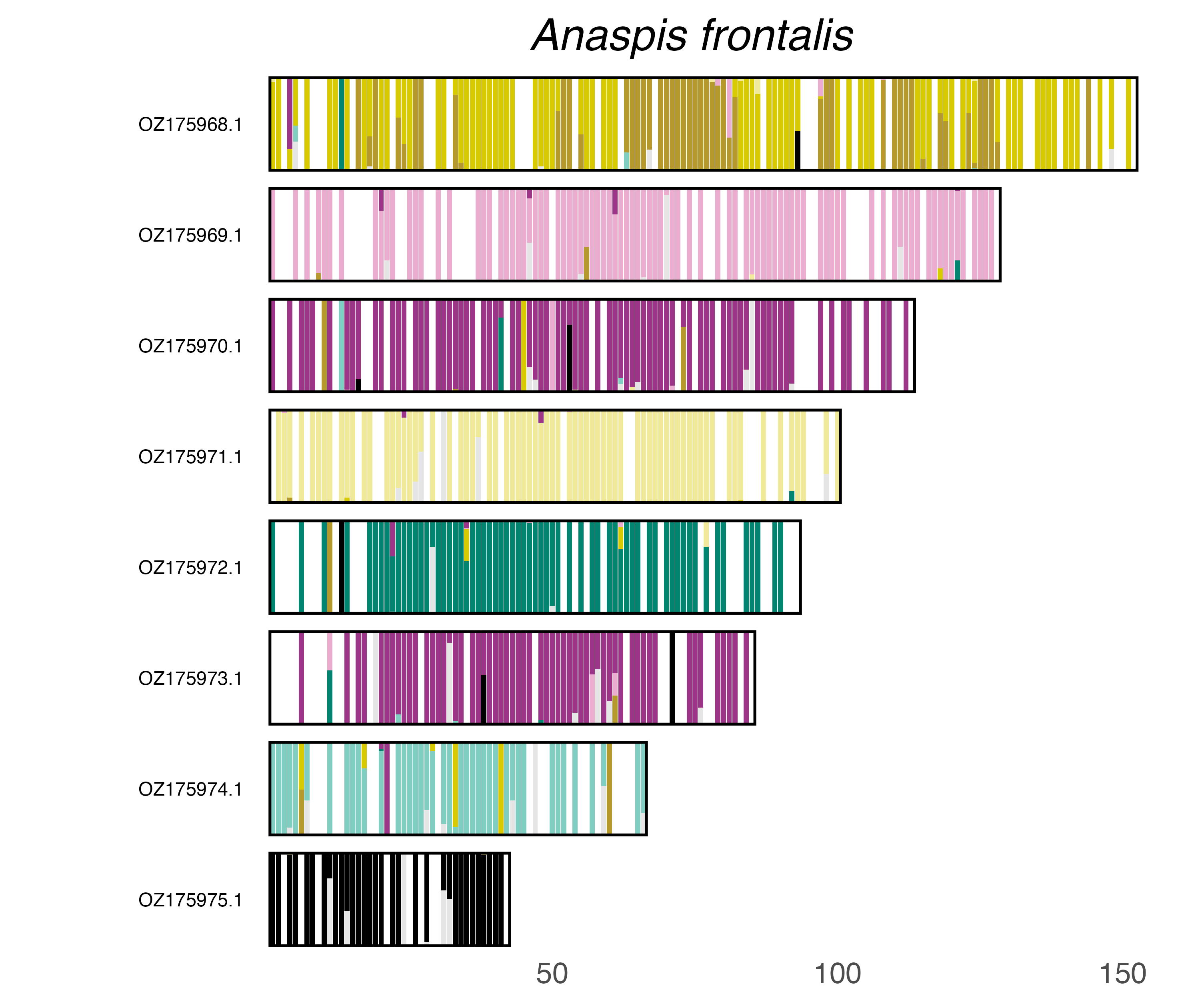

### plot.syngraph_windowbp_1Mb_rel_vertical_chrsize_Anaspis_regimbarti.png

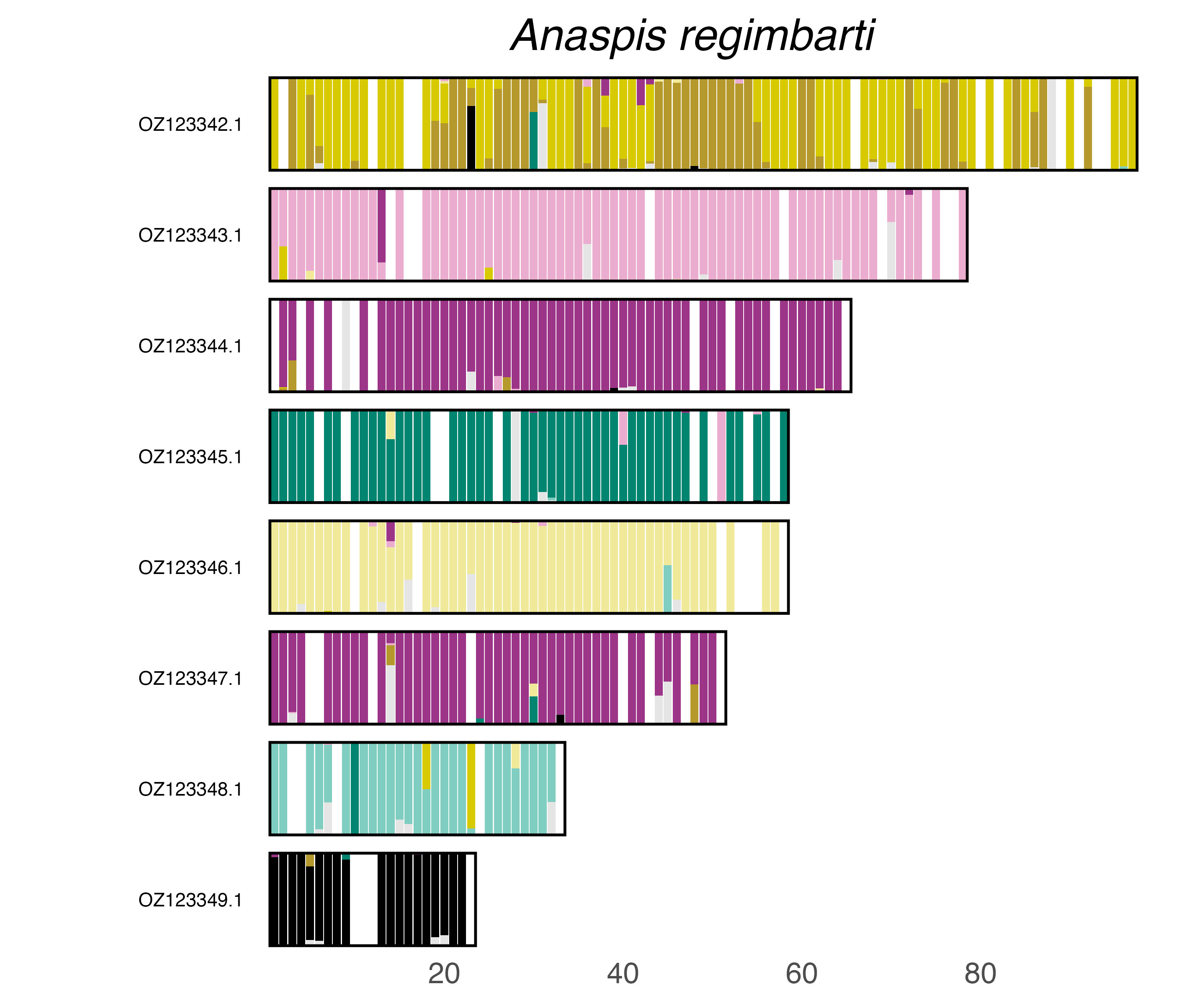

### plot.syngraph_windowbp_1Mb_rel_vertical_chrsize_Anoplophora_glabripennis.png

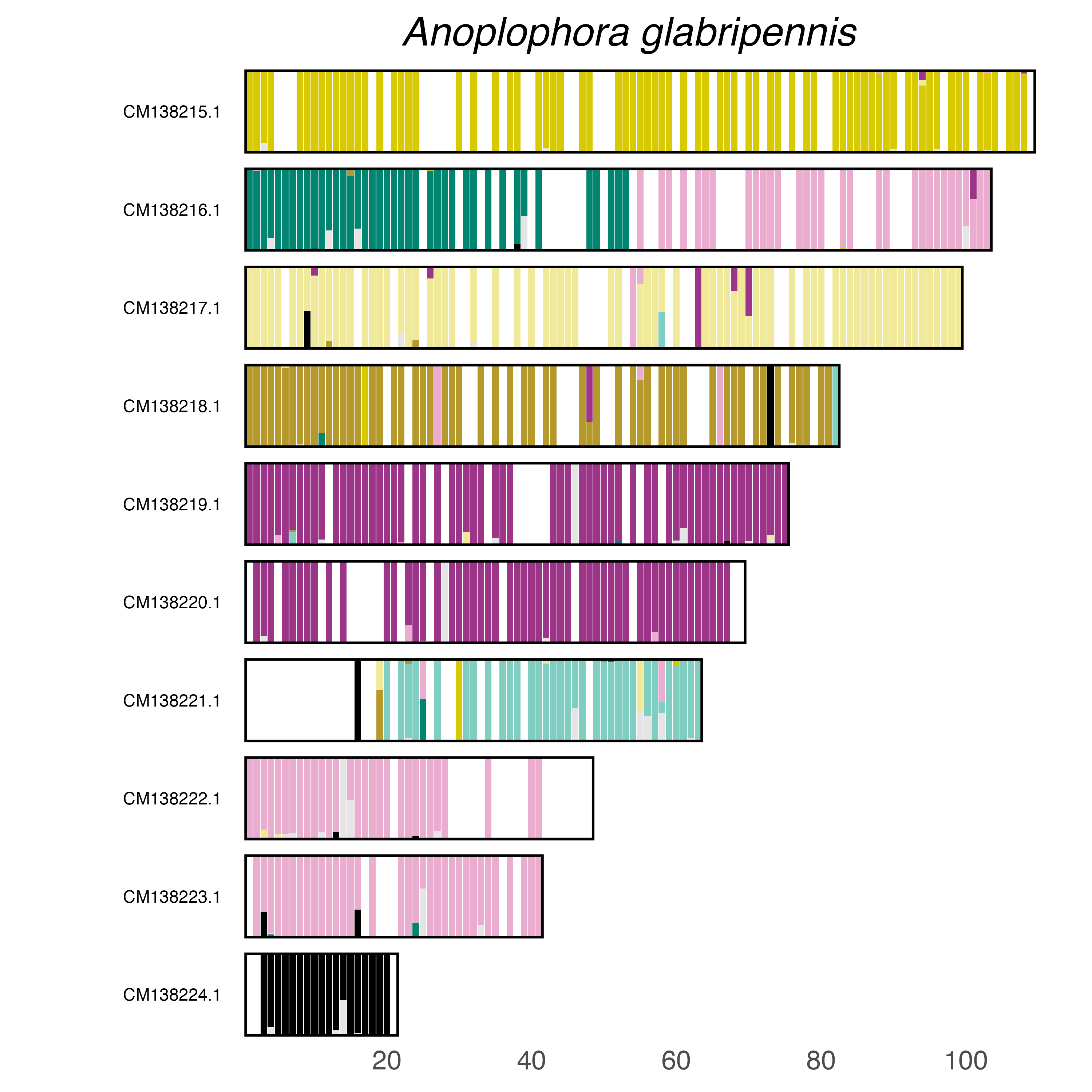

### plot.syngraph_windowbp_1Mb_rel_vertical_chrsize_Aphodius_granarius.png

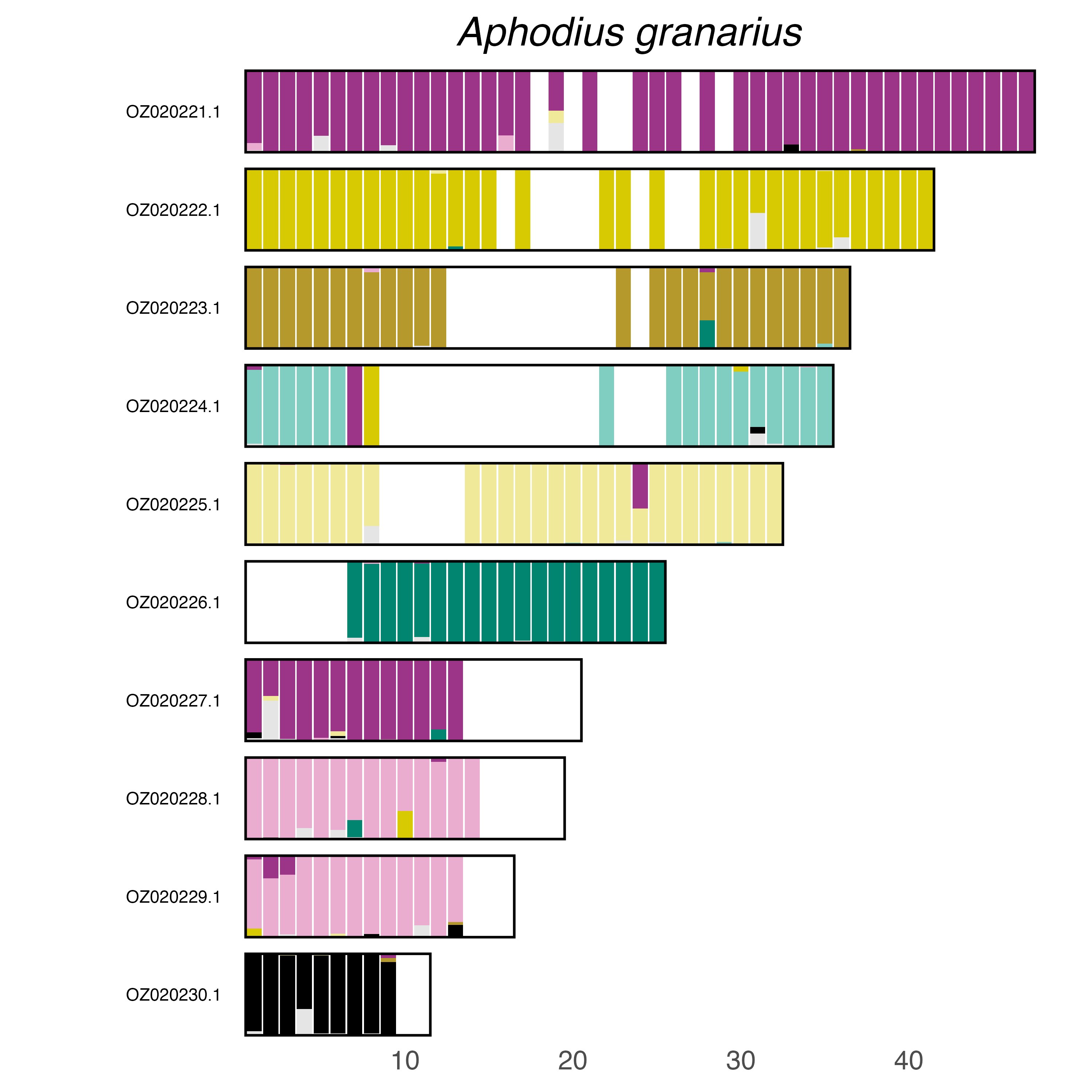

### plot.syngraph_windowbp_1Mb_rel_vertical_chrsize_Aspidapion_aeneum.png

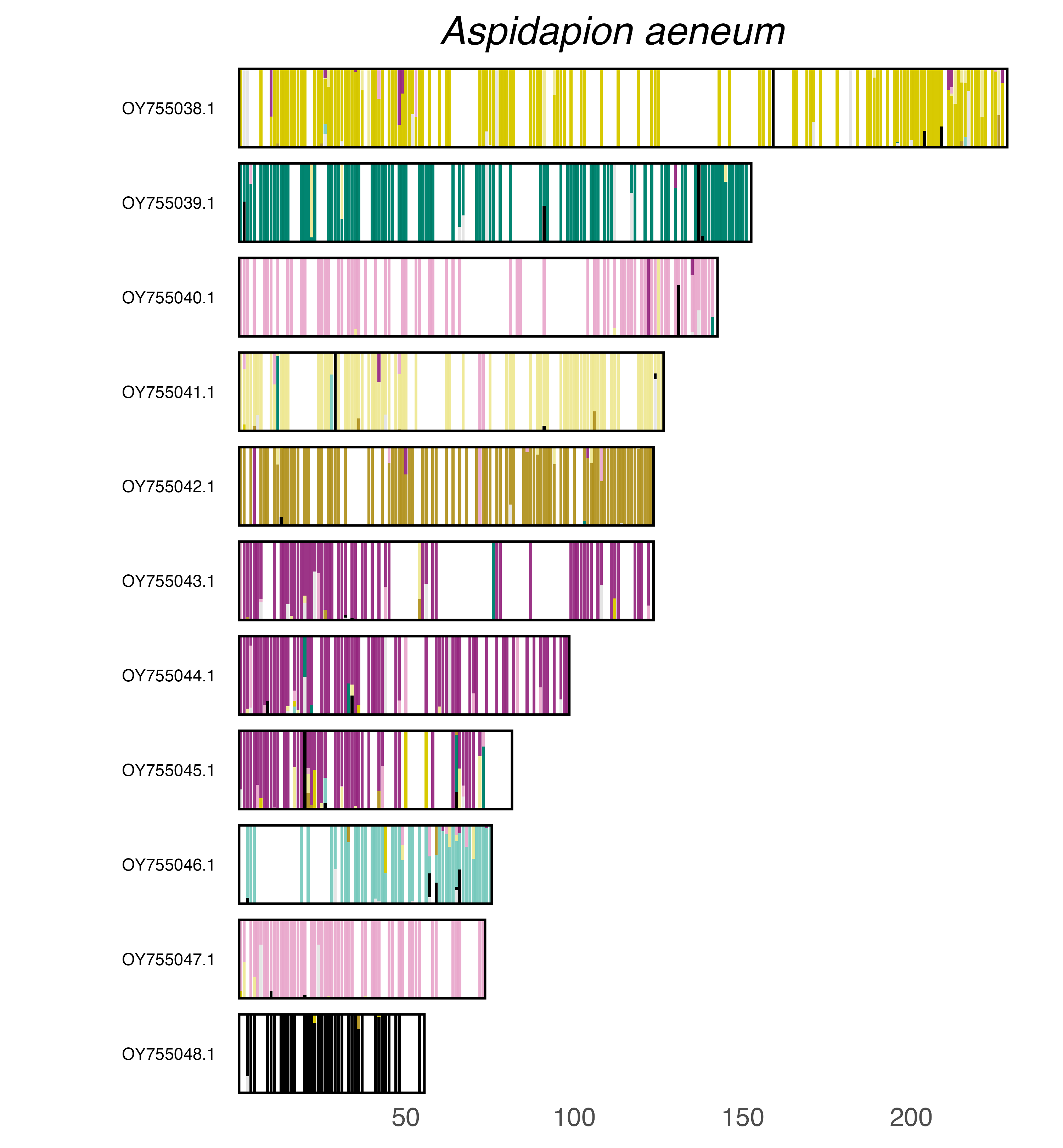

### plot.syngraph_windowbp_1Mb_rel_vertical_chrsize_Aulacophora_lewisii.png

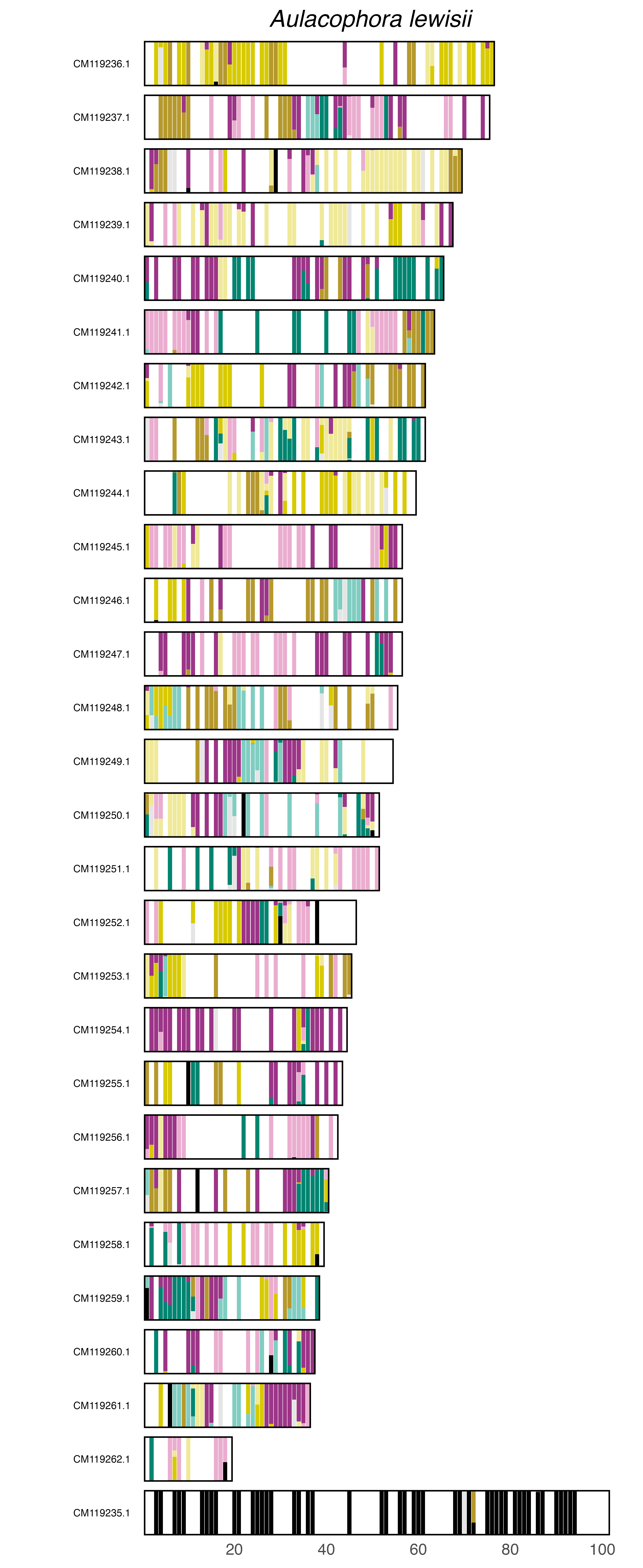

### plot.syngraph_windowbp_1Mb_rel_vertical_chrsize_Betulapion_simile.png

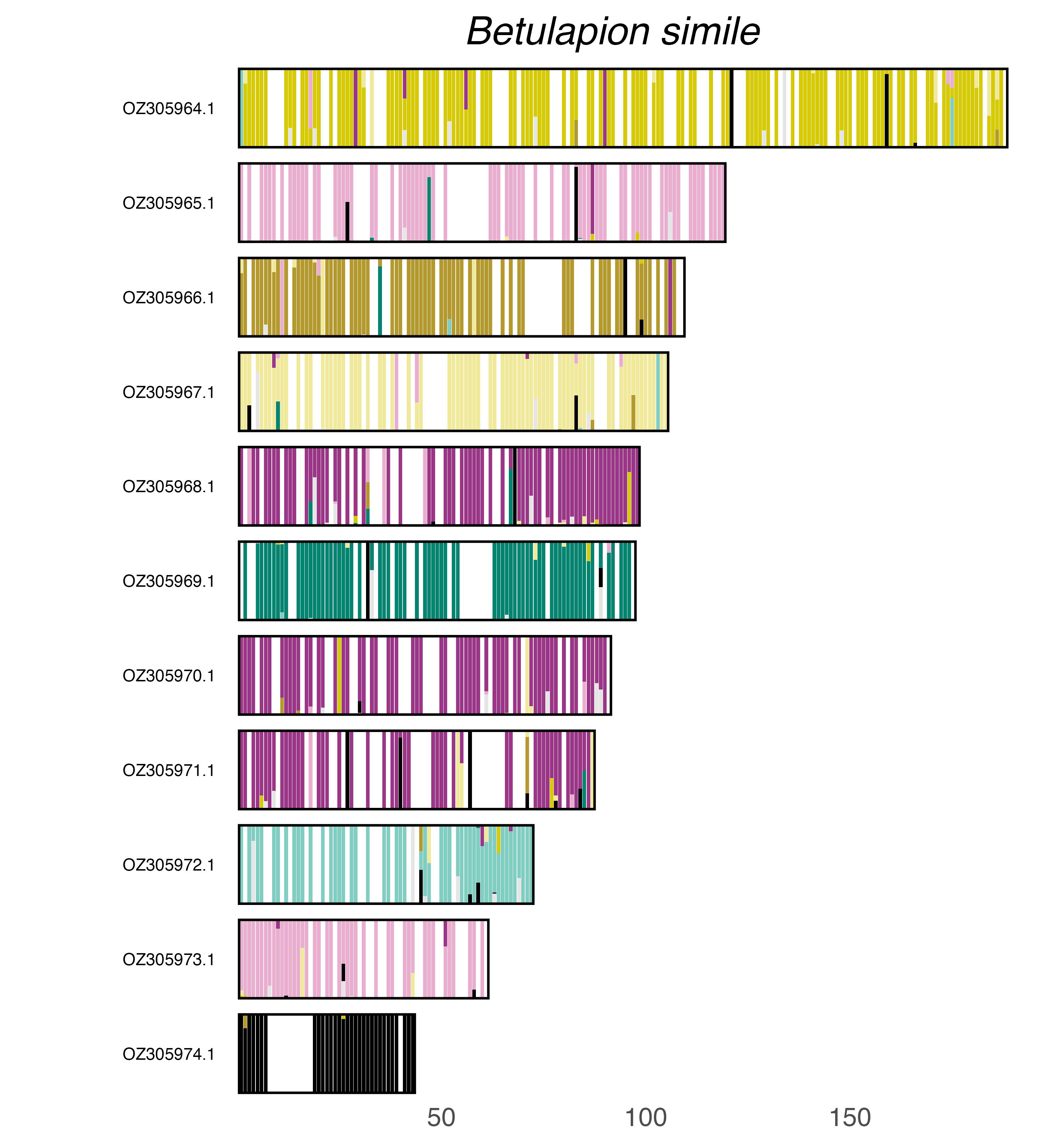

### plot.syngraph_windowbp_1Mb_rel_vertical_chrsize_Biphyllus_lunatus.png

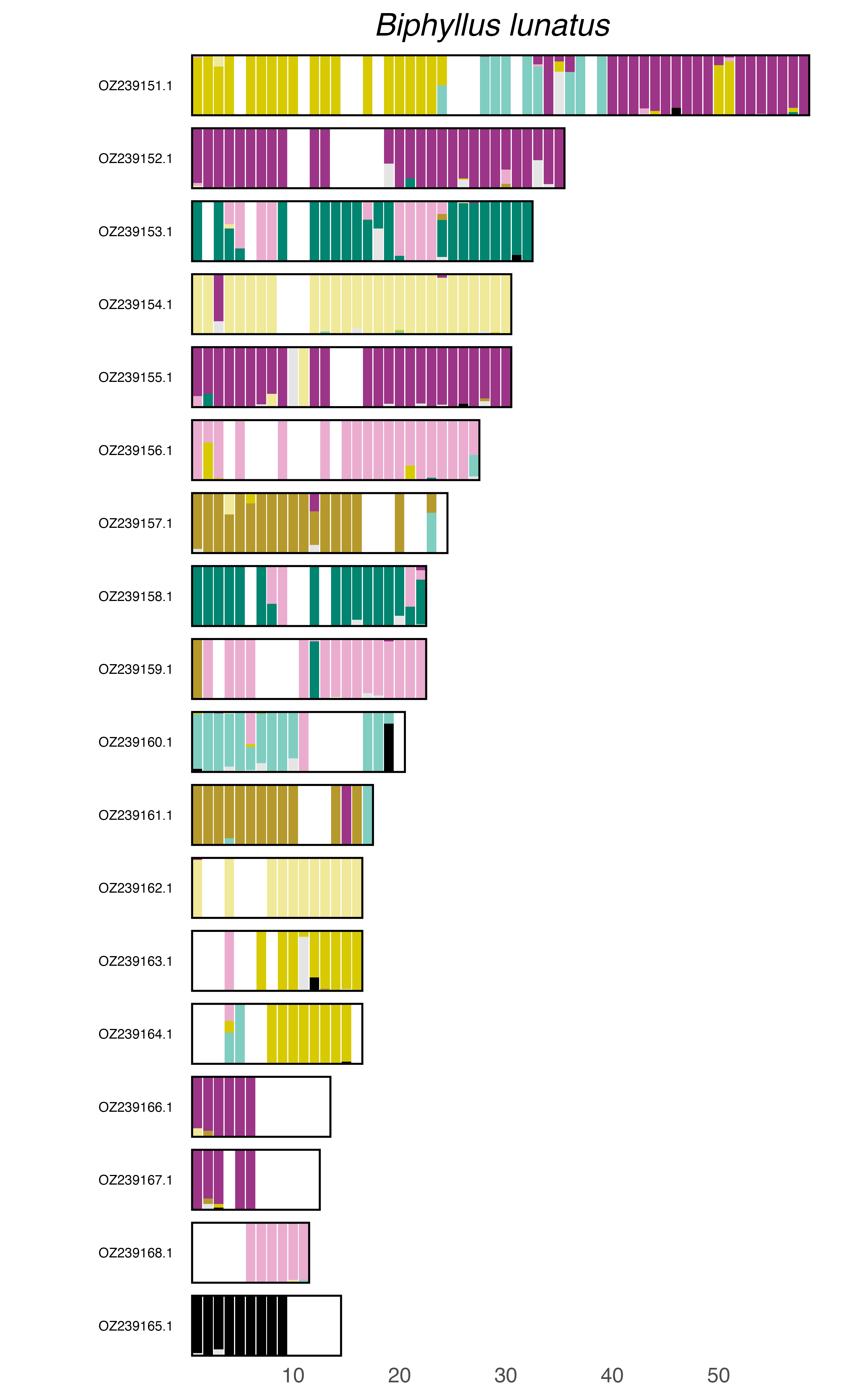

### plot.syngraph_windowbp_1Mb_rel_vertical_chrsize_Blaps_rhynchoptera.png

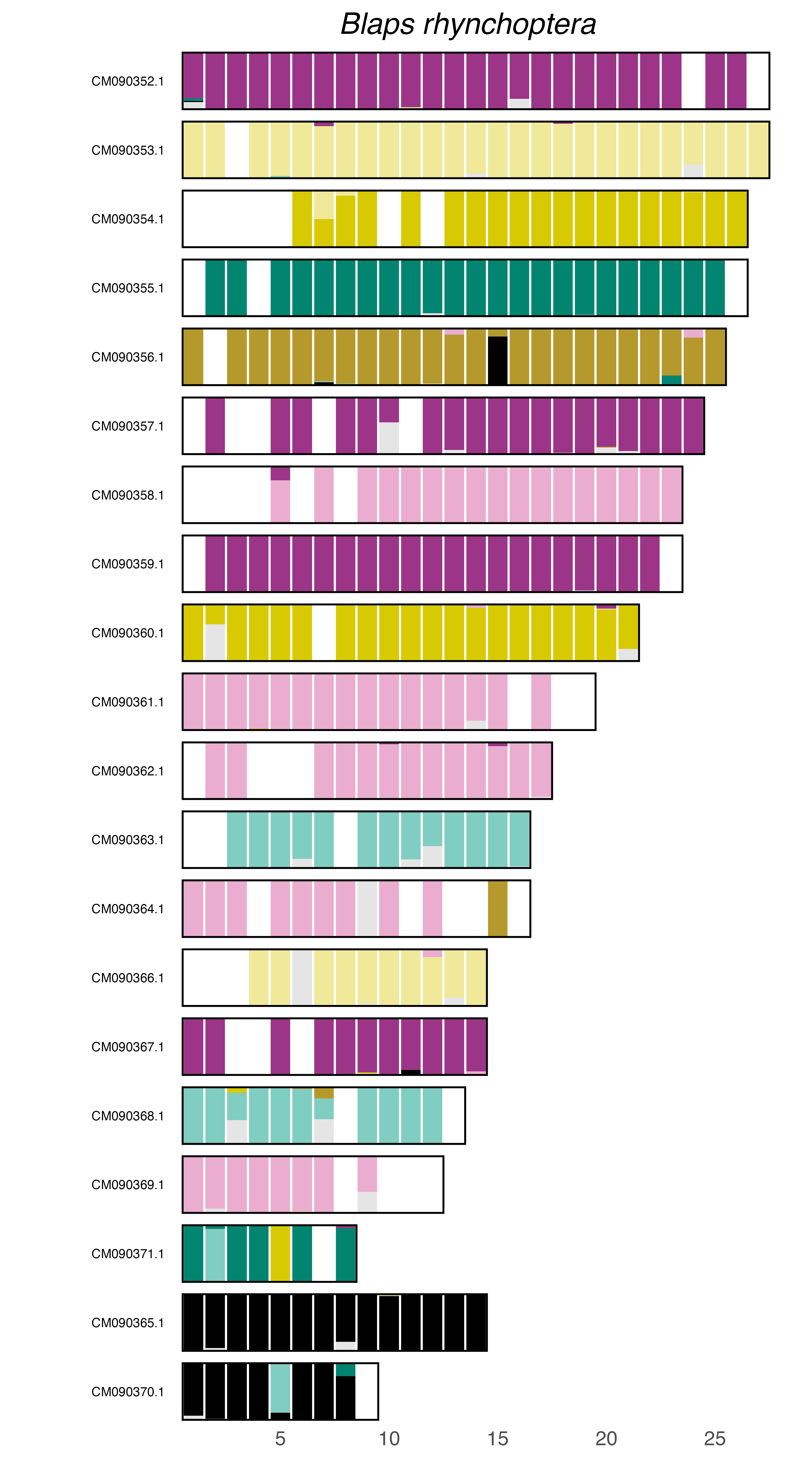

### plot.syngraph_windowbp_1Mb_rel_vertical_chrsize_Bruchidius_varius.png

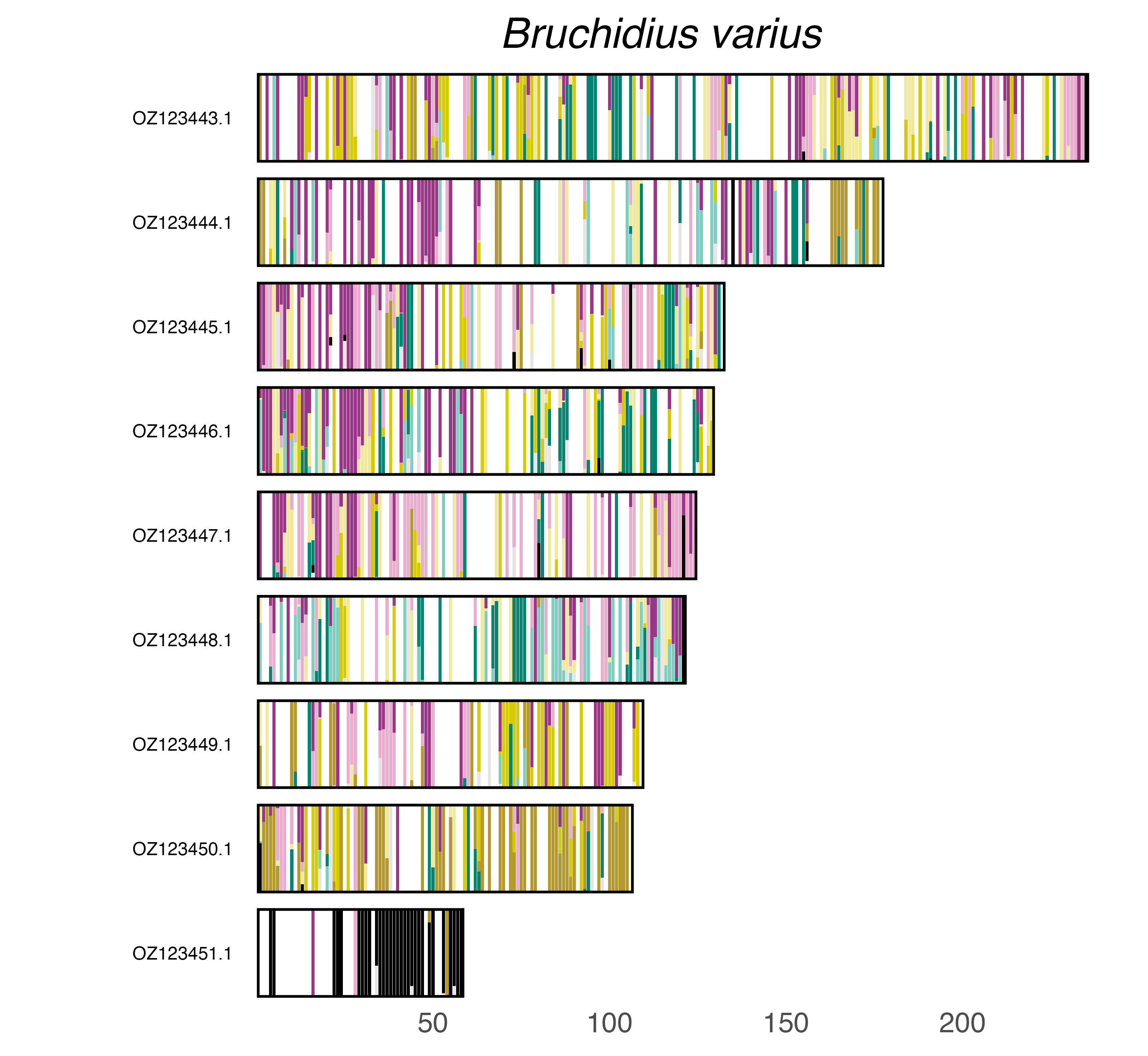

### plot.syngraph_windowbp_1Mb_rel_vertical_chrsize_Callosobruchus_maculatus.png

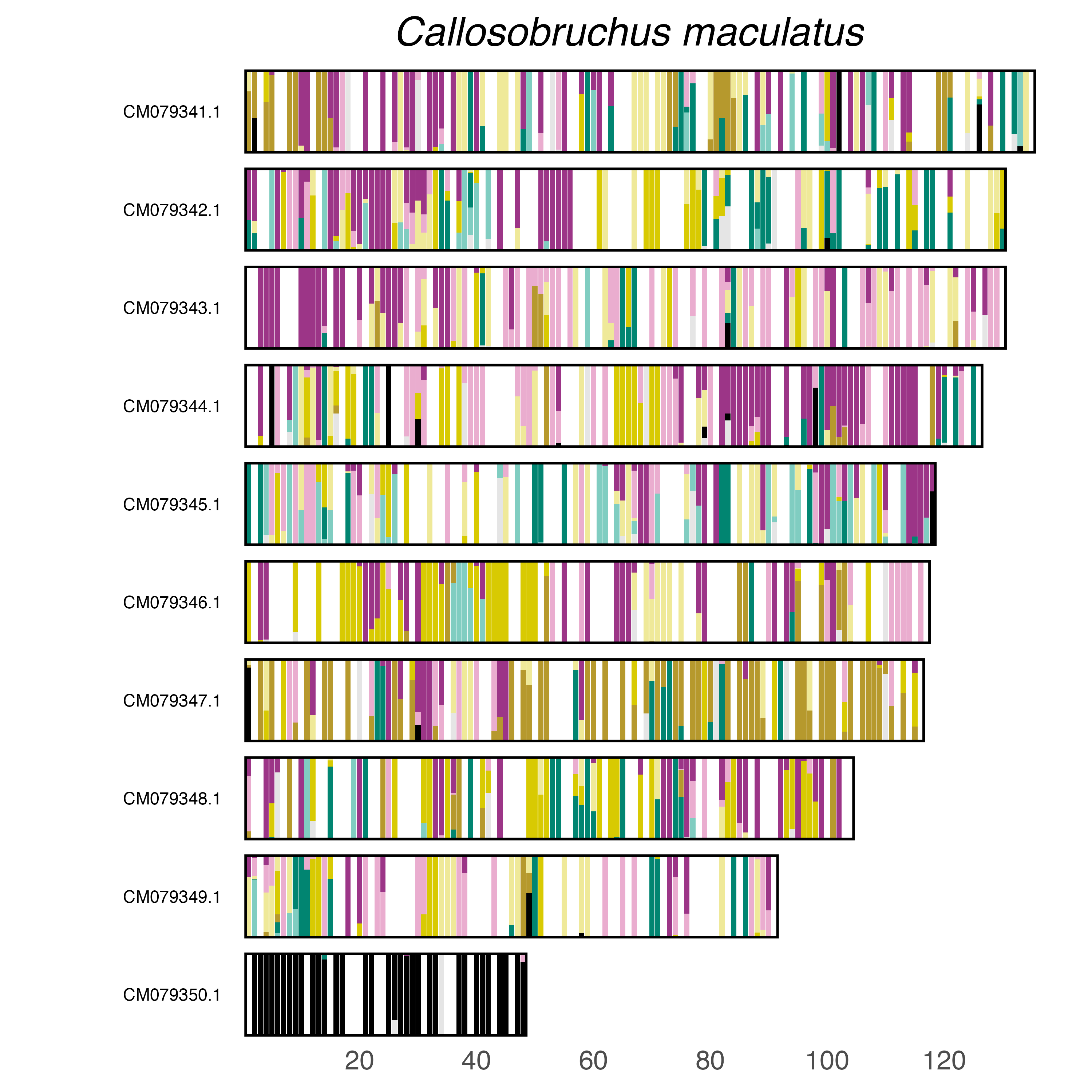

### plot.syngraph_windowbp_1Mb_rel_vertical_chrsize_Cantharis_decipiens.png

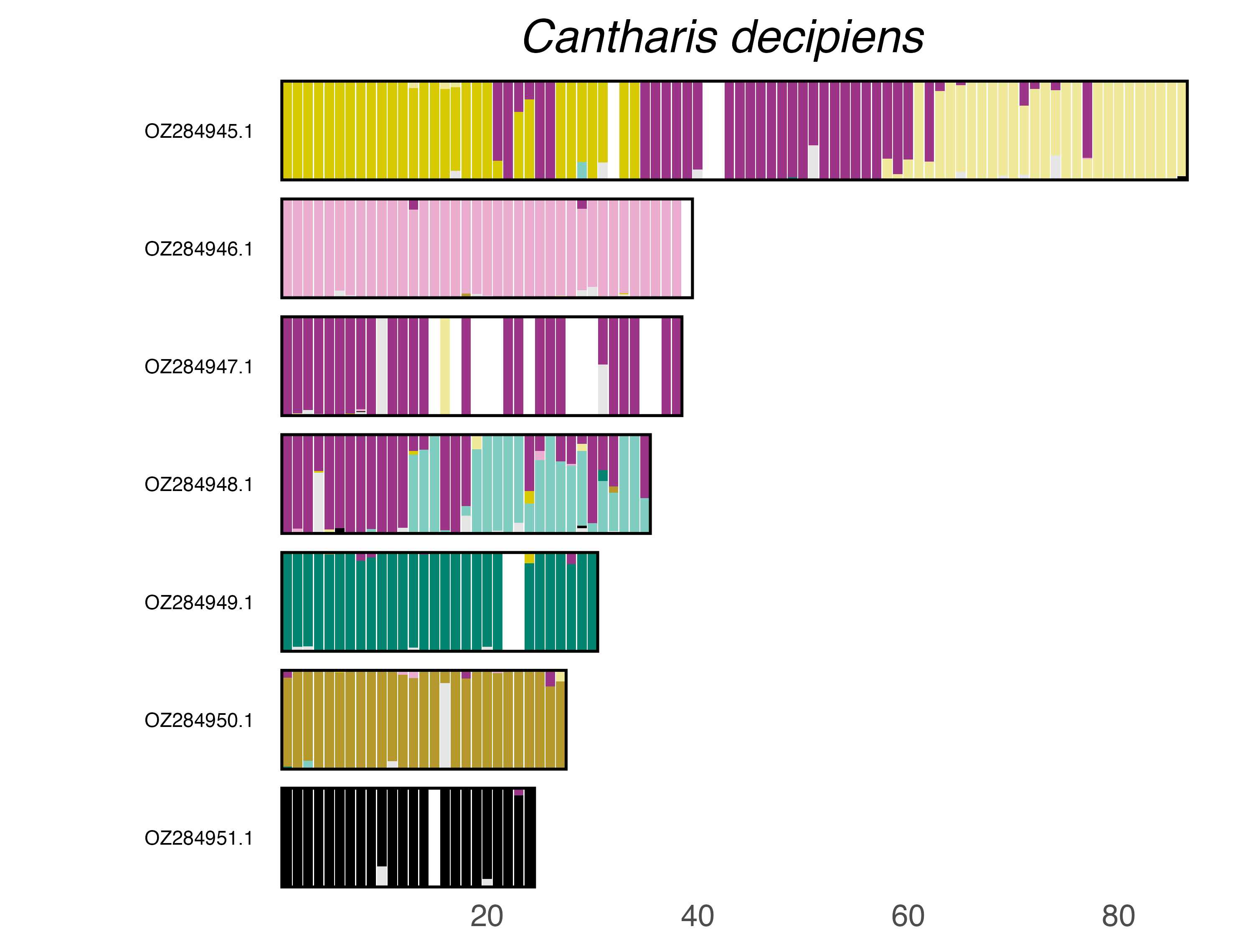

### plot.syngraph_windowbp_1Mb_rel_vertical_chrsize_Cantharis_lateralis.png

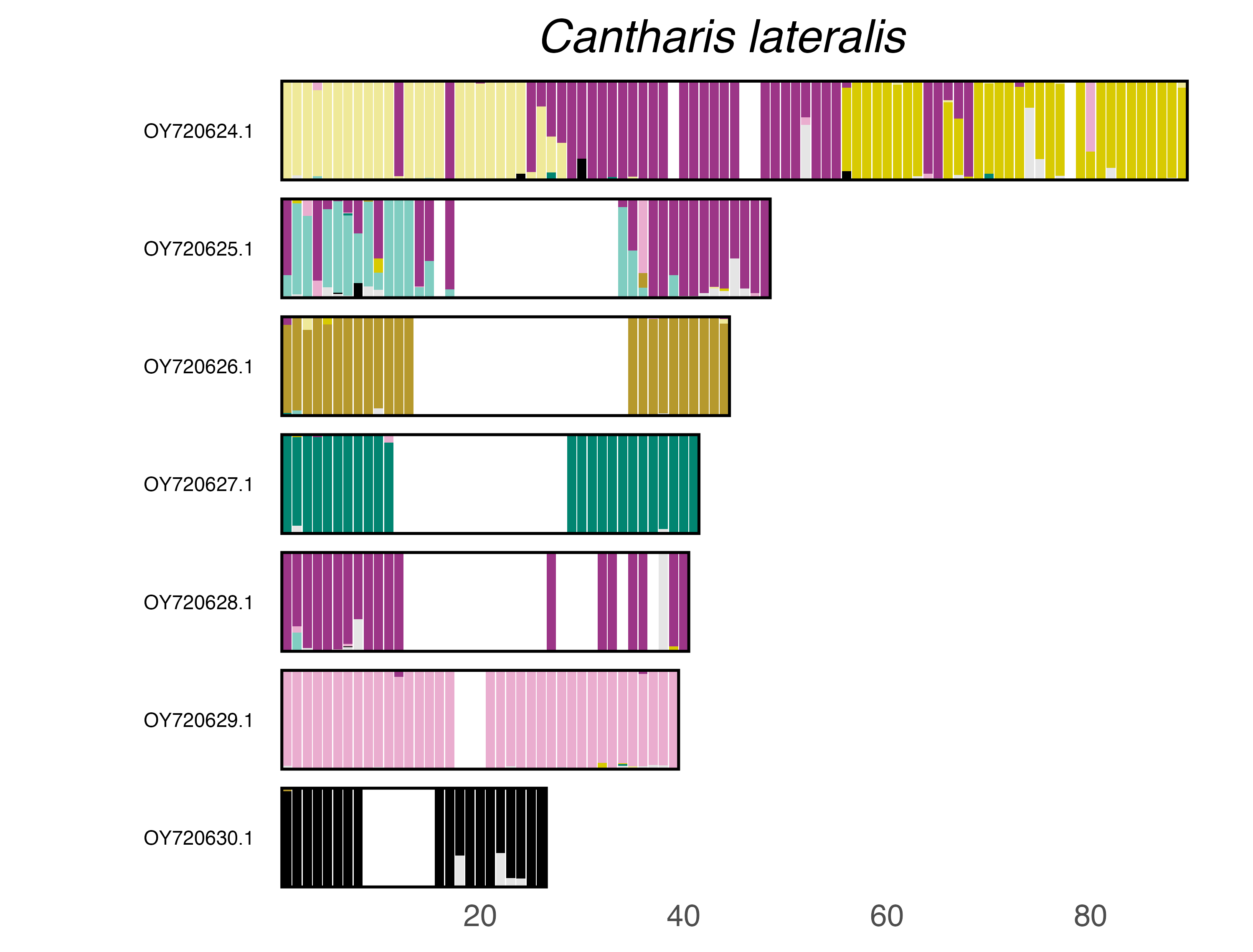

### plot.syngraph_windowbp_1Mb_rel_vertical_chrsize_Cantharis_nigra.png

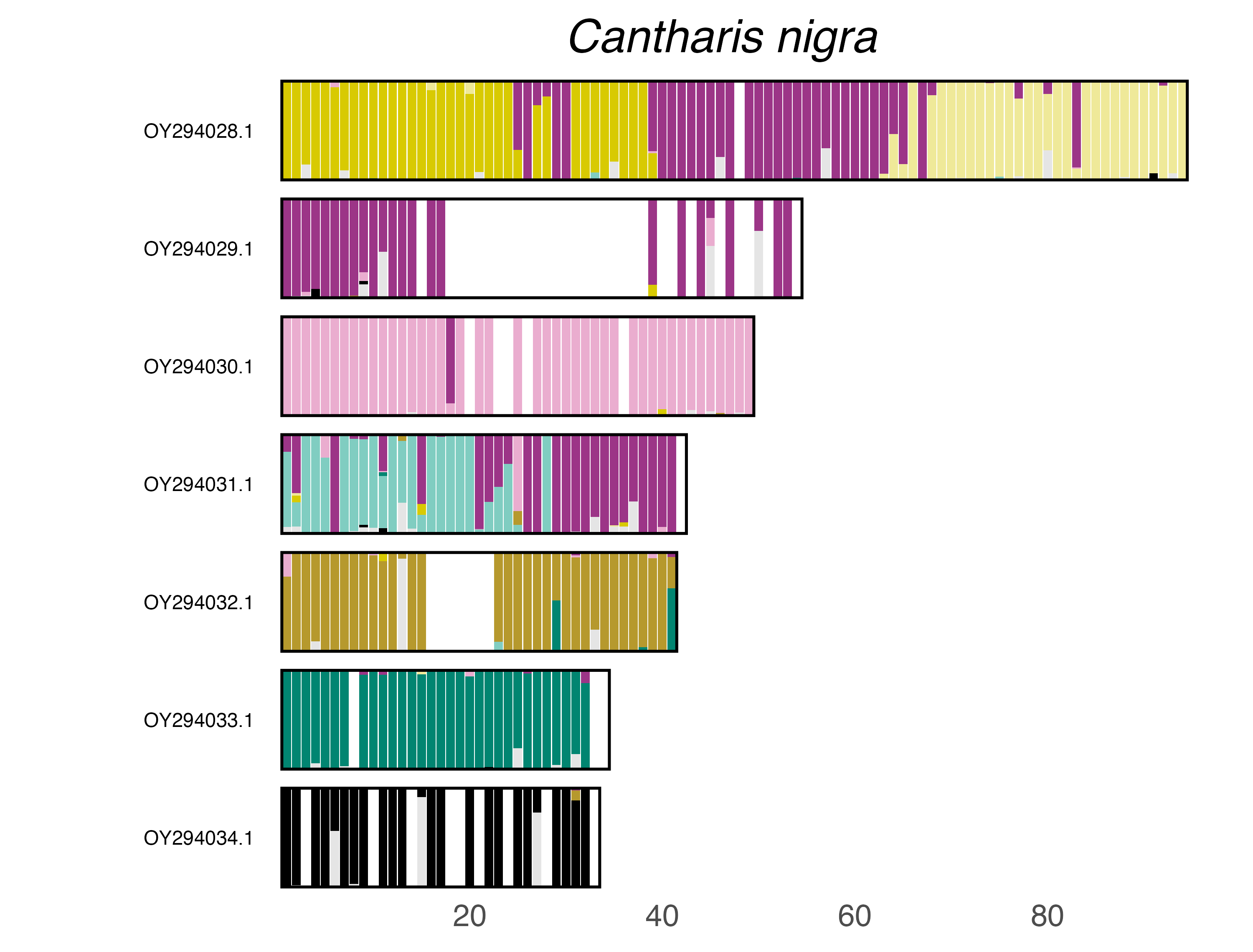

### plot.syngraph_windowbp_1Mb_rel_vertical_chrsize_Catops_nigricans.png

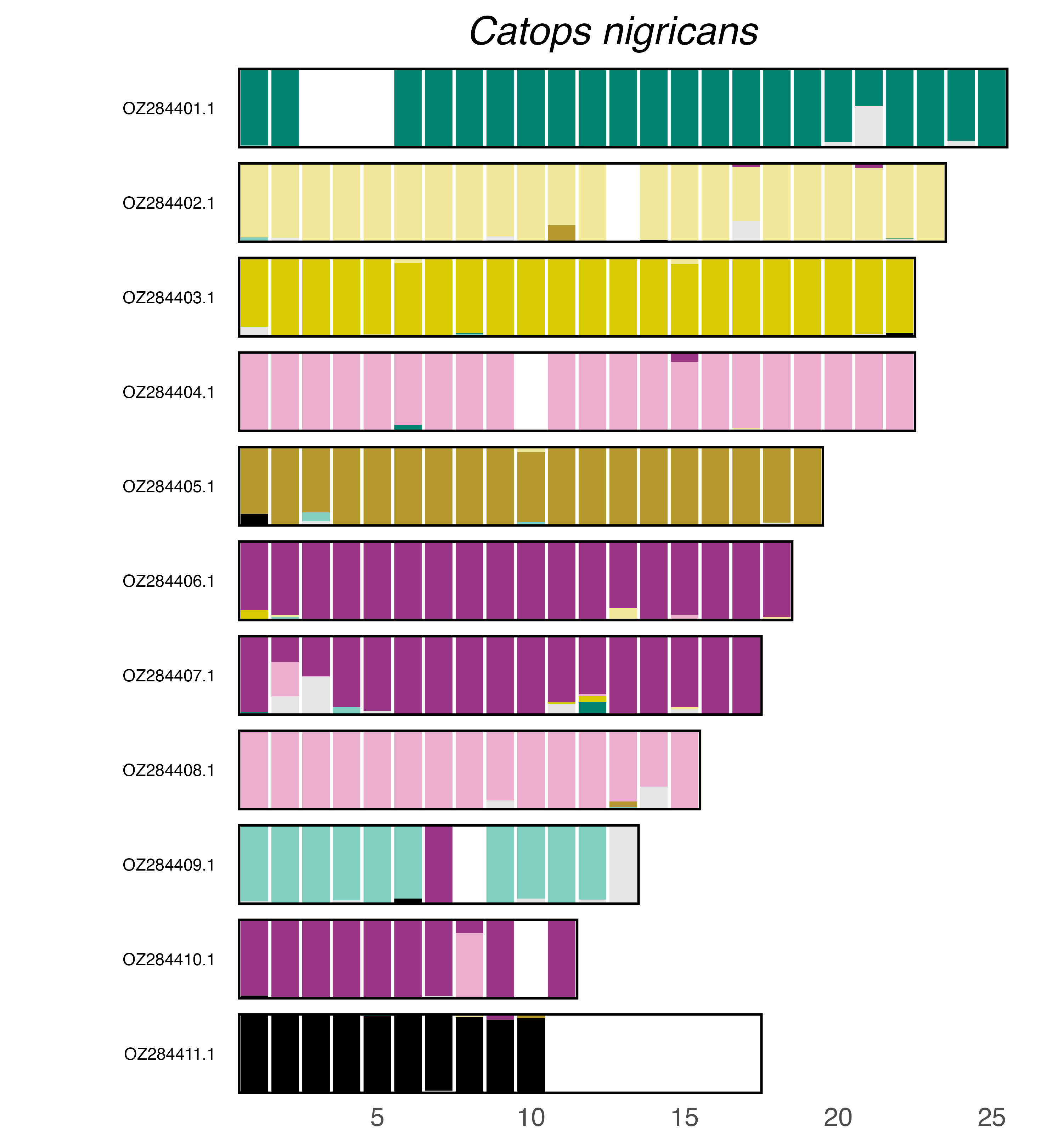

### plot.syngraph_windowbp_1Mb_rel_vertical_chrsize_Cedrorum_azoricus.png

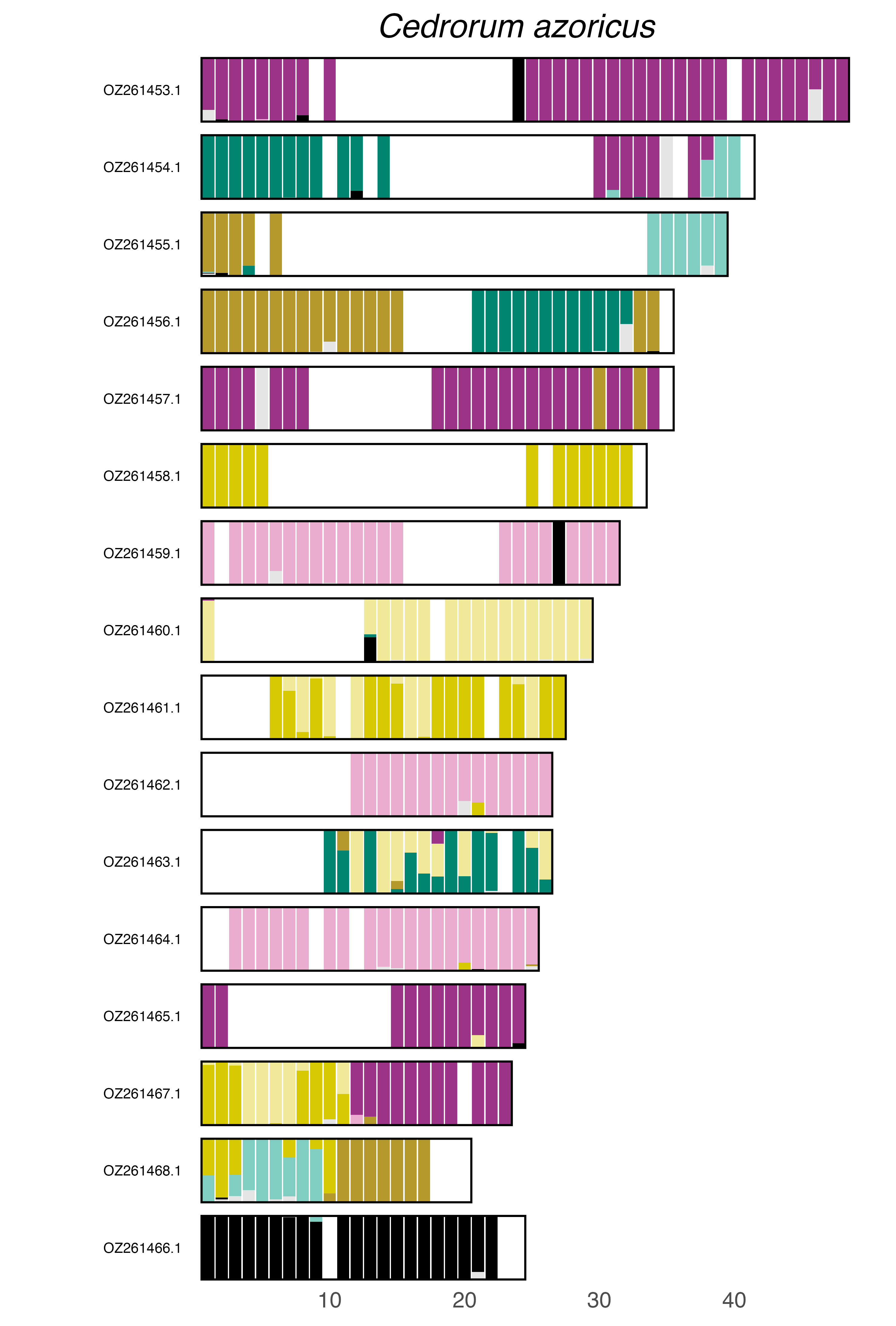

### plot.syngraph_windowbp_1Mb_rel_vertical_chrsize_Chaetocnema_hortensis.png

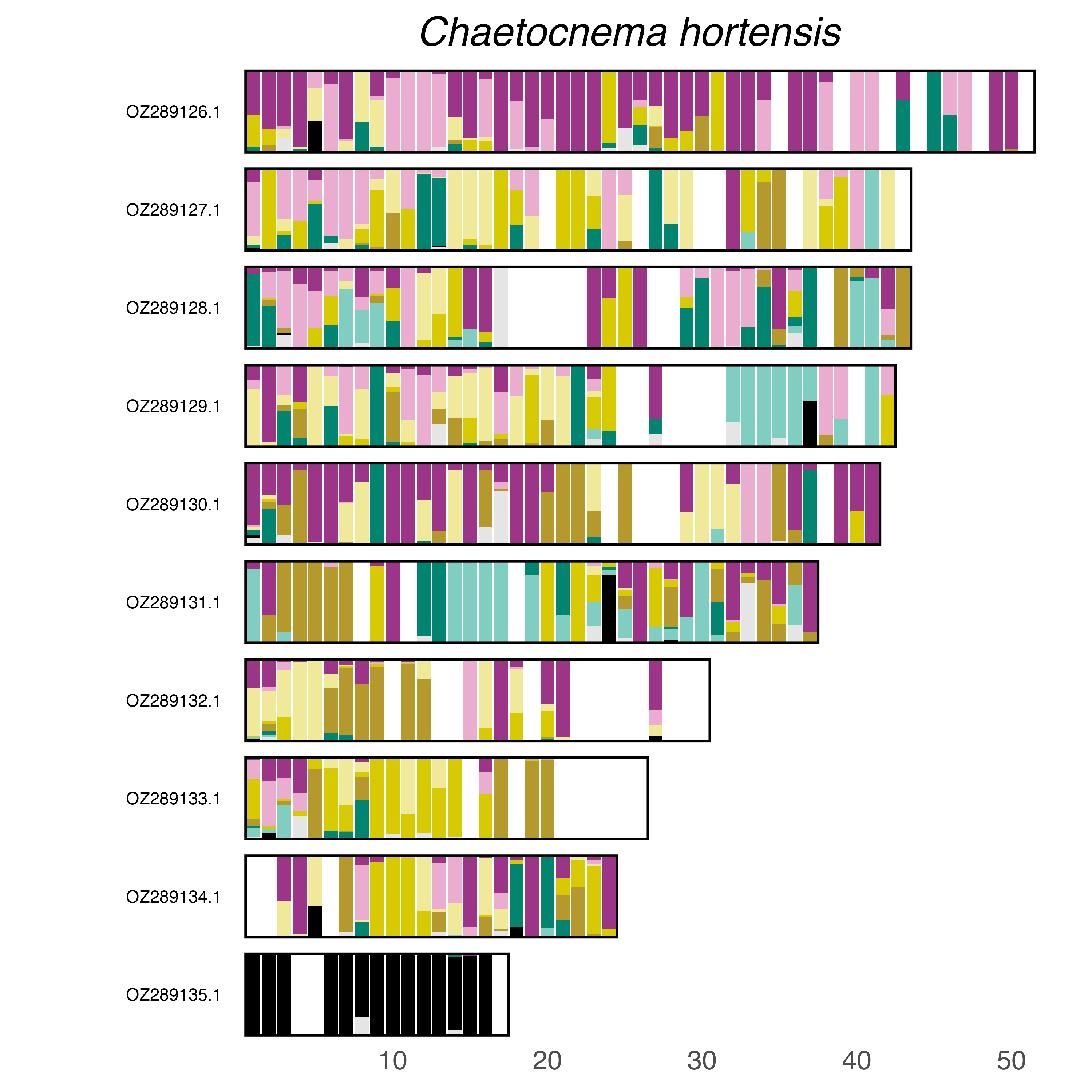

### plot.syngraph_windowbp_1Mb_rel_vertical_chrsize_Chrysomela_aeneicollis.png

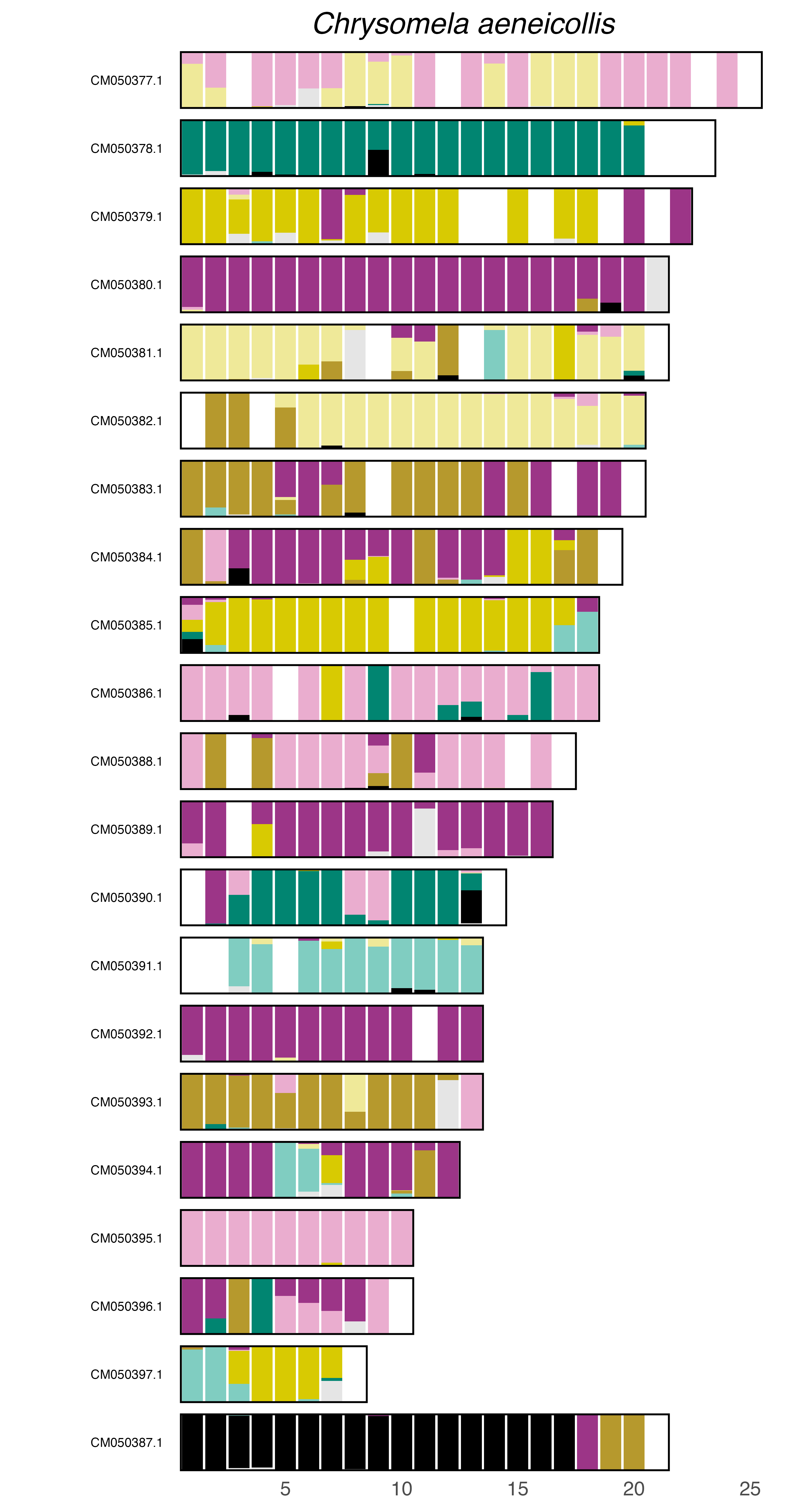

### plot.syngraph_windowbp_1Mb_rel_vertical_chrsize_Coccinella_septempunctata.png

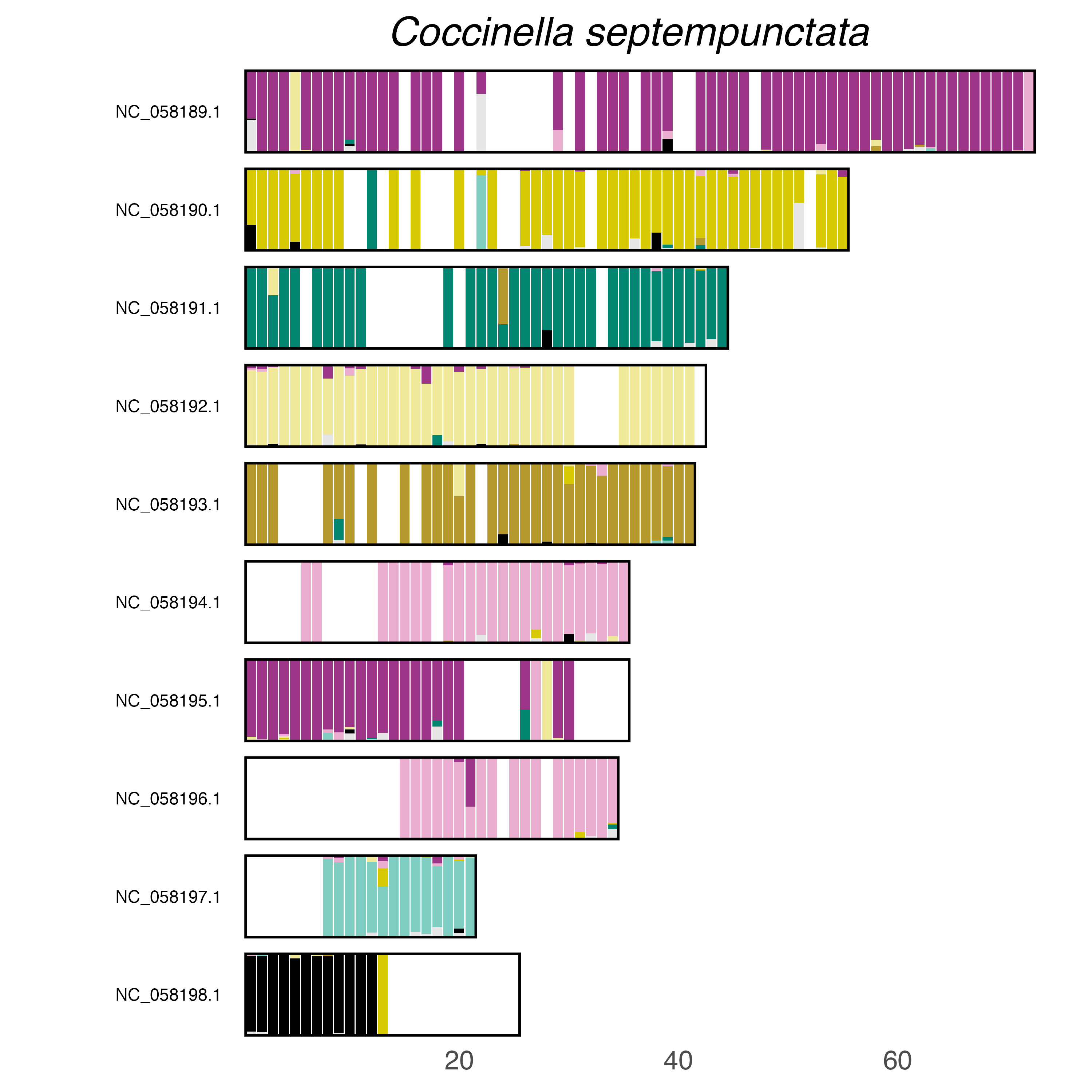

### plot.syngraph_windowbp_1Mb_rel_vertical_chrsize_Conwentzia_psociformis.png

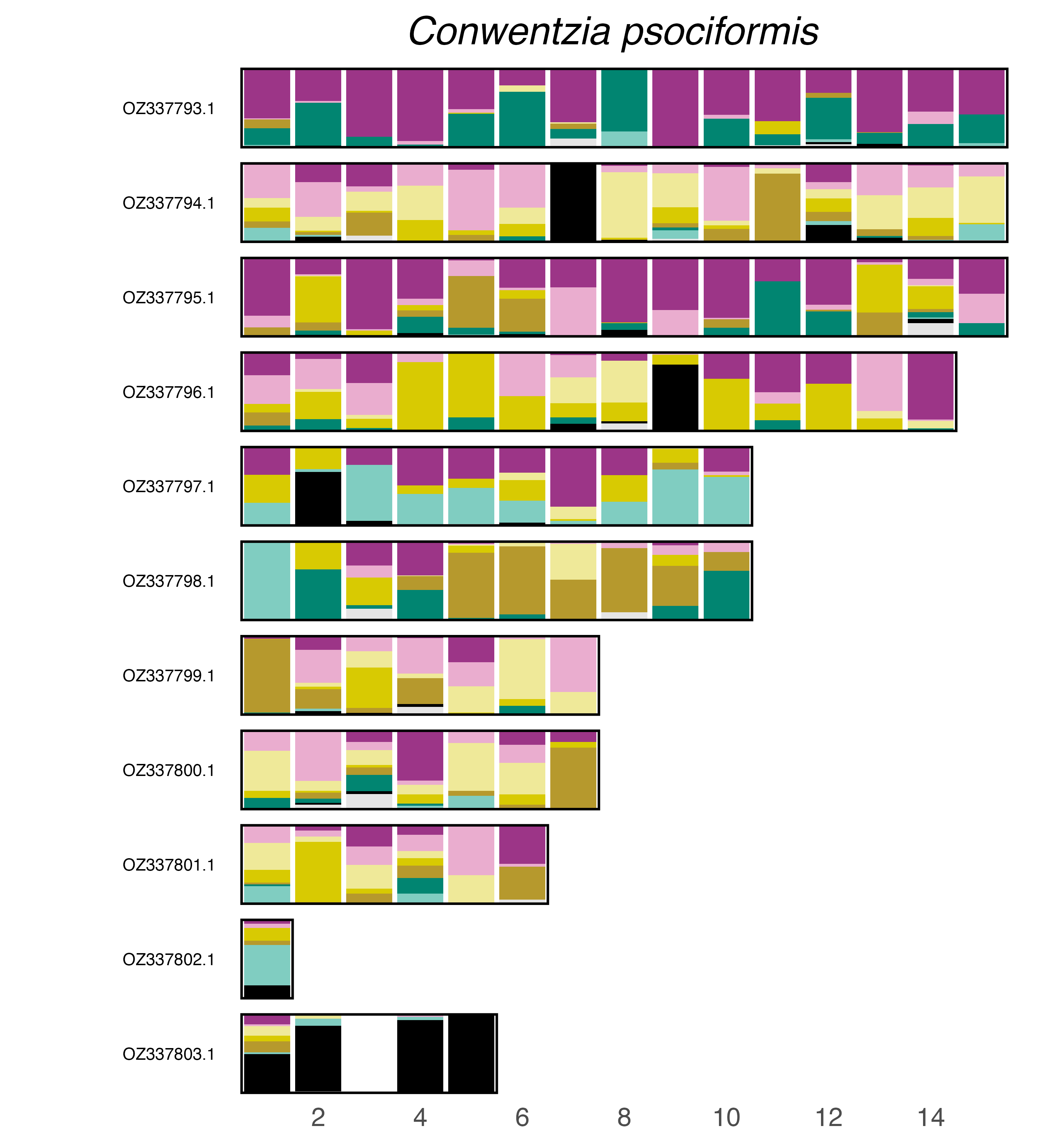

### plot.syngraph_windowbp_1Mb_rel_vertical_chrsize_Crepidodera_aurata.png

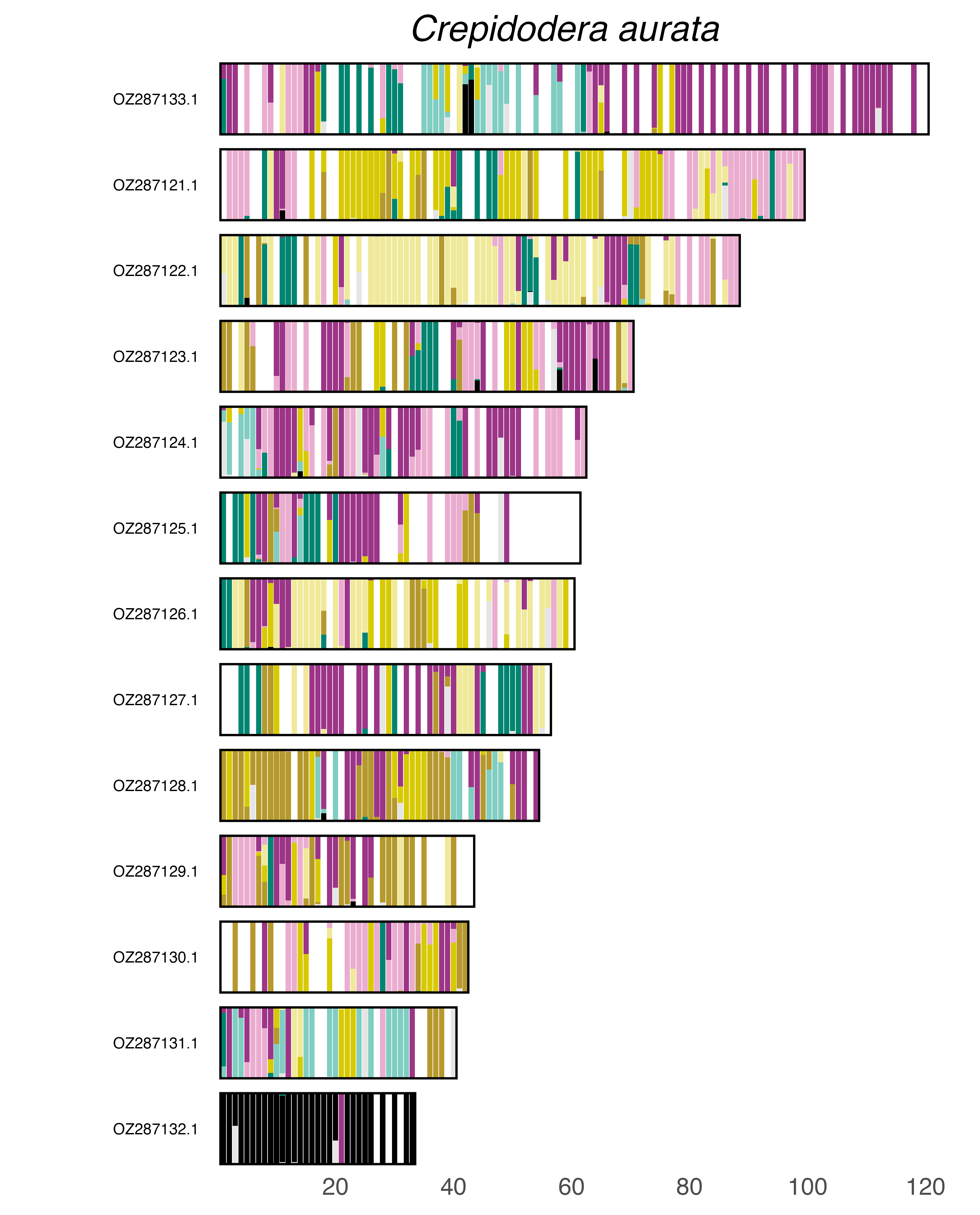

### plot.syngraph_windowbp_1Mb_rel_vertical_chrsize_Crepidodera_aurea.png

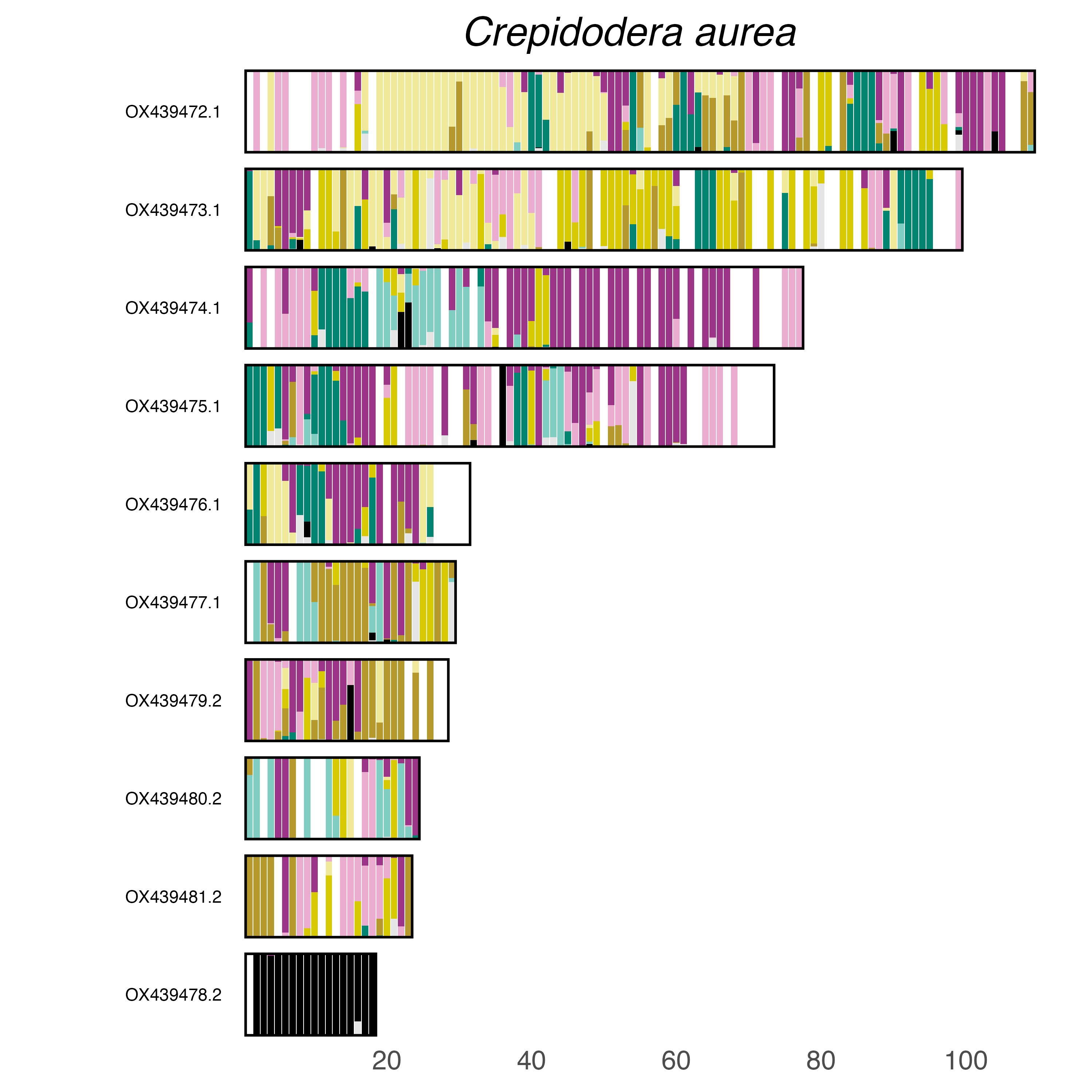

### plot.syngraph_windowbp_1Mb_rel_vertical_chrsize_Crioceris_asparagi.png

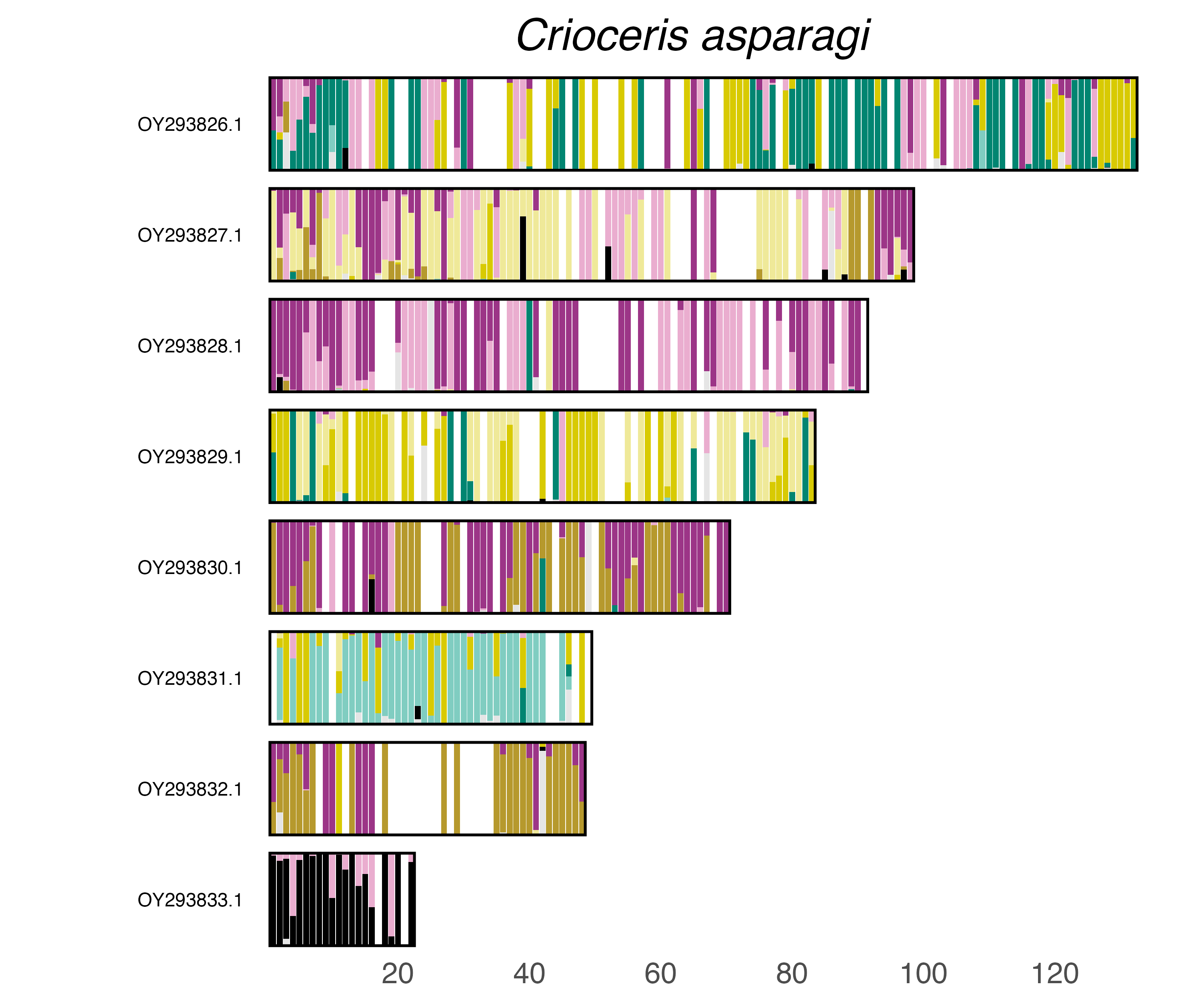

### plot.syngraph_windowbp_1Mb_rel_vertical_chrsize_Crudosilis_ruficollis.png

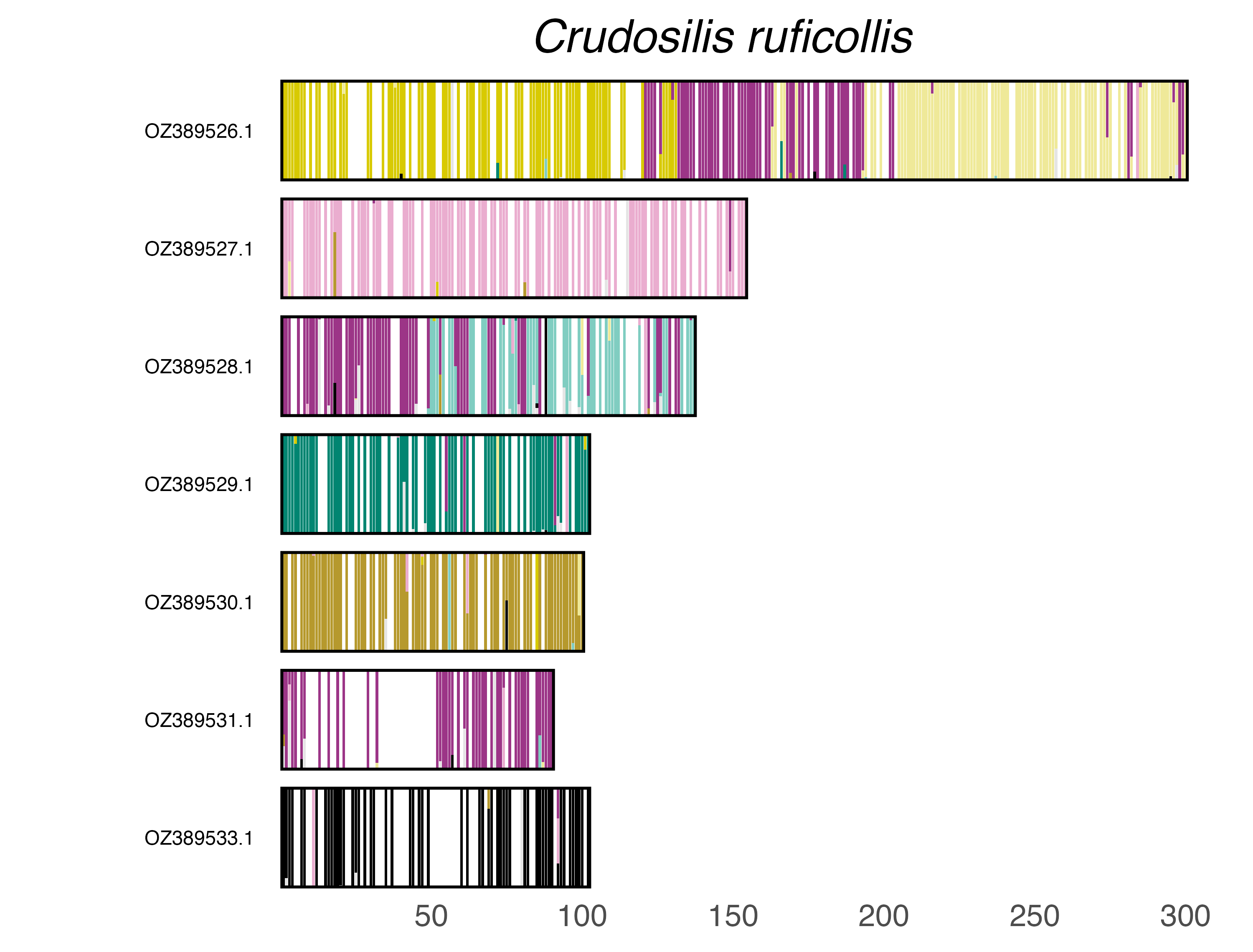

### plot.syngraph_windowbp_1Mb_rel_vertical_chrsize_Cryptocephalus_primarius.png

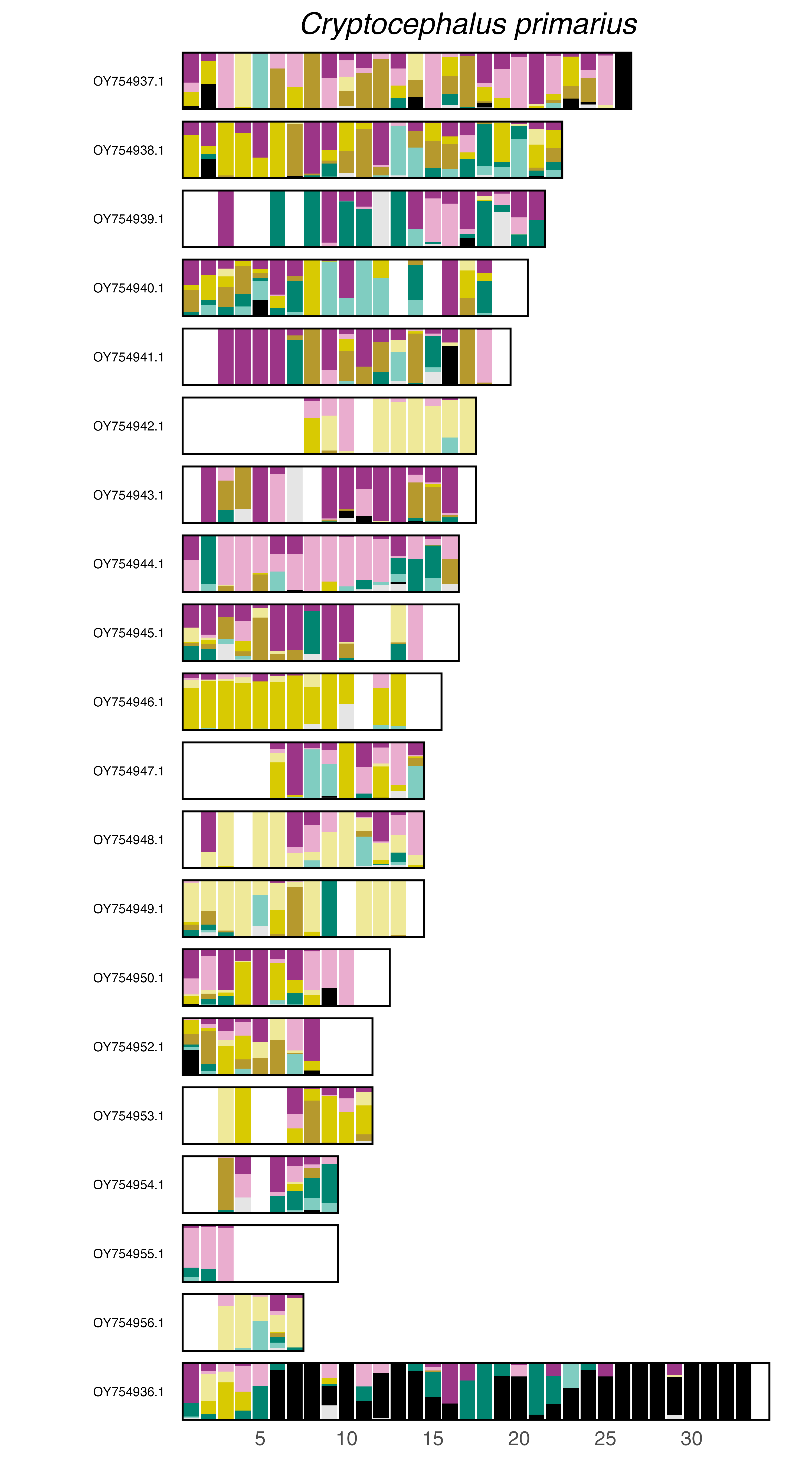

### plot.syngraph_windowbp_1Mb_rel_vertical_chrsize_Cryptophagus_acutangulus.png

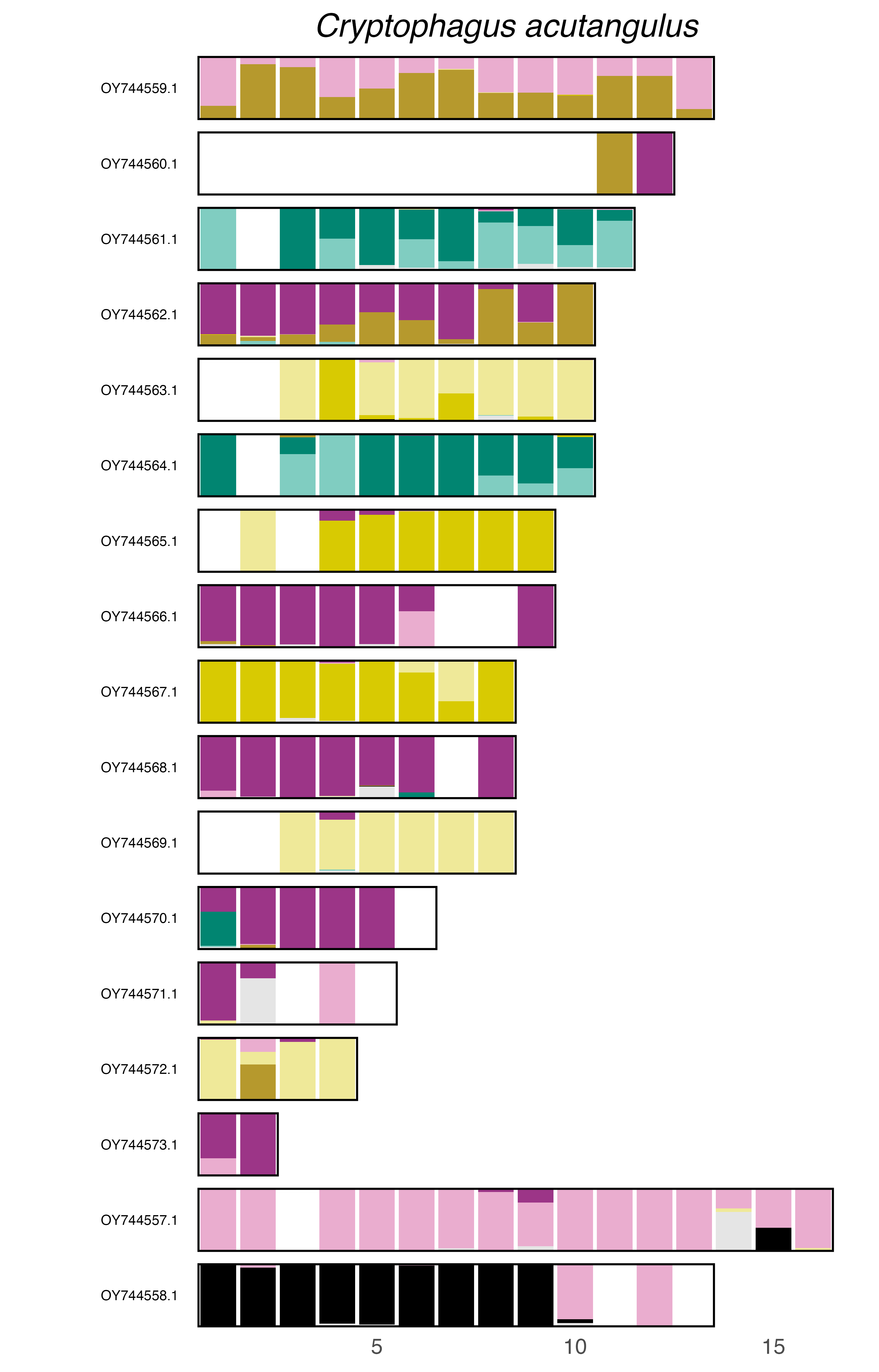

### plot.syngraph_windowbp_1Mb_rel_vertical_chrsize_Cryptorhynchus_lapathi.png

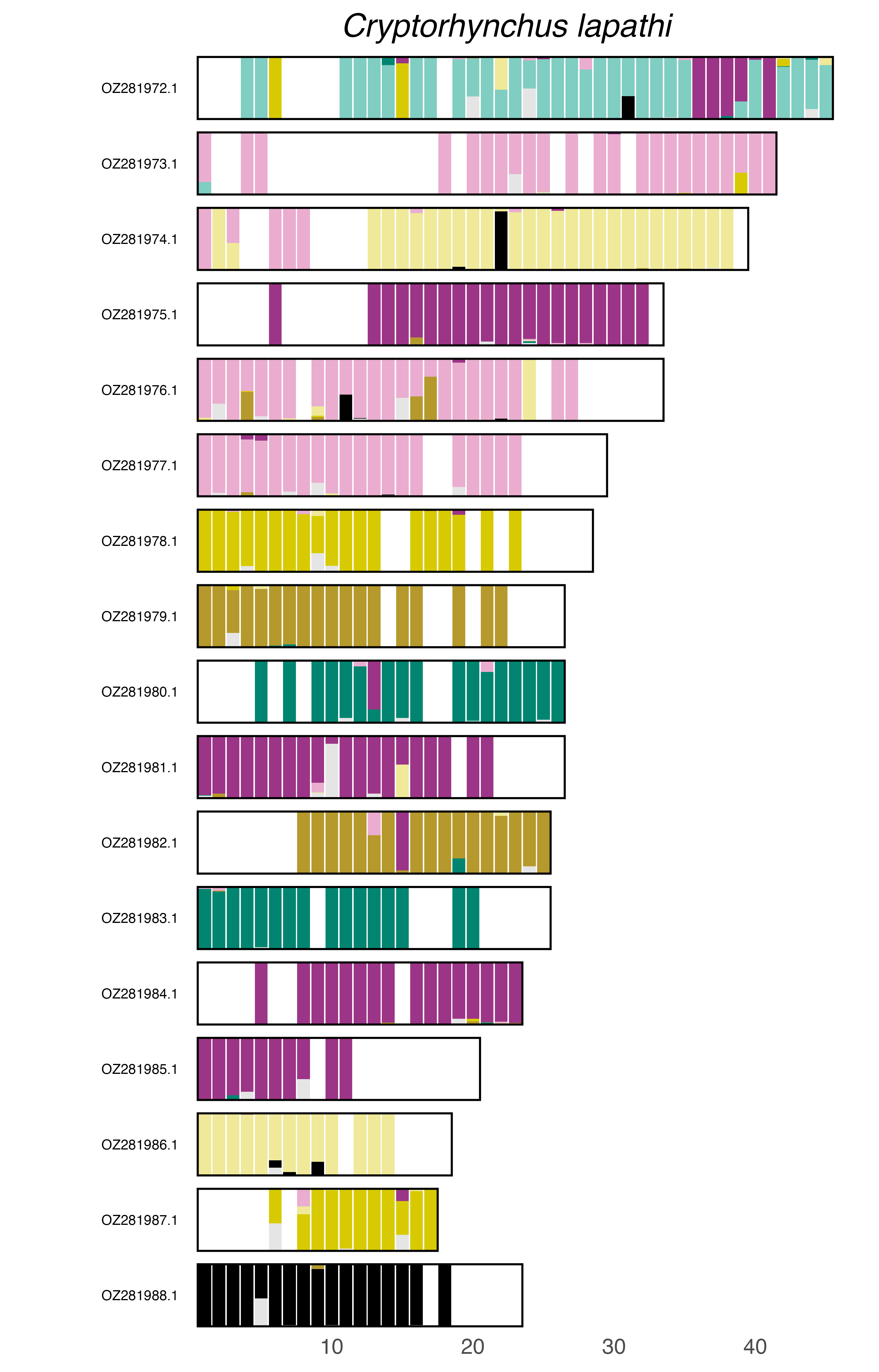

### plot.syngraph_windowbp_1Mb_rel_vertical_chrsize_Curculio_caryae.png

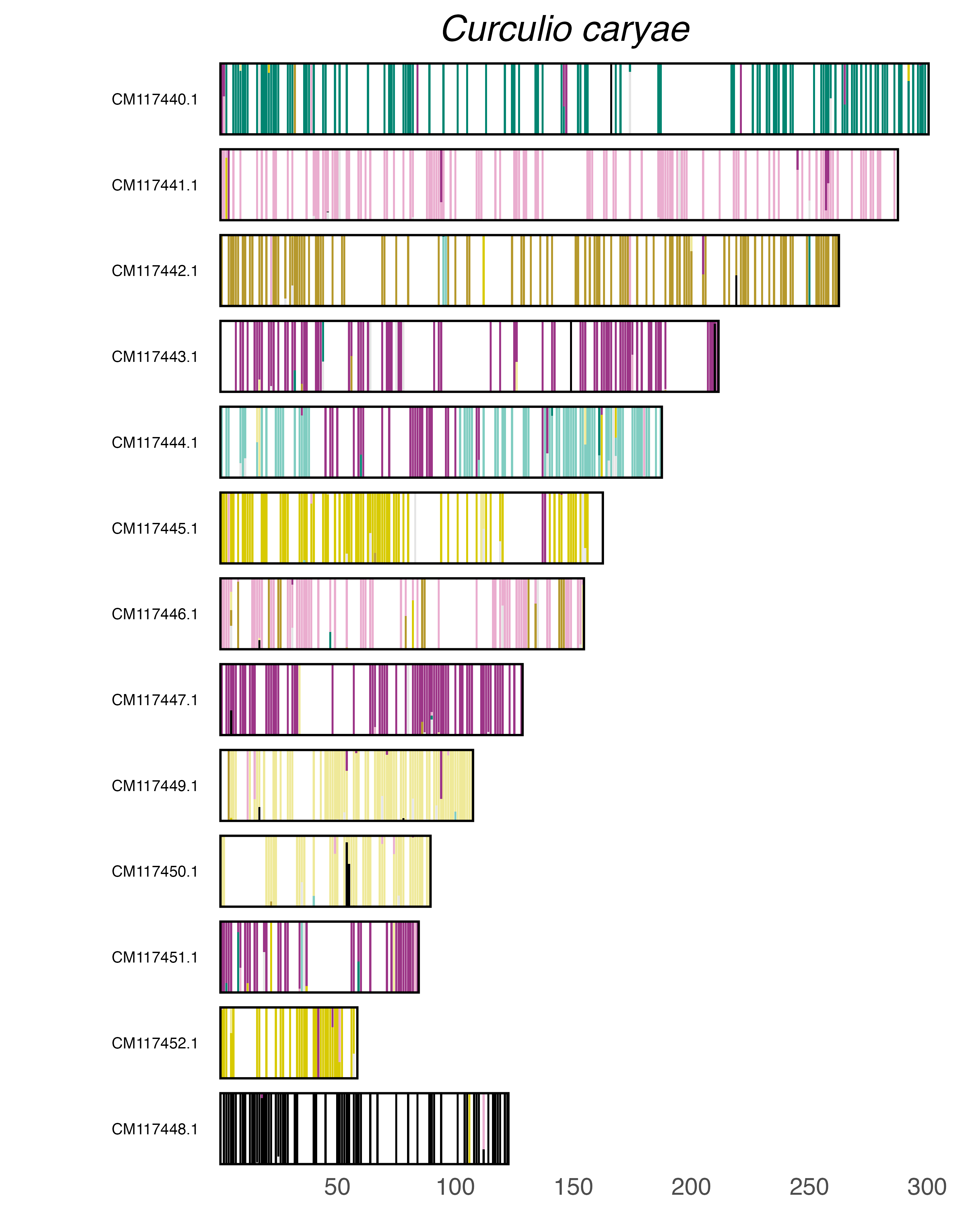

### plot.syngraph_windowbp_1Mb_rel_vertical_chrsize_Dailognatha_quadricollis.png

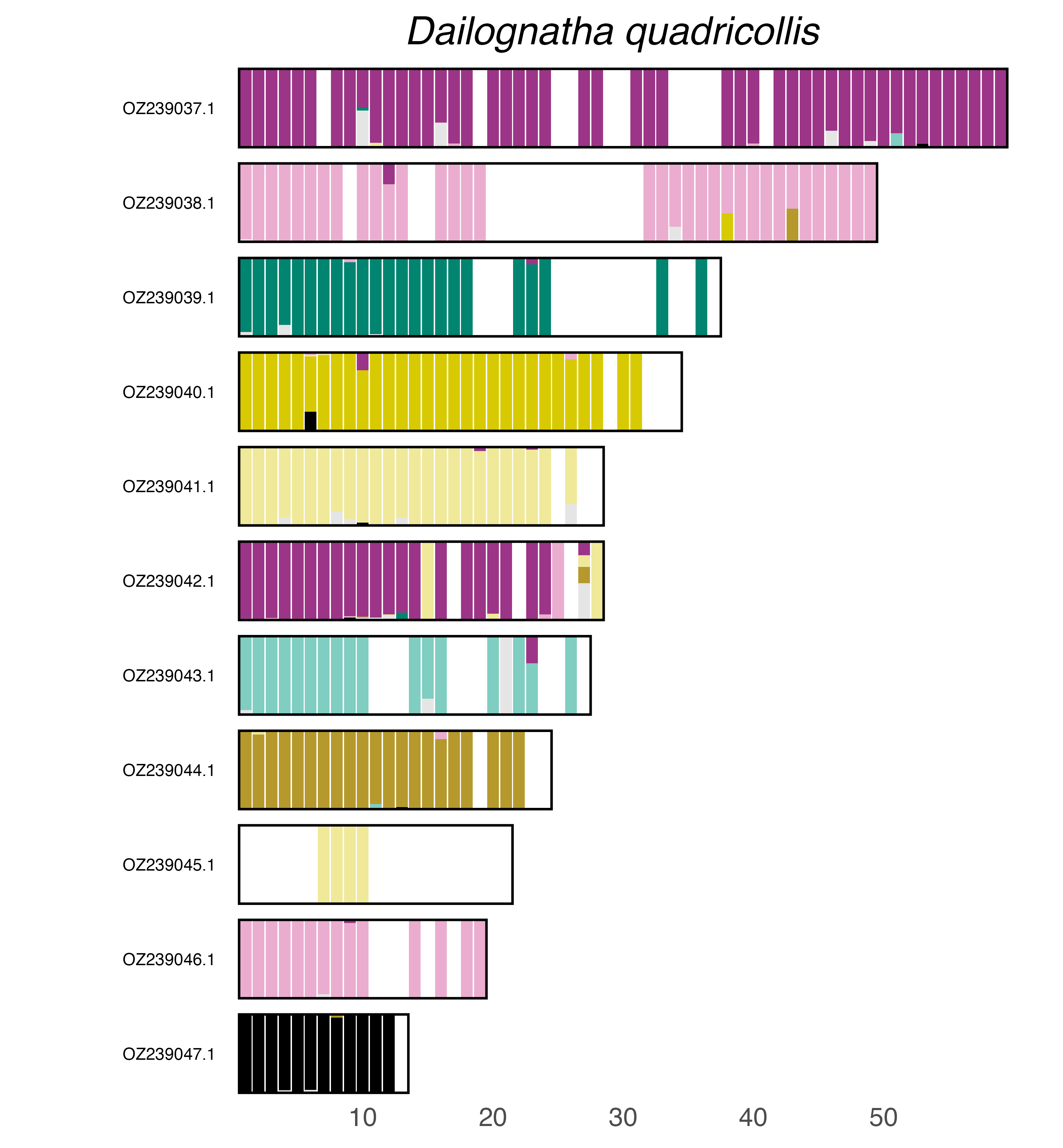

### plot.syngraph_windowbp_1Mb_rel_vertical_chrsize_Dalopius_marginatus.png

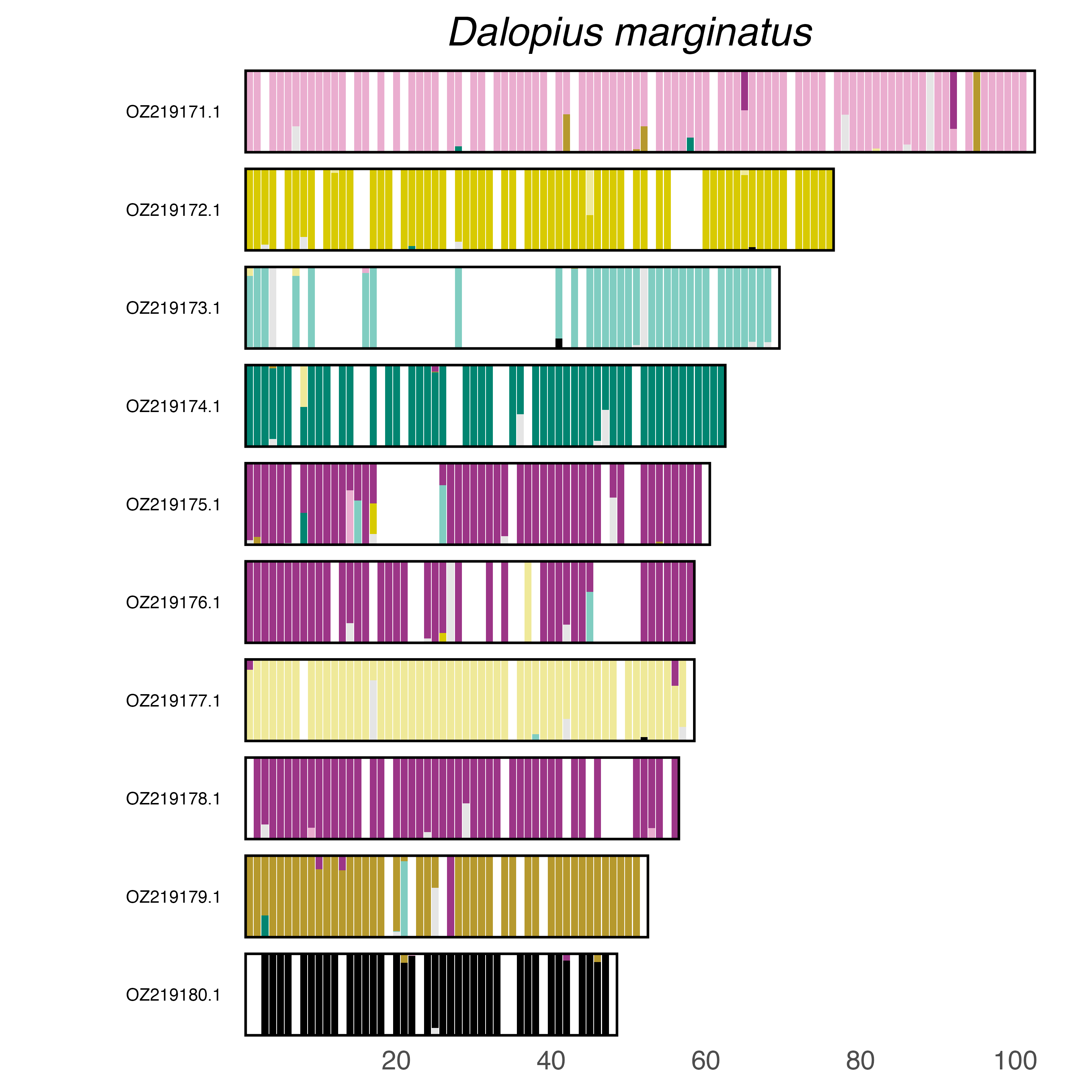

### plot.syngraph_windowbp_1Mb_rel_vertical_chrsize_Dastarcus_helophoroides.png

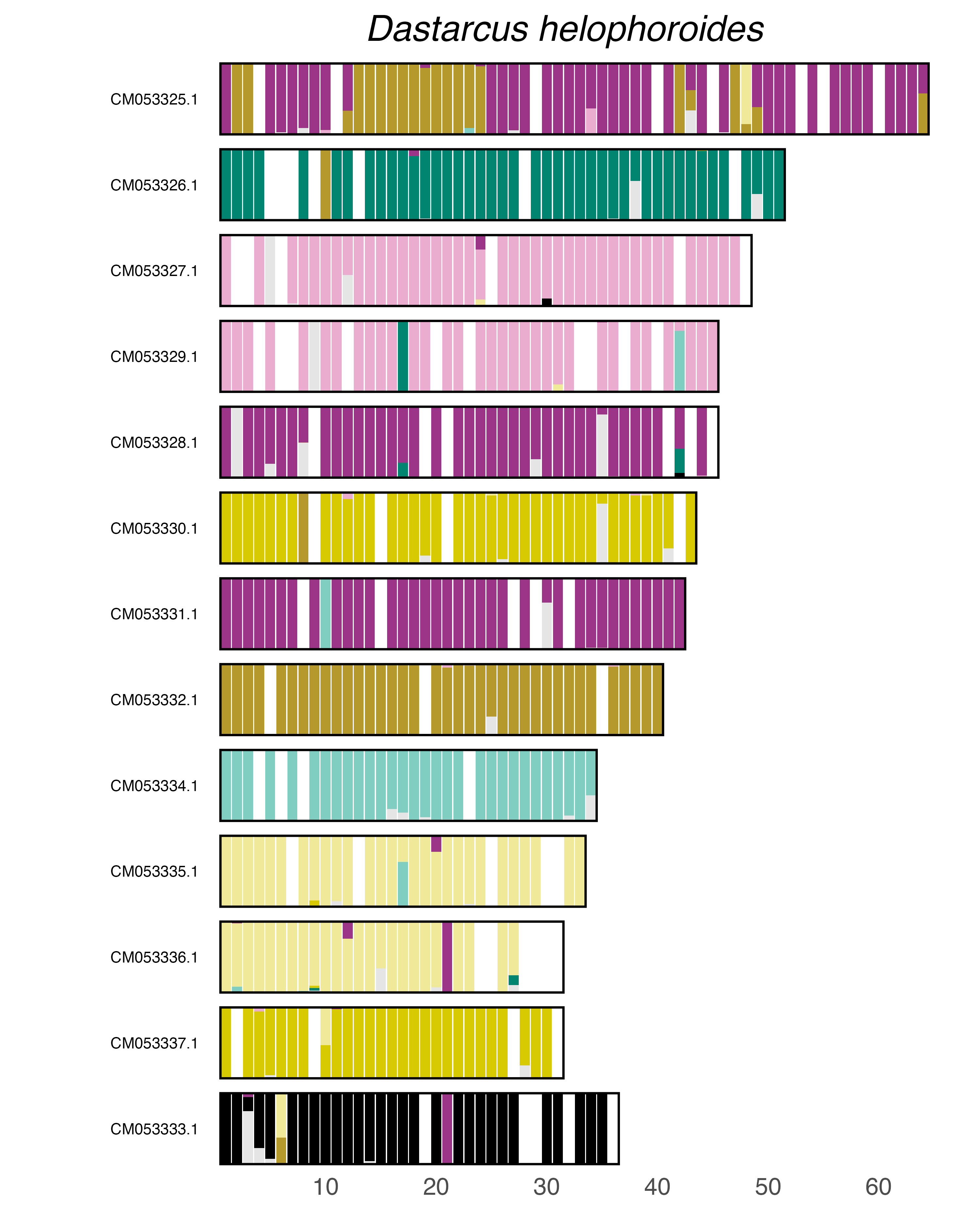

### plot.syngraph_windowbp_1Mb_rel_vertical_chrsize_Denticollis_linearis.png

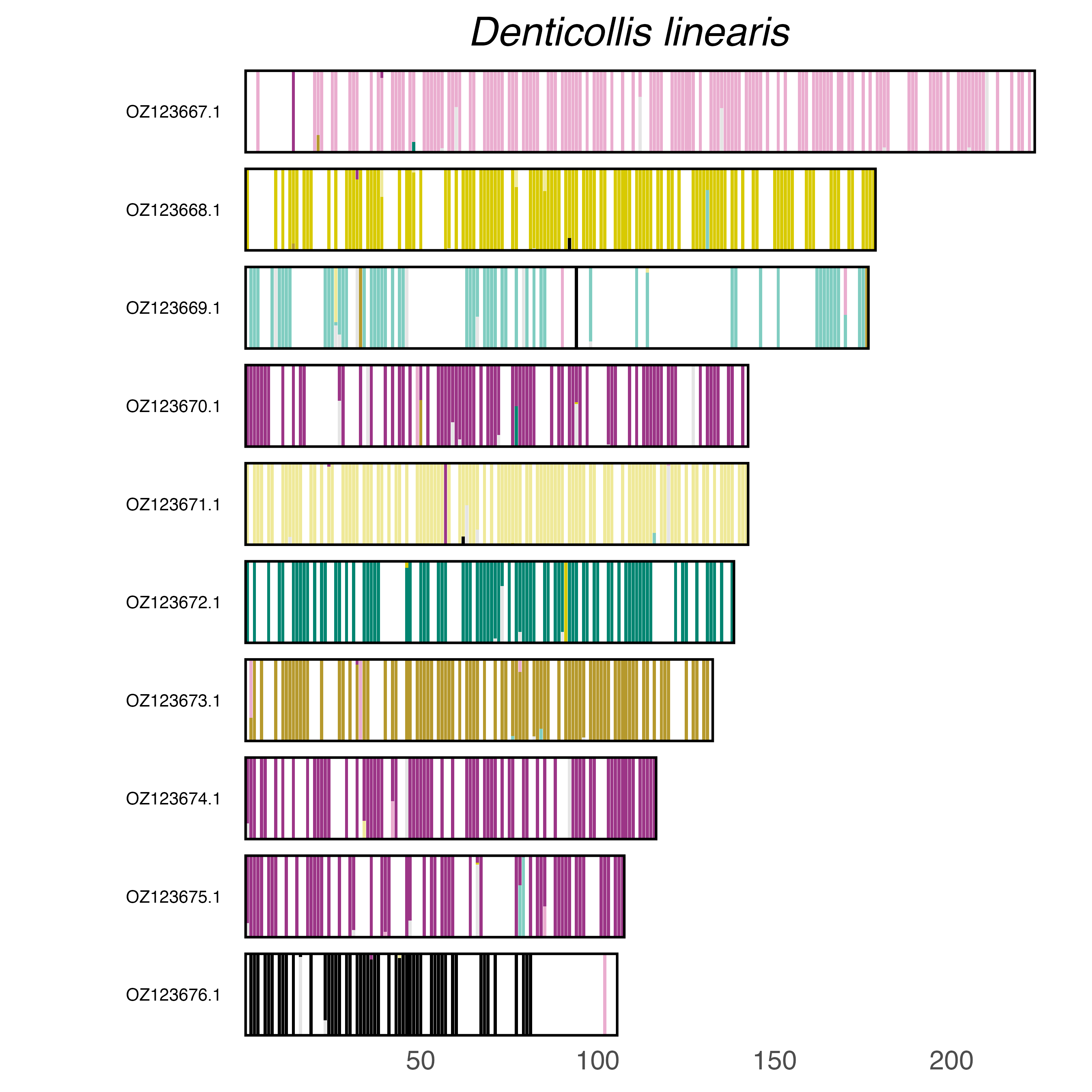

### plot.syngraph_windowbp_1Mb_rel_vertical_chrsize_Diabrotica_virgifera.png

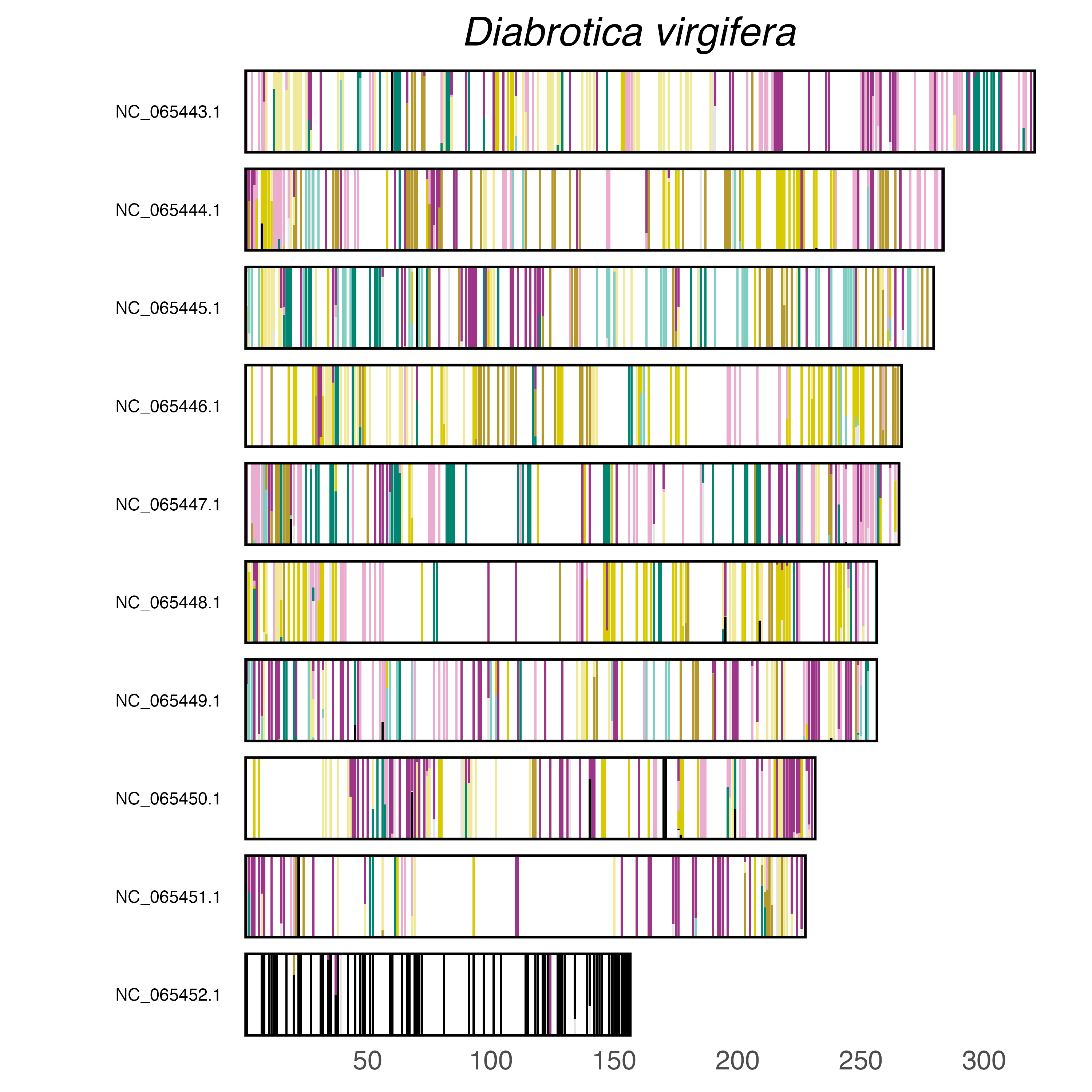

### plot.syngraph_windowbp_1Mb_rel_vertical_chrsize_Diorhabda_carinulata.png

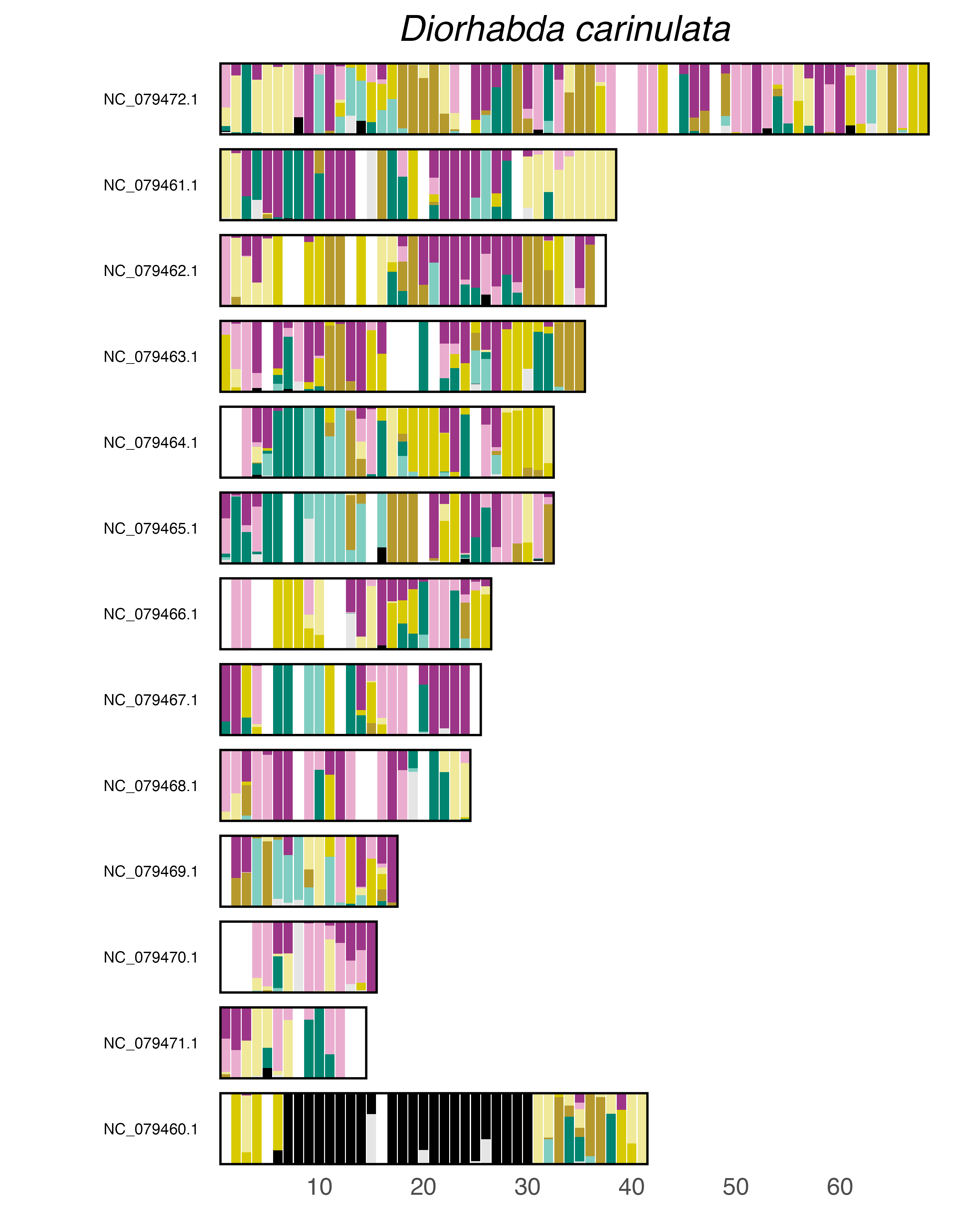

### plot.syngraph_windowbp_1Mb_rel_vertical_chrsize_Dorcus_parallelipipedus.png

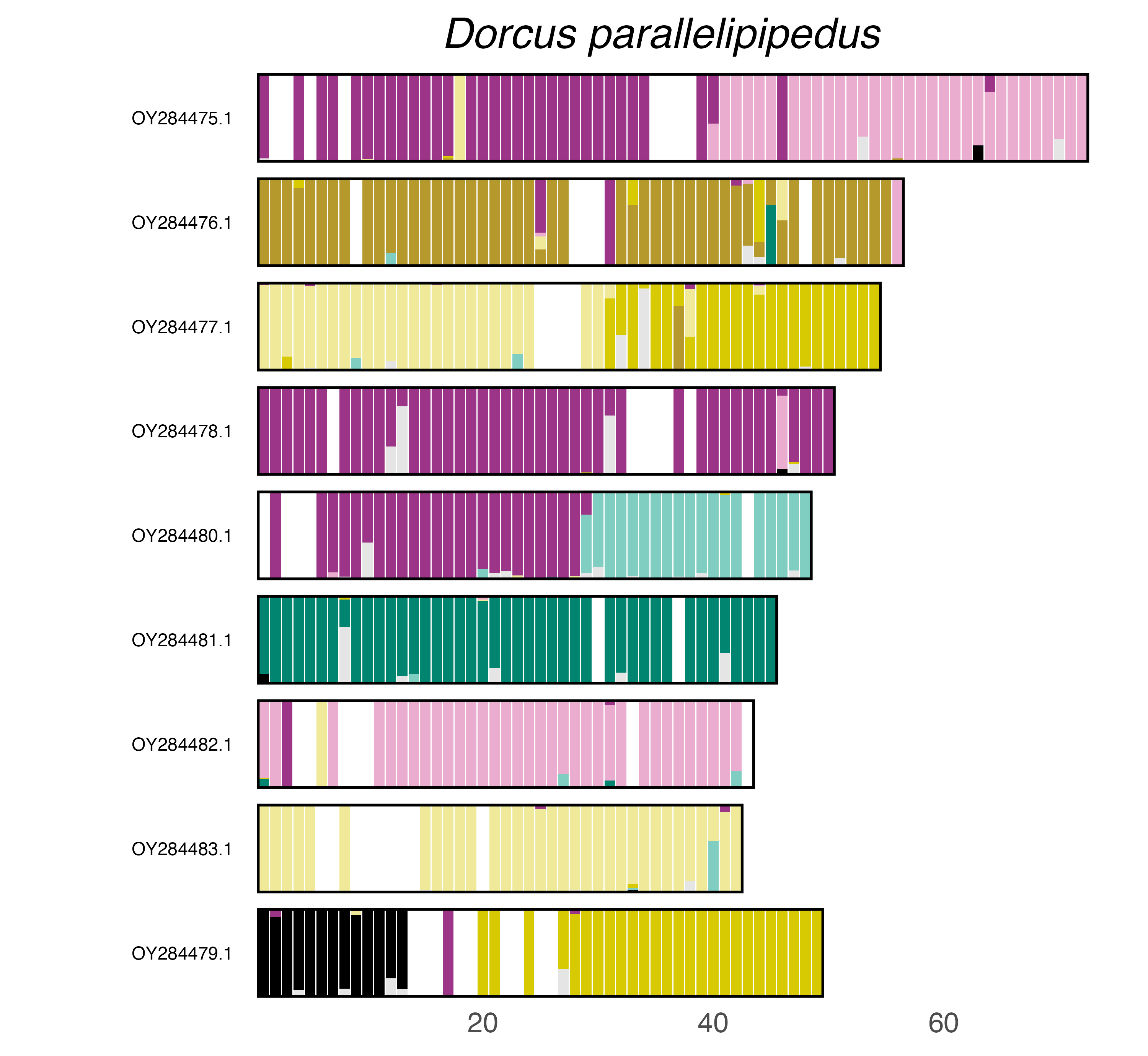

### plot.syngraph_windowbp_1Mb_rel_vertical_chrsize_Drilus_flavescens.png

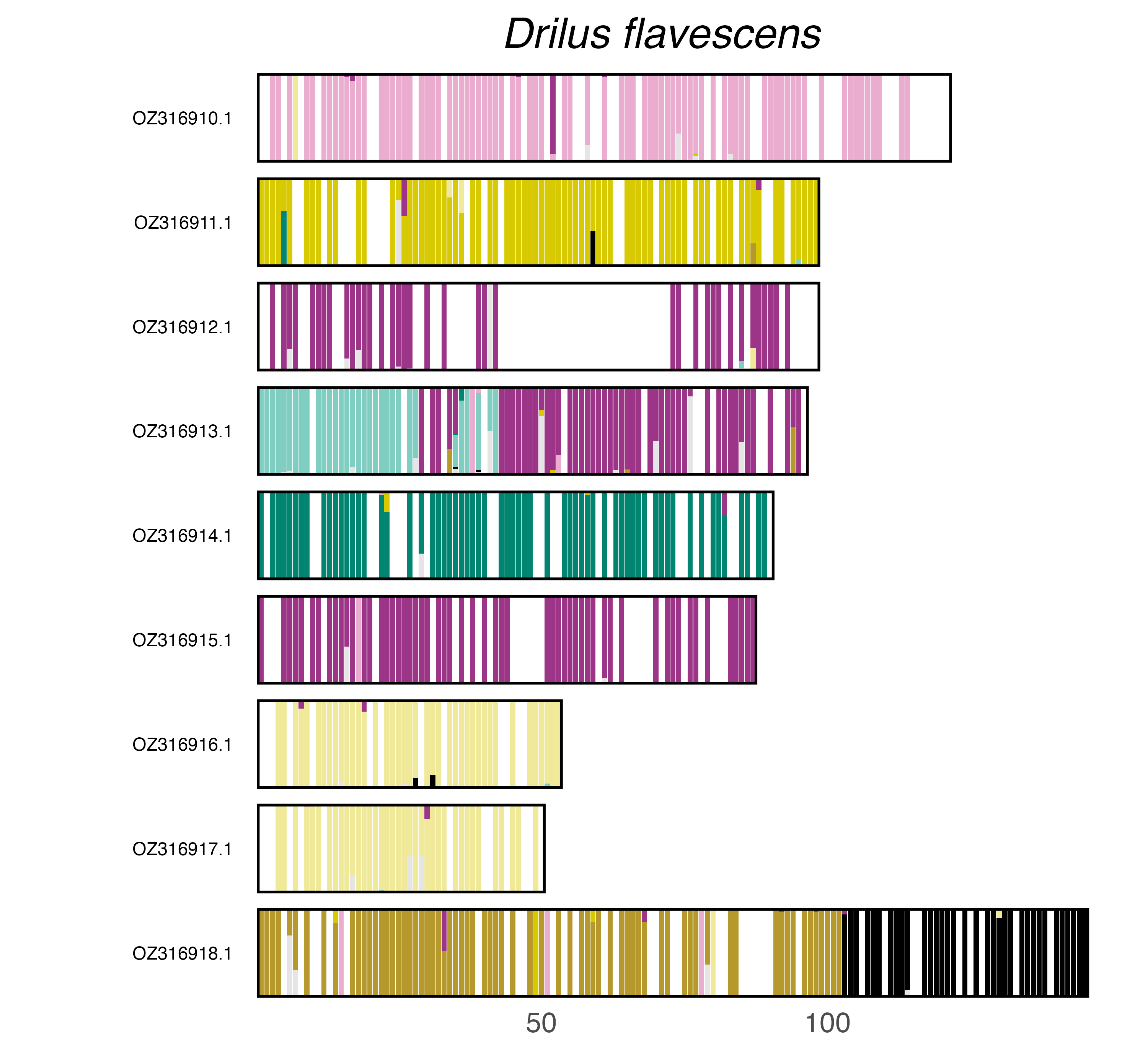

### plot.syngraph_windowbp_1Mb_rel_vertical_chrsize_Dromius_quadrimaculatus.png

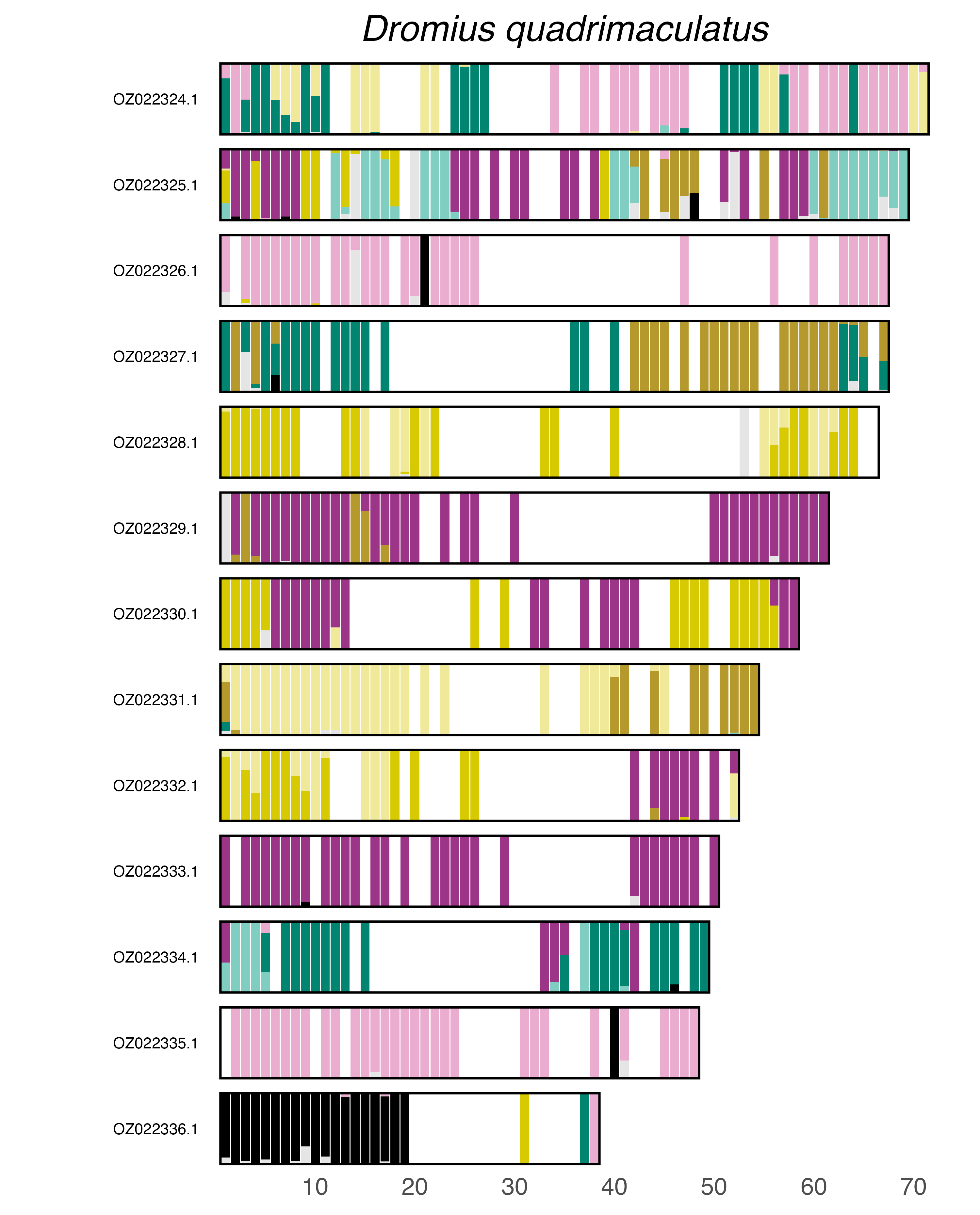

### plot.syngraph_windowbp_1Mb_rel_vertical_chrsize_Elmis_aenea.png

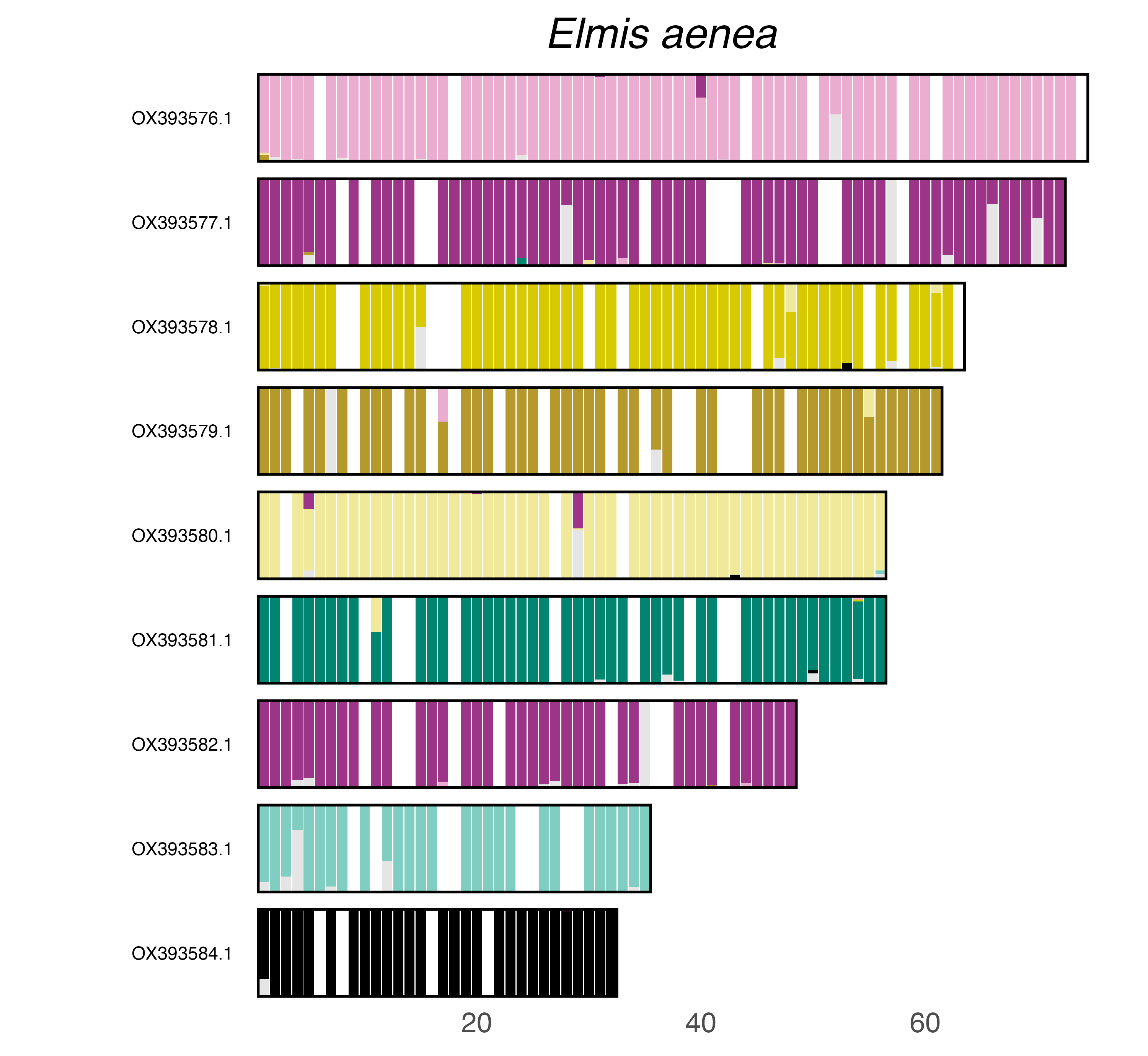

### plot.syngraph_windowbp_1Mb_rel_vertical_chrsize_Eucryptorrhynchus_scrobiculatus.png

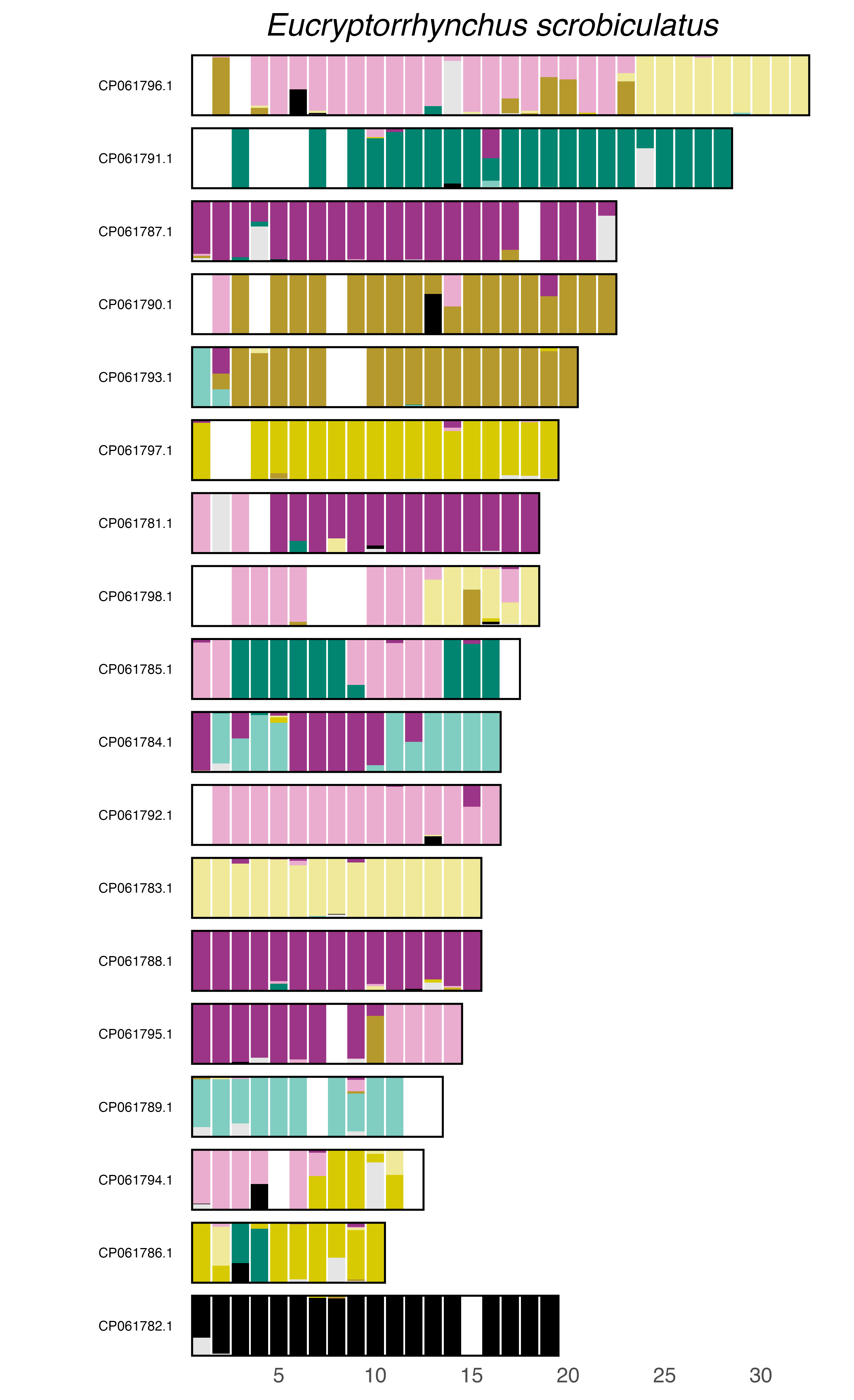

### plot.syngraph_windowbp_1Mb_rel_vertical_chrsize_Euwallacea_fornicatus.png

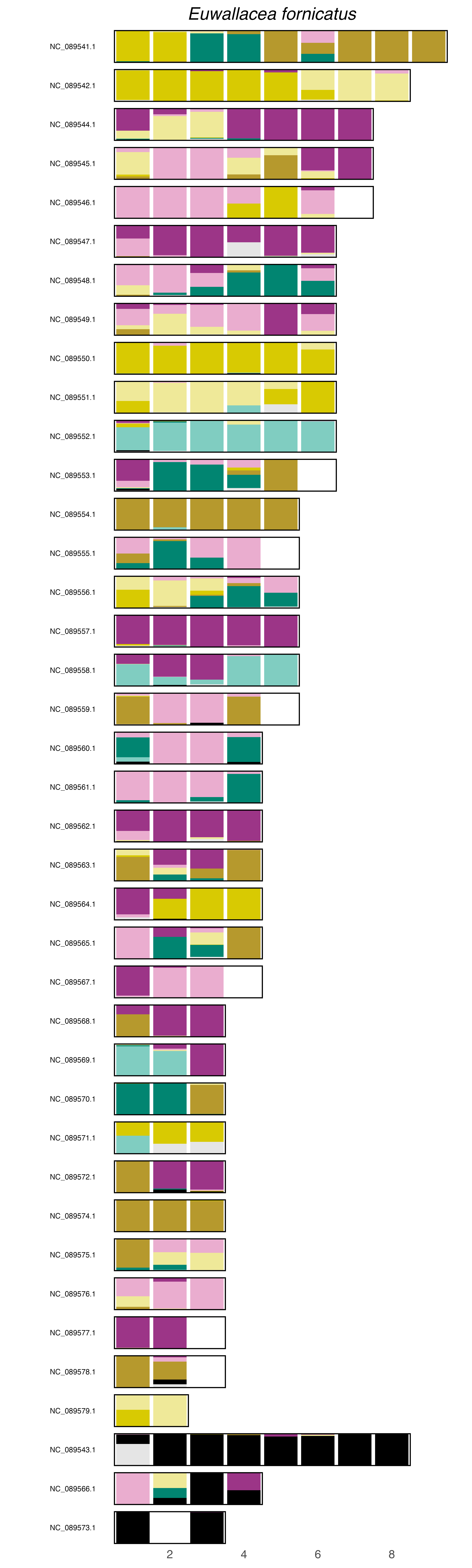

### plot.syngraph_windowbp_1Mb_rel_vertical_chrsize_Galerucella_nymphaeae.png

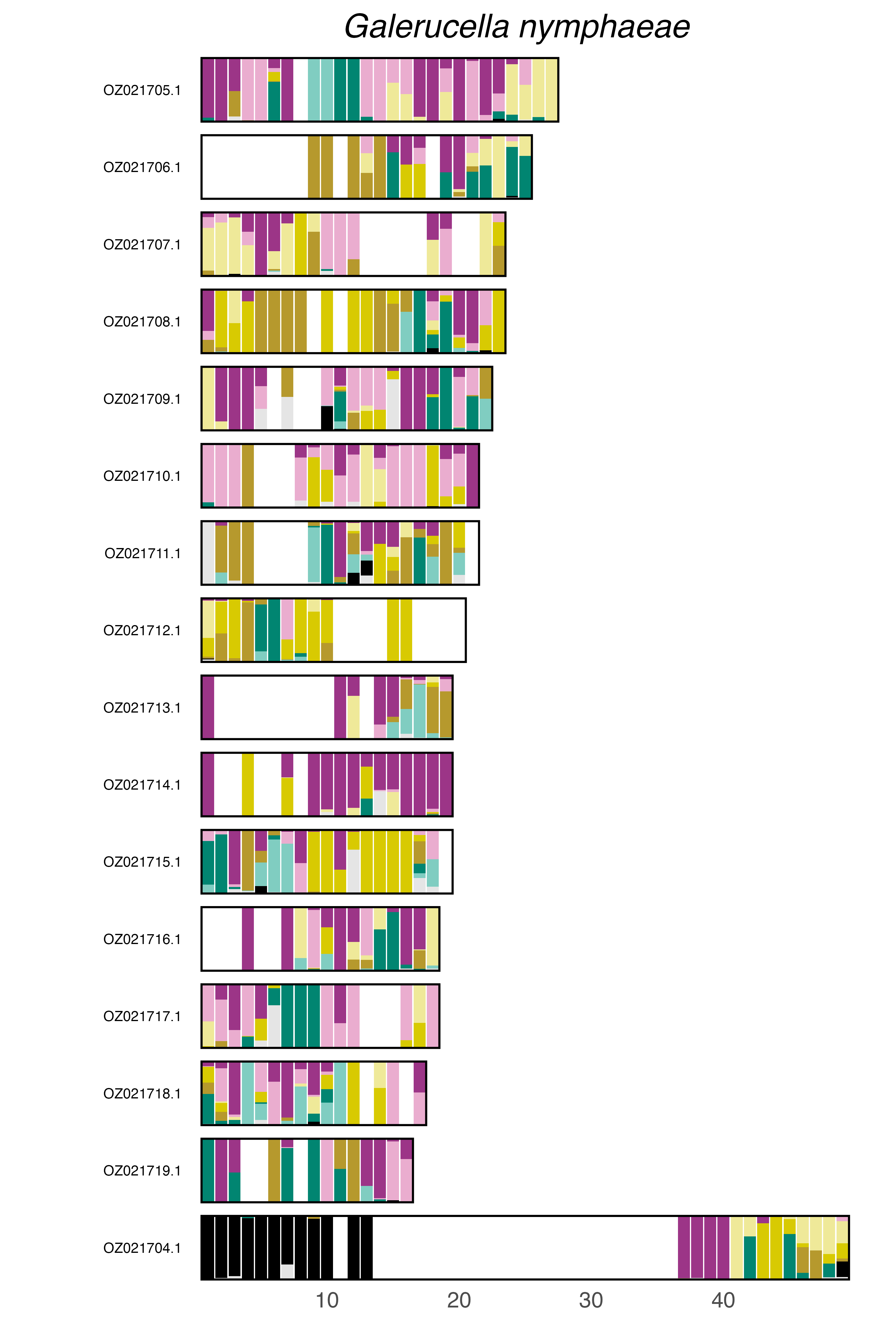

### plot.syngraph_windowbp_1Mb_rel_vertical_chrsize_Gastrophysa_polygoni.png

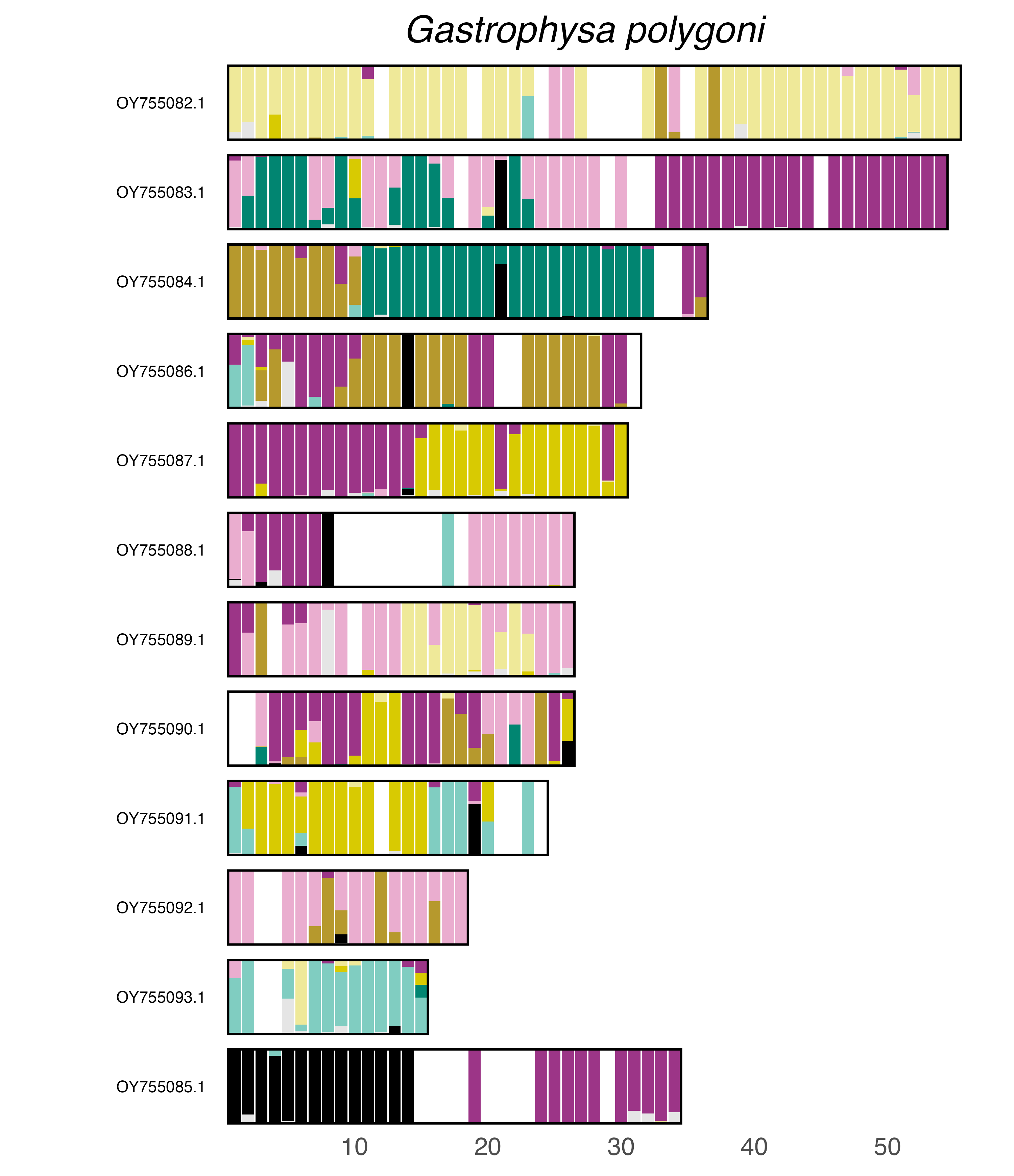

### plot.syngraph_windowbp_1Mb_rel_vertical_chrsize_Hallomenus_binotatus.png

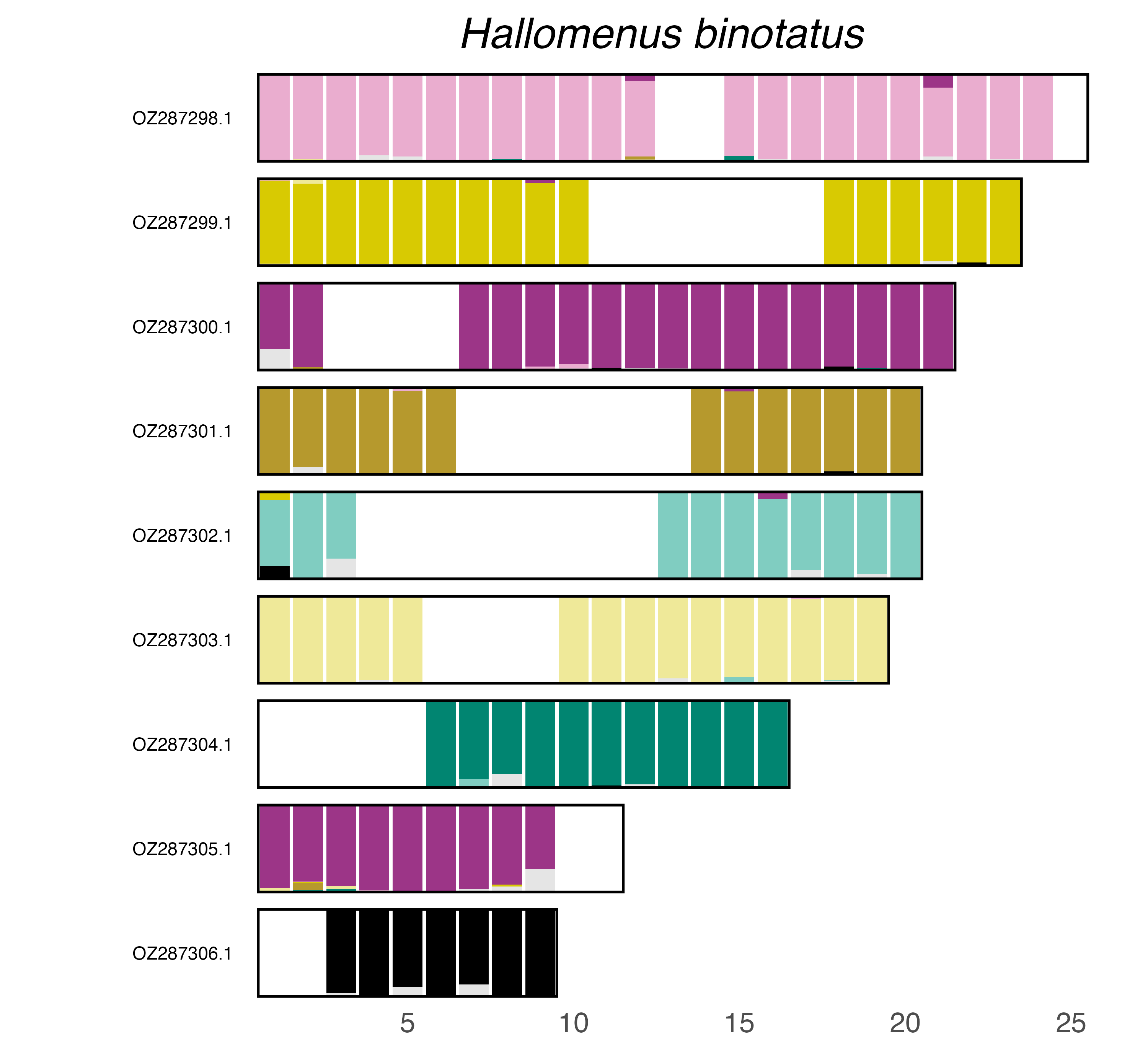

### plot.syngraph_windowbp_1Mb_rel_vertical_chrsize_Halyzia_sedecimguttata.png

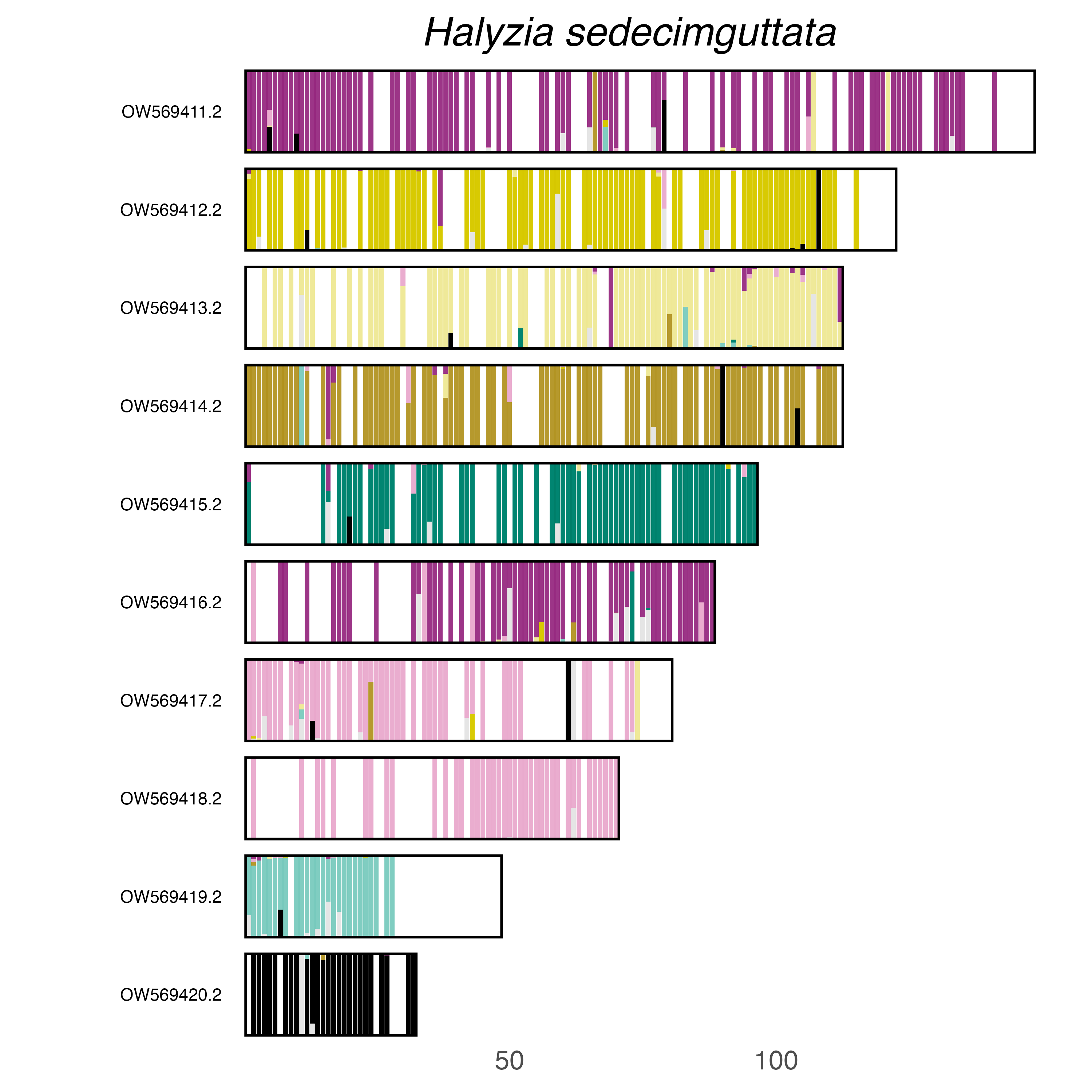

### plot.syngraph_windowbp_1Mb_rel_vertical_chrsize_Harmonia_axyridis.png

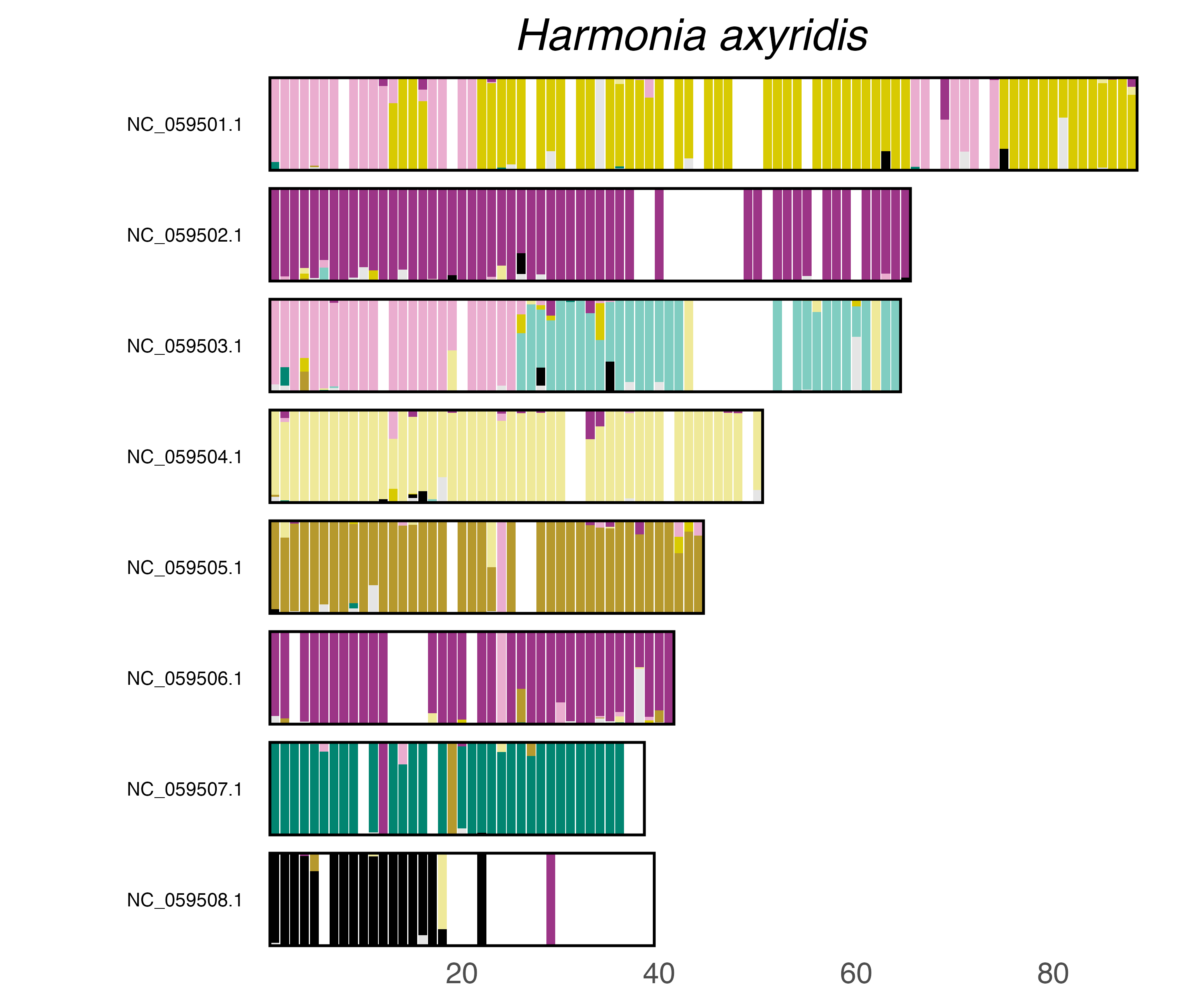

### plot.syngraph_windowbp_1Mb_rel_vertical_chrsize_Henosepilachna_vigintioctomaculata.png

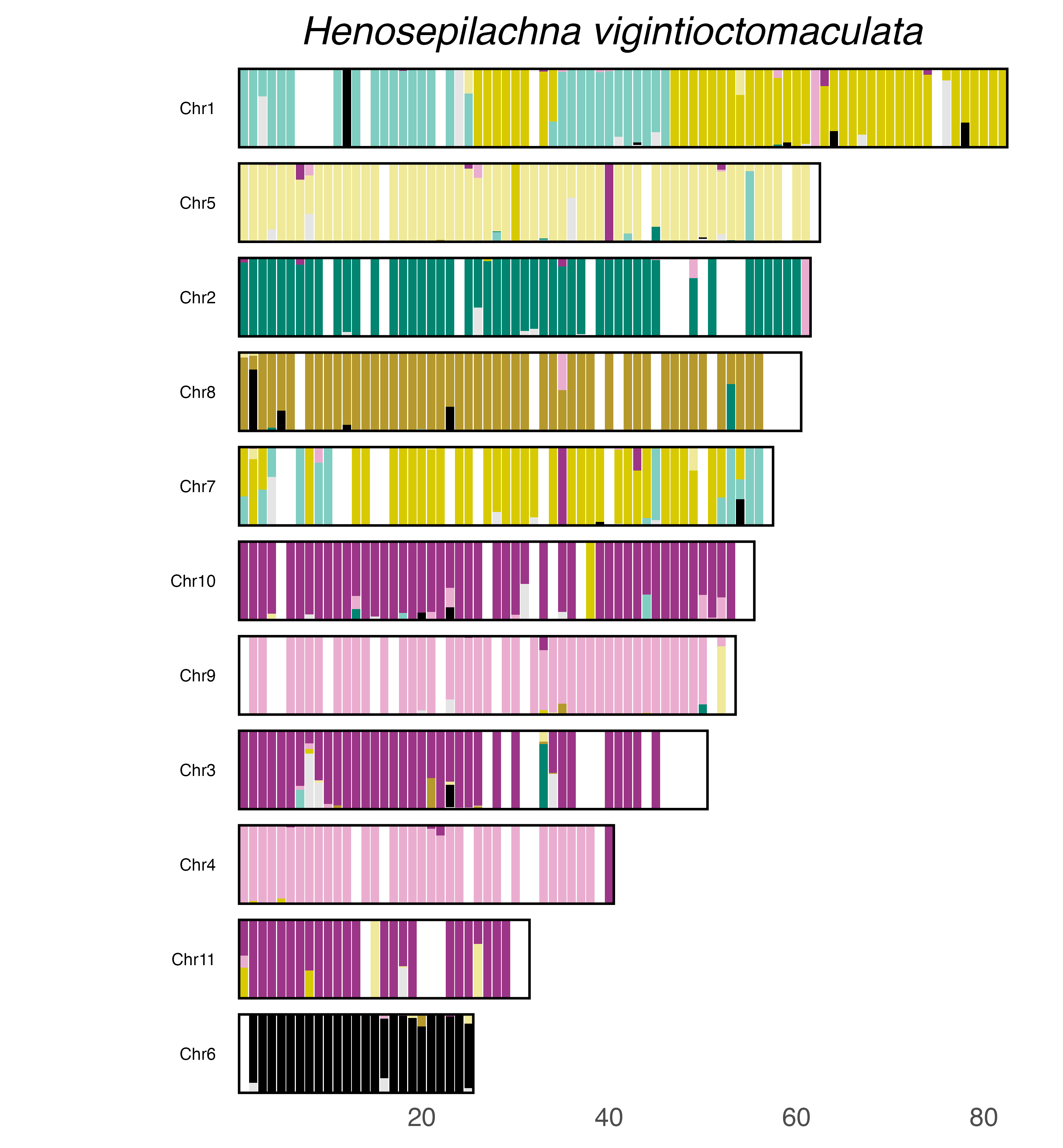

### plot.syngraph_windowbp_1Mb_rel_vertical_chrsize_Hippodamia_convergens.png

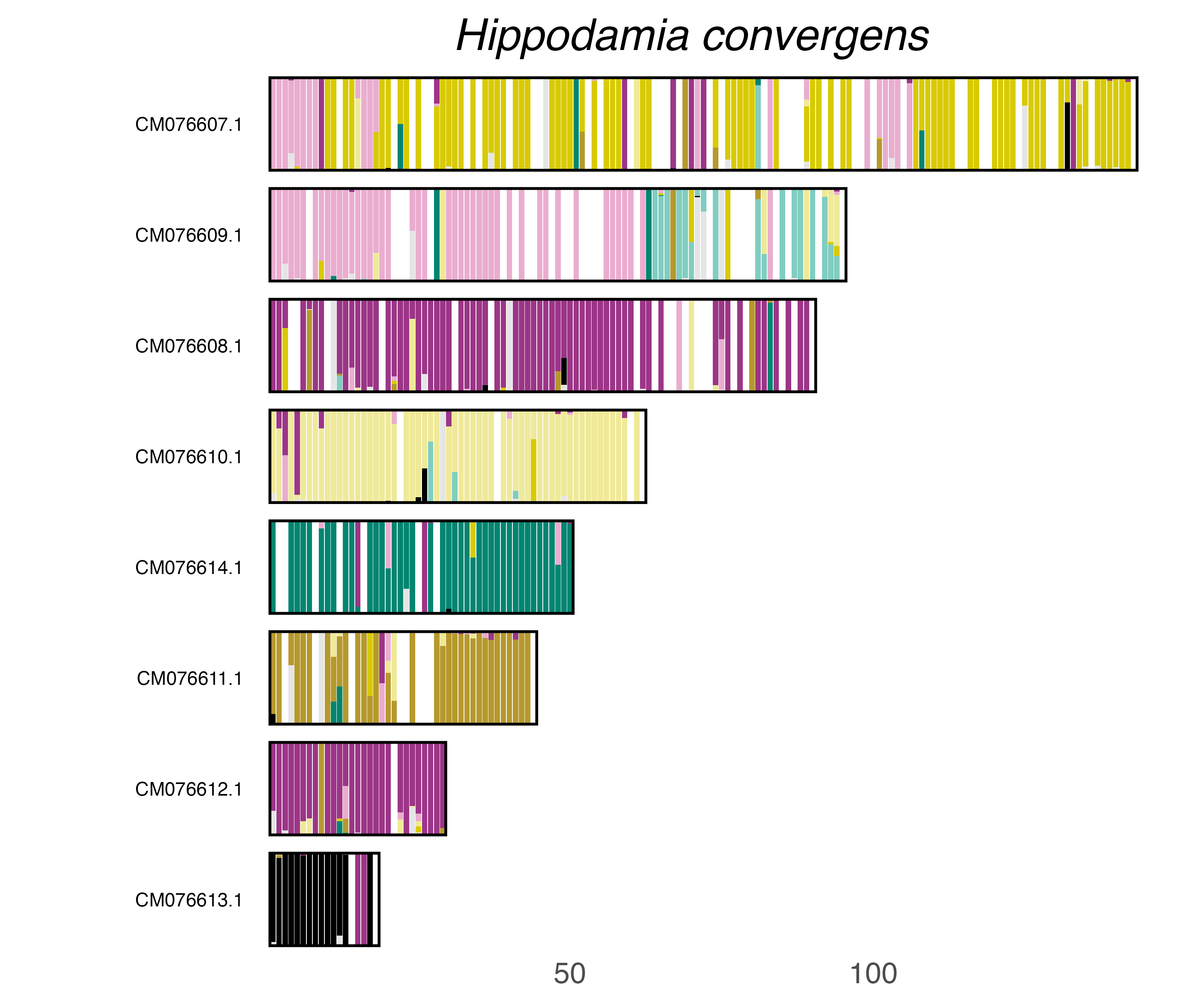

### plot.syngraph_windowbp_1Mb_rel_vertical_chrsize_Hypera_rumicis.png

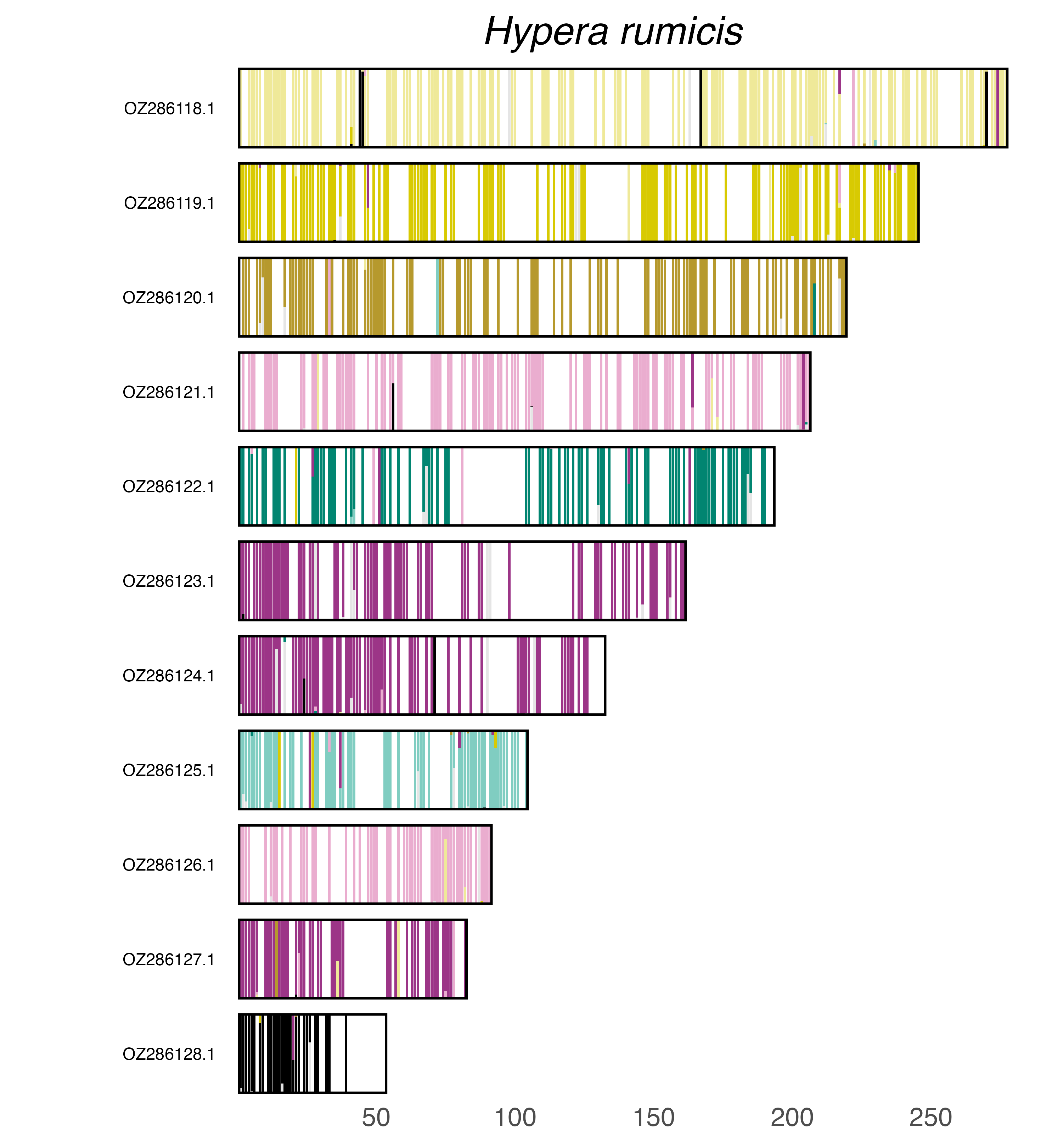

### plot.syngraph_windowbp_1Mb_rel_vertical_chrsize_Ips_sexdentatus.png

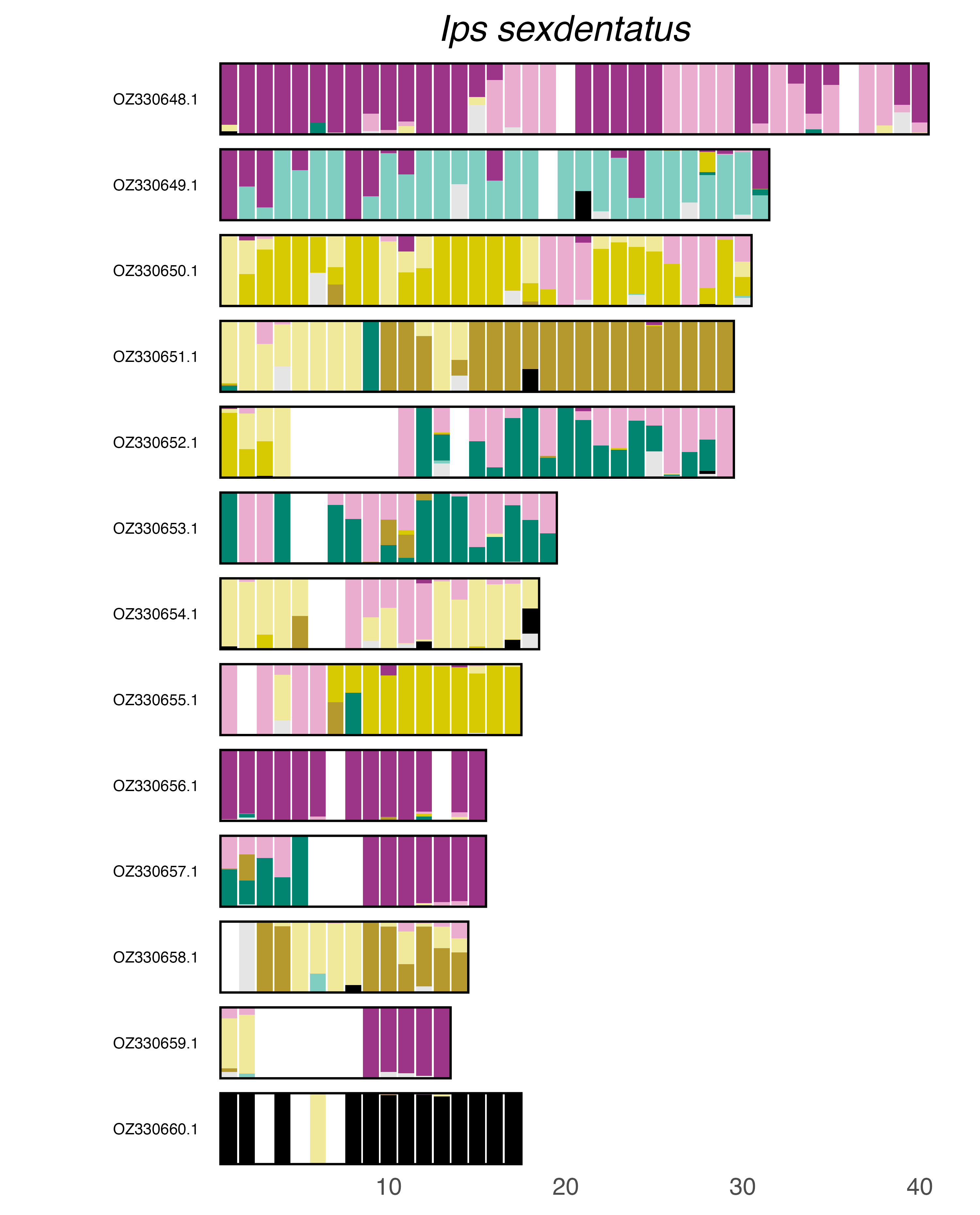

### plot.syngraph_windowbp_1Mb_rel_vertical_chrsize_Ips_typographus.png

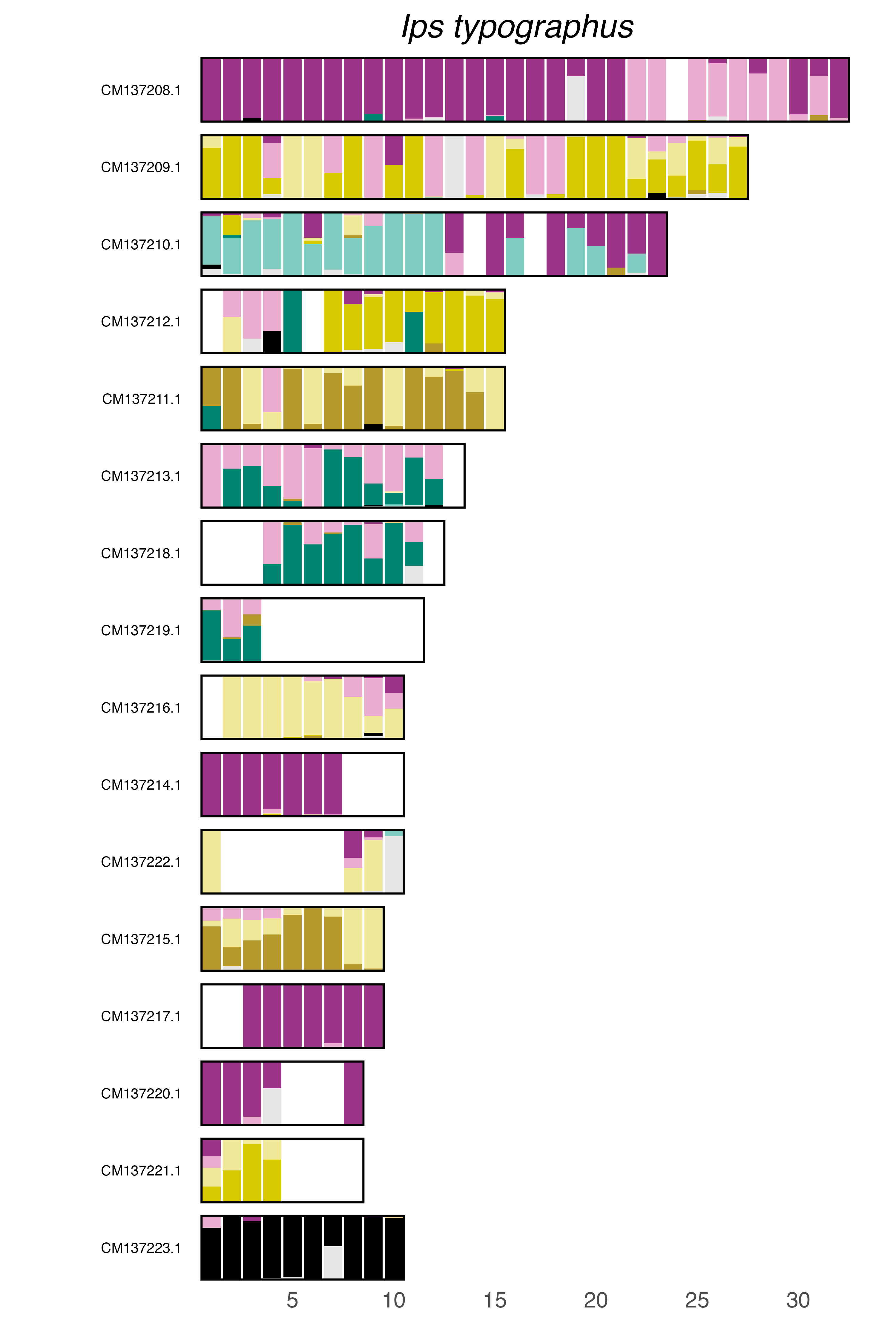

### plot.syngraph_windowbp_1Mb_rel_vertical_chrsize_Laccobius_striatulus.png

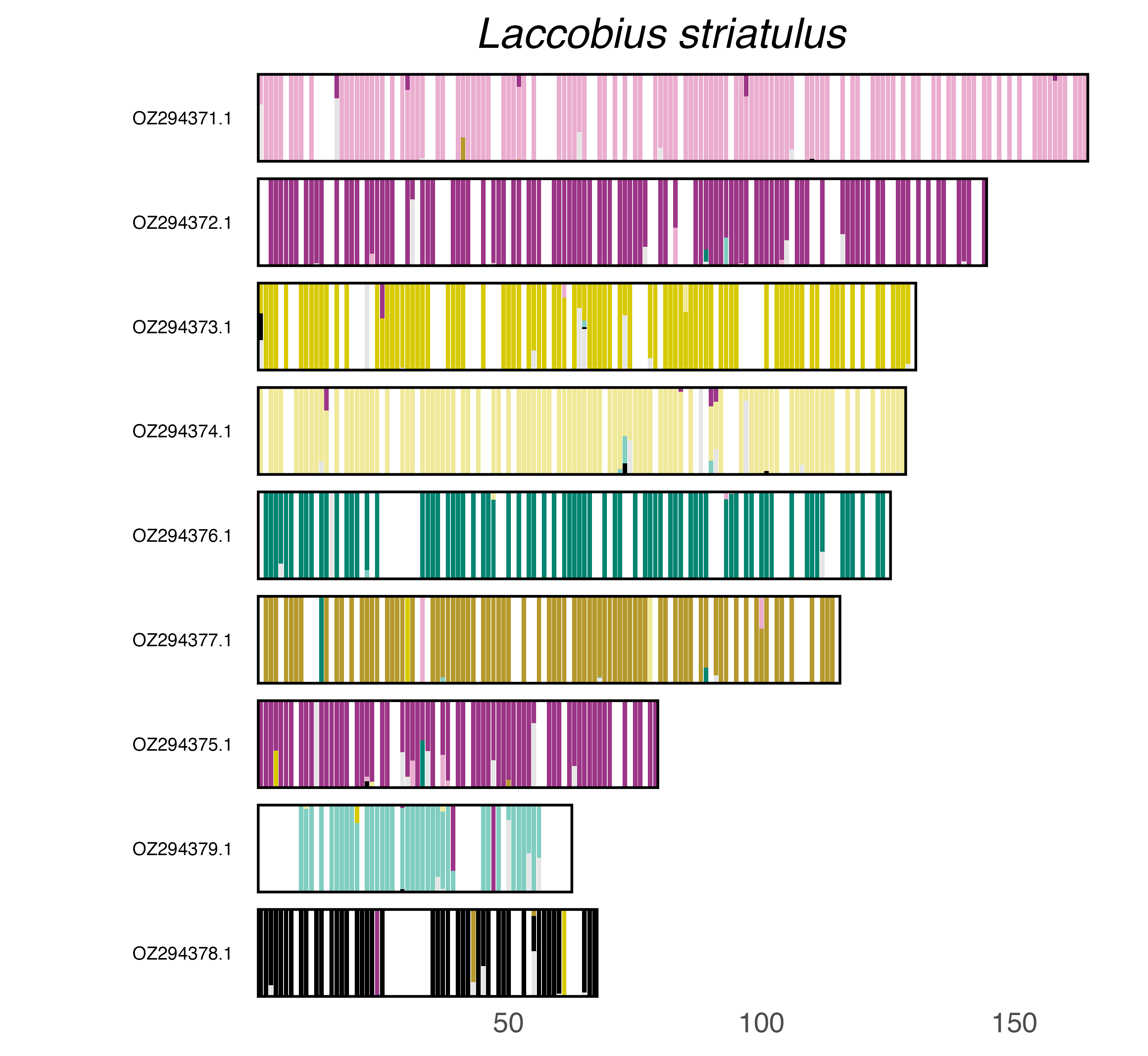

### plot.syngraph_windowbp_1Mb_rel_vertical_chrsize_Lasiorhynchites_cavifrons.png

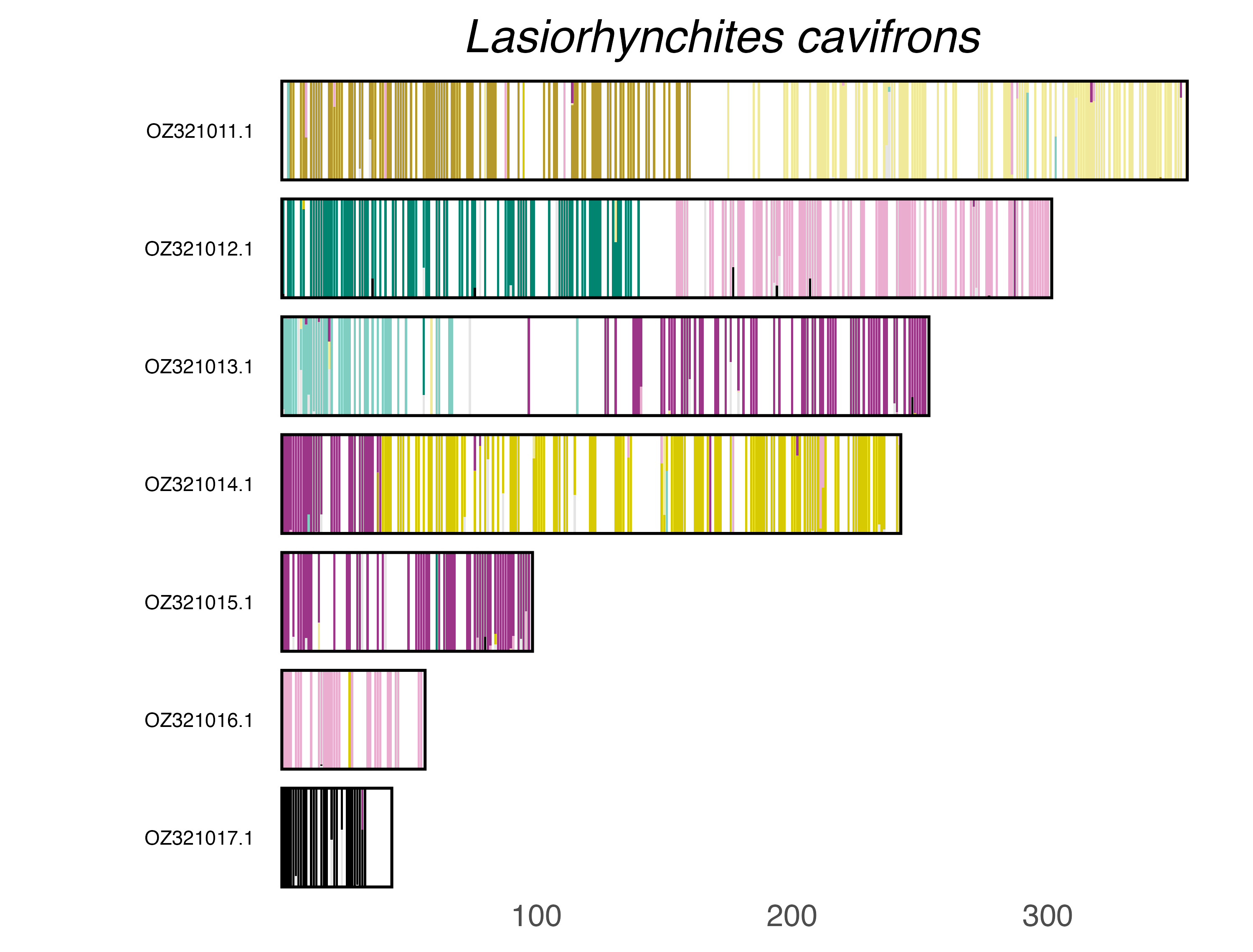

### plot.syngraph_windowbp_1Mb_rel_vertical_chrsize_Leptura_quadrifasciata.png

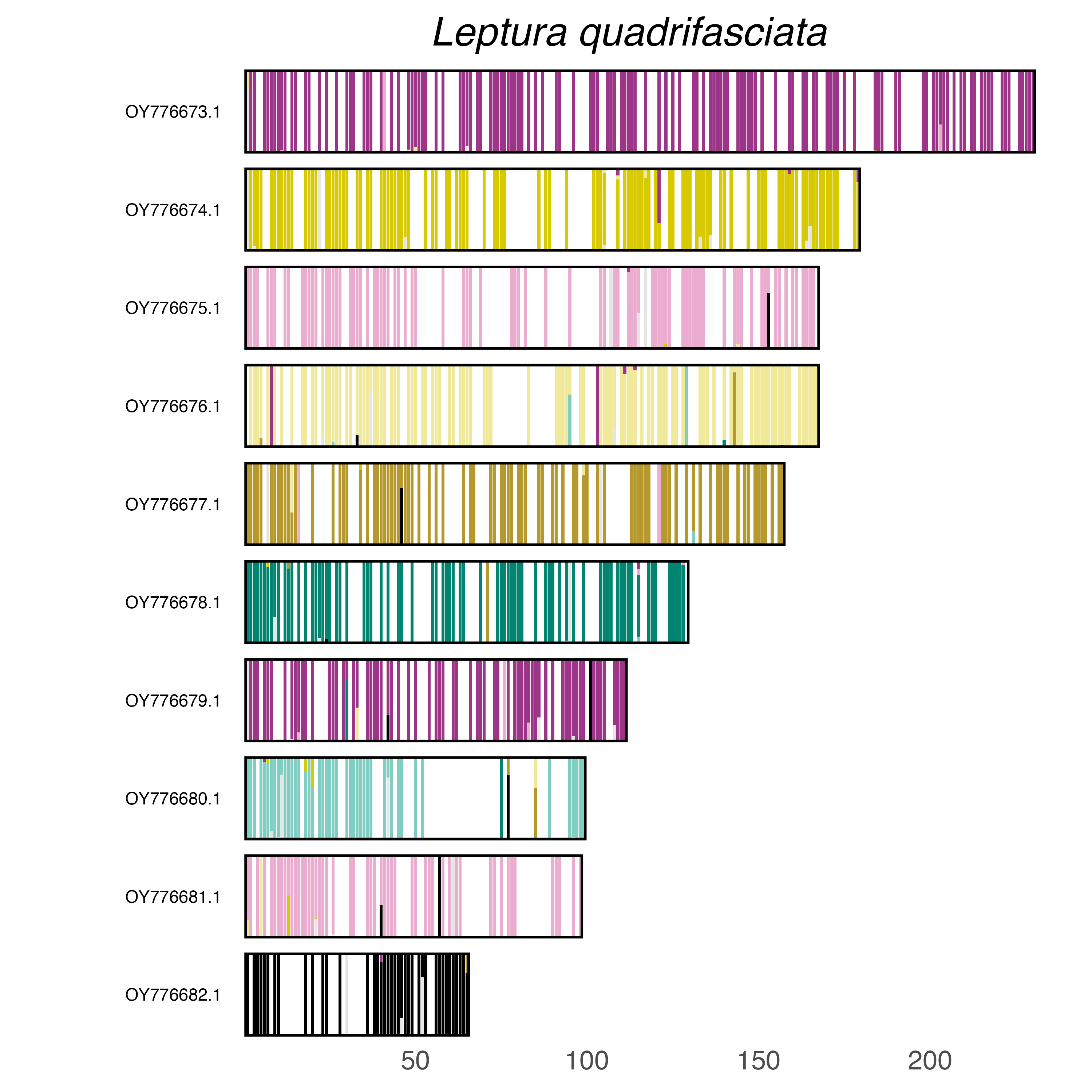

### plot.syngraph_windowbp_1Mb_rel_vertical_chrsize_Librodor_japonicus.png

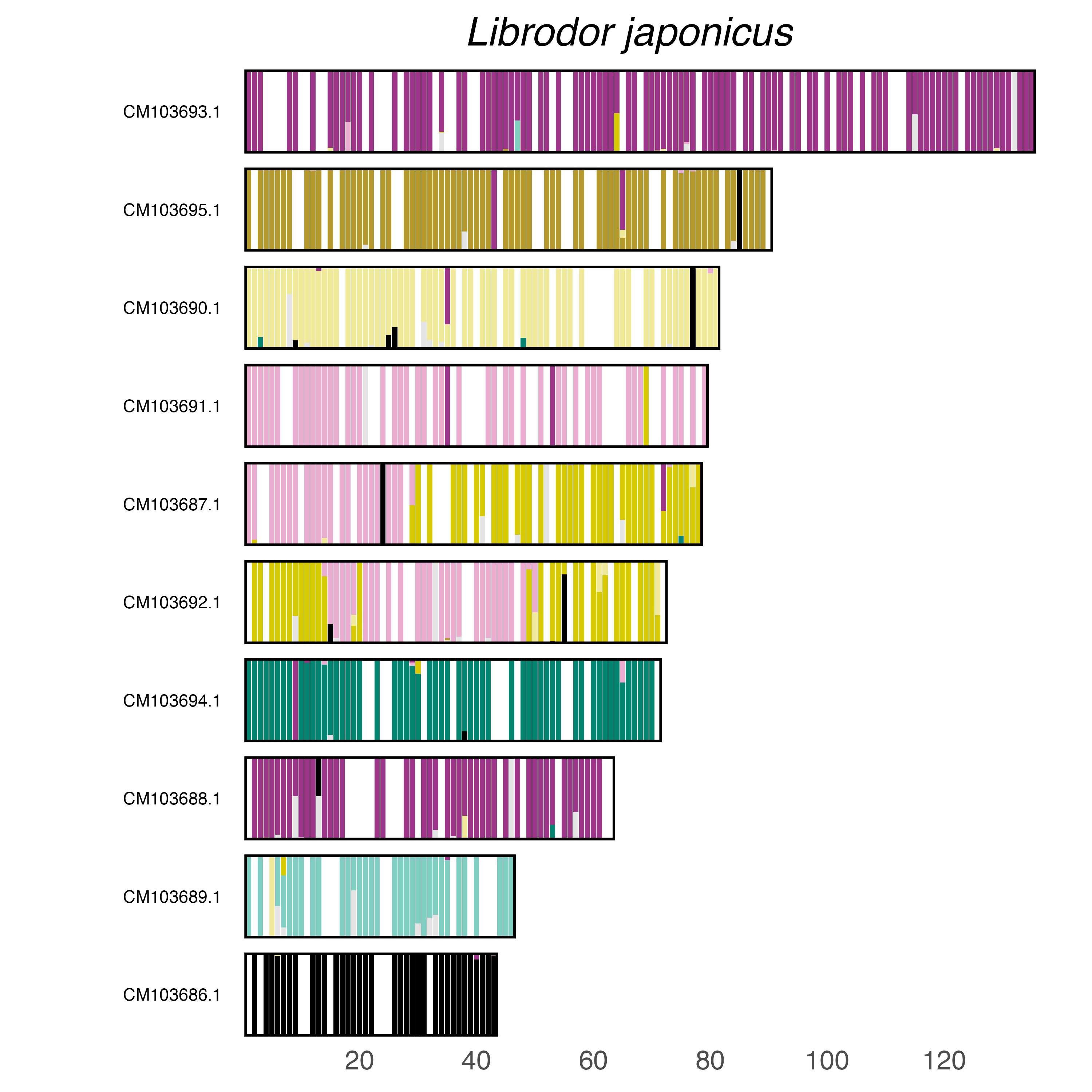

### plot.syngraph_windowbp_1Mb_rel_vertical_chrsize_Longitarsus_dorsalis.png

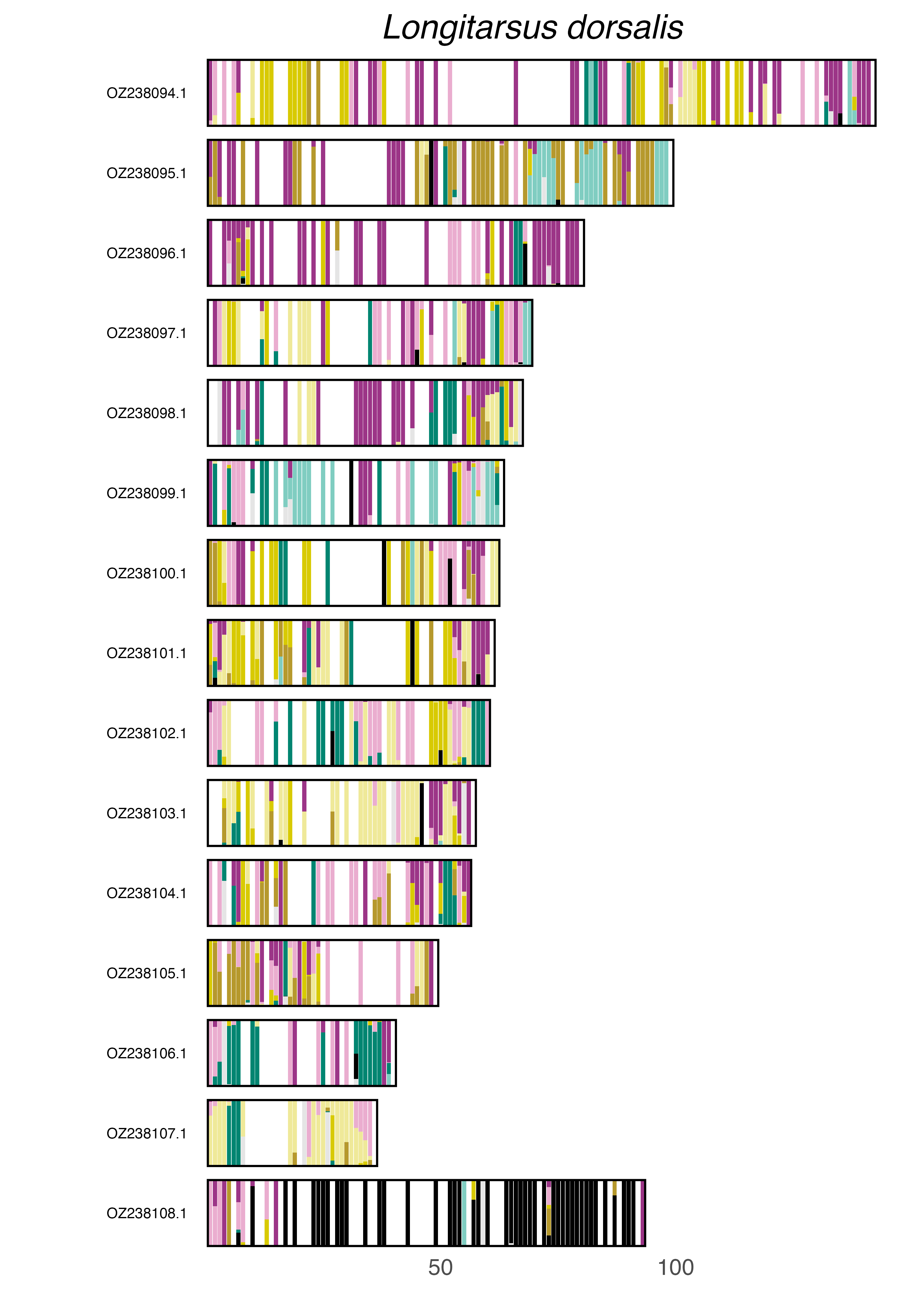

### plot.syngraph_windowbp_1Mb_rel_vertical_chrsize_Lordithon_lunulatus.png

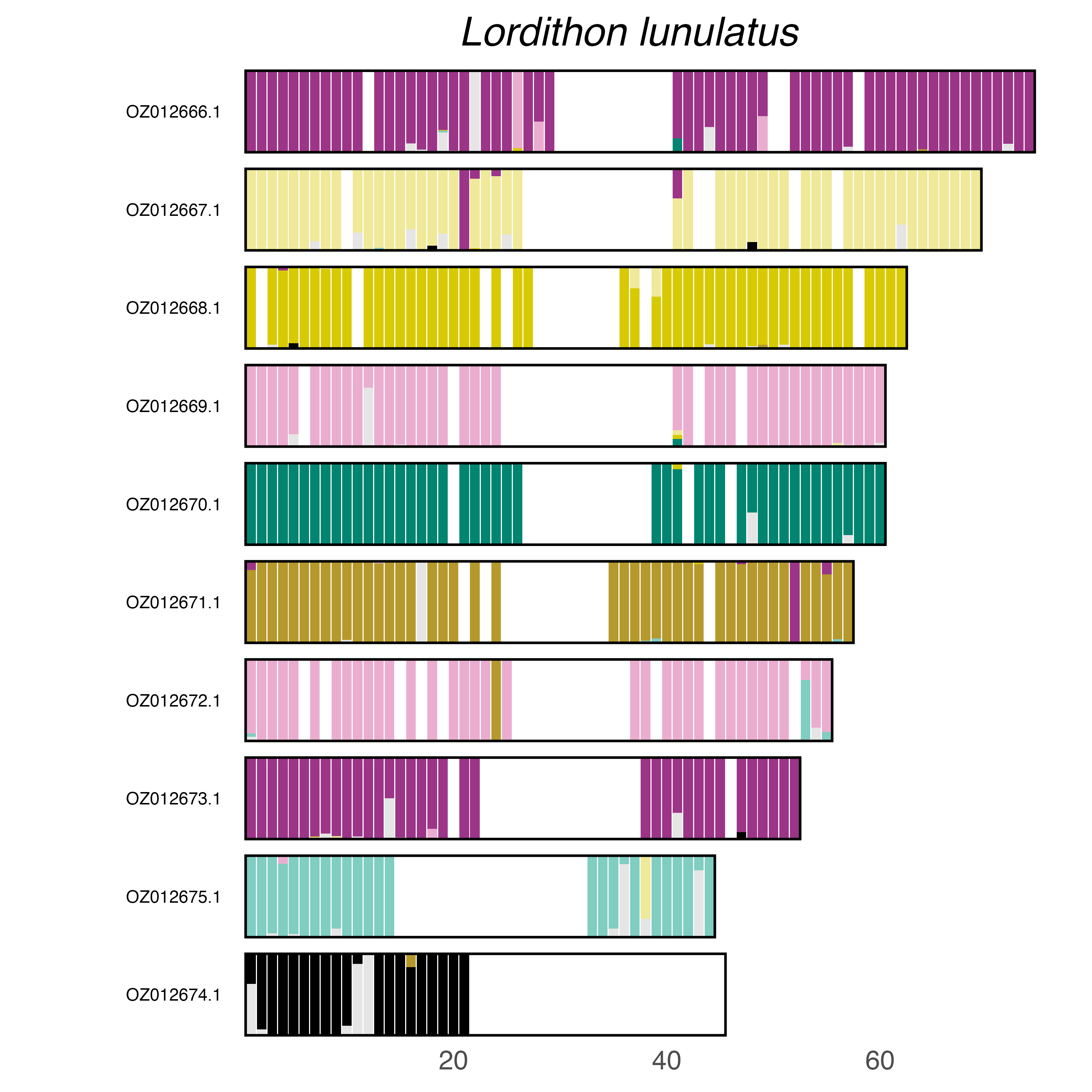

### plot.syngraph_windowbp_1Mb_rel_vertical_chrsize_Magdalis_barbicornis.png

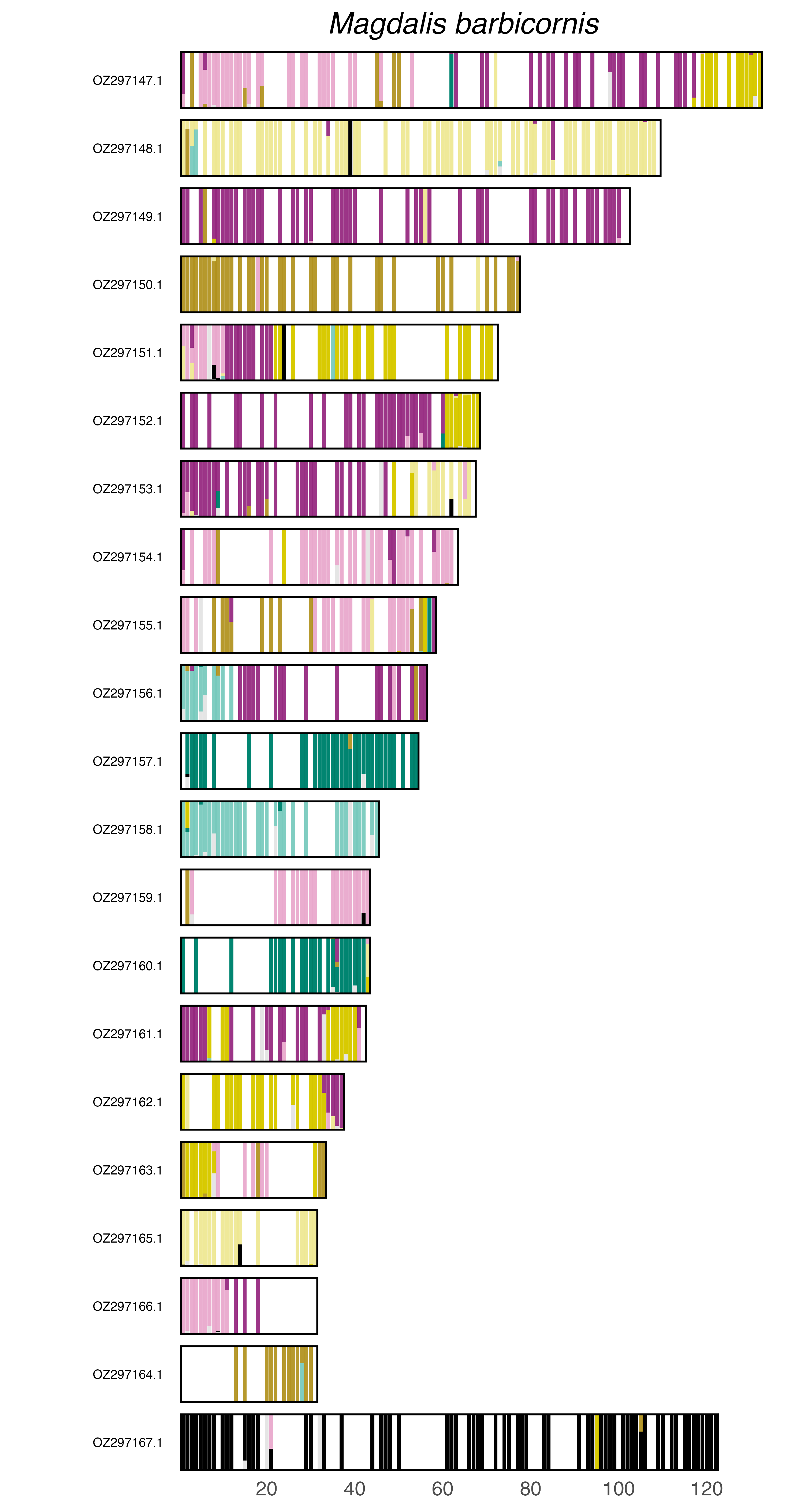

### plot.syngraph_windowbp_1Mb_rel_vertical_chrsize_Magdalis_cerasi.png

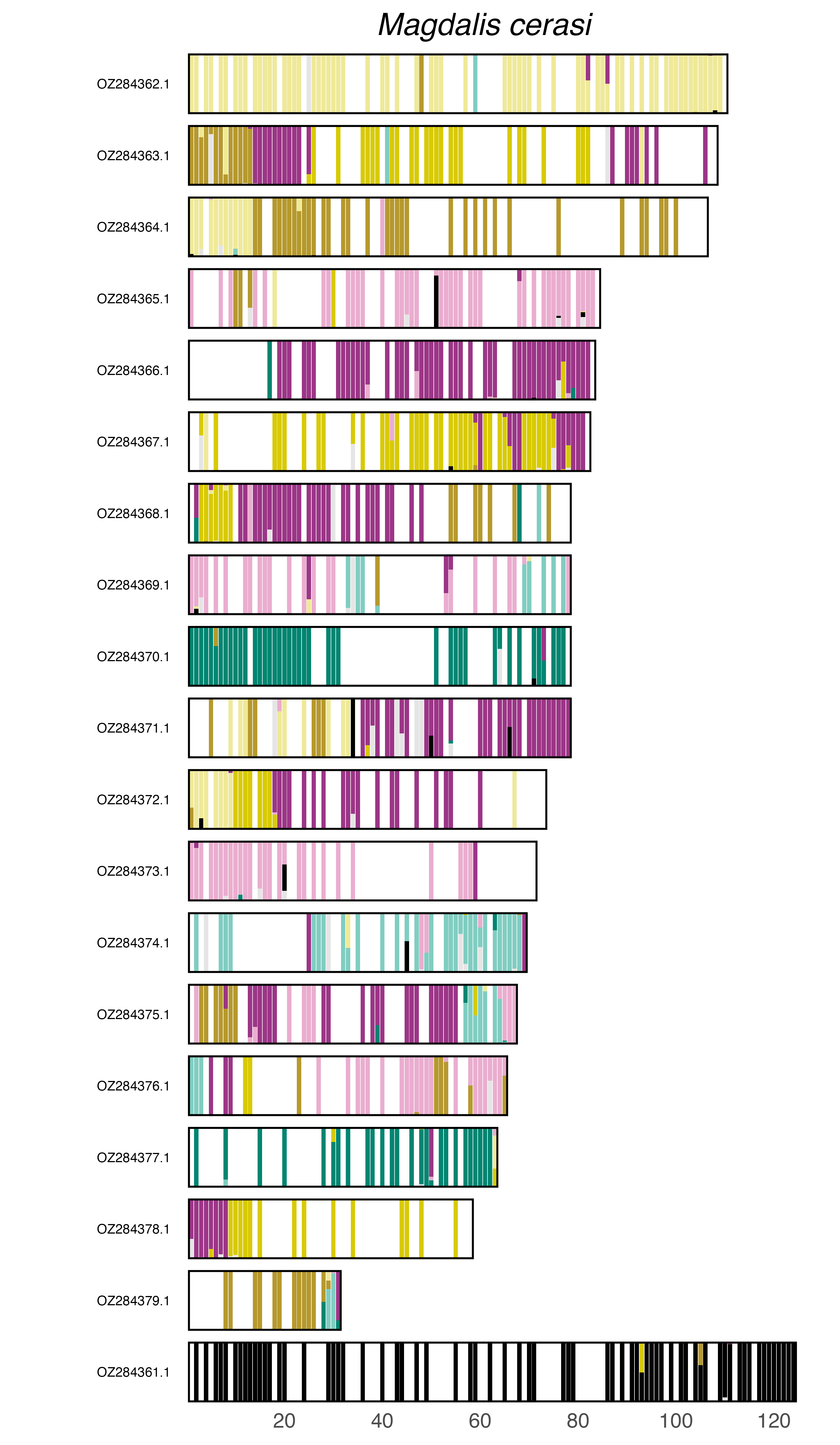

### plot.syngraph_windowbp_1Mb_rel_vertical_chrsize_Malachius_bipustulatus.png

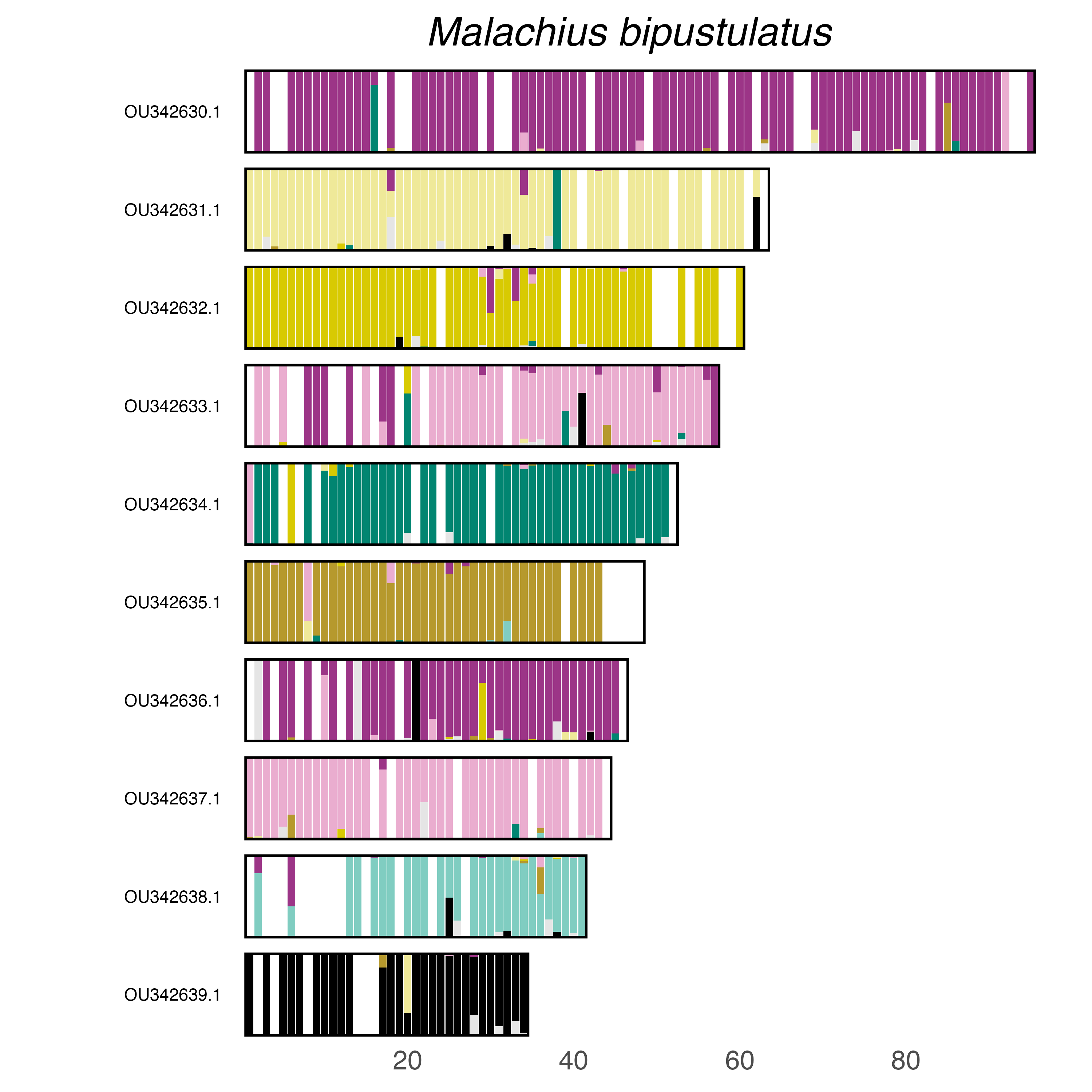

### plot.syngraph_windowbp_1Mb_rel_vertical_chrsize_Malthodes_minimus.png

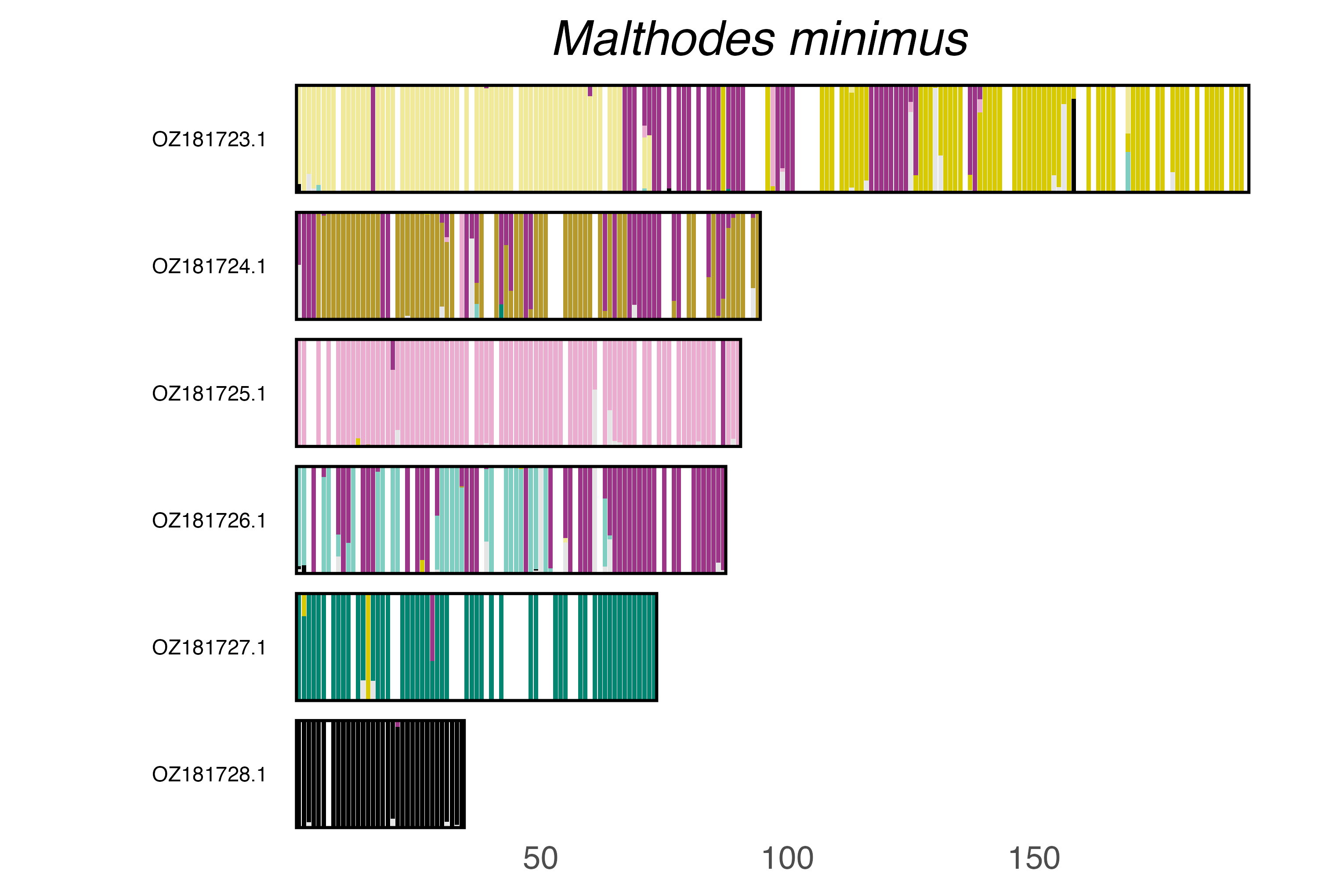

### plot.syngraph_windowbp_1Mb_rel_vertical_chrsize_Megalocaria_dilatata.png

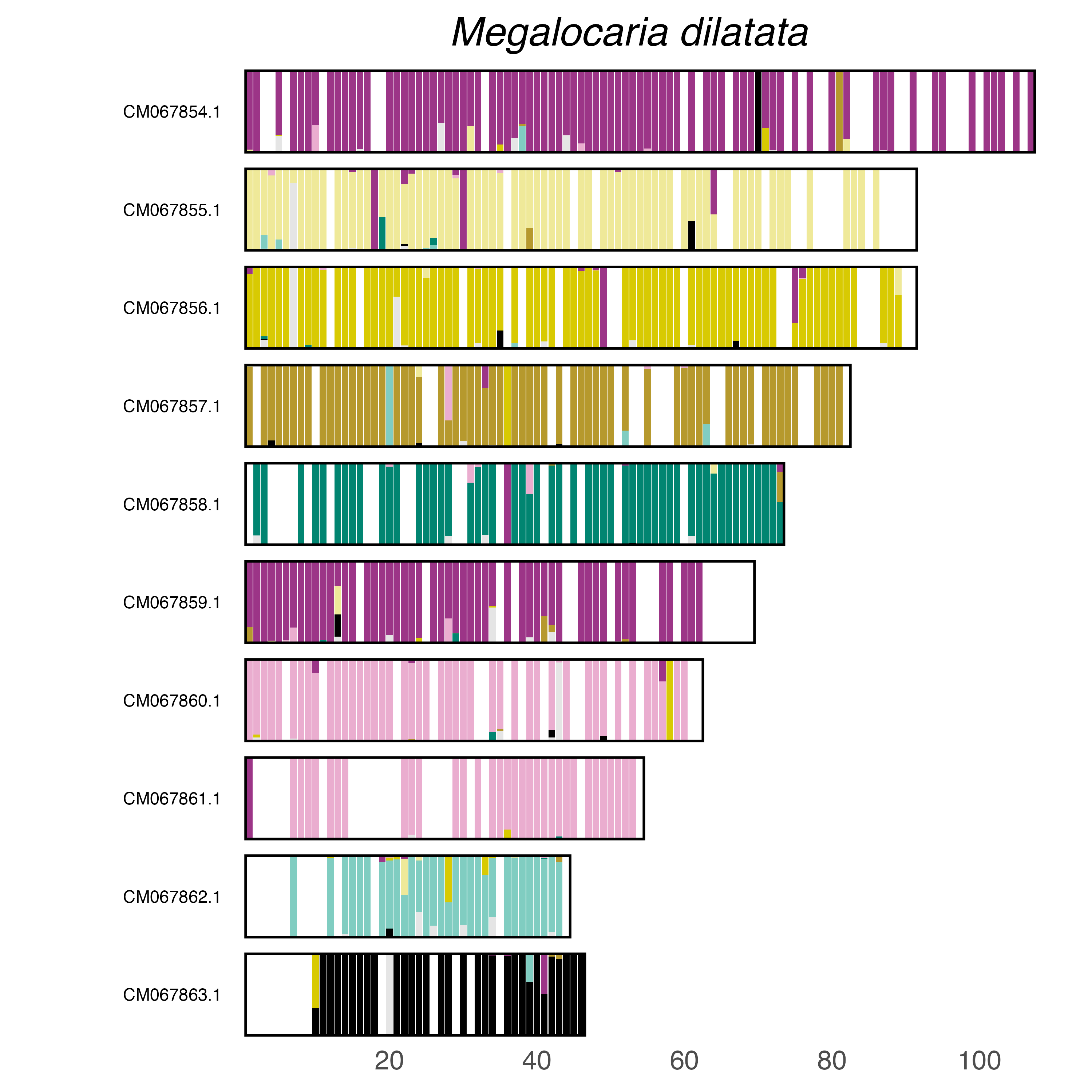

### plot.syngraph_windowbp_1Mb_rel_vertical_chrsize_Monochamus_saltuarius.png

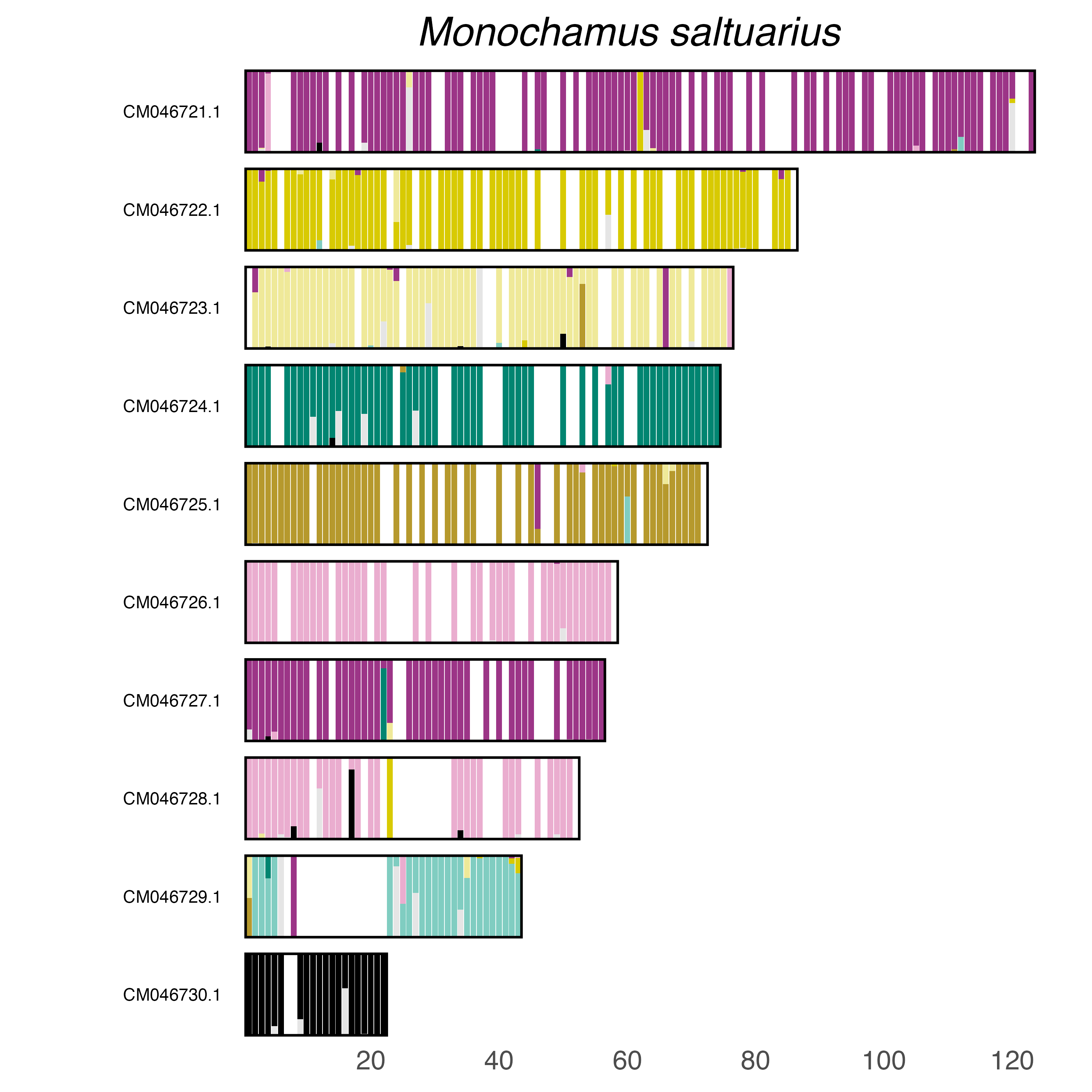

### plot.syngraph_windowbp_1Mb_rel_vertical_chrsize_Monochamus_scutellatus.png

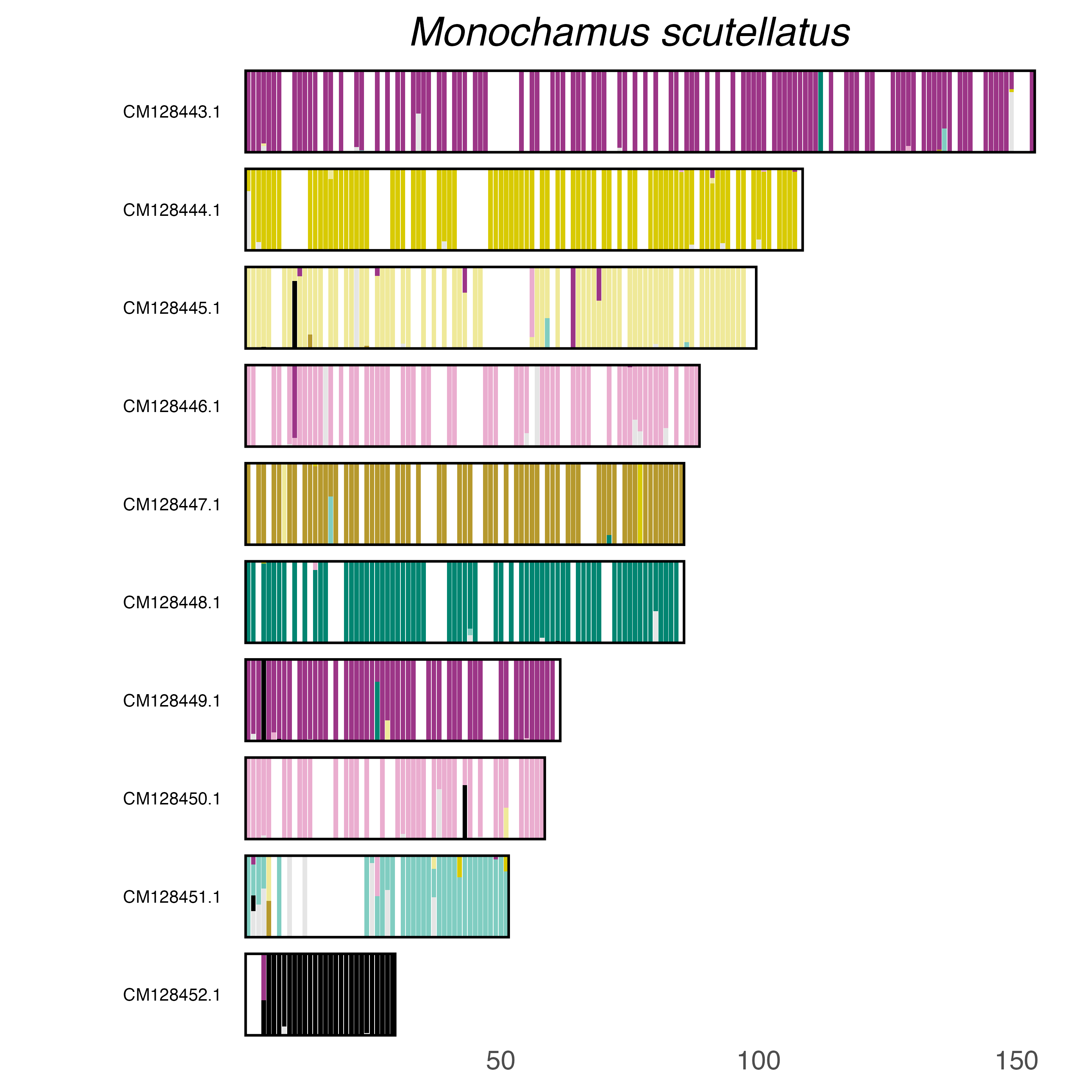

### plot.syngraph_windowbp_1Mb_rel_vertical_chrsize_Monochamus_sutor.png

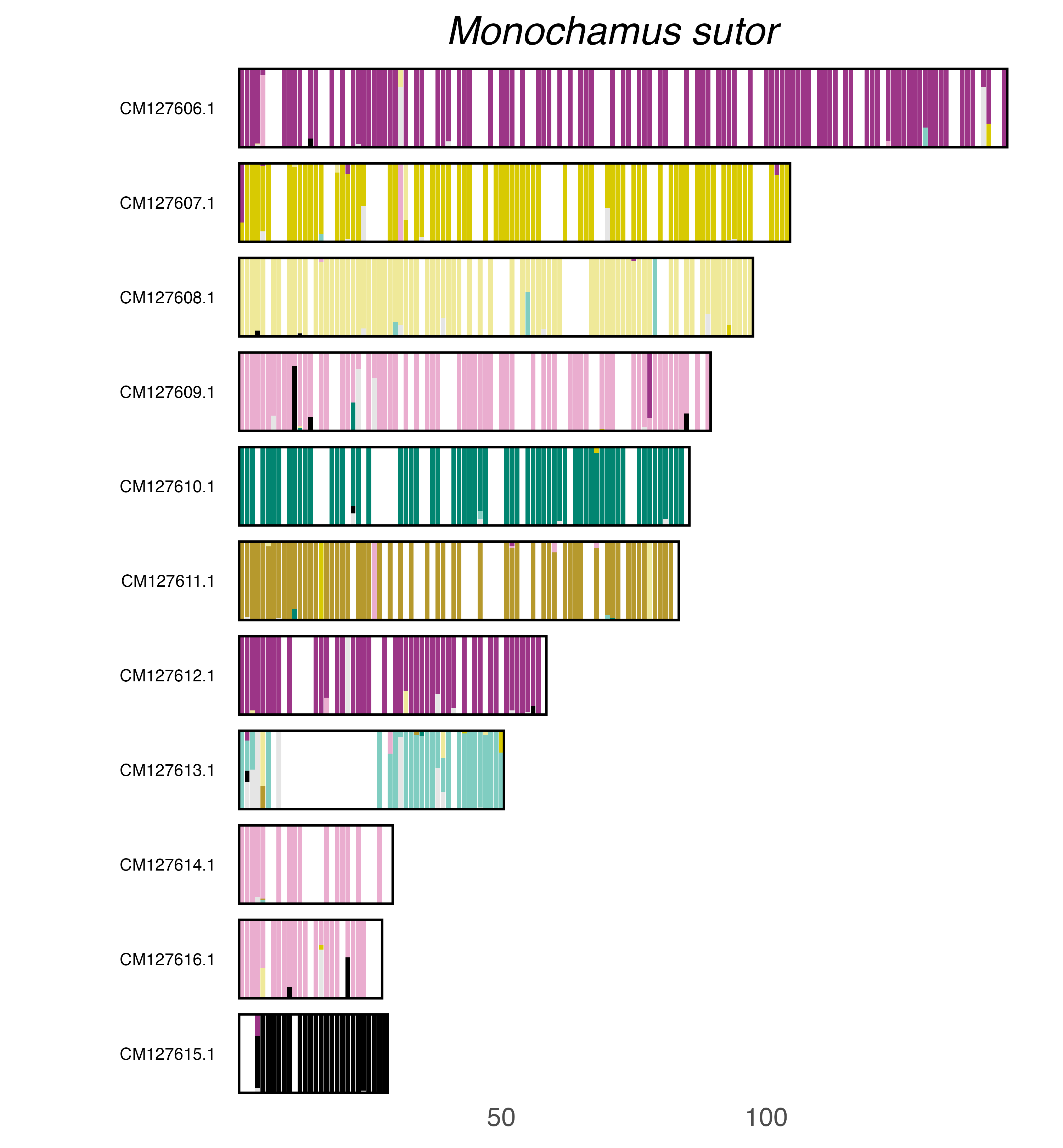

### plot.syngraph_windowbp_1Mb_rel_vertical_chrsize_Monolepta_hieroglyphica.png

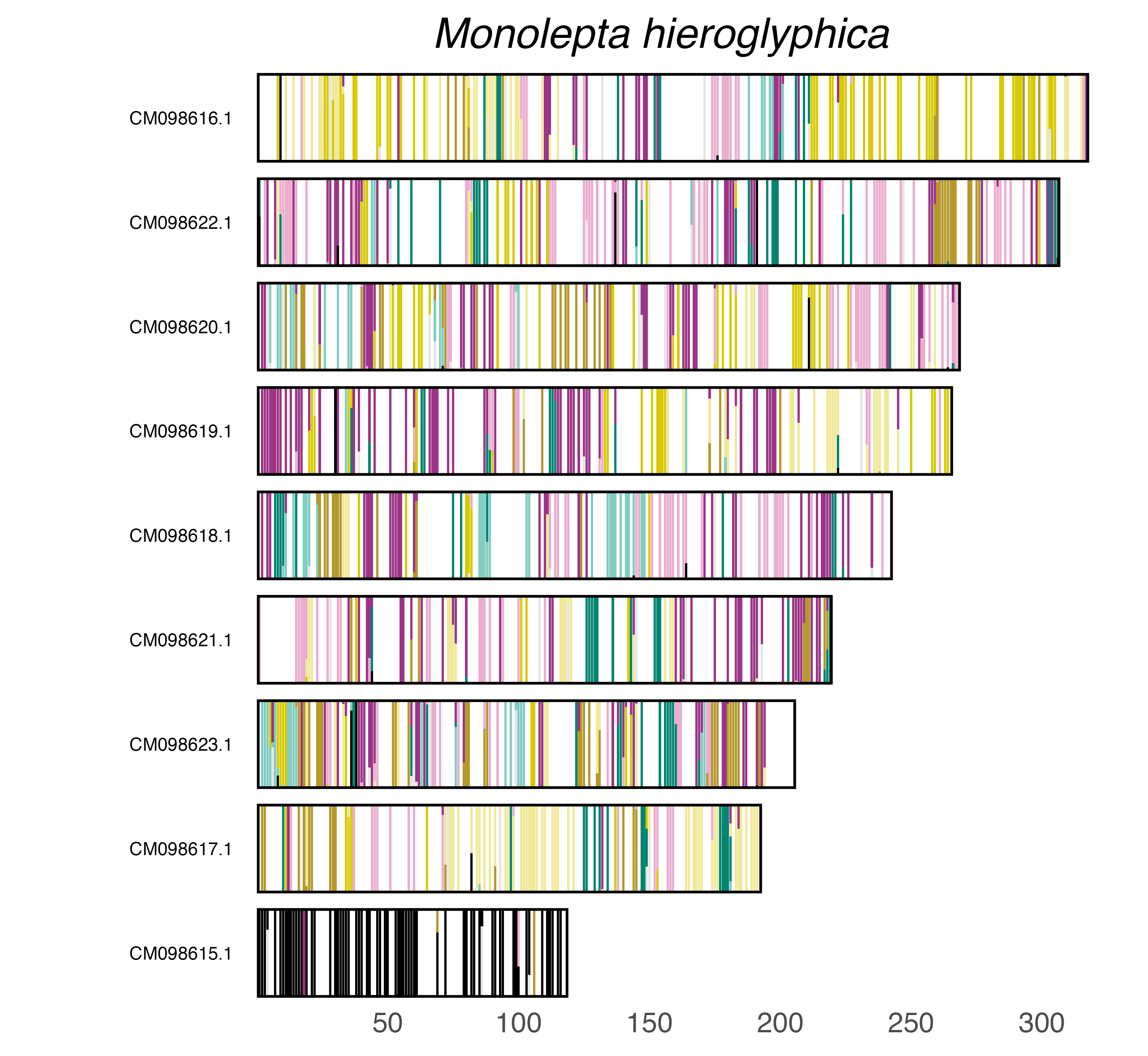

### plot.syngraph_windowbp_1Mb_rel_vertical_chrsize_Nalassus_laevioctostriatus.png

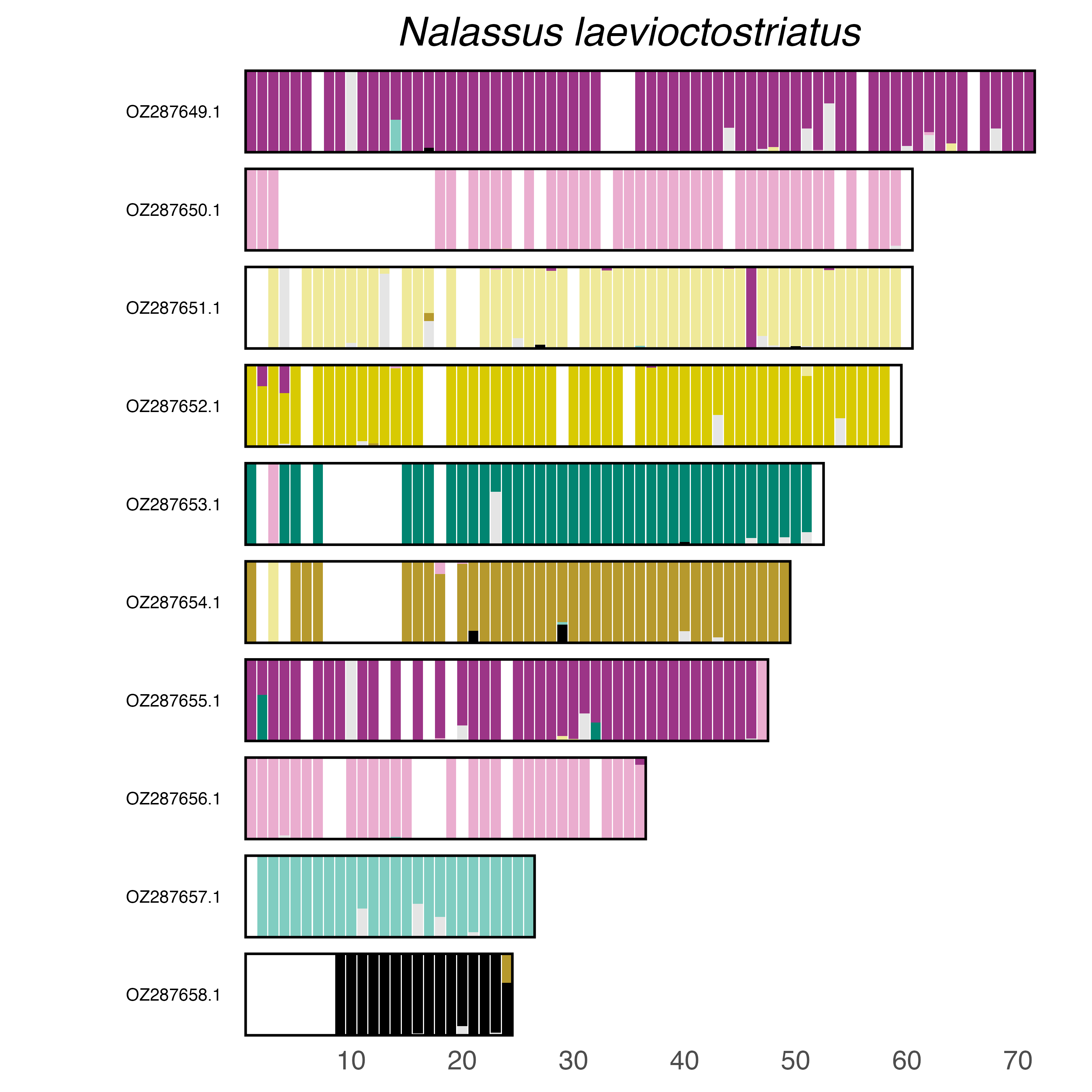

### plot.syngraph_windowbp_1Mb_rel_vertical_chrsize_Nedyus_quadrimaculatus.png

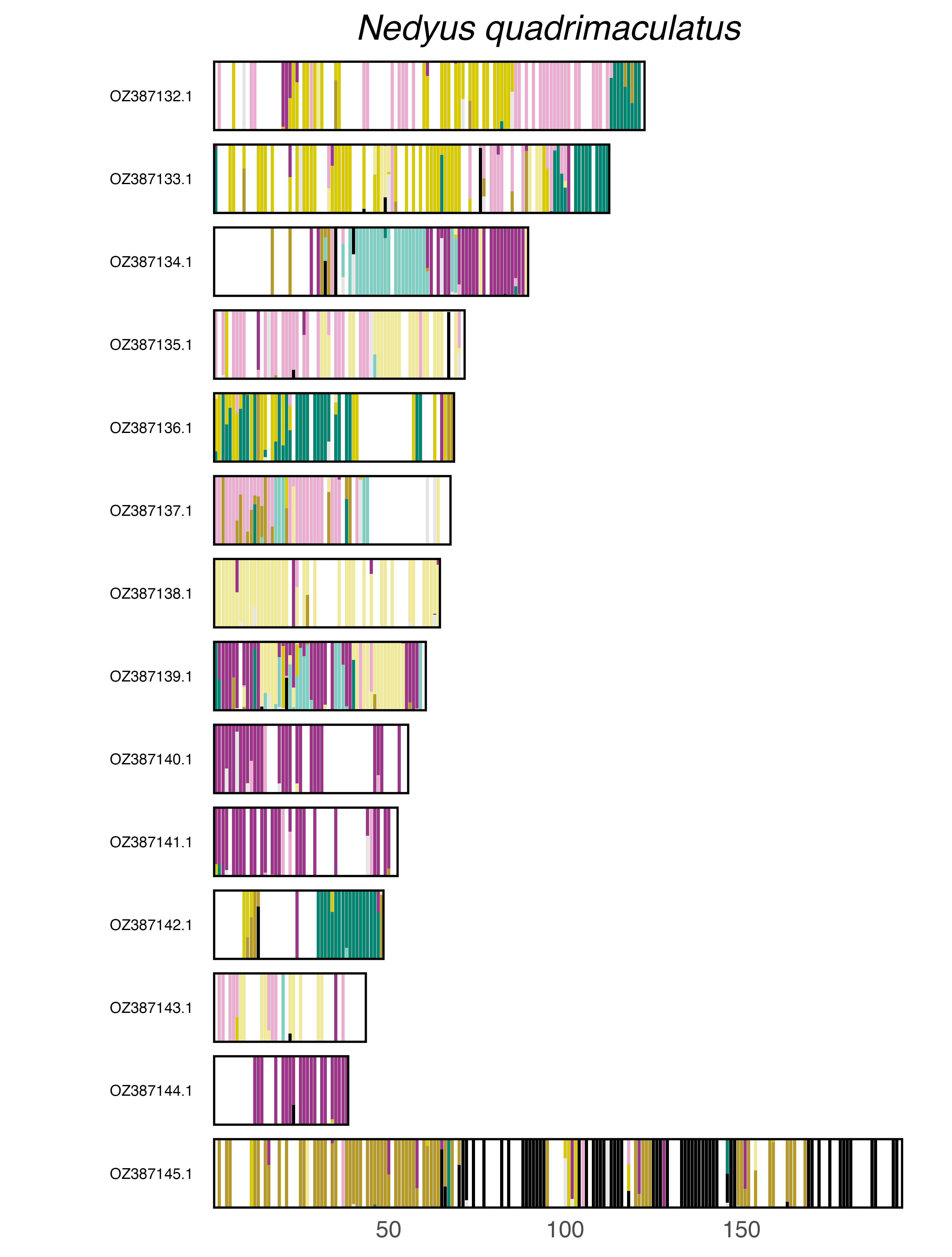

### plot.syngraph_windowbp_1Mb_rel_vertical_chrsize_Neocrepidodera_transversa.png

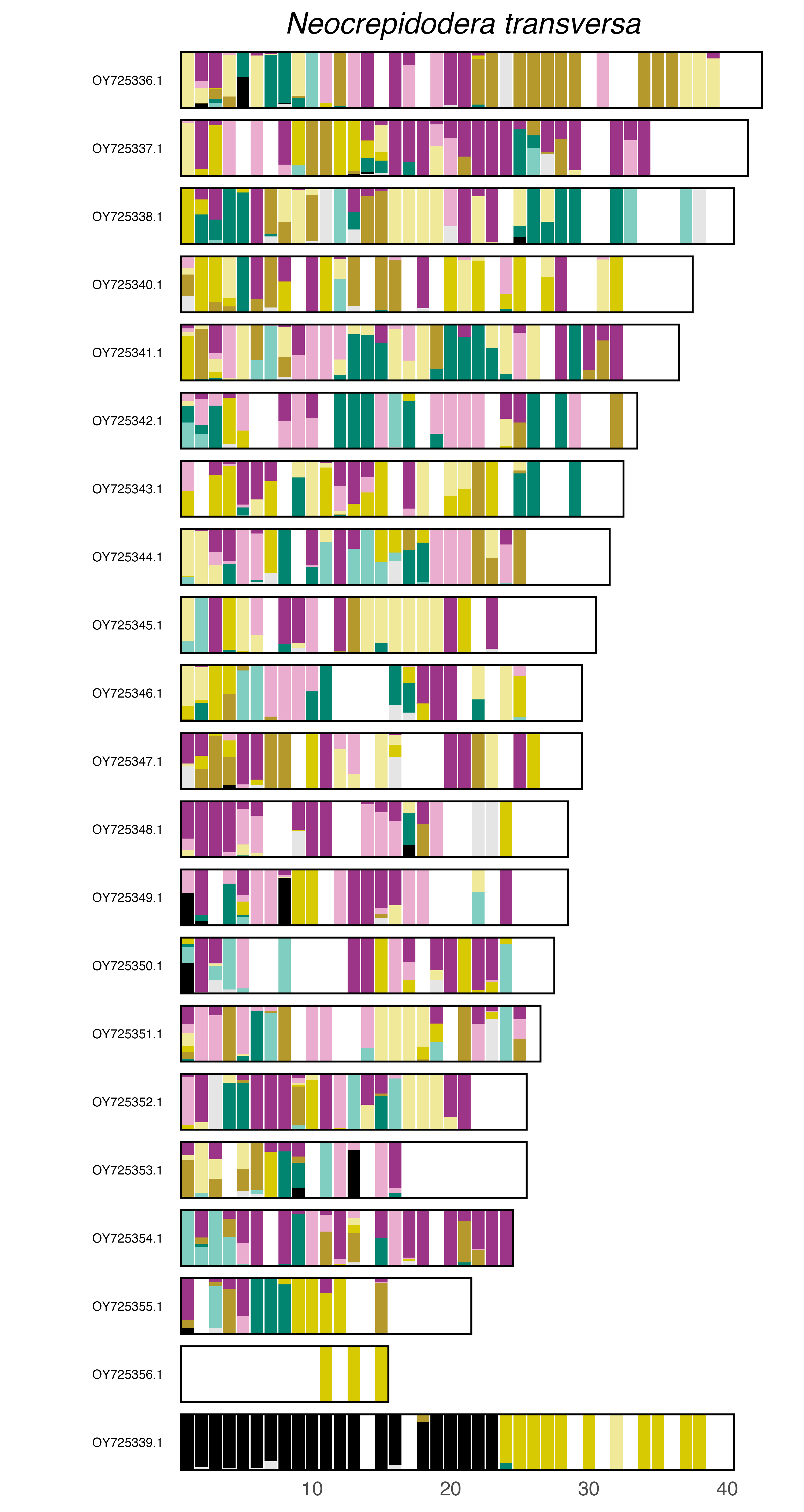

### plot.syngraph_windowbp_1Mb_rel_vertical_chrsize_Neoneuromus_ignobilis.png

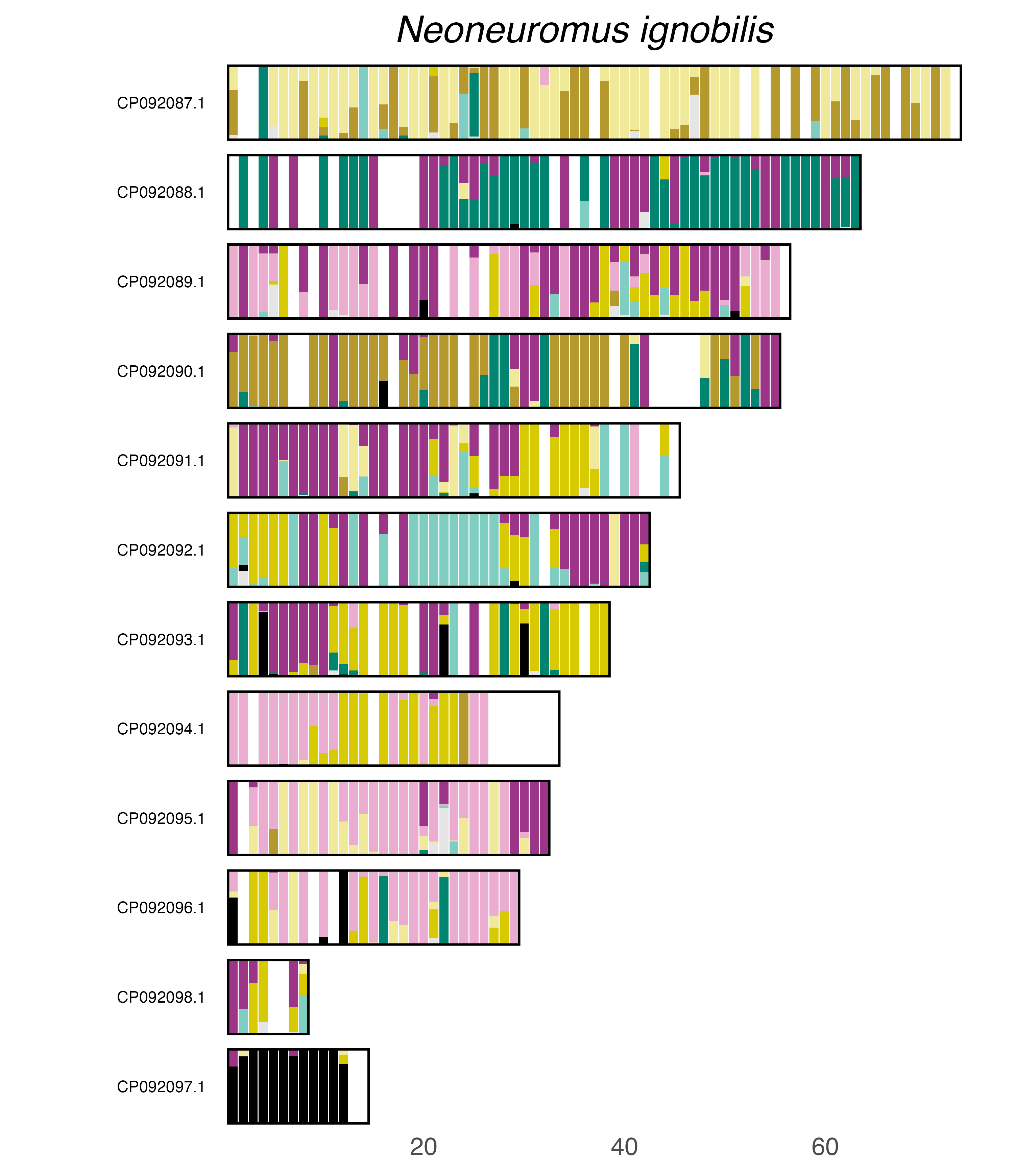

### plot.syngraph_windowbp_1Mb_rel_vertical_chrsize_Onthophagus_taurus.png

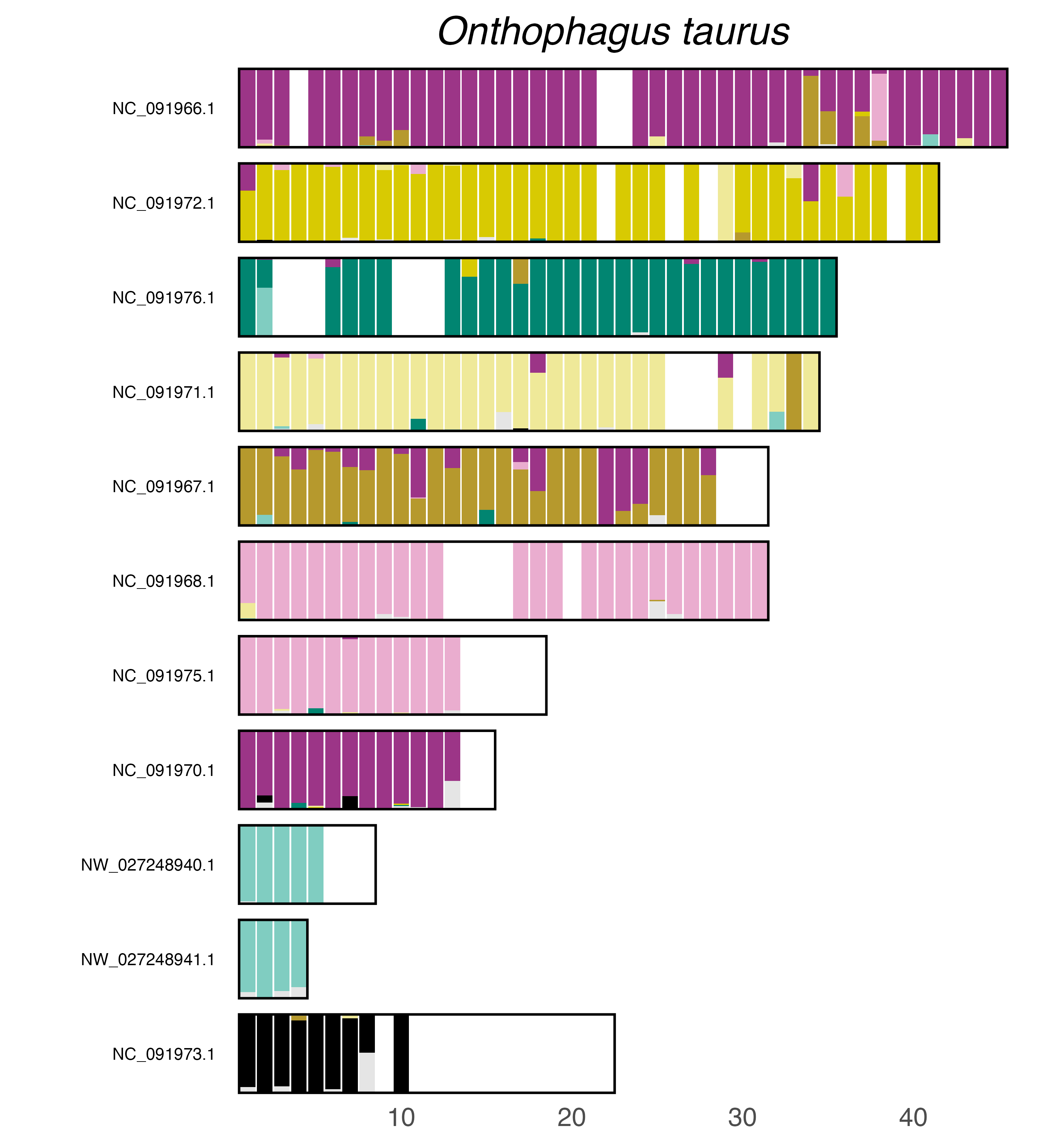

### plot.syngraph_windowbp_1Mb_rel_vertical_chrsize_Othius_punctulatus.png

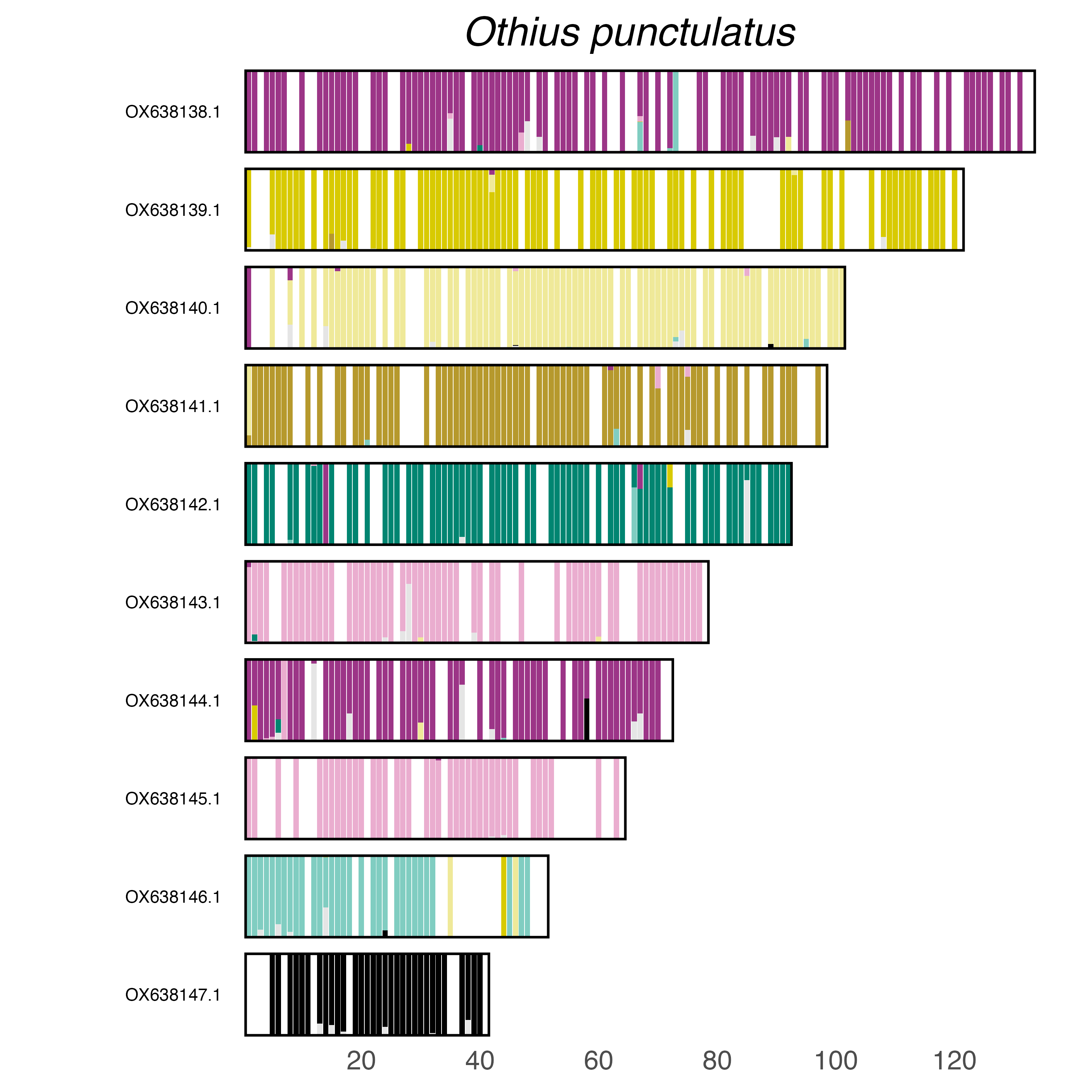

### plot.syngraph_windowbp_1Mb_rel_vertical_chrsize_Phyllotreta_striolata.png

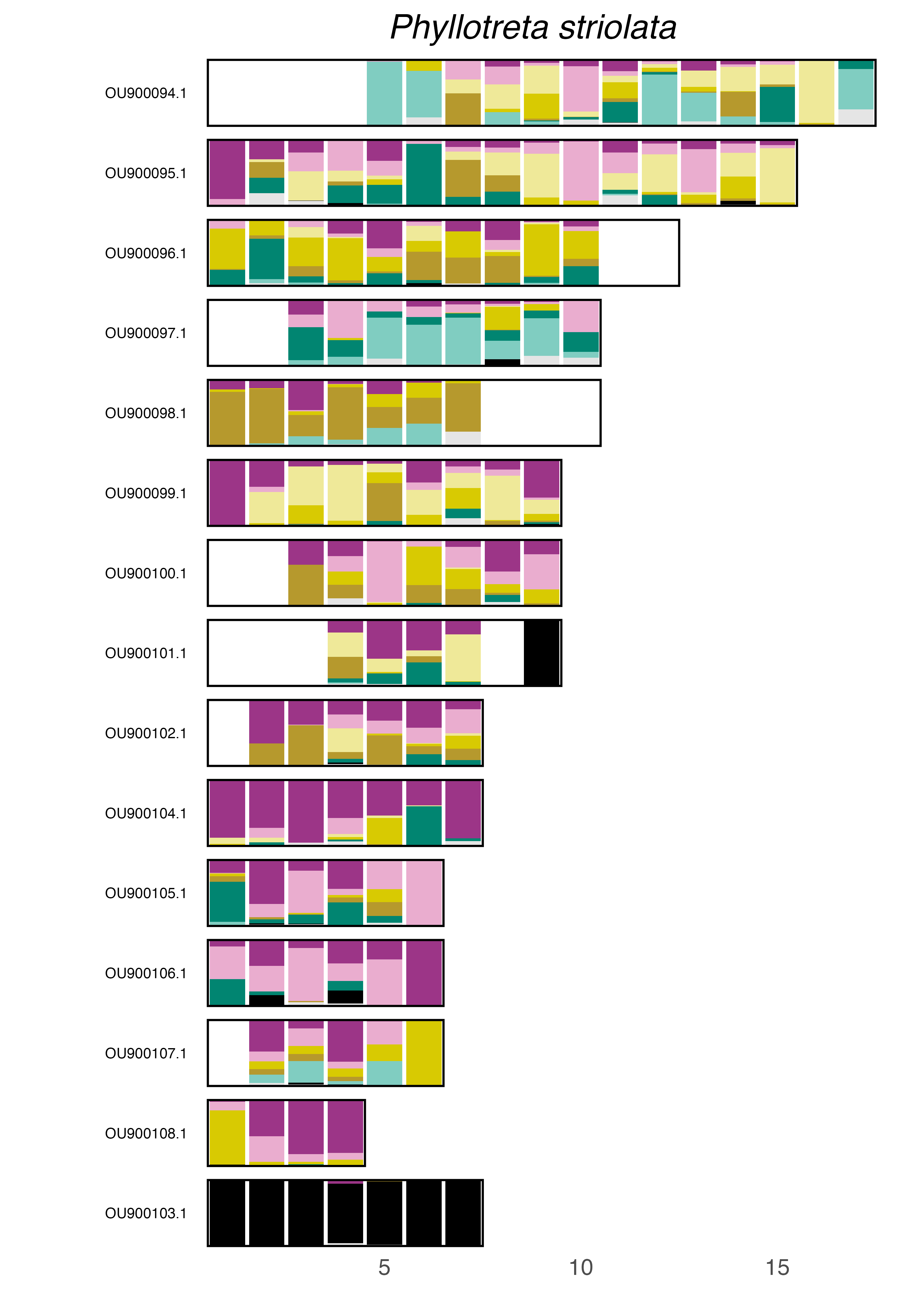

### plot.syngraph_windowbp_1Mb_rel_vertical_chrsize_Podabrus_alpinus.png

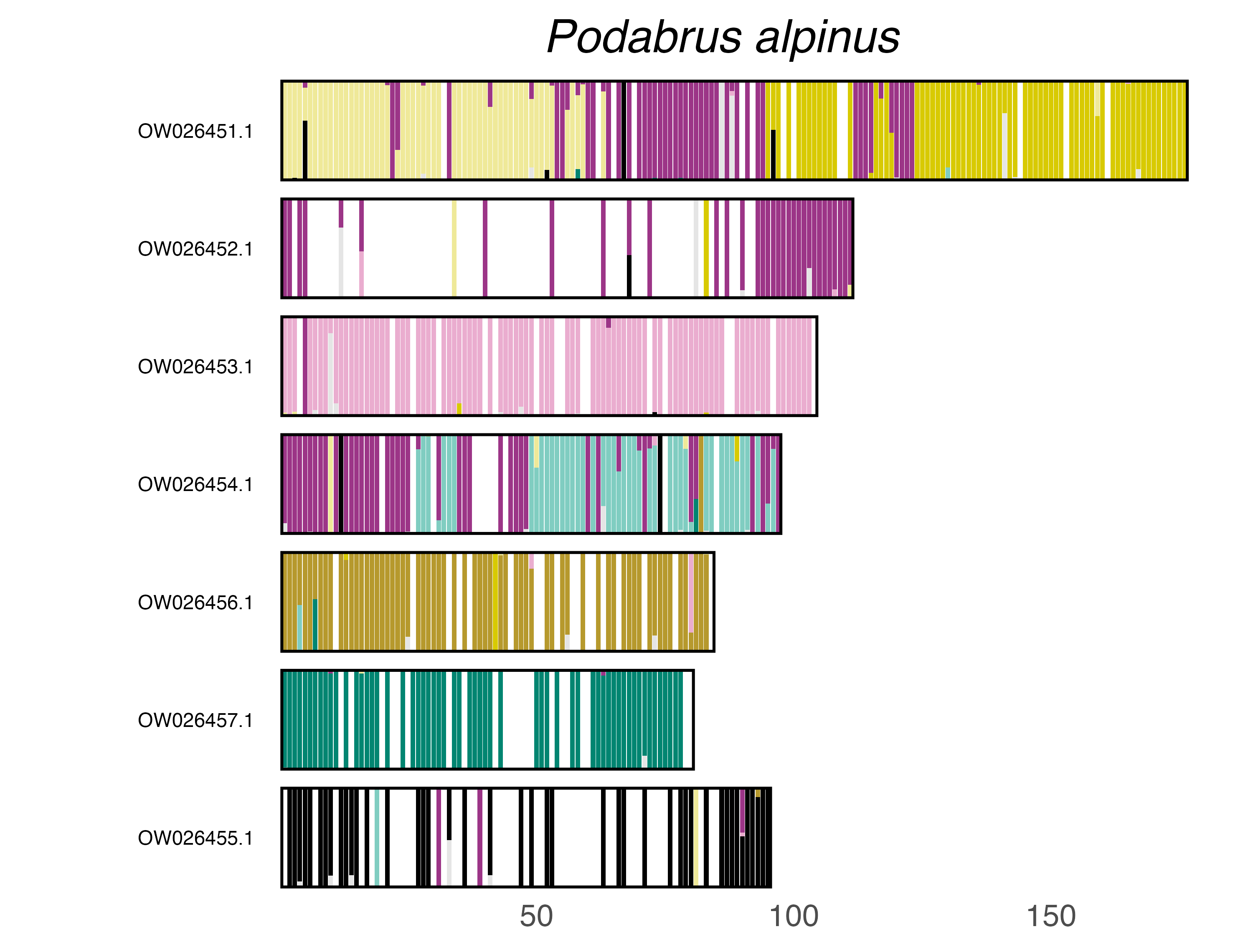

### plot.syngraph_windowbp_1Mb_rel_vertical_chrsize_Polydrusus_impressifrons.png

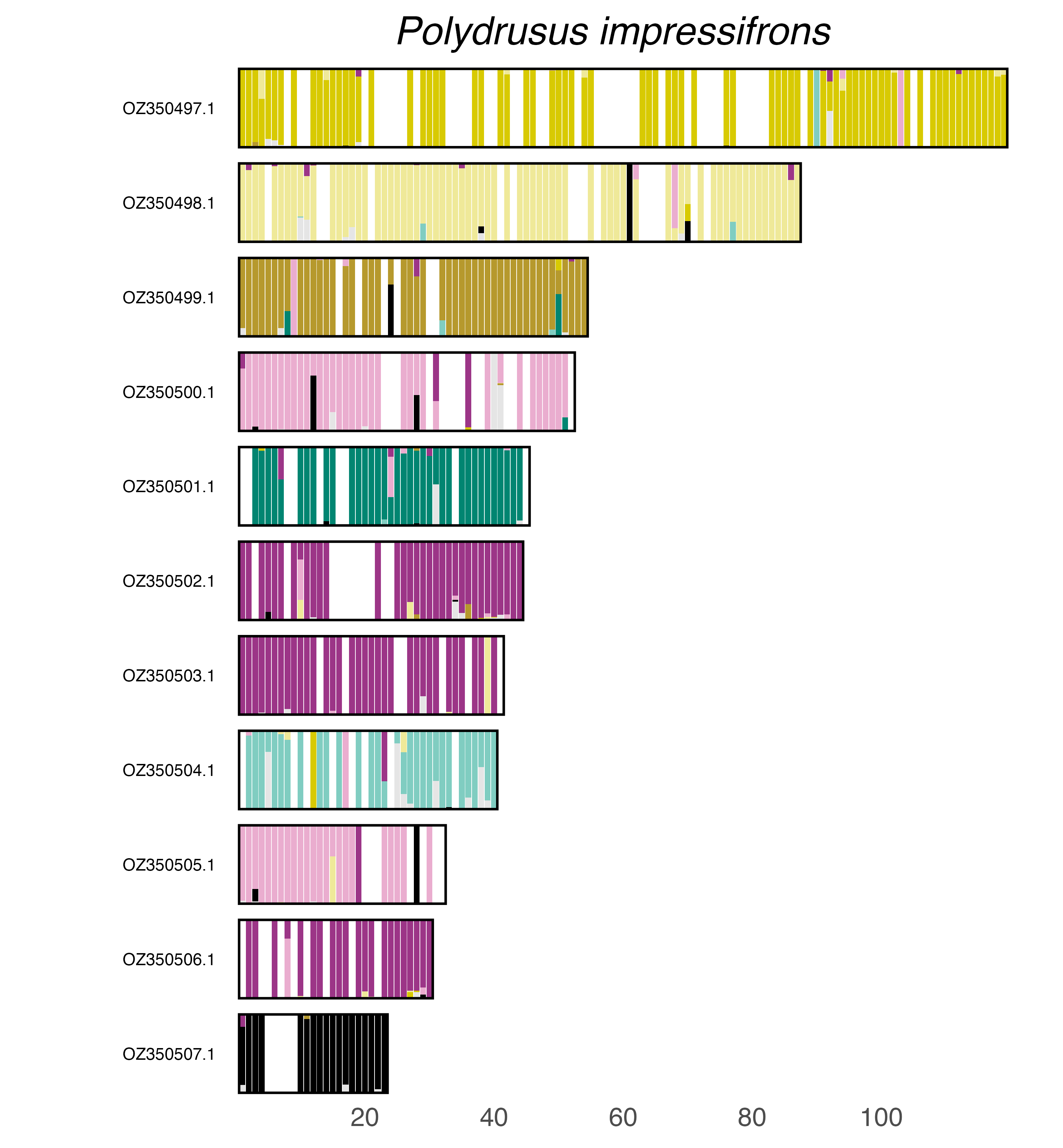

### plot.syngraph_windowbp_1Mb_rel_vertical_chrsize_Prionychus_ater.png

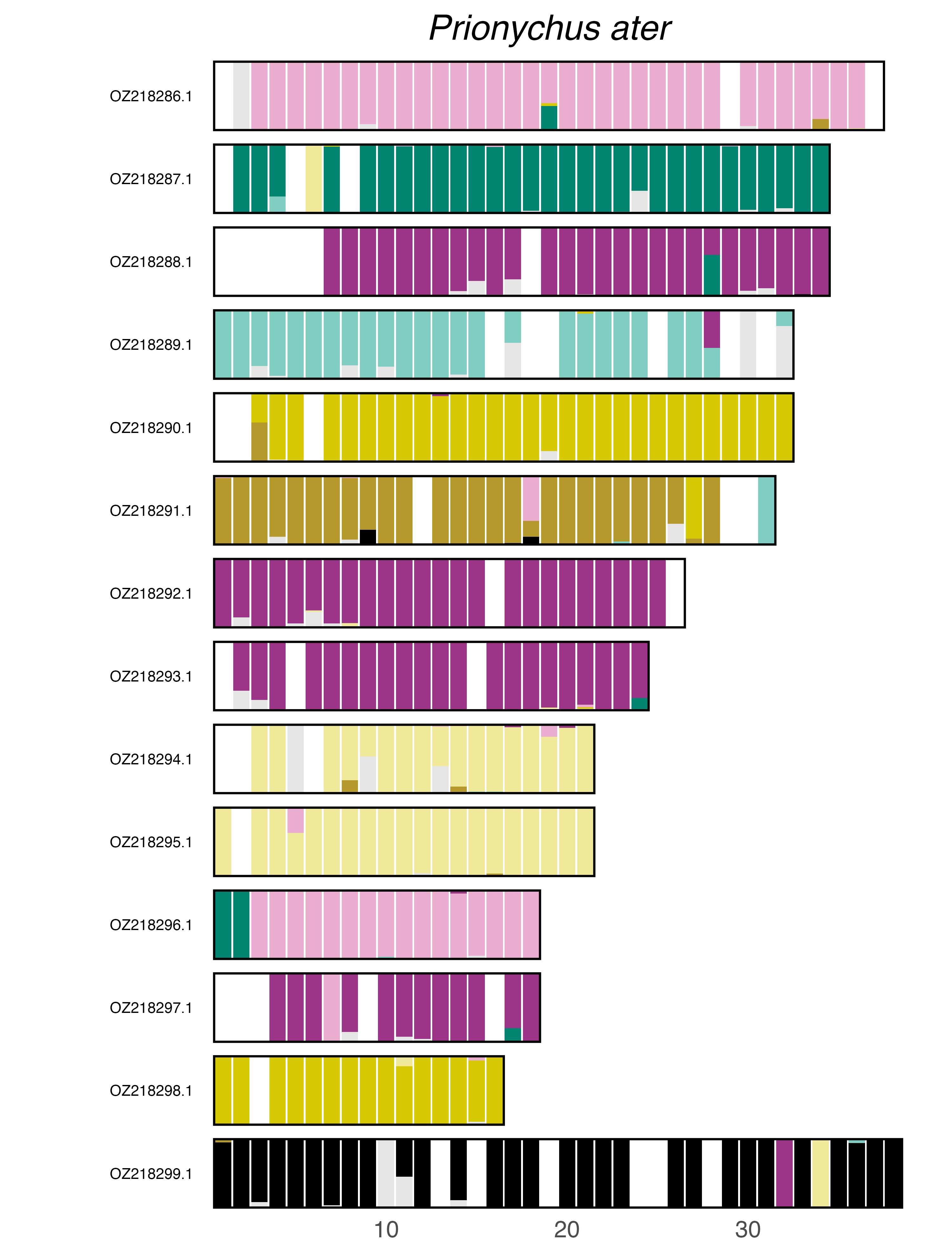

### plot.syngraph_windowbp_1Mb_rel_vertical_chrsize_Propomacrus_cypriacus.png

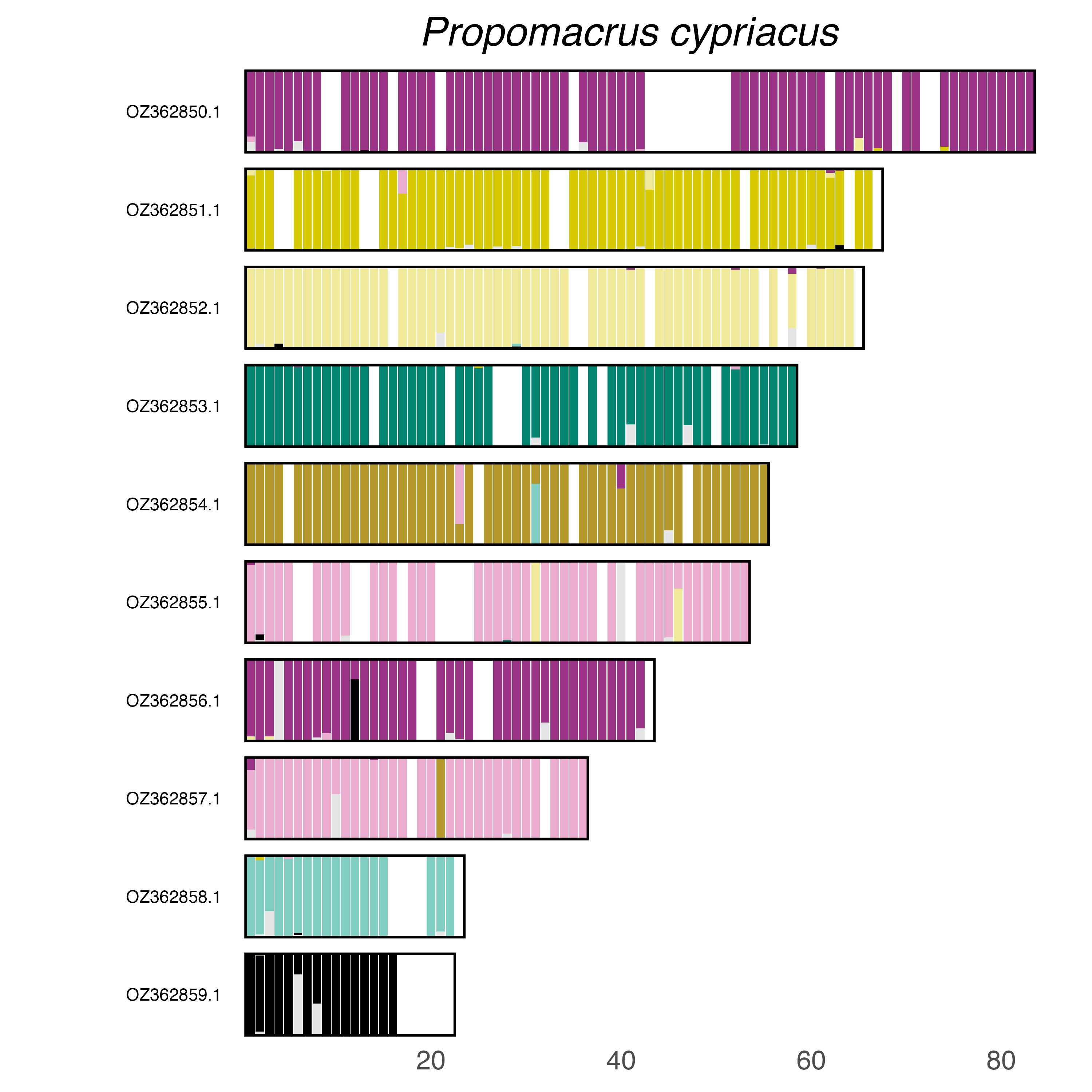

### plot.syngraph_windowbp_1Mb_rel_vertical_chrsize_Pseudocistela_ceramboides.png

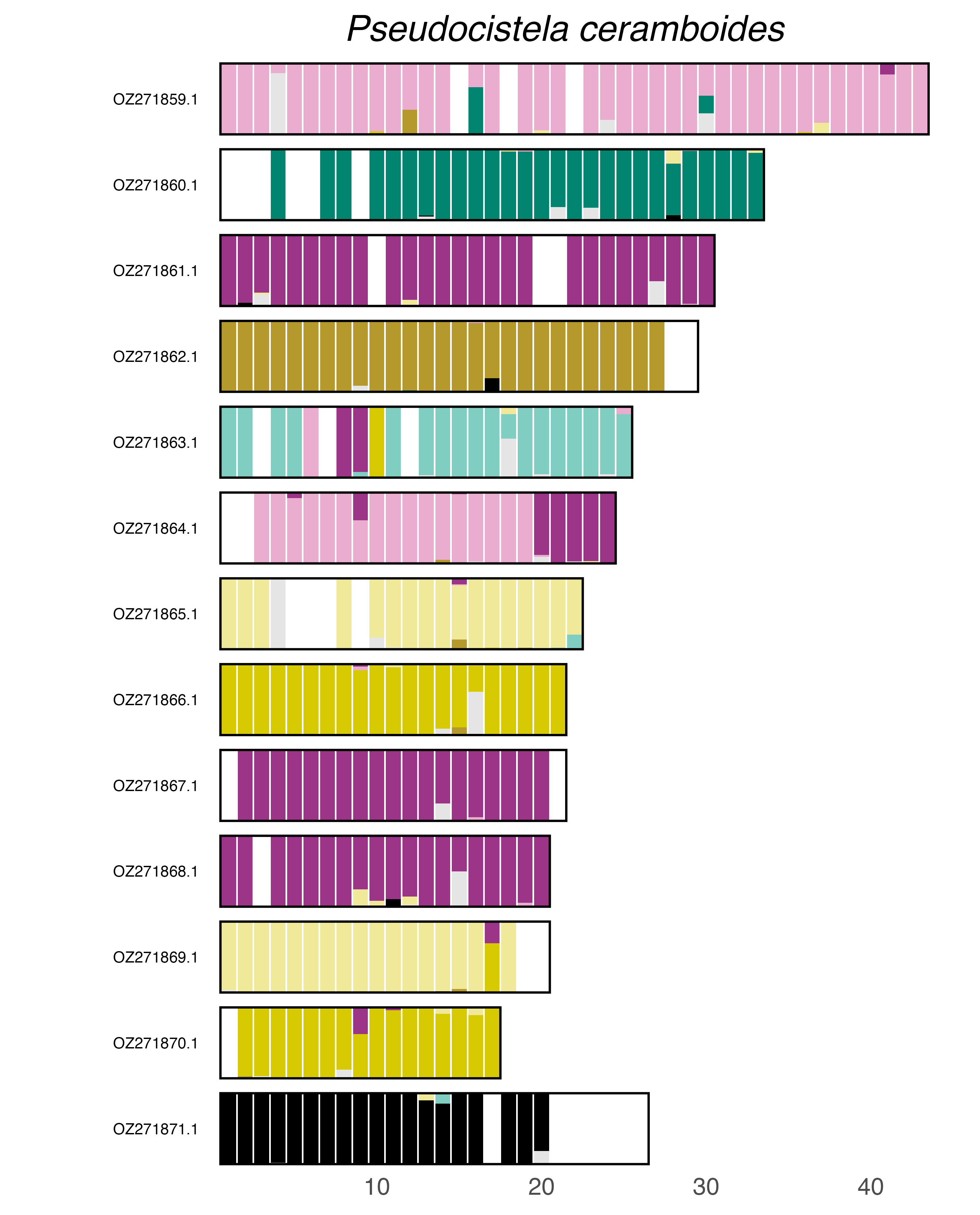

### plot.syngraph_windowbp_1Mb_rel_vertical_chrsize_Pterostichus_niger.png

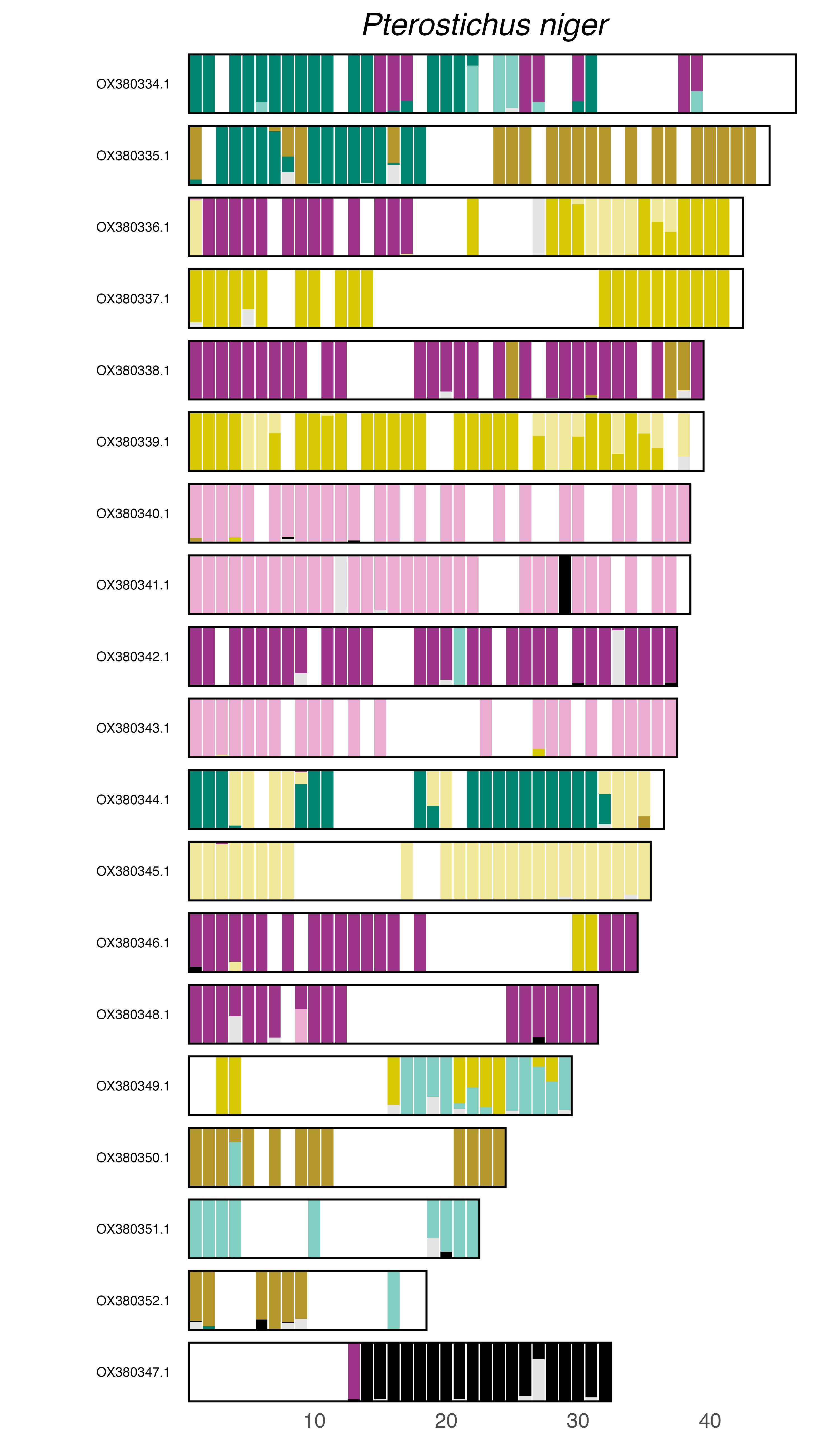

### plot.syngraph_windowbp_1Mb_rel_vertical_chrsize_Rhagium_mordax.png

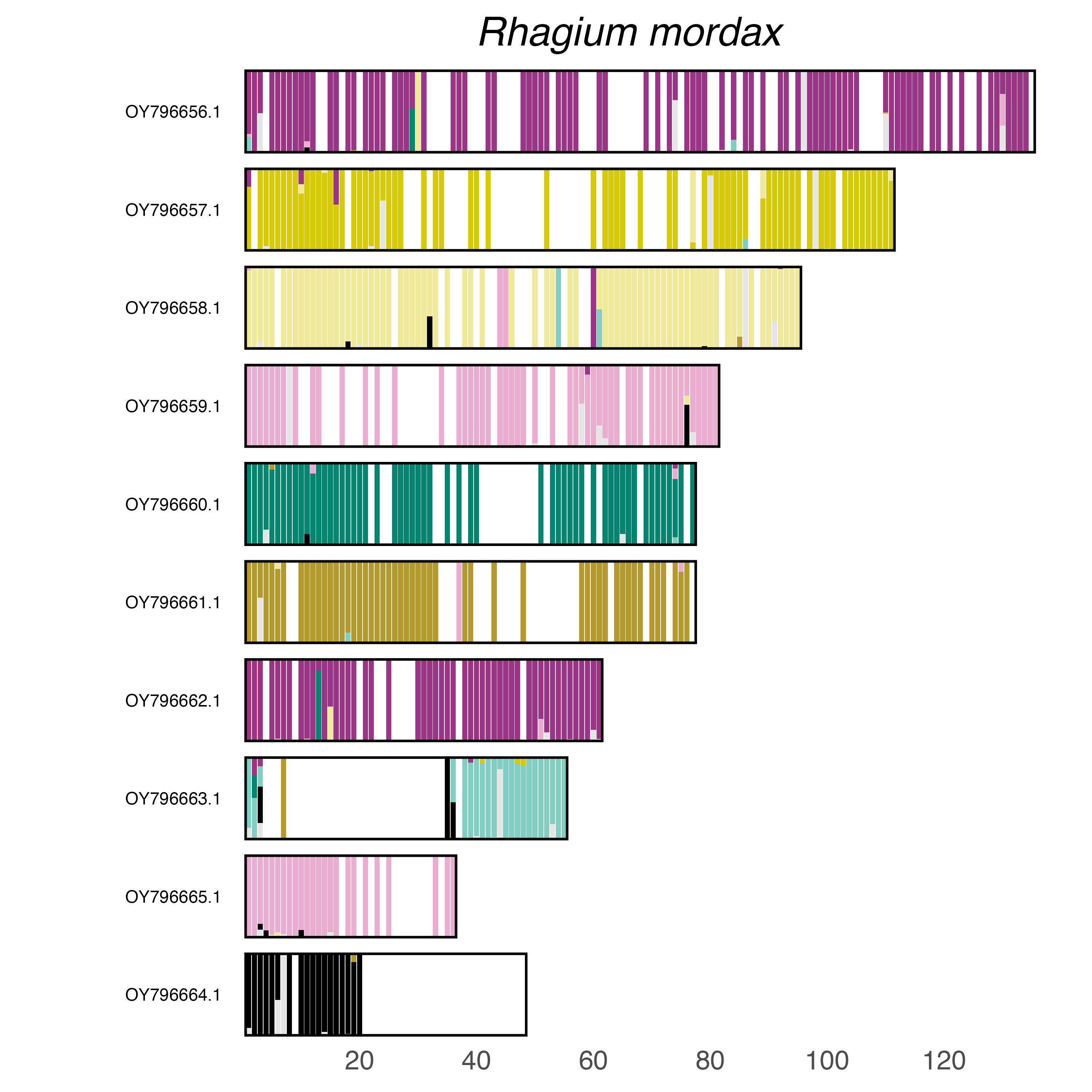

### plot.syngraph_windowbp_1Mb_rel_vertical_chrsize_Salpingus_planirostris.png

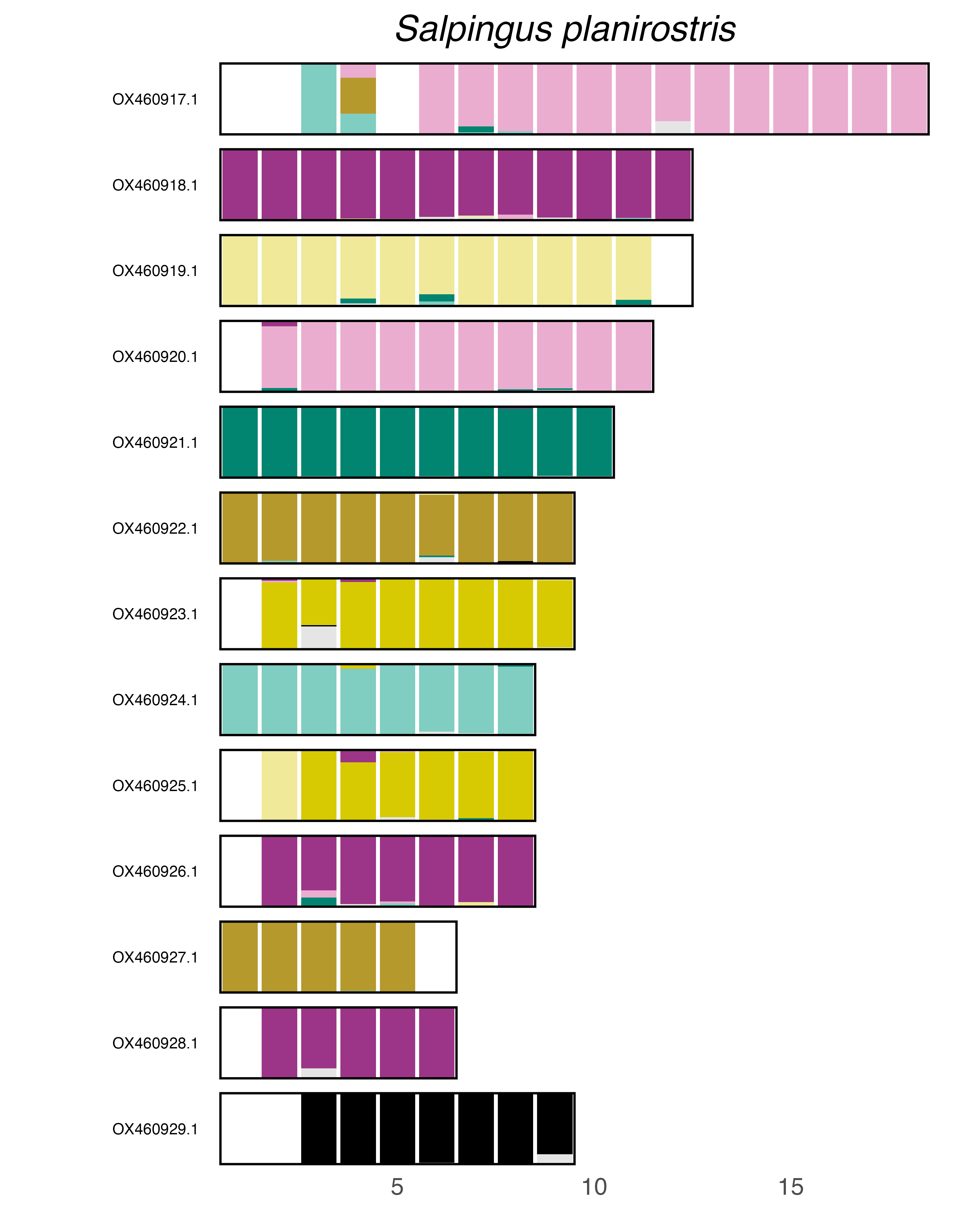

### plot.syngraph_windowbp_1Mb_rel_vertical_chrsize_Scaphidium_quadrimaculatum.png

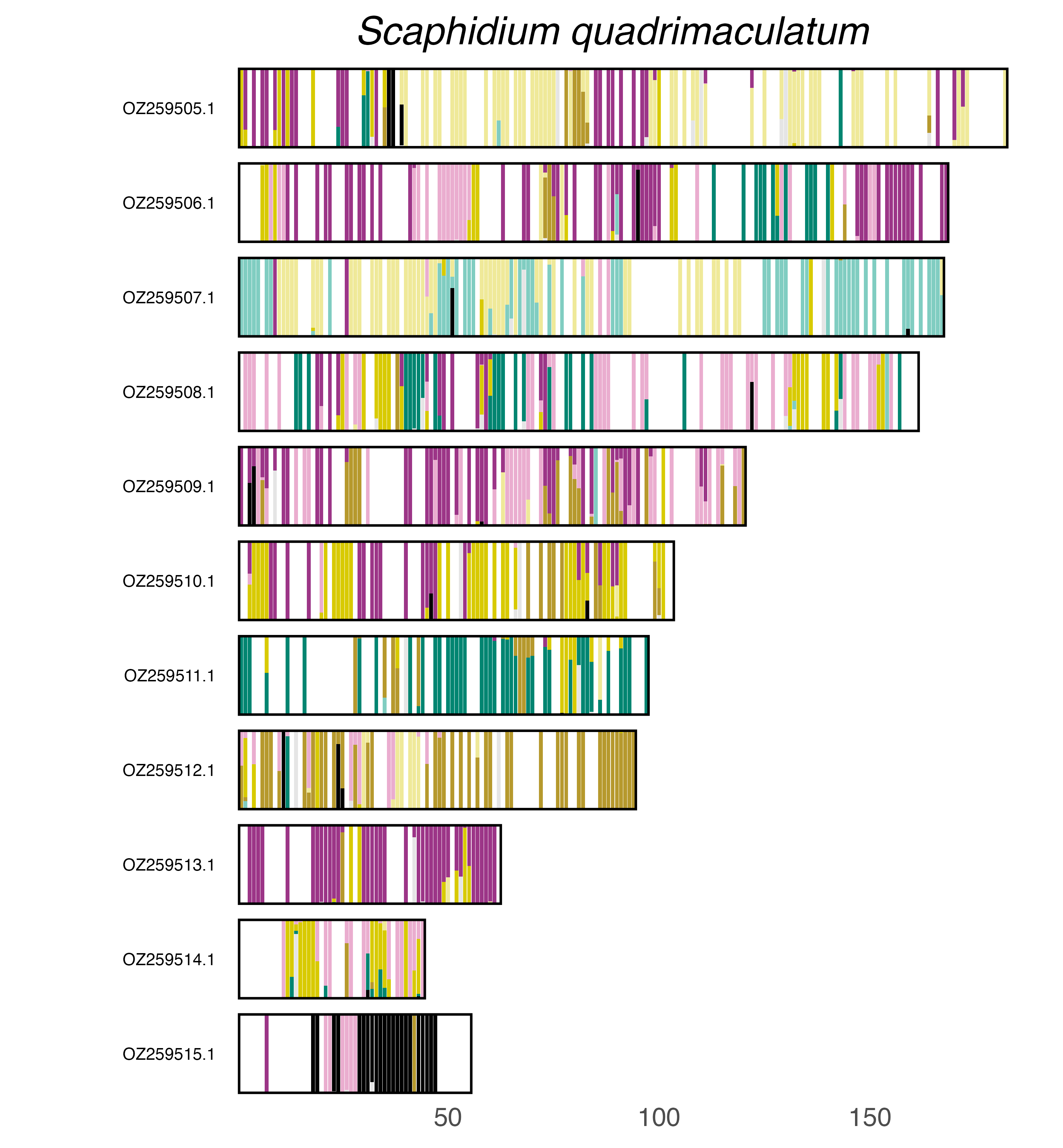

### plot.syngraph_windowbp_1Mb_rel_vertical_chrsize_Schizotus_pectinicornis.png

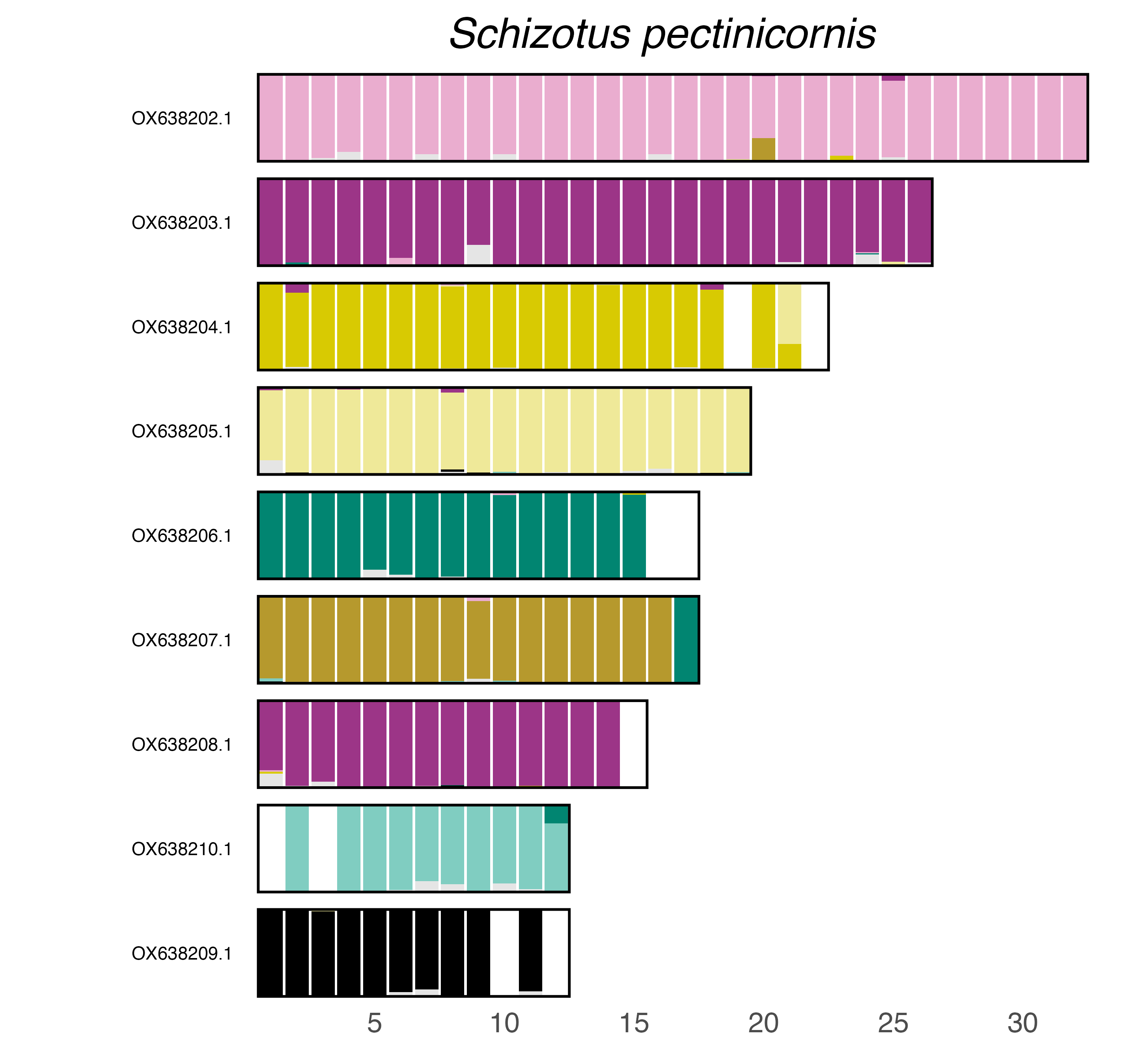

### plot.syngraph_windowbp_1Mb_rel_vertical_chrsize_Serica_brunnea.png

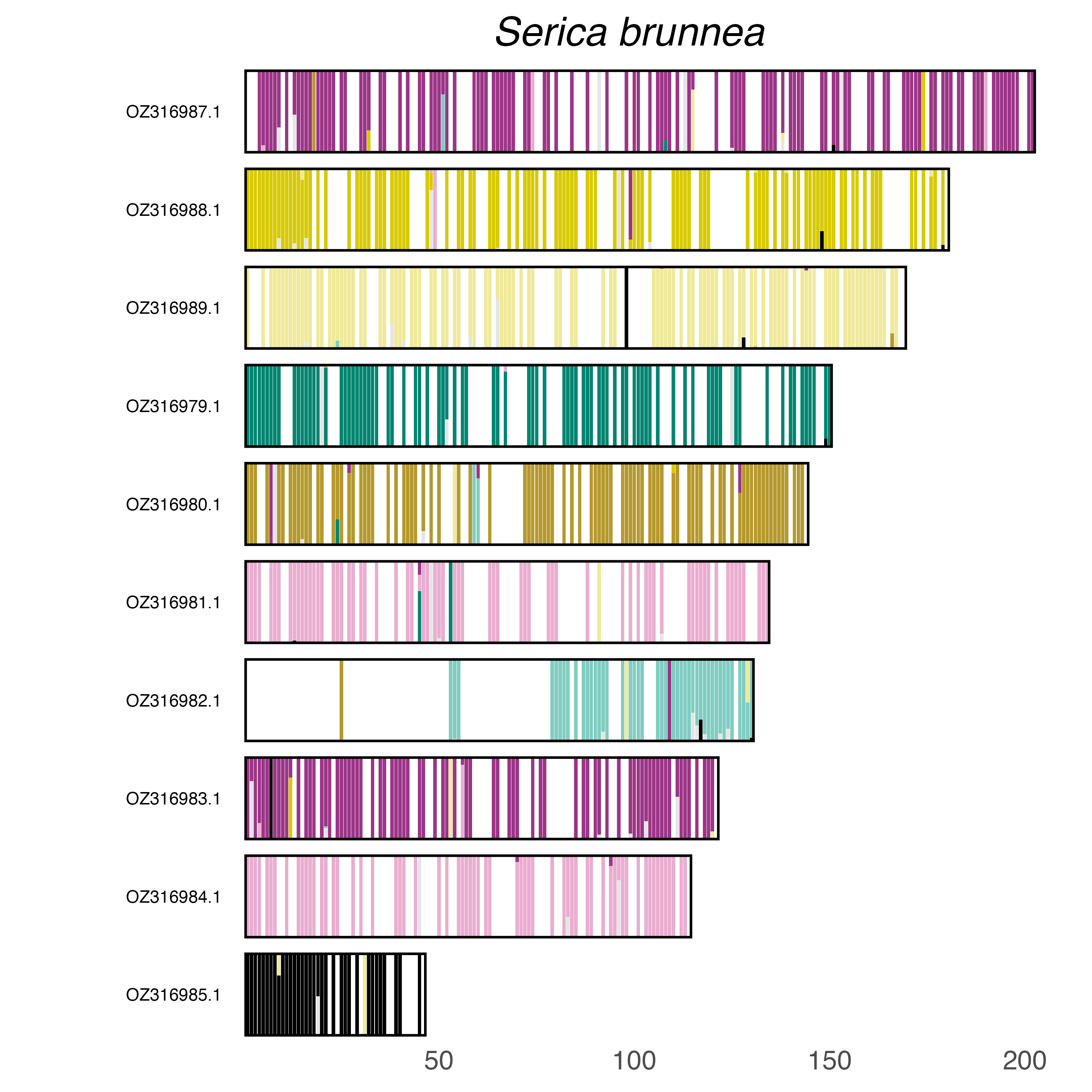

### plot.syngraph_windowbp_1Mb_rel_vertical_chrsize_Sialis_lutaria.png

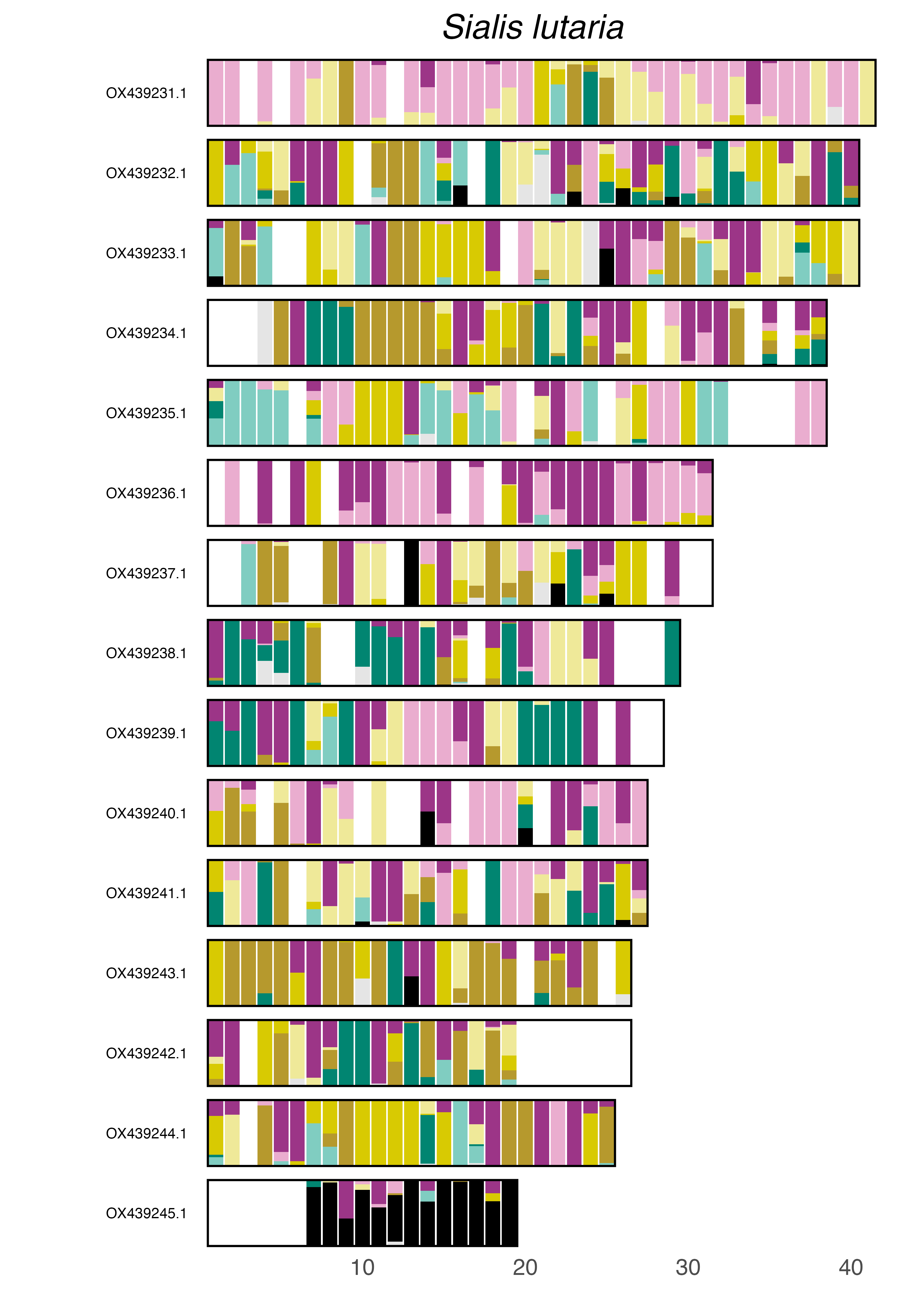

### plot.syngraph_windowbp_1Mb_rel_vertical_chrsize_Sitophilus_zeamais.png

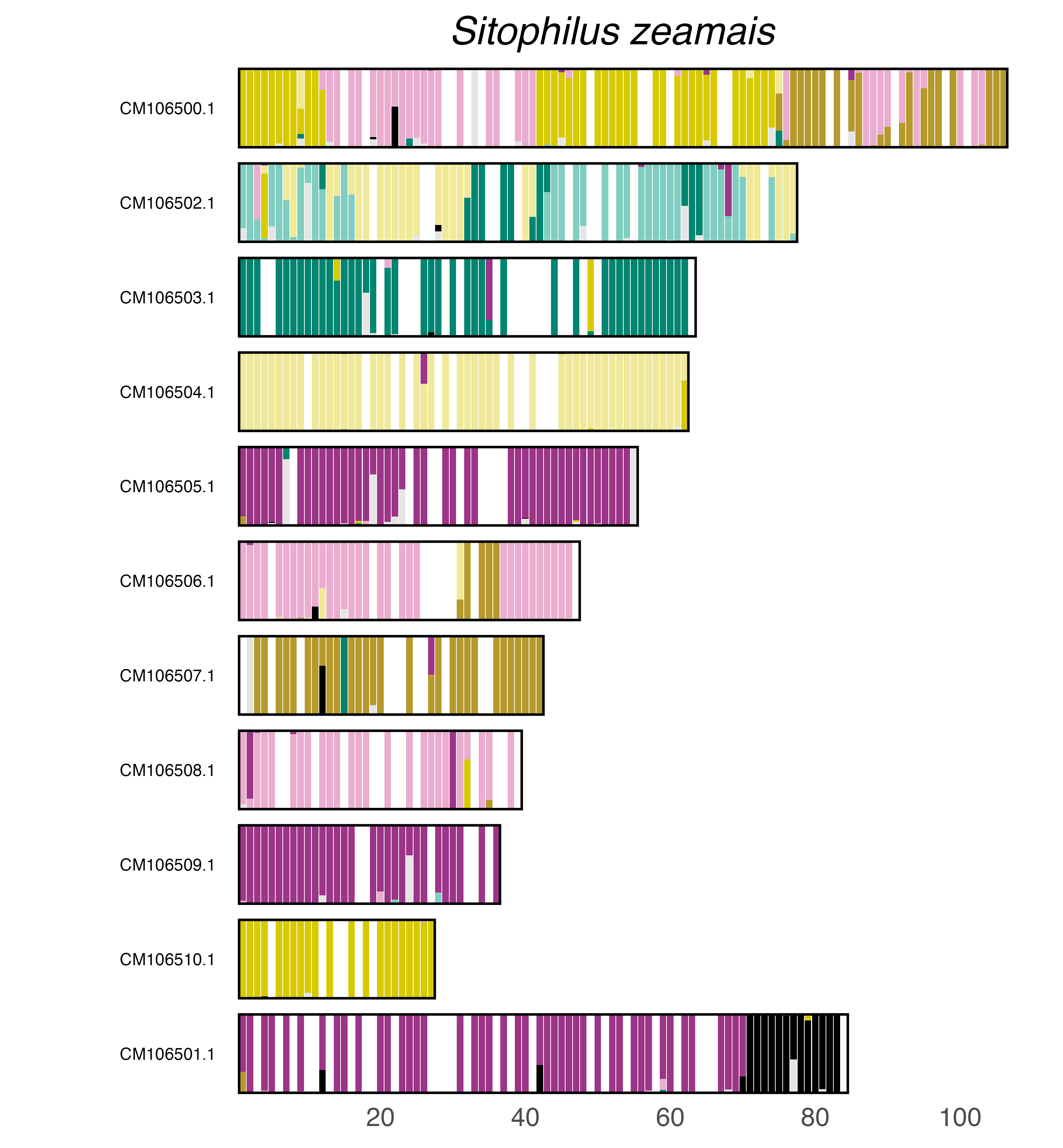

### plot.syngraph_windowbp_1Mb_rel_vertical_chrsize_Sphaeroderma_testaceum.png

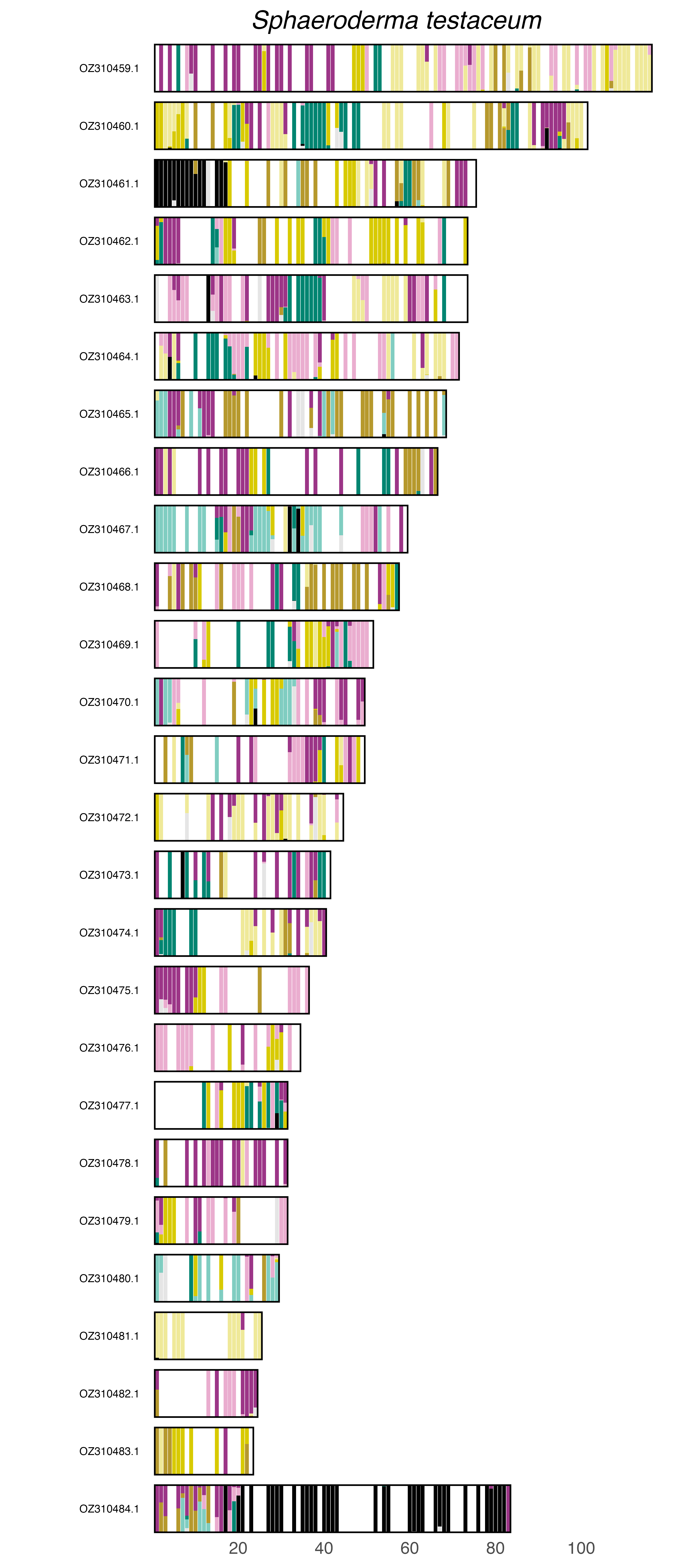

### plot.syngraph_windowbp_1Mb_rel_vertical_chrsize_Staphylinus_erythropterus.png

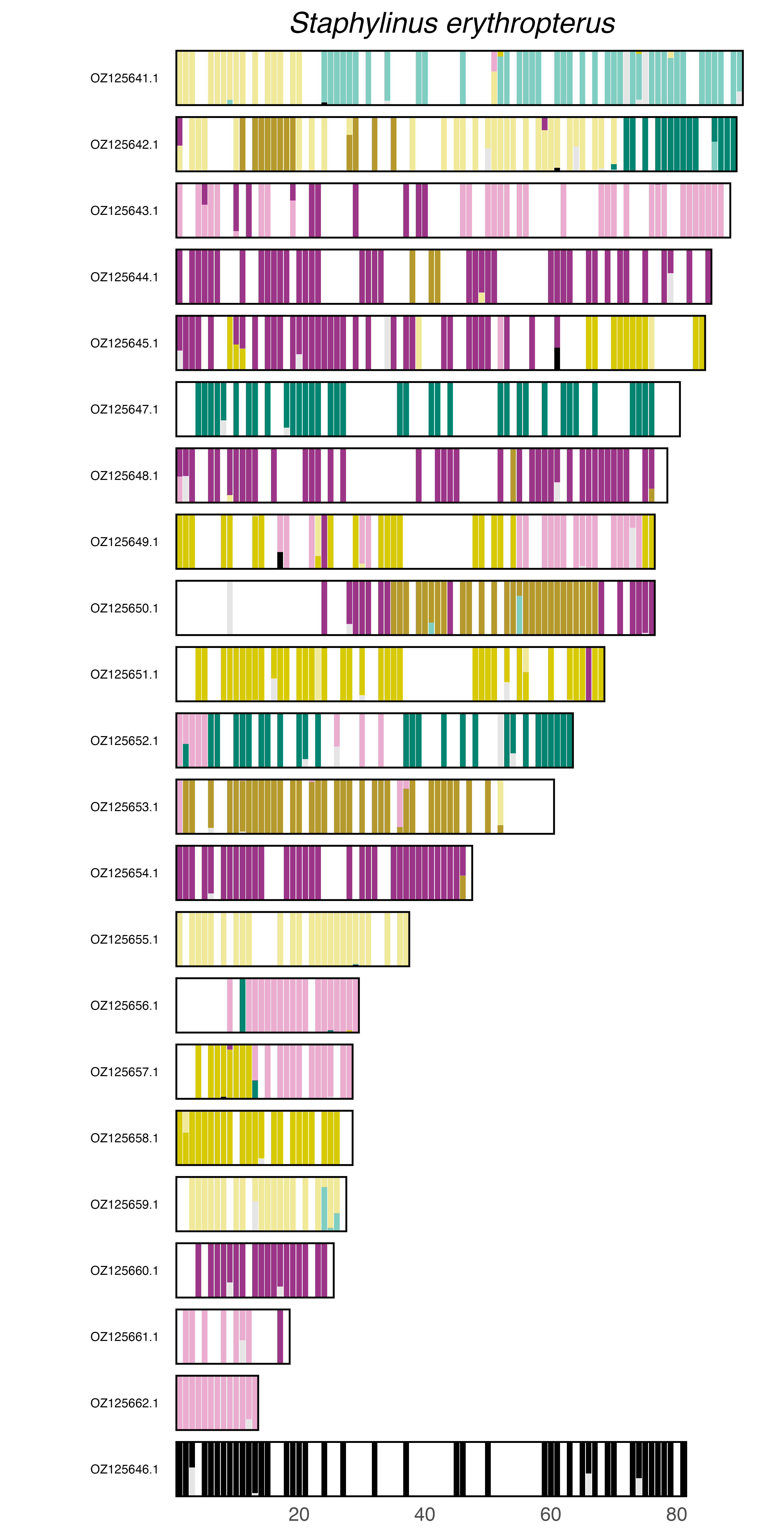

### plot.syngraph_windowbp_1Mb_rel_vertical_chrsize_Stylops_aterrimus.png

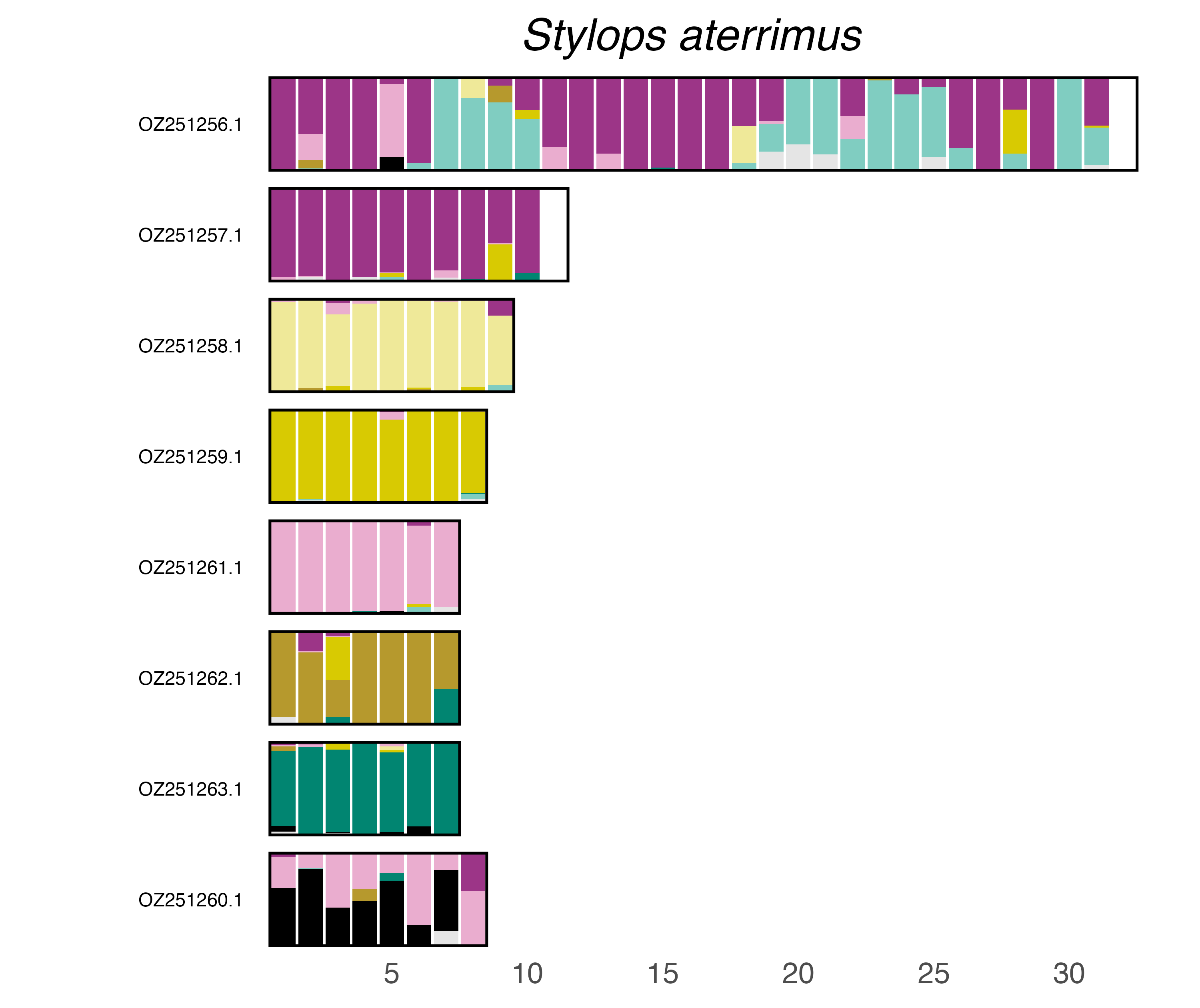

### plot.syngraph_windowbp_1Mb_rel_vertical_chrsize_Subcoccinella_vigintiquattuorpunctata.png

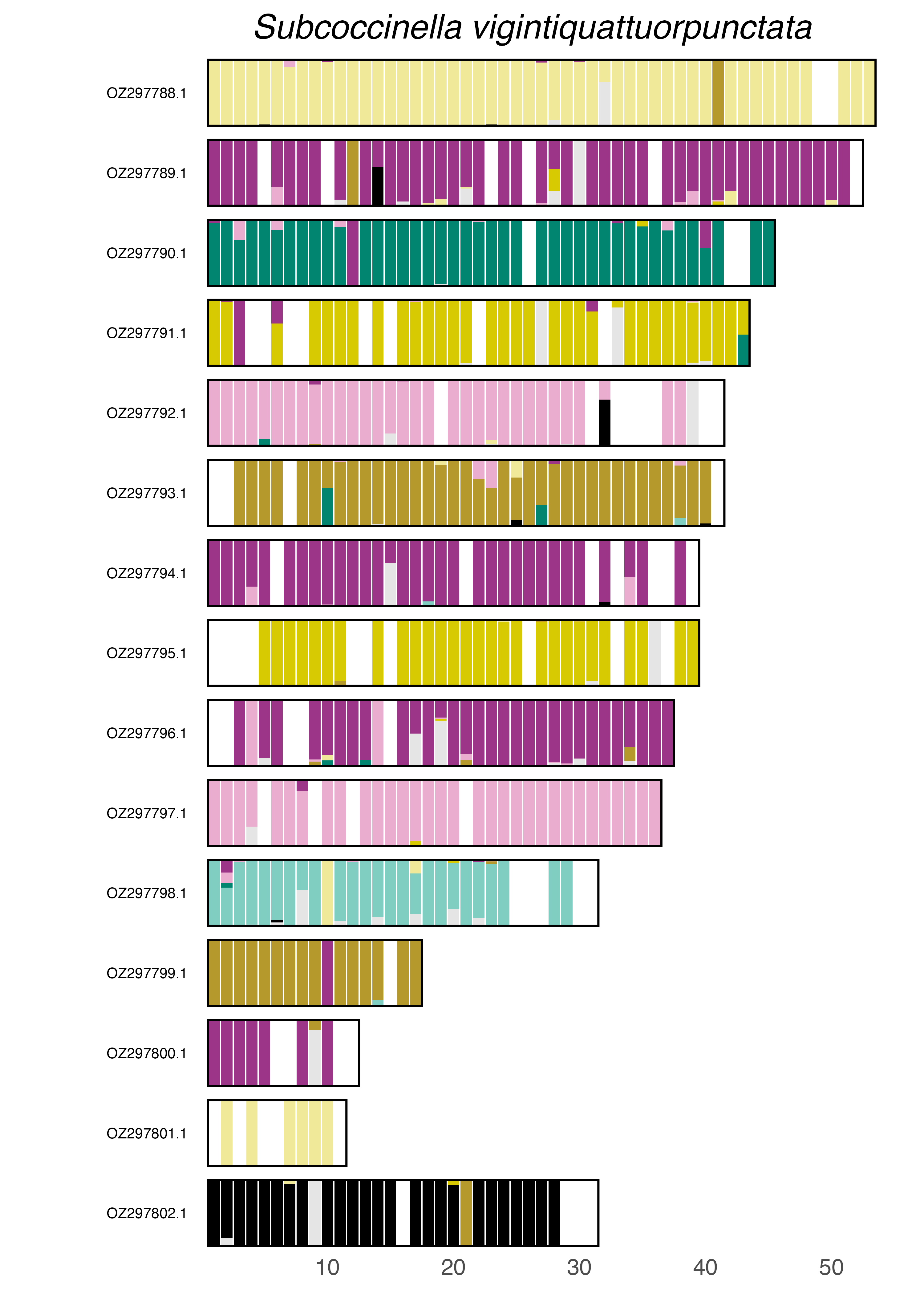

### plot.syngraph_windowbp_1Mb_rel_vertical_chrsize_Tarphius_canariensis.png

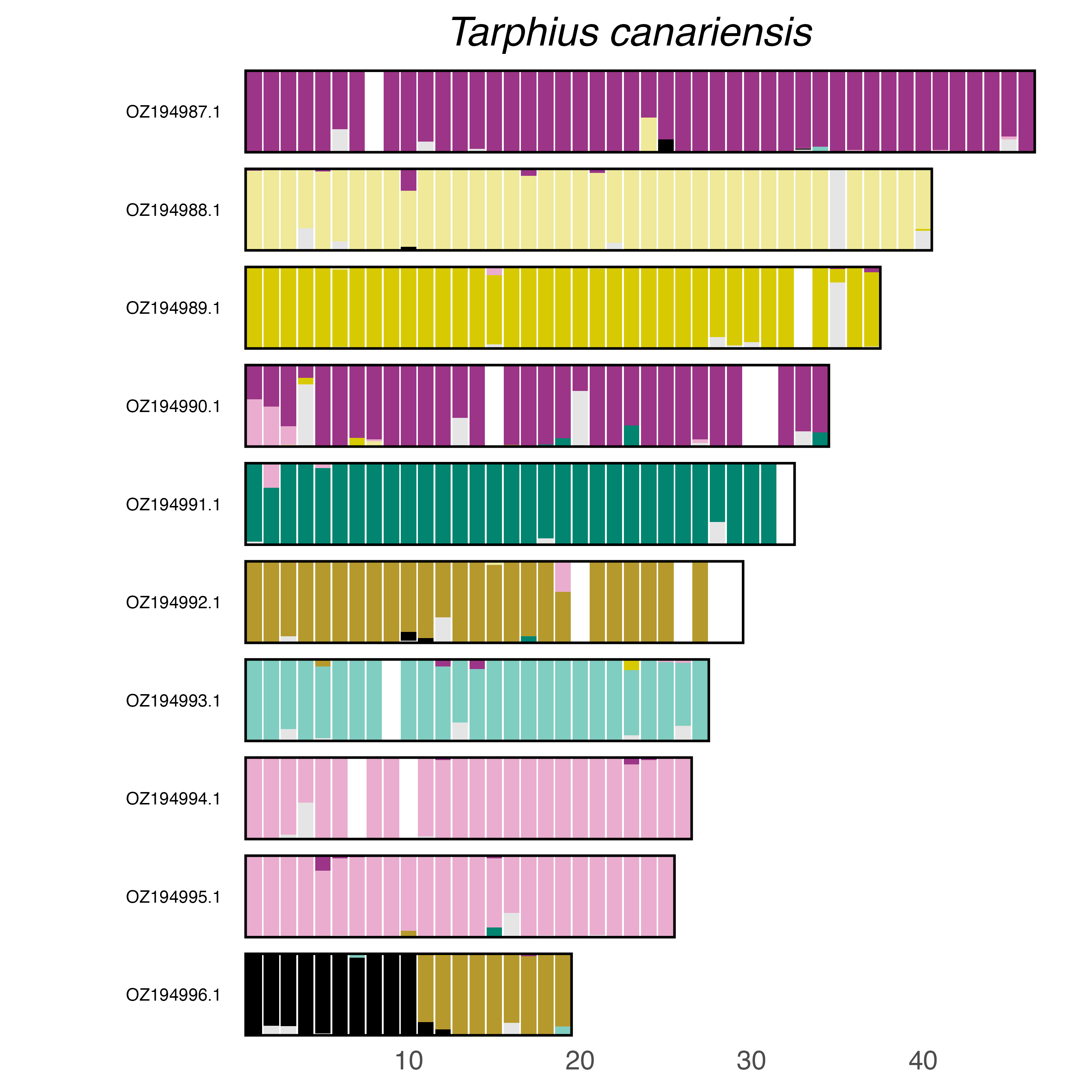

### plot.syngraph_windowbp_1Mb_rel_vertical_chrsize_Tetraopes_melanurus.png

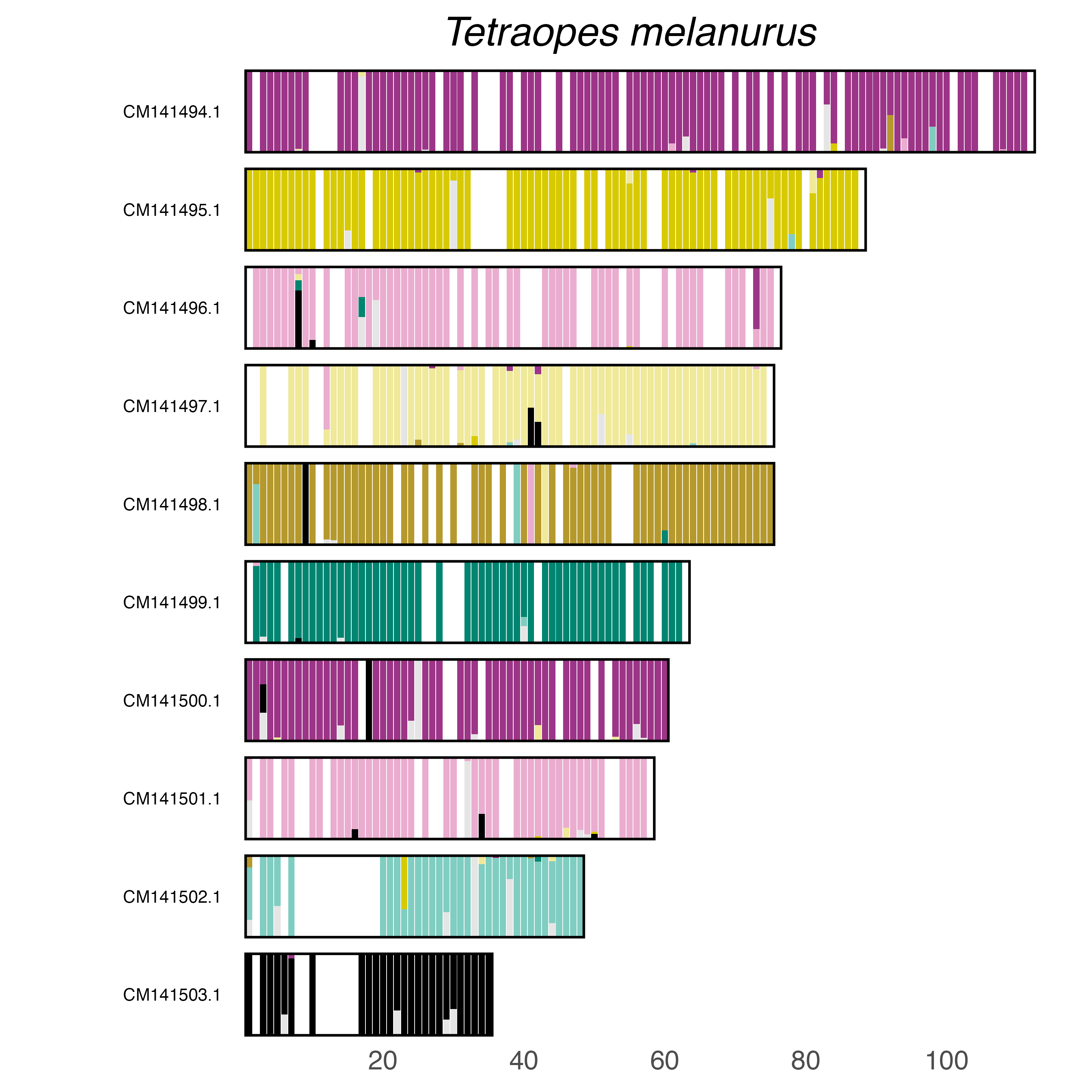

### plot.syngraph_windowbp_1Mb_rel_vertical_chrsize_Tribolium_castaneum.png

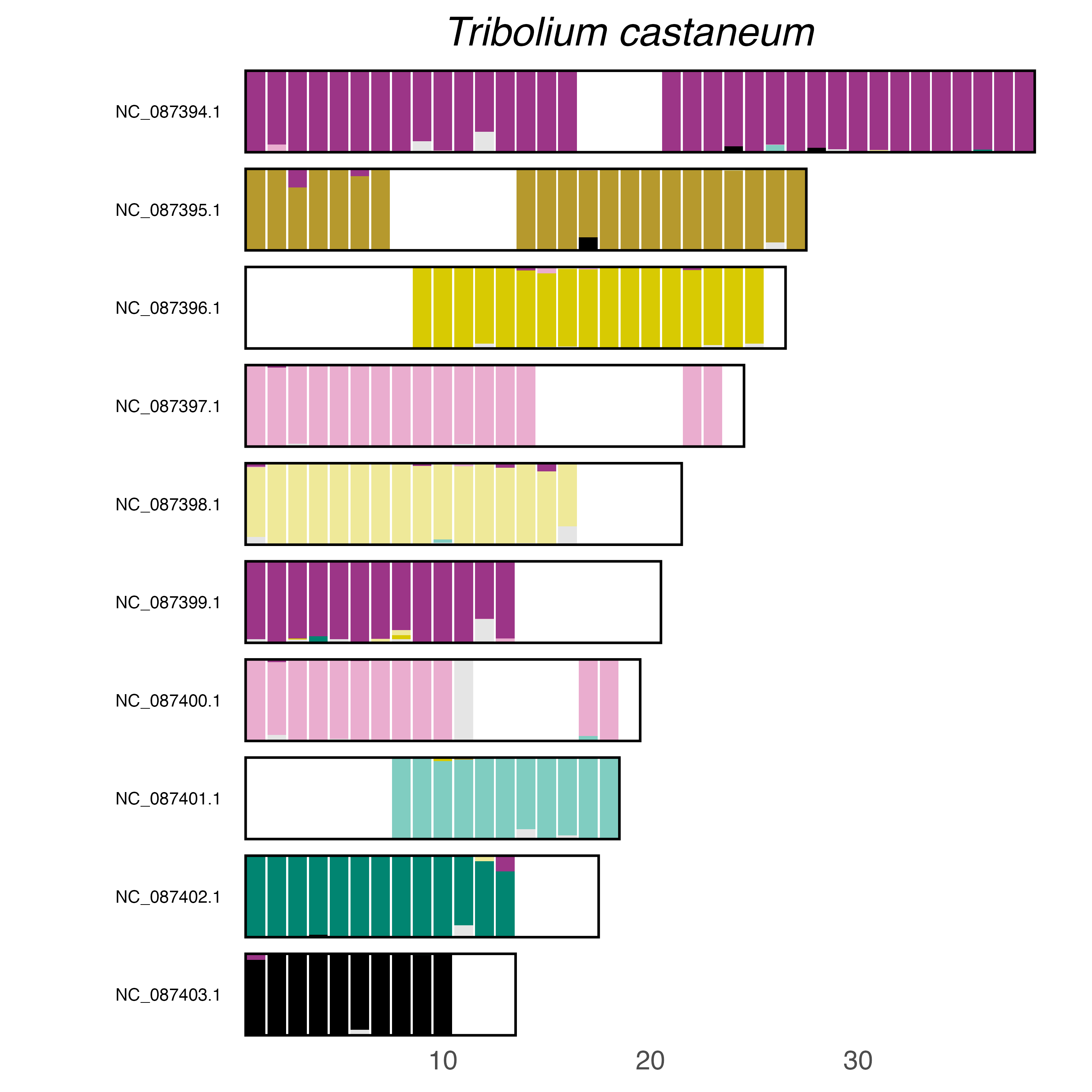

### plot.syngraph_windowbp_1Mb_rel_vertical_chrsize_Ulomoides_dermestoides.png

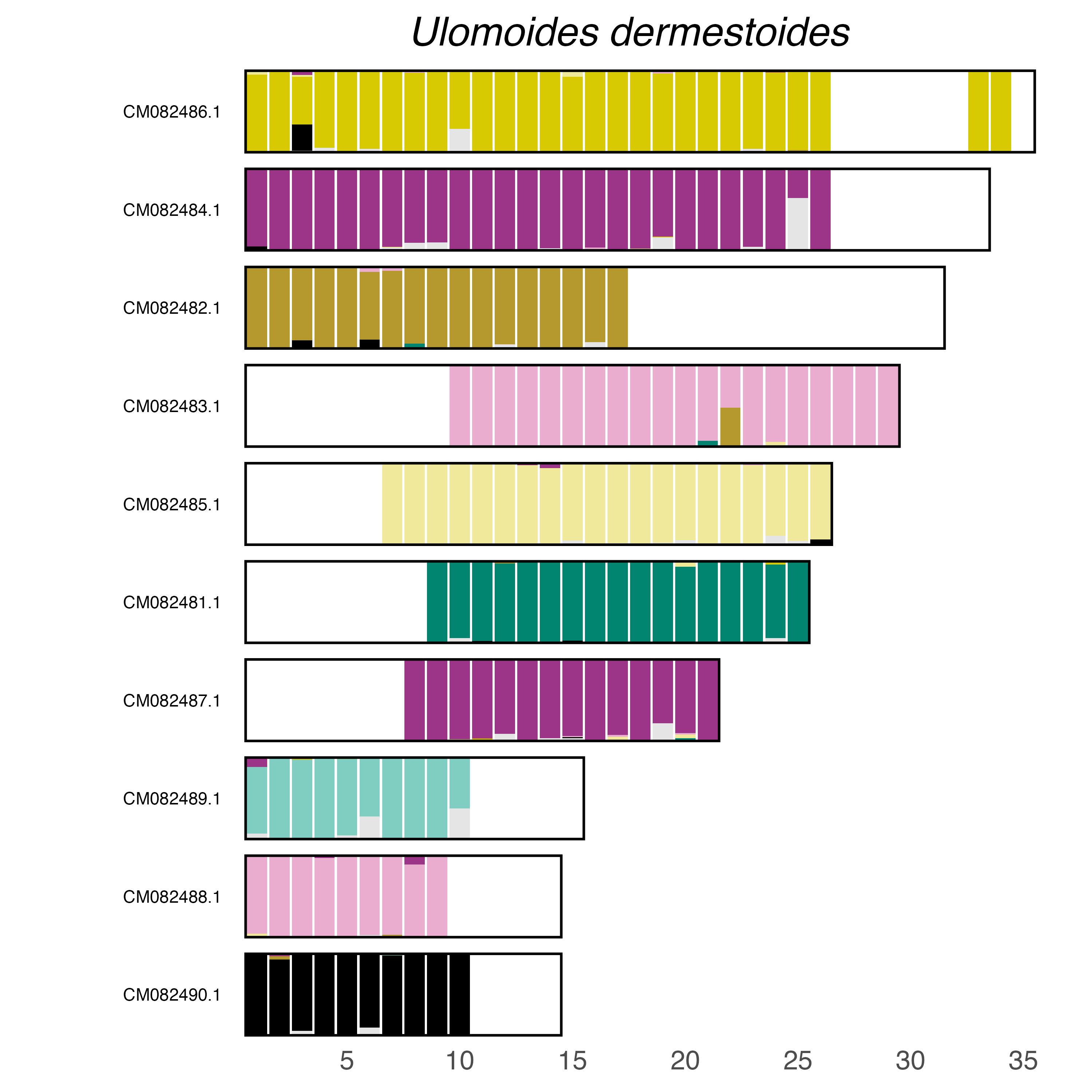

### plot.syngraph_windowbp_1Mb_rel_vertical_chrsize_Xanthostigma_xanthostigma.png

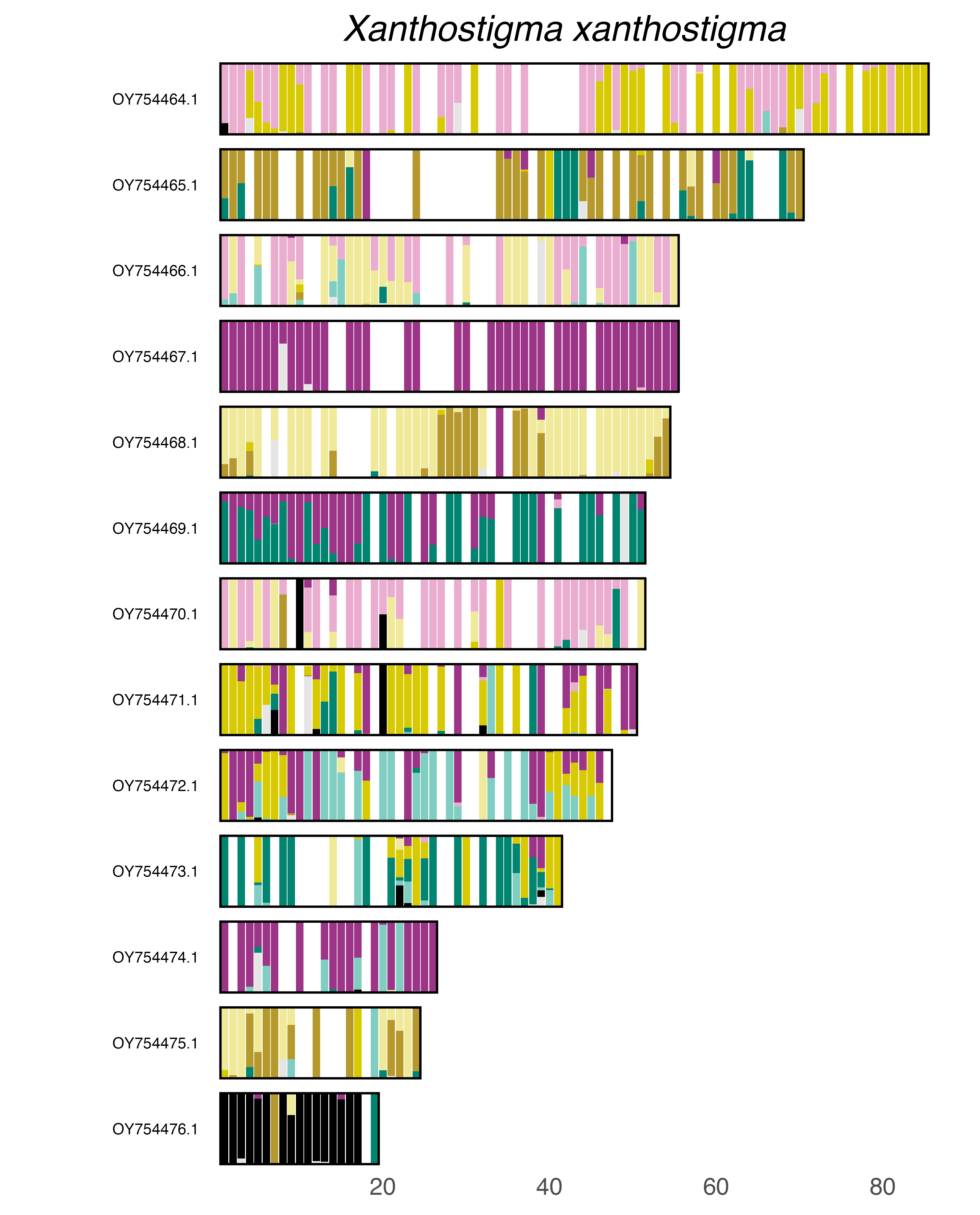

### plot.syngraph_windowbp_1Mb_rel_vertical_chrsize_Zophobas_atratus.png

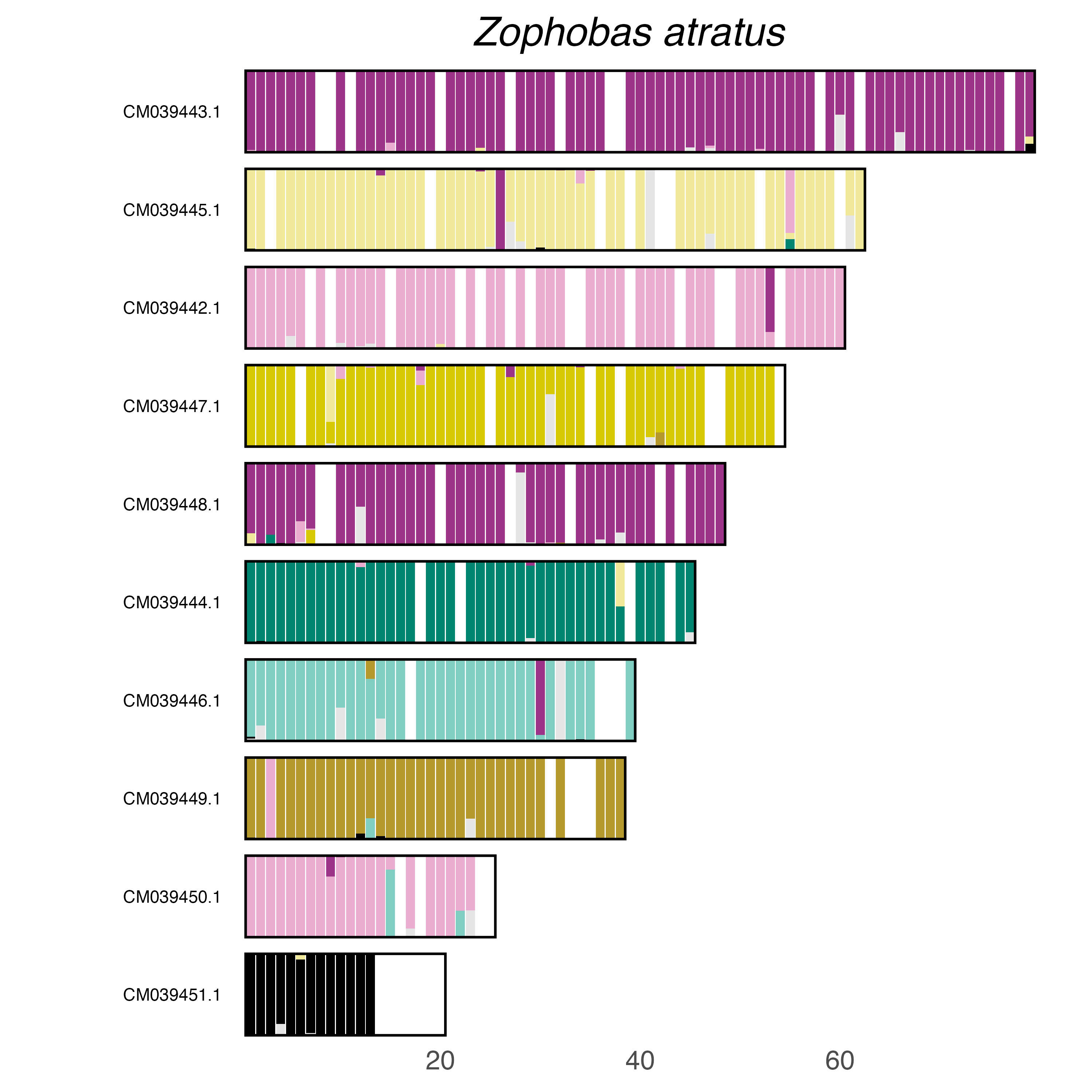
